# Functional metagenomics reveals an alternative, broad-specificity pathway for metabolism of carbohydrates in human gut commensal bacteria

**DOI:** 10.1101/2024.03.25.586180

**Authors:** Seyed Amirhossein Nasseri, Aleksander C. Lazarski, Imke L. Lemmer, Chloe Y. Zhang, Eva Brencher, Hong-Ming Chen, Lyann Sim, Leo Betschart, Liam J. Worrall, Natalie C. J. Strynadka, Stephen G. Withers

**Affiliations:** Department of Chemistry, University of British Columbia, Vancouver, Canada; Michael Smith Laboratories, University of British Columbia, Vancouver, Canada; Department of Biochemistry, University of British Columbia, Vancouver, Canada

## Abstract

The vast majority of the glycosidases characterised so far follow one of the variations of the “Koshland” mechanisms to hydrolyse glycosidic bonds. Herein we describe a large-scale screen of a human gut microbiome metagenomic library using an assay that selectively identifies non-Koshland glycosidase activities. This screen led to identification of a commonly occurring cluster of enzymes with unprecedentedly broad substrate specificities that is thoroughly characterised, mechanistically and structurally. Not only do these enzymes break glycosidic linkages of both α and β stereochemistry and multiple connectivities, but also substrates that are not cleaved by standard glycosidases. These include thioglycosides such as glucosinolates and pseudo-glycosidic bonds of pharmaceuticals such as acarbose. This is achieved via a distinct mechanism of hydrolysis that involves stepwise oxidation, elimination and hydration steps, each catalysed by enzyme modules that are in many cases interchangeable between organisms and substrate classes. These appear to constitute a substantial alternative pathway for glycan degradation.

## Introduction

Most glycosidic bonds are remarkably stable under typical biological conditions, with half-lives for degradation of cellulose and starch being around 5 million years^1^. In order to hydrolyse such substrates on biologically useful timescales, glycosidases have evolved to be among the most proficient of catalysts^1^. For the vast majority of cases, hydrolysis of the glycosidic bond is achieved by one of two well-characterised “Koshland” mechanisms^2^ involving acid/base catalysed reactions via oxocarbenium ion-like transition states (Fig. 1a). Minor variations exist in the identities of the catalytic residues^3^ or the recruitment of intramolecular catalysis^4^. Achievement of such high proficiencies in catalysis by these enzymes requires the evolution of an enzyme active site that exquisitely stabilises the transition state of the reaction, and this is realized through complementary interactions of all types with essentially all parts of the substrate on both sides of the scissile bond^5^. As a consequence, glycosidases tend to be quite specific for their natural substrates.

**Fig. 1:**
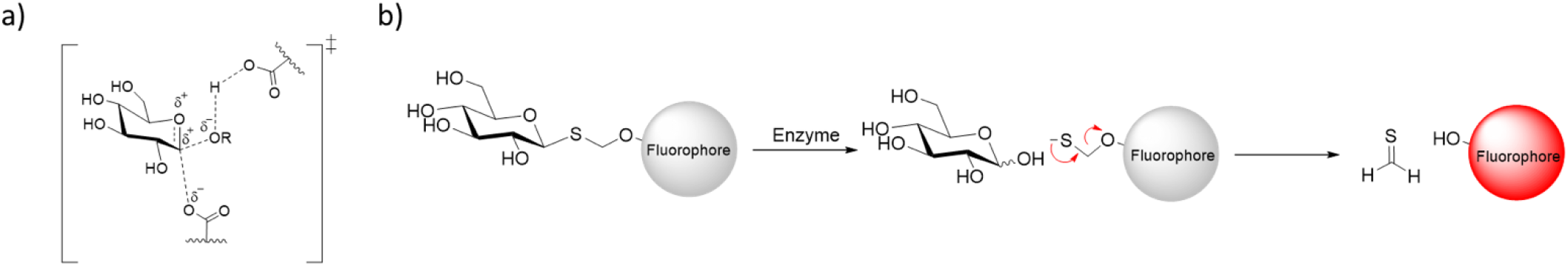
**a)** Oxocarbenium like transition state of a typical retaining Koshland glycosidases b) structure of the selective substrates used in this study and the mechanism of the self-immolative linker disintegration.

The question then arises as to whether alternative, less-demanding and thus less-specific mechanisms for degradation of glycosides may have evolved that would endow organisms with the ability to degrade pools of diverse substrates, otherwise inaccessible to them. This could include complex O-glycans of unusual linkage type that are not cleaved by specific Koshland glycosidases or glycosides linked by atoms other than oxygen, such as carbon, nitrogen or sulfur.

Intrigued by this idea we explored its possibility in the bacterial world through a functional metagenomic approach. The large number of unassigned genes, representing up to 25% of those in known bacterial genomes^6^, certainly allows room for this possibility. As screening substrates for the discovery of new mechanistic classes of glycosidases, we used the fluorogenic alkylthioglycosides (Fig. 1b) that we had previously^7^ shown to not be hydrolysed by Koshland glycosidases, yet cleaved efficiently by non-Koshland glycosidases from CAZy^8^ families GH4/109^9^ and GH88/105^10^. These substrates will therefore detect not only thioglycosidases but also other possible glycoside-cleaving enzymes that do not use Koshland mechanisms.

In this report, we apply this strategy to a large-scale screen of a human gut microbiome metagenomic library using a diverse set of thioglycoside substrates. Twelve unique hits were identified, with at least one hit for each of the substrates used. Through kinetic, mechanistic and structural analysis of two of these hits and a homologous system in another bacterium, we have identified and characterised an operon that carries out the hydrolysis of a remarkably broad set of substrates. Analogues of these enzymes are present in a wide variety of organisms and the components of this system are, in many cases, interchangeable between organism and substrate type. These enzyme systems represent a previously largely unrecognised mode of glycan degradation.

## Results

### Screening of a human gut metagenome library with multiple thioglycosides

We synthesized the seven fluorogenic thioglycoside substrates shown in Fig. 2, based on β-glucose, β-galactose, α-mannose, β-glucuronic acid, 4,5-unsaturated β-glucuronic acid, β-N-acetylglucosamine and β-6-phosphoglucose, based on previously reported methods^7,11^ and as described in the SI (Schemes S1-4). These substrates cover a broad spectrum of commonly occurring glycosides as well as two (**5** and **7**) as controls since enzymes from GH4 (6-phospho-β-glucosidase) and/or GH88/105 (α-4-5-ene-glucuronidase) were likely to be present in our libraries.

**Fig. 2:**
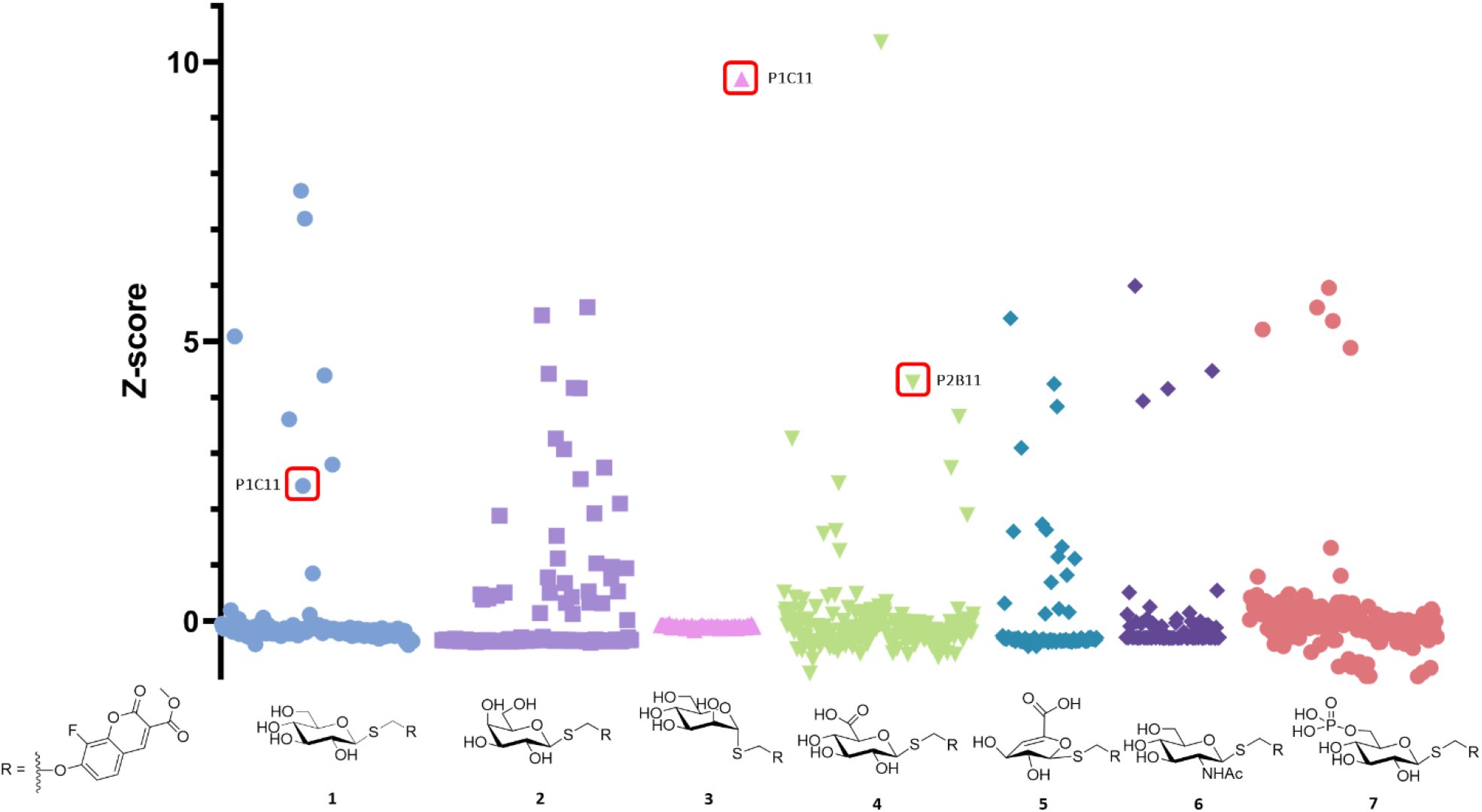
Structures for the substrates used in this study, and the deconvoluted Z-score plots for the 2^nd^ round of screening

Our 20,000 member human gut metagenomic fosmid library^12^ was chosen for the screen, since the vast amount of genome sequence available for the human gut microbiome simplifies and enriches the downstream analysis. Anticipating few hits, we pooled the substrates for our first screen, which to our surprise led to approximately 120 hits (Fig. S2); a hit rate comparable to those of metagenomic screens for well-established enzymatic activities^13^. Hits were picked and arrayed into two 96-well “hit plates”, copies of which were then screened separately with the seven synthesized substrates (Fig. 2), as well as the equivalent O-glycosides. All the results are summarized in Table S1. At least one hit was found for each thioglycoside substrate.

Sequencing of the hit fosmids revealed that many of them were derived from the same regions of bacterial genomes. After removal of redundant hits about a dozen candidates remained. Table S2 summarises these and their most probable source organisms. The majority of these fosmids contain no known glycosidase genes, yet hydrolytic activity was unequivocal. In this report, we focus on two of the hits for which we have fully characterised the sets of proteins responsible for the activity. Detailed investigation of the other hits is ongoing.

### Investigation of the α-thiomannosidase hit P1C11

A particularly interesting hit was P1C11, the single clone that was active against the α-S-mannoside substrate **3** and the β-S-glucoside substrate **1**, hinting at an unusual mechanism that is able to hydrolyse both α and β glycosides (Fig. 2 and Table S1). Sequencing of this fosmid revealed its origin from an *Alistipes* species with open reading frames (ORF) encoding many uncharacterised genes along with several annotated as carbohydrate-related enzymes, but no known glycosidases (Fig. 3a). The closest was a domain of unknown function (DUF)1080-containing gene encoding a protein with structural similarity to GH16 endo-1,3-1,4-β glucanases^14^ along with a nearby protein annotated as a sugar phosphate isomerase, two Gfo/Idh/MocA-like oxidoreductases and a DUF6377 predicted to be a transcriptional regulator. *In silico* analysis of these proteins, henceforth called AL1, AL2, AL3, AL4 and AL5 (Fig. 3b) predicts that in addition to AL5 which is membrane associated^15^, three of the other proteins have signal peptides^16^: AL1 has a secretion signal peptide, AL2 a lipoprotein signal peptide and AL3 a twin-arginine signal peptide suggesting periplasmic locations while AL4 is cytosolic.

**Fig. 3:**
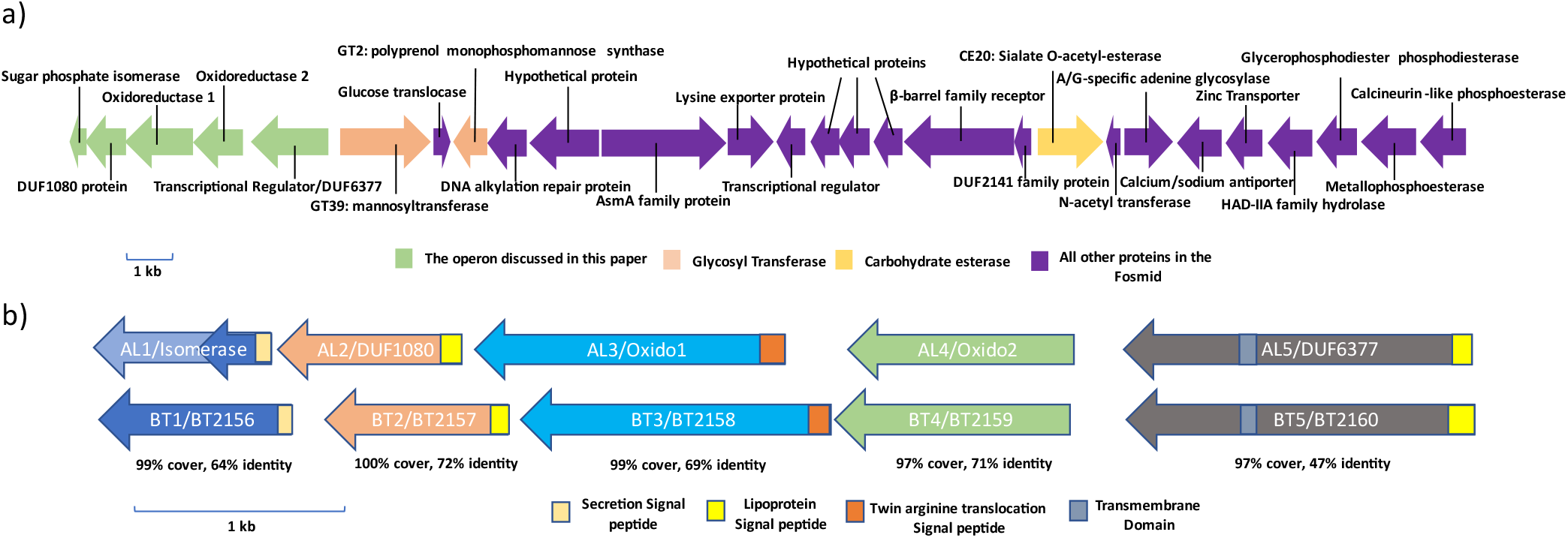
a) ORFs present in the hit, P1C11, these include glycosyltransferases from GT2 and GT39 families and a carbohydrate esterase, the five genes discussed in this report are shown in green b) The organization of the genes in the two clusters showing the percent identity of the AL enzymes with their BT counterparts. The ORF for AL1, present in the fosmid was truncated (dark blue); the full length ORF found in multiple *Alistipes* sp. genomes is shown in light blue.

### Parallels with recent reports in *Bacteroides thetaiotaomicron*

Intriguingly, a very similar operon (BT2156-2160, Fig. 3b) in another prevalent gut commensal bacterium, *Bacteroides thetaiotaomicron*, was recently identified as responsible for metabolism of glucosinolates, naturally occurring β-thioglucosides in plants.^17^ At least three of the four enzymes, as well as the regulator, were needed for activity in vivo while preliminary data suggested that only two of the enzymes were required to observe the activity in vitro, though the roles of these proteins were not elucidated. The same cluster was also identified through a functional genetics study on *B. thetaiotaomicron*^18^ as being responsible for degradation of several α-linked disaccharides including α,α-trehalose, leucrose and maltitol.

The similarity of our *Alistipes* sp. enzymes to those in *B. thetaiotaomicron* and the limited mechanistic and structural insights into how this *B. thetaiotaomicron* cluster functions to degrade such diverse glycans, led us to embark on parallel structural and mechanistic investigations of the two systems. To that end we cloned each of the enzyme-encoding genes and expressed and purified them from *E. coli* as described in the SI.

### Mechanistic study of the AL and BT enzyme clusters

With all these proteins in hand, we demonstrated that the four BT enzymes, used together, hydrolyse both our fluorogenic β-S-glycoside substrate **1** and the α-S-mannoside **3** as well as both the α and β anomers of methylumbelliferyl O-glucoside and mannoside, with a decided preference for the β-glycosides. The four AL enzymes together likewise hydrolyse both anomers of both the glucoside and mannoside, but with a preference for the α anomers (SI section 2.5.2.3.). As implied by Liou *et al*^17^, neither AL4 or BT4 are essential for activity in vitro (SI section 2.5.2.2.). Importantly, we showed (SI section 2.5.1.2.) that enzymes AL/BT2 and AL/BT3 are largely interchangeable between the two systems, suggesting similar roles and also that they do not work as obligate multi-enzyme complexes.

While the cloned enzymes were active together in vitro, the overall rates under the conditions used were extremely low, as also reported by Liou *et al*^17^, making mechanistic studies challenging and more importantly suggesting a missing component. However, addition of various metal ions, varying the buffer salts, ionic strength, etc. led to only minor rate improvements, (SI sections 2.5.1.3 and 2.5.1.4). By contrast addition of EDTA ablates the reaction for both series of enzymes indicating important previously bound metals.

Despite the fact that the clusters contain a pair of oxidoreductases, addition of NAD(P)(H) had no pronounced effect on the reaction rates, nor did any other tested cofactors (SI section 2.5.1.5). Assuming that NAD(H) is indeed involved, based on the precedent of other Gfo/Idh/MocA oxidoreductases, this would suggest a tightly bound cofactor carrying out an oxidation of the substrate and getting recycled back into its oxidised form by some means other than cofactor exchange, consistent with the periplasmic location. Based upon our previous studies of the GH4 enzyme family^19^, the most likely mechanism involves an initial oxidation by AL/BT3 to generate a 3-keto-sugar intermediate, as suggested by Liu et al^18^. This acidifies the proton at C2, and facilitates glycoside cleavage via an elimination reaction: entirely different from the substitution reaction of Koshland glycosidases.

In the absence of NAD(H) exchange in the in vitro system, AL/BT3 will get stuck in the reduced form, unable to perform another catalytic cycle. In order to bypass this restriction we added methyl β-3-ketoglucoside (synthesized using a site-selective palladium catalyst^20^) to the reaction mixtures of both series of enzymes as a stoichiometric re-oxidant. This indeed increases the hydrolysis rates of our fluorogenic substrates by more than 100-fold (Fig. S9) indicating that this was the rate-limiting step and opening up the opportunity of detailed mechanistic investigations.

### Reactions and roles of individual enzymes

Roles of individual enzymes were next identified by ^1^H-NMR analyses of their reactions. We first confirmed the roles of AL/BT3 as the initial oxidants by showing that addition of AL/BT3 to equimolar mixtures of a glucoside and a 3-keto-glucose derivative (Glc-α-pNP and 3-keto-1,5-anhydoglucitol in Fig. 4a) results in oxidation of the glucoside to its 3-ketoglucoside, yielding an equilibrium mixture (54:46) of the reduced:oxidised species. Performing the same reaction with the reverse reduced/oxidized pair of substrates gave the same equilibrium mix (SI section 2.7.6.).

**Fig. 4:**
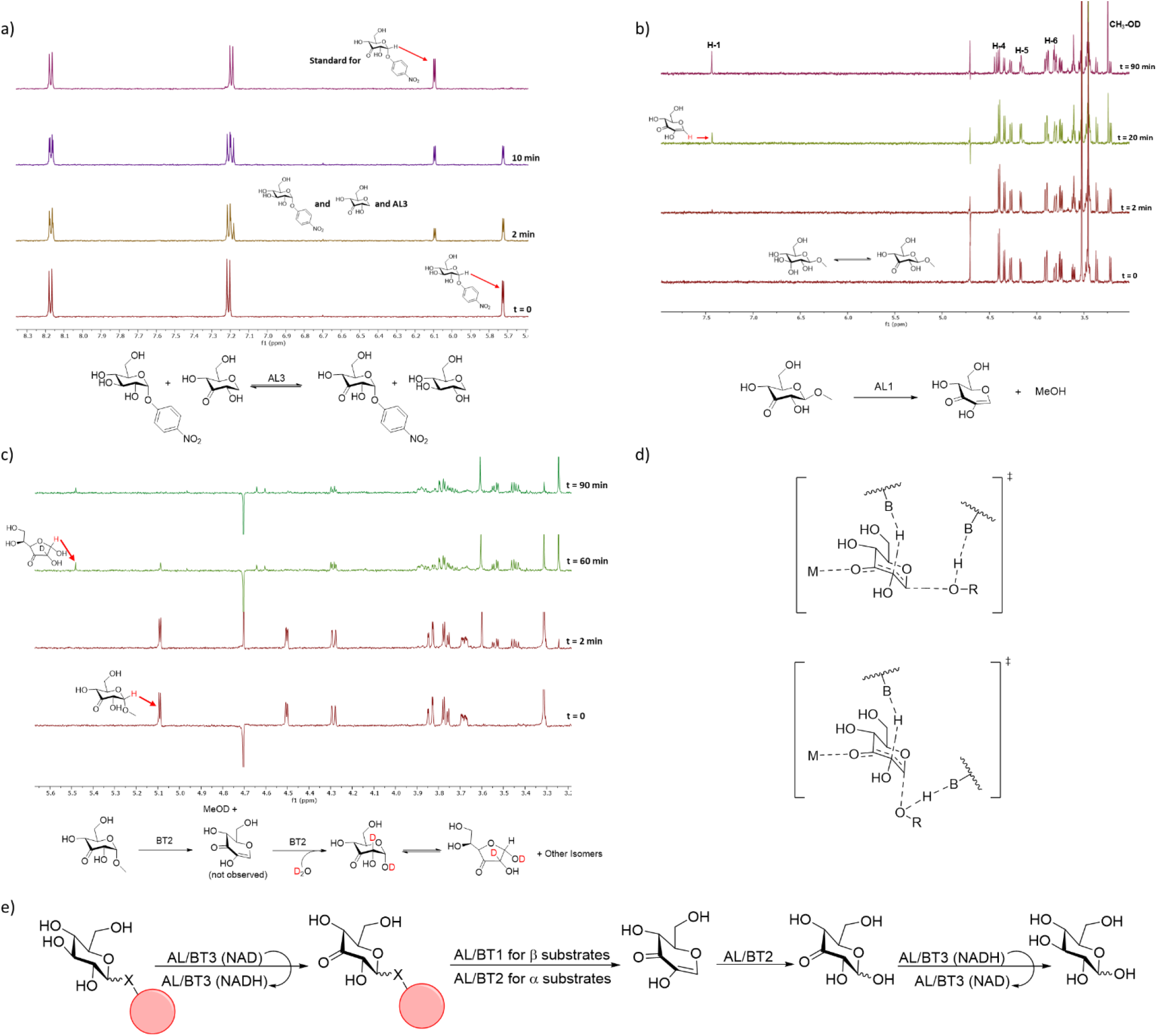
a) ^1^H-NMR characterisation of the reactions of a) AL3 and b) AL1 c) BT2 and d) proposed transition states for AL/BT1 (top) and AL/BT2 (bottom), B refers to protein residues capable of acid/base catalysis and M refers to the metal ions that coordinate the 3-keto group. e) Mechanism of stepwise hydrolysis of α and β glycosides by the enzymes AL and BT 1-3.

Roles for the other components were assigned by monitoring reactions of chemically synthesized 3-ketoglycoside versions of the minimalist substrates, methyl α- and β glucoside. As seen in Figs. 4b and c, AL/BT1 carry out the elimination reaction on the β anomer of 3keto-GlcOMe, while AL/BT2 do so on the α anomer. Hydration of the resultant 3-keto-2-hydroxy-glucal is then performed efficiently by AL/BT2 while AL/BT1 only does so much more slowly (SI section 2.7.2. to 2.7.5.). Hydration leads to a mixture of isomers of 3-keto-glucose, dominant among which is the 3-keto-gluco-furanose shown in Fig. 4c^21^. Thus, the enzymes AL/BT1, previously annotated as sugar phosphate isomerases, are in fact 3-keto-β-glucoside-1,2-lyases while AL/BT2, members of DUF1080 family, are both 3-keto-α-glucoside-1,2-lyases and 3-keto-2-hydroxy-glucal-hydratases, and not 3-keto-glucoside hydrolases as previously suggested^18^. The dual activity of AL/BT1 and AL/BT2 is not surprising, considering the very similar transition states for the elimination and hydration reactions (Fig. 4d). Control experiments (Fig. S93-94) confirm that these enzymes alone have no effect on simple glucoside substrates such as para-nitrophenyl glucosides. Moreover, by manipulating enzyme concentrations or by stepwise addition of the enzymes, the intermediates of each step in the overall reaction for hydrolysis of both anomers of para-nitrophenyl glucosides are detected by NMR (Figs. S91-92 and 95-98). Based on these findings, the overall pathway for hydrolysis of glycosides by the enzymes AL/BT 1-3 is depicted in Fig. 4e.

### X-ray crystallography

To gain further insights into the mechanisms of these enzymes, high resolution models of AL1, AL2, AL3, BT1, and BT2 were determined by X-ray crystallography (see SI section 2.8. for details). Resulting models of AL1 and BT1 displayed a classical overall α/β (TIM) barrel fold, typical of the isomerases they are related to, with an eight-stranded core barrel surrounded by eight helices on the outer layer (Fig. 5a). The active site contains a conserved catalytic Glu (249 and 254 in AL1 and BT1 respectively) for proton extraction (Fig. 5b). In agreement with the results of kinetic assays (SI section 2.5.4.2), a Co^2+^ ion was observed and confirmed by XRF scans of AL1 and BT1 crystals (Fig. S103a,b). This is coordinated by His 178/182, Asp 211/214, and His 237/240 as well as the 3-keto group of the substrates in the substrate-bound models (Fig. 5b,c). In addition to this ion, the substrate binds to His 103/107, Trp 214/217, and His 180/184 while the predicted catalytic Glu 272/275 side chain OE2 is positioned 2.7 Å above carbon 2 of the bound substrate analogue, 3-keto-1,5-anhydroglucitol 157/163 (Fig. 5b).

**Fig. 5:**
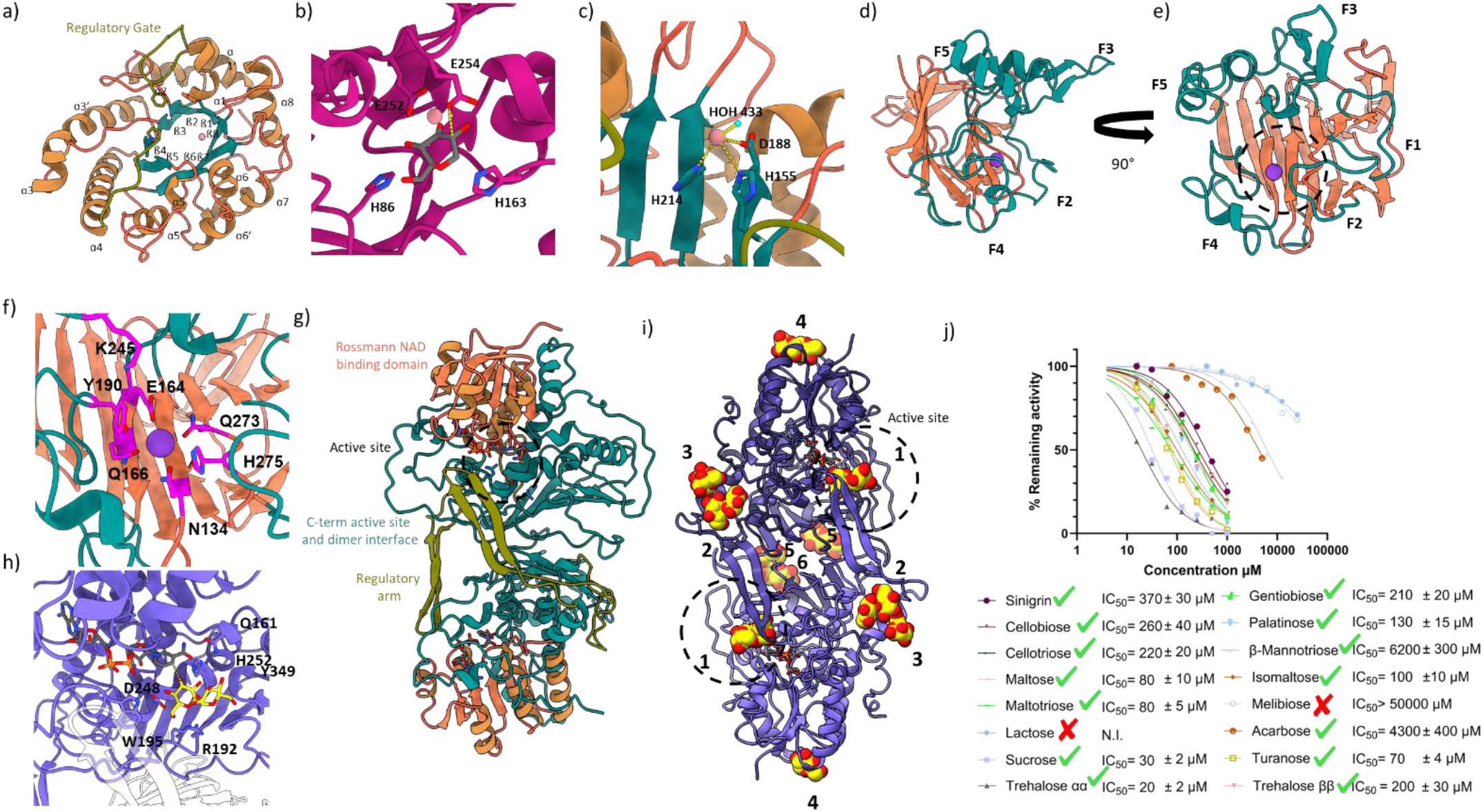
a) Cartoon representation of AL1 with secondary structure, and regulatory loop labelled. Barrel strands (cyan) helices (orange), and regulatory gate (olive). b) Active site of BT1 bound with substrate analogue 3k-1,5-anhydroglucitol with coordinating residues labelled c) Metal binding site of apo AL1 with coordinating residues labeled d) Front view and e) Side view of overall fold of BT2 with jelly roll in orange, fingers (cyan) labelled F1-F5, and active site denoted with a dashed circle. f) BT2 active site with conserved possible active site and metal binding residues in medium magenta g) Overall fold of AL3 with Rossmann-like NAD fold (orange), the C-terminal active site region and dimerization interface (cyan) and regulatory arm (olive). h) Active site of AL3 bound with trehalose with key residues labelled i) Binding sites of trehalose observed in AL3 crystal soaked with trehalose. Active site is labelled as 1 and circled, secondary sites are labelled 2-6 j) Inhibition of AL enzymes by a select number of disaccharides: those that were hydrolyzed are shown with green check marks and those that are not hydrolyzed are marked with red x marks.

Apo BT2 and AL2 formed monomers adopting a ß jelly roll fold that creates a catalytic channel flanked by five finger domains comparable to proteins from the GH16 family^14^ (Fig. 5d,e). BT2 and AL2 displayed highly similar folds with an RMSD of 0.5 Å across 229 Cα pairings (Fig. S110). Notable divergences from GH16 structures were observed within the catalytic channel where no homologous E-[ILV]-D-[IVAF]-[VILMF] (0,1)-E motif^22^ was observed indicating that while the overall fold is similar, BT2 contains different catalytic residues. A potassium ion was observed in the channel (Fig. S103i), which likely replaced the functional metal (proposed to be Ca^2+^ based on the kinetic data, SI 2.5.4.2.) during crystallization due to its high concentration (Fig. 5f, Table S6). To identify possible catalytic and metal coordinating residues, BT2 crystals were soaked with α,α-trehalose prior to diffraction. Soaked crystals contained a glucose-like molecule in the active site coordinated at the 2, 3, 4, and 6 OH groups by His 275, the metal ion, Gln 273, and Lys 245 respectively (Fig. S107). Consistent with this, sequence alignments confirmed conserved amino acids in the channel where Glu 103 (>95% conserved as Glu/Asp), Asn 134, Glu 164, Gln 166, Tyr 190, Lys 245, Gln 273 and His 275 were all highly conserved across 3183 homologous sequences and likely comprise the metal binding and active site of BT2 (Figs. 5f, S104b).

AL3 structures crystallized as dimers with an N-terminal Rossmann like fold, and a C-terminal domain containing the active site, dimerization interface, and a regulatory loop that blocks the opposing monomer active site in the dimer pair (Fig. 5g). Both apo and trehalose-bound enzymes crystallized with NAD co-factors (Fig. S108c), however, despite XRF scans indicating the presence of Co^2+^ (Fig. S103e), no Co^2+^ was observed. Trehalose-bound AL3 identified the primary glycoside binding pocket as comprised of Gln 161 (that mediates the adjacent NAD cofactor), as well as Arg 192 (coordinating O-5 and OH-6 of the glucose ring), Trp 195, Asp 248, His 252 (OH-3), and Tyr 349 (OH-2) (Fig. 5h). Numerous additional distinct trehalose binding sites were observed beyond that of the active site just described, with some arranged in a chain-like configuration. It is tempting to suggest this continuous path of sugar binding may show how AL3 and BT3 bind larger oligosaccharides for catalysis (Fig. 5i, see below for further discussion), but given the high concentration of trehalose present in the soaking experiment, non-specific binding may be responsible.

### Broad substrate specificities

The finding that enzyme systems of this class are able to hydrolyse simple non-activated substrates such as methyl glucosides of both α and β configurations is unprecedented. We were therefore interested in exploring whether this broad specificity extends to commonly occurring oligosaccharides, since this might illuminate the reasons for the parallel evolution of these enzymes in the presence of highly efficient Koshland glycosidases.

As an initial screen, using a mixture of all the enzymes, we assayed inhibition of the hydrolysis of our fluorogenic substrates by a series of oligosaccharides, since this should detect both competitive substrates and competitive inhibitors (SI section 2.5.5.). As shown in Fig. 5j, with the exception of galactosides, a wide range of α- and β-glycosides of varying linkage were inhibitory. Further analysis by TLC and ^1^H-NMR (SI sections 2.6.2, 2.7.9. and 2.7.10.) reveals, remarkably, that all of the inhibitory oligosaccharides are indeed completely hydrolysed. This includes both β- and α-glucosides with essentially any glycosidic linkage to the next sugar unit. The next sugar unit can be any monosaccharide (examples include sucrose (Glcα(→)β ruc) and maltose (Glcα(→)Glc), or indeed larger saccharide units since cello- and malto-oligosaccharides are all efficiently hydrolysed. This broad activity also extends to glucosides with aglycones such as phenols (salicin, fluorogenic MU-glycosides) as well as thio-glycosides such as sinigrin. The enzymes appear to be strict exo-glycosidases degrading from the non-reducing end and thus have little effect on polysaccharides in vitro. In vivo, however they would work in concert with suitable endo-glycosidases to degrade such substrates.

Since the mechanism of action of these enzymes does not require an endocyclic oxygen, we hypothesized that pseudo-glycosides could also be cleaved. Indeed, the enzyme systems also break the C-N bond of the pseudo-oligosaccharide acarbose (Fig S70 & 72), a Type II diabetes therapeutic, hinting at a possible degradation pathway for this drug molecule and presumably for other compounds of this class in human gut environments that harbour these enzymes. This provides another example of how the human gut microbiome can metabolise drugs, this time discovered via an indirect mechanistic approach. Indeed, essentially any cyclohexyl ring with the right configuration of hydroxy groups for oxidation of the 3-hydroxy should undergo a subsequent elimination reaction to break the (pseudo)glycosidic bonds, provided that the substrate fits into the enzyme active sites.

Extension of the inhibition assay on AL/BT3 to monosaccharides (SI section 2.5.5.) reveals that the only essential hydroxy groups in the case of AL/BT3 are those at the 3 and 4 positions. Modifications at other positions are allowed as long as their size does not preclude active site binding. Thus, for example, quinovose and 6-phospho glucose are accepted but not melibiose (Galα(→)Glc).

A consequence of this mechanism is that glycosides modified at the 3-position are not hydrolysed, because oxidation does not occur. Interestingly, many of the aminoglycoside antibiotics have C-3 modifications and none of those we tested are substrates of these enzymes (Figs. S71 & S72, also SI section 2.6.1.3.3.). 3-Amino modified monosaccharides can also be found among many other natural products^23,24^, while 3-*O*-methylation, deoxygenation and branching occurs in some other cases^25,26^. It therefore seems quite probable that modifications at the 3-position have evolved as a means of combatting the degradation of glycosylated natural products such as antibiotics via a 3-keto mediated pathway.

### Investigation of the β-thioglucuronidase hit P2B11

Another of our fosmid hits, P2B11, from an unknown Gram-positive bacterium is active on thio-glucuronide substrate **4**. This fosmid encodes a cluster of four enzymes in which, unlike in the above cases, the two oxidoreductase genes are flanked by two genes that are annotated as sugar-phosphate isomerases with no apparent nearby transporter/regulator (Fig. 6a). While several other genes annotated as glycosidases are found in this fosmid, the similarity of this set of four genes to those previously studied (Fig. 6b) suggested that they were responsible for the activity. Biochemical characterisation confirmed this, revealing significant similarity between these proteins and those discussed above, despite modest sequence similarity. Here too, Oxido1 proved to be the most important for activity, while Iso1 and Iso2 both enhance the rate decidedly and Oxido2 has minor effects (SI section 2.5.6.5.). Catalytic activity is enhanced in the presence of Mn^2+^ and Co^2+^ ions, while the presence of NAD(P)(H) cofactors again has no effect on reaction rate. These four proteins together also have a relatively broad substrate specificity, hydrolysing both β- and α-glucuronide substrates at similar rates (SI section 2.5.6.6.). While glucosides are not hydrolysed, glycosides bearing negative charges at the 6-position, like 6-PO_4_, 6-SO_4_ and sulfoquinovoside substrates are all cleaved, albeit slowly compared to glucuronides. Some of the components of this cluster can be swapped with those found in P1C11, despite different substrate specificities. Thus, AL3 can efficiently oxidise glucuronide substrates, and a mixture of AL3 and Iso 1 and Iso 2 from P2B11 hydrolyses the glucuronide substrate (SI section 2.5.6.9.), confirming the mechanistic parallels.

**Fig. 6:**
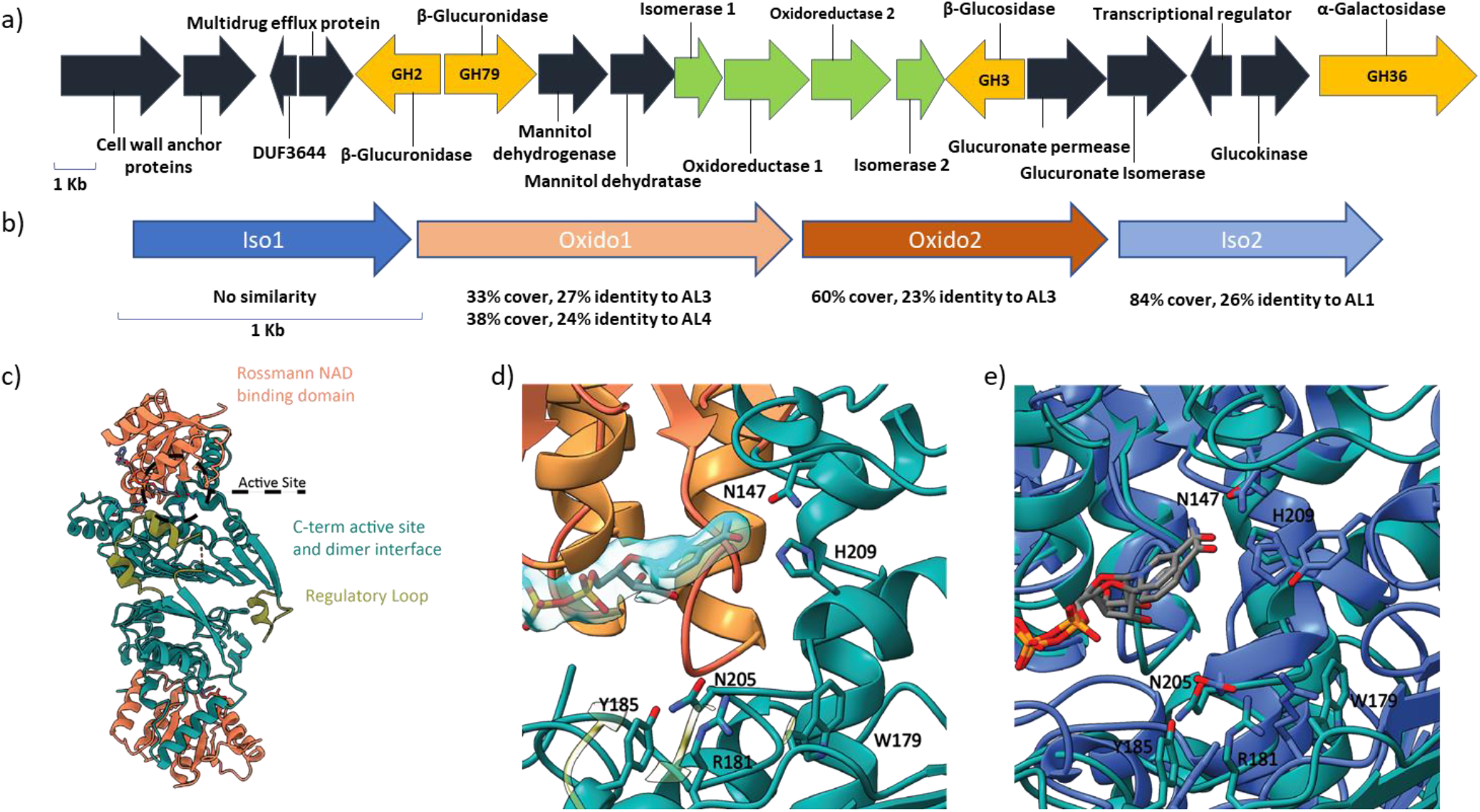
a) ORFs present in the hit, P2B11 b) The organization of the genes, with the percent identity of the proteins with AL and BT counterparts. c) Overall fold of P2B11-Oxi1 dimer pair with Rossmann like fold (orange), C-terminal active site and dimerization interface (cyan), and regulatory loop (olive) denoted. d) P2B11-Oxi1 apo active site with core residues labelled and 2mFo-dFc omit map around NAD shown as a surface at map level 0.3 in ChimeraX. e) Alignment of P2B11-Oxi1 apo active site with AL3 apo RMSD 1.2 Å over 91 Cα pairs, core CT3 residues are labelled.

P2B11-Oxido1 crystallized as a dimer (six dimers in the asymmetric unit of the P21 cell) with the core N-terminal Rossmann domain, C-terminal active site region, dimerization interface, and inhibitory loop for opposing monomers displaying an overall fold comparable to that of AL3 (Fig. 6c). XRF scans of the crystals confirmed the presence of Co^2+^ (Fig. S103f) suggesting metal binding and specificity; however, as with AL3 no ordered anomalous electron density was evident in the active site maps. Similar predominantly hydrophobic interfaces were observed between each inhibitory loop and its regulatory partner in both enzymes; however, the relative orientation of the fold and disposition of the active sites on opposite faces in P2B11-Oxido-1 dimer differed compared to the same faces in AL3 (Fig. S111b). Glycoside binding pockets appear to contain the same core in Asn 147 for NAD coordination, while Trp 179, Arg 181, Tyr 185, Asn 205, and His 209 comprise the active site pocket (Fig. 6d). Most residues appeared in similar positions except for Tyr 185 and Trp 179 lying on opposite sides of Arg 181 relative to the analogous residues in AL3 (Fig. 6e).

## Discussion and conclusions

The operons described in this study endow the bacteria that encode them with the ability to not only cleave otherwise recalcitrant thioglycosides, but also both the α and β glycosidic bonds of a wide range of commonly occurring oligosaccharides. This versatility derives from their completely different chemical mechanism, involving anionic transition states rather than the cationic oxocarbenium ion-like transition states of typical Koshland glycosidases, as well as the separation of the reaction into several steps. We established the functions of the individual enzyme components by NMR monitoring of their reactions, detected each intermediate in the overall reaction, and identified the key residues involved in binding and catalysis by solving the structures of the enzymes.

As depicted in Fig. 4e, the tightly bound NAD cofactor of AL/BT3 is most likely recycled in vivo through reduction of the 3-ketoglucose generated by AL/BT2. Addition of the 3-ketoglucose derivative was necessary for initial rate kinetics in vitro, since isomerisation of 3-ketoglucose to its furanose form reduces the concentration of the active form. Re-oxidation of the enzyme in vivo might also occur via specialized proteins, such as cytochromes^27,28^ or via other small molecule hydride acceptors. To assess this possibility, we screened a range of common carbonyl-containing chemicals and found that D-erythrose and dehydroascorbic acid can also serve as co-substrates, (SI section 2.5.7.). The reaction rates are significantly lower than with 3-ketosugars however, and the relevance of these specific molecules in vivo when 3-keto-glucose itself will be generated by AL/BT2 remains to be seen.

As shown by Liou et al^17^, homologues of the BT enzymes are widely prevalent in healthy human gut microbiome samples. Moreover, our identification of the glucuronide-hydrolysing hit shows that bacterial breakdown of glycosides through a 3-keto intermediate is not limited to glucosides and likely extends well beyond. The relatively low sequence similarity of the Gram-positive sourced enzymes from P2B11 with their Gram-negative counterparts from *Alistipes* and *Bacteroides*, as well as the absence of a DUF1080 containing protein, foretell that other operons utilising this step-wise process may indeed have varied architectures and distant sequence similarities.

There are several consequences of the differences of this mechanism from that of the Koshland glycosidases. Since the hydroxyl at C3 is the linchpin of this mechanism rather than the anomeric centre, both α and β- anomers can be accommodated. Indeed, comparable pseudo-Michaelis-Menten kinetic parameters are found for the reactions catalyzed by individual or collective enzymes with both anomers of simple aryl glycosides. Further, by breaking down the reaction coordinate into a series of lower energy steps, less evolutionary optimisation is required to arrive at useful reaction rates for diverse and challenging substrates. Moreover, the interchangeability of these subunits should make the evolution of “new” activities more facile. This versatility does, however, come at the cost of expressing multiple proteins to perform a reaction that Koshland glycosidases carry out efficiently in one step.

Keto glycosides were first discovered in the 1960s, when bacterial strains that oxidise disaccharides into 3-keto sugars were reported^29^. The best studied example was the plant pathogen *Agrobacterium tumefaciens*, which converts sucrose, among other disaccharides, to 3-keto-sucrose^30,31^. However, the identity of the enzymes responsible was not known and no chemical rationale was provided for the oxidation: the enzymes responsible for breaking the glycosidic bonds were assumed to be regular glycosidases. More recently, ketosugars have been reported as intermediates in the breakdown of anhydrosugars such as levoglucosan^32,33^, as well as C-glycoside-cleaving systems that operate via similar oxidation and elimination reactions.^34–38^ As mentioned, other glycosidases that transiently oxidize sugar hydroxy groups as part of their mechanism have been reported by us and others^9,39–41^. However, no similarly broad specificity was observed in any of these cases. While levoglucosan is not a substrate of our enzymes, AL/BT3 are capable of oxidising C-glycosides but the lyase enzymes AL/BT1 do not further degrade the 3-keto-C-glycoside (See SI section 2.6.1.3.2.), presumably because they lack the required residues for aromatic enolization^42^.

Our findings consolidate and considerably extend these disparate reports and lay out substantive structural and mechanistic support for this common, highly flexible alternative pathway that is employed by bacteria for degradation of a wide range of glycans. Glycosidases are arguably one of the most extensively studied classes of enzymes. Elaboration of a new biochemical pathway for metabolism of one of the best-studied class of biomolecules, in the well-trodden ground of the human gut microbiome serves as a reminder that there remains an immense amount to be learned about the chemistry of biological systems.

## Experimental methods

All the experimental methods are explained in the supplementary information file.

## Acknowledgements

This work was supported by operating grants from the Natural Sciences and Engineering Research Council of Canada (NSERC) and the Canadian Institutes of Health Research (CIHR) as well as infrastructure funds from the Canada Foundation for Innovation and BC Knowledge Development Fund. S.A.N. was supported by a Vanier scholarship from NSERC. N.C.J.S. is a Tier I Canada Research Chair in Structure-guided Antibiotic Discovery. Part of the research described in this paper was performed using beamline CMCF-BM at the Canadian Light Source, a national research facility of the University of Saskatchewan, which is supported by the Canada Foundation for Innovation (CFI), the NSERC, the National Research Council (NRC), CIHR, the Government of Saskatchewan, and the University of Saskatchewan. We also thank GM/CA beamline staff at beamline 23-ID-B at the APS for access and support. GM/CA@APS has been funded in whole or in part with Federal funds from the National Cancer Institute (ACB-12002) and the National Institute of General Medical Sciences (AGM-12006).

We would like to thank Dr. Peter Rahfeld for guidance about the screening process, Dr. Deepesh Panwar and Prof. Harry Brumer for critical reading of the manuscript, Avery Noonan and Prof. Steven Hallam for assistance with processing of sequencing data, Dr. Maria Ezhova for assistance with the NMR experiments and Dr. Feng Liu, Dr. Spence Macdonald, Dr. Philip Danby, Benjamin Herring, Dr. Jacob Wardman, Dr. Charlotte Olagnon, Rhea Bains and Dr. Yuqing Tian for useful suggestions and discussions. This work was inspired by and is dedicated to Professor Emeritus Tony Warren of the UBC Department of Microbiology and Immunology.

## Notes

### Competing Interest Statement

The authors have declared no competing interest.

