## Supplementary information for "Functional metagenomics reveals an alternative, broad-specificity pathway for metabolism of carbohydrates in human gut commensal bacteria"

**Abstract:** The vast majority of the glycosidases characterised so far follow one of the variations of the "Koshland" mechanisms to hydrolyse glycosidic bonds. Herein we describe a large-scale screen of a human gut microbiome metagenomic library using an assay that selectively identifies non-Koshland glycosidase activities. This screen led to identification of a commonly occurring cluster of enzymes with unprecedentedly broad substrate specificities that is thoroughly characterised, mechanistically and structurally. Not only do these enzymes break glycosidic linkages of both α and β stereochemistry and multiple connectivities, but also substrates that are not cleaved by standard glycosidases. These include thioglycosides such as glucosinolates and pseudo-glycosidic bonds of pharmaceuticals such as acarbose. This is achieved via a distinct mechanism of hydrolysis that involves stepwise oxidation, elimination and hydration steps, each catalysed by enzyme modules that are in many cases interchangeable between organisms and substrate classes. These appear to constitute a substantial alternative pathway for glycan degradation.

**List of the abbreviations**

| | |
|---|---|
| **Ac** | Acetyl- |
| **ACN** | Acetonitrile, also abbreviated as MeCN |
| **BLAST** | Basic Local Alignment Search Tool |
| **BGLK** | β-glucoside kinase from *Klebsiella pneumoniae* |
| ***B.T.*** | *Bacteroides thetaiotaomicron* |
| **δ** | Chemical shift |
| **DBU** | 1,8-Diazabicyclo[5.4.0]undec-7-en |
| **DCE** | 1,2-Dichloroethane |
| **DCM** | Dichloromethane |
| **DIPEA** | N,N-Diisopropylethylamine |
| **DMF** | Dimethylformamide |
| **DMSO** | Dimethylsulfoxide |
| **EDTA** | Ethylenediaminetetraacetic acid |
| ***E. coli*** | *Escherichia coli* |
| **eq** | Equivalent(s) |
| **ESI** | Electrospray-Ionisation |
| **EtOAc** | Ethyl acetate |
| **EtOH** | Ethanol |
| **HEPES** | (4-(2-hydroxyethyl)-1-piperazineethanesulfonic acid) |
| **HPLC** | High performance liquid chromatography |
| **HRMS** | High resolution mass spectrometry |
| **IPTG** | Isopropyl β-*D*-thiogalactoside |
| ***J*** | Coupling constant |
| **JB** | Jericho blue // 3-carboxy-8-fluoro-7-hydroxycoumarin |
| **JB-OMe** | 3-methylcarboxlate-8-fluoro-7-hydroxycoumarin |
| **LB** | Luria-Bertani broth, also known as Lysogeny broth |
| **MES** | 2-(*N*-morpholino)ethanesulfonic acid |
| **MeOH** | Methanol |
| **MU** | 4-Methylumbelliferone |
| **NaPi** | Sodium Phosphate |
| **NMR** | Nuclear magnetic resonance |
| **O/N** | overnight |
| **PBS** | Phosphate saline buffer |
| **PE** | petroleum ether |
| **PMHS** | Poly(methylhydrosiloxane) |
| **$R_f$** | Retardation factor |
| **Rpm** | Revolutions per minute |
| **rt** | Room temperature |
| **SD** | Standard deviation |
| **TRIS** | Tris(hydroxymethyl)aminomethane |
| **TBA** | Tetrabutylammonium |
| **THF** | Tetrahydrofuran |
| **TLC** | Thin layer chromatography |

### 1. Synthesis and characterization of compounds

#### 1.1. Reagents, solvents and instruments

All chemicals utilized were purchased from Sigma-Aldrich, Toronto research chemicals or Biosynth unless otherwise noted. As mentioned in the procedures, some of the reactions were conducted using dry solvents, which were made anhydrous by distillation under a nitrogen atmosphere: DCM and MeCN over $CaH_2$ and MeOH over Mg. DCE and $CH_2Br_2$ were dried over molecular sieves. All reactions were conducted under inert conditions (argon/nitrogen atmosphere) unless otherwise noted. The glassware was dried in an oven at 120 °C. All commercially available reagents and solvents were used without further purification. TLC was performed on silica plates 60 F254 aluminum sheets (Merck, Germany). TLC spots were visualized via UV light and/or through staining with 10% ammonium molybdate in 2 M $H_2SO_4$. Column chromatography was carried out using silica gel with 230-400 mesh size. When working up reaction mixtures, solutions of organic solvents were washed with approximately the same volumes of aqueous solutions. $^{1}H$, $^{13}C$, $^{19}F$ and $^{31}P$ NMR spectra were recorded in $(CD_3)_2CO$, $(CD_3)_2SO$, $D_2O$ or $CD_3OD$, are reported in δ scale in ppm and are referenced to one of the following: $(CD_3)_2CO$ (δ 2.05 ppm for $^{1}H$, δ 206.26 ppm for $^{13}C$), $(CD_3)_2SO$ (δ 2.50 ppm for $^{1}H$, δ 39.52 ppm for $^{13}C$), $CD_3OD$ (δ 3.31 ppm for $^{1}H$, δ 49.00 ppm for $^{13}C$). For $^{19}F$ and $^{31}P$ spectra, $CFCl_3$ and 85% $H_3PO_4$ were used as external references. Data for the NMR spectra are reported as follows: s = singlet, d = doublet, dd = doublet of doublets, t = triplet, q = quartet, m = multiplet, *J* = coupling constant in Hertz. The coupling constants are accurate within ± 0.3 Hertz range. Low resolution mass spectra (LRMS) were obtained using a Waters ZQ mass spectrometer equipped with ESCI ion source and High-resolution mass spectra (HRMS) were recorded by the University of British Columbia mass spectrometry facility in a Waters/Micromass LCT with time of flight detection and electrospray ionization.

#### 1.2. Synthesis of S-glycosides

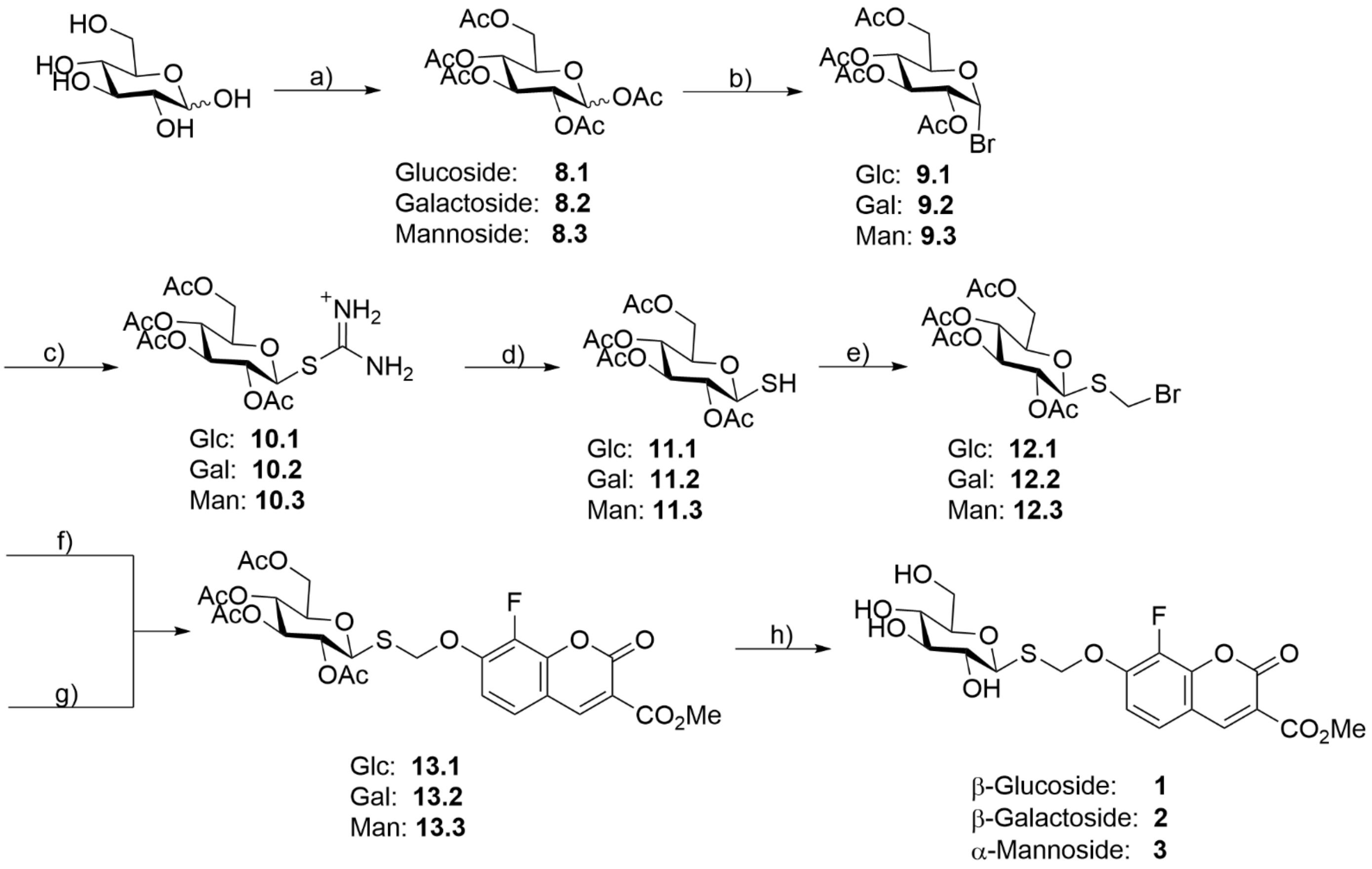

**Scheme S1:** Synthesis route for substrates **1-3** via 7 steps: a) $Ac_2O$ (5.0 eq), $I_2$ (0.01 eq), rt, 16 h or $Ac_2O$ (15 eq), pyridine (15 eq), 16 h **8.1**: 92%, **8.2**: 96%,**8.3**: 94%; b) 33% HBr in AcOH (38.0 eq), $Ac_2O$ (5.0 eq), 0 °C → 4 °C, 16 h, DCM, **9.1**: 87%,**9.2**: 91%, **9.3**: 86% ; c) Thiourea (1.4 eq), rt, 16 h, MeCN, **10.1**: 47%, **10.2**: 52%, **10.3**: 56%, d) $Na_2S_2O_5$ (5.0 eq), 60 °C, 1 h, DCM/$H_2O$; e) $CH_2Br_2$ (580 eq), DIPEA (1.0 eq), 80 °C, 16 h, **12.1**: 82%, **12.2**: 41%, **12.3**: 86%, over two steps; f) Jericho blue-OMe (1.0 eq), DIPEA (1.10 eq), 55 °C, 10 d, MeCN, **13.1**: 67% , **13.2**: 63%, **13.3**: 74%; g) TBA-Jericho blue-OMe (1.0 eq), NaI (0.2 eq), 55 °C, 9 d, MeCN, **13.1**: 71%, **13.2**: 65%, **13.3**: 75%; h) NaOMe (0.5 M) (catalytic amounts), 0 °C → rt, 16 h, DCM/MeOH, **1**: 92%, **2**: 89%, **3**: 81%.

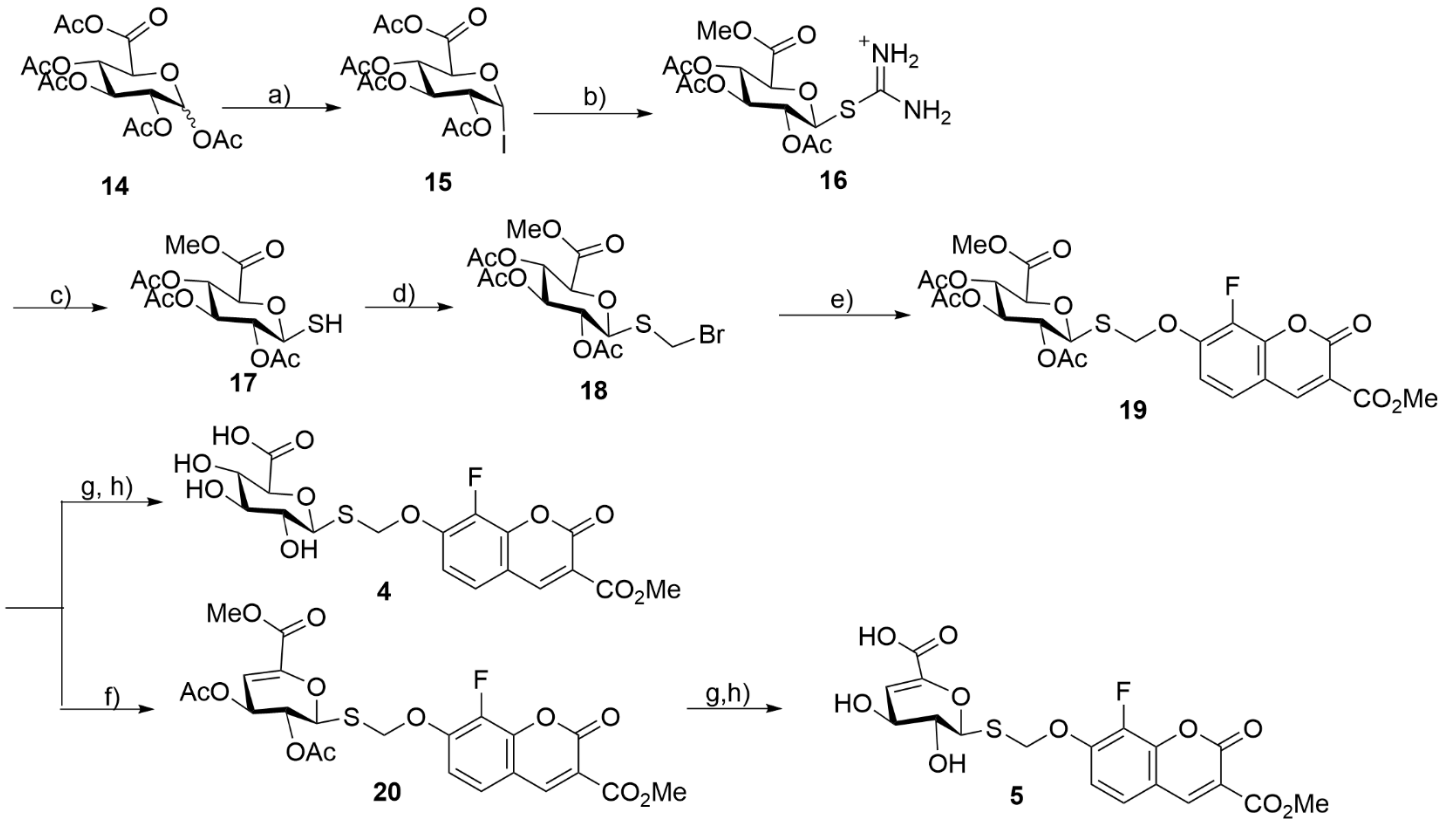

**Scheme S2:** Synthesis route for substrate **4**-**5**: a) PMHS (1.4 eq), $I_2$ (1.4 eq), 50°C, 1 h, DCE, 47%; b) Thiourea (1.4 eq), rt, 16 h, MeCN, 56%; c) $Na_2S_2O_5$ (5.0 eq), 60 °C, 1 h, DCM/$H_2O$; d) $CH_2Br_2$ (580 eq), DIPEA (1.0 eq), 80 °C, 16 h, 50%; e) Jericho blue-OMe (1.0 eq), DIPEA (1.1 eq), 55 °C, 10 d, MeCN, 32%; f) DBU (1.5 eq), rt, 4 h, DCM, 57%; g) NaOMe (0.5 M) (catalytic amounts), 0 °C → rt, 16 h, DCM/MeOH; h) LiOH (0.01M, 1.0 eq), rt, 10 min., THF **4**: 90%, **5**: 95%, over two steps.

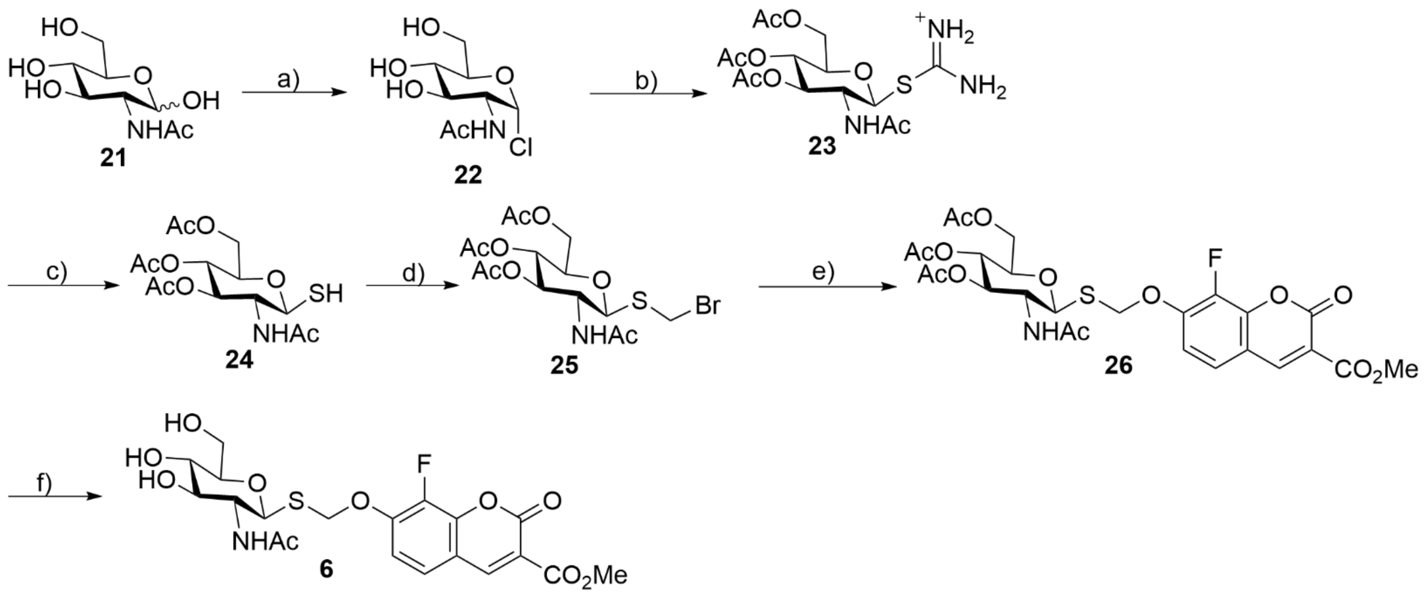

**Scheme S3:** Synthesis route for substrate **6** via 6 steps: a) $CH_3COCl$ (9.0 eq), 60 °C (15 min) → rt, 16 h, 90%; b) Thiourea (1.4 eq), rt, 16 h, MeCN, 81% c) $Na_2S_2O_5$ (5.0 eq), 60 °C, 1 h, DCM/$H_2O$; d) $CH_2Br_2$ (580 eq), DIPEA (1.0 eq), 80 °C, 16 h, 50% over two steps; e) Jericho blue-OMe (1.0 eq), DIPEA (1.1 eq), 55 °C, 10 d, MeCN, 99%; f) NaOMe (0.5 M) (catalytic amounts), 0 °C → rt, 16 h, DCM/MeOH, 97%.

##### 1.2.1. General procedures for synthesis of common starting materials

Global protection of sugars:
**Method I)** The sugar (1.00 eq) was dissolved in $Ac_2O$ (5.00 eq). Iodine (0.01 eq) was added and the mixture stirred at rt for 16 h. Afterwards, the reaction mixture was diluted with DCM and washed with 1 M HCl (3x), water and brine[1]. The product was used without any further purification.
**Method II)** To a suspension of the sugar (1 eq) in pyridine (10-15 eq) was added $Ac_2O$ (12-15 eq) and the mixture was stirred overnight. When TLC indicated the end of the reaction, the reaction mixture was concentrated by evaporation and the residual pyridine was co-evaporated with toluene. The mixture was dissolved in EtOAc and washed with 1 M solution of HCl, water, saturated $NaHCO_3$ solution and brine. The organic layers were dried and concentrated to yield the product, which was used without any further purification.

Synthesis of Glycosyl halides:
*Glycosyl bromides:*
The globally protected sugar was dissolved in DCM. 33% HBr in AcOH (38.0 eq) and $Ac_2O$ (9.0 eq) were added at 0 °C and the mixture was stirred for a few hours at 4 °C. When TLC indicated the consumption of starting material, the reaction was quenched with ice water. The aqueous phase was extracted with DCM (3x). The combined organic layers were washed with ice-cold water, saturated solution of $NaHCO_3$ (3x) and brine. The organic layer was dried over $MgSO_4$ and evaporated under reduced pressure to yield a white foam. The product was used without any further purification.

*Glycosyl iodide[2]:*
The globally protected β-D-glucopyranuronic acid was dissolved in DCE. $I_2$ (1.4 eq) and PMHS (Poly(methylhydrosiloxane)) (1.4 eq) were added and the mixture was refluxed at 80 °C for 1 h. Subsequently the mixture was diluted with DCM and the organic layer was washed with ice-cold solution of $NaHCO_3$ and $Na_2S_2O_3$. The aqueous phase was re-extracted with DCM (2x). The combined organic phases were washed with ice-cold brine, dried over $MgSO_4$, filtered and evaporated. The crude product was purified by chromatography on a short silica column using 2:1 PE/EtOAc as eluant.

*Glycosyl chloride:*
*N*-Acetyl-D-glucosamine (1.0 eq) was dissolved in acetyl chloride (9.0 eq). The mixture was refluxed for 15 min, then stirred at rt for 16 h. Subsequently the reaction was diluted with DCM and quenched with ice water. The aqueous phase was extracted with DCM (3x). The combined organic layers were washed with ice-cold $H_2O$, brine and saturated $NaHCO_3$ solution, dried over $MgSO_4$, filtered and evaporated. The product was used without any further purification.

##### 1.2.2. Synthesis of the thiourea-glycosides

The glycosyl halide (1.0 eq) was dissolved in MeCN (c = 0.85 M). Thiourea (1.4 eq) was added and the reaction was stirred for 16 h at 60 °C. The reaction mixture was concentrated by evaporation followed by addition of ethyl acetate that precipitates out most of the product from the solution as a white solid. The resulting products were found to be sufficiently pure in most cases and were therefore used in the next step without further purification.

*2,3,4,6-Tetra-O-acetyl-β-D-glucosyl-1-isothiouronium bromide[3]* **10.1**

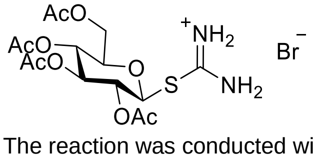

The reaction was conducted with 2.00 g (4.88 mmol) of **9.1** and yielded 1.11 g (2.28 mmol, 47%) **10.1** as white solid.
$R_f$ = 0.03 (PE/EtOAc 1:1), visualization with UV and molybdate.
**HRMS** calculated for $C_{15}H_{23}N_2O_9S^+$ ($[M]^+$): 407.1124, found: 407.1122
**$^1$H NMR** (400 MHz, $D_2O$) δ 5.50 (d, *J* = 10.0 Hz, 1H, H-1), 5.48 (t, *J* = 9.2 Hz, 1H), 5.40 – 5.31 (m, 1H), 5.25 (t, *J* = 9.7 Hz, 1H), 4.42 (dd, *J* = 12.8, 4.4 Hz, 1H), 4.30 (dd, *J* = 12.7, 2.2 Hz, 1H), 4.27 – 4.20 (m, 0H), 2.16 (s, 3H), 2.15 (s, 3H), 2.12 (s, 3H), 2.10 (s, 2H).
**$^{13}$C NMR** (101 MHz, $D_2O$) δ 173.5, 172.9, 172.6, 172.4, 167.5, 81.1, 75.8, 73.3, 69.1, 67.5, 61.8, 20.1, 20.0 (two peaks), 19.9.
*2,3,4,6-Tetra-O-acetyl-β-D-galactosyl-1-isothiouronium bromide[3]* **10.2**

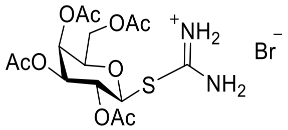

The reaction was performed with 800 mg (1.95 mmol) of **9.2** and 492 mg (1.01 mmol, 52%) **10.2** was obtained as a white solid.
***R*f** = 0.03 (PE/EtOAc 1:1), visualization with UV and molybdate.
**HRMS** calculated for $C_{15}H_{23}N_2O_9S+$ ([M]+): 407.1124, found: 407.1123
**$^1$H NMR** (400 MHz, Deuterium Oxide) δ 5.59 (d, *J* = 3.2 Hz, 1H, H-4 ), 5.51 (t, *J* = 9.8 Hz, 1H, H-2), 5.41 (d, *J* = 10.0 Hz, 1H, H-1), 5.35 (dd, *J* = 9.7, 3.2 Hz, 1H, H-3), 4.44 (t, *J* = 6.1 Hz, 1H, H-5), 4.29 (dd, *J* = 6.1, 2.8 Hz, 2H, H-6), 2.23 (s, 3H, OAc), 2.16 (s, 3H, OAc), 2.11 (s, 3H, OAc), 2.04 (s, 3H, OAc).

**$^{13}$C NMR** (101 MHz, D2O) δ 173.4, 172.8, 172.8, 172.4, 167.7, 81.4, 75.4, 71.5, 67.8, 66.6, 62.1, 20.5, 20.1, 20.0, 20.0.

*2,3,4,6-Tetra-O-acetyl-α-D-mannosyl-1-isothiouronium bromide*[4] **10.3**

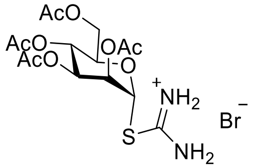

The reaction was conducted with 2.00 g (4.88 mmol) of **9.3** and yielded 1.33 g (2.73 mmol, 56%) of **10.3** as a white solid.
***R*f** = 0.03 (PE/EtOAc 1:1), visualization with UV and molybdate.
**HRMS** calculated for $C_{15}H_{23}N_2O_9S^+$ ($[M]^+$): 407.1124, found: 407.1127
**$^1$H NMR** (400 MHz, Deuterium Oxide) δ 6.34 (d, *J* = 1.7 Hz, 1H, H-1), 5.56 (dd, *J* = 3.3, 1.8 Hz, 1H, H-2), 5.34 (t, *J* = 9.7 Hz, 1H, H-4), 5.27 (dd, *J* = 9.8, 3.3 Hz, 1H, H-3), 4.56 (dddd, *J* = 29.2, 9.9, 5.1, 2.3 Hz, 1H, H-5), 4.41 (dd, *J* = 12.5, 5.2 Hz, 1H, H-6), 4.30 (dd, *J* = 12.7, 2.4 Hz, 1H, H-6), 2.24 (s, 3H, OAc), 2.14 (d, *J* = 2.8 Hz, 6H, 2 OAc), 2.06 (s, 3H, OAc).

**$^{13}$C NMR** (101 MHz, D2O) δ 182.0, 173.4, 172.4, 172.0, 168.0, 82.2, 71.1, 69.4, 69.4, 65.7, 62.2, 20.7, 20.6 (two signals), 20.5.

*Methyl (2,3,4-tri-O-acetyl-β-D-glucopyranosyl-1-isothiouronium iodide)uronate* **16**

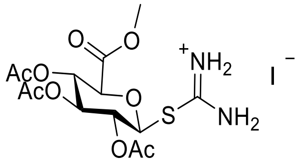

The reaction was performed with 1.10 g (2.48 mmol) of **15** and yielded 720 mg (1.38 mmol, 56%) **16** as a white solid.
***R*f** = 0.02 (PE/EtOAc 1:1), visualization with UV and molybdate.
**HRMS** calculated for $C_{14}H_{21}N_2O_9S^+$ ($[M]^+$): 393.0968, found: 393.0973
**NMR** spectra correspond to those presented in *Nasseri* et al.[5]

*2,3,4,6-Tetra-O-acetyl-β-D-2-acetamido-2-deoxy-glucopyranosyl-1-isothiouronium chloride*[3] **23**

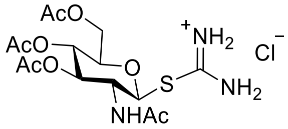

2.00g (5.06 mmol) of compound **22** was used as starting material. The reaction yielded 1.81 g (4.10 mmol, 81%) of **23** as white solid.
***R*f** = 0.01 (PE/EtOAc 1:2), visualization with UV and molybdate.
**HRMS** calculated for $C_{15}H_{24}N_3O_8S^+$ ($[M]^+$): 406.1284, found: 406.1280
**$^1$H NMR** (400 MHz, Deuterium Oxide) δ 5.50 (d, *J* = 10.6 Hz, 1H, H-1), 5.39 (t, *J* = 9.7 Hz, 1H, H-3), 5.20 (t, *J* = 9.7 Hz, 1H, H-4), 4.43 (dd, *J* = 12.7, 4.5 Hz, 1H, H-6), 4.37 (t, *J* = 10.3 Hz, 1H, H-2), 4.31 (dd, *J* = 12.8, 2.3 Hz, 1H, H-6), 4.24 (ddd, *J* = 10.3, 4.6, 2.3 Hz, 1H, H-5), 2.16 (s, 3H, OAc), 2.13 (s, 3H, OAc), 2.10 (s, 3H, OAc), 2.03 (s, 3H, NHAc).
**$^{13}$C NMR** (101 MHz, $D_2O$) δ 174.8, 173.8, 173.2, 172.8, 168.0, 82.2, 76.0, 73.1, 68.2, 62.1, 52.0, 22.1, 20.4, 20.3, 20.2.

##### 1.2.3. Synthesis of 1-thiosugars

The starting material (1.0 eq) was dissolved in DCM/$H_2O$ 1:1 (c = 0.15 M). $Na_2S_2O_5$ (5.0 eq) was added and the biphasic mixture was stirred vigorously and heated to 60 °C for 1 h. The reaction mixture was cooled to rt and diluted with DCM. The aqueous phase was extracted with DCM (3x). The combined organic layers were dried over $MgSO_4$, filtered and evaporated. No further purification was performed and the products were directly used for the next step since the resulting thiols are prone to oxidation and formation of dimers.

*2,3,4,6-Tetra-O-acetyl-1-β-thio-D-glucose* **11.1**

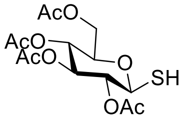

1.00 g (2.05 mmol) of compound **10.1** was reduced to compound **11.1**, which was used in the following step without any further purification.
***R*f** = 0.29 (PE/EtOAc = 2:1), visualization with molybdate.
**HRMS** calculated for $C_{14}H_{20}O_9SNa$ ($[M+Na]^+$): 387.0726, found: 387.0725

*2,3,4,6-Tetra-O-acetyl-1-β-thio-D-galactose* **11.2**

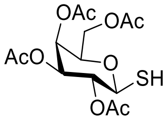

340 mg (699 µmol) of compound **10.2** was reduced to compound **11.2**, which was used in the following step without any further purification.
***R*f** = 0.29 (PE/EtOAc = 2:1), visualization with molybdate.
**HRMS** calculated for $C_{14}H_{20}O_9SNa$ ($[M+Na]^+$): 387.0726, found: 387.0723

2,3,4,6-Tetra-O-acetyl-1-α-thio-D-mannose **11.3**

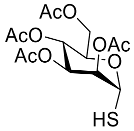

100 mg (205 µmol) of compound **10.3** was reduced to compound **11.3**, which was used in the following step without any further purification.
***R*f** = 0.29 (PE/EtOAc = 2:1), visualization with molybdate.
**HRMS** calculated for $C_{14}H_{20}O_9SNa$ ($[M+Na]^+$): 387.0726, found: 387.0724

*Methyl (2,3,4-tri-O-acetyl-1-β-thio-D-glucopyranoside)uronate* **17**

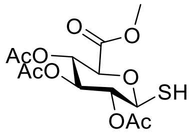

678 mg (1.30 mmol) of compound **16** was reduced to **17**. This compound was used in the following step without any further purification.
***R*f** = 0.28 (PE/EtOAc = 2:1), visualization with molybdate.
**HRMS** calculated for $C_{13}H_{18}O_9SNa$ ($[M+Na]^+$): 373.0569, found: 373.0567

*2,3,4,6-Tetra-O-acetyl-1-β-thio-D-(Acetylamino)-2-deoxy-glucopyranose* **24**

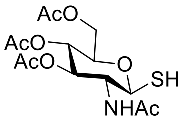

618 mg (1.40 mmol) of compound **23** was reduced to **24**, which was used in the following step without any further purification.
***R*f** = 0.27 (PE/EtOAc = 2:1), visualization with molybdate.
**HRMS** calculated for $C_{14}H_{21}NO_8SNa$ ($[M+Na]^+$): 386.0886, found: 386.0887

##### 1.2.4. Introduction of the *S,O*-acetal linker

The thiosugar (1.0 eq) was dissolved in degassed $CH_2Br_2$ (580 eq). DIPEA (1.0 eq) was added and the reaction mixture was stirred for 16 h at 80 °C (reflux). Afterwards, the reaction mixture was evaporated and the product was purified via column chromatography (PE/EtOAc, 2:1).

*Bromomethylthio 2,3,4,6-tetra-O-acetyl-β-D-glucopyranoside* **12.1**

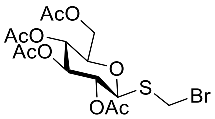

896 mg of crude **11.1** was used for the reaction and yielded 769 mg (1.69 mmol, 82%, over two steps) of **12.1** as a light-yellow foam.
***R*f** = 0.34 (PE/EtOAc 1:2), visualization with UV and molybdate.
**HRMS** calculated for $C_{15}H_{21}BrO_9SNa$ ($[M+Na]^+$): 478.9987, found: 478.9985
**NMR** spectra were found to match those presented in *Qing* et al.[6]

*Bromomethylthio 2,3,4,6-tetra-O-acetyl-β-D-galactopyranoside* **12.2**

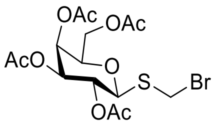

304 mg of crude **11.2** was used. This reaction yielded 131 mg (287 µmol, 41%, over two steps) of **12.2** as a light-yellow foam.
***R*f** = 0.34 (PE/EtOAc 1:2), visualization with UV and molybdate.
**HRMS** calculated for $C_{15}H_{21}BrO_9Sna$ ($[M+Na]^+$): 478.9987, found: 478.9986
**$^1$H NMR** (400 MHz, Acetone-d$_6$) δ 5.47 (dd, *J* = 3.4, 1.2 Hz, 1H, H-4), 5.28 (dd, *J* = 9.9, 3.4 Hz, 1H, H-3), 5.20 (t, J = 9.9 Hz, 1H, H-2), 5.06 (d, *J* = 9.9 Hz, 1H, H-1), 4.95 (d, J = 10.9 Hz, 1H, $CH_2$), 4.89 (d, *J* = 10.9 Hz, 1H, $CH_2$), 4.34 (ddd, *J* = 7.2, 6.3, 1.2 Hz, 1H, H-6), 4.19 (dd, *J* = 11.4, 6.7 Hz, 1H, H-5), 4.13 (dd, *J* = 11.3, 6.1 Hz, 1H, H-6), 2.14 (s, 3H, Oac), 2.03 (s, 3H, Oac), 2.00 (s, 3H, Oac), 1.93 (s, 3H, Oac).

**$^{13}$C NMR** (101 MHz, Acetone) δ 170.0, 169.8, 169.4, 169.3, 82.4, 74.7, 71.6, 67.7, 67.4, 61.5, 33.2, 19.9, 19.9, 19.8, 19.8.

*Bromomethylthio 2,3,4,6-tetra-O-acetyl-α-D-mannopyranoside* **12.3**

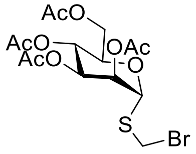

75.7 mg of **11.3** was used and yielded 80.4 mg (176 µmol, 86%, over two steps) of **12.3** as a light-yellow foam.
***R*f** = 0.34 (PE/EtOAc 1:2), visualization with UV and molybdate.
**HRMS** calculated for $C_{15}H_{21}BrO_9Sna$ ($[M+Na]^+$): 478.9987, found: 478.9988

**$^1$H NMR** (400 MHz, Acetone-$d_6$) δ 5.59 (d, *J* = 1.5 Hz, 1H, H-1), 5.37 – 5.27 (m, 2H, H-2, H-4), 5.12 (dd, *J* = 10.1, 3.5 Hz, 1H, H-3), 4.92 (d, *J* = 11.0 Hz, 1H, $CH_2$), 4.81 (d, *J* = 11.0 Hz, 1H, $CH_2$), 4.30 – 4.22 (m, 2H, H-6), 4.14 (d, *J* = 9.9 Hz, 1H, H-5), 2.14 (s, 3H, Oac), 2.03 (s, 3H, Oac), 2.02 (s, 3H, Oac), 1.94 (s, 3H, Oac).

**$^{13}$C NMR** (101 MHz, Acetone) δ 170.0, 169.6, 169.6, 169.4, 81.2, 70.1, 69.9, 69.7, 65.8, 62.1, 32.9, 20.0, 19.9 (two signals), 19.8.

Methyl (bromomethylthio 2,3,4-tri-O-acetyl-β-D-glucopyranoside)12rinate **18**

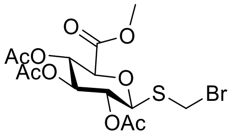

602 mg of **17** was used as starting material. This reaction yielded 429 mg (971 µmol, 75%, over two steps) of **18** as light-yellow foam.
***R*f** = 0.33 (PE/EtOAc 1:2), visualization with UV and molybdate.
**HRMS** calculated for $C_{14}H_{19}BrO_9Sna$ ([M+Na]$^+$): 464.9831, found: 464.9830
**NMR** spectra correspond to the ones presented in *Nasseri* et al.[5]

Bromomethylthio 3,4,6-tri-O-acetyl-β-D-acetamido-2-deoxy-glucopyranoside **25**

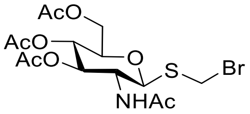

552 mg **24** was used as starting material and 318 mg (0.70 mmol, 50%, over two steps) **25** were obtained as white foam.
**Rf** = 0.30 (PE/EtOAc 1:2), visualization with UV and molybdate.
**HRMS** calculated for $C_{14}H_{22}BrNO_8Sna$ ([M+Na]+): 478.0147, found: 478.0144
**$^1$H NMR** (400 MHz, Acetone-$d_6$) δ 7.25 (d, *J* = 9.5 Hz, 1H, NH), 5.32 (dd, *J* = 10.2, 9.3 Hz, 1H, H-3), 5.07 (d, *J* = 10.7 Hz, 1H, H-1), 5.02 (t, *J* = 9.8, 9.1 Hz, 1H, H-4), 4.95 (d, *J* = 10.7 Hz, 1H, $CH_2$), 4.87 (d, *J* = 10.7 Hz, 1H, $CH_2$), 4.28 (dd, *J* = 12.3, 5.4 Hz, 1H, H-3), 4.20 – 4.08 (m, 2H, H-2, H-6), 3.90 (ddd, *J* = 10.1, 5.4, 2.4 Hz, 1H, H-5), 2.02 (s, 3H, Oac), 2.00 (s, 3H, Oac), 1.95 (s, 3H, Oac), 1.84 (s, 3H, Oac).

**$^{13}$C NMR** (101 MHz, Acetone) δ 169.9, 169.8, 169.4, 169.3, 82.6, 76.0, 73.7, 69.0, 62.2, 52.7, 33.5, 22.2, 19.9, 19.9, 19.8.

##### 1.2.5. Substitution of bromide with fluorophore

**Procedure A:** The protected thioglycoside (1.0 eq) was dissolved in MeCN (c = 0.02 M). Jericho blue-OMe (1.0 eq) and DIPEA (2.00 eq) were added and the reaction mixture was stirred until TLC indicated the consumption of starting material.
**Procedure B**: The protected thioglycoside (1.0 eq) was dissolved in MeCN (c = 0.02 M). TBA-Jericho blue-OMe[7] (1.0 eq) was added and the reaction mixture was stirred until TLC indicated the consumption of starting material.
Note: This reaction takes several days to complete. The reaction is faster when conducted in DMF instead of MeCN, or with addition of catalytic amounts of NaI, however the above conditions result in better yields.

*(3-Methylcarboxy-8-fluoro-7-hydroxycoumarin)methylthio 2,3,4,6-tetra-O-acetyl-β-D-glucoside* **13.1**

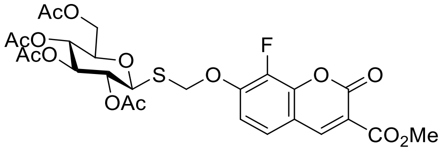

A) 150 mg (329 µmol) of compound **12.1** was used for this reaction. It yielded 136 mg (221 µmol, 67%) of the product **13.1** as a white solid.
B) 150 mg (329 µmol) of compound **12.1** was used for this reaction. It yielded 143 mg (234 µmol, 71%) of the product **13.1** as a white solid.
$\boldsymbol{R}_f$ = 0.58 (PE/EtOAc 1:3), visualization with UV and molybdate.
**HRMS** calculated for $C_{26}H_{27}FO_{14}SNa$ ([M+Na]$^+$): 637.1003, found: 637.0999
$^1$**H NMR** (400 MHz, Acetone-$d_6$) δ 8.64 (d, *J* = 1.4 Hz, 1H, H-4'), 7.68 (dd, $J_{H\text{-}H}$ = 8.9, 2.0 Hz, 1H, Ar), 7.32 (dd, $J_{H\text{-}H}$ = 8.9, $J_{H\text{-}F}$= 7.0 Hz, 1H, Ar), 5.76 (d, *J* = 12.2 Hz, 1H, $CH_2$), 5.64 (d, *J* = 12.2 Hz, 1H, $CH_2$), 5.31 (t, *J* = 9.4 Hz, 1H, H-3), 5.12 (d, *J* = 10.1 Hz, 1H, H-1), 5.07 (t, *J* = 9.8 Hz, 1H, H-4), 5.01 – 4.94 (m, 1H, H-2), 4.28 – 4.21 (m, 1H, H-6), 4.12 (dd, *J* = 12.3, 2.4 Hz, 1H, H-6), 4.08 – 4.03 (m, 1H, H-5), 3.86 (s, 3H, OMe), 2.01 (s, 3H, Oac), 1.99 (s, 3H, Oac), 1.94 (s, 3H, Oac), 1.86 (s, 3H, Oac).

$^{13}$**C NMR** (101 MHz, Acetone-$d_6$) δ 170.0, 169.5, 169.3, 169.0, 163.3, 154.8, 149.6 (d, $J_{C\text{-}F}$ = 7.6 Hz), 148.6 (d, $J_{C\text{-}F}$ = 2.6 Hz), 144.5 (d, $J_{C\text{-}F}$ = 8.9 Hz), 139.8 (d, $J_{C\text{-}F}$ = 249.6 Hz), 125.3 (d, $J_{C\text{-}F}$= 4.4 Hz), 116.0, 113.9, 112.8, 81.6, 75.8, 73.5, 70.5, 69.9, 68.4, 62.1, 52.0, 19.9, 19.9, 19.8, 19.7.
$^{19}$**F NMR** (377 MHz, Acetone-$d_6$) δ -155.80.

*(3-Methylcarboxy-8-fluoro-7-hydroxycoumarin)methylthio 2,3,4,6-tetra-O-acetyl-β-D-galactoside* **13.2**

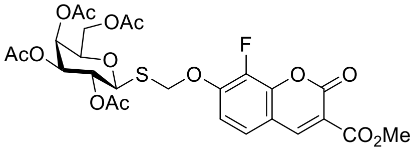

A) This procedure was executed with 70 mg (154 µmol) of **12.2** and yielded 60 mg (97.7 µmol, 63%) of the desired compound **13.2** as a white solid.
A) 150 mg (329 µmol) of the starting material **12.2** were used and yielded 131 mg (214 µmol, 65%) of the desired compound **13.2** as a white solid.
$\boldsymbol{R}_f$ = 0.58 (PE/EtOAc 1:3), visualization with UV and molybdate.
**HRMS** calculated for $C_{26}H_{27}FO_{14}SNa$ ([M+Na]$^+$): 637.1003, found: 637.1001

$^1$**H NMR** (400 MHz, Acetone-$d_6$) δ 8.65 (d, *J* = 1.5 Hz, 1H, H-4'), 7.70 (dd, $J_{H\text{-}H}$ = 8.9, 2.0 Hz, 1H, Ar), 7.34 (dd, $J_{H\text{-}H}$ = 8.9, $J_{H\text{-}F}$= 7.1 Hz, 1H, Ar), 5.78 (d, *J* = 12.2 Hz, 1H, $CH_2$), 5.65 (d, *J* = 12.2 Hz, 1H, $CH_2$), 5.45 (dd, *J* = 3.1, 1.2 Hz, 1H, H-6), 5.20 – 5.19 (m, 1H, H-2), 5.16 (d, *J* = 9.6 Hz, 1H, H-6), 5.10 (dd, *J* = 8.8, 1.5 Hz, 1H, H-1), 4.32 (td, *J* = 6.5, 1.3 Hz, 1H, H-3), 4.10 (d, *J* = 6.4 Hz, 2H, H-4, H-5), 3.86 (s, 3H, OMe), 2.13 (s, 3H, Oac), 1.99 (s, 3H, Oac), 1.91 (s, 3H, Oac), 1.87 (s, 3H, Oac).

$^{13}$**C NMR** (101 MHz, Acetone-$d_6$) δ 170.0, 169.8, 169.4, 169.2, 163.3, 154.8, 149.6 (d, $J_{C\text{-}F}$ = 7.7 Hz), 148.6 (d, $J_{C\text{-}F}$ = 2.6 Hz), 144.5 (d, $J_{C\text{-}F}$ = 9.0 Hz), 139.8 (d, $J_{C\text{-}F}$ = 249.6 Hz), 125.3 (d, $J_{C\text{-}F}$ = 4.5 Hz), 116.0, 113.9, 112.9, 82.2, 74.6, 71.6, 70.1, 67.7, 67.7, 61.5, 52.0, 19.8, 19.8 (two signals), 19.8.
$^{19}$**F NMR** (377 MHz, Acetone-$d_6$) δ -156.07.

*(3-Methylcarboxy-8-fluoro-7-hydroxycoumarin)methylthio 2,3,4,6-tetra-O-acetyl-α-D-mannoside* **13.3**

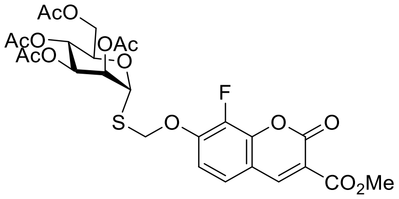

A) 70 mg (154 µmol) of the compound **12.3** yielded 70 mg (114 µmol, 74%) of the desired compound **13.3** as a white solid.
B) 100 mg (219 µmol) of the starting material **12.3** yielded 101 mg (164 µmol, 75%) of the desired compound **13.3** as a white solid.
***R*f** = 0.58 (PE/EtOAc 1:3), visualization with UV and molybdate.
**HRMS** calculated for $C_{26}H_{27}FO_{14}SNa$ ([M+Na]$^+$): 637.1003, found: 637.1002

**$^{1}$H NMR** (400 MHz, Acetone-$d_6$) δ 8.64 (d, *J* = 1.5 Hz, 1H, H-4'), 7.69 (dd, $J_{H\text{-}H}$ = 8.9, 2.0 Hz, 1H, Ar), 7.35 (dd, $J_{H\text{-}H}$= 8.9, $J_{H\text{-}F}$=7.0 Hz, 1H, Ar), 5.78 – 5.65 (m, 3H, H-1, H-3, H-4), 5.37 – 5.27 (m, 2H, H-2, $CH_2$), 5.17 (dd, *J* = 10.2, 3.4 Hz, 1H, $CH_2$), 4.34 – 4.22 (m, 2H, H-5, H-6), 4.15 – 4.08 (m, 1H, H-6), 3.86 (s, 3H, OMe), 2.13 (s, 3H, Oac), 2.03 (d, *J* = 0.9 Hz, 6H, 2 Oac), 1.94 (s, 3H, Oac).

**$^{13}$C NMR** (101 MHz, Methanol-$d_4$) δ 171.1, 170.8, 170.7, 170.5, 164.4, 155.9, 150.6 (d, $J_{C\text{-}F}$ = 7.8 Hz), 149.7 (d, $J_{C\text{-}F}$ = 2.6 Hz), 145.7, 139.8 (d, $J_{C\text{-}F}$ = 249.6 Hz), 126.6 (d, $J_{C\text{-}F}$ = 4.5 Hz), 117.2, 115.1, 113.5, 82.0, 71.3, 71.2, 70.5, 67.0, 63.2, 53.1, 21.1, 21.0 (two signals), 20.9.

**$^{19}$F NMR** (377 MHz, Methanol-$d_4$) δ -155.06.

*Methyl ((3-methylcarboxy-8-fluoro-7-hydroxycoumarin)methylthio 2,3,4-tri-O-acetyl-β-D-glucopyranoside)14rinate* **19**

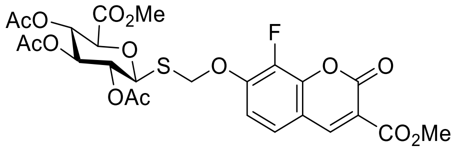

A) Using 170 mg (452 µmol) of compound **18** the reaction yielded 85 mg (141 µmol, 32%) of compound **19** as a white solid.
***R*f** = 0.57 (PE/EtOAc 1:3), visualization with UV and molybdate.
**HRMS** calculated for $C_{25}H_{25}FO_{14}SNa$ ([M+Na]$^+$): 623.0847, found: 623.0850
**$^{1}$H NMR** (400 MHz, Acetone-$d_6$) δ 8.66 (d, *J* = 1.5 Hz, 1H, H-4'), 7.69 (m, $J_{H\text{-}H}$ = 8.9, 2.0 Hz, 1H, Ar), 7.35 (dd, $J_{H\text{-}H}$ = 8.9, $J_{H\text{-}F}$ = 7.0 Hz, 1H, Ar), 5.77 (d, *J* = 12.2 Hz, 1H, $CH_2$), 5.68 (d, *J* = 12.2 Hz, 1H, $CH_2$), 5.38 (t, *J* = 9.4 Hz, 1H, H-3), 5.21 (d, *J* = 10.2 Hz, 1H, H-1), 5.12 (t, *J* = 9.8 Hz, 1H, H-4), 5.01 (dd, *J* = 10.2, 9.3 Hz, 1H, H-2), 4.41 (d, *J* = 10.0 Hz, 1H, H-5), 3.86 (s, 3H, OMe), 3.70 (s, 3H, OMe), 1.97 (s, 3H, Oac), 1.95 (s, 3H, Oac), 1.86 (s, 3H, Oac).

**$^{13}$C NMR** (101 MHz, Acetone-$d_6$) δ 169.2, 169.0, 168.8, 166.9, 163.1, 154.7, 149.3 (d, $J_{C\text{-}F}$ = 7.7 Hz), 148.5 (d, $J_{C\text{-}F}$ = 2.6 Hz), 139.7 (d, $J_{C\text{-}F}$ = 249.6 Hz), 125.1 (d, $J_{C\text{-}F}$ = 4.4 Hz), 115.9, 113.8, 112.8, 82.1, 75.5, 72.5, 70.1, 70.0, 69.4, 52.0, 51.8, 19.6 (two signals), 19.5.

**$^{19}$F NMR** (377 MHz, Acetone-$d_6$) δ -155.88.

*(3-Methylcarboxy-8-fluoro-7-hydroxycoumarin)methylthio 3,4,6-tri-O-acetyl-β-D-2-acetamido-2-deoxy-glucopyranoside* **26**

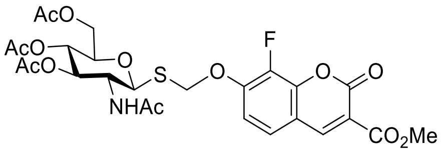

A) Starting with 70 mg (154 µmol) of compound **25** the reaction yielded 96 mg (152 µmol, 99%) of the desired compound **26** in form of a white solid**.**
***R*f** = 0.54 (PE/EtOAc 1:3), visualization with UV and molybdate.
**HRMS** calculated for $C_{26}H_{28}FNO_{13}Sna$ ([M+Na]$^+$): 636.1163, found: 636.1163
**$^1$H NMR** (400 MHz, Acetone-$d_6$) δ 8.67 (s, 1H, H-4'), 7.70 (d, *J* = 8.8 Hz, 1H, NH), 7.35 (t, *J* = 8.0 Hz, 1H, Ar), 7.16 (d, *J* = 9.6 Hz, 1H, Ar), 5.82 (d, *J* = 12.2 Hz, 1H, $CH_2$), 5.58 (d, *J* = 12.2 Hz, 1H, $CH_2$), 5.24 (t, *J* = 9.7 Hz, 1H, H-3), 5.11 (d, *J* = 10.6 Hz, 1H, H-1), 5.01 (t, *J* = 9.8 Hz, 1H, H-4), 4.23 (dd, *J* = 12.3, 5.1 Hz, 1H, H-2), 4.13 – 4.06 (m, 2H, H-5, H-6), 3.86 (s, 3H, OMe), 2.77 (s, 1H, H-6), 1.99 (d, *J* = 6.4 Hz, 6H, 2 Oac), 1.93 (s, 3H, Oac), 1.73 (s, 3H, NHAc).

**$^{13}$C NMR** (101 MHz, Acetone-$d_6$) δ 170.0, 169.8, 169.3, 169.2, 163.3, 154.8, 149.8 (d, $J_{C\text{-}F}$ = 7.3 Hz), 148.6 (d, $J_{C\text{-}F}$ = 2.8 Hz), 144.5, 139.8 (d), 125.3 (d, $J_{C\text{-}F}$ = 4.4 Hz), 115.9, 113.8, 112.9, 82.5, 75.9, 73.7, 69.7, 68.9, 62.3, 53.2, 51.9, 22.1, 22.1, 19.9, 19.8.

**$^{19}$F NMR** (377 MHz, DMSO-$d_6$) δ -155.66.

##### 1.2.6. Elimination of the C4-acetyl group using DBU:

*Methyl((3-methylcarboxy-8-fluoro-7-hydroxycoumarin)methylthio 2,3-di-O-acetyl-4-deoxy-α-L-threo-hex-4-enopyranosid)uronate* **20**

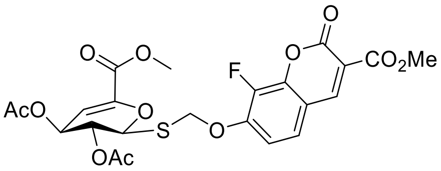

160 mg (1.00 eq, 267 µmol) of compound **19** was dissolved in DCM (c = 0.05 M). 59.8 µL DBU (1.5 eq, 400 µmol) were added dropwise and the reaction mixture was stirred until TLC indicated the starting material was consumed (approx. 4 h). The reaction mixture was then loaded directly on a silica column and purified by flash column chromatography (PE/EtOAc, 1:3). The reaction yielded 82 mg (152 µmol, 57%) of compound **20** as a white solid.
***R*f** = 0.54 (PE/EtOAc 1:3), visualization with UV and molybdate.
**HRMS** calculated for $C_{23}H_{21}FO_{12}SNa$ ([M+Na]$^+$): 563.0635 , found: 563.0631
**$^1$H NMR** (400 MHz, Acetone-$d_6$) δ 8.65 (d, *J* = 1.5 Hz, 1H, H-4'), 7.68 (dd, $J_{H\text{-}H}$ = 8.9, 2.0 Hz, 1H, Ar), 7.35 (dd, $J_{H\text{-}H}$ = 8.8, , $J_{H\text{-}F}$ = 7.0 Hz, 1H, Ar), 6.22 – 6.18 (m, 1H, H-1), 5.95 (dd, *J* = 2.4, 1.6 Hz, 1H, H-3), 5.76 (d, *J* = 12.1 Hz, 1H, $CH_2$), 5.67 (d, *J* = 12.0 Hz, 1H, $CH_2$), 5.14 (dd, *J* = 4.0, 1.5 Hz, 2H, H-2, H-4), 3.86 (s, 3H, OMe), 3.79 (s, 3H, OMe), 2.06 (s, 3H, OAc), 2.04 (s, 3H, OAc).
**$^{13}$C NMR** (101 MHz, Acetone-$d_6$) δ 169.0, 168.9, 163.1, 161.6, 154.7, 149.3 (d, $J_{C\text{-}F}$ = 7.7 Hz), 148.5 (d, $J_{C\text{-}F}$ = 2.5 Hz), 144.4 (d, $J_{C\text{-}F}$ = 8.9 Hz), 143.2, 139.7 (d, $J_{C\text{-}F}$ = 249.6 Hz), 125.3 (d, $J_{C\text{-}F}$ = 4.4 Hz), 115.9, 113.8, 112.6, 106.7, 78.8, 70.8, 68.6, 63.7, 52.0, 51.9, 19.8, 19.8.
**$^{19}$F NMR** (377 MHz, Acetone-$d_6$) δ -155.82.

##### 1.2.7. Deprotection of the compounds

**Zemplén deprotection:**

The globally protected sugar compound was dissolved in DCM/methanol 1:1 (c = 0.01 M) and cooled to 0 ºC. A catalytic amount of freshly prepared sodium methoxide was added, and the reaction mixture was stirred until TLC indicated the end of the reaction. The reaction mixture was neutralized with Amberlite IR-120 ($H^+$ form) and filtered. The solvents were evaporated to yield the product.

Note: Depending on the purity of the resulting product, the resulting glycosides might need to be purified. This can be achieved via precipitation of the product from methanolic solutions using ice cold $Et_2O$, or alternatively using flash column chromatography.

**Hydrolysis of the methyl ester protecting group of the uronic acids using aqueous LiOH solution:**

The de-acetylated uronic acid compound resulting from Zemplén deprotection was dried then dissolved in THF/d$H_2O$ 1:1 (c = 0.01 M), 1 M LiOH (1.1 eq) were added and the reaction mixture was stirred until TLC indicated the end of the reaction (typically 10 min). The reaction mixture was neutralized with Amberlite IR-120 ($H^+$ form) and filtered. The solvents were evaporated to yield the product.

Note: The methyl ester of the Jericho-blue group is mostly left untouched in these conditions, however the small amount of byproduct with both methyl esters hydrolyzed was not readily separable from the main product. Since both the products with JBOMe and JB fluorophores are good screening substrates[7], we used the mixture in our experiments. The NMR spectra for compounds **4** and **5** is therefore reported both before the final methyl ester deprotection and for a small fraction of the final product that was separated from most of the impurities.

*(3-Methylcarboxy-8-fluoro-7-hydroxycoumarin)methylthio β-D-glucoside* ***1***

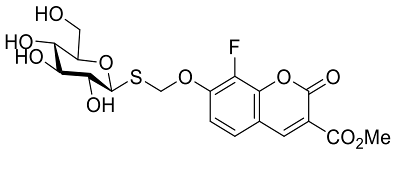

80.0 mg (130 µmol) of the substrate **13.1** were deprotected using the Zemplén method. This yielded 53.5 mg (120 µmol, 92%) of the desired substrate **1** as a white solid.

***R*f** = 0.13 (DCM/MeOH 8:1), visualization with UV and molybdate.

**HRMS** calculated for $C_{18}H_{19}FO_{10}SNa$ ([M+Na]$^+$): 469.0581, found: 469.0583

**$^1$H NMR** (400 MHz, DMSO-$d_6$) δ 8.70 (d, *J* = 3.2 Hz, 1H, H-4'), 7.65 (d, $J_{H\text{-}H}$ = 8.8 Hz, 1H, Ar), 7.30 (dd, $J_{H\text{-}H}$ = 8.8, $J_{H\text{-}F}$ = 7.0 Hz, 1H, Ar), 5.76 (d, *J* = 12.2, 1H, $CH_2$), 5.57 (d, *J* = 12.1, 1H, $CH_2$), 4.43 (d, *J* = 9.6 Hz, 1H, H-1), 3.79 (d, *J* = 1.9 Hz, 3H, H-6), 3.65 (s, 1H, H-6), 3.43 (dd, *J* = 12.1, 6.1 Hz, 1H, H-6), 3.14 (d, *J* = 8.5 Hz, 2H, H-4, H-5), 3.08 (d, *J* = 9.2 Hz, 1H, H-3), 3.02 (d, *J* = 9.0 Hz, 1H, H-2).

**$^{13}$C NMR** (101 MHz, DMSO-$d_6$) δ 163.5, 155.6, 150.0 (two carbon signals, one of them a doublet with $J_{C\text{-}F}$ = 7.3 Hz), 144.0 (d, $J_{C\text{-}F}$ = 8.6 Hz), 139.1 (d, $J_{C\text{-}F}$ = 246.4 Hz), 126.2 (d, $J_{C\text{-}F}$ = 3.9 Hz), 114.9, 113.4, 112.8, 83.1, 81.5, 78.1, 73.7, 70.2, 69.3, 61.5, 52.9.

**$^{19}$F NMR** (377 MHz, DMSO-$d_6$) δ -156.00.

*(3-Methylcarboxy-8-fluoro-7-hydroxycoumarin)methylthio β-D-galactoside* **2**

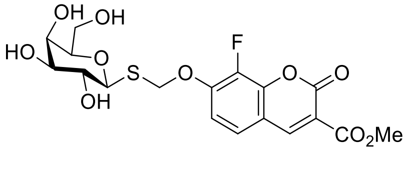

95.0 mg (155 µmol) of the substrate **13.2** were deprotected via Zemplén method. This yielded 61.5 mg (138 µmol, 89%) of the desired substrate **2** as a white solid.

$\boldsymbol{R_f}$ = 0.13 (DCM/MeOH 8:1), visualization with UV and molybdate.
**HRMS** calculated for $C_{18}H_{19}FO_{10}SNa$ ($[M+Na]^+$): 469.0581 , found: 469.0578
**1H NMR** (400 MHz, DMSO-d$_6$) δ 8.75 (d, *J* = 1.4 Hz, 1H, H-4'), 7.70 (dd, $J_{H\text{-}H}$ = 8.9, 1.8 Hz, 1H, Ar), 7.34 (dd, $J_{H\text{-}H}$ = 8.9, JH-F= 7.1 Hz, 1H, Ar), 5.75 (d, *J* = 12.2 Hz, 1H, $CH_2$), 5.59 (d, *J* = 12.1 Hz, 1H, $CH_2$), 4.41 (d, *J* = 9.4 Hz, 1H, H-1), 3.81 (s, 3H, OMe), 3.68 (d, *J* = 3.2 Hz, 1H, H-4), 3.48 (d, *J* = 6.6 Hz, 2H, H-6), 3.43 – 3.38 (m, 1H, H-5), 3.34 (d, *J* = 9.3 Hz, 1H, H-2), 3.28 (dd, *J* = 9.2, 3.2 Hz, 1H, H-3).

**$^{13}$C NMR** (101 MHz, DMSO-*d*$_6$) δ 163.7, 155.7, 150.2 (d, $J_{C\text{-}F}$ = 7.2 Hz), 149.9, 144.2 (d, $J_{C\text{-}F}$ = 9.1 Hz), 139.3 (d, $J_{C\text{-}F}$ = 247.7 Hz), 126.3 (d, $J_{C\text{-}F}$ = 4.2 Hz), 115.2, 113.6, 113.0, 84.0, 80.0, 74.9, 70.8, 69.7, 68.8, 61.0, 53.0.

**$^{19}$F NMR** (377 MHz, DMSO-*d*$_6$) δ -156.07.

*(3-Methylcarboxy-8-fluoro-7-hydroxycoumarin)methylthio α-D-mannoside **3***

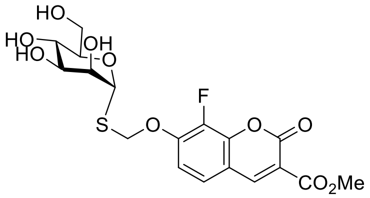

110 mg (179 µmol) of the substrate **13.3** were deprotected using the Zemplén method. This reaction yielded 64.7 mg (145 µmol, 81%) of the desired substrate **3** as a white solid.
$\boldsymbol{R_f}$ = 0.13 (DCM/MeOH 8:1), visualization with UV and molybdate.
**HRMS** calculated for $C_{18}H_{19}FO_{10}SNa$ ($[M+Na]^+$): 469.0581, found: 469.0586
**$^1$H NMR** (400 MHz, DMSO-*d*$_6$) δ 8.74 (d, *J* = 1.4 Hz, 1H, H-4'), 7.71 (dd, $J_{H\text{-}H}$ = 8.9, 1.8 Hz, 1H, Ar), 7.35 (dd, $J_{H\text{-}H}$= 8.9, $J_{H\text{-}F}$ = 7.1 Hz, 1H, Ar), 5.59 (d, *J* = 12.0 Hz, 1H, $CH_2$), 5.56 (d, *J* = 11.6 Hz, 1H, $CH_2$), 4.41 (d, *J* = 9.4 Hz, 1H, H-1), 3.81 (s, 3H, OMe), 3.68 (d, *J* = 3.2 Hz, 1H, H-4), 3.48 (d, *J* = 6.6 Hz, 2H, H-6), 3.43 – 3.38 (m, 1H, H-5), 3.34 (d, *J* = 9.3 Hz, 1H, H-2), 3.28 (dd, *J* = 9.2, 3.2 Hz, 1H, H-3).

**$^{13}$C NMR** (101 MHz, DMSO-*d*$_6$) δ 163.5, 155.5, 149.7 (d, $J_{C\text{-}F}$ = 6.8 Hz, two signals), 144.0 (d, $J_{C\text{-}F}$ = 8.4 Hz), 139.2 (d, $J_{C\text{-}F}$ = 248.0 Hz), 126.2, 115.2, 113.6, 112.8, 83.3, 75.6, 71.7, 71.6, 69.8, 67.2, 61.1, 52.9.

**$^{19}$F NMR** (377 MHz, DMSO-*d*$_6$) δ -156.36.

*(3-Methylcarboxy-8-fluoro-7-hydroxycoumarin)methylthio β-D-glucopyranoside)uronate **4***

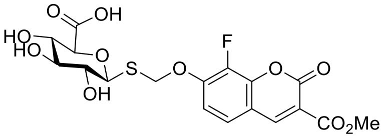

For deprotecting 150 mg (250 µmol) of starting material **19** the Zemplén method as well as hydrolysis of the methyl ester via LiOH were used. This yielded 104 mg (225 µmol, 90%, over two steps) of the desired substrate **4** as a white solid.
$\boldsymbol{R_f}$ = 0.12 (DCM/MeOH 8:1), visualization with UV and molybdate.
**HRMS** calculated for $C_{18}H_{18}FO_{11}S$ ($[M+H^+]$): 461.0554, found: 461.0551

**$^1$H NMR** for product before methyl ester deprotection (400 MHz, $D_2O$) δ 8.62 (t, *J* = 1.3 Hz, 1H, H-4'), 7.47 (dd, *J* = 8.8, 2.0 Hz, 1H, Ar), 7.18 (dd, *J* = 8.9, 6.9 Hz, 1H, Ar), 5.62 (d, *J* = 12.2 Hz, 1H, $CH_2$), 5.54 (d, *J* = 12.3 Hz, 1H, $CH_2$), 4.60 (d, *J* = 9.7 Hz, 1H, H-1), 3.83 (s, 3H, OMe), 3.80 (d, *J* = 9.8 Hz, 1H, H-5), 3.74 – 3.72 (m, 1H, H-2), 3.71 (s, 3H, OMe), 3.46 (dd, *J* = 9.8, 9.0 Hz, 1H, H-4), 3.32 (t, *J* = 8.9 Hz, 1H, H-3).

**$^{19}$F NMR** for product before methyl ester deprotection (377 MHz, $D_2O$) δ -155.00.

**$^1$H NMR** for final product (600 MHz, DMSO-d$_6$) δ 8.13 (s, 1H), 7.49 (d, *J* = 8.9 Hz, 1H), 7.22 (t, *J* = 8.1 Hz, 1H), 5.68 (d, *J* = 12.2 Hz, 1H), 5.52 (d, *J* = 12.2 Hz, 1H), 4.46 (d, *J* = 9.9 Hz, 1H), 3.49 – 3.41 (m, 2H), 3.23 (t, *J* = 9.3 Hz, 1H), 3.18 (t, *J* = 8.7 Hz, 1H), 3.05 (t, *J* = 9.2 Hz, 1H), 1.18 (s, 3H).
**$^{13}$C NMR** for final product (151 MHz, DMSO-d$_6$) δ 159.0, 147.4, 142.5, 138.91 (d, $J_{C\text{-}F}$ = 246.4 Hz), 124.0, 119.8, 119.0, 117.8, 115.9, 114.0, 112.3, 83.4, 78.4, 77.2, 72.8, 71.4, 69.6, 69.2.

*(3-Methylcarboxy-8-fluoro-7-hydroxycoumarin)methylthio 4-deoxy-α-L-threo-hex-4-enopyranosiduronic acid* (**5**)

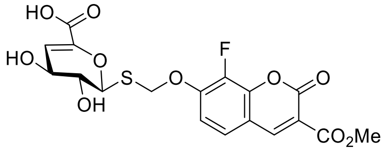

140 mg (260 µmol) of compound **20** were deprotected via the Zemplén method and the methyl ester hydrolyzed using LiOH. This yielded 105 mg (247 µmol, 95%, over two steps) of the desired substrate **5** as a white solid.
***R*$_f$** = 0.11 (DCM/MeOH 8:1), visualization with UV and molybdate.
**HRMS** calculated for $C_{18}H_{16}FO_{10}S$ ([M+H]$^+$): 443.0448, found: 443.0446

**$^1$H NMR** for product before methyl ester deprotection (600 MHz, MeOD) δ 8.73 (d, *J* = 1.3 Hz, 1H, H-4'), 7.58 (dd, *J* = 8.8, 1.8 Hz, 1H, Ar), 7.29 (dd, $J_{H\text{-}H}$ = 8.9, $J_{H\text{-}F}$ = 6.9 Hz, 1H, Ar), 6.17 (dd, *J* = 4.4, 1.1 Hz, 1H, H-4), 5.74 (d, *J* = 12.0 Hz, 1H, CH$_2$), 5.62 (dd, *J* = 4.0, 1.2 Hz, 1H, H-1), 5.52 (d, *J* = 12.0 Hz, 1H, CH$_2$), 4.03 (ddd, *J* = 4.5, 3.2, 1.2 Hz, 1H, H-3), 3.91 (s, 3H, OMe), 3.83 (ddd, *J* = 4.2, 3.2, 1.2 Hz, 1H, H-2), 3.79 (s, 3H, OMe).

**$^{13}$C NMR** for product before methyl ester deprotection (151 MHz, MeOD) δ 163.0, 162.4, 155.8, 149.6, 148.9, 140.5, 139.2 (d, $J_{C\text{-}F}$ = 250.5 Hz), 124.6, 114.2, 113.0, 112.3, 110.9, 81.0, 70.0, 69.9, 64.6, 51.2, 51.1.

**$^{19}$F NMR** for product before methyl ester deprotection (377 MHz, MeOD) δ -156.68.

**$^1$H NMR** for final product (400 MHz, MeOD-d$_4$) δ 8.72 (d, *J* = 1.5 Hz, 1H, H-4'), 7.57 (dd, *J* = 8.9, 2.0 Hz, 1H, Ar), 7.30 (dd, *J* = 8.9, 6.9 Hz, 1H, Ar), 6.19 (dd, *J* = 4.4, 1.2 Hz, 1H, H-4), 5.77 (d, *J* = 12.0 Hz, 1H, CH$_2$), 5.63 (dd, *J* = 4.0, 1.2 Hz, 1H, H-1), 5.51 (d, *J* = 12.0 Hz, 1H, CH$_2$), 4.04 (ddd, *J* = 4.4, 3.2, 1.2 Hz, 1H, H-3), 3.90 (s, 3H, OMe), 3.83 (ddd, *J* = 4.0, 3.3, 1.1 Hz, 1H, H-2)

**$^{13}$C NMR** for final product (101 MHz, MeOD-d$_4$) δ 165.4, 164.9, 157.6, 152.2, 151.5, 150.7, 145.6, 142.7, 139.7 (d, *J* = 36.8 Hz), 126.5 (d, *J* = 4.4 Hz), 116.1, 115.0, 114.2, 112.6, 82.6, 71.8, 71.8, 66.6, 53.1.

*(3-Methylcarboxy-8-fluoro-7-hydroxycoumarin)methylthio β-D-2-acetamido-2-deoxy-glucopyranoside* **6**

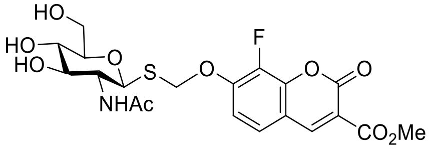

The Zemplén deprotection was started with 96 mg of compound **25** (152 µmol) and yielded 75 mg (148 µmol, 97%) of the desired substrate **6** as a white solid.
***R*$_f$** = 0.10 (DCM/MeOH 8:1), visualization with UV and molybdate.
**HRMS** calculated for $C_{20}H_{22}FNO_{10}SNa$ ([M+Na]$^+$): 510.0846, found: 510.0845
**$^1$H NMR** (400 MHz, DMSO-*d*$_6$) δ 8.72 (s, 1H, H-4'), 7.81 (d, *J* = 9.3 Hz, 1H, NH), 7.66 (d, $J_{H\text{-}H}$ = 8.8 Hz, 1H, Ar), 7.30 (d, $J_{H\text{-}H}$ = 8.9 Hz, 1H, Ar), 5.76 (d, *J* = 12.2 Hz, 1H, CH$_2$), 5.47 (d, *J* = 12.0 Hz, 1H, CH$_2$), 4.54 (d, *J* = 10.4 Hz, 1H, H-1), 3.80 (s, 3H, OMe), 3.57 (s, 1H, H-6), 3.52 (t, *J* = 9.6 Hz, 1H, H-2), 3.44 (dd, *J* = 12.0, 5.9 Hz, 1H, H-6), 3.27 (t, *J* = 9.0 Hz, 1H, H-3), 3.13 (d, *J* = 8.7 Hz, 2H, H-4, H-5), 1.68 (s, 3H, NHAc).

**$^{13}$C NMR** (101 MHz, DMSO-*d*$_6$) δ 170.1, 163.5, 155.6, 150.0 (d, $J_{C\text{-}F}$ = 7.4 Hz), 149.7, 144.0 (d, $J_{C\text{-}F}$ = 8.4 Hz), 139.1 (d, $J_{C\text{-}F}$ = 248.1 Hz), 126.1, 115.0, 113.5, 113.0, 82.6, 81.7, 75.4, 70.7, 69.9, 61.6, 55.0, 52.9, 23.2.

**$^{19}$F NMR** (377 MHz, DMSO-*d*$_6$) δ -155.62

##### 1.2.8. Enzymatic phosphorylation

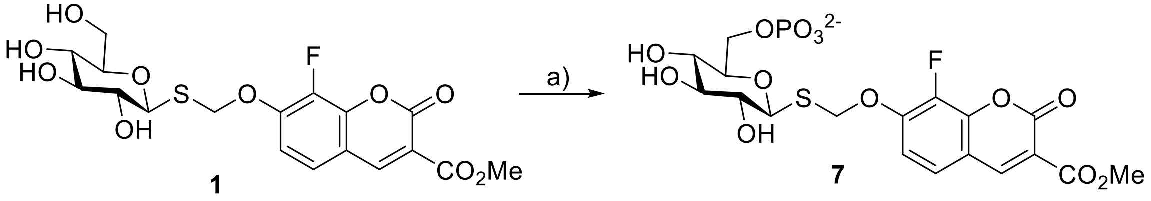

**Scheme S4:** Enzymatic synthesis of substrate **7**: a) BglK (1 mg/mL), ATP (2.0 eq), $MgCl_2$ (0.03 eq), rt, 2 h, ~3% DMSO in HEPES 25 mM pH 7.5, 90%

*(3-Methylcarboxy-8-fluoro-7-hydroxycoumarin)methylthio β-D-glucoside-6-phosphate* **7**

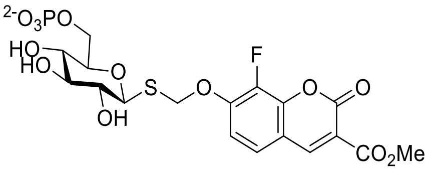

The starting material for this reaction was 28.0 mg (62.8 µmol) of substrate **1** which was dissolved in 500 µL of DMSO and added to 14.25 mL of HEPES buffer (pH 7.5, 25 mM) followed by 150 µL of $MgCl_2$ solution (1.0 M) and 83 mg of ATP (disodium salt, 150 µmol, 2.4 eq, pH of the solution might need adjustment after adding ATP). Finally, 100 µL of the enzyme BglK[8] (c = 1 mg/mL) was added and the reaction mixture was stirred at room temperature until completion (TLC) after 2 h. The solution was then lyophilized, and the product was purified via a Sep-Pak (C8) column. This reaction yielded 29.7 mg (56.5 µmol, 90%) of the desired substrate **7,** as a white solid.
***R*f** = 0.45 (t-BtOH/AcOH/$H_2O$ 5:2:3), visualization with UV and molybdate.
**HRMS** calculated for $C_{18}H_{19}FO_{13}PS$ ($[M-H]^-$): 525.0268, found: 525.0269
**$^1$H NMR** (400 MHz, Deuterium Oxide) δ 8.73 (d, *J* = 1.4 Hz, 1H, H-4'), 7.55 (dd, $J_{H-H}$ = 8.8, 1.8 Hz, 1H, Ar), 7.27 (dd, $J_{H-H}$ = 8.9, $J_{H-F}$ = 7.0 Hz, 1H, Ar), 5.76 (d, *J* = 12.2 Hz, 1H, $CH_2$), 5.63 (d, *J* = 12.2 Hz, 1H, $CH_2$), 4.77 (d, 1H, H-1), 4.09 (dt, *J* = 6.2, 3.0 Hz, 2H, H-6), 3.93 (s, 3H, OMe), 3.61 – 3.54 (m, 2H, H-5, H-4), 3.55 – 3.49 (m, 1H, H-3), 3.42 (dd, *J* = 9.8, 8.5 Hz, 1H, H-2).

**$^{13}$C NMR** (101 MHz, Deuterium Oxide) δ 164.9, 158.6, 151.4 (d, $J_{C-F}$ = 2.4 Hz), 150.0 (d, $J_{C-F}$ = 7.5 Hz), 143.7 (d, $J_{C-F}$ = 9.0 Hz), 139.4 (d, $J_{C-F}$ = 249.5 Hz), 126.1 (d, $J_{C-F}$= 4.2 Hz), 113.8, 113.7, 113.4, 84.3, 79.1 (d, $J_{C-P}$ = 8.0 Hz), 77.0, 72.6, 70.3, 68.9, 64.0 (d, $J_{C-P}$ = 4.9 Hz), 53.3, 38.9.

**$^{19}$F NMR** (377 MHz, Deuterium Oxide) δ -155.53.

**$^{31}$P NMR** (162 MHz, Deuterium Oxide) δ 1.55.

#### 1.3. Synthesis of 3-keto-glycosides and 3-keto-glucose analogues

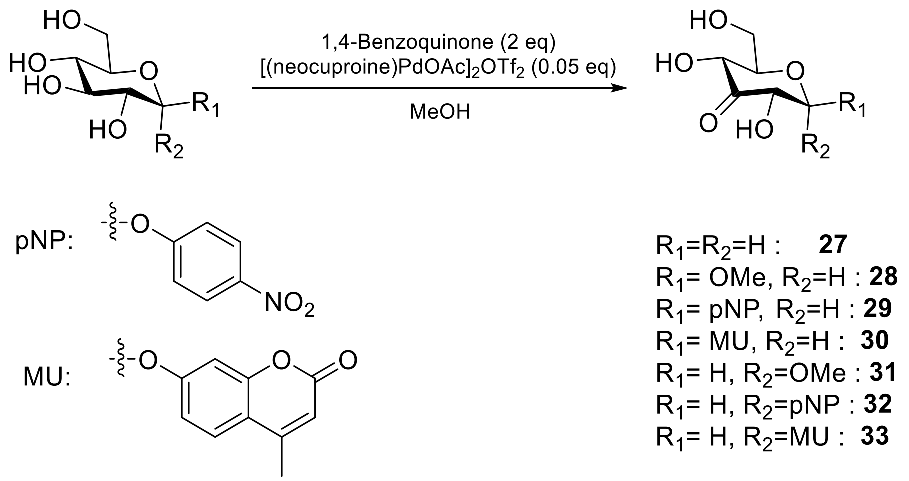

**Scheme S5:** Synthesis scheme for 3-keto glycoside substrates and 3-keto-1,5-anhydroglucitol

Based on the procedure from Jäger et al[9] with slight modifications, the glycoside starting material and 1,4-benzoquinone (0.8 to 2 eq.) were suspended in methanol (final concentration of glycoside: 0.03 M). To this mixture was added 0.05 eq of the catalyst, $[(neocuproine)PdOAc]_2OTf_2$ and the suspension was stirred at room temperature until TLC indicated the consumption of the starting material (typically a few hours). The reaction mixture was then dried *in vacuo*, and purified via flash column chromatography (dry loading, DCM/Acetone) to yield pure product. (For the products that are water soluble, most of the organic impurities can be separated by suspending the reaction mixture in water and filtering it over Celite, prior to the final purification by flash column chromatography).

Note: using more equivalents of benzoquinone or extending the reaction time leads to formation of over-oxidized by-products, notably as a result of oxidation at the C-6 position. If the starting material glycoside is readily available, it is desirable to use less than 1 eq. of benzoquinone, since separation of starting material and product is often easier than separation of product from undesirable by-products. In this case, reaction should be stirred for not more than 2 hours at room temperature.

Also note that some of these compounds exist as equilibria of the keto and hydrate forms in the presence of water. Where applicable, the reported NMR spectra therefore reflect the signals for these two forms which sometimes appear as unassignable peaks due to overlap of similar signals. Interestingly however, for compounds **31** to **33** that are 3-keto-α-glucosides, only the keto form is observed, since large groups in axial positions destabilize the hydrate form sufficiently that they are not readily observable by NMR.

*3-keto-1,5-anhydro-D-glucitol* ***27***

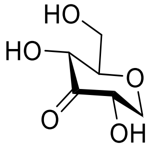

Compound **27** was synthesized from 1,5-anhydroglucitol (100 mg, 0.61 mmol) using the general procedure for synthesis of 3-keto-glycosides 1.8 to yield 80 mg of product (0.49 mmol, 81%).

$R_f$ = 0.4 (DCM/Acetone 1:4), visualization with molybdate stain.

**$^1$H NMR** (400 MHz, Acetone-$d_6$) δ 4.40 (ddd, *J* = 16.9, 7.0, 3.5 Hz, 2H, H-4 and OH), 4.32 – 4.21 (m, 2H, H-1 and H-2), 3.93 – 3.83 (m, 2H, H-6 and OH), 3.73 (ddd, *J* = 8.9, 6.6, 3.7 Hz, 1H, H-6), 3.34 – 3.22 (m, 2H, H-1 and H-5), 2.99 (s, 1H, OH).

**$^{13}$C NMR** (101 MHz, Acetone-$d_6$) δ 208.9, 85.7, 73.7, 73.2, 73.0, 62.8.

*Methyl β-D-ribo-hexopyranoside-3-ulose **28***

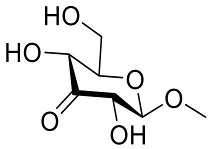

Compound **28** was synthesized from methyl β-glucopyranoside (500 mg, 2.57 mmol) using the general procedure for synthesis of 3-keto-glycosides to yield 300 mg of product (1.56 mmol, 61%).
The spectra were in agreement with those reported previously[9].
***R*f** = 0.4 (DCM/Acetone 1:4), visualization with molybdate stain.

**$^{1}$H NMR** (600 MHz, Acetone-$d_6$) δ 4.44 (d, *J* = 4.9 Hz, 1H, OH), 4.41 (d, *J* = 5.2 Hz, 1H, OH), 4.30 (d, *J* = 7.9 Hz, 1H, H-1), 4.31 – 4.26 (m, 1H, H-4), 4.13 (ddd, *J* = 7.8, 4.9, 1.9 Hz, 1H, H-2), 4.01 (t, *J* = 6.5 Hz, 1H, OH), 3.93 (ddd, *J* = 12.0, 5.9, 2.2 Hz, 1H, H-6), 3.79 (ddd, *J* = 11.8, 6.7, 4.9 Hz, 1H, H-6), 3.52 (s, 3H, OMe), 3.33 (ddd, *J* = 10.1, 4.9, 2.2 Hz, 1H, H-5).

**$^{13}$C NMR** (151 MHz, Acetone-$d_6$) δ 207.0, 106.4, 77.9, 77.7, 73.2, 62.3, 56.8.

*Para-nitrophenyl β-D-ribo-hexopyranoside-3-ulose **29***

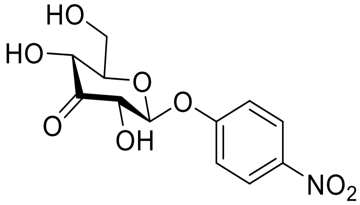

Compound **29** was synthesized from para-nitrophenyl β-glucopyranoside (215 mg, 0.71 mmol) using the general procedure for synthesis of 3-keto-glycosides to yield 170 mg of product (0.56 mmol, 79%).
***R*f** = 0.3 (DCM/Acetone 1:1), visualization with UV and molybdate stain.
This compound exists in two forms in MeOD. The reported peaks correspond to the major form which was presumed to be the keto compound.

**$^{1}$H NMR** (400 MHz, MeOD-$d_4$) δ 8.24 (d, *J* = 9.3 Hz, 2H, Ar), 7.28 (d, *J* = 9.3 Hz, 2H, Ar), 5.20 (d, *J* = 7.8 Hz, 1H, H-1), 4.48 (dd, *J* = 7.8, 1.7 Hz, 1H, H-2), 4.37 (dd, *J* = 10.2, 1.7 Hz, 1H, H-4), 3.99 (dd, *J* = 12.3, 2.1 Hz, 1H, H-6), 3.84 (dd, *J* = 12.3, 4.8 Hz, 1H, H-6), 3.62 (ddd, *J* = 10.2, 4.8, 2.1 Hz, 1H, H-5).

**$^{13}$C NMR** (101 MHz, MeOD-$d_4$) δ 205.0, 162.1, 142.96, 125.4, 116.8, 101.6, 77.4, 74.2, 72.2, 61.0.

*Methylumbelliferyl β-D-ribo-hexopyranoside-3-ulose **30***

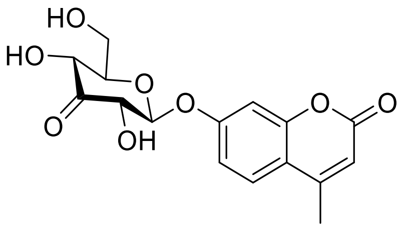

Compound **30** was synthesized from methylumbelliferyl β-glucopyranoside (100 mg, 0.30 mmol) using the general procedure for synthesis of 3-keto-glycosides to yield 65 mg of product (0.19 mmol, 64%).
Note: The peaks for the hydrate form are most visible in the C-NMR, and those assigned for the hydrate are identified based on HSQC and HMBC spectra.

***R*f** = 0.4 (DCM/Acetone 1:1), visualization with UV and molybdate stain.
**$^{1}$H NMR** (400 MHz, DMSO-$d_6$) δ 7.73 (d, *J* = 8.7 Hz, 1H, Ar), 7.12 (s, 1H, Ar), 7.08 (d, *J* = 8.8 Hz, 1H, Ar), 6.27 (d, *J* = 1.4 Hz, 1H, H-4'), 5.79 (s, 1H, OH), 5.53 (s, 1H, OH), 5.24 (d, *J* = 7.9 Hz, 1H, H-1), 4.91 (d, *J* = 6.0 Hz, 1H, OH), 4.33 (d, *J* = 7.8 Hz, 1H, H-2), 4.19 (d, *J* = 9.8 Hz, 1H, H-4), 3.77 (dd, *J* = 10.1, 4.7 Hz, 1H, H-6), 3.67 – 3.56 (m, *J* = 7.0 Hz, 2H, H-6 & H-5), 2.41 (d, *J* = 1.2 Hz, 3H, Me).

**$^{13}$C NMR** (101 MHz, DMSO-d$_6$) δ 205.6, 191.0 (hydrate), 160.4 (hydrate), 160.1, 159.6, 154.9 (hydrate), 154.3, 153.7 (hydrate), 153.4, 147.4, 134.3, 126.7 (hydrate), 126.6, 114.5, 113.4, 113.0 (hydrate), 112.0, 110.2 (hydrate), 103.3, 102.2 (hydrate), 100.6, 83.6 (hydrate), 76.7, 76.2, 72.0, 67.8 (hydrate), 60.5, 60.0 (hydrate), 18.2.

*Methyl α-D-ribo-hexopyranoside-3-ulose **31***

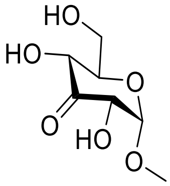

Compound **31** was synthesized from methyl-α-glucopyranoside (500 mg, 2.57 mmol) using the general procedure for synthesis of 3-keto-glycosides to yield 380 mg of product (1.98 mmol, 77%).
***R*f** = 0.5 (DCM/MeOH 5:1), visualization with molybdate stain.
**$^{1}$H NMR** (400 MHz, Acetone-d$_6$) δ 5.03 (d, *J* = 4.3 Hz, 1H, H-1), 4.39 (dd, *J* = 4.1, 1.6 Hz, 1H, H-2), 4.28 (dd, *J* = 9.7, 1.6 Hz, 1H, H-2), 3.85 (dd, *J* = 12.0, 2.3 Hz, 1H, H-6), 3.77 (dd, *J* = 12.0, 4.7 Hz, 1H, H-6), 3.60 (ddd, *J* = 9.5, 4.6, 2.2 Hz, 1H, H-5), 3.33 (s, 3H, OMe).

**$^{13}$C NMR** (101 MHz, Acetone-d$_6$) δ 206.9, 103.6, 76.9, 75.8, 73.2, 62.4, 55.5.

*Para-nitrophenyl α-D-ribo-hexopyranoside-3-ulose **32***

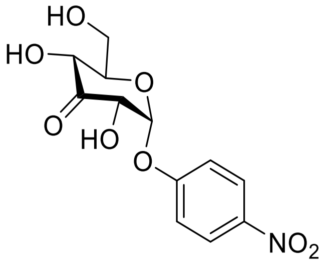

Compound **29** was synthesized from para-nitrophenyl *α*-glucopyranoside (500 mg, 1.66 mmol) using the general procedure for synthesis of 3-keto-glycosides to yield the product in a semi-pure form. To purify it further, this was crystalized from iso-propanol, yielding 350 mg of pure product (1.17 mmol, 70%).
***R*f** = 0.4 (DCM/Acetone 1:1), visualization with UV and molybdate stain.
**$^{1}$H NMR** (400 MHz, MeOD-d$_4$) δ 8.27 – 8.19 (m, 2H, Ar), 7.36 – 7.26 (m, 2H, Ar), 6.03 (d, *J* = 4.2 Hz, 1H, H-1), 4.65 (dd, *J* = 4.2, 1.6 Hz, 1H, H-2), 4.41 (dd, *J* = 9.8, 1.5 Hz, 1H, H-4), f3.80 (d, *J* = 3.3 Hz, 2H, H-6), 3.70 (dt, *J* = 9.8, 3.3 Hz, 1H, H-5).

**$^{13}$C NMR** (101 MHz, MeOD-d$_4$) δ 205.0, 161.44, 143.0, 125.4, 116.7, 100.3, 77.0, 74.2, 71.8, 60.9.

*Methylumbelliferyl α-D-ribo-hexopyranoside-3-ulose **33***

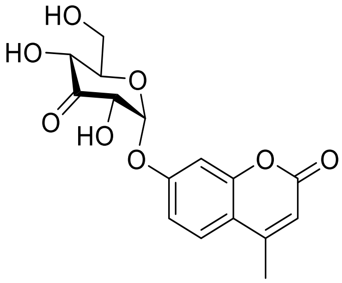

Compound **30** was synthesized from methylumbelliferyl *α*-glucopyranoside (50 mg, 0.15 mmol) using the general procedure for synthesis of 3-keto-glycosides to yield 30 mg of product (0.089 mmol, 60%).
***R*f** = 0.4 (DCM/Acetone 1:1), visualization with UV and molybdate stain.

**$^{1}$H NMR** (400 MHz, DMSO-$d_6$) δ 7.72 (d, *J* = 8.7 Hz, 1H), 7.11 (d, *J* = 2.4 Hz, 1H), 7.06 (dd, *J* = 8.8, 2.4 Hz, 1H), 6.27 (s, 1H), 6.01 (d, *J* = 4.2 Hz, 1H), 5.67 (d, *J* = 7.6 Hz, 1H), 5.55 (d, *J* = 6.5 Hz, 1H), 4.87 (t, *J* = 5.5 Hz, 1H), 4.55 (t, *J* = 6.0 Hz, 1H), 4.25 (t, *J* = 8.0 Hz, 1H), 3.58 (d, *J* = 12.9 Hz, 3H), 2.40 (s, 3H).

**$^{13}$C NMR** (101 MHz, DMSO-$d_6$) δ 205.6, 160.0, 159.0, 154.2, 153.2, 126.6, 114.6, 113.8, 112.0, 103.9, 100.2, 76.7, 74.0, 71.7, 60.5, 18.1.

#### 1.4. Synthesis of O-glycosides

In addition to the S-glycoside substrates described above, a diverse set of O-glycoside substrates has been used in the course of this study. The structures of all these substrates, many of which are commercially available, are presented in section 1.5. The synthetic scheme for those compounds that have not been previously reported is described in this section.

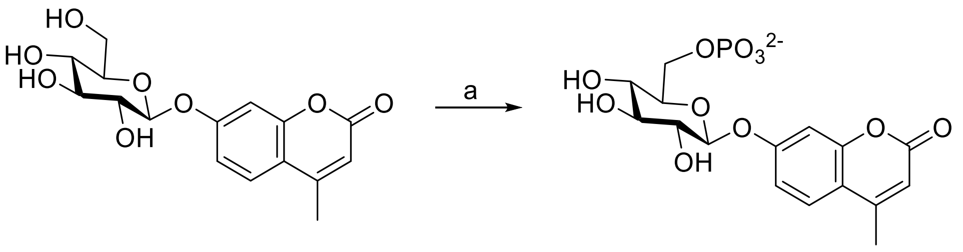

**34**

**Scheme S6:** Enzymatic synthesis of substrate 6PGlc-β-MU: a) BglK (1 mg/mL), ATP (2.0 eq), $MgCl_2$ (0.03 eq), rt, 2 h, ~4% DMSO in HEPES 25 mM pH 7.5, 85%

*Methylumbelliferyl β-D-glucoside-6-phosphate* **34**

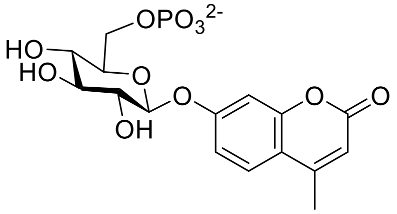

Following the procedure detailed in section 1.2.8. with the starting material Methylumbelliferyl β-D-glucoside (28 mg, 82.7 µmol) yielded 29.3 mg (70.3 µmol, 85%) of the desired substrate **34,** as a white solid.
***$R_f$*** ≈ 0.5 (t-BtOH/AcOH/$H_2O$ 5:2:3), visualization with UV and molybdate.
**$^1$H NMR** (400 MHz, Deuterium Oxide) δ 7.42 (d, *J* = 8.9 Hz, 1H), 6.95 (d, *J* = 8.9 Hz, 1H), 6.78 (s, 1H), 5.96 (s, 1H), 5.11 (d, *J* = 7.3 Hz, 1H), 4.28 – 4.07 (m, 2H), 3.80 (d, *J* = 8.4 Hz, 1H), 3.65 (dq, *J* = 16.9, 5.0 Hz, 4H), 2.22 (s, 4H).
**$^{13}$C NMR** (101 MHz, Deuterium Oxide) δ 164.0, 159.3, 155.9, 153.3, 126.5, 114.7, 113.8, 110.9, 103.2, 99.7, 75.1, 75.0, 72.8, 68.6, 63.4 (d, $J_{C\text{-}P}$ = 4.8 Hz), 17.9.
**$^{31}$P NMR** (162 MHz, Deuterium Oxide) δ 0.67.

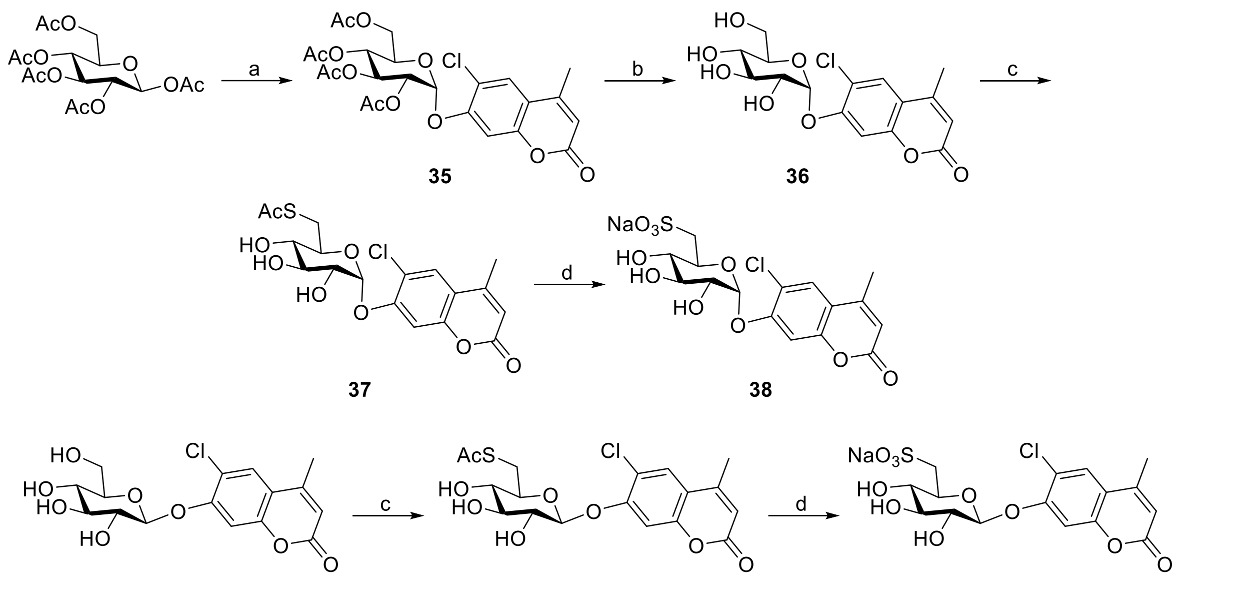

**39** **40**

**Scheme S7:** Synthesis of sulfoquinovoside substrates. Reagents and conditions: a) $CHCl_3$, $Cl_2CHOCH_3$, $BF_3 \cdot Et_2O$, rt; CMuONa, hexamethylphosphoramide, rt; 34%. b) DCM, MeOH, Na, rt, 93%. c) DMF, THF, AcSH, DIAD, $Ph_3P$, 0°C; α: 58%; β: 62%. d) KOAc, 30% $H_2O_2$, 50°C; MeOH, Na, rt; α: 40%; β: 64%.

*6-Chloromethylumbelliferyl 2,3,4,6-tetra-O-acetyl-α-D-glucopyranoside*[10,11] **35**

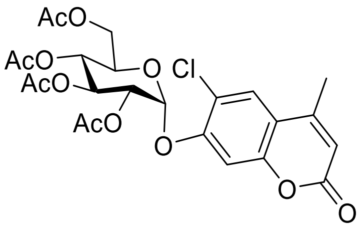

To a solution of 1,2,3,4,6-penta-*O*-acetyl β-D-glucopyranose (5 g, 12.82 mmol) in chloroform (10 mL, dried with molecular sieves) and dichloromethyl methyl ether (10 mL, dried with molecular sieves) was added $BF_3 \cdot Et_2O$ (0.5 mL) as one portion with stirring under $N_2$ at room temperature, the reaction mixture was stirred for 5 h at the same condition. The solvents were evaporated, and the resulting residue was redissovled in diethyl ether (400 mL), washed with cold $H_2O$ (2 x 200 mL), cold saturated $NaHCO_3$ (200 mL), cold brine (200 mL), dried over $MgSO_4$, filtered and evaporated, dried under vacuum for 0.5 h to afford a foam. To the foam were added 6-chlorocoumarin sodium salt (4.47 g (19.22 mmol, 1.5 equiv) and hexamethylphosphoramide (45 mL), the suspension was stirred vigorously for 66 h at room temperature under $N_2$. After completion, cold water (200 mL) was added, extracted with EtOAc (3 x 250 mL). The organic phase was washed with $H_2O$ (300 mL), 1 M NaOH (300 mL), saturated $NaHCO_3$ (300 mL) and brine (300 mL), dried over $MgSO_4$, filtered and concentrated. The resulting residue was purified by flash column chromatography (PE/EtOAc = 2/1 and 3/2) to afford the product as a white solid (2.34 g, 34%).

**ESI-HRMS** (*m/z*): $[M+H]^+$ calcd for $C_{24}H_{26}ClO_{12}$ 541.1113, found 541.1100.

**$^{1}$H NMR** (400 MHz, $CDCl_3$) δ 7.61 (s, 1 H, Ar), 7.23 (s, 1 H, Ar), 6.24 (s, 1 H, Ar), 5.81 (d, *J* = 3.6 Hz, 1 H, H-1), 5.73 (t, *J* = 10.0 Hz, 1 H, H-3), 5.16 (t, *J* = 10.0 Hz, 1 H, H-4), 5.05 (dd, *J* = 4.0, 10.0 Hz, 1 H, H-2), 4.28 (dd, *J* = 4.8, 12.0 Hz, 1 H, H-6), 4.17 (m, 1 H, H-5), 4.07 (dd, *J* = 2.0, 12.0 Hz, 1 H, H-6), 2.40 (d, *J* = 0.8 Hz, 3 H, Ar-$CH_3$), 2.10 (s, 3 H, OAc), 2.08 (s, 3 H, OAc), 2.07 (s, 3 H, OAc), 2.05 (s, 3 H, OAc).

**$^{3}$C NMR** (100 MHz, $CDCl_3$) δ 170.7, 170.4, 170.1, 169.7, 160.1, 154.3, 153.2, 151.2, 125.9, 120.6, 116.4, 114.5, 105.9, 96.1, 70.5, 69.9, 68.9, 68.2, 61.7, 20.83, 20.77, 20.7, 18.8.

*6-Chloromethylumbelliferyl α-D-glucopyranoside* **36**

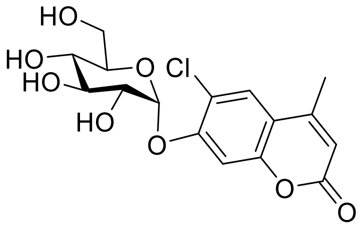

To a solution of 6-chloromethylumbelliferyl 2,3,4,6-tetra-*O*-acetyl-α-D-glucopyranoside (2.16 g, 4.00 mmol) in dry DCM (10 mL) and dry methanol (100 mL) was added a catalytic amount of sodium under $N_2$, the mixture was stirred at the same condition for 6 h. The formed white precipitate was filtered, washed with cold methanol to afford the product. The mother liquor was neutralized Amberlite 120H, evaporated to give another portion of product, repeated twice. The white solid was dried under vacuum overnight (1.381 g, 93%).

**ESI-HRMS** (*m/z*): $[M+Na]^+$ calcd for $C_{16}H_{17}ClO_8$ 395.0510, found 395.0503.

**$^1$H NMR** (400 MHz, $d_6$-DMSO + $D_2O$) δ 7.83 (s, 1 H, Ar), 7.35 (s, 1 H, Ar), 6.29 (d, *J* = 1.2 Hz, 1 H, Ar), 5.77 (d, *J* = 3.2 Hz, 1 H, H-1), 3.70 (t, *J* = 9.2 Hz, 1 H, H-3), 3.52 (dd, *J* = 1.2, 11.2 Hz, 1 H, H-6), 3.47-3.36 (m, 3 H, H-2, H-5 & H-6), 3.21 (t, *J* = 9.6 Hz, 1 H, H-4), 2.51 (s, 3 H, Ar-$CH_3$).

**$^{13}$C NMR** (100 MHz, $d_6$-DMSO + $D_2O$) δ 160.4, 155.1, 153.42, 153.37, 126.5, 119.4, 115.1, 113.1, 104.6, 98.6, 75.2, 73.1, 71.7, 70.0, 61.0, 18.7.

*6-Chloromethylumbelliferyl 6-S-acetyl-6-thio-α-D-glucopyranoside*[12] **37**

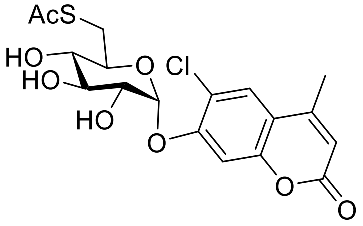

To a clear solution of 6-chloromethylumbelliferyl α-D-glucopyranoside (0.943 g, 2.53 mmol) in dry DMF (10 mL) was added dry THF (20 mL), stirred at 0°C under $N_2$, thioacetic acid (0.22 mL, 3.1 equiv) was added as one portion. A mixture of $Ph_3P$ (1.99 g, 7.6 mmol, 3 equiv) and DIAD (0.98 mL, 5.0 mmol, 2 equiv) in dry THF (20 mL) was added dropwise to above solution. The reaction mixture was stirred for 5 h at 0°C, quenched by addition of methanol and water, evaporated and co-evaporated with toluene to remove DMF to give a solid. To the solid was added methanol and EtOAc to give first portion of product, repeated these procedures three times. The mother liquor was concentrated and purified by flash column chromatography (DCM/MeOH = 100/1, 50/1, 25/1 and 15/1). The product was obtained as a white solid (628 mg, 58%).

**ESI-HRMS** (*m/z*): $[M+Na]^+$ calcd for $C_{18}H_{19}ClO_8S$ 453.0387, found 453.0394.

**$^1$H NMR** (400 MHz, $d_6$-DMSO + $D_2O$) δ 7.85 (s, 1 H, Ar), 7.31 (s, 1 H, Ar), 6.32 (d, *J* = 0.8 Hz, 1 H, Ar), 5.77 (d, *J* = 3.2 Hz, 1 H, H-1), 3.65 (t, *J* = 9.2 Hz, 1 H, H-3), 3.48 (dd, *J* = 3.6, 9.6 Hz, 1 H, H-2), 3.42 (m, 1 H, H-5), 3.36 (dd, *J* = 2.4, 14.0 Hz, 1 H, H-6), 3.12 (t, *J* = 9.2 Hz, 1 H, H-4), 2.81 (dd, *J* = 9.2, 14.0 Hz, 1 H, H-6), 2.41 (d, *J* = 1.6 Hz, 3 H, Ar-$CH_3$), 2.12 (s, 3 H, SAc).

**$^{13}$C NMR** (100 MHz, $d_6$-DMSO + $D_2O$) δ 195.1, 160.4, 154.5, 153.4, 153.3, 126.5, 119.4, 115.3, 113.2, 105.1, 98.3, 73.3, 72.9, 72.7, 71.6, 31.1, 30.7, 18.7.

*Sodium 6-Chloromethylumbelliferyl 6-deoxy-6-sulfonato-α-D-glucopyranoside* **38**

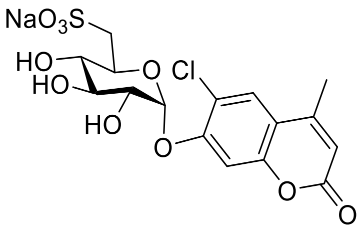

To a suspension of 6-chloromethylumbelliferyl 6-S-acetyl-6-thio-α-D-glucopyranoside (1.03 g, 2.39 mmol) and KOAc (0.26 g, 2.65 mmol, 1.1 equiv) was added 30% $H_2O_2$ (13 mL), the reaction mixture was stirred overnight at 50°C. The reaction mixture was cooled to rt and further cooled to 0°C, diluted with water (50 mL), quenched by dropwise addition of $Ph_3P$ in ether (1 M, 30 mL), separated two layers. The organic phase was extracted with water (2 x 50 mL). The combined aqueous phase was extracted with ether (2 x 100 mL). Water was evaporated to afford a solid. The solid was dried under vacuum overnight, treated with dry methanol and sodium. To the reaction mixture were added water and silica gel, evaporated. The product was purified by flash column chromatography (EtOAc/MeOH/$H_2O$ = 17/2/1, 12/2/1, 10/2/1 and 9/2/1) (450 mg, 40%).
**ESI-HRMS** (*m/z*): $[M-H]^-$ calcd for $C_{16}H_{16}ClO_{10}S$ 435.0153, found 435.0147.
**$^1$H NMR** (400 MHz, $D_2O$) δ 7.70 (s, 1 H, Ar), 7.25 (s, 1 H, Ar), 6.19 (d, *J* = 1.2 Hz, 1 H, Ar), 5.78 (d, *J* = 3.6 Hz, 1 H, H-1), 4.09 (m, 1 H, H-5), 4.06 (t, *J* = 9.2 Hz, 1 H, H-3), 3.85 (dd, *J* = 3.6, 10.0 Hz, 1 H, H-2), 3.43 (t, *J* = 9.2 Hz, 1 H, H-4), 3.32 (dd, *J* = 1.2, 14.4 Hz, 1 H, H-6), 3.08 (dd, *J* = 9.2, 14.8 Hz, 1 H, H-6), 2.34 (s, 3 H, Ar-$CH_3$).
**$^{13}$C NMR** (100 MHz, $D_2O$) δ 164.2, 155.4, 154.0, 152.2, 126.1, 120.7, 115.6, 112.2, 105.2, 97.3, 72.9, 72.4, 71.2, 69.8, 52.2, 18.2.

*6-Chloromethylumbelliferyl 6-S-acetyl-6-thio-β-D-glucopyranoside* **39**

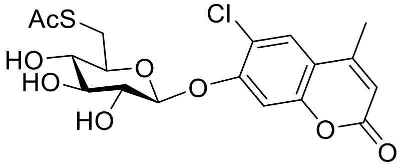

To a clear solution of 6-chloromethylumbelliferyl β-D-glucopyranoside[13] (1.088 g, 2.92 mmol) in dry DMF (11 mL) was added dry THF (21 mL), stirred at 0°C under $N_2$, thioacetic acid (0.25 mL, 3.55 mmol, 1.2 equiv) was added as one portion. A mixture of $Ph_3P$ (2.83 g, 10.8 mmol, 3.7 equiv) and DIAD (1.16 mL, 5.90 mmol, 2 equiv) in dry THF (21 mL) was added dropwise to above solution. The reaction mixture was stirred for 24 h at 0°C, quenched by addition of methanol and water, evaporated and co-evaporated with toluene to remove DMF to give a solid. To the solid was added methanol, filtered and washed with methanol, and the filtrate was concentrated. The resulting residue was purified by flash column chromatography (DCM/MeOH = 100/1, 50/1, 25/1 and 15/1) to afford the product as a white solid (0.78 g, 62%).
**ESI-HRMS** (*m/z*): $[M+Na]^+$ calcd for $C_{18}H_{19}ClO_8S$ 453.0387, found 453.0387.
**$^1$H NMR** (400 MHz, $d_6$-DMSO + $D_2O$) δ 7.81 (s, 1 H, Ar), 7.24 (s, 1 H, Ar), 6.29 (d, *J* = 1.2 Hz, 1 H, Ar), 5.16 (d, *J* = 7.6 Hz, 1 H, H-1), 3.57 (m, 1 H, H-5), 3.48 (dd, *J* = 2.4, 14.0 Hz, 1 H, H-6), 3.37 – 3.27 (m, 2 H, H-2 & H-3), 3.10 (t, *J* = 8.8 Hz, 1 H, H-4), 2.83 (dd, *J* = 9.2, 14.0 Hz, 1 H, H-6), 2.39 (d, *J* = 0.4 Hz, 3 H, Ar-$CH_3$), 2.29 (s, 3 H, SAc).
**$^{13}$C NMR** (100 MHz, $d_6$-DMSO + $D_2O$) δ 195.3, 160.4, 155.1, 153.4 (2 C), 126.5, 118.6, 115.3, 113.3, 104.7, 100.4, 76.6, 75.5, 73.5, 73.3, 31.1, 30.9, 18.7.

*Sodium 6-Chloromethylumbelliferyl 6-deoxy-6-sulfonato-β-D-glucopyranoside* **40**

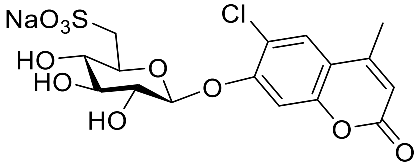

To a suspension of 6-chloromethylumbelliferyl 6-S-acetyl-6-thio-β-D-glucopyranoside (0.9 g, 2.1 mmol) and KOAc (0.225 g, 2.30 mmol, 1.1 equiv) was added 30% $H_2O_2$ (11 mL), the reaction mixture was stirred overnight at 50°C. The reaction mixture

was cooled to rt and further cooled to 0°C, diluted with water (50 mL), quenched by dropwise addition of $Ph_3P$ in ether (1 M, 30 mL), separated two layers. The organic phase was extracted with water (2 x 50 mL). The combined aqueous phase was extracted with ether (2 x 100 mL). Water was evaporated to afford a solid. The solid was dried under vacuum overnight, treated with dry methanol and sodium. To the reaction mixture were added water and silica gel, evaporated. The product was purified by flash column chromatography (EtOAc/MeOH/$H_2O$ = 12/2/1, 10/2/1 and 9/2/1) (638 mg, 64%).
**ESI-HRMS** (*m*/*z*): $[M-H]^-$ calcd for $C_{16}H_{16}ClO_{10}S$ 435.0153, found 435.0158.
**$^1$H NMR** (400 MHz, $D_2O$) δ 7.75 (s, 1 H, Ar), 7.38 (s, 1 H, Ar), 6.21 (d, *J* = 1.2 Hz, 1 H, Ar), 5.11 (d, *J* = 7.6 Hz, 1 H, H-1), 3.96 (m, 1 H, H-5), 3.65 (dd, *J* = 7.6, 9.2 Hz, 1 H, H-2), 3.58 (t, *J* = 9.6 Hz, 1 H, H-3), 3.39 (dd, *J* = 1.2, 15.2 Hz, 1 H, H-6), 3.32 (t, *J* = 9.2 Hz, 1 H, H-4), 3.02 (dd, *J* = 9.6, 15.2 Hz, 1 H, H-6), 2.34 (s, 3 H, Ar-$CH_3$).
**$^{13}$C NMR** (100 MHz, $D_2O$) δ 163.8, 155.0, 154.7, 152.1, 125.8, 119.7, 115.3, 112.2, 105.0, 100.8, 75.3, 72.8 (2 C), 72.1, 52.2, 18.1.

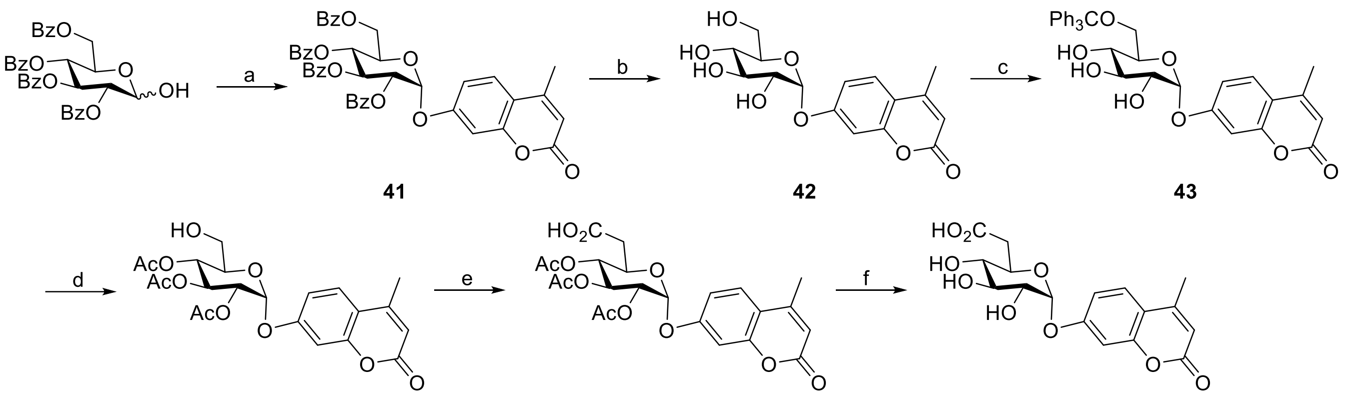

41 42 43

44 45 46

**Scheme S8:** Synthesis of α-glucuronide substrate. Reagents and conditions: a) DCM, trichloroacetonitrile, DBU, 0°C, 76%; $CH_3CN$, MuOH, 4Å molecular sieves, TMSOTf, -20°C, 61%. b) DCM, MeOH, Na, rt, 83%. c) Pyridine, trityl chloride, 95°C; $Ac_2O$, 0°C-rt; 38%. d) HCOOH, $Et_2O$, rt, 80%. e) DCM/$H_2O$=1/1, NaBr, TBAB, $NaHCO_3$, TEMPO, bleach, 0°C, 62%. f) MeOH, NaOMe/MeOH, 0°C, 92%.

*Methylumbelliferyl 2,3,4,6-tetra-O-benzoyl-α-D-glucopyranoside* **41**

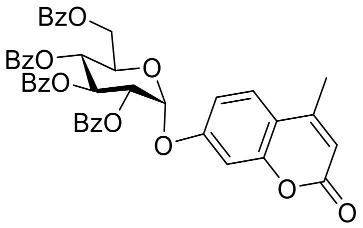

To a solution of 2,3,4,6-tetra-*O*-benzoyl-D-glucopyranose (7.19 g, 12.07 mmol), stirred at 0°C under $N_2$, were added trichloroacetonitrile (12 mL, 119.67 mmol, 9.9 eq) and DBU (0.14 mL), and the reaction mixture stirred for 1.5 h at 0°C and concentrated. The resulting residue was purified with flash column chromatography (petroleum ether / EtOAc = 6/1) to afford 2,3,4,6-tetra-*O*-benzoyl-D-glucosyl trichloroacetimidate as a white foam (6.788 g, 9.17 mmol, 76%). A mixture of the white foam, 4-methylumbelliferone (3.879 g, 22.04 mmol, 2.4 eq) and 4Å molecular sieves (5 g) in dry acetonitrile (120 mL) was stirred for 0.5 h at room temperature under $N_2$, cooled down to -20°C, TMSOTf (3.6 mL, 19.86 mmol, 2.2 eq) was added as one portion. The reaction mixture was continued to stir for 12 h at the same condition, filtered through Celite and washed with acetone, evaporated. The resulting residue was re-dissolve in DCM (400 mL), washed with 1 M NaOH (2 x 200 mL), saturated $NaHCO_3$ (2 x 200 mL) and brine (200 mL), dried over $MgSO_4$, filtered and concentrated. The resulting residue was purified by flash column chromatography (petroleum ether / EtOAc = 3/1 and 2/1) to afford the product as a white solid (4.214 g, 61%).
ESI-MS (*m/z*): $[M+Na]^+$ calcd for $C_{44}H_{34}O_{12}$ 777.2, found 777.4.
**$^1$H NMR** (400 MHz, $CDCl_3$) δ 7.90 – 7.15 (m, 21 H, Ar), 7.04 (d, *J* = 2.4 Hz, 1 H, Ar), 6.98 (dd, *J* = 2.4, 8.8 Hz, 1 H, Ar), 6.33 (t, *J* = 9.9 Hz, 1 H, H-3), 5.99 (d, *J* = 1.1 Hz, 1 H, Ar), 6.04 (d, *J* = 3.6 Hz, 1 H, H-1), 5.69 (t, *J* = 9.9 Hz, 1 H, H-4), 5.47 (dd, *J* = 3.6, 10.2 Hz, 1 H, H-2), 4.50 – 4.36 (m, 3 H, H-5 & H-6), 2.17 (s, 3 H, Ar-$CH_3$).
**$^{13}$C NMR** (100 MHz, $CDCl_3$) δ 166.0, 165.9, 165.8, 165.3, 160.7, 158.6, 154.7, 152.0, 133.7, 133.66, 133.4, 133.2, 129.95, 129.94, 129,8, 129.77, 129.4, 129.0, 128.7, 128.6, 128.57, 128.5, 128.4, 128.3, 115.4, 113.4, 113.2, 104.8, 94.6, 71.4, 70.3, 69.2, 69.1, 62.7, 18.6.

*Methylumbelliferyl α-D-glucopyranoside* **42**

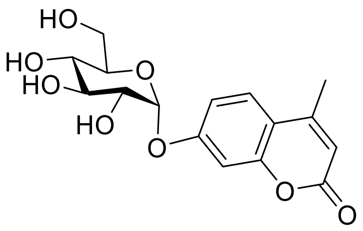

To a solution of methylumbelliferyl 2,3,4,6-tetra-*O*-benzoyl-α-D-glucopyranoside (3.812 g, 5.06 mmol) in dry DCM (10 mL) and dry methanol (150 mL) was added catalytic amount of sodium at room temperature, and reaction mixture was stirred for 4 h under $N_2$. The reaction mixture was neutralized with Amberlite 120 H, filtered and evaporated. The white solid was filtered and washed with cold MeOH, the filtrate was evaporated and precipitate was filtered, repeated three times. The product was obtained as a white solid (1.424 g, 83%).
ESI-MS (*m/z*): $[M+Na]^+$ calcd for $C_{16}H_{18}O_8$ 361.1, found 361.1.
$^1$**H NMR** (400 MHz, $d_6$-DMSO + $D_2O$) δ 7.71 (d, *J* = 8.8 Hz, 1 H, Ar), 7.10 -7.07 (m, 2 H, Ar), 6.23 (d, *J* = 0.8 Hz, 1 H, Ar), 5.56 (d, *J* = 3.6 Hz, 1 H, H-1), 3.57 (m, 1 H, H-5), 3.65 – 3.15 (m, 6 H, rest protons on sugar ring), 2.40 (d, *J* = 0.8 Hz, 3 H, SAc).

*Methylumbelliferyl 2,3,4-tri-O-acetyl-6-O-trityl-α-D-glucopyranoside* **43**

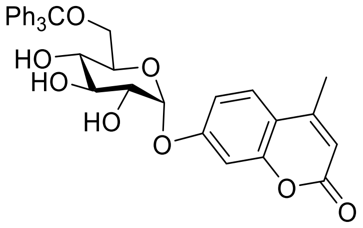

A mixture of methylumbelliferyl α-D-glucopyranoside (0.5 g, 1.48 mmol) and trityl chloride (0.495 g, 1.78 mmol, 1.2 eq) in dry pyridine (3 mL) was stirred for 6 h at 95°C under $N_2$, cooled to room temperature and further cooled with an ice bath, was added $Ac_2O$ (1.5 mL). The reaction mixture was stirred at 0°C for 1 h, and then overnight at room temperature. The reaction mixture was poured into ice and stirred for 1 h, extracted with EtOAc (2 x 50 mL). The organic phase was washed with 1 M HCl (50 mL), saturated $NaHCO_3$ (2 x 50 mL) and brine (50 mL), dried over $MgSO_4$, filtered and concentrated. The resulting residue was purified by flash column chromatography (petroleum ether / EtOAc = 2/1) to afford the product as a white solid (396 mg, 38%).
ESI-MS (*m/z*): $[M+Na]^+$ calcd for $C_{41}H_{38}O_{11}$ 729.2, found 729.2.
$^1$**H NMR** (400 MHz, $CDCl_3$) δ 7.54– 7.12 (m, 18 H, Ar), 6.22 (d, *J* = 1.1 Hz, 1 H, Ar), 5.87 (d, *J* = 3.7 Hz, 1 H, H-1), 5.63 (t, *J* = 9.8 Hz, 1 H, H-3), 5.16 (t, *J* = 9.6, 10.2 Hz, 1 H, H-4), 5.09 (dd, *J* = 3.7, 10.3 Hz, 1 H, H-2), 3.95 (m, 1 H, H-5), 3.17 (dd, *J* = 2.4, 10.6 Hz, 1 H, H-6), 3.12 (dd, *J* = 5.2, 10.7 Hz, 1 H, H-6), 2.42 (d, *J* = 1.0 Hz, 3 H, Ar-$CH_3$), 2.09 (s, 3 H, OAc), 2.05 (s, 3 H, OAc), 1.76 (s, 3 H, OAc).

*Methylumbelliferyl 2,3,4-tri-O-acetyl-α-D-glucopyranoside* **44**

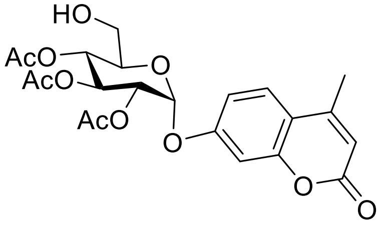

A mixture of methylumbelliferyl 2,3,4-tri-*O*-acetyl-6-*O*-trityl-α-D-glucopyranoside (0.38 g, 0.54 mmol), formic acid (20 mL) and diethyl ether (20 mL) was stirred for 1.5 h at room temperature, diluted with EtOAc (75 mL), washed with water (3 x 50 mL),

saturated $NaHCO_3$ (50 mL) and brine (50 mL), dried over $MgSO_4$, filtered and concentrated. The resulting residue was purified by flash column chromatography (petroleum ether / EtOAc = 3/2 & 1/1) to afford the product as a white solid (200 mg, 80%).
ESI-MS (*m/z*): $[M+Na]^+$ calcd for $C_{22}H_{24}O_{11}$ 487.1, found 487.1.
**$^1$H NMR** (400 MHz, $CDCl_3$) δ 7.55 (d, *J* = 8.4 Hz, 1 H, Ar), 7.05 (m, 2 H, Ar), 6.20 (d, *J* = 1.0 Hz, 1 H, Ar), 5.85 (d, *J* = 3.6 Hz, 1 H, H-1), 5.78 (t, *J* = 9.9 Hz, 1 H, H-3), 5.18 (t, *J* = 9.9 Hz, 1 H, H-4), 5.04 (dd, *J* = 3.6, 10.2 Hz, 1 H, H-2), 3.87 (m, 1 H, H-5), 3.68 (dd, *J* = 2.1, 13.0 Hz, 1 H, H-6), 3.57 (dd, *J* = 3.5, 13.0 Hz, 1 H, H-6), 2.42 (d, *J* = 1.0 Hz, 3 H, Ar-$CH_3$), 2.088 (s, 3 H, OAc), 2.087 (s, 3 H, OAc), 2.08 (s, 3 H, OAc).

*Methylumbelliferyl 2,3,4-tri-O-acetyl-α-D-glucuronic acid* **45**

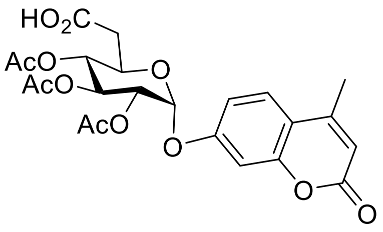

A mixture of methylumbelliferyl 2,3,4-tri-*O*-acetyl-α-D-glucopyranoside (197 mg, 0.42mmol), NaBr (12 mg) and TBAB (12 mg) in DCM/$H_2O$ (1/1, 4 mL) was cooled with an ice bath, and then saturated $NaHCO_3$ (3 mL), TEMPO (12 mg) and bleach (3.8 mL) were added. The reaction mixture was stirred vigorously for 3 h at 0°C, two layers were separated. The aqueous layer was extracted with DCM (3 x 5 mL). To the aqueous phase was added DCM (5 mL), cooled to 0°C, 1 M HCl was added to adjust pH about 3. Two layers were separated, water phase was extracted with DCM (3 x 5 mL). The combined organic phase was washed with water, dried over $MgSO_4$, filtered and concentrated. The resulting residue was purified by flash column chromatography (DCM / MeOH = 10/1) to afford the product as a white solid (126 mg, 62%).
ESI-MS (*m/z*): $[M-H]^-$ calcd for $C_{22}H_{22}O_{12}$ 477.1, found 487.0.
**$^1$H NMR** (400 MHz, $CDCl_3$) δ 7.56 (d, *J* = 8.8 Hz, 1 H, Ar), 7.13 -7.06 (m, 2 H, Ar), 6.22 (s, 1 H, Ar), 5.91 (d, *J* = 3.3 Hz, 1 H, H-1), 5.74 (t, *J* = 9.6 Hz, 1 H, H-3), 5.33 (t, *J* = 9.6 Hz, 1 H, H-4), 5.10 (dd, *J* = 3.6, 10.1 Hz, 1 H, H-2), 4.42 (d, *J* = 10.1 Hz, 1 H, H-5), 2.42 (brs, 3 H, Ar-$CH_3$), 2.08 (s, 6 H, 2 x OAc), 2.06 (s, 3 H, OAc).

*Methylumbelliferyl α-D-glucuronic acid* **46**

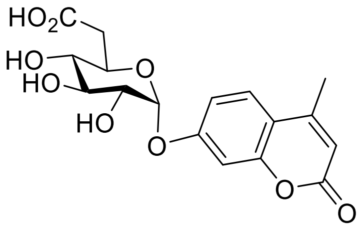

A solution of methylumbelliferyl 2,3,4-tri-*O*-acetyl-α-D-glucuronic acid (115 mg, 0.24 mmol) was cooled with an ice bath, and then freshly made MeONa in MeOH (1 M, 1 mL) was added. The reaction mixture was stirred for 1 h at 0°C under $N_2$, white precipitates came out. After evaporation, the white solid was re-dissolved in water, neutralized with Amberlite 120 H, filtered and washed with water, concentrated and freeze dried to afford the product as a white solid (78 mg, 92%).
ESI-MS (*m/z*): $[M-H]^-$ calcd for $C_{16}H_{16}O_9$ 351.1, found 351.1.
**$^1$H NMR** (400 MHz, $d_6$-DMSO + $D_2O$) δ 7.71 (d, *J* = 9.4 Hz, 1 H, Ar), 7.09 (m, 2 H, Ar), 6.23 (s, 1 H, Ar), 5.67 (d, *J* = 3.2 Hz, 1 H, H-1), 3.74 (d, *J* = 9.9 Hz, 1 H, H-5), 3.64 (t, *J* = 9.2 Hz, 1 H, H-3), 3.46 (dd, *J* = 3.3, 9.4Hz, 1 H, H-2), 3.36 (t, *J* = 9.3 Hz, 1 H, H-4), 2.39 (s, 3 H, Ar-$CH_3$).
**$^{13}$C NMR** (100 MHz, $d_6$-DMSO + $D_2O$) δ 171.5, 160.8, 160.0, 127.2, 114.8, 114.3, 112.3, 104.0, 97.7, 73.0, 72.8, 72.2, 71.4, 18.7.

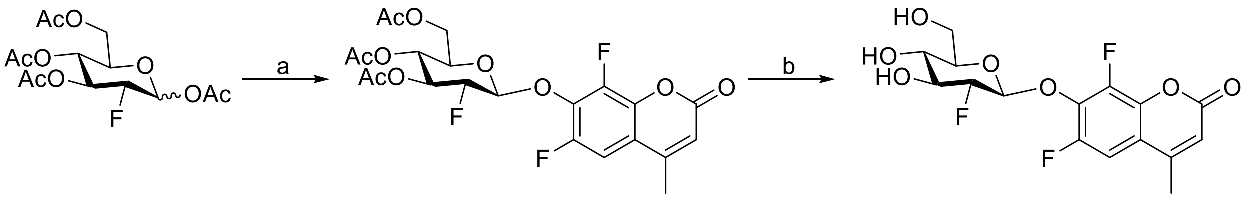

**47** **48**

**Scheme S9:** Synthesis scheme for 2F-Glc-β-$F_2$MU, Reagents and conditions: a) DCM, $Ac_2O$, HBr/AcOH, rt; $CH_3CN$, difluoromethylumbelliferone, $Ag_2O$, 4Å molecular sieves, 2,6-lutidine, rt; 63%. b) MeOH, Na, rt, 85%.

*6,8-Difluoromethylumbelliferyl 3,4,6-tri-O-acetyl-2-fluoro-2-deoxy-β-D-glucopyranoside* **47**

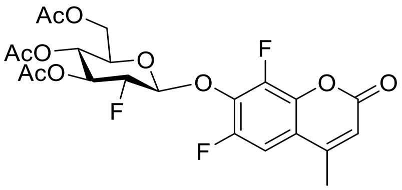

To a solution of 1,3,4,6-tetra-*O*-acetyl-2-fluoro-2-deoxy-D-glucopyranose[14] (533 mg, 1.52 mmol) in anhydrous DCM (6 mL) were added acetic anhydride (1 mL) and HBr/AcOH (2 mL) at room temperature, and the reaction mixture was stirred at the same condition overnight. After completion, the reaction mixture was diluted with DCM (50 mL), washed with cold water (30 mL), cold saturated $NaHCO_3$ (2 x 30 mL) and cold brine (30 mL), dried over $MgSO_4$, filtered and concentrated, dried under vacuum to afford a slightly yellow foam in quantitative yield (565 mg). A suspension of crude bromide (266 mg, 0.72 mmol), difluorocoumarin[15,16] (240 mg, 70% purity), $Ag_2O$ (0.18 g) and molecular sieves (0.2 g) in dry $CH_3CN$ (10 mL) was stirred for 10 minutes under $N_2$. To the mixture was added lutidine (1 mL), and the reaction mixture was stirred for 26 h at room temperature under $N_2$. The reaction mixture was filtered through Celite and washed with acetone, concentrated. The resulting residue was re-dissolved in DCM (50 mL), washed with 1 M HCl (20 mL), saturated $NaHCO_3$ (2 x 20 mL) and brine (20 mL), dried over $MgSO_4$, filtered and evaporated. The resulting residue was purified by flash column chromatography (petroleum ether / EtOAc = 2/1 and 3/2) to afford the product as a white solid (225 mg, 63%).
ESI-MS (*m/z*): $[M+Na]^+$ calcd for $C_{22}H_{21}F_3O_{10}$ 525.1, found 525.3**.**
**$^1$H NMR** (400 MHz, $CDCl_3$) δ 7.20 (dd, J = 2.0 Hz & 10.8 Hz, 1 H, Ar), 6.38 (s, 1 H, Ar), 5.46 (dt, *J* = 8.8, 14.8 Hz, 1 H, H-3), 5.41 (dd, *J* = 2.8, 7.2 Hz, 1 H, H-1), 5.20 (t, 1 H, H-4), 4.70 (dt, *J* = 8.0, 50.4 Hz, 1 H, H-2), 4.29 (dd, *J* = 4.8, 12.4 Hz, 1 H, H-6), 4.14 (dd, *J* = 2.4 Hz, 1 H, H-6), 3.83 (m, 1 H, H-5), 2.44 (s, 3 H, Ar-$CH_3$), 2.16 (s, 3 H, OAc), 2.091 (s, 3 H, OAc), 2.086 (s, 3 H, OAc).
**$^{19}$F NMR** (282 MHz, $CDCl_3$) δ 131.5, 145.3, 199.9.

*6,8-Difluoromethylumbelliferyl 2-fluoro-2-deoxy-β-D-glucopyranoside* **48**

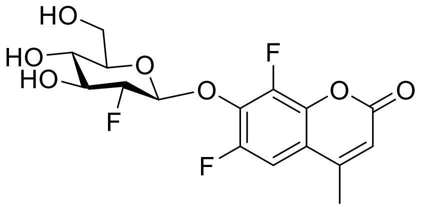

To a solution of 6,8-Difluoromethylumbelliferyl 3,4,6-tri-*O*-acetyl-2-fluoro-2-deoxy-β-D-glucopyranoside (173 mg, 0.34 mmol) in dry DCM (2 mL) and dry methanol (15 mL) was added a catalytic amount of sodium under $N_2$, the mixture was stirred at the same condition for 2 h. After completion, the reaction mixture was neutralized with Amberlite 120 H, filtered and concentrated, recrystallized from methanol and diethyl ether to afford the product as a white solid (110 mg, 85%).
ESI-MS (*m/z*): $[M+Na]^+$ calcd for $C_{16}H_{15}F_3O_7$ 399.1, found 399.3.
**$^1$H NMR** (400 MHz, MeOD) δ 7.45 (dd, *J* = 2.0, 10.6 Hz, 1 H, Ar), 6.39 (s, 1 H, Ar), 5.38 (dd, *J* = 2.4, 7.2 Hz, 1 H, H-1), 4.30 (dt, *J* = 8.4, 51.6 Hz, 1 H, H-2), 3.82 (dd, *J* = 1.6, 12.0 Hz, 1 H, H-6), 3.73 (m, 1 H, H-5), 3.67 (dd, *J* = 5.2 Hz, 1 H, H-6), 3.46 (t, *J* = 9.2 Hz, H-4), 3.38 (m, 1 H, H-3), 2.61 (s, 3 H, Ar-$CH_3$).
**$^{19}$F NMR** (282 MHz, $CDCl_3$) δ 133.1, 148.5, 201.6.

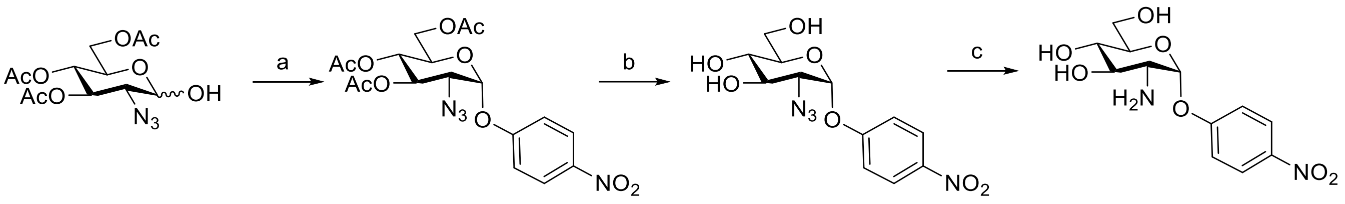

**49** **50** **51**

**Scheme S10:** Synthesis of 2-amino-2-deoxy-glucoside substrate. Reagents and conditions: a) DCM, trichloroacetonitrile, DBU, 0°C, 87%; DCM, 4-nitrophenol, 3Å molecular sieves, TMSOTf, -20°C, 56%. b) MeOH, Na, rt, 84%. c) THF, $H_2O$, silica gel, $Ph_3P$, 50°C, 78%.

*4-Nitrophenyl 3,4,6-tri-O-acetyl-2-azido-2-deoxy-α-D-glucopyranoside* **49**

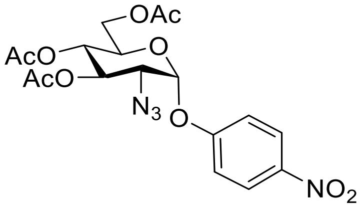

After flash column chromatography (PE / EtOAc = 4/1 and 3/1), the product was obtained as a white solid in 56% yield following the literature procedures[17].
ESI-MS (*m*/*z*): $[M+Na]^+$ calcd for $C_{18}H_{20}N_4O_{10}$ 475.1, found 475.1.
**$^1$H NMR** (400 MHz, $CDCl_3$) δ 8.26 (d, *J* = 9.2 Hz, 1 H, Ar), 7.33 (d, 1 H, Ar), 5.80 (d, *J* = 3.6 Hz, 1 H, H-1), 5.72 (dd, *J* = 9.6, 10.4 Hz, 1 H, H-3), 5.20 (t, 1 H, H-4), 4.30 (dd, *J* = 4.4, 12.4 Hz, 1 H, H-6), 4.10 (m, 2 H, H-5 & H-6), 3.64 (dd, *J* = 3.6, 10.4 Hz, 1 H, H-2), 2.17 (s, 3 H, OAc), 2.09 (s, 3 H, OAc), 2.08 (s, 3 H, OAc).
**$^{13}$C NMR** (100 MHz, $CDCl_3$) δ 170.5, 170.1, 169.7, 160.5, 143.5, 126.1, 116.8, 96.7, 70.3, 69.2, 68.1, 61.5, 60.7, 20.79, 20.75, 20.7.

*4-Nitrophenyl 2-azido-2-deoxy-α-D-glucopyranoside* **50**

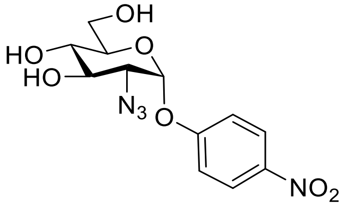

To a solution of 4-nitrophenyl 3,4,6-tri-*O*-acetyl-2-azido-2-deoxy-α-D-glucopyranoside (230 mg, 0.51 mmol) in dry MeOH (15 mL) was added a catalytic amount of sodium at room temperature under $N_2$, and the reaction mixture was stirred for 1 h. The reaction mixture was neutralized with Amberlite 120H, filtered, washed with methanol, concentrated. The resulting residue was purified by flash column chromatography (DCM/MeOH = 15/1), the product was obtained as a white solid (140 mg, 84%).
ESI-MS (*m*/*z*): $[M+Na]^+$ calcd for $C_{12}H_{14}N_4O_7$ 349.1, found 349.1.
**$^1$H NMR** (400 MHz, MeOD) δ 8.25 (d, *J* = 9.2 Hz, 1 H, Ar), 7.31 (d, 1 H, Ar), 5.78(d, *J* = 3.2 Hz, 1 H, H-1), 4.05 (dd, *J* = 8.8, 10.4 Hz, 1 H, H-3), 3.76 (dd, *J* = 2.4, 12.0 Hz, 1 H, H-6), 3.70 (dd, *J* = 4.8 Hz, 1 H, H-6), 3.60 (m, 1 H, H-5), 3.50 (dd, *J* = 8.8, 10.0 Hz, 1 H, H-4), 3.42 (dd, *J* = 3.6, 10.4 Hz, 1 H, H-2).
**$^{13}$C NMR** (100 MHz, MeOD) δ 161.6, 142.9, 125.5, 116.7, 97.2, 74.2, 71.5, 70.3, 63.0, 60.9.

*4-Nitrophenyl 2-amino-2-deoxy-α-D-glucopyranoside* **51**

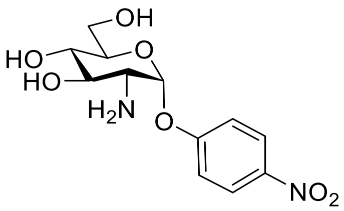

A suspension of starting material (50 mg, 0.15 mmol), $Ph_3P$ (56 mg, 0.21 mmol, 1.4 eq), silica gel (20 mg) in THF (6 mL) and $H_2O$ (1.5 mL) was stirred overnight at 50°C[18]. The reaction mixture was evaporated. The resulting residue was purified by flash column chromatography (dry loaded, DCM/MeOH = 9/1 and 6/1 with 1% TEA), the product was obtained as a white solid (36 mg, 78%).

ESI-MS (*m/z*): $[M+H]^+$ calcd for $C_{12}H_{16}N_2O_7$ 301.1, found 301.1.

**$^1$H NMR** (400 MHz, $D_2O$) δ 8.26 (d, *J* = 9.2 Hz, 1 H, Ar), 7.31 (d, 1 H, Ar), 5.77(d, *J* = 3.2 Hz, 1 H, H-1), 3.83 (t, *J* = 9.6 Hz, 1 H, H-3), 3.76 (d, *J* = 3.6 Hz, 2 H, H-6), 3.70 (m, 1 H, H-5), 3.53 (t, *J* = 9.6 Hz, 1 H, H-4), 3.00 (dd, *J* = 3.6, 10.0 Hz, 1 H, H-2).

**$^{13}$C NMR** (100 MHz, $D_2O$) δ 161.8, 142.5, 126.2, 116.9, 97.6, 74.0, 73.4, 69.6, 60.5, 54.9.

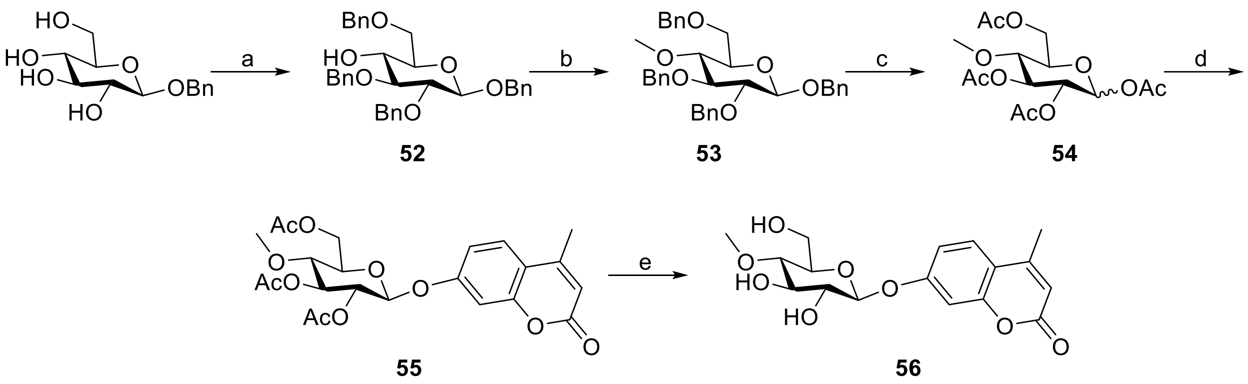

**Scheme S11:** Synthesis of 4-O-methyl-glucoside substrate. Reagents and conditions: a) DMF, p-TSA, benzaldehyde dimethyl acetal, 50 °C; DMF, NaH, BnBr, 0 °C~rt; THF, Na(CN)$BH_3$, HCl/$Et_2O$, rt; 60%. b) THF, NaH, MeI, rt, 100%. c) EtOAc, MeOH, Pd/C, rt; Pyridine, $Ac_2O$, rt, 100%. d) $CH_2Cl_2$, $Ac_2O$, HBr/AcOH, 0 °C. f) $CH_2Cl_2$, 5% NaOH, TBAB, Mu, rt, 70%. e) MeOH, Na, rt, 93%.

*Benzyl 2,3,6-Tri-O-benzyl β-D-glucopyranoside* **52**

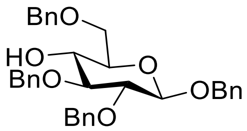

A flask with benzyl β-D-glucopyranoside (5.78 g, 21.4 mmol), DMF (100 mL), *p*-TSA (350 mg) and benzaldehyde dimethyl acetal (4 mL, 26.7 mmol, 1.2 eq) was attached to a Büchi evaporator, rotated, evacuated and lowered into a water bath at 50 ºC so that DMF refluxed in the vapor duct. The mixture was rotated for 2 h at the same condition, and monitored by TLC. The reaction mixture was neutralized with trimethylamine (3 mL), and evaporated under reduced pressure to afford a residue. The resulting residue was purified by a silica gel plug (3:1 and 3:2, petroleum ether-EtOAc) to afford benzyl 4,6-*O*-benzylidene β-D-glucopyranoside as a white solid. To the solution of the white solid in dry DMF (100 mL) was added NaH (4.3 g, 60 %, 107.5 mmol, 5 eq), and stirred for 1.5 h room temperature. The reaction mixture was cooled down with an ice bath, and BnBr (12.7 mL, 106.8 mmol, 5 eq) was added dropwise, stirred and gradually warmed to room temperature. The reaction was monitored by TLC. After completion, the reaction was quenched by addition of methanol at 0 °C. The solvents were evaporated under reduced pressure to afford a residue. The residue was dissolved in EtOAc (500 mL), washed with water (300 mL) and brine (300 mL), dried over $MgSO_4$, filtered and evaporated. The resulting residue was purified by a silica gel plug (15:1 and 9:1, petroleum ether-EtOAc) to afford benzyl 2,3-di-*O*-benzyl-4,6-*O*-benzylidene β-D-glucopyranoside as a white solid. A mixture of the white solid and 4Å molecular sieves (2.5 g) in dry THF (150 mL) was stirred at room temperature under $N_2$. To the mixture were added sodium cyanoborohydride (4.9 g, 78.0 mmol, 3.6 eq) as one portion and dropwise 2 M HCl in ether till no gas generating, and the mixture was stirred for another hour at the same condition, filtered through a short pad of Celite and washed with THF. After evaporation, the residue was dissolved in $CH_2Cl_2$ (500 mL), washed with water (250 mL), saturated $NaHCO_3$ (250 mL) and brine (250 mL), dried over $MgSO_4$. The crude product was purified by flash column chromatography (PE/ EtOAc = 5/1 & 4/1,) to afford the product as a white solid (6.93 g, 60%).
EIS-MS (*m*/*z*): $[M+Na]^+$ calcd for $C_{34}H_{36}O_6$ 563.2, found 563.5.
**$^1$H NMR** (300 MHz, $CDCl_3$) δ 7.35 (m, 20 H, Ar), 5.00 - 4.58 (m, 8 H, Ar-$CH_2$), 4.55 (d, *J* = 7.4 Hz, 1 H, H-1), 3.82 (dd, *J* = 3.8, 10.4 Hz, 1 H, H-6), 3.75 (dd, *J* = 5.2 Hz, 1 H, H-6), 3.67 - 3.44 (m, 4 H, H-2, H-3, H-4 & H-5).
**$^{13}$C NMR** ($CDCl_3$, 75 MHz) δ 138.8, 138.5, 138.2, 137.5, 128.7, 128.6, 128.5, 128.3, 128.2, 128.1, 128.0, 127.9, 102.8, 84.3, 81.9, 75.4, 75.0, 74.3, 73.8, 71.7, 71.3, 70.4.

*Benzyl 2,3,6-tri-O-benzyl-4-O-methyl-β-D-glucopyranoside* **53**

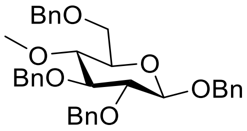

A suspension of tetra-*O*-benzyl-β-D-glucopyranoside (1.0 mmol), anhydrous THF (30 mL) and NaH (200 mg, 60%, 5.0 mmol, 5.0 eq) was stirred for 1 h at room temperature under $N_2$. To the suspension was added MeI (0.25 mL), and the reaction mixture was stirred overnight at the same condition, quenched by addition of methanol. The solvents were removed under reduced pressure. The resulting residue was dissolved in EtOAc (60 mL), washed with water (30 mL) and brine (30 mL), dried over $MgSO_4$, filtered and concentrated. The resulting residue was purified by flash column chromatography (petroleum ether / EtOAc = 9/1 & 5/1) to afford the product as a slightly yellow syrup in 98% yield.
EIS-HRMS (*m/z*): $[M+Na]^+$ calcd for $C_{35}H_{38}O_6$ 577.2566, found 577.2591.
**$^1$H NMR** (300 MHz, $CDCl_3$) δ 7.46 (m, 20 H, Ar), 5.10 - 4.69 (m, 8 H, Ar-$CH_2$), 4.61 (d, 1 H, *J* = 7.6 Hz, H-1), 3.89 (dd, *J* = 2.0, 10.8 Hz, 1 H, H-6), 3.83 (dd, *J* = 4.4 Hz, 1 H, H-6), 3.63 (m, 2 H, H-3 & H-2), 3.60 (s, 3 H, $OCH_3$), 3.51 (m, 1 H, H-5), 3.45 (dd, *J* = 9.6, 8.0 Hz, 1 H, H-4).
**$^{13}$C NMR** ($CDCl_3$, 75 MHz) δ 138.9, 138.7, 138.6, 137.8, 128.7, 128.6 (3 C), 128.58, 128.4, 128.2 (2 C), 128.0, 127.93, 127.88, 127.85, 127.84, 102.8, 84.9, 82.4, 80.1, 75.8, 75.3, 75.1, 73.8, 71.3, 69.3, 60.9.

*1,2,3,6-Tetra-O-acetyl-4-O-methyl-D-glucopyranose* **54**

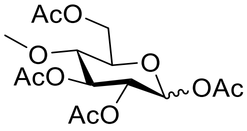

To a solution of tetra-*O*-benzyl-*O*-methyl-β-D-glucopyranoside (1.0 mmol) in EtOAc (3 mL, HPLC grade) and MeOH (30 mL, HPLC grade) was added Pd/C (85 mg, 10% w/w), and the suspension was evacuated and filled with hydrogen three times. The suspension was stirred for 3 days under hydrogen atmosphere at room temperature. After completion, the mixture was filtered through a short pad of Celite, and washed with MeOH. The solvents were evaporated under reduced pressure and dried under vacuum for 0.5 h. The resulting residue was treated with pyridine (5 mL) and acetic anhydride (2 mL) overnight at room temperature. The solvents were evaporated, and water (10 mL) was added and stirred for 0.5 h at room temperature, extracted with ethyl acetate (3 x 20 mL), washed with 1 M HCl (30 mL), saturated $NaHCO_3$ (2 x 30 mL) and brine (30 mL), dried over $MgSO_4$. The resulting residue was purified by flash column chromatography (petroleum ether / EtOAc = 3/1 & 2/1) afford the product as a syrup in quantative yield.
EIS-MS (*m/z*): $[M+Na]^+$ calcd for $C_{15}H_{22}O_{10}$ 385.1, found 385.4.
**$^1$H NMR** ($CDCl_3$, 300 MHz, β-anomer) δ 5.66 (d, *J* = 8.4 Hz, 1 H, H-1), 5.19 (t, *J* = 9.4 Hz, 1 H, H-3), 4.99 (dd, 1 H, H-2), 4.30 (dd, *J* = 2.0, 11.9 Hz, 1 H, H-6), 4.24 (brd, 1 H, H-6), 3.65 (m, 1 H, H-5), 3.43 (s, 3 H, $OCH_3$), 3.39 (t, 1 H, *J* = 9.6 Hz, H-4), 2.09 (s, 3 H, OAc), 2.08 (s, 3 H, OAc), 2.06 (s, 3 H, OAc), 1.98 (s, 3 H, OAc).

*Methylumbelliferyl 2,3,6-tri-O-acetyl-4-O-methyl-β-D-glucopyranoside* **55**

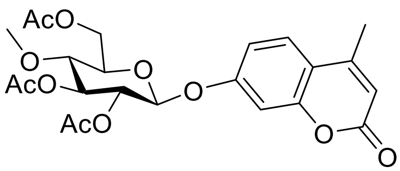

1,2,3,6-Tetra-*O*-acetyl-4-*O*-methyl-D-glucopyranose (1.0 mmol) was treated with DCM (1 mL), acetic anhydride (0.1 mL) and HBr/AcOH (0.8 mL, 30 % w/w) at 0 °C overnight. The reaction mixture was diluted with DCM (10 mL), washed with cold water (2 x 5 mL), cold $NaHCO_3$ (2 x 5 mL) and cold brine (5 mL), dried over $MgSO_4$. After evaporation, crude α-D-glycosyl bromide was obtained. A mixture of α-glycosyl bromide (1.0 mmol), tetrabutyl ammonium bromide (TBAB, 322 mg, 1.0 eq), DCM (3 mL), 5% NaOH (1.5 mL) and 7-hydroxyl-4-methyl-coumarin (Mu, 352 mg, 2.0 eq) was stirred vigorously overnight at room temperature, diluted with DCM (50 mL), washed with 1 M NaOH (2 x 20 mL), water (25 mL) and brine (25 mL), dried over

MgSO$_4$, filtered and concentrated. The resulting residue was purified by flash column chromatography (petroleum ether / EtOAc = 3/2 & 1/1) to afford the product as a white solid in 70% yield.
**EIS-HRMS**(*m/z*): [M+Na]$^+$ calcd for $C_{23}H_{26}O_{11}$ 501.1373, found 501.1360.
**$^1$H NMR** ($CDCl_3$, 400 MHz) δ 7.49 (d, *J* = 8.8 Hz, 1 H, Ar), 6.94 (d, *J* = 2.4 Hz, 1 H, Ar), 6.90 (dd, *J* = 2.4, 8.7 Hz, 1 H, Ar), 6.16 (d, *J* = 1.1 Hz, 1 H, Ar), 5.26 (t, *J* = 8.7, 9.0 Hz, 1 H, H-3), 5.18 (t, *J* = 7.7, 9.1 Hz, 1 H, H-2), 5.12 (d, *J* = 7.7 Hz, 1 H, H-1), 4.41 (dd, *J* = 2.5, 12.0 Hz, 1 H, H-6), 4.26 (dd, *J* = 6.0, 12.0 Hz, 1 H, H-6), 3.75 (m, 1 H, H-5), 3.45 (s, 3 H, $OCH_3$), 3.44 (t, *J* = 9.4 Hz, 1 H, H-4), 2.38 (d, *J* = 1.0 Hz, 3 H, Ar-$CH_3$), 2.11 (s, 3 H, OAc), 2.10 (s, 3 H, OAc), 2.02 (s, 3 H, OAc).
**$^{13}$C NMR** ($CDCl_3$, 100 MHz) δ 170.8, 170.1, 169.8, 160.9, 159.4, 155.0, 152.3, 125.8, 115.5, 113.9, 113.2, 104.3, 98.4, 77.7, 74.8, 73.6, 71.5, 62.7, 60.6, 21.0, 20.9, 20.8, 18.8.

*Methylumbelliferyl 4-O-methyl-β-D-glucopyranoside* **56**

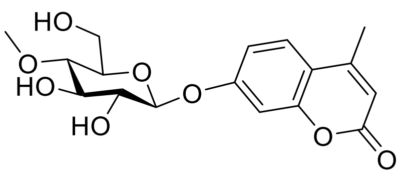

A solution of methylumbelliferyl 2,3,6-tri-*O*-acetyl-4-*O*-methyl-β-D-glucopyranoside (1.0 mmol) in dry MeOH (60 mL) was treated with a catalytic amount of sodium under $N_2$ at room temperature. The white precipitate was filtered and washed with methanol. The mother liquor was neutralized with Amberlite 120 H, concentrated, and precipitated out with methanol and diethyl ether. The combined white solid was dried under vacuum to afford the product as a white solid in 93% yield.
**EIS-HRMS**(*m/z*): [M+Na]$^+$ calcd for $C_{17}H_{20}O_8$ 375.1056, found 375.1057.
**$^1$H NMR** ($CD_3OD$, 300 MHz) δ 7.72 (d, *J* = 8.8 Hz, 1 H, Ar), 7.12 (d, 1 H, Ar), 7.08 (s, 1 H, Ar), 6.22 (s, 1 H, Ar), 5.04 (d, *J* = 7.6 Hz, 1 H, H-1), 3.90 (dd, *J* = 1.4, 12.2 Hz, 1 H, H-6), 3.73 (dd, *J* = 4.6 Hz, 1 H, H-6), 3.61 (s, 3 H, $OCH_3$), 3.65 - 3.50 (m, 3 H, H-2, H-3 & H-5), 3.23 (t, 1 H, *J* = 9.3 Hz, H-4), 2.47 (s, 3 H, Ar-$CH_3$).
**$^{13}$C NMR** ($CD_3OD$, 75 MHz) δ 160.1 (2 C), 154.4, 153.3, 126.4, 114.1, 113.3, 111.7, 103.1, 99.7, 78.9, 76.2, 75.7, 73.3, 60.2, 59.7, 18.1.

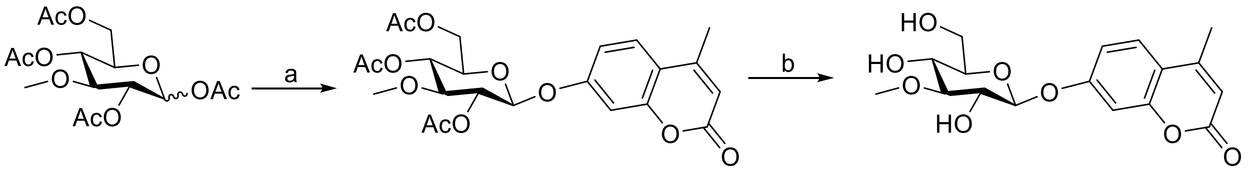

**57** **58**

**Scheme S12:** Synthesis of 3-O-methyl-glucoside substrate Reagents and conditions: a) DCM, 5% NaOH, TBAB, MuOH, rt, 63%. b) MeOH, Na, rt, 95%.

*Methylumbelliferyl 2,4,6-tri-O-acetyl-3-O-methyl-β-D-glucopyranoside* **57**

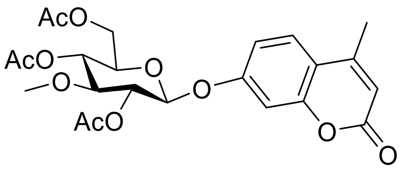

1,2,4,6-Tetra-*O*-acetyl-3-*O*-methyl-D-glucopyranose (1.0 mmol) was treated with DCM (1 mL), acetic anhydride (0.1 mL) and HBr/AcOH (0.8 mL, 30 % w/w) at 0 °C overnight. The reaction mixture was diluted with DCM (10 mL), washed with cold water (2 x 5 mL), cold $NaHCO_3$ (2 x 5 mL) and cold brine (5 mL), dried over $MgSO_4$. After evaporation, crude α-D-glycosyl bromide was obtained. A mixture of α-glycosyl bromide (1.0 mmol), tetrabutyl ammonium bromide (TBAB, 322 mg, 1.0 eq), DCM (3 mL), 5% NaOH (1.5 mL) and 7-hydroxyl-4-methyl-coumarin (Mu, 352 mg, 2.0 eq) was stirred vigorously overnight at room

temperature, diluted with DCM (50 mL), washed with 1 M NaOH (2 x 20 mL), water (25 mL) and brine (25 mL), dried over $MgSO_4$, filtered and concentrated. The resulting residue was purified by flash column chromatography (petroleum ether / EtOAc = 3/2 & 1/1) to afford the product as a white solid in 63% yield.
**EIS-HRMS**(*m*/*z*): $[M+Na]^+$ calcd for $C_{23}H_{26}O_{11}$ 501.1373, found 501.1370.
**$^{1}$H NMR** ($CDCl_3$, 300 MHz) δ 7.52 (d, *J* = 9.0 Hz, 1 H, Ar), 6.92 (m, 2 H, Ar), 6.19 (d, 1 H, *J* = 1.1 Hz, Ar), 5.30 (dd, *J* = 9.3 Hz, 1 H, H-2), 5.13 (t, *J* = 9.5 Hz, 1 H, H-4), 5.08 (d, *J* = 7.7 Hz, 1 H, H-1), 4.23 (dd, *J* = 6.0, 12.3 Hz, 1 H, H-6), 4.16 (dd, *J* = 2.7 Hz, 1 H, H-6), 3.82 (m, 1 H, H-5), 3.61 (t, *J* = 9.3 Hz, 1 H, H-3), 3.47 (s, 3 H, $OCH_3$), 2.41 (d, *J* = 1.0 Hz, 3 H, Ar-$CH_3$), 2.14 (s, 3 H, OAc), 2.13 (s, 3 H, OAc), 2.12 (s, 3 H, OAc).
**$^{13}$C NMR** ($CDCl_3$, 100 MHz) δ 170.9, 169.6, 169.3, 161.0, 159.6, 155.1, 152.4, 125.9, 115.5, 114.4, 113.3, 103.9, 98.8, 81.4, 72.9, 71.7, 69.0, 62.4, 59.3, 21.0 (2 C), 20.9, 18.9.

*Methylumbelliferyl 3-O-methyl-β-D-glucopyranoside* **58**

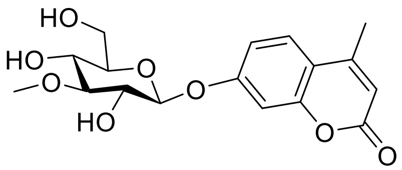

A solution of methylumbelliferyl 2,4,6-tri-*O*-acetyl-3-*O*-methyl-β-D-glucopyranoside (1.0 mmol) in dry MeOH (60 mL) was treated with a catalytic amount of sodium under $N_2$ at room temperature. The white precipitate was filtered and washed with methanol. The mother liquor was neutralized with Amberlite 120 H, concentrated, and precipitated out with methanol and diethyl ether. The combined white solid was dried under vacuum to afford the product as a white solid in 95% yield.
**EIS-HRMS**(*m*/*z*): $[M+Na]^+$ calcd for $C_{17}H_{20}O_8$ 375.1056, found 375.1056.
**$^{1}$H NMR** ($d_6$-DMSO + $D_2O$, 400 MHz) δ 7.71 (d, *J* = 9.6 Hz, 1 H, Ar), 7.03 (m, 2 H, Ar), 6.24 (d, *J* = 0.8 Hz, 1 H, Ar), 5.05 (d, *J* = 7.6 Hz, 1 H, H-1), 3.68 (dd, *J* = 4.8, 14.4 Hz, 1 H, H-6), 3.51 (s, 3 H, $OCH_3$), 3.45 (m, 2 H, H-5 & H-6), 3.35 (dd, 1 H, H-2), 3.25 (t, *J* = 8.8 Hz, 1 H, H-4), 3.11 (t, *J* = 9.0 Hz, 1 H, H-3), 2.40 (d, *J* = 1.0 Hz, 3 H, Ar-$CH_3$).
**$^{13}$C NMR** ($d_6$-DMSO + $D_2O$, 100 MHz) δ 160.0, 154.4, 153.3, 126.4, 114.1, 113.3, 111.7, 103.2, 99.8, 86.3, 76.9, 72.6, 68.9, 60.4, 60.1, 18.1.

#### 1.5. Structures and acronyms for all the substrates

The structures for all the substrates used in the course of this study are presented in this section. Several of these compounds are commercially available. We have already reported the synthesis of 3-, 4-, and 6- amino-glucosides[19]. In addition to the compounds discussed herein, we have also used a diverse set of compounds as potential cofactors and inhibitors, as discussed in sections 2.5.1.5., 2.5.5. and 2.5.7.; these are all commercially available with the exception of dehydro-ascorbic acid and dehydro-isoascorbic acid, which were prepared according to the method previously described[20].

For the substrate we used in the following sections that do not have a common name, we have used an intuitive naming convention instead of numbers, in order to avoid confusion. These consist of a prefix (where applicable), the abbreviated name for the sugar unit followed by the linkage type and the aglycone name. The structures and names of all the substrates are depicted in Fig. S1.

The abbreviation used are: Glc: Glucose, Gal: Galactose, Man: Mannose, 6PGlc: Glucose-6-phosphate, GA: Glucuronic acid, ΔGA: 4,5-unsaturated-glucuronic acid, Xyl: Xylose, Fuc: L-Fucose, GlcNAc: N-Acetylglucosamine, GalNAc: N-Acetylgalactosamine, SQ: sulfoquinovose, $6SO_4$Glc: glucose-6-sulfate, Cell: cellobiose, SC: Thio-acetal linker, 3K: 3keto, MU: 4-Methylumbelliferone, ClMU: 6-Chloro-4-methylumbelliferone, diFMU: 6,8-difluoro-4-methylumbelliferone, JBOMe: Jericho Blue[7] methyl ester, OMe: Methoxy, pNP: paranitrophenol

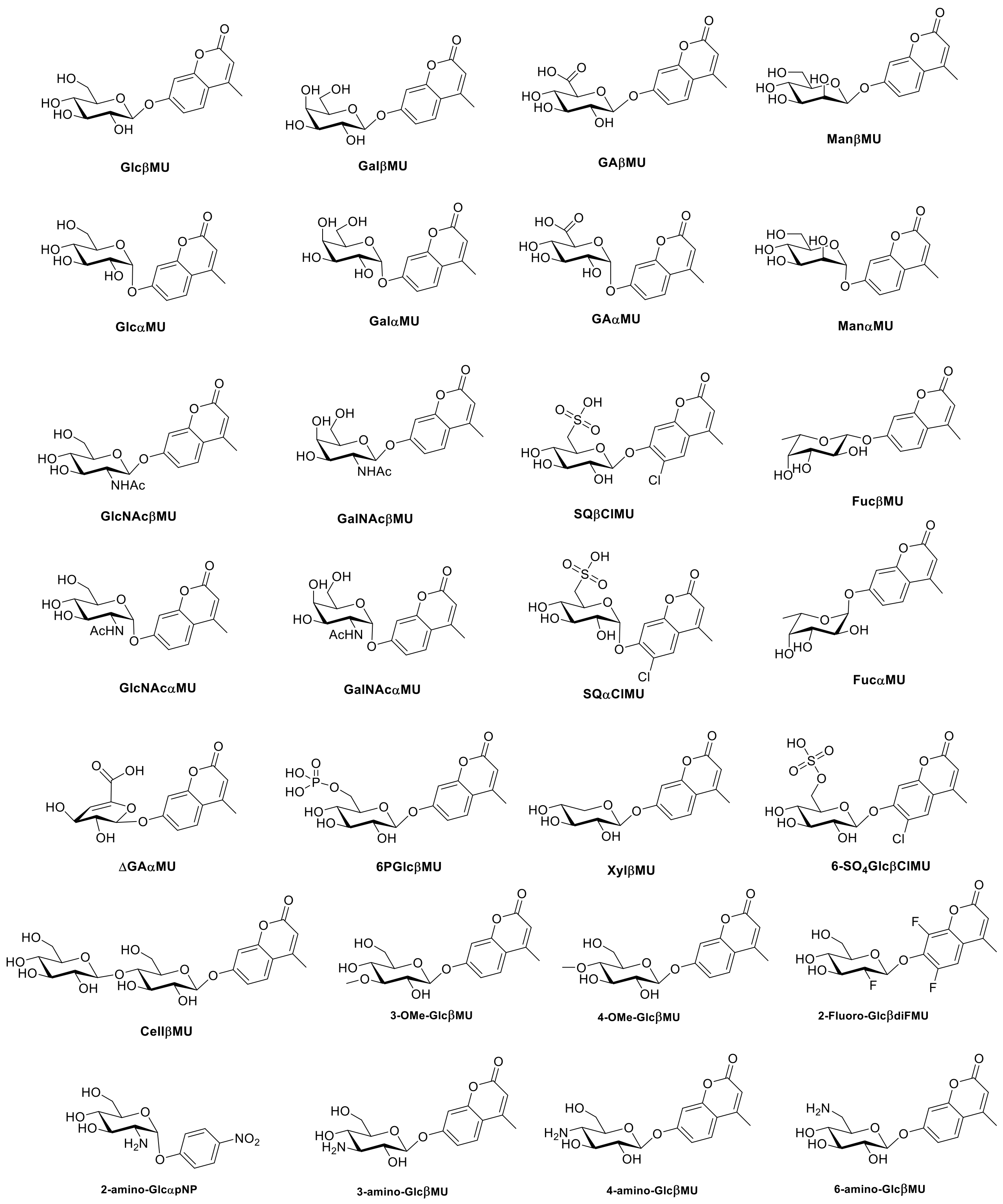

**Figure S1**: structures and acronyms of the substrates used in this study – continued on the next page

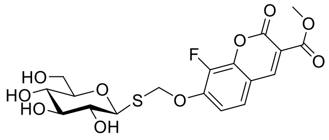

**1: GlcSCJBOMe**

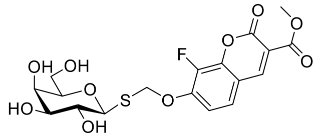

**2: GalSCJBOMe**

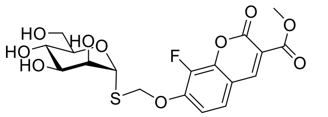

**3: ManSCJBOMe**

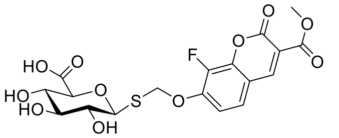

**4: GASCJBOMe**

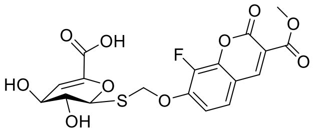

**5: ΔGASCJBOMe**

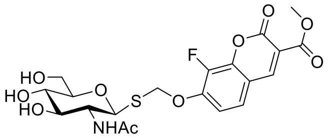

**6: GlcNAcSCJBOMe**

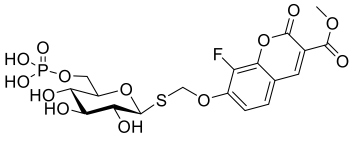

**7: 6PGlcSCJBOMe**

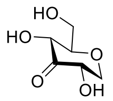

**3K-GlcH**

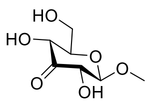

**3K-GlcβOMe**

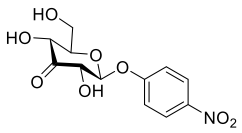

**3K-GlcβpNP**

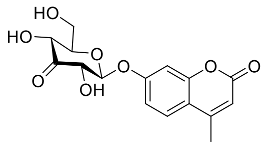

**3K-GlcβMU**

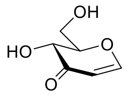

**3K-Glucal**

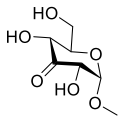

**3K-GlcαOMe**

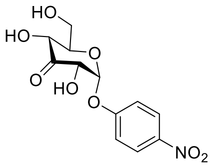

**3K-GlcαpNP**

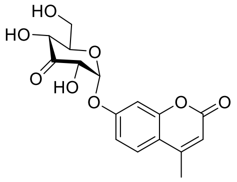

**3K-GlcαMU**

**Figure S1:** structures and acronyms of the substrates used in this study - cont'd

### 2. Biochemical methods

#### 2.1. Metagenomic screen

The screening procedure is based on that reported by Armstrong *et al*[21] with minor changes. Appropriate numbers of 384-well plates [Corning] were filled with LB media (50 µL/well) containing 100 µg/mL L-Arabinose and 12.5 µg/mL chloramphenicol. These plates were next inoculated with bacteria from the metagenomic library storage plates using a Qpix2 xt robot [Genetix] and then incubated overnight at 37 °C in closed containers to minimize evaporation. Next, 45 µL of screening buffer (100 mM phosphate buffer pH = 7 with 2% Triton X-100 containing 60 µM of each of the substrates **1**, **3**, **5**, **6** and **7**) was added to each well and the plates were again incubated at 37 °C in closed containers. This process was repeated for compounds **2** and **4** separately.
The fluorescence of each well was measured after initial addition of the screening buffer and after 2 and 5 hours and overnight incubation using a Synergy H1 plate reader [BioTek] with an excitation wavelength of 405 nm and an emission wavelength of 454 nm[7].

The Z-score for each well was calculated on a per-plate basis via the equation below (E.1):

$$\text{Z-Score} = (\text{Fluorescence}_{\text{Well}} - \text{Mean Fluorescence}_{\text{All wells}}) / \text{Standard deviation}_{\text{All wells}} \qquad \text{E.1}$$

Figure S1 below shows the plot of the Z-score values for each well.

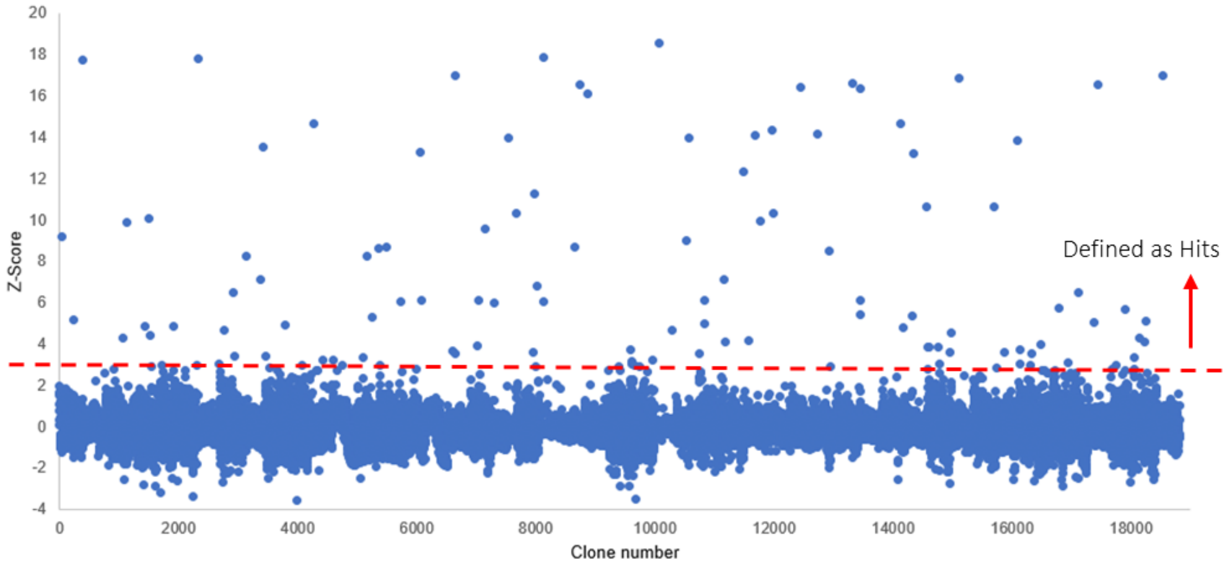

**Figure S2:** Z-score plots for the initial human gut microbiome screening

All the hits with a Z-score value above the depicted threshold were next picked and arrayed into two 96-well plates named “Hit plates”. The clones in the Hit plates were next re-grown and screened with each of the substrates individually in order to find the deconvoluted activity for each well, as well as to validate the hits. The results for this secondary screen for the seven thio-glycoside substrates are shown in figure 2 in the manuscript.

In addition to these, the hit plates were also screened using conventional methylumbelliferyl glycoside substrates. As expected, several of the wells show activity towards multiple substrates. All these results are summarized in Table S1 below. The following abbreviations are used in naming the glycoside substrates: Glc: Glucose, Gal: Galactose, GA: Glucuronic acid, Man: Mannose, 6PGlc: 6-Phosphoglucose, ΔGA: 4,5-unsaturated-glucuronic acid, Xyl: Xylose, Fuc: Fucose, GlcNAc: N-Acetylglucosamine, GalNAc: N-Acetylgalactosamine, SQ: sulfoquinovose.

**Table S1:** Screening results

| Well-ID | Hit substrates | Max Z-score | Substrate of Max Z-score |
|---|---|---|---|
| #1: P1C11 | Compound1; Compound3; Man-α-MU | 12.66 | Compound 3 |
| #2 | Compound5; ΔGAMU; Compound4; 6PGlc-β-MU | 11.29 | ΔGAMU |
| #3 | Gal-α-MU; Fuc-α-MU | 10.92 | Fuc-α-MU |
| #4 | SQ-β-MU; Compound4; GA-β-MU | 10.35 | Compound4 |
| #5 | SQ-β-MU; ΔGAMU; Compound4; GA-β-MU | 10.35 | SQ-β-MU |
| #6 | Xyl-β-MU; Fuc-β-MU; Compound1; Compound7; Glc-β-MU; Compound7 | 9.08 | Compound7 |
| #7 | Xyl-β-MU; Fuc-β-MU; Compound1; Compound7; Glc-β-MU; Compound7 | 8.84 | Compound7 |
| #8 | Fuc-α-MU | 7.98 | Fuc-α-MU |
| #9 | Compound2; Gal-β-MU | 7.47 | Compound2 |
| #10 | Gal-α-MU | 7.44 | Gal-α-MU |
| #11 | Gal-α-MU | 6.93 | Gal-α-MU |
| #12 | Compound2 | 6.88 | Compound2 |
| #13 | SQ-β-MU; SQ-α-MU; GlcNAc-α-MU; 6PGlc-β-MU; Man-α-MU; Glc-α-MU | 6.58 | SQ-α-MU |
| #14 | Fuc-β-MU; Compound1; Glc-β-MU | 6.47 | Glc-β-MU |
| #15 | Xyl-β-MU; Fuc-β-MU; Compound1; Compound7; Glc-β-MU; Compound7 | 6.47 | Xyl-β-MU |
| #16 | Compound2 | 6.38 | Compound2 |
| #17 | Compound5 | 6.02 | Compound5 |
| #18 | GalNAc-β-MU; Compound6; GlcNAc-β-MU; Man-β-MU | 5.99 | Compound6 |
| #19 | Xyl-β-MU; Fuc-β-MU; Compound1; Compound7; Glc-β-MU; Compound7 | 5.99 | Compound1 |
| #20 | Gal-β-MU; 6PGlc-β-MU; GlcNAc-β-MU | 5.79 | Gal-β-MU |
| #21 | Xyl-β-MU; Fuc-β-MU; Compound1; Compound7; Glc-β-MU | 5.71 | Fuc-β-MU |
| #22 | Gal-α-MU; Compound5; Compound4; Glc-β-MU | 5.47 | Compound5 |
| #23 | GalNAc-β-MU; Compound6; GlcNAc-β-MU | 4.64 | GlcNAc-β-MU |
| #24 | Compound5; ΔGAMU | 4.46 | Compound5 |
| #25 | GalNAc-β-MU; GalNAc-α-MU; GlcNAc-β-MU; Man-β-MU | 4.45 | GlcNAc-β-MU |
| #26 | GalNAc-β-MU; GalNAc-α-MU; Compound6; 6PGlc-β-MU; GlcNAc-β-MU | 4.35 | GlcNAc-β-MU |
| #27 | Compound4 | 4.27 | Compound4 |
| #28 | Compound2 | 4.17 | Compound2 |
| #29 | Compound2 | 4.16 | Compound2 |
| #30 | SQ-α-MU; GlcNAc-α-MU; Compound6; Man-α-MU | 4.15 | Compound6 |
| #31 | Gal-α-MU | 3.82 | Gal-α-MU |
| #32: P2B11 | Compound4 | 3.65 | Compound4 |
| #33 | GalNAc-β-MU; GlcNAc-β-MU | 3.59 | GlcNAc-β-MU |
| #34 | Compound2 | 3.26 | Compound2 |
| #35 | GalNAc-β-MU; GalNAc-α-MU; GlcNAc-β-MU; Man-β-MU | 3.24 | GlcNAc-β-MU |

| | | | |
|---|---|---|---|
| #36 | GalNAc-β-MU; GalNAc-α-MU; Man-β-MU | 3.20 | Man-β-MU |
| #37 | GalNAc-β-MU; GalNAc-α-MU; GlcNAc-β-MU; Man-β-MU | 3.18 | GlcNAc-β-MU |
| #38 | GalNAc-β-MU; GalNAc-α-MU; Man-β-MU | 3.14 | Man-β-MU |
| #39 | Compound2 | 3.08 | Compound2 |
| #40 | GalNAc-β-MU; GlcNAc-β-MU; Man-β-MU | 3.08 | GlcNAc-β-MU |
| #41 | SQ-α-MU; GlcNAc-α-MU; Man-α-MU | 3.07 | SQ-α-MU |
| #42 | Gal-β-MU; 6PGlc-β-MU | 3.07 | Gal-β-MU |
| #43 | GalNAc-β-MU; Man-β-MU | 3.04 | Man-β-MU |
| #44 | ΔGAMU | 3.02 | ΔGAMU |
| #45 | Gal-β-MU | 3.00 | Gal-β-MU |
| #46 | Gal-β-MU | 2.89 | Gal-β-MU |
| #47 | Compound2 | 2.75 | Compound2 |
| #48 | GalNAc-β-MU; GalNAc-α-MU; GlcNAc-β-MU; Man-β-MU | 2.65 | GalNAc-β-MU |
| #49 | GalNAc-β-MU; GlcNAc-β-MU; Man-β-MU | 2.64 | Man-β-MU |
| #50 | GalNAc-β-MU; GalNAc-α-MU; GlcNAc-β-MU; Man-β-MU | 2.63 | GalNAc-β-MU |
| #51 | Gal-β-MU | 2.63 | Gal-β-MU |
| #52 | GalNAc-β-MU; GalNAc-α-MU | 2.59 | GalNAc-β-MU |
| #53 | Compound5 | 2.58 | Compound5 |
| #54 | Compound2 | 2.54 | Compound2 |
| #55 | SQ-α-MU; GlcNAc-α-MU; Man-α-MU | 2.48 | SQ-α-MU |
| #56 | GalNAc-β-MU; GalNAc-α-MU; Man-β-MU | 2.46 | Man-β-MU |
| #57 | Compound5 | 2.44 | Compound5 |
| #58 | Gal-β-MU | 2.43 | Gal-β-MU |
| #59 | SQ-α-MU; GlcNAc-α-MU | 2.41 | GlcNAc-α-MU |

#### 2.2. Sequencing results and analysis

Aliquots from the selected wells, containing *E. coli* clones carrying the fosmids, were plated on LB/Agar plates with chloramphenicol (35 mg/mL) and incubated at 37 °C overnight. Single colonies were picked for growth in 10 mL LB media containing chloramphenicol (12.5 µg/mL) and L-arabinose (100 µg/mL) at 37 °C with shaking overnight. The fosmid was then isolated using GeneJET plasmid Miniprep Kit [Thermoscientific] using the manufacturer's protocol with the only adjustment being that the samples were incubated at 65 °C for 3 min with the elution buffer prior to elution.
Miniprepped fosmids were then further purified for sequencing. Therefore, 2 µL 25 mM ATP solution, 6 µL 10X Plasmid-Safe Reaction Buffer and 2 µL Plasmid-Safe ATP-dependent DNase were added to the fosmid sample and incubated at 37 °C for 1 h. Afterwards, the sample was purified using GeneJET Gel Extraction Kit [Thermoscientific]. 60 µL binding buffer were added to each sample, the mixture was transferred to a column and centrifuged for 1 min. 750 µL wash buffer was next added and the sample was spun down again. 20 µL elution buffer were added and the sample was incubated at 65 °C for 3 minutes. Elution was done at 12000 rpm for 2 min. This was done a second time to yield 40 µL eluted DNA. Concentration was measured using a nanodrop photometer and the DNA was stored at -20 °C before being sent for sequencing [Plasmidsaurus].
The sequencing results were then analyzed using the tools available online from National Center for Biotechnology Information website. The most likely sources of the fosmid hits were identified using standard nucleotide Basic Local Alignment Search Tool (BLAST) searches. The results for selected verified hits after removal of the duplicates are shown in Table S2. The open reading frames present on each fosmid were detected using the Open Reading Frame Finder tool (https://www.ncbi.nlm.nih.gov/orffinder/) and each ORF was then analyzed using protein BLAST (https://blast.ncbi.nlm.nih.gov/Blast.cgi). The annotation for each gene was assigned based on the most prevalent similar genes found using BLAST searches. The nucleotide sequences of the hits described in this paper are included in section 5.

**Table S2:** Source organism for select hits

| Well-ID | Hit substrates | Most likely source organism, % cover & % identity of nucleotide sequence |
|---|---|---|
| #1: P1C11 | Compound1; Compound3; Man-α-MU | *Alistipis onderdonkii subsp. vulgaris*, 100% & 99.84% |
| #2 | Compound5; ΔGAMU; Compound4; 6PGlc-β-MU | *Phocaeicola (Bacteroides) vulgatus*, 100% & 99.8% |
| #4 | SQ-β-MU; Compound4; GA-β-MU | *Phocaeicola (Bacteroides) vulgatus*, 99% & 99.75% |
| #5 | SQ-β-MU; ΔGAMU; Compound4; GA-β-MU | *Lawsonibacter asaccharolyticus,* 100% & 99.98% |
| #7 | Xyl-β-MU; Fuc-β-MU; Compound1; Compound7; Glc-β-MU | *Oscillospiraceae bacterium*, 65% & 97.6% |
| #9 | Compound2; Gal-β-MU | *Prevotella copri,* 77% & 98.75% |
| #14 | Fuc-β-MU; Compound1; Glc-β-MU | *Prevotella copri*, 100% & 95.63% |
| #18 | GalNAc-β-MU; Compound6; GlcNAc-β-MU; Man-β-MU | *Phocaeicola (Bacteroides) vulgatus*, 78% & 99.71% |
| #21 | Xyl-β-MU; Fuc-β-MU; Compound1; Compound7; Glc-β-MU | *Prevotellacopri*, 76% and 97.43% |
| #26 | GalNAc-β-MU; GalNAc-α-MU; Compound6; 6PGlc-β-MU; GlcNAc-β-MU | *Phocaeicola (Bacteroides) xylanisolvens*, 56% & 99.11% |
| #27 | Compound4 | No hits |
| #33: P2B11 | Compound4 | *Collinsella stercoris*, 17% & 93.82%, (in terms of protein sequence, the genes are all similar to those found in *Collinsella tanakaei*) |

#### 2.3. Cloning

All the ORFs were cloned into pET28 with a C-terminal His-TAG (or N-terminal His-TAG for the clones specified as pET28(a)). Signal peptides were predicted using SignalP[22] and when present, the DNA bases for their expression were omitted from the cloned construct. Cloning was performed using the polymerase incomplete primer extension (PIPE) cloning method[23] (The only exception being P1C11-DUF1080 which was cloned using the Golden Gate[24] method). Briefly, separate PCR reactions were set up for the vector (v) and the insert (i) using 0.2 µM of each primer (F and R for forward and reverse primers respectively), 5 ng of template (pET28 for the vector and the purified fosmid or genomic DNA for the insert), 5 µL of GC or HF buffer [Thermofisher], 120 µM dNTP mix (30 µM each, final concentration), 0.5 µL Phusion DNA polymerase [Thermofisher] and dd$H_2O$ to a final volume of 25 µL. The program was set to initial denaturation at 98 °C for 2 minutes, then 25 cycles of denaturation at 98 °C (30 s each), annealing was done at a temperature 5 °C lower than the calculated $T_m$ of the primers for 30 s and elongation was at 72 °C for 45 s per kbp. PCR products were controlled via 1% agarose gel electrophoresis and directly transformed into chemical competent DH5α *E. coli* cells using 2 µL of each PCR product and 50 µL of chemical competent cells. All primers were ordered from Thermofisher Scientific and are listed in Table S3.

**Table S3:** Sequence of primers used for cloning

| Primer | Sequence |
|---|---|
| P1C11 ISO-MBP -- IF | AGAAGGAGATATACCATGATCGAAGAAGGTAAACTGG |
| P1C11 ISO-MBP – IR | GCTGTAAACAGTTTACGTTTTGAAATCCTTCCCTCGA |
| P1C11 ISO-MBP – VF | TCGAGGGAAGGATTTCAAAACGTAAACTGTTTACAGCGGT |
| P1C11 ISO-MBP -- VR | AGTTTACCTTCTTCGATCATGGTATATCTCCTTCTTAAAGTTAAACAAAAT |
| P1C11 DUF1080 – IF | ATTGGTCTCGAATGGGCGGCCCGGGGGCCAAA |
| P1C11 DUF1080 – VR | ATTGGTCTCCCATTGGTATATCTCCTTCTTAAAGTTAAACAAA |
| P1C11 DUF1080 – VF | ATTGGTCTCCCACCATCACCATCACCATTGAGATC |
| P1C11 DUF1080 – IR | ATTGGTCTCCGGTGGTCGAGCACCTTGATGCGGATGTTGCGG |
| P1C11 Oxido1 – IF | ACTTTAAGAAGGAGATATACCATGAAGAAGTTCGGGCACACGGC |
| P1C11 Oxido1 – IR | CAATGGTGATGGTGATGGTGACGCGGCATGTCGGGC |
| P1C11 Oxido1 – VF | CTGCCCGACATGCCGCGTCACCATCACCATCACCATTG |
| P1C11 Oxido1 – VR | TGTGCCCGAACTTCTTCATGGTATATCTCCTTCTTAAAGTTAAACAA |
| P1C11 Oxido2 – VR | GACTTTAATTGGATTCATGGTATATCTCCTTCTTAAAGTTAAACAA |
| P1C11 Oxido2 – IR | CAATGGTGATGGTGATGGTGTGCGCGCTTCCTCAGCAAC |
| P1C11 Oxido2 – IF | AAGAAGGAGATATACCATGAATCCAATTAAAGTCGGTCTGATCGGTTTTGG |
| P1C11 Oxido2 – VF | TTGCTGAGGAAGCGCGCACACCATCACCATCACCATTG |
| BT2156 – IF | TAAGAAGGAGATATACCATGCAGTCTAAAACAAAAGCG |
| BT2156 – IR | ATGGTGATGGTGATGGTGTTTAGCGTATGAGCTTTTAA |
| BT2156 – VF | AAAAGCTCATACGCTAAACACCATCACCATCACCATTG |
| BT2156 – VR | TTTTGTTTTAGACTGCATGGTATATCTCCTTCTTAAAGTTAAACAA |
| pET28(a)- BT2157 – IF | CAGCGGCCTGGTGCCGCGCGGGAGTGCTCAGGAAGAG |
| pET28(a)- BT2157 – IR | GCTAGCCATATGGCTGCCTCAGTCCAGAACTTTCACG |
| pET28(a)- BT2157 – VF | CGCGTGAAAGTTCTGGACTGAGGCAGCCATATGGCTAGCATGACTGG |
| pET28(a)- BT2157 – VR | CTCTTCCTGAGCACTCCCGCGCGGCACCAGGCCGC |
| pET28(a)- BT2158 – IF | CTGGTGCCGCGCGGCAGCCACGTTTCCCCAACCGACAA |
| pET28(a)- BT2158 – IR | GTCATGCTAGCCATATGTTATCGTGGCATATCAGGCAACT |
| pET28(a)- BT2158 – VF | CCTGATATGCCACGATAACATATGGCTAGCATGACTGGTGGACAGCAAATGGGT |
| pET28(a)- BT2158 – VR | GTCGGTTGGGGAAACGTGGCTGCCGCGCGGCACCA |
| BT2159 – IF | AAGAAGGAGATATACCATGGATGCACAATTTTCACTTG |
| BT2159 – IR | CAATGGTGATGGTGATGGTGTTTTAAGGAGTTAGTTATTTGTG |
| BT2159 – VF | ATAACTAACTCCTTAAAACACCATCACCATCACCATTG |
| BT2159 – VR | AATTGTGCATCCATGGTATATCTCCTTCTTAAAGTTAAACAAAATT |
| P2B11-Oxido1 – IF | AAGAAGGAGATATACCATGGAAAAGGTGCGTTACGGC |
| P2B11-Oxido1 – IR | CAATGGTGATGGTGATGGTGCTCGCGAACGGGGAAC |
| P2B11-Oxido1 – VF | AAGTTCCCCGTTCGCGAGCACCATCACCATCACCATTG |
| P2B11-Oxido1 – VR | GTAACGCACCTTTTCCATGGTATATCTCCTTCTTAAAGTTAAACAA |
| P2B11-Iso1 – IF | AAGAAGGAGATATACCATGTTAGGAGGCTCCATGGGTAGCG |
| P2B11-Iso1 – IR | AATGGTGATGGTGATGGTGCGACGCGGTCGCGCCA |

| | |
|---|---|
| P2B11-Iso1 – VF | ATTGGCGCGACCGCGTCGCACCATCACCATCACCATTG |
| P2B11-Iso1 – VR | CATGGAGCCTCCTAACATGGTATATCTCCTTCTTAAAGTTAAACAA |
| P2B11-Iso2 – IF | AAGAAGGAGATATACCATGGGATTGATTGGCGTTCA |
| P2B11-Iso2 – IR | CAATGGTGATGGTGATGGTGGAACCAATCCTCATAGCC |
| P2B11-Iso2 – VF | GGCTATGAGGATTGGTTCCACCATCACCATCACCATTG |
| P2B11-Iso2 – VR | AACGCCAATCAATCCCATGGTATATCTCCTTCTTAAAGTTAAACAA |
| P2B11-Oxido2 – VR | AAGAAGGAGATATACCATGATTGTGGGTATGGGAG |
| P2B11-Oxido2 – VR | CAATGGTGATGGTGATGGTGTACATTCGCCCATGAG |
| P2B11-Oxido2 – VR | GGCTCATGGGCGAATGTACACCATCACCATCACCATTG |
| P2B11-Oxido2 – VR | CATACCCACAATCATGGTATATCTCCTTCTTAAAGTTAAACAAAAT |

It was later found out that two of these constructs do not result in expression of soluble or stable proteins. Expression of the protein encoded by P1C11-Iso-MBP yields soluble protein, but the product is extremely unstable. Since only a partial sequence was present in our fosmid, we reasoned that this instability is due to the lack of a full-length protein. In addition, expression of the protein encoded by P2B11-Oxido2 yielded only insoluble products in all conditions tested. We reasoned that this might be fixed by changing the His-TAG position to the N-terminal. We therefore ordered plasmids for expression of the full-length Isomerase from Alistipes species (NCBI Reference Sequence: WP_022333038.1), named AL-Iso-FL and for expression of P2B11-Oxi2, with N-terminal His-TAG, named pET28(a)-P2B11 oxido2 from Twist biosciences. Both of these constructs fixed the issues with their protein products (see section 2.4.).

#### 2.4. Protein purification

All proteins were expressed in *E. coli* BL21(DE3) cells in either LB media (induced with 0.1 mM IPTG) or in LBE-5052 Auto-induction media[25]. The specific growth condition for each protein can be found in Table S4. Cells were harvested at 4000 rpm for 30 min and stored at -20 °C until further purification. Culture pellets were resuspended in lysis buffer (20 mL for 500 mL of culture, Lysis buffer: 50 mM Tris pH 7.0, 150 mM NaCl, 1% glycerol, 20 mM Imidazole, 10 mM $MgSO_4$, 0.2 µL benzonase, 0.5 mg Hen Egg White lysozyme, 1 Tablet of Pierce EDTA-free Protease Inhibitor per 50 mL of buffer). Cell disruption was done via sonication at 35% intensity, 5/10 pulse for 3 min. Insoluble fractions were pelleted via centrifugation at 16000 rpm, 4°C, 30 min. Supernatant was loaded onto Ni-NTA columns, equilibrated in Buffer A (50 mM Tris pH 7.0, 150 mM NaCl, 1% Glycerol). The column was washed with an imidazole gradient from 20 mM to 400 mM over 50 mL of eluent. The fractions were analyzed using SDS-PAGE and fractions containing the protein of interest were concentrated and buffer exchanged into storage buffer, 50 mM Tris pH 7.0, 1% Glycerol, using an ultracentrifugation device (Amicon, Canada).
The proteins were further purified using size exclusion chromatography (Sepharose, Superdex 200) using an ÄKTA go liquid chromatography system (Cytiva) eluting with Tris buffer (50 mM Tris pH 7.0). Fractions showing UV-activity were analyzed using SDS-PAGE, concentrated and directly used for crystallization screens.
Protein concentrations are calculated based on the absorbance of purified proteins at 280 nm, with the extinction coefficients calculated based on the protein sequences using ProtParam expasy tool (https://web.expasy.org/protparam/)[26].

**Table S4:** Expression conditions for all the proteins

| Protein | **AL1/Isomerase** | **AL2/DUF1080** | **AL3/Oxido1** | **AL4/Oxido2** |
|---|---|---|---|---|
| Size (kDa) | 32.7 | 30.9 | 52.4 | 41.4 |
| Plasmid | pET28-AL-Iso-FL | pET28-P1C11 Duf1080 | pET28-P1C11 Oxido1 | pET28-P1C11 Oxido2 |
| Expression condition | 37 °C for 24 hr in LBE-5052 | 30 °C for 36hr in LBE-5052 | 37 °C, O.N. in LB | 37 °C, O.N. in LB |
| Yield (per 1L culture) | 76 mg | 4.5 mg | 40 mg | 100 mg |
| Protein | **BT1/BT2156** | **BT2/BT2157** | **BT3/BT2158** | **BT4/BT2159** |
| Size (kDa) | 33.2 | 31.4 | 52.9 | 43.6 |
| Plasmid | pET28-BT2156 | pET28(a)-BT2157 | pET28(a)-BT2158 | pET28-BT2159 |
| Expression condition | 37 °C, O.N. in LB | 30 °C for 36hr in LBE-5052 | 30 °C for 36hr in LBE-5052 | 37 °C, O.N. in LB |
| Yield (per 1L culture) | 13.5 mg | 100 mg | 2.5 mg | 40 mg |
| Protein | **P2B11- Iso1** | **P2B11-Oxido1** | **P2B11-Oxido2** | **P2B11-Iso2** |
| Size (kDa) | 34.3 | 45.2 | 40.6 | 33.1 |
| Plasmid | pET28-P2B11 Iso1 | pET28-P2B11 Oxido1 | pET28(a)-P2B11 Oxido2 | pET28-P2B11 Iso2 |
| Expression condition | 37 °C, O.N. in LB | 37 °C for 24 hr in LBE-5052 | 37 °C for 24 hr in LBE-5052 | 37 °C, O.N. in LB |
| Yield (per 1L culture) | 20 mg | 100 mg | 3 mg | 22 mg |

#### 2.5. Enzymatic kinetics

Kinetic studies on enzymatic reactions were monitored using a Synergy H1 Hybrid plate reader [BioTek]. The reactions were all performed at room temperature and were done in either black or clear half-area flat-bottom 96-well plates [Corning]. For reactions where the fluorophore is JBOMe, reactions were followed by monitoring fluorescence of the wells using $\lambda_{ex}/\lambda_{em}$ of 405/450 nm, where MU was used as the fluorophore, the values were set to 365/455 nm[7], while for pNP absorbance was monitored at 405 nm. Unless otherwise stated, the rates are reported as concentration of released aglycone per specified unit of time, calculated based on the calibration curve made for the fluorophore/chromophore in the assay buffer and under the same conditions as in the assay. All the assays described herein were based on initial rates of the reaction, that are calculated for the first 1-10 minutes of the reaction, depending on the time necessary to observe the response, as well as the linear response range of the fluorophore/chromophore signal. All the data analysis is performed using GraphPad [Prism]. Wherever error bars are reported, they represent standard deviation (SD) of the reported value measured from at least three (n=3) distinct samples, while the reported values are the mean of these measurements.

Whenever different enzymes are referred to within a series, these abbreviations are used:

1: AL1/Isomerase from *Alistipes sp*., BT1/BT2156 from *Bacteroides thetaiotaomicron,* Iso2 from P2B11
2: AL2/DUF1080 from *Alistipes sp*., BT2/BT2157 from *Bacteroides thetaiotaomicron,* Iso1 from P2B11
3: AL3/Oxidoreductase 1 from *Alistipes sp*., BT3/BT2158 from *Bacteroides thetaiotaomicron,* Oxi1 from P2B11
4: AL4/Oxidoreductase 2 from *Alistipes sp*., BT4/BT2159 from *Bacteroides thetaiotaomicron,* Oxi2 from P2B11

Some of the early assays are repeated twice, once before finding out the effect of adding 3-ketoglycoside co-substrates on rate of the reaction (section 2.5.1.) and once after this (section 2.5.2.). Even though the latter assays are more insightful than the former and essentially replace them, the data for both are included here since they are used as part of the discussion in the manuscript.
Note that the initial assays performed with enzymes from the P1C11 hit, those in sections 2.5.1 and 2.5.2, are done using the protein that was present on P1C11 fosmid, cloned as P1C11-Iso-MBP and not the full-length isomerase analogue that is present in multiple *Alistipes* genomes and which was produced later, called AL-Iso-FL (section 2.3.). In general, the full-length enzyme is significantly more stable than the other and therefore is noticeably more active in catalysing the elimination reaction with its preferred substrates, 3-keto-β-glucosides (section 2.5.4.3.). Using the truncated enzyme however does not significantly alter the results of the initial exploratory assays. For all the assays in later sections where stability of the enzyme was a concern, the full-length AL1 enzyme was used.

##### 2.5.1. Initial assays for AL and BT enzymes: substrate specificity test

To a mixture of four enzymes from either of the two organisms (1 μM each) in 50 mM Tris buffer in pH 7.0, was added 50 μM of the substrates specified. Reactions were followed by measuring the fluorescence of released fluorophores and the initial rates were calculated for the first 10 minutes of the reaction.

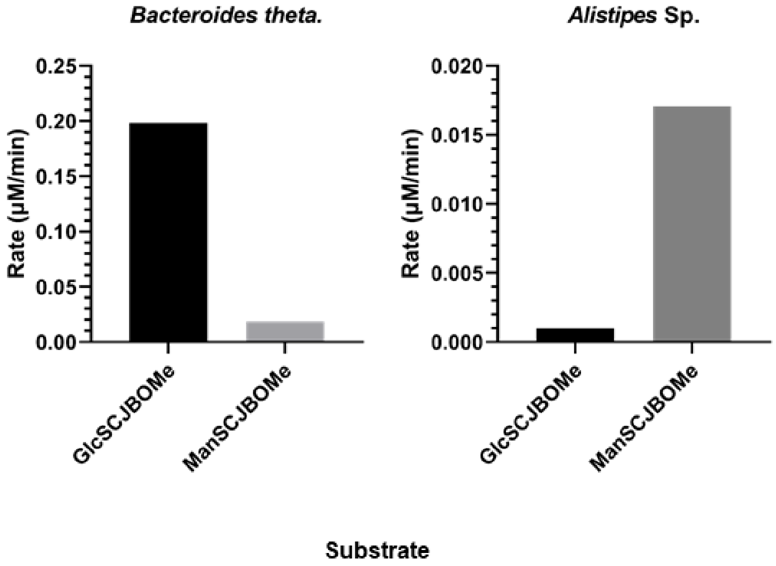

**Figure S3:** Substrate specificity

###### 2.5.1.1. Essential enzymes

To a mixture of 2 μM of each of the specified enzyme in 50 mM Tris buffer pH 7.0, was added 50 μM of the substrates (compound **1** for the BT enzymes and compound **3** for the AL enzymes) and initial rates were calculated for the first 10 minutes of the reactions as depicted below.

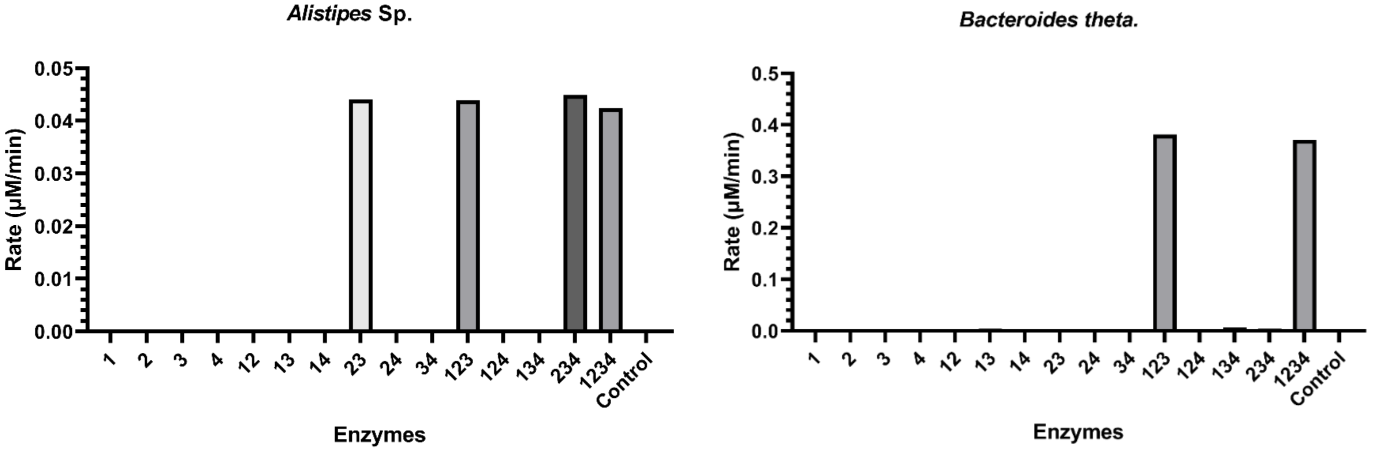

**Figure S4:** Essential enzymes

###### 2.5.1.2. Swapping the enzymes

Solutions of a mixture of the three first enzymes of each series were made in 50 mM Tris buffer pH 7.0, where the concentration of all enzymes was 0.2 µM and the specified enzymes were replaced with their counterpart from the other series, one at a time. The replaced enzyme is specified for each reaction. To these solutions was added 50 µM of the substrates (compound **1** for the BT enzymes and compound **3** for the AL enzymes). Reactions were then followed by measuring the fluorescence of the wells and the initial rates are depicted below. Control denotes wells with no enzymes present.

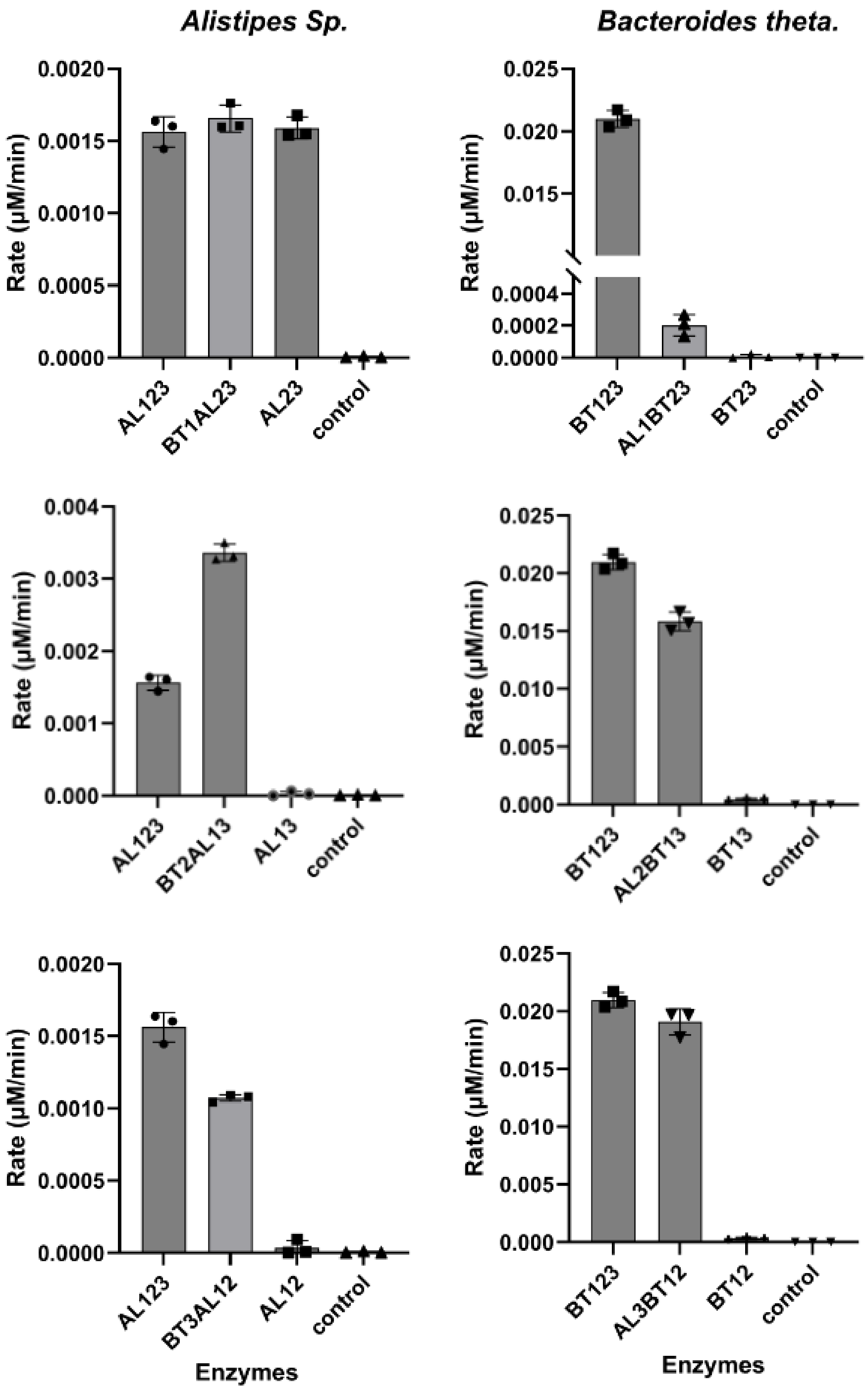

**Figure S5:** Swapping the enzymes, (bars indicate the mean, and error bars the SD of technical replicates (n=3)).

##### 2.5.1.3. Screening for metal cofactors

To a solution containing 0.2 μM each of the four enzymes of either series in 50 mM Tris buffer pH 7.0, 100 μM of the specified metal ion and (when specified) 50 μM of EDTA, was added 50 μM of the substrates (compound **1** for the BT enzymes and compound **3** for the AL enzymes), and the reaction was followed by measuring the fluorescence of released JBOMe over time. Initial rates were calculated for the first 10 minutes of the reactions and are shown below. As controls, solutions containing only enzymes ("no additive") and solutions containing enzymes and 50 μM of EDTA were used, showing that while the activity of the enzymes is metal dependent, none of the added metals is the missing component and the reason for the slow reaction.

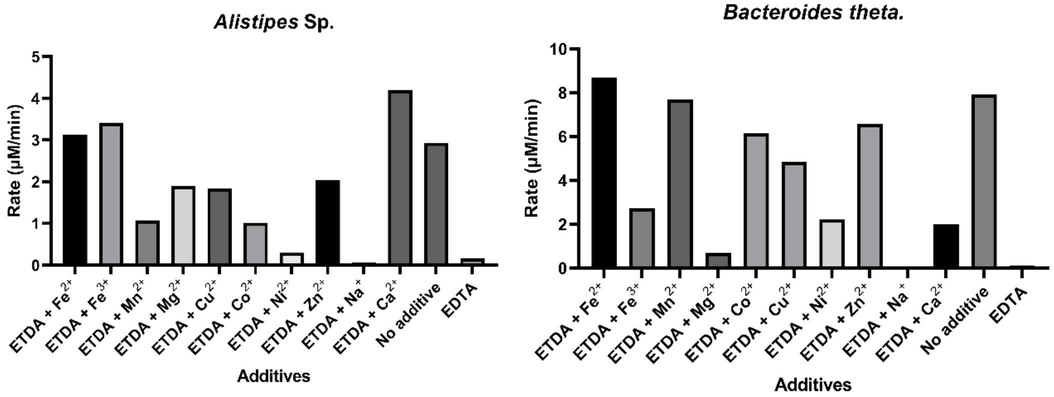

**Figure S6:** Screening for metal cofactors

##### 2.5.1.4. Screening for buffer conditions and ionic strength

To solutions containing 0.2 μM each of the four enzymes of either series in various buffers (all at 50 mM and all adjusted to pH 7.0) was added 50 μM of the substrates (compound **1** for the BT enzymes and compound **3** for the AL enzymes), and the reaction was followed by measuring the fluorescence of released JBOMe over time. Initial rates were calculated for the first 10 minutes of the reactions and are reported below.

Even though HEPES was found to be the best buffer, the rate enhancement was not pronounced. Moreover, we noticed that enzymes are prone to precipitation when stored in any buffer other than TRIS; therefore, we continued using TRIS buffer for storage as well as the rest of the assays in this section.

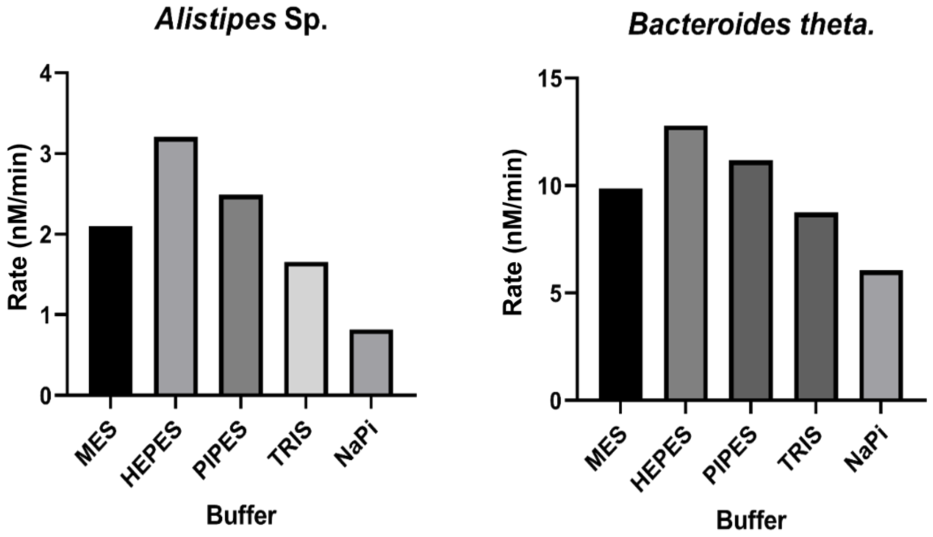

**Figure S7:** Buffer conditions

###### 2.5.1.5. Screening for co-factors

To a solution containing 0.2 µM of each of the four enzymes of either series in TRIS buffers (50 mM at pH 7.0) and 200 µM of the specified cofactor, was added 50 µM of the substrates (compound **1** for the BT enzymes and compound **3** for the AL enzymes), and the reaction was followed by measuring the fluorescence of wells over time. Initial rates were calculated for the first 10 minutes of the reactions and are reported as nM of released fluorophore per minute, calculated based on the calibration curve made for JBOMe separately in buffers containing 200 µM of each cofactor. Making separate calibration curves is necessary in this case, since riboflavin cofactors are colored and hence reduce the fluorescence of the fluorophore. Since none of these cofactors had a dramatic effect in increasing the rate, we also screened a set of other less common cofactors in the same way as above.

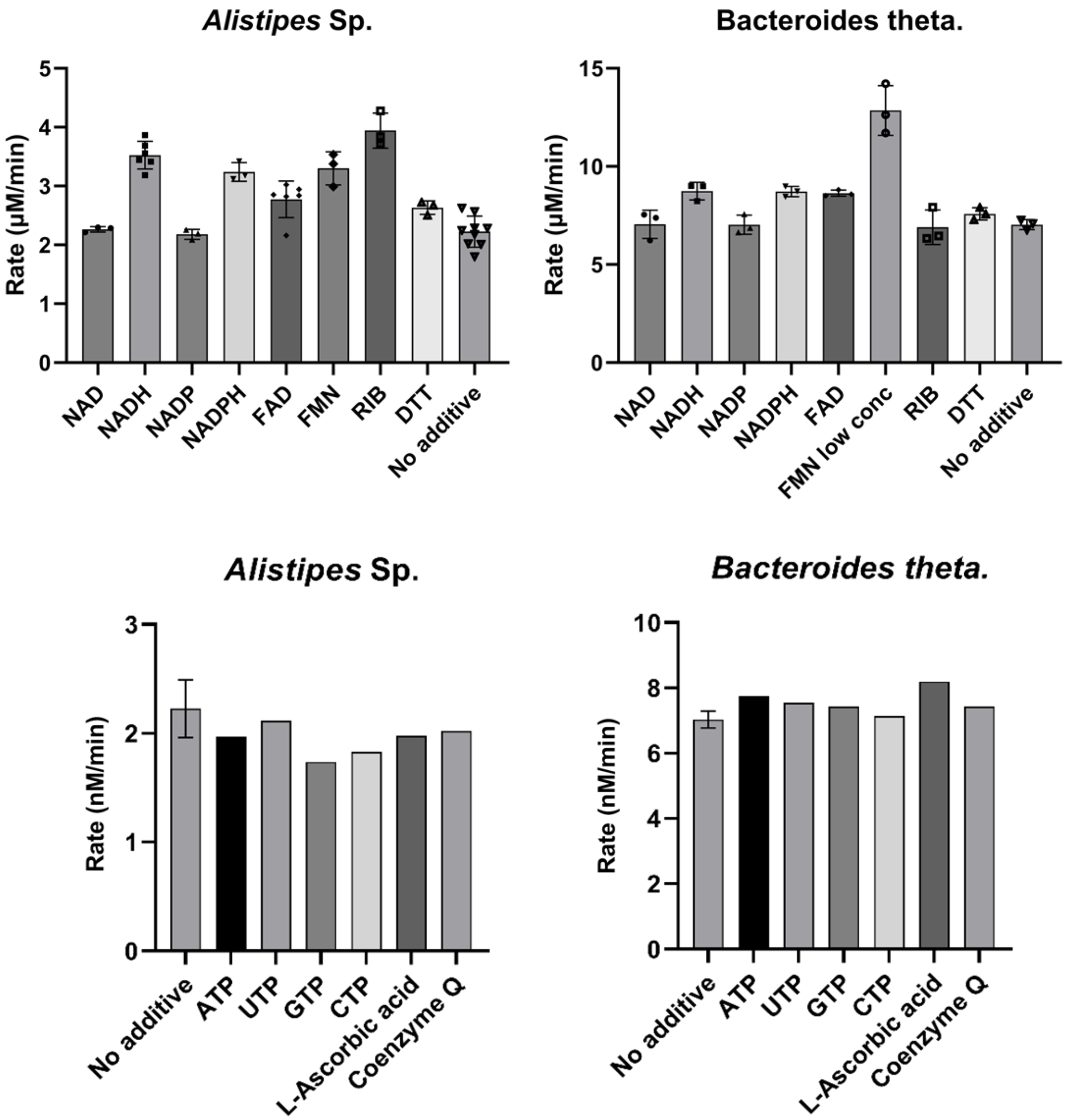

**Figure S8:** Screening for co-factors ((where present, bars indicate the mean, and error bars the SD of technical replicates (n=3)).

##### 2.5.2. Assays for AL and BT operons utilizing 3-Keto-glucosides

To a solution containing 0.02 μM each of the four enzymes of either series in TRIS buffers (50 mM at pH 7.0) and 50 μM of either 3KGlcβOMe or GlcβOMe was added 50 μM of the substrates (compound **1** for the BT enzymes and compound **3** for the AL enzymes).

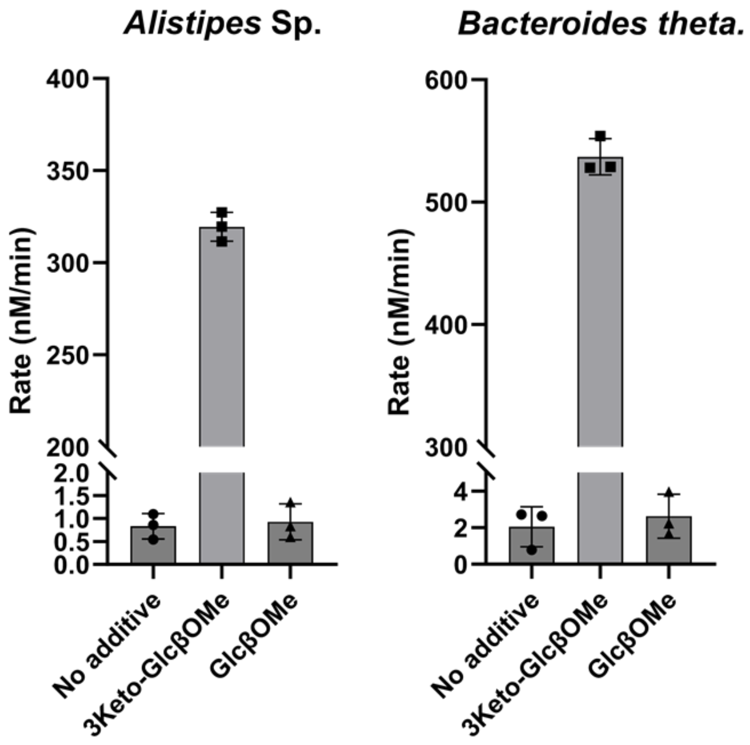

**Figure S9:** Effect of 3-keto co-substrate on rate of reaction (bars indicate the mean, and error bars the SD of technical replicates (n=3)).

Initial rates of the reactions show a dramatic rate increase upon addition of 3KGlcβOMe. Note that all of the results presented prior to this section are indeed dealing with the minor improvements upon the slow rate observed in the "no additive" controls in the graphs above. We therefore first determined the dependence of rate on the concentration of 3KGlcβOMe and then repeated all the experiments above, this time including the optimal amount of 3KGlcβOMe in the reaction buffers.

###### 2.5.2.1. 3-Keto-glucoside co-substrate pseudo Michaelis-Menten kinetics

To a solution containing 0.002 μM each of the four enzymes of either series in TRIS buffers (50 mM at pH 7.0) and varying concentrations of 3KGlcβOMe was added 20 μM of the substrates (compound **1** for the BT enzymes and compound **3** for the AL enzymes).

In both cases, the rate of the reaction is strongly affected by the concentration of 3KGlcβOMe, with substrate inhibition observed at higher concentrations. The graphs below show the apparent $K_m$ and $K_i$ for this compound and both the systems, derived using the built-in auto-inhibition kinetic model in GraphPad [Prism]. For all the rest of the assays, the concentration of 3KGlcβOMe was kept at 50 μM, approximately the optimum amount for rate enhancement.

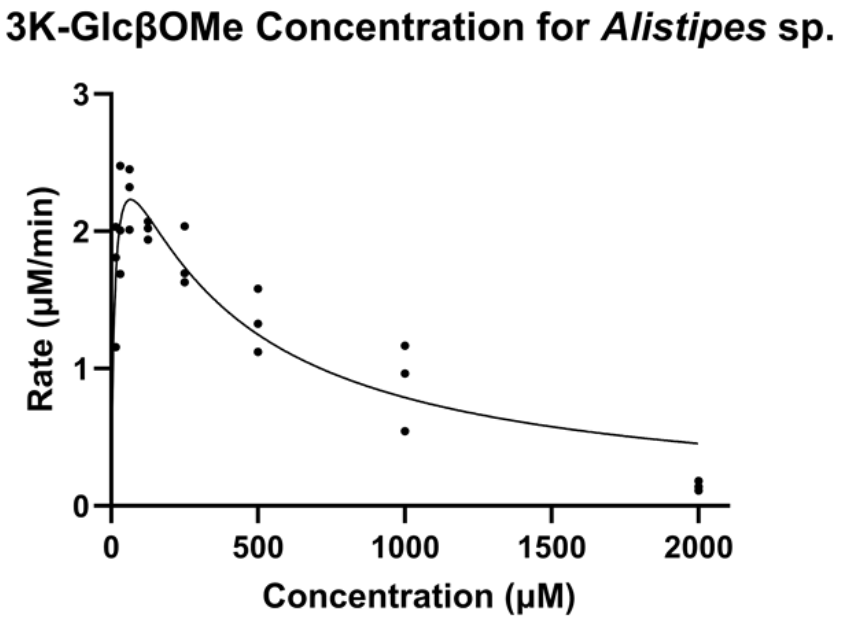

$K_m$ = 13 ± 5 μM

$K_i$ = 350 ± 100 μM

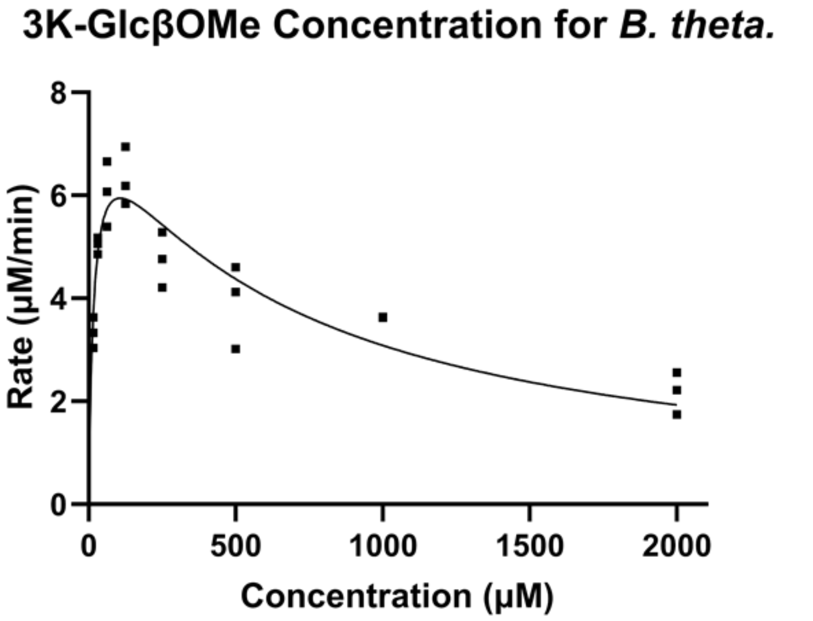

$K_m$ = 18 ± 5 μM

$K_i$ = 650 ± 150 μM

**Figure S10:** Pseudo-Michaelis-Menten kinetics for 3K-GlcβOMe, each data point represents that rate measured for a replicate (n=3).

A similar experiment was also repeated with another synthesized 3-keto substrate, as follows. To a solution containing 0.0002 μM each of the four enzymes of either series in NaPi buffer (25 mM at pH 7.0) and varying concentrations of 3KGlcH was added 20 μM of the substrates (compound **1** for the BT enzymes and compound **3** for the AL enzymes), and the reaction was followed by measuring the fluorescence of released JBOMe over time. Initial rates were calculated for the first 5 minutes of the reactions.

In both cases, the rate of the reaction is strongly affected by the concentration of 3KGlcH, with substrate inhibition observed at higher concentration. The graphs below show the apparent $K_m$ and $K_i$ for this compound and both the systems.

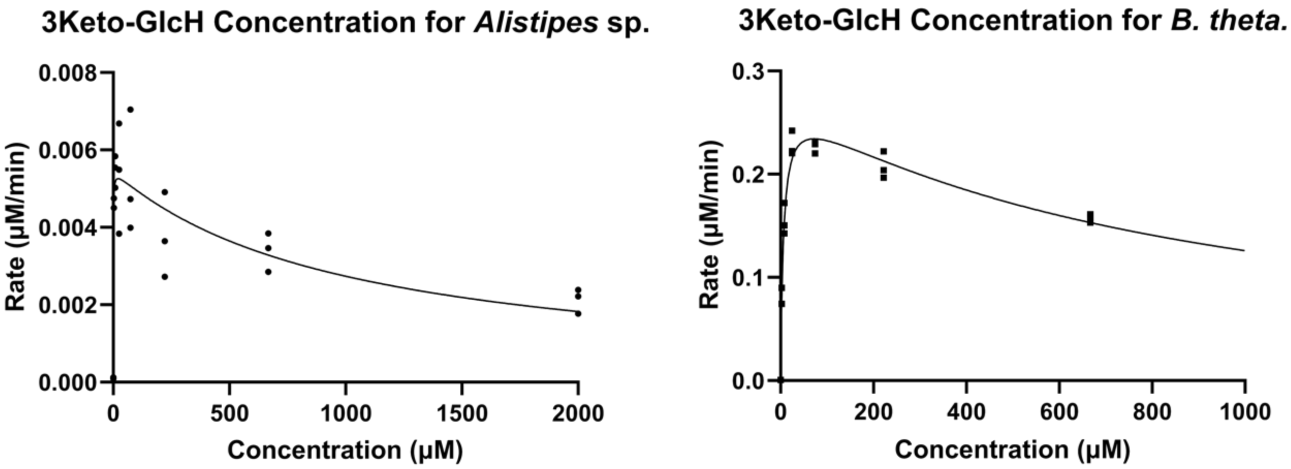

$K_m \leq$ 5 μM
$K_i$ = 350 ± 300 μM

$K_m$ = 6 ± 5 μM
$K_i$ = 860 ± 80 μM

**Figure S11:** Pseudo-Michaelis-Menten kinetics for 3K-GlcH, each data point represents that rate measured for a replicate (n=3).

###### 2.5.2.2. Essential enzymes

To a solution of 0.06 μM of each of the specified enzyme and 50 μM of 3KGlcβOMe in 50 mM Tris buffer pH 7.0, was added 20 μM of the substrates (compound **1** for the BT enzymes and compound **3** for the AL enzymes), and the rate of reaction was followed by measuring the fluorescence of released JBOMe over time.

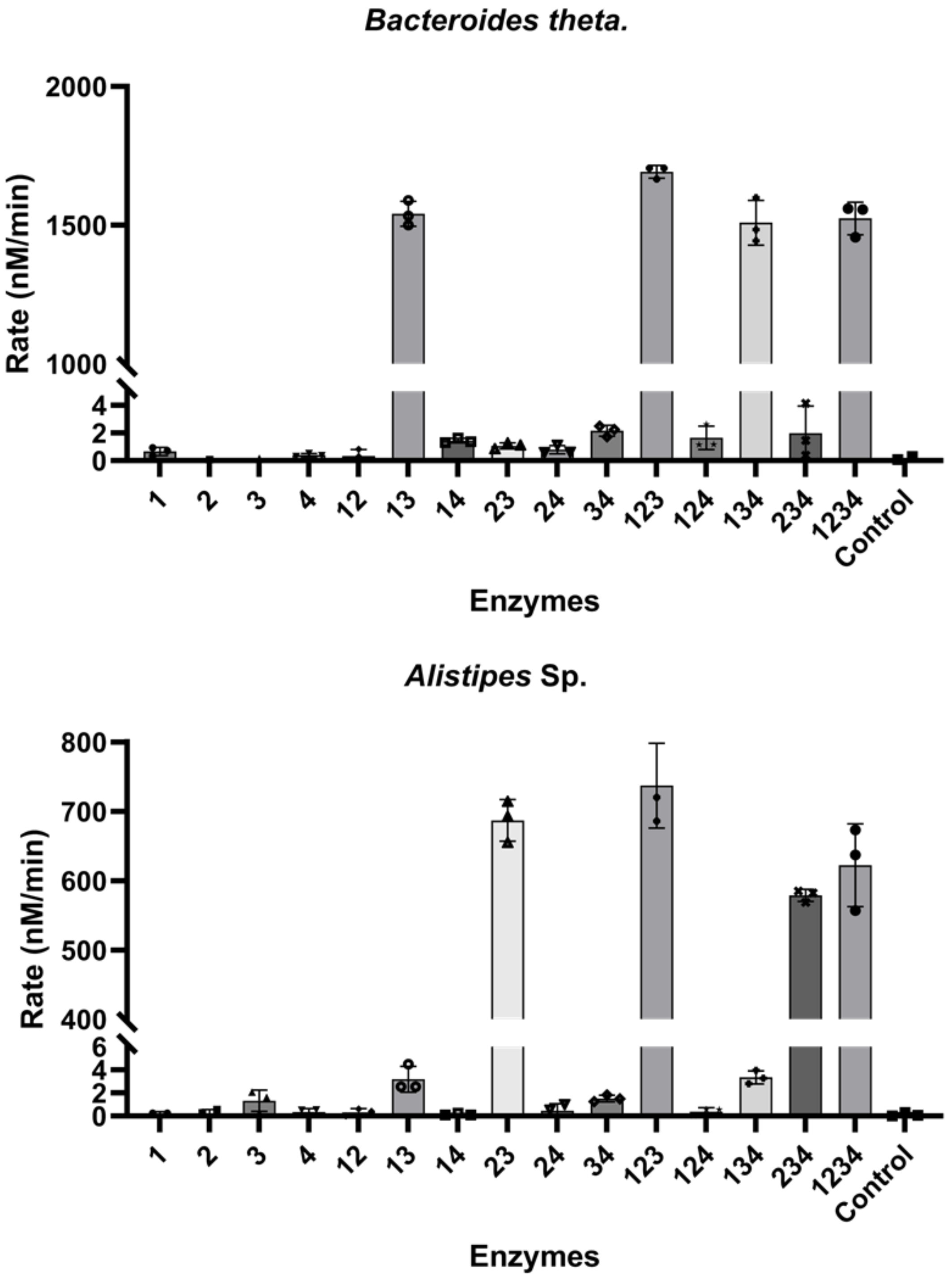

**Figure S12:** Essential enzymes, (bars indicate the mean, and error bars the SD of technical replicates (n=3)).

###### 2.5.2.3. Substrate specificity test

To a solution of 0.02 μM of each of the four enzymes from specified organisms and 50 μM of 3KGlcβOMe in 50 mM Tris buffer pH 7.0, was added 50 μM of the specified substrates, and the rate of reaction was followed by measuring the fluorescence of released fluorophores (JBOMe or MU) over time. Initial rates were calculated for the first 4 minutes of the reactions.

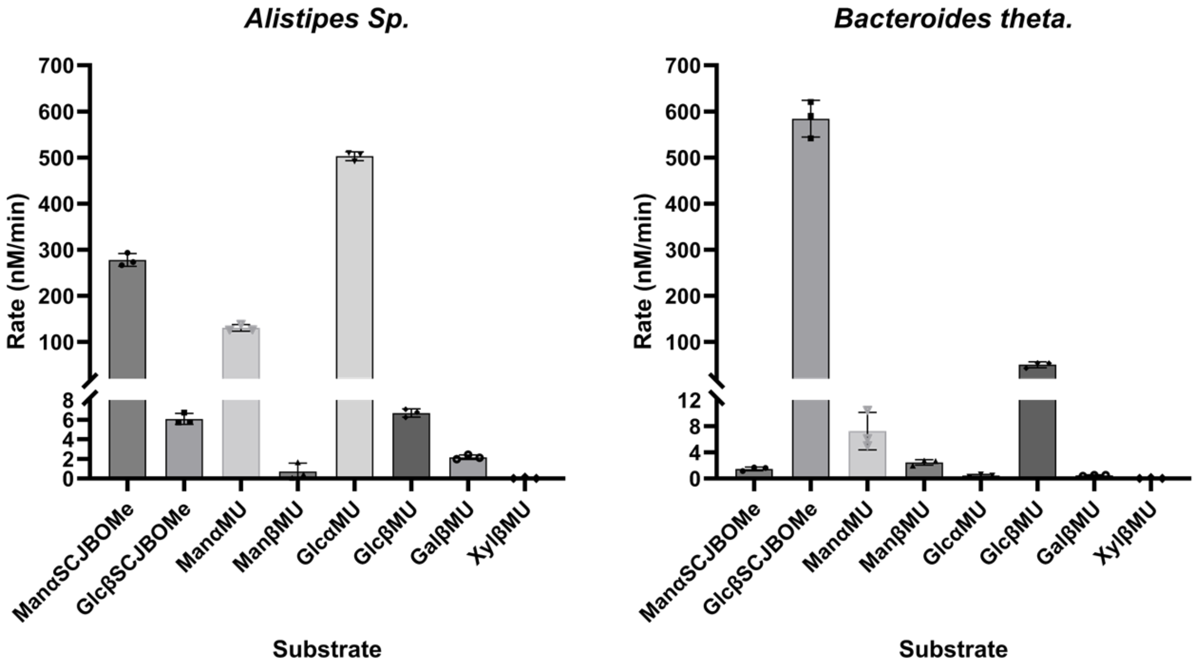

**Figure S13:** Substrate specificity, (bars indicate the mean, and error bars the SD of technical replicates (n=3)).

Here since the stability of all the enzymes makes a difference in their ability to catalyze reactions for β vs α substrates, we later repeated the assay with the full-length AL1 from *Alistipes.* The full-length enzyme is significantly more active in catalyzing the elimination reaction with β-glucosides and therefore the activity on β-glucoside substrates is markedly increased. Still, the four enzymes together are more active on α-glucosides, followed by β-glucosides and α-mannosides.

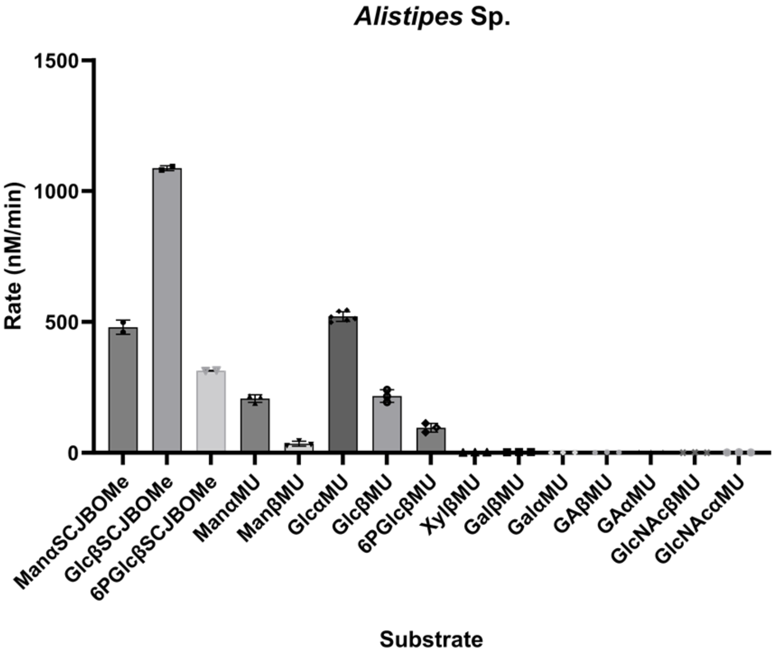

**Figure S14:** Substrate specificity for *Alistipes* enzymes, (bars indicate the mean, and error bars the SD of technical replicates (n=3)).

###### 2.5.2.4. Swapping the enzymes

Solutions of a mixture of the three first enzymes of each series were made in the reaction buffer (50 mM Tris buffer in pH 7.0 containing 50 µM of 3KGlcβOMe), where the concentration of each enzyme was 0.06 µM and the specified enzymes were replaced with their respective counterpart from the other series, one at a time. The replaced enzyme is specified for each reaction, for instance BT1AL23 contains BT1 plus AL2 and AL3, thus in the AL series AL1 is replaced by BT1. To these solutions was added 20 µM of the substrates (compound **1** for the BT enzymes and compound **3** for the AL enzymes) and the initial rates of the reaction are calculated based on change in fluorescence of the solution. Control denotes wells with no enzymes added.

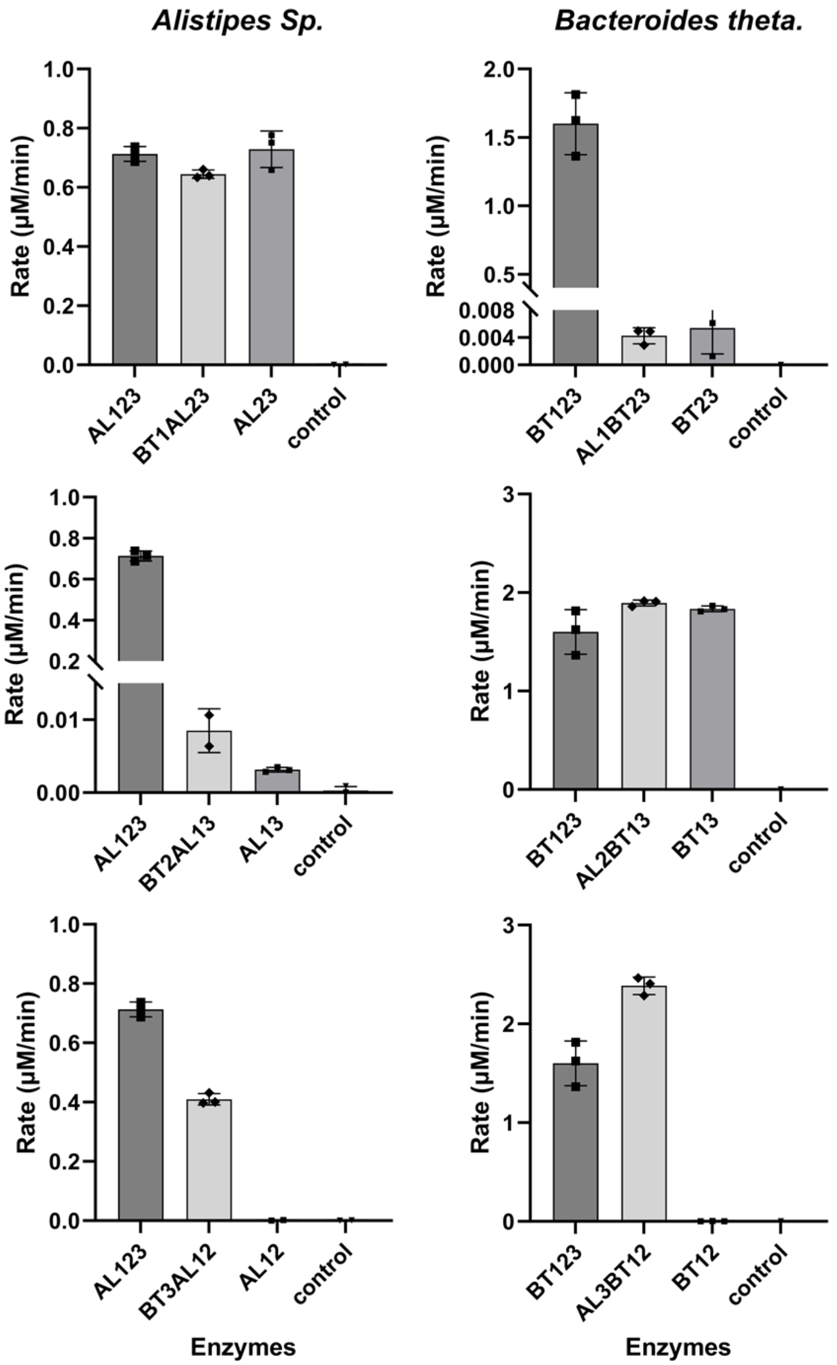

**Figure S15:** Swapping the enzymes, (bars indicate the mean, and error bars the SD of technical replicates (n=3)).

###### 2.5.2.5. Screening for buffer conditions and ionic strength

To solutions containing 0.01 μM each of the four enzymes of either series in various buffers (all at 50 mM and adjusted to pH 7.0) was added 50 μM of the substrates (compound **1** for the BT enzymes and compound **3** for the AL enzymes), and the reaction was followed by measuring the fluorescence of released JBOMe over time.

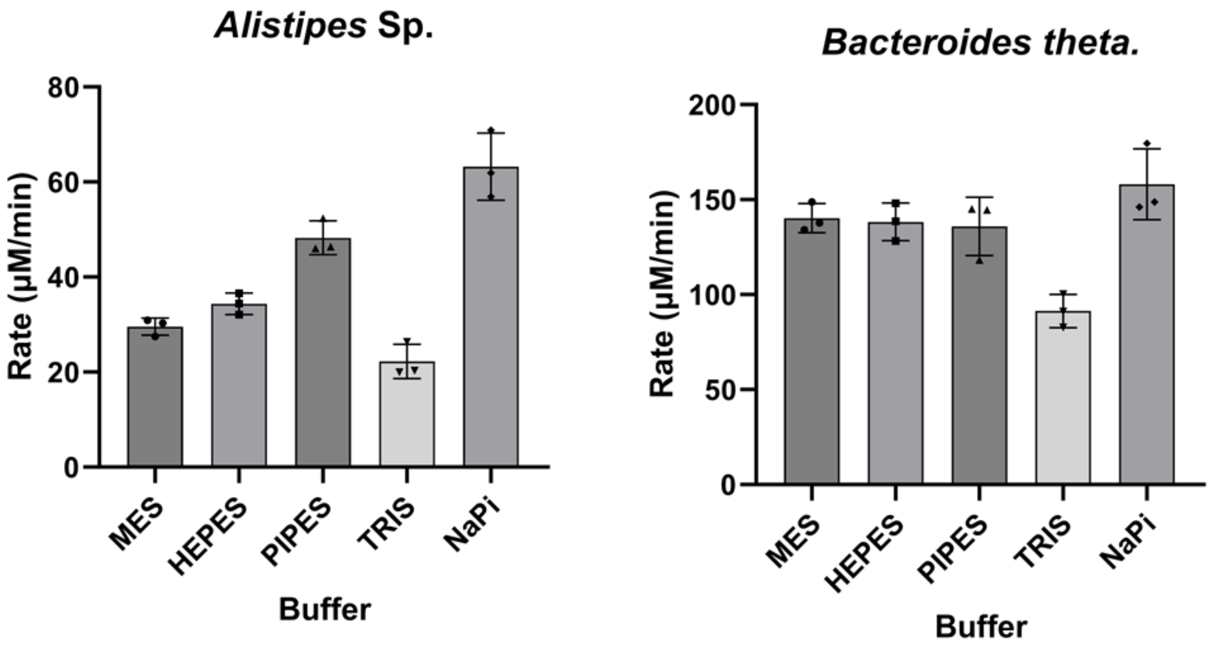

**Figure S16:** Buffer conditions, (bars indicate the mean, and error bars the SD of technical replicates (n=3)).

To solutions containing 0.005 μM of each of the four enzymes of either series in NaPi buffer at different concentrations and/or in the presence of additives, was added 50 μM of the substrates (compound **1** for the BT enzymes and compound **3** for the AL enzymes), and the initial rates were monitored, as shown below.

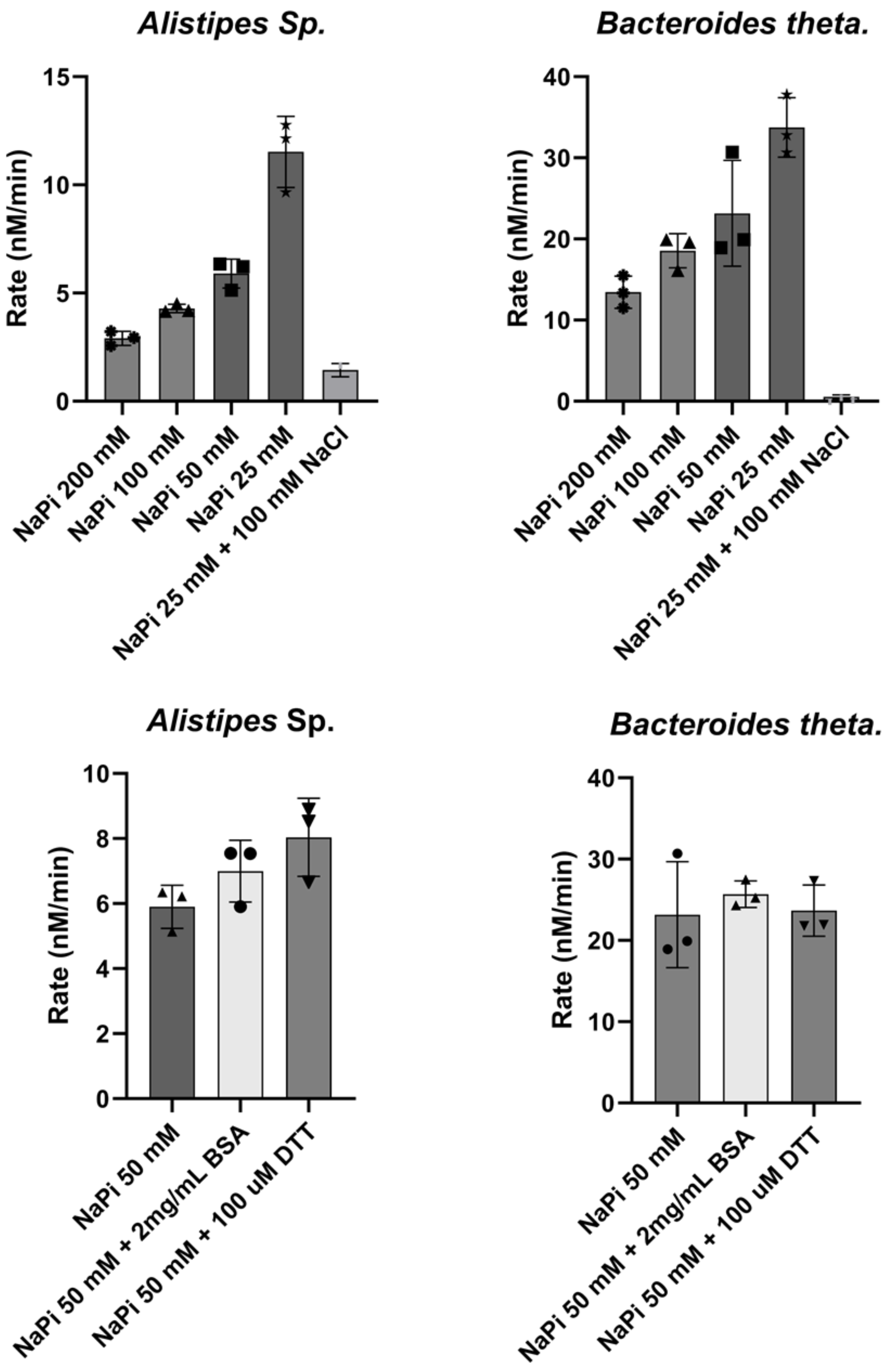

**Figure S17:** Buffer conditions – cont'd
(bars indicate the mean, and error bars the SD of technical replicates (n=3)).

###### 2.5.2.6. Screening for metal cofactors

To a solution containing 0.005 µM of each of the four enzymes of either series in 50 mM HEPES buffer pH 7.0 (Note: We used HEPES buffer, despite it not being the best buffer, since unlike TRIS or NaPi it is less inclined to chelate metal ions or cause them to precipitate[27]), 50 µM of 3KGlcβOMe, 50 µM of the specified metal ion or EDTA, was added 20 µM of the substrates (compound **1** for the BT enzymes and compound **3** for the AL enzymes), and the reaction was followed by measuring the fluorescence of released JBOMe over time.

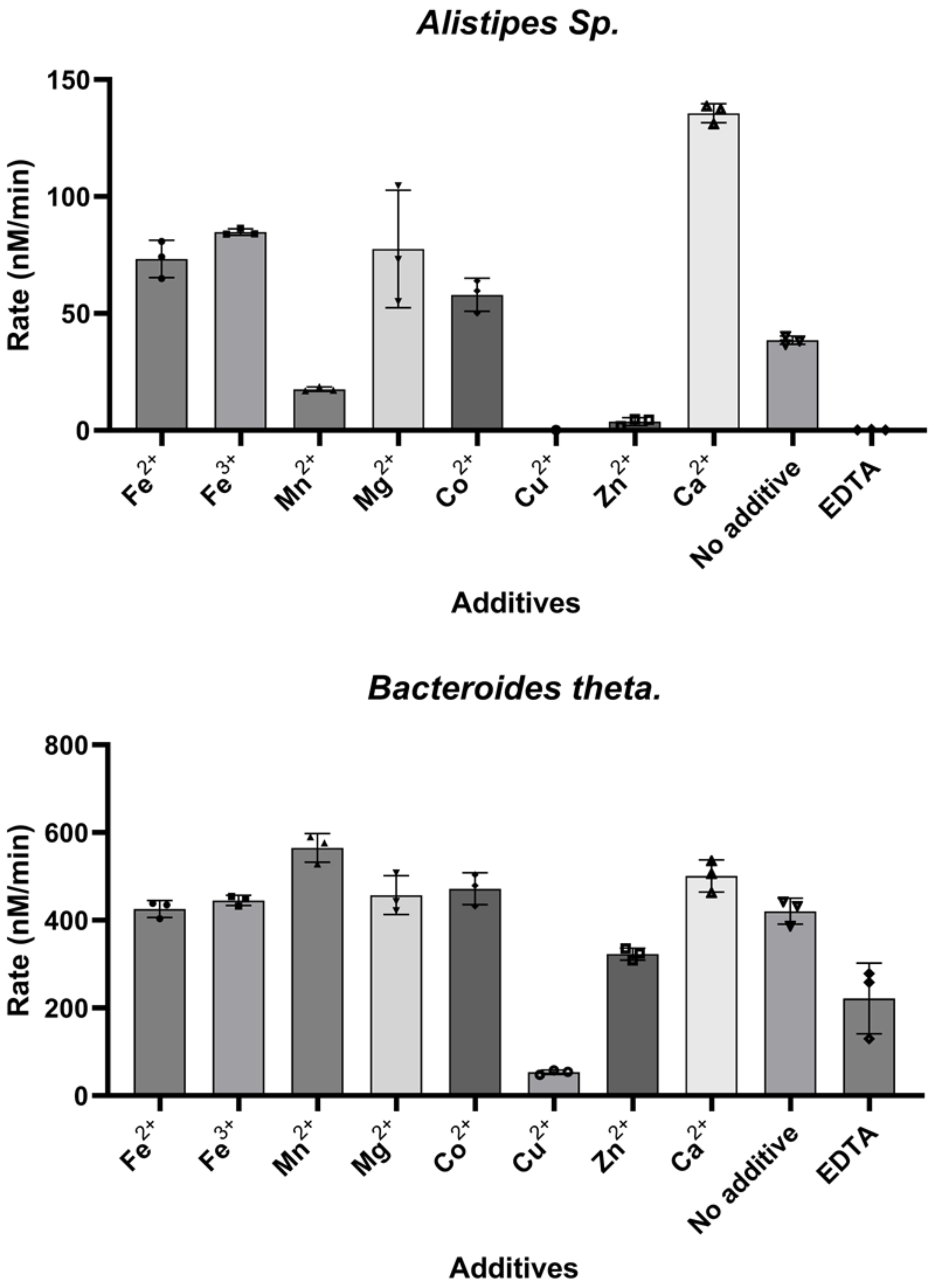

**Figure S18:** Screening for metal ions, (bars indicate the mean, and error bars the SD of technical replicates (n=3)).

To a solution containing 0.002 μM of each of the four enzymes of either series in the same buffer as above and 50 μM of each of the two specified metal ions or 100 μM EDTA, was added 20 μM of the substrates (compound **1** for the BT enzymes and compound **3** for the AL enzymes) and the reaction was followed by measuring the fluorescence of released JBOMe over time.

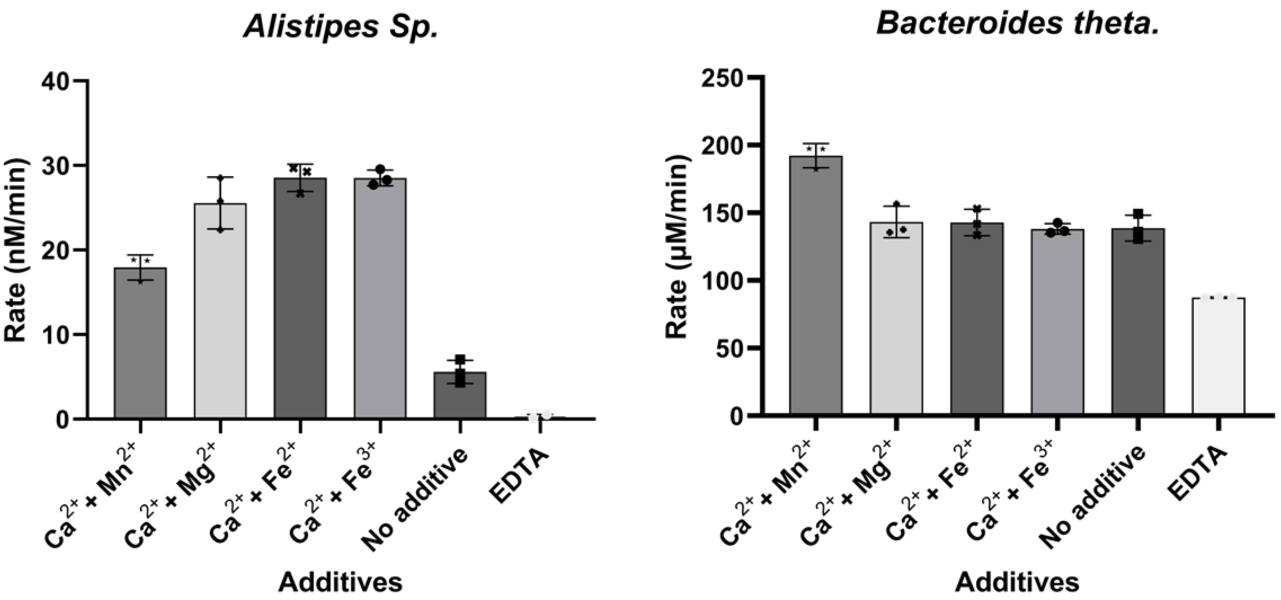

**Figure S19:** Screening for metal ions – cont'd, (bars indicate the mean, and error bars the SD of technical replicates (n=3)).

Next, we repeated these experiment in the 25 mM NaPi buffer to make sure this trend is the same in the best buffer found in the earlier assay.

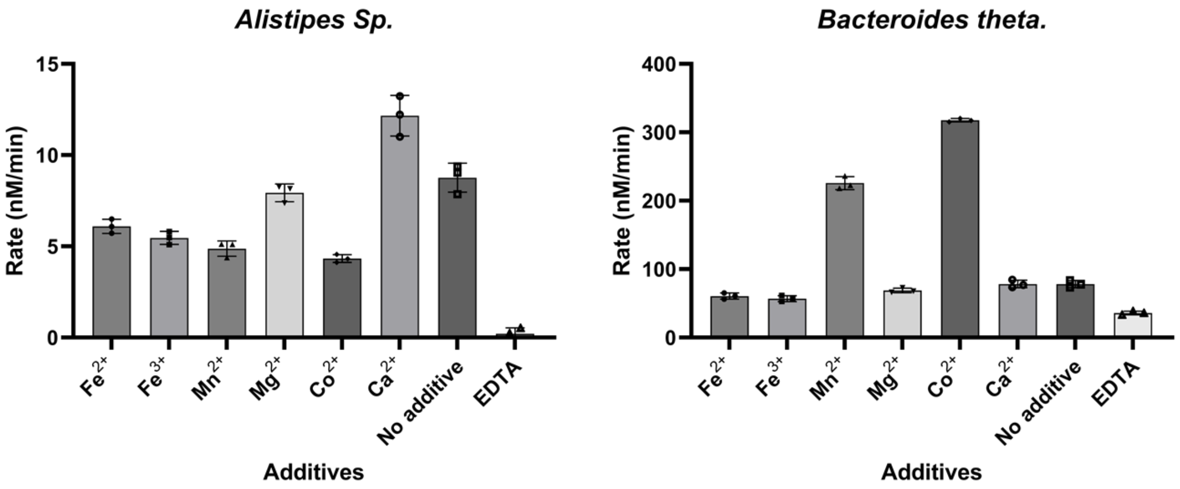

**Figure S20:** Screening for metal ions – cont'd, (bars indicate the mean, and error bars the SD of technical replicates (n=3)).

And then checked the concentration-dependent effects of the select few metal ions in the same conditions.

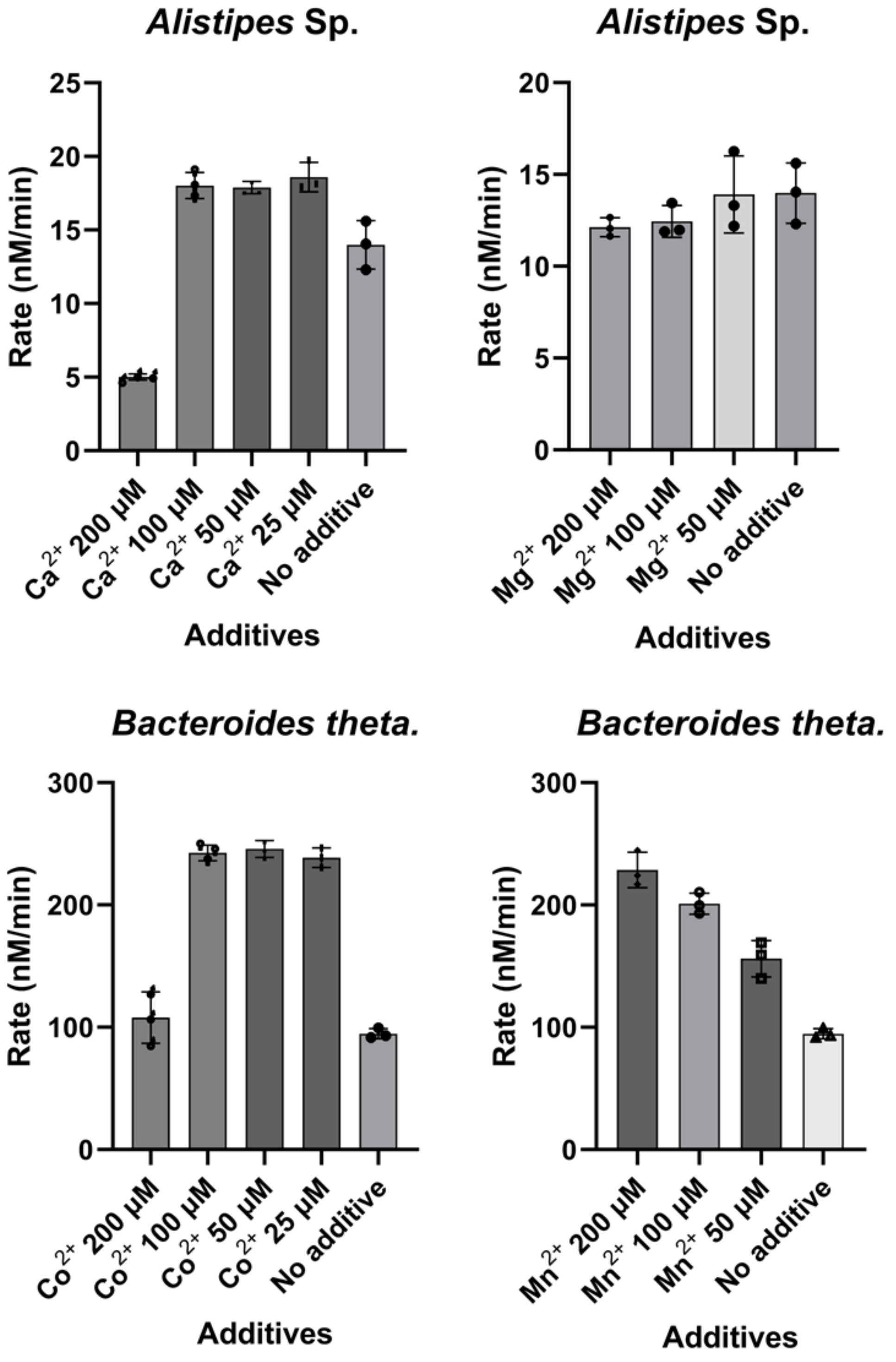

**Figure S21:** Screening for metal ions – cont'd, (bars indicate the mean, and error bars the SD of technical replicates (n=3)).
We therefore selected the following conditions for our future assays:
For BT enzymes: $CoCl_2$ 50 μM and $MnCl_2$ 200 μM.
For AL enzymes: $CaCl_2$ 25 μM.
Note that we later found more detailed information about the specific metal ions that each enzyme binds. (See section 2.5.3.)

###### 2.5.2.7. Screening for cofactors

To a solution containing 0.004 µM each of the first three enzymes of either series in NaPi buffer (25 mM at pH 7.0) and varying concentration of the specified cofactor was added 20 µM of the substrates (compound **1** for the BT enzymes and compound **3** for the AL enzymes), and the reaction was followed by measuring the fluorescence of released JBOMe over time. Initial rates were calculated for the first 5 minutes of the reactions.
The results show that none of the cofactors are able to increase the rate of the reaction.

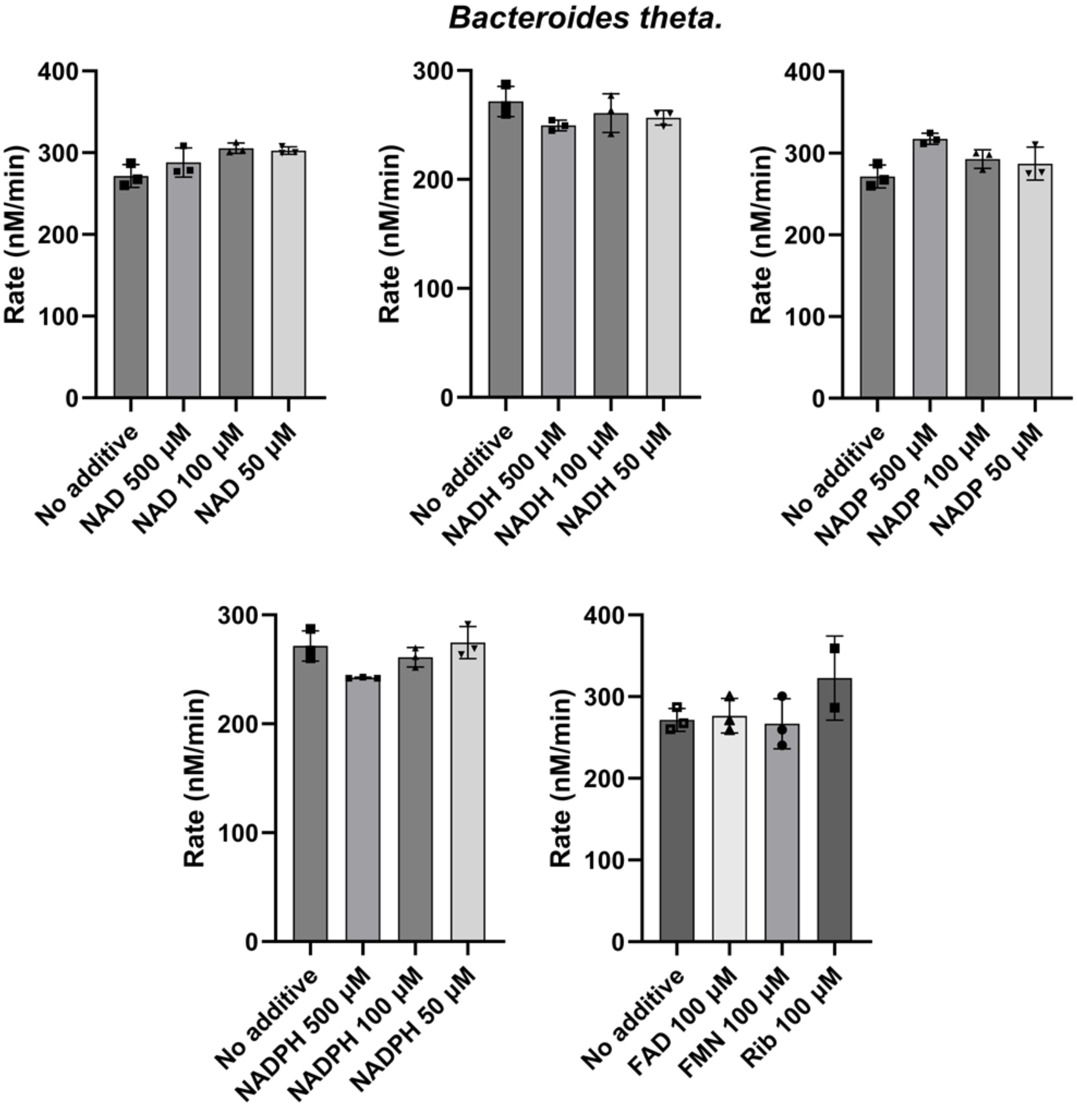

**Figure S22:** Screening for cofactors – BT enzymes, (bars indicate the mean, and error bars the SD of technical replicates (n=3)).

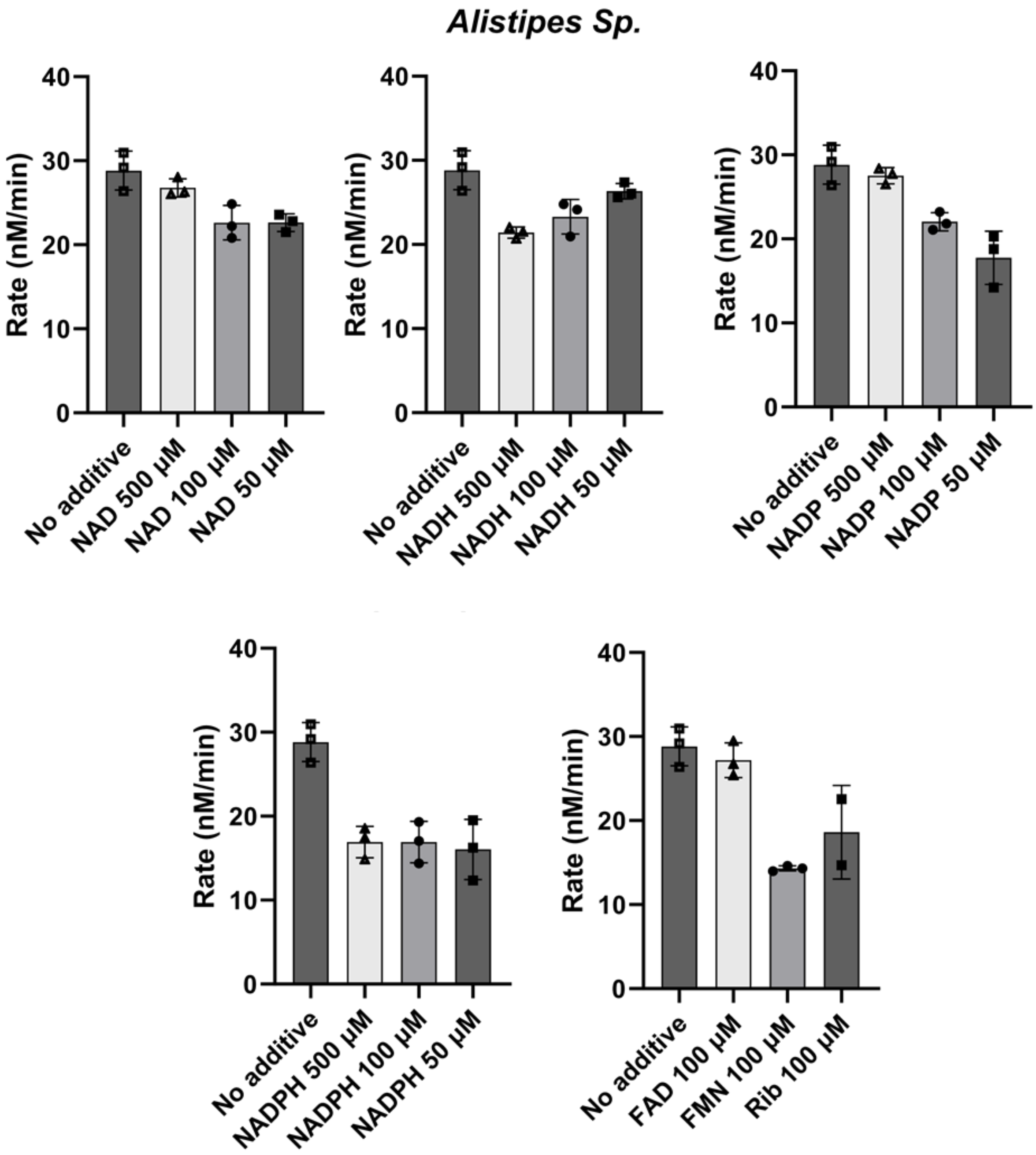

**Figure S23:** Screening for cofactors – AL enzymes, (bars indicate the mean, and error bars the SD of technical replicates (n=3)).

##### 2.5.3. Pseudo-Michaelis-Menten Kinetics

The reactions of varying concentrations of the substrates and the four enzymes (0.002 µM each for the BT enzymes and 0.001 µM each for AL enzymes – the full length AL1, AL-iso-FL was used for this experiment) in NaPi buffer (25 mM, pH 7.0), containing 50 µM of 3KGlcβOMe and the appropriate amounts of metals (for BT enzymes: 50 µM $CoCl_2$ and $MnCl_2$ 200 µM, for AL enzymes: 25 µM $CaCl_2$) were monitored by measuring the fluorescence of released fluorophore over time. The initial rates were then fit to a Michaelis-Menten model. Note that the reported values are all pseudo-Michaelis-Menten values, since a mixture of enzymes is used here.

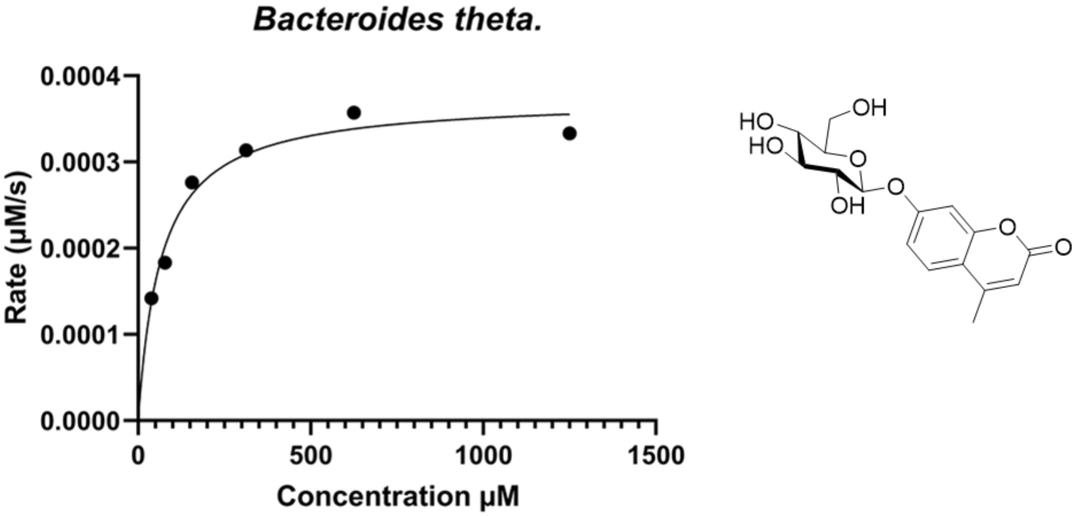

$K_m$ = 66 ± 12 µM

$k_{cat}$ = 0.19 ± 0.01 $s^{-1}$

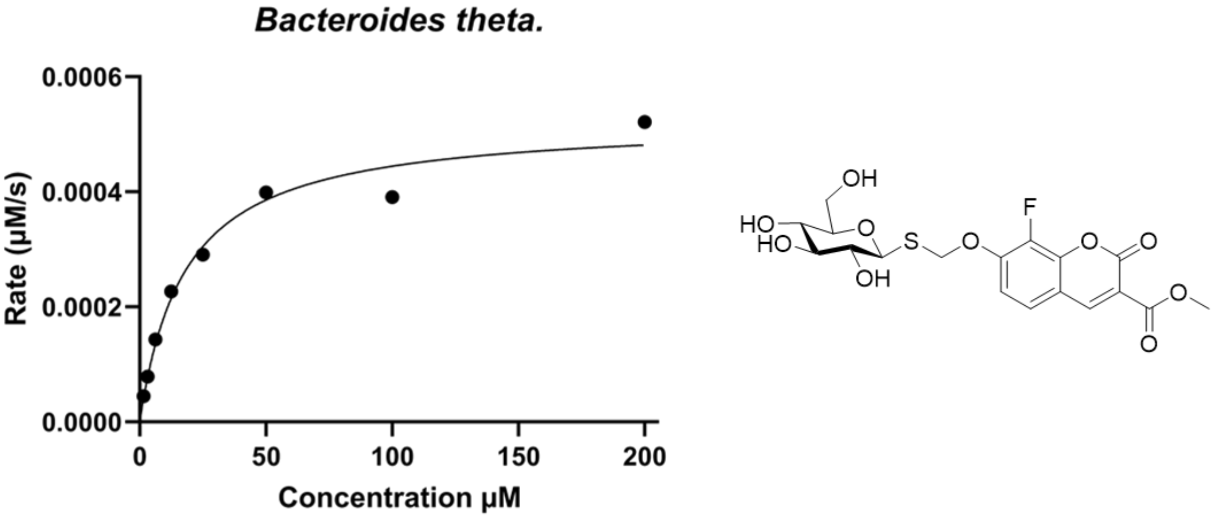

$K_m$ = 18 ± 3 µM

$k_{cat}$ = 0.26 ± 0.02 $s^{-1}$

**Figure S24:** Pseudo-Michaelis-Menten kinetics – BT enzymes

#### *Alistipes Sp.*

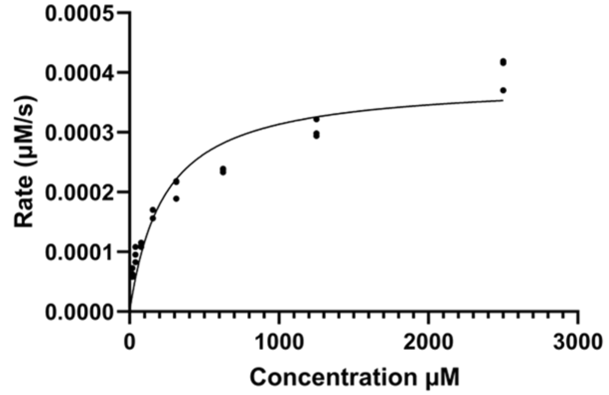

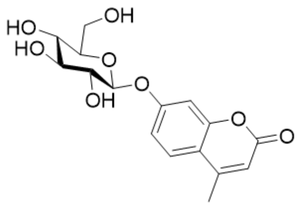

$K_m$ = 231 ± 42 µM
$k_{cat}$ = 0.39 ± 0.02 $s^{-1}$

#### *Alistipes Sp.*

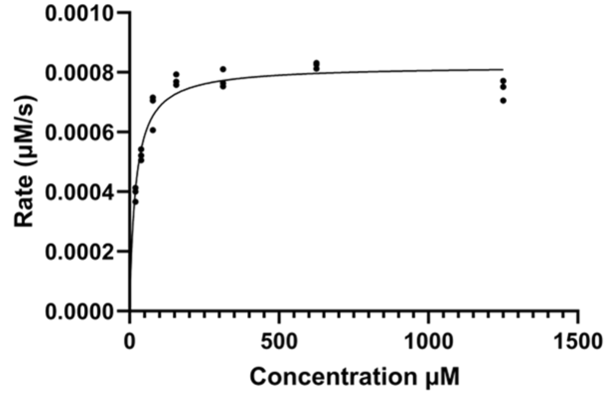

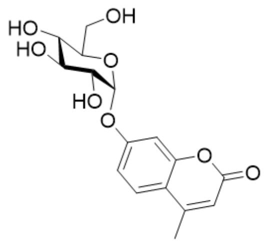

$K_m$ = 20 ± 2 µM
$k_{cat}$ = 0.82 ± 0.02 $s^{-1}$

#### *Alistipes Sp.*

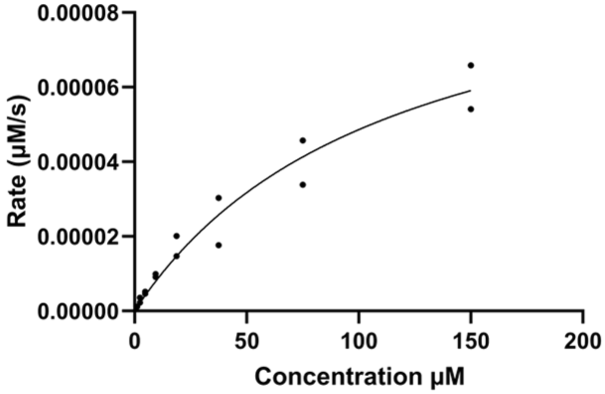

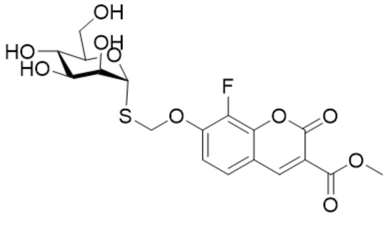

$K_m$ = 114 ± 33 µM
$k_{cat}$ = 0.10 ± 0.02 $s^{-1}$

**Figure S25:** Pseudo-Michaelis-Menten kinetics – AL enzymes, each data point represents that rate measured for a replicate (n=3).

##### 2.5.4. Kinetic assays for individual enzymes from BT and AL operons

Since enzymes AL/BT1 and AL/BT2 can catalyze the elimination reaction of 3-keto-glycosides, we were able to readily study their Michaelis-Menten kinetics with chromogenic and fluorogenic 3-keto-glycoside substrates. As discussed in the paper, we chemically synthesized these substrates (see section 1.8.) and checked the activity of the enzymes with them as follows.

###### 2.5.4.1. Testing the preference and activity of enzymes

3-Keto glycosides and especially those that have activated aglycones, such as our synthetic substrates, are not completely stable in the reaction buffer. They slowly decompose to yield the products of the elimination reaction, the free aglycone and the enone 3-keto-2-hydroxy-glucal (see section 2.7.1.). We therefore compared the rates of hydrolysis with and without addition of enzymes. Solutions of the fluorogenic substrates, 3KGlc-α and β-MU at 25 µM in 25 mM NaPi buffer, pH 7.0, were thus incubated with the enzymes, one at a time. The results are shown below. The concentration of all enzymes used for these assays was 40 nM.

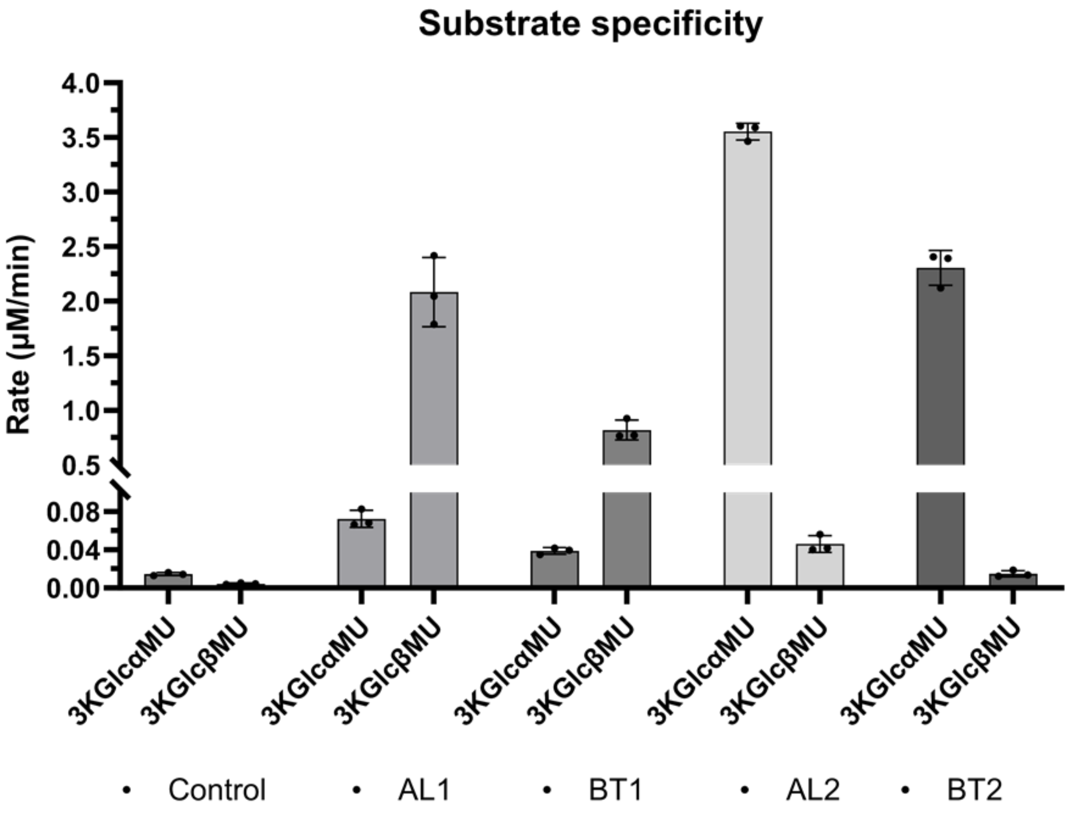

**Figure S26:** substrate specificity for individual enzymes, (bars indicate the mean, and error bars the SD of technical replicates (n=3)).

###### 2.5.4.2. Screening for metal cofactors

Next, using these substrates to monitor individual enzymes, we examined the effect of metal additives on the rates of the catalyzed reactions.

The rates of the reaction of 100 µM of chromogenic substrate 3KGlcαpNP catalyzed by 40 nM of AL2 or BT2 in 25 mM NaPi buffer pH 7.0 in the presence of 50 µM of the specified metals and 25 µM of EDTA were therefore monitored. The results show that AL2 and BT2 are both $Ca^{2+}$-dependent but are also activated by $Mg^{2+}$.

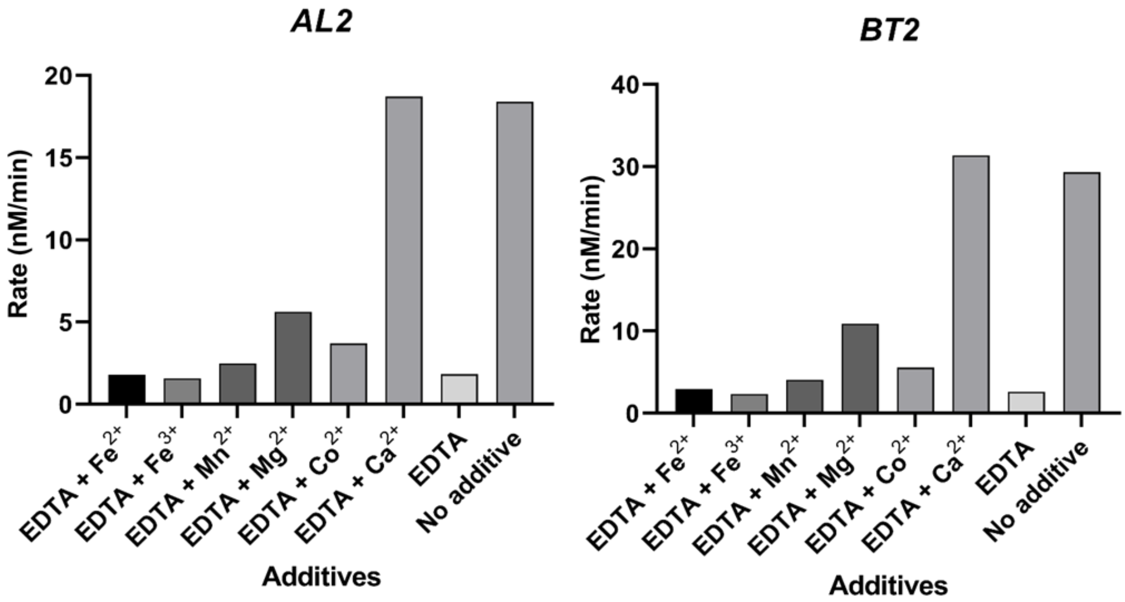

**Figure S27:** Screening for metal cofactors – AL2 and BT2

The above experiment was not conclusive for AL1 and BT1 in NaPi buffer and therefore was repeated in HEPES buffer, with 10 µM of 3KGlcβMU and 4 nM of each enzyme, 50 µM of the specified metals and 25 µM of EDTA where mentioned. The results suggest that these enzymes, and BT1 to a greater extent, are activated by both $Co^{2+}$ and $Mn^{2+}$. Next, for BT1, the dependence of the rate on the concentration of each metal was also determined to distinguish the most likely metal bound in

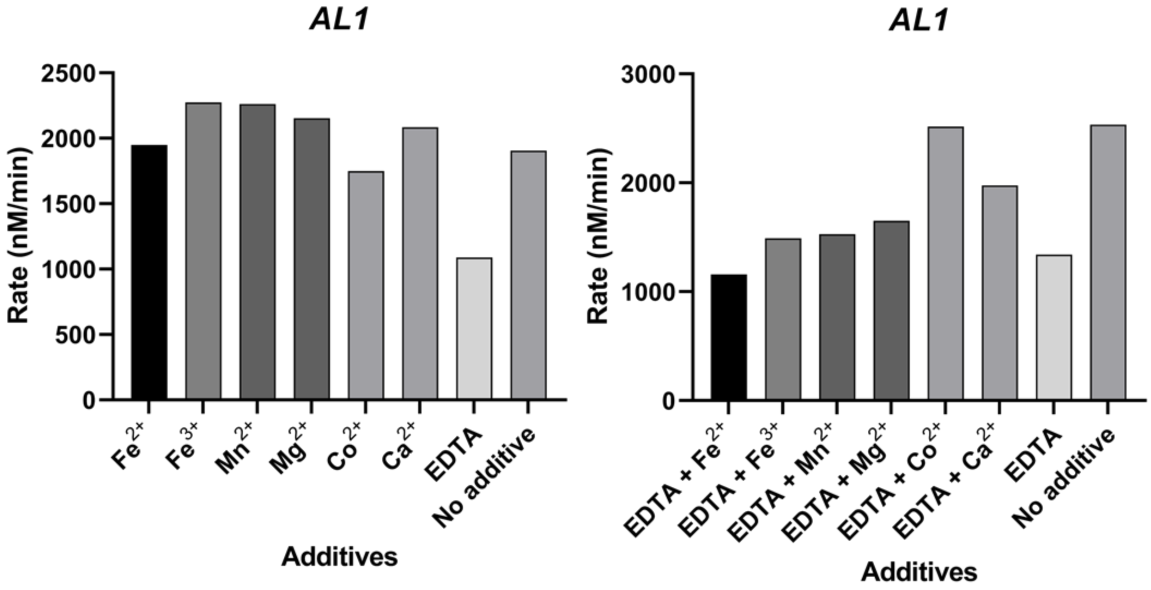

**Figure S28:** Screening for metal cofactors – AL1 and BT1

the active site under native conditions. BT1 binds more tightly to $Co^{2+}$ but reaches a higher rate enhancement upon addition of high concentrations of $Mn^{2+}$.

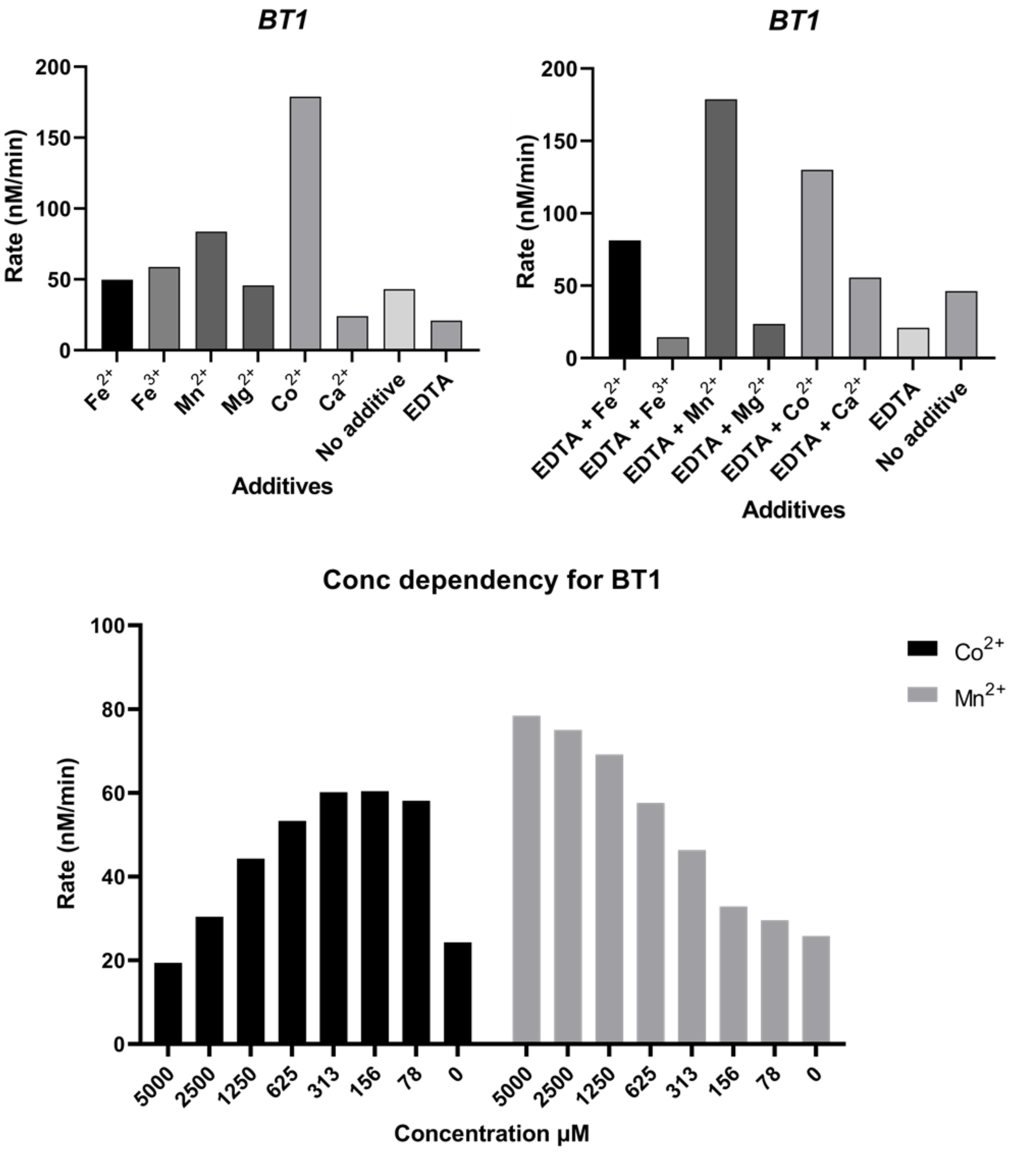

**Figure S29:** extended screening for metal cofactors – BT1

###### 2.5.4.3. Michaelis-Menten Kinetics for individual enzymes

Solutions of substrates at different concentrations were made in 25 mM NaPi buffer. Reactions were started by addition of a fixed amount of enzyme to the plates containing the above solutions. The data from the initial rates of the reactions were used to fit a Michaelis-Menten model as depicted below. Note that the observed kinetic parameters for BT1 and AL1 are most likely different from their actual values ($k_{cat}$ should be higher and $K_m$ lower than the observed values), since the 3-keto-β-glucoside substrates in solution are present as an equilibrium between the keto and hydrate forms (the portion of the keto-form in equilibrium is roughly half the total amount) and the equilibrium is not established instantly (see section 2.7.).

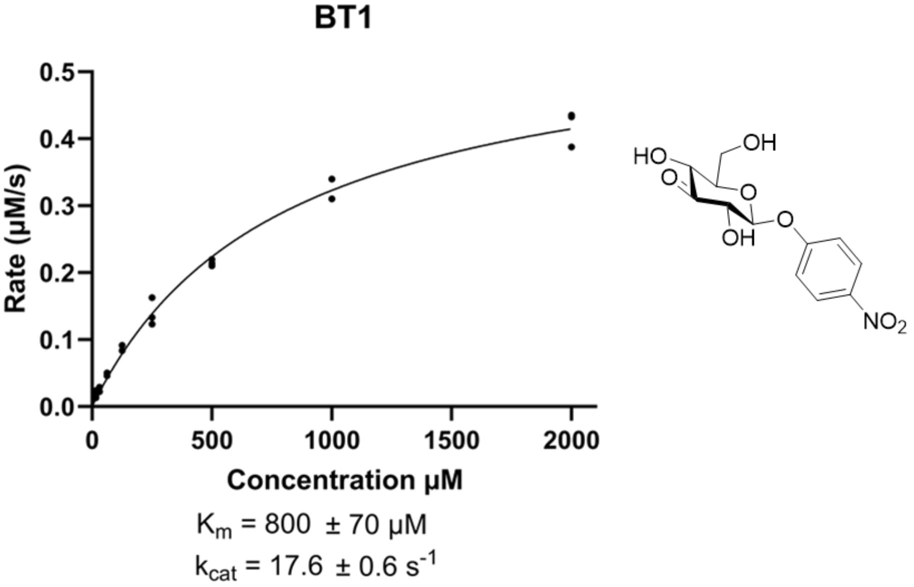

**Figure S30:** Michaelis-Menten kinetics for BT1, each data point represents that rate measured for a replicate (n=3).

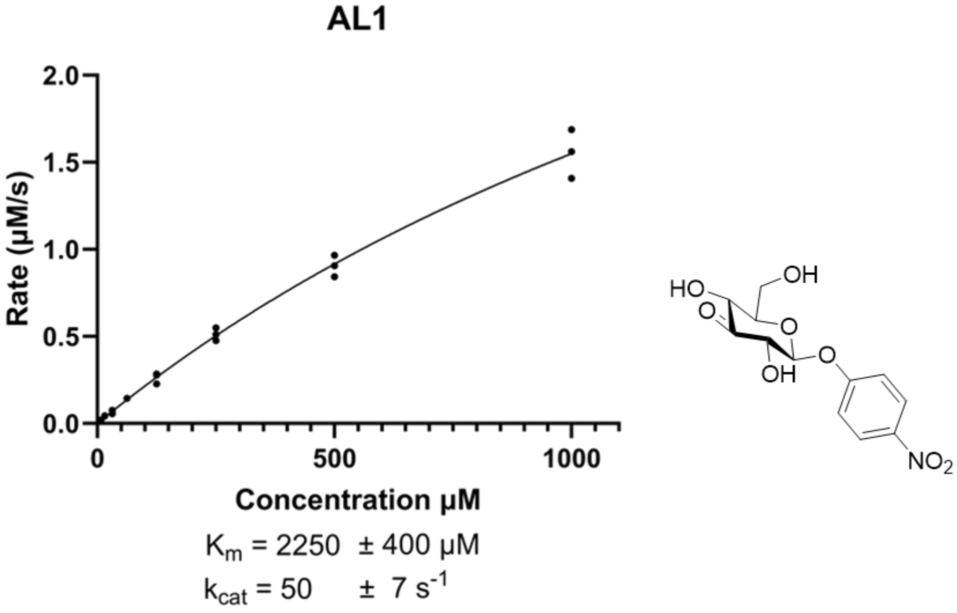

**Figure S31:** Michaelis-Menten kinetics for AL1, each data point represents that rate measured for a replicate (n=3).

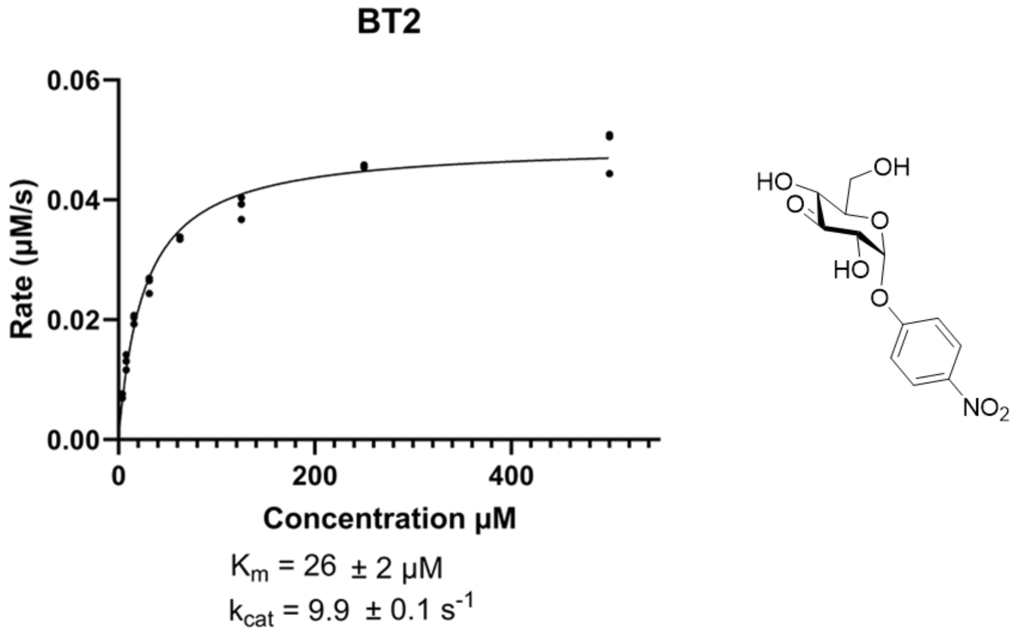

**Figure S32:** Michaelis-Menten kinetics for BT2, each data point represents that rate measured for a replicate (n=3).

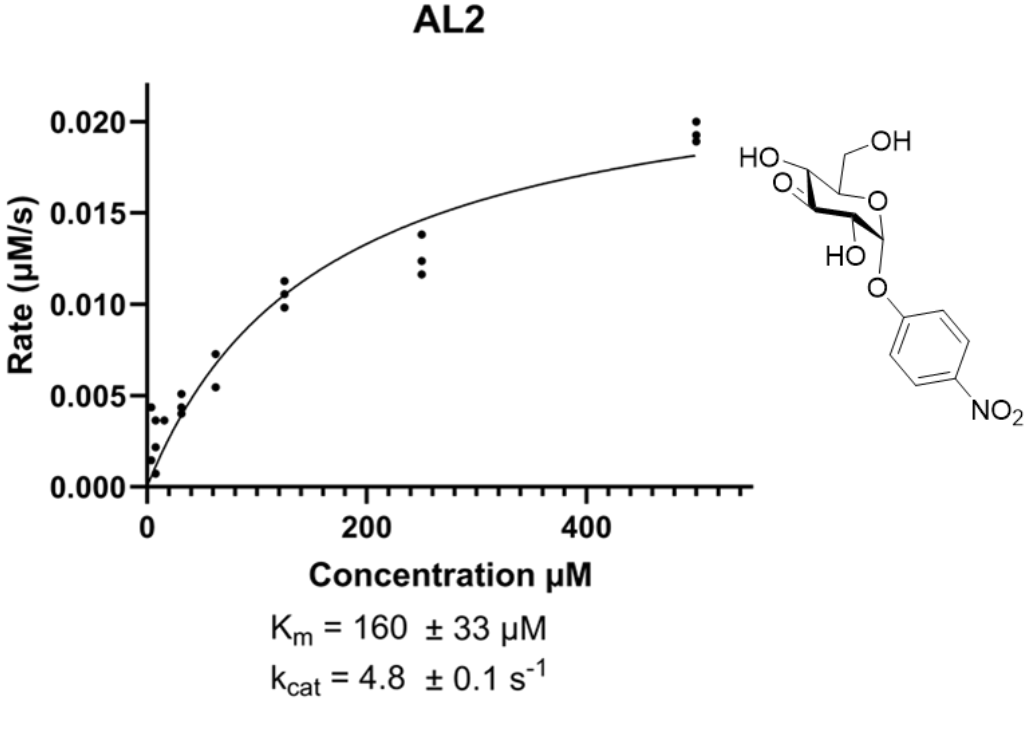

**Figure S33:** Michaelis-Menten kinetics for BT2, each data point represents that rate measured for a replicate (n=3).

##### 2.5.5. Assay for competitive Inhibitor/substrates

A set of compounds were first tested as competitive inhibitors of enzymes AL1, AL2, BT1 and BT2, as these enzymes can be assayed individually using the chemically synthesized 3-keto-glucoside substrates. In each assay, to solutions of varying concentrations of the possible inhibitors and 2 nM of the specified enzyme in NaPi buffer (25 mM at pH 7.0), was added the substrates (final concentration 10 µM, 3KGlcβMU for the AL1/BT1 and 3KGlcαMU for AL2/BT2) and the reaction was followed by measuring the fluorescence of released fluorophore over time. The $IC_{50}$ values are calculated based on the initial rates of the reactions and are reported in µM:

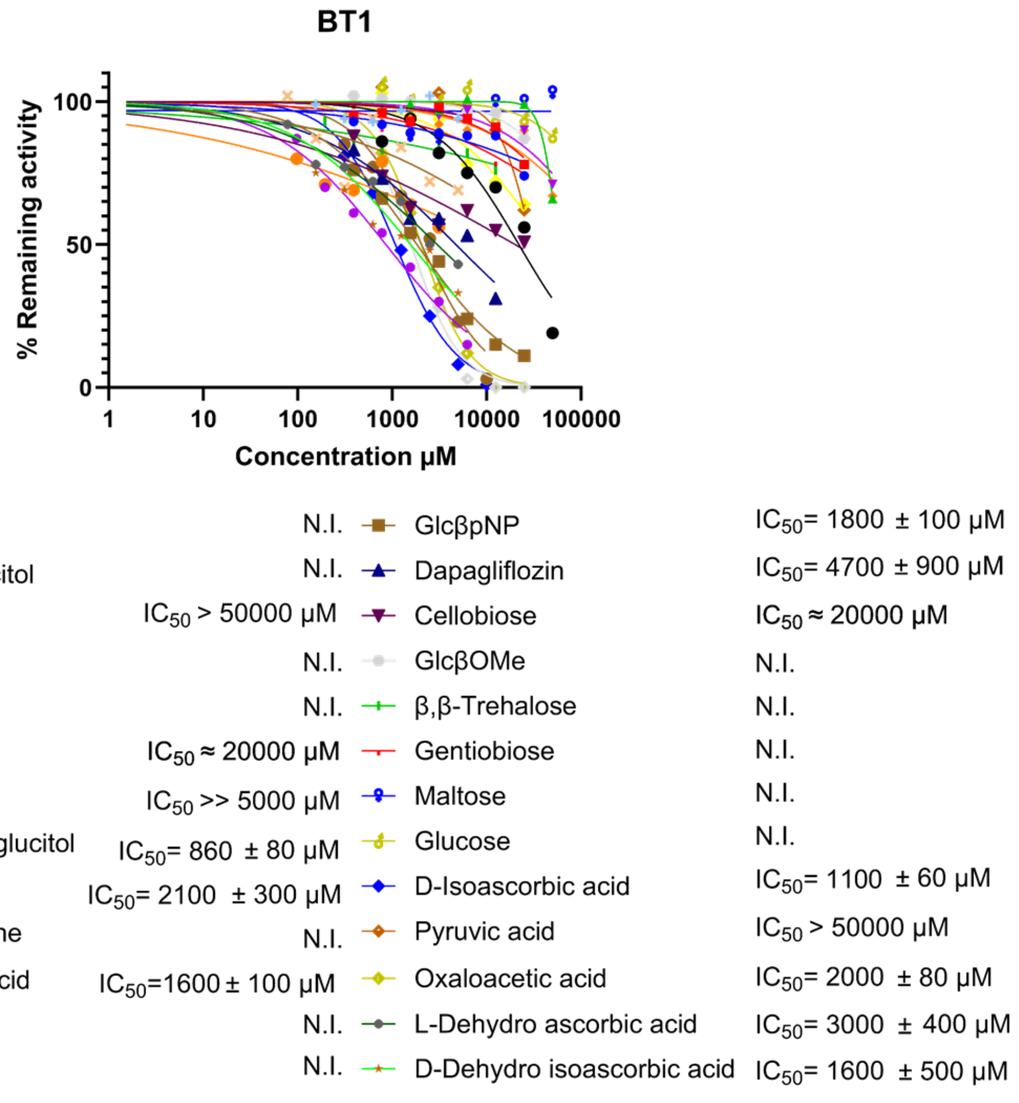

**Figure S34:** Inhibition assay for BT1

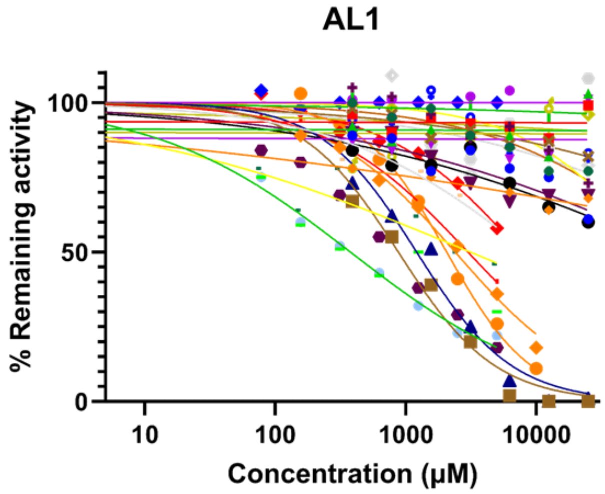

| | | | |
|---|---|---|---|
| D-Erythrose | $IC_{50}$ > 25000 µM | Oxaloacetic acid | $IC_{50}$= 880 ± 80 µM |
| D-Arabinose | N.I. | α-Ketoglutaric acid | $IC_{50}$= 1200 ± 130 µM |
| D-Glucose | N.I. | Dihydroxyacetone | $IC_{50}$ > 25000 µM |
| D-Fructose | N.I. | Hydroxyacetone | N.I. |
| DL-Glyceraldehyde | $IC_{50}$ > 25000 µM | Acetaldehyde | N.I. |
| Pyruvic acid | $IC_{50}$ > 25000 µM | Acetone | N.I. |
| Glucal | N.I. | GlcβpNP | $IC_{50}$ >> 10000 µM |
| Sinigrin | $IC_{50}$ > 50000 µM | Maltose | N.I. |
| α,α-Trehalsoe | N.I. | Cellobiose | N.I. |
| Dapagliflozin | N.I. | L-Ascorbic acid | $IC_{50}$= 2400 ± 150 µM |
| 3K-1,5-anhydroglucitol | $IC_{50}$= 3000 ± 200 µM | D-Isoascorbic acid | $IC_{50}$= 2100 ± 100 µM |
| 1,5-anhydroglucitol | N.I. | 1,4-anhydroglucitol | N.I. |
| D-Threose | $IC_{50}$= 8800 ± 2600 µM | L-Dehydro ascorbic acid | $IC_{50}$= 610 ± 160 µM |
| D-Ribulose | $IC_{50}$= 3100 ± 1500 µM | D-Dehydro isoascorbic acid | $IC_{50}$= 380 ± 30 µM |

**Figure S35:** Inhibition assay for AL1

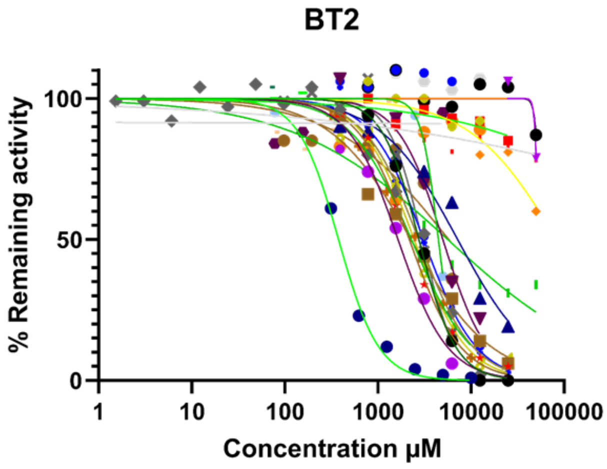

| | | | |
|---|---|---|---|
| Glucal | N.I. | GlcαpNP | $IC_{50}$= 2300 ± 200 μM |
| 1,5-anhydroglucitol | N.I. | Dapagliflozin | $IC_{50}$= 7200 ± 900 μM |
| Sinigrin | N.I. | Cellobiose | $IC_{50}$= 5000 ± 400 μM |
| Sucrose | N.I. | GlcβOMe | N.I. |
| α,α-Trehalsoe | N.I. | 3K-1,5-anhydroglucitol | $IC_{50}$= 2400 ± 200 μM |
| Glc-1-F | N.I. | GlcβpNP | $IC_{50}$= 3100 ± 200 μM |
| Glucose | N.I. | β,β-Trehalose | N.I. |
| Maltose | $IC_{50}$= 5600 ± 1500 μM | 3K-Glc-β-OMe | $IC_{50}$ >5000 μM |
| Gentiobiose | N.I. | Dihydroxyacetone | N.I. |
| L-Ascorbic acid | $IC_{50}$= 2700 ± 200 μM | D-Isoascorbic acid | $IC_{50}$= 3800 ± 20 μM |
| Oxaloacetic acid | $IC_{50}$= 2800 ± 100 μM | α-Ketoglutaric acid | $IC_{50}$= 1600 ± 100 μM |
| Pyruvic acid | $IC_{50}$= 2200 ± 100 μM | L-Dehydro ascorbic acid | N.I. |
| D-Ribulose | N.I. | D-Dehydro isoascorbic acid | $IC_{50}$= 4500 ± 400 μM |
| | | D-Threose | N.I. |

**Figure S36:** Inhibition assay for BT2

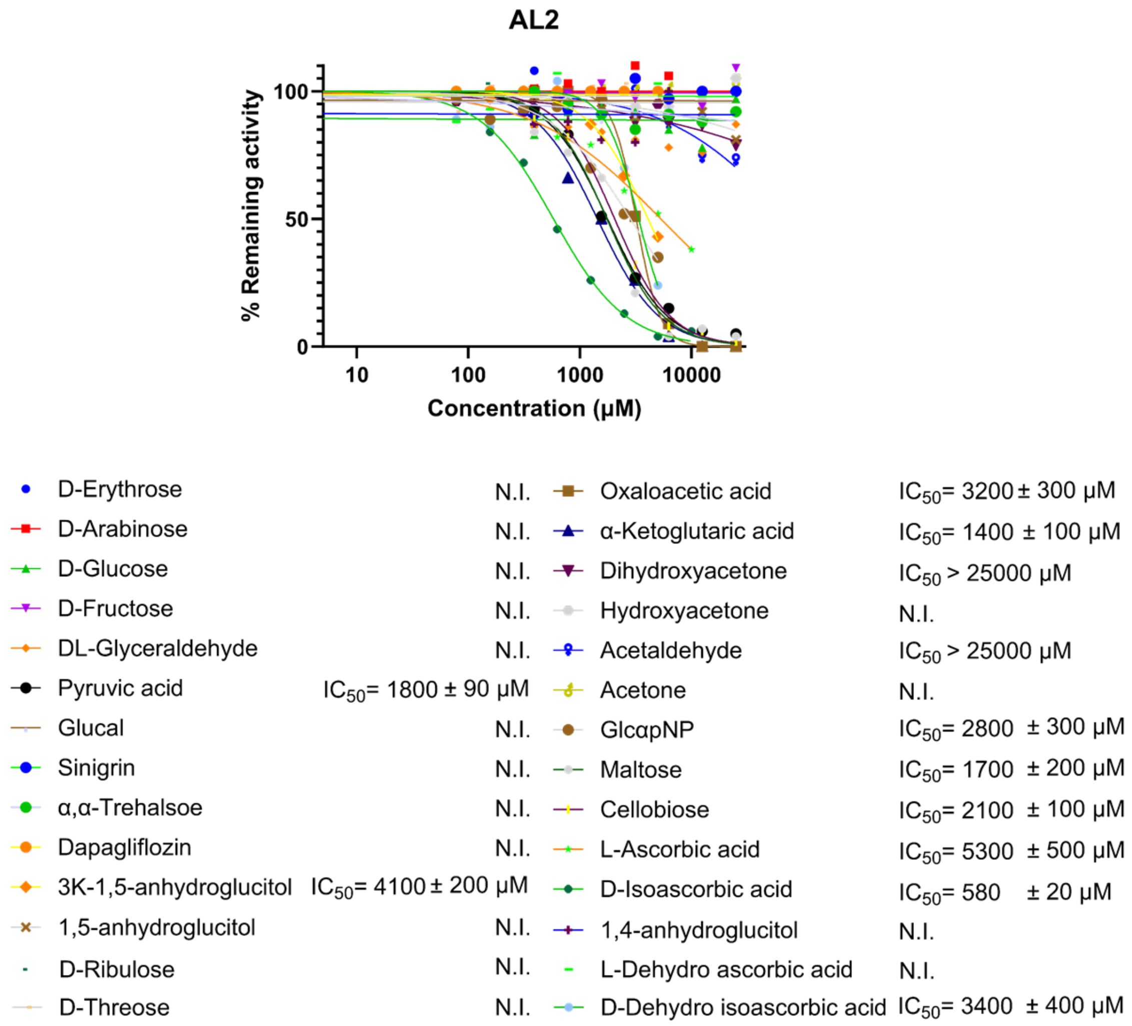

**Figure S37:** Inhibition assay for AL2

Next, we tested a more diverse set of compounds for their effect on the rates of the reactions catalyzed by the mixture of all the enzymes from either series. To pre-incubated solutions of varying concentrations of a set of possible inhibitors/competitive substrates and 2 nM each of the four enzymes of either series in NaPi buffer (25 mM at pH 7.0), containing 50 μM of 3KGlcβOMe and the appropriate amounts of metals (for BT enzymes: 50 μM $CoCl_2$ and $MnCl_2$ 200 μM, for AL enzymes 25 μM $CaCl_2$) was added the substrates (final concentration 5 μM, compound **1** (GlcSCJBOMe) for the BT enzymes and compound **3** (ManSCJBOMe) for the AL enzymes) and the reaction was followed by measuring the fluorescence of released JBOMe over time. This experiment was done for the BT enzymes first and the discussion below is based on those data. The data for AL enzymes is also included below and follows the same trends as that of BT enzymes.

Note that we have four enzymes in the mixture and therefore any observed inhibition can be caused by the effect of the compounds on any one (or multiple) of these four enzymes (BT4 is not necessary for observing the activity however and is not likely to be the reason for observed inhibition). More importantly, note that the presence of a 3-keto substrate is required to observe the reaction of these enzymes with the fluorogenic substrate. A likely reason for observing slower rates of reaction (e.g. inhibition) here is therefore that the compounds might be substrates that are oxidized by AL3/BT3 first and thus deplete the supply of 3KGlcβOMe. In this case, rather than being competitive inhibitors, these compounds would be substrates for the enzymes. In addition, it is also possible that the oxidized versions of these compounds could be inhibitors of downstream enzymes BT1 and BT2.

Observed $IC_{50}$ values for a set of monosaccharides were determined based on the initial rates of the reactions are reported here in µM (Fig S38).

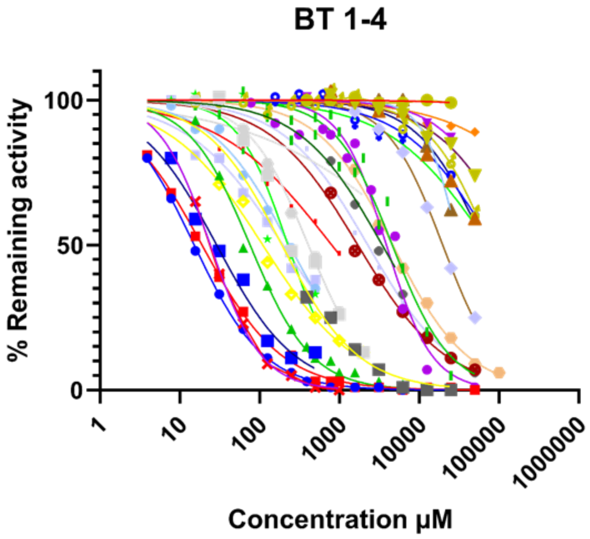

| | | | |
|---|---|---|---|
| Glucose | $IC_{50}$= 19.0 ± 0.5 µM | Levoglucosan | N.I. |
| Glucal | $IC_{50}$= 15.2 ± 0.2 µM | 2-H-Glucose | $IC_{50}$= 29 ± 3 µM |
| 1,5-Anhydrolglucitol | $IC_{50}$= 76 ± 4 µM | 2-F-Glucose | $IC_{50}$= 2300 ± 100 µM |
| Mannose | $IC_{50}$= 410 ± 23 µM | Glucosamine | $IC_{50}$= 110 ± 8 µM |
| Galactose | N.I. | N-Ac-Glucosamine | $IC_{50}$= 1700 ± 100 µM |
| Allose | N.I. | 4-F-Glucose | N.I. |
| Quinovose | $IC_{50}$= 800 ± 150 µM | Myo-Inositol | N.I. |
| 6-$PO_4$-Glucose | $IC_{50}$ = 4606 ± 450 µM | Scyllo-Inositol | N.I. |
| Glucuronic acid | $IC_{50}$ = 3900 ± 300 µM | Glc-α-F | $IC_{50}$= 200 ± 30 µM |
| Fructose | $IC_{50}$= 2900 ±40 µM | Glc-α-OMe | $IC_{50}$= 240 ± 40 µM |
| Xylose | $IC_{50}$= 4700 ± 200 µM | Glc-β-OMe | $IC_{50}$= 230 ± 15 µM |
| Sorbitol | $IC_{50}$= 19700 ± 500 µM | L-Ascorbic acid | $IC_{50}$= 190 ± 30 µM |
| D-Arabinose | N.I. | D-Isoascorbic acid | $IC_{50}$= 24.2 ± 0.5 µM |
| L-Arabinose | N.I. | | |

**Figure S38:** Inhibition assay for BT1-4 --monosaccharides

Since most of these compounds are not inhibitors of BT1 or BT2 (See figs S36 and S34, with the exception of L-ascorbic acid and D-isoascorbic acid), their observed inhibition is likely the result of their interaction with BT3, be it simple binding in the active site or being oxidized by BT3 and depleting the supply of the 3-keto co-substrate. The above data therefore indicate that the crucial interactions in the BT3 active site are made with the hydroxyl groups at the 3 and 4 positions, as galactose and allose are the only compounds showing no inhibitory activity. Binding at the 6-OH is important but not crucial (quinovose is an inhibitor, and so are glucuronic acid and glucose-6-phosphate, though with marked decrease in potency in the case of the latter compounds with bulky and charged substitutions at C-6). This is also the case for the hydroxyl at position 2, with 2-deoxy-glucose, mannose, glucosamine and N-acetylglucosamine all serving as inhibitors, though again the bulkier substitution of GlcNAc results in lower observed $IC_{50}$. Interestingly, 2-deoxy-2-fluoro-glucose also functions as an inhibitor but with a markedly lower potency. We attribute this to the destabilization of the oxidation transition state as a result of the presence of the electronegative fluorine group (for further discussion, see section 2.6.3.). It is also evident that the cyclic pyranose form is

what binds the best in the active site of BT3, since sorbitol, fructose and D-arabinose are all poor inhibitors compared with glucose.
We next tested a set of larger glycoside/saccharides in the same manner, with $IC_{50}$ values reported in µM:

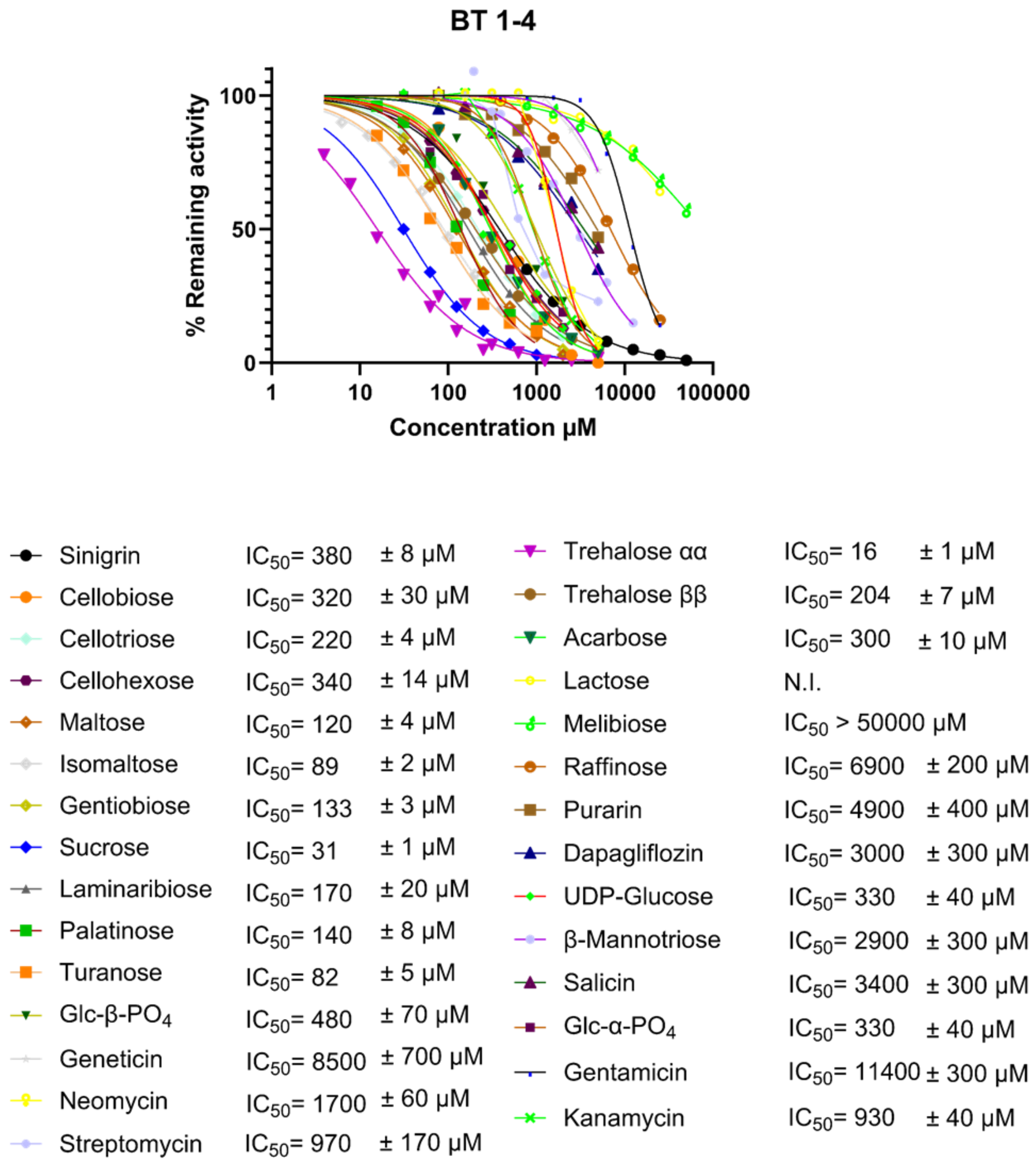

**Figure S39:** Inhibition assay for BT1-4 – glycosides/larger saccharides

Again, the most striking observation is that BT3 is very forgiving in terms of substrate preferences. Substitutions at the anomeric position are accepted regardless of their anomeric stereochemistry (Glc-α and β-OMe), charge (Glc-α and β-$PO_4$) and interestingly their size (Glc-α-F and UDP-Glucose). All of the disaccharides tested are inhibitory to the enzyme, regardless of the linkage position and type (See the values for maltose (Glc α(1→4) Glc), isomaltose (Glc α(1→6)Glc), gentiobiose (Glc β(1→6)Glc), laminaribiose (Glc β(1→3)Glc), sucrose (Glc α(1→2)βFruc),, cellobiose Glc β(1→4)Glc), trehalose αα (Glc α(1→1)αGlc), trehalose ββ (Glc β(1→1)βGlc), palatinose (also known as isomaltulose Glc α(1→6)Fruc), turanose (Glc α(1→3)Fruc)). Also of note, is that the binding pocket of BT3 must be open for binding to longer polymers as well, since cellotriose and cellohexaose bind with similar potency to cellobiose. The only disaccharides not acting as inhibitors are lactose (Gal β(1→4)Glc) and melibiose (Gal α(1→6)Glc), showing that the active site does not accommodate large substitutions on the glucose moiety nor is galactose an acceptable substrate, as already shown.

The results of the assays on AL enzymes are similar:

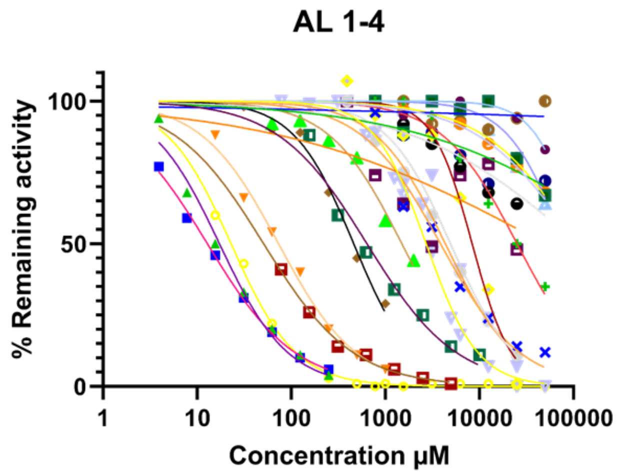

| | | | | | |
|---|---|---|---|---|---|
| Glucose | $IC_{50}$= 12.9 | ± 0.4 µM | D-Arabinose | $IC_{50}$> 50000 µM | |
| Glucal | $IC_{50}$= 17 | ± 8 µM | 1,4-Anhydrolglucitol | $IC_{50}$> 50000 µM | |
| 1,5-Anhydrolglucitol | $IC_{50}$ = 73 | ± 5 µM | Levoglucosan | N.I. | |
| Mannose | $IC_{50}$= 460 | ± 60 µM | Myo-Inositol | $IC_{50}$> 50000 µM | |
| Galactose | N.I. | | Scyllo-Inositol | $IC_{50}$ ≈ 25000 µM | |
| Allose | N.I. | | Glucosamine | $IC_{50}$= 51 | ± 3 µM |
| Quinovose | $IC_{50}$= 1500 | ± 330 µM | N-Acetyl-Glucosamine | $IC_{50}$= 640 | ± 70 µM |
| 6-$PO_4$-Glcucose | $IC_{50}$= 4930 | ± 620 µM | Erythritol | $IC_{50}$> 50000 µM | |
| Glucuronic acid | $IC_{50}$= 8380 | ± 750 µM | L-Ascorbic acid | $IC_{50}$= 2720 | ± 100 µM |
| Xylose | $IC_{50}$= 3860 | ± 530 µM | D-Isoascorbic acid | $IC_{50}$= 23.0 | ± 0.6 µM |
| Sorbitol | $IC_{50}$≈ 25000 µM | | | | |
| Fructose | $IC_{50}$= 4350 | ± 460 µM | | | |

**Figure S40:** Inhibition assay for AL1-4 --monosaccharides

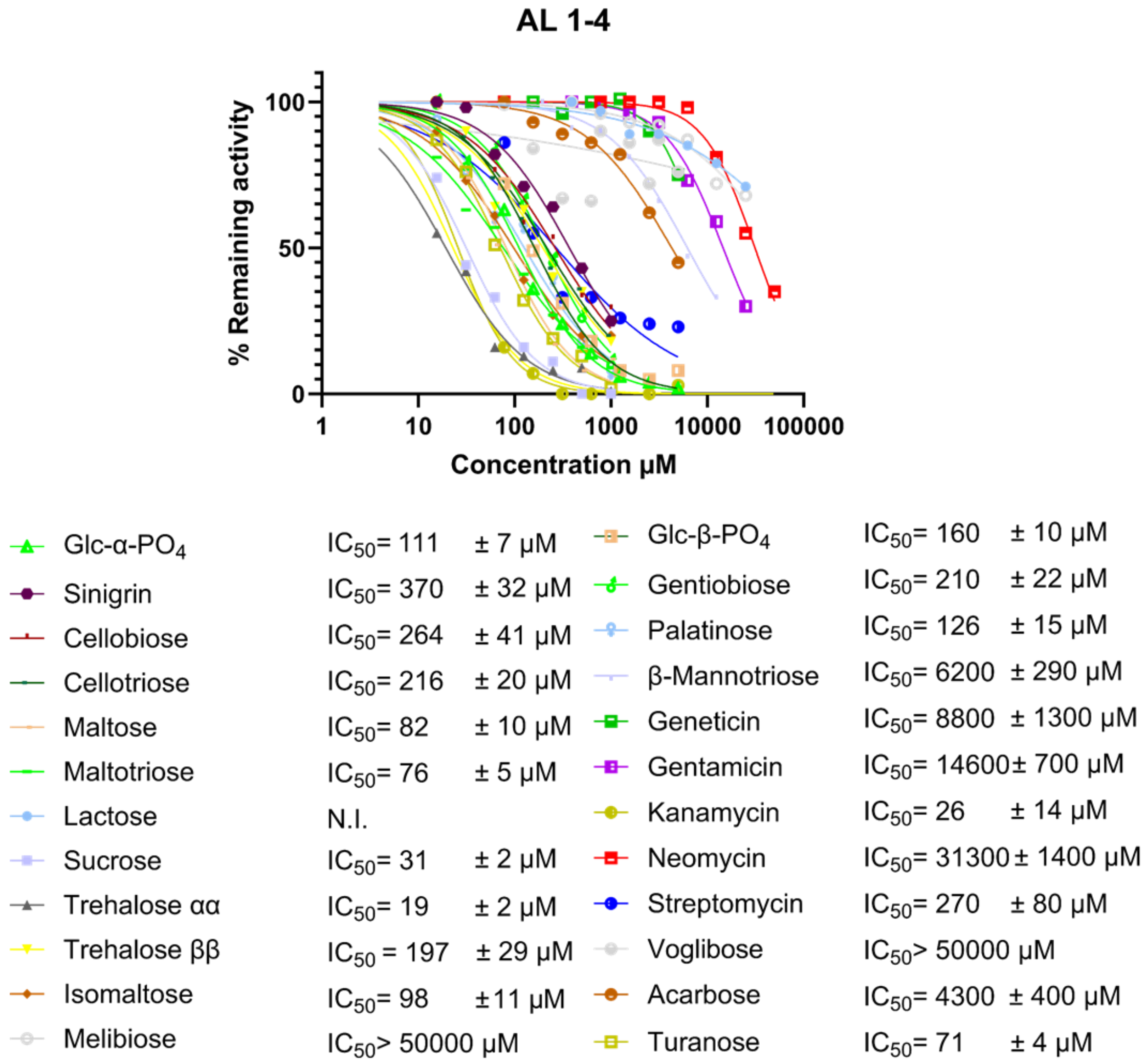

**Figure S41:** Inhibition assay for AL1-4 – glycosides/larger saccharides

Note that with these assays, we are only observing the inhibition caused by these disaccharides as they compete with our fluorogenic compound. We therefore used chromatography and NMR spectroscopy to see whether these disaccharides are indeed oxidized by AL/BT3 and later hydrolysed by the other enzymes, as is detailed in sections 2.6. and 2.7. These results showed that this 4-enzyme system has likely evolved to confer the ability of metabolizing a wide range of substrates to the bacteria that harbor them, rather than having one specific substrate.

##### 2.5.6. Assays for P2B11 enzymes

Similar to the assays described above, we ran assays to characterize the four enzymes from the P2B11 fosmid. In general, the characteristics of these enzymes are very similar to those described above, even though they are only distantly similar at the protein sequence level.

###### 2.5.6.1. Screening for co-factors

To solutions containing 0.7 μM each of the enzymes Iso1 and Iso2 and 2.8 μM of Oxi1 in HEPES buffers (50 mM at pH 7.0) and 50 μM of the specified cofactor was added 50 μM of the substrates (compound **4**) and the reaction was followed by measuring the fluorescence of released JBOMe over time. The results show that, as seen with the above systems, addition of the cofactors does not result in any rate enhancement.

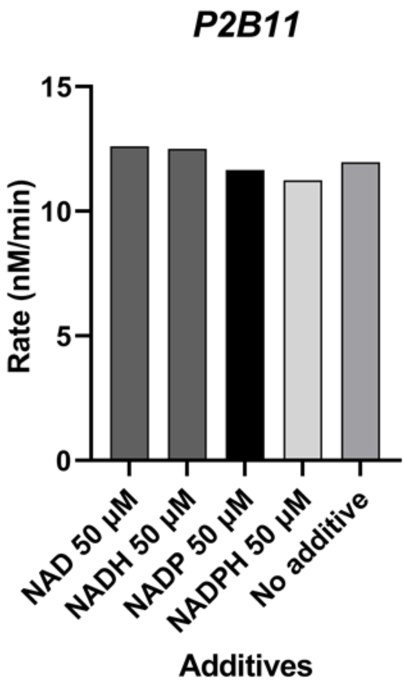

**Figure S42:** Effect of cofactors on rate of hydrolysis

###### 2.5.6.2. Screening for Metals

To solutions containing 0.4 μM of each of the enzymes Iso1, Iso2 and 2.8 μM of Oxi1 in HEPES buffers (50 mM at pH 7.0) and 50 μM of the specified metal ions, was added 50 μM of the substrate (compound **4**) and the reaction was followed by measuring the fluorescence of released JBOMe over time. The results show that $Mn^{2+}$ and $Co^{2+}$ have the most prominent effect in enhancing the rate of reaction, and further that these effects are not cumulative. For all the assays conducted in HEPES buffer after this we therefore added 2.5 mM $MnCl_2$ to the buffer. In NaPi buffer however, these amounts of metal ions are not soluble and therefore we did not add any metal ions to the assays conducted in that buffer (see section 2.5.6.4.)

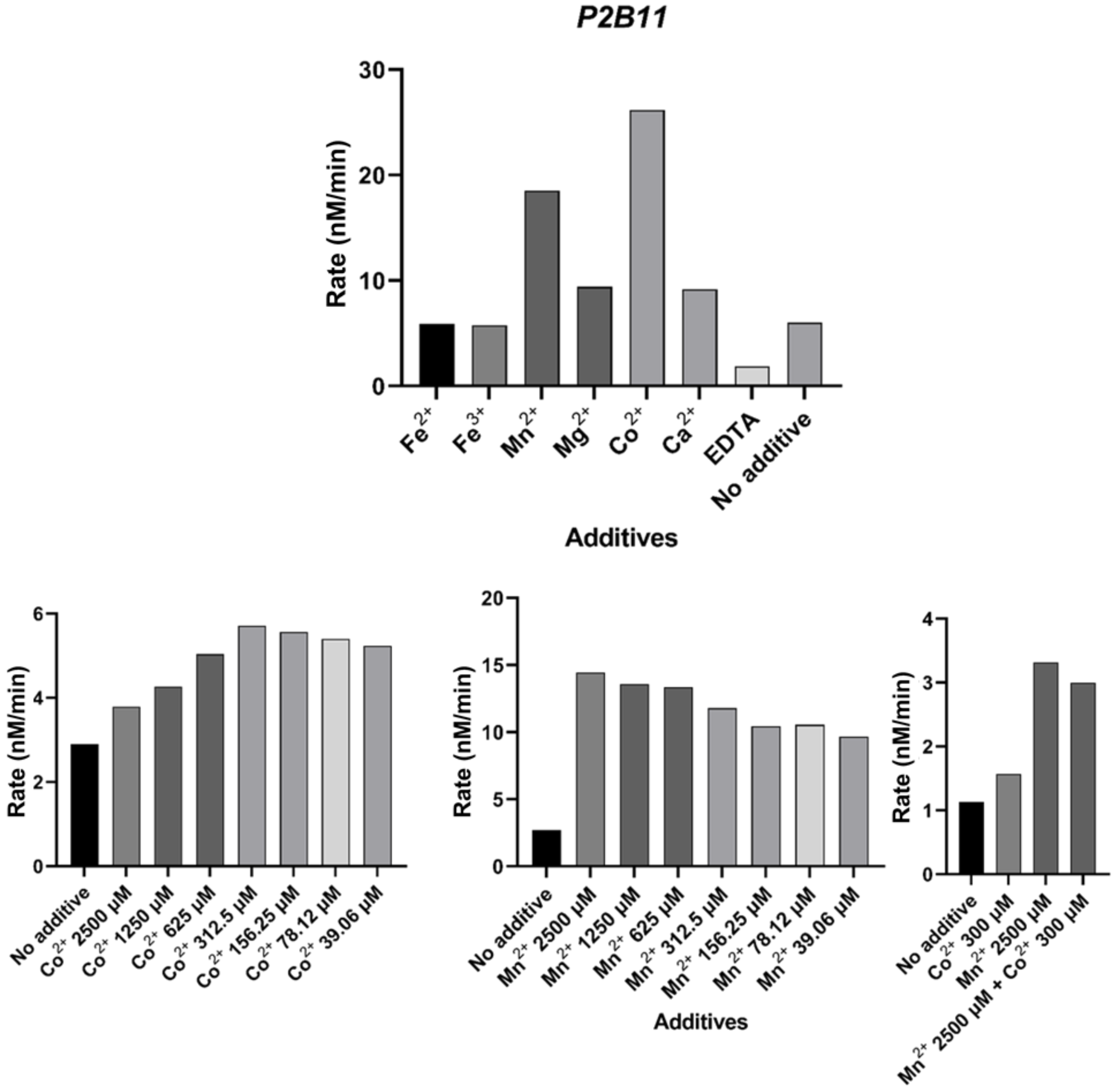

**Figure S43:** Effect of metal ions on rate of hydrolysis

###### 2.5.6.3. Effect of 3K-glucosides on rate of hydrolysis

To solutions containing 0.4 μM each of the enzymes Iso1 and Iso2 and 2.5 μM of Oxi1 in HEPES buffers (50 mM at pH 7.0) with 2.5 mM $MnCl_2$ and varying concentrations of either 3KGlcβOMe or 3KGlc-H, was added 100 μM of the substrate (compound **4**) and the reaction was followed by measuring the fluorescence of released JBOMe over time.

Addition of these 3-keto-glucosides has a prominent effect in enhancing the rate, similar to the case of the enzymes from *Alistipes* sp. and *B. theta*, even though the actual substrate here should be a 3-keto-glucuronide. Unlike the enzymes from *Alistipes* sp. and *B. theta* however, P2B11 enzymes work in the absence of the 3-keto co-substrate as well. We postulate that this is because in this case 3-keto-glucuronic acid, which is one of the products, can be used as the co-substrate. By contrast 3-keto-glucose exists primarily as a furanose isomer in solution[28] and is therefore a less useful co-substrate. Also note that in the case of the assays with P2B11 enzymes, we only observe activity when using high concentrations of Oxi1. In the case of this assay for instance, each enzyme only needs to turn over 50 molecules of substrates for a complete turn-over.

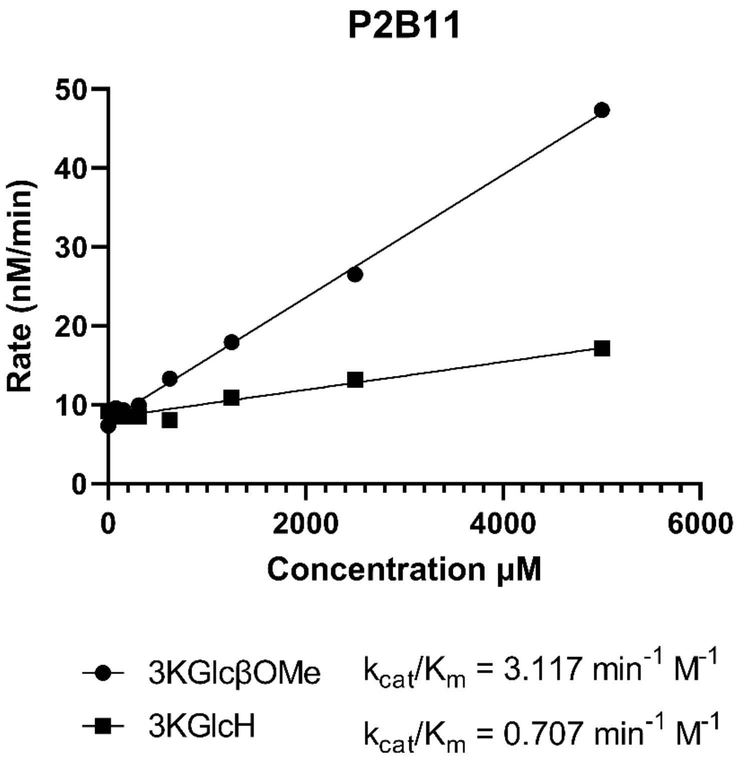

**Figure S44:** Effect of 3K-glucosides on rate of hydrolysis

###### 2.5.6.4. Effect of buffer

To solutions containing 0.4 μM of each of the enzymes Iso1 and Iso2 and 2.5 μM of Oxi1 in specified buffers (all 50 mM at pH 7.0) with 300 μM $CoCl_2$ and 1 mM of 3KGlcβOMe, was added 50 μM of the compound **4** and the reaction was followed by measuring the fluorescence of released JBOMe over time. Similar to the enzymes characterized before, the reaction is fastest in NaPi buffer, while the buffer concentration does not play a significant role here.

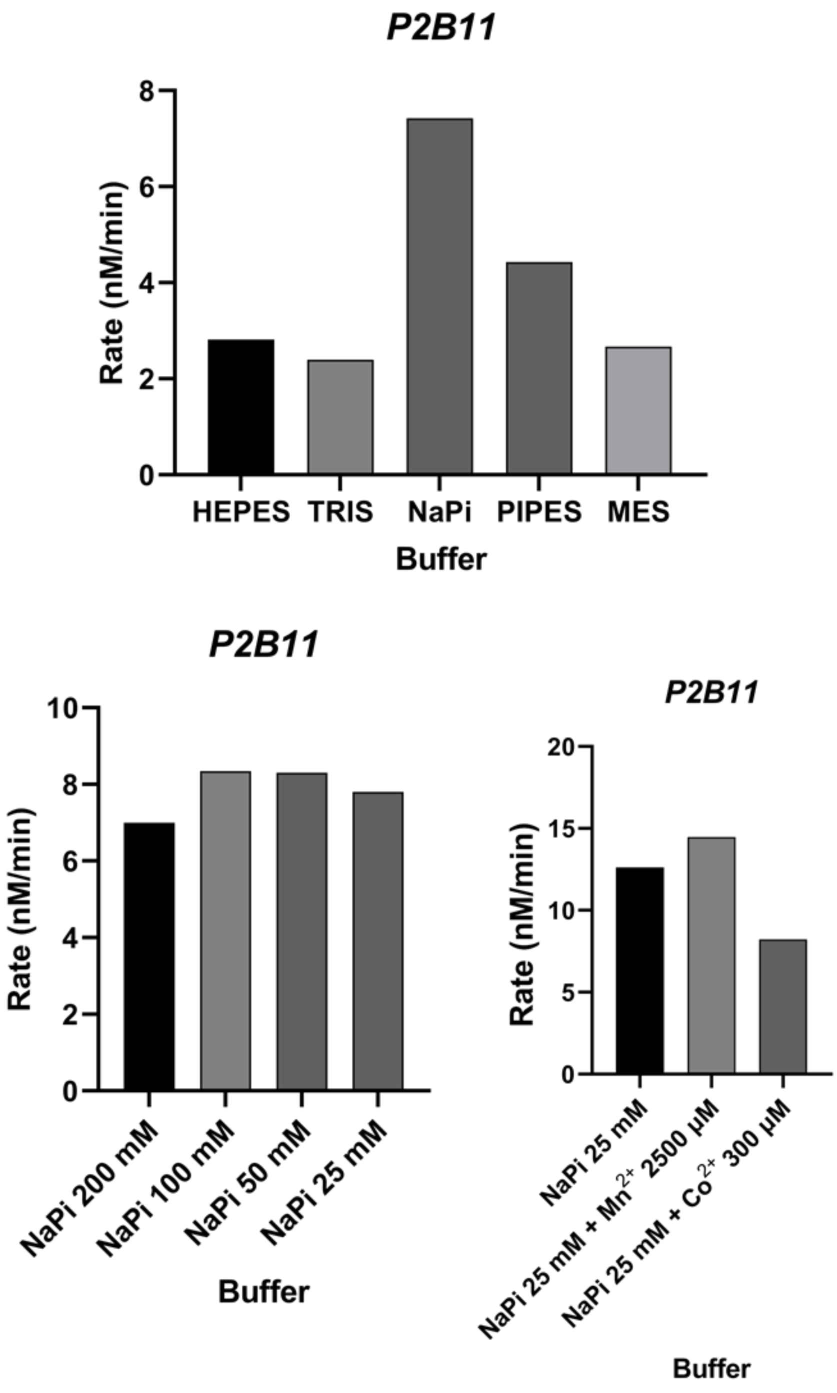

**Figure S45:** Effect of buffer conditions on rate of hydrolysis

###### 2.5.6.5. Essential enzymes

To solutions containing 1 μM each of the specified enzymes in HEPES buffers (50 mM, pH 7.0) with 2.5 mM $MnCl_2$ and 1 mM of 3KGlcβOMe was added 100 μM of compound **4** and the rate of the reaction was followed by measuring the fluorescence of released JBOMe over time. The results show that similar to the AL and BT enzymes, the presence of Oxi1 along with at least one of the enzymes labelled as an isomerase is required for observation of activity. Unlike the latter cases however, the presence of Oxi2 increases the rate of reaction here. Oxi2 is also capable of functioning in place of Oxi1, though it is significantly slower than Oxi1.

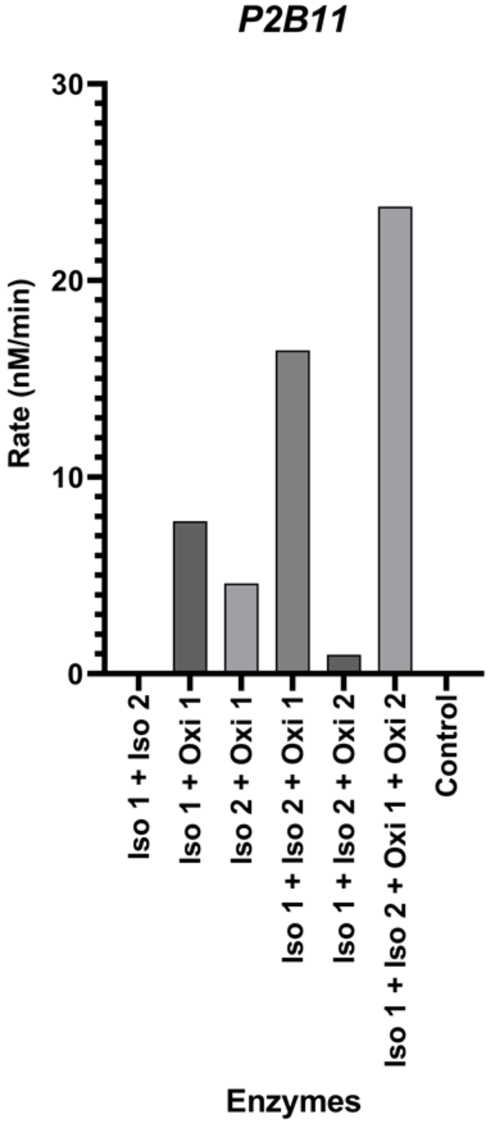

**Figure S46:** Essential enzymes

##### 2.5.6.6. Substrate specificity

To solutions containing 0.5 μM each of enzymes Iso1, Iso2, Oxi2 and 1 μM of Oxi1 in HEPES buffers (50 mM at pH 7.0) with 2.5 mM $MnCl_2$ and 1 mM of 3KGlcβOMe, was added 100 μM of the substrates specified below and the initial rate of each reaction was followed by measuring the fluorescence of released fluorophore over time.

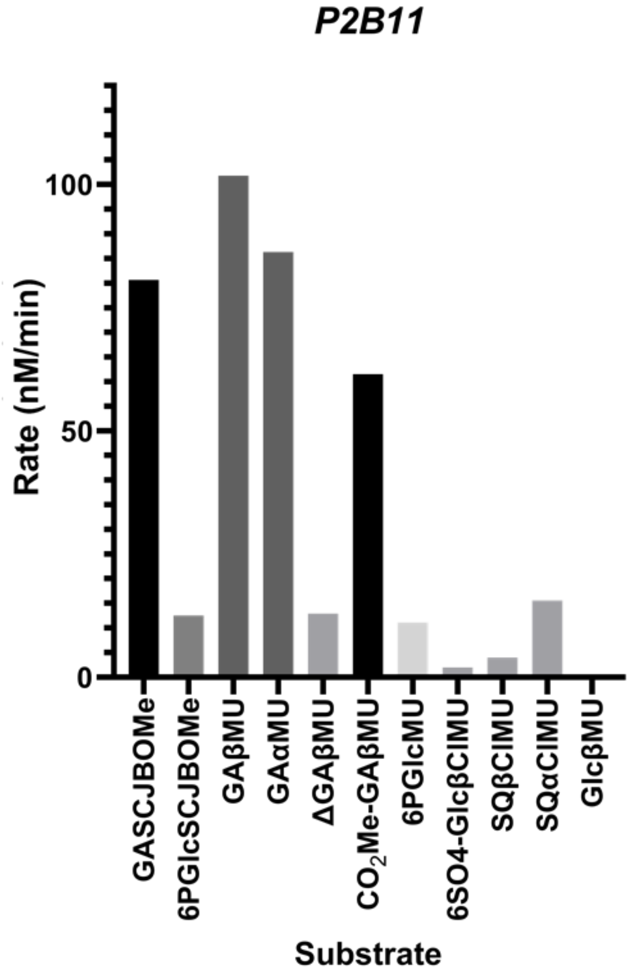

**Figure S47:** Screening for substrate specificity – P2B11 enzymes

###### 2.5.6.7. Michaelis-Menten Kinetics

To solutions containing 0.5 μM each of four enzymes in NaPi buffer (50 mM at pH 7.0) with 1 mM of 3KGlcβOMe, was added varying concentrations of the substrates and the initial rate of the reactions was followed by measuring the fluorescence of released fluorophore over time. The results were fit to a Michaelis-Menten kinetic model to yield pseudo Michaelis-Menten kinetic parameters for the four-enzyme catalysis of glycosidic bond cleavage. The $k_{cat}$ value may seem significantly lower compared to the enzymes from *Alistipes* sp. and BT but the comparison is flawed since the appropriate co-substrate is not added for P2B11 enzymes.

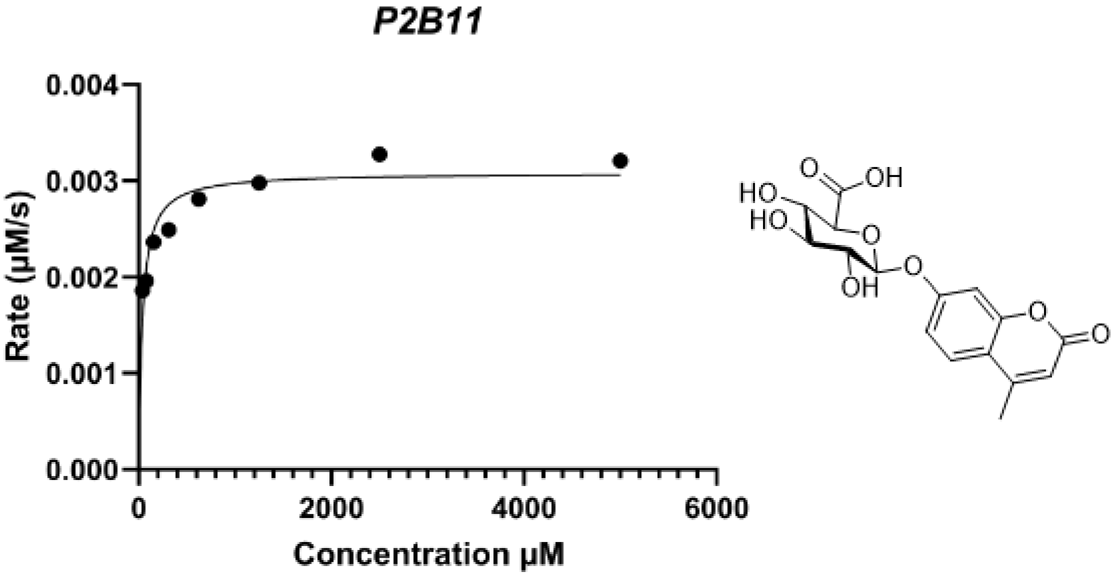

$K_m = 38 \pm 9$ μM

$k_{cat} = 0.0062 \pm 0.0002$ $s^{-1}$

**Figure S48:** Pseudo-Michaelis-Menten kinetics – P2B11 enzymes

###### 2.5.6.8. Metal dependency for individual enzymes

Similar to the case of individual enzymes from the systems discussed above, the enzymes from P2B11 also showed catalytic activity for the elimination reaction of 3-keto-glucoside substrates. The activity here however is naturally orders of magnitude lower, since the 3-keto-glucosides are not the native substrates. In comparison to the assays described in section 2.5.4., about ten times higher enzyme concentrations were used to run these assays (final enzyme concentrations: 0.7 μM) and the activity was only observable using the fluorogenic substrate (3KGlcβMU, final concentration 25 μM). Both the enzymes prefer the β anomer substrate and are activated by addition of $Co^{2+}$ and $Mn^{2+}$. Concentration of all the metal ions and EDTA in the assays is 50 μM. When EDTA and metal ions are both present, the concentration of EDTA is 25 μM. All assays are conducted in 50 mM HEPES buffer at pH 7.0.

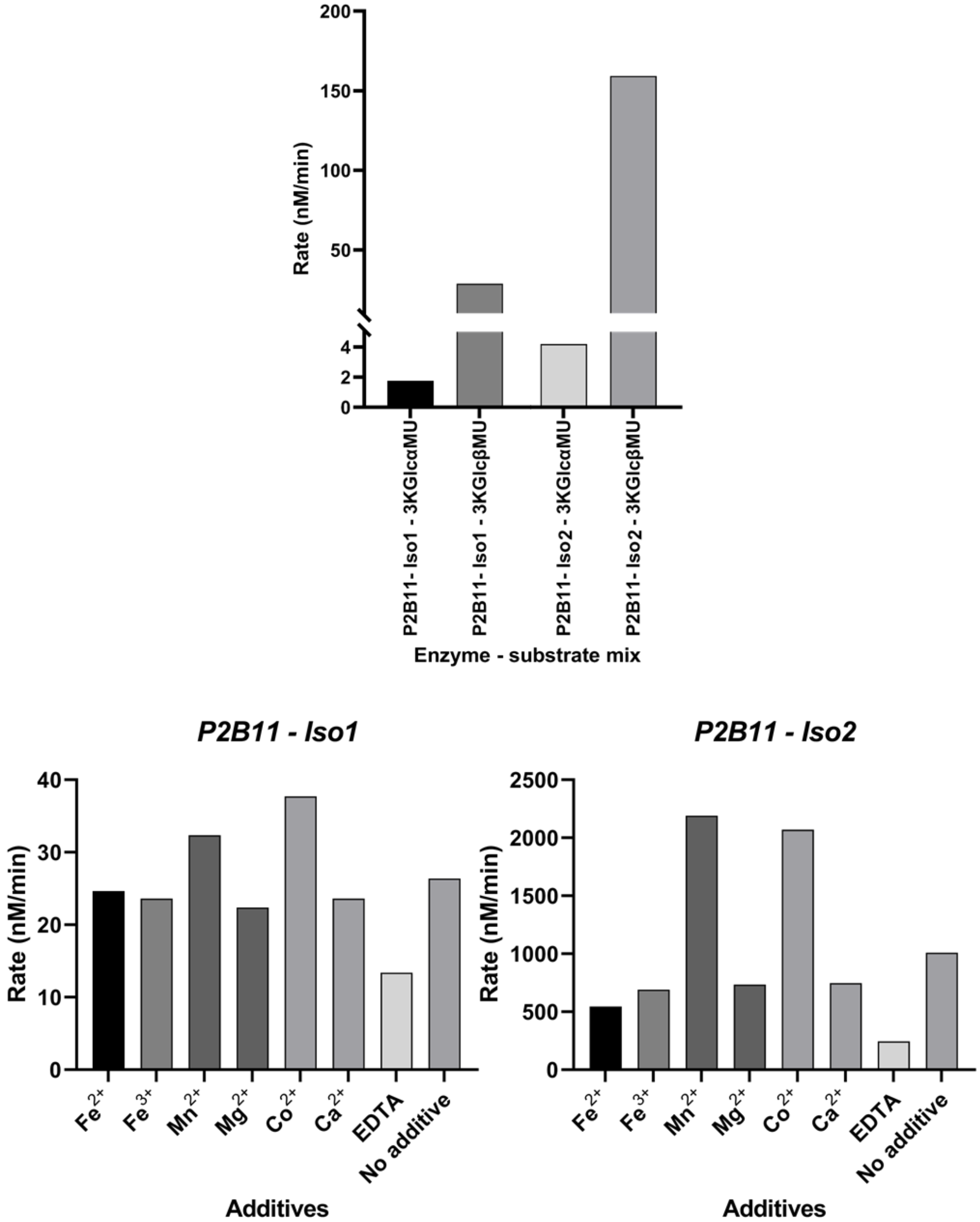

**Figure S49:** Effect of metal ions on individual P2B11 enzymes

###### 2.5.6.9. Swapping the enzymes

Solutions of a mixture of the first three enzymes of P2B11 and *Alistipes* series were made in the reaction buffer (50 mM NaPi buffer in pH 7.0 containing 250 µM of 3KGlcβOMe), where the concentrations of all enzymes were 0.25 µM and where the specified enzymes were replaced from the other series, one at the time. The replaced enzyme is specified for each reaction. To these solutions was added 500 µM of the substrates (GAβMU or GlcβMU) and the reaction was followed by measuring the fluorescence of released MU over time. Initial rates were calculated for the first 2 minutes of the reactions. Based on the protein sequence similarity of the proteins from the two series, it was assumed that AL1 is the counterpart to P2B11-Iso2, AL3 is the counterpart to P2B11-Oxi1 and this leaves AL2 to be the counterpart to P2B11-Iso1, despite lack of similarity.

The results show that the enzymes from the two series are interchangeable in some but not all cases. Specifically, it is clear that while AL3 is capable of oxidizing glucuronide substrates, P2B11-oxido1 is not able to oxidize glucosides effectively. This also points to the general observation that P2B11 enzymes are significantly slower than the other series in the *in vitro* assays. This is at least partially because we are trying to recycle the cofactor of P2B11 series via addition of 3-keto glucosides, which are not good co-substrates in this case.

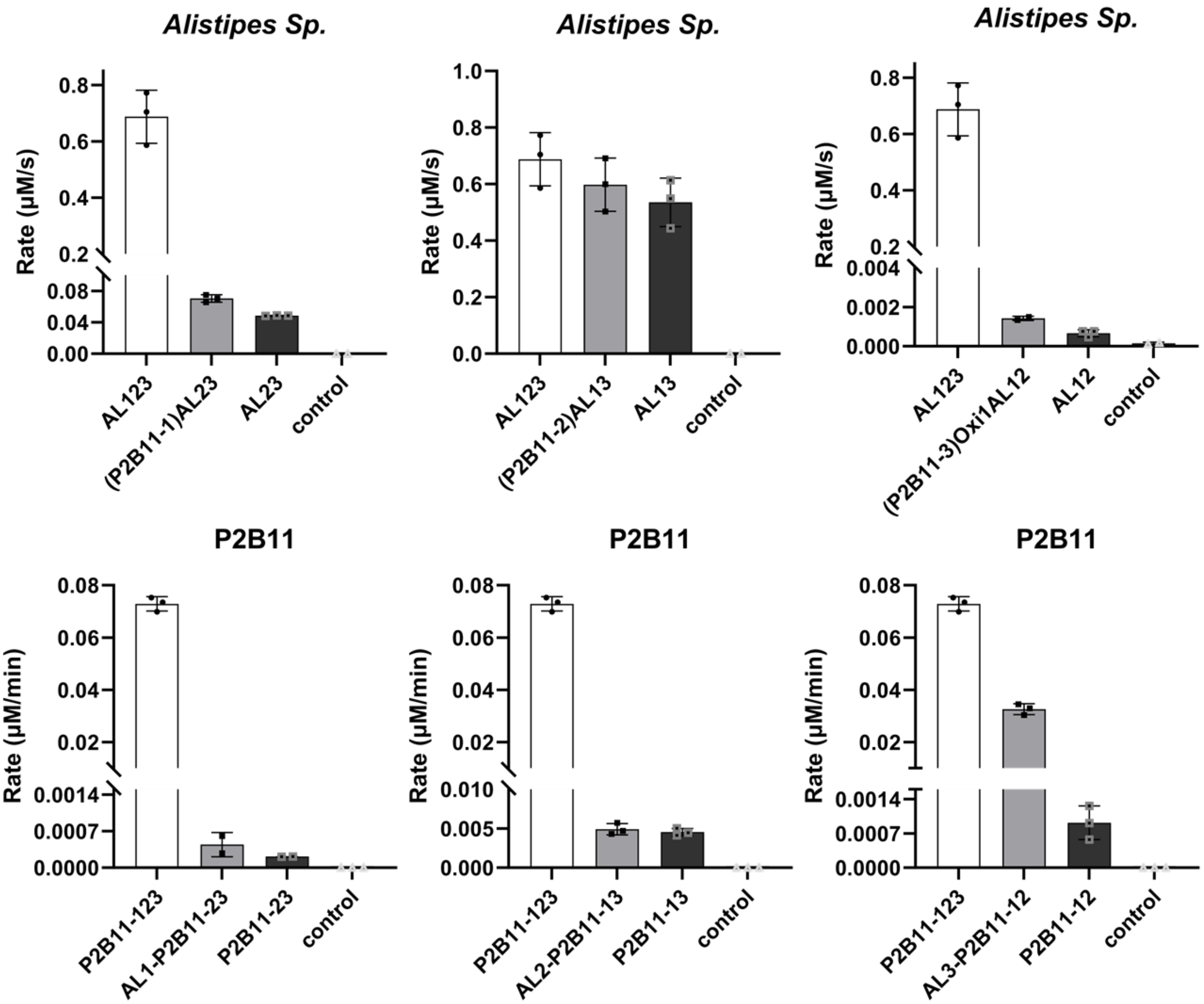

**Figure S50:** Swapping the enzymes between P2B11 and *Alistipes* series, (bars indicate the mean, and error bars the SD of technical replicates (n=3)).

##### 2.5.7. Assays for Identifying Co-substrate

Solutions of a set of various carbonyl-containing compounds all at 500 µM and a mixture of the enzymes of the *Alistipes* series were made in reaction buffer (25 mM NaPi buffer, pH 7.0 containing 50 µM of $CaCl_2$), where the concentration of each enzyme was 0.02 µM. To these solutions was added 500 µM of the substrate GlcβMU and the initial rates of the reactions were calculated based on change in fluorescence of the solution.

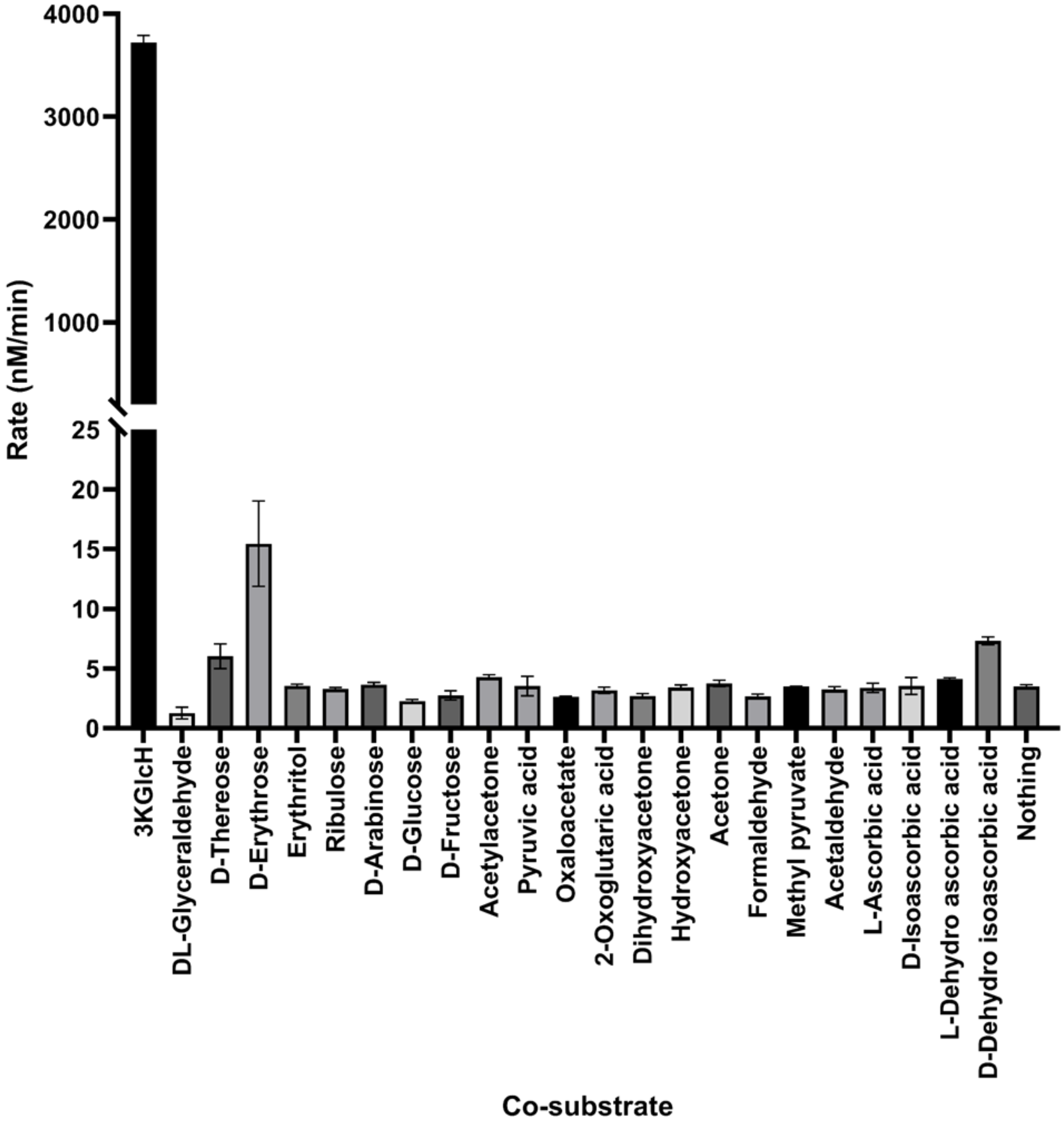

**Figure S51:** Screening for co-substrates, (bars indicate the mean, and error bars the SD of technical replicates (n=3)).

#### 2.6. Chromatography assisted assays of enzymatic reactions

##### 2.6.1. LC-MS studies

All experiments were done using an Agilent 1260 Infinity HPLC equipped with a UV detector and an Agilent 6120 Quadrupole LC/MS detector. The buffer for these experiments was the same buffer used in kinetic experiments, 25 mM NaPi buffer pH = 7.0 with the appropriate metal ions. In all the experiments, the reactions were set up using 1 mM of the specified substrates and 0.2 µM of enzymes unless otherwise specified. Upon addition of the enzyme(s), reaction mixtures were incubated at 37° C for the specified amount of time. 10 µL of the reaction solutions was then diluted into 90 µL of acetonitrile to precipitate the enzyme. The solutions were then centrifuged for 3 minutes, and the top soluble portion was used to prepare LC-MS samples. For each LC-MS run, 5 µL of the sample was injected into the column (Agilent Eclipse XBD C-18) which was washed with water (solvent A) and acetonitrile (Solvent B) using the following steps: 0 to 1 min, 95% A, 1 to 11 min gradual change to 5% A, 11 to 12 min, 5% A, 12 to 15 min, gradual change to 95% A. The identity of the peaks is verified by assessing the MS spectra, and when possible, with the use of standard samples assayed under the same conditions.

###### 2.6.1.1. Characterization of reaction catalysed by BT3

Treatment of a mixture of CellβMU and 3KGlcH with BT3 results in formation of a new peak with the M.W. of the product, 3KCellβMU.

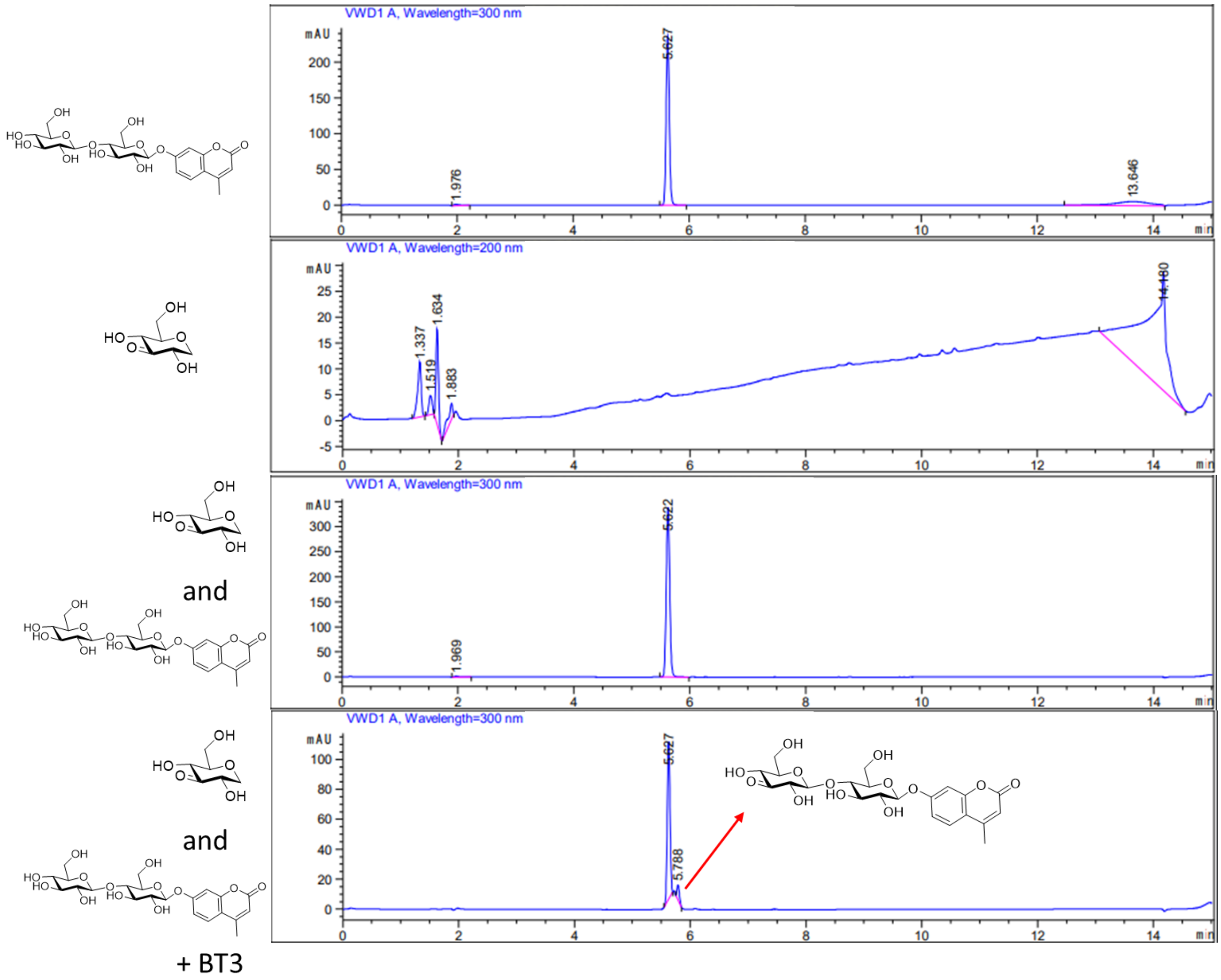

**Figure S52:** LCMS trace for reaction of Cell βMU and 3KGlcH catalyzed by BT3

##### 2.6.1.2. Characterization of reaction catalysed by BT1, BT2 and BT3

Treatment of a mixture of GlcβMU (1 mM) and 3KGlcH (500 µM) with BT1, BT2 and BT3 results in consumption of the peak for GlcβMU and emergence of new peaks corresponding to the product, MU. No separate peak for the 3Keto intermediates was detected in the mixture, indicating that the oxidation step is the rate determining step in the reaction. Note that conversion is not complete since the co-substrate is the limiting reagent.

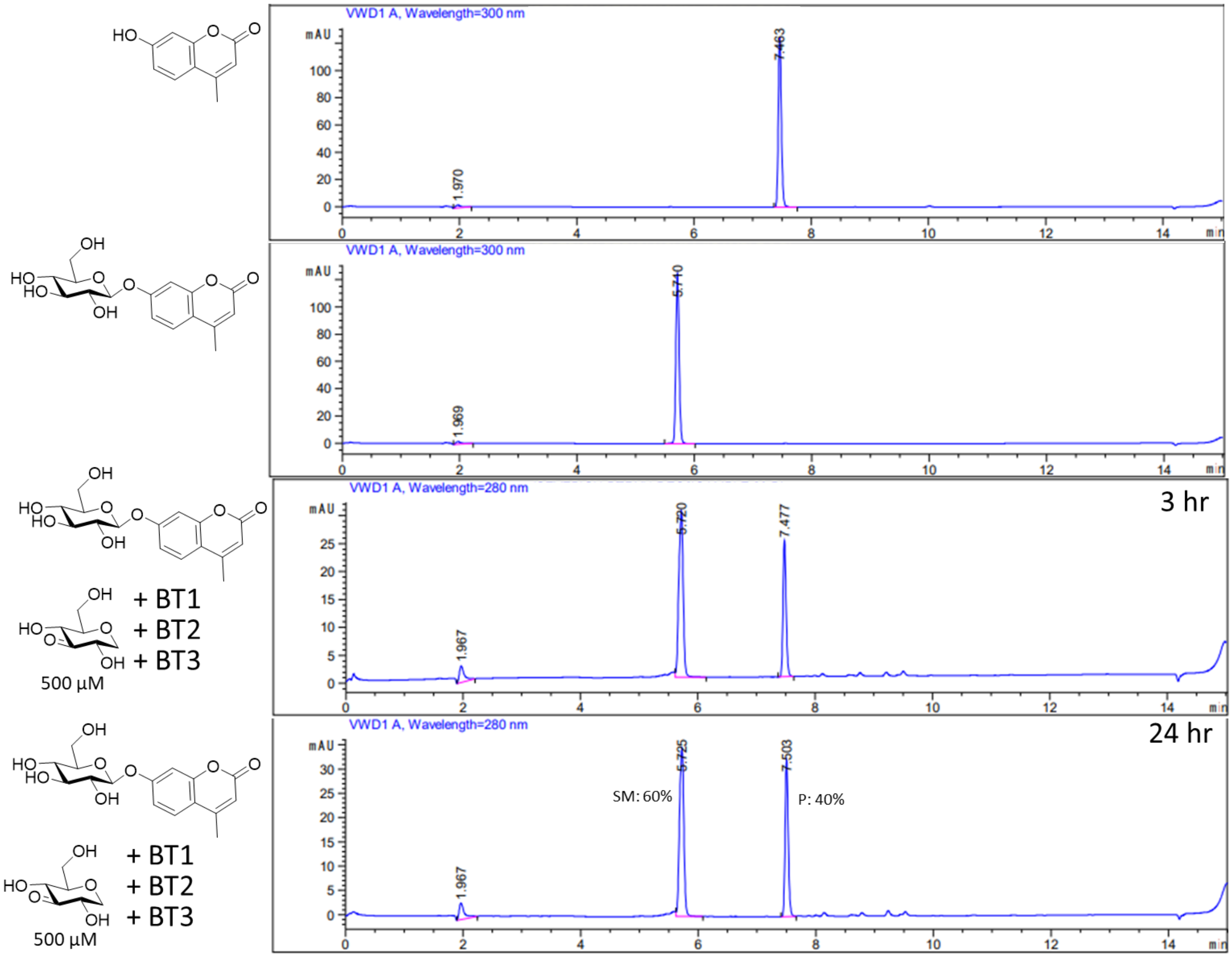

**Figure S53:** LCMS trace for hydrolysis of GlcβMU catalyzed by BT123 - 1

Treatment of a mixture of GlcβMU (1mM) and 3KGlc-H (2 mM) with BT1, BT2 and BT3 results in consumption of the peak for GlcβMU and emergence of new peaks corresponding to the product MU. Note that conversion is almost complete in this case since more than 1 eq. of the co-substrate is used.

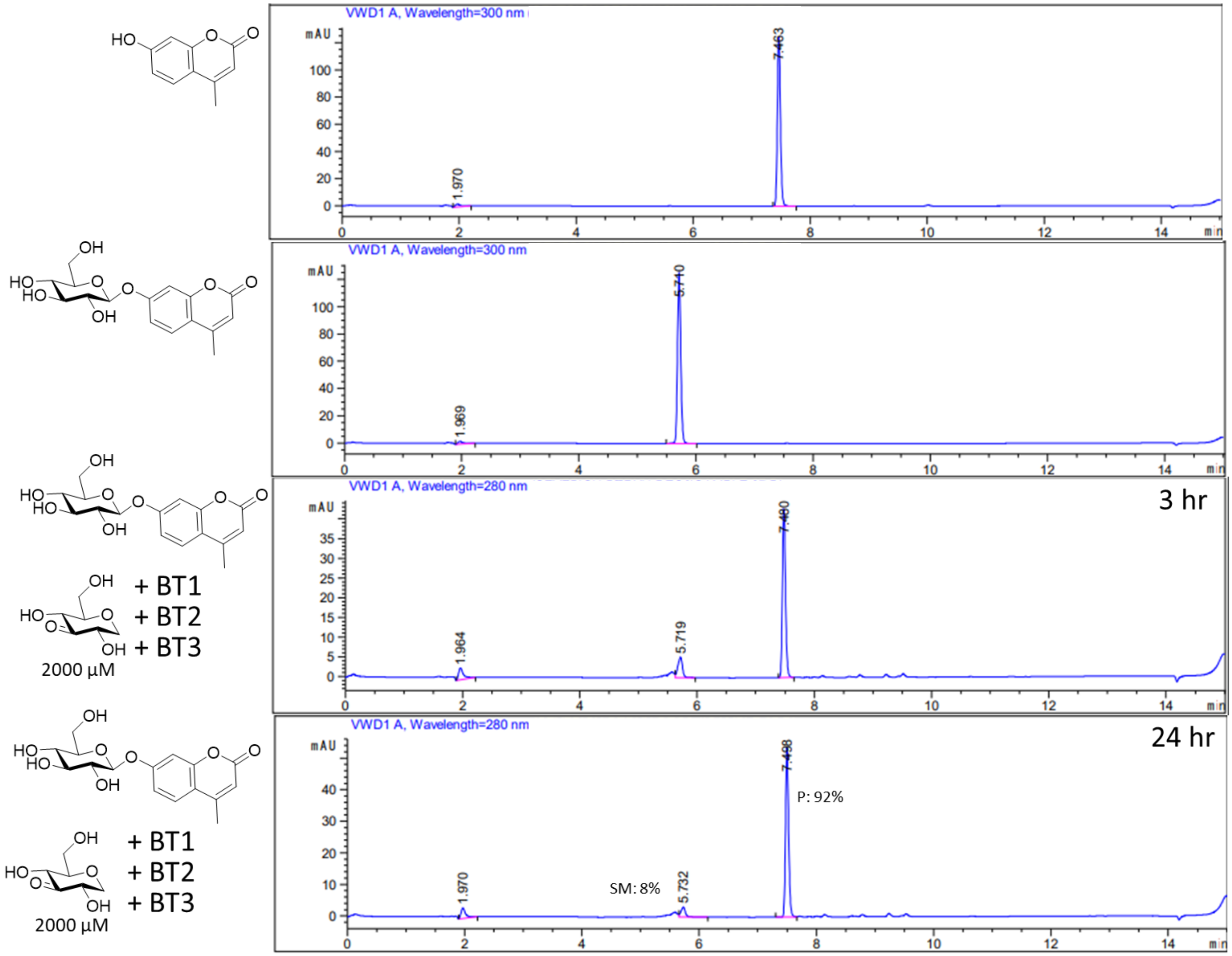

**Figure S54:** LCMS trace for hydrolysis of GlcβMU catalyzed by BT123 - 2

Treatment of a mixture of GlcβMU (1 mM) and 3KGlcH (500 µM) with BT1, BT2 and BT3 results in consumption of the peak for GlcβMU and emergence of a new peak corresponding to the product MU. Note that conversion is not complete since the co-substrate is the limiting reagent, however in the presence of NAD, the final conversion is slightly higher than in its absence (48% compared to 40%). Even though addition of NAD cofactor does not have any measurable effect on the rate of the reaction, it is presumably able to exchange with the NAD bound in BT3 slowly over time and this results in increased total turn-over with the reaction that is incubated for 24 hours.

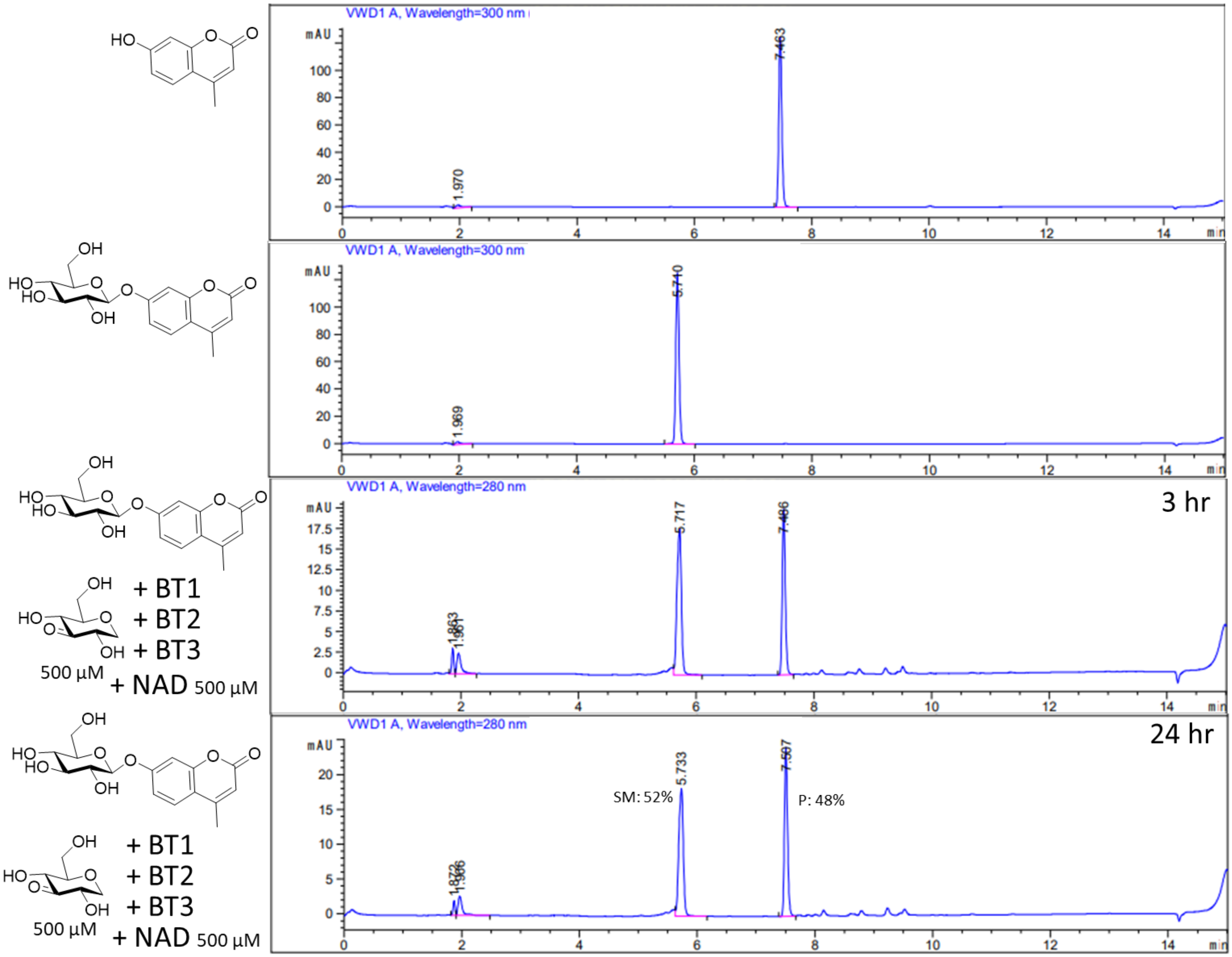

**Figure S55:** LCMS trace for hydrolysis of GlcβMU catalyzed by BT123 - 3

Treatment of a mixture of CellβMU (1 mM) and 3KGlcH (500 µM) with BT1, BT2 and BT3 results in formation of new peaks corresponding to the products GlcβMU and MU within one hour. No separate peak for the 3Keto intermediates was detected in the mixture, indicating that the oxidation step is the rate determining step in the reaction. Note that here the reaction was monitored at early stages to detect the intermediates. Given enough time, the reaction goes to completion (see section 2.6.2).

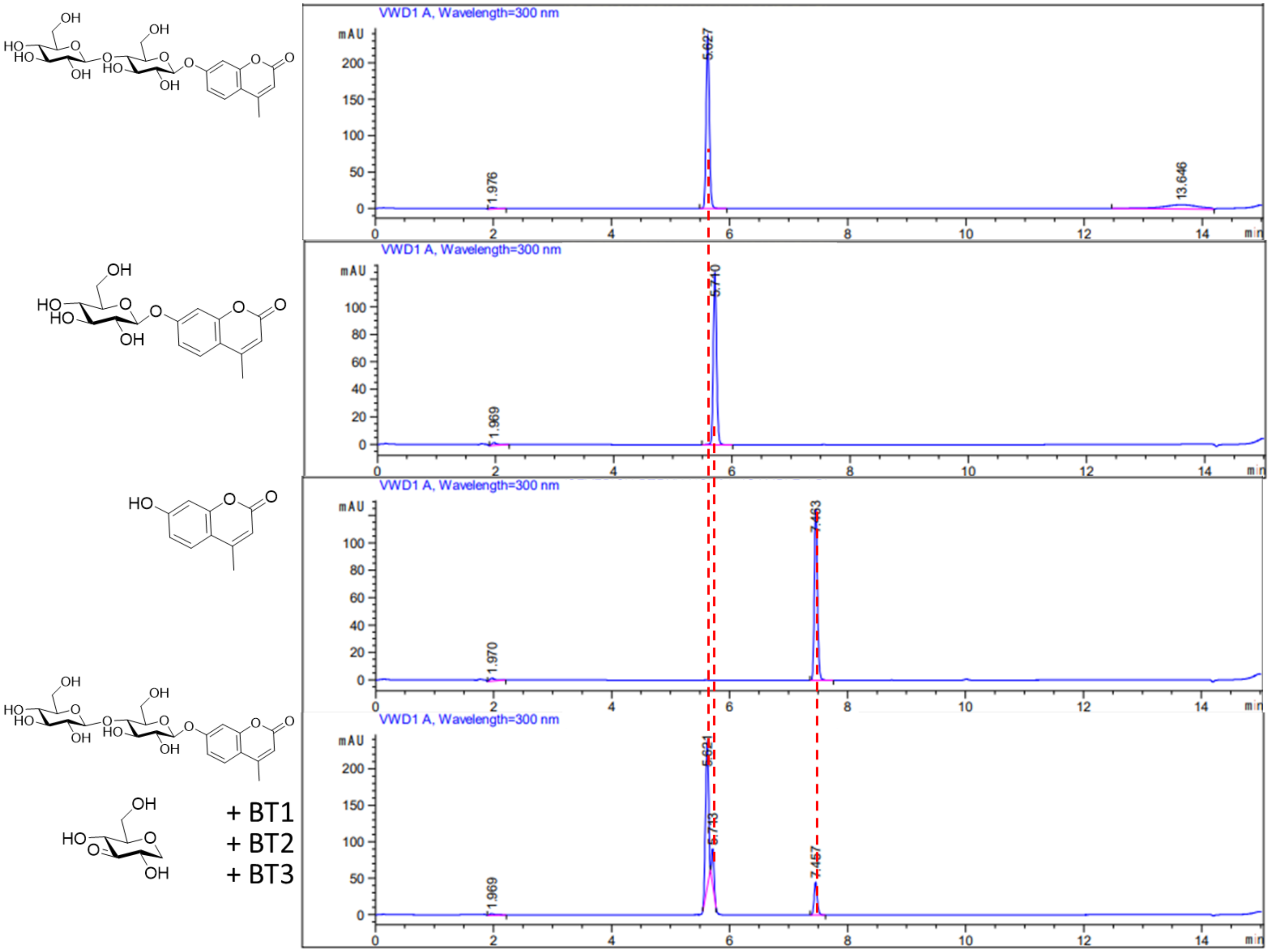

**Figure S56:** LCMS trace for hydrolysis of CellβMU catalyzed by BT123

###### 2.6.1.3. Inspection of reactions with challenging substrates

All reactions in this section were performed to investigate whether any hydrolysis can be observed with challenging substrates, those that did not show any sign of hydrolysis when inspected with other methods. 2-Fluoro-glucosides, C-glycosides and amino-glucosides were found to not be substrates of our enzymes, with the exception of 2-amino-α-glucoside. As well, in agreement with the previous observations, the OH groups at 3 and 4 positions are strictly required to observe the activity – 3 and 4 methylated glucosides are also not accepted as substrates.

###### 2.6.1.3.1. Inspection of reaction with 2-fluoro-Glc-β-6,8-difluoroMU

Treatment of a mixture of 2-fluoro-Glc-β-$F_2$MU substrate (1 mM) and 3KGlcH (500 µM) with BT1, BT2 and BT3 in the same conditions as the reactions above results in release of a very small amount of the fluorophore.

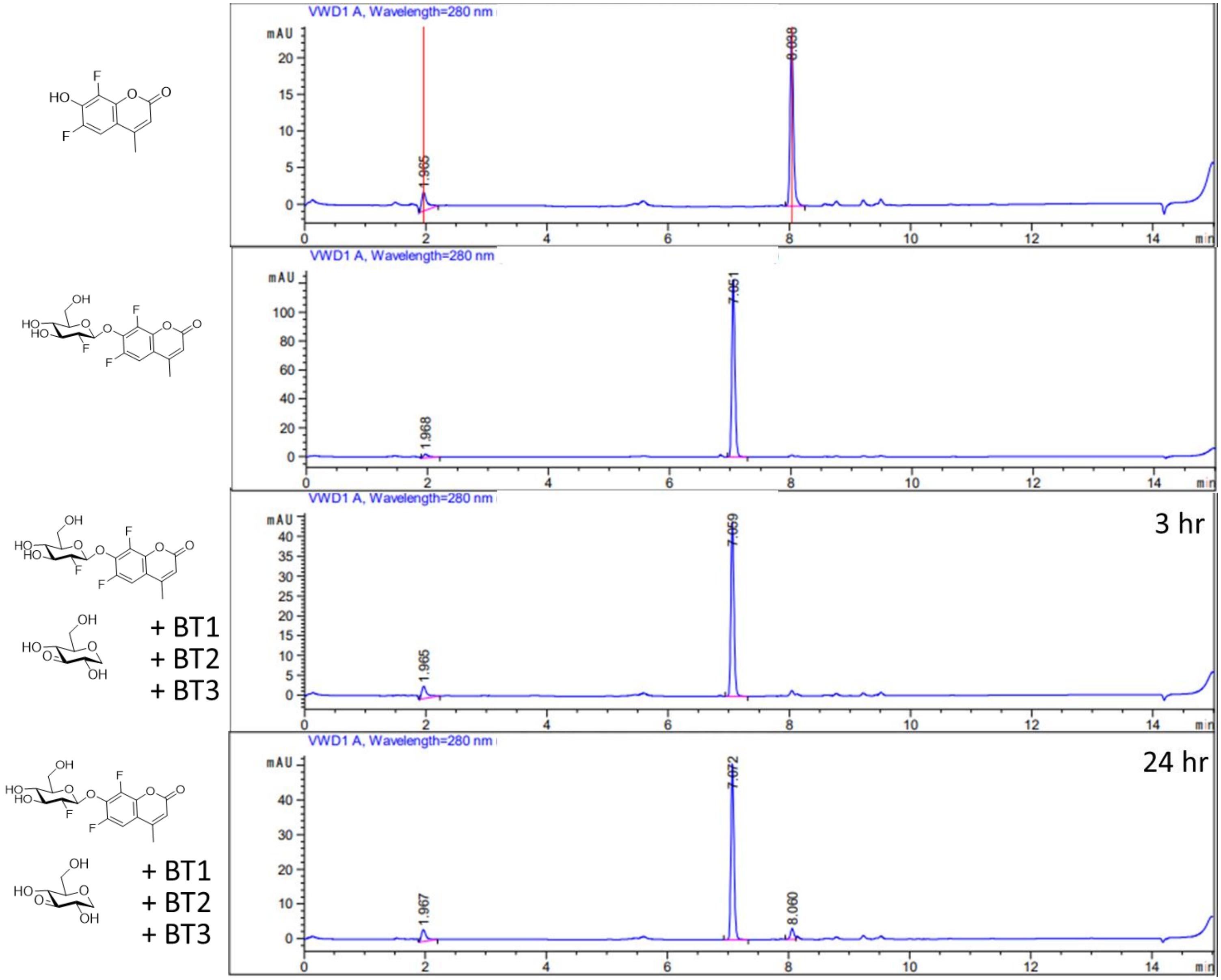

**Figure S57:** LCMS trace for attempted hydrolysis of 2FGlcβMU catalyzed by BT123

###### 2.6.1.3.2. Inspection of reaction with C-glycosides

Treatment of a mixture of two different C-glycoside substrates (Puerarin and Dapagliflozin, 1 mM each) and 3K-Glc-H (500 µM) with BT1, BT2 and BT3 in the same conditions as the reactions above results in formation of 3-keto glycosides but no observable cleavage of the C-glycosidic bond.

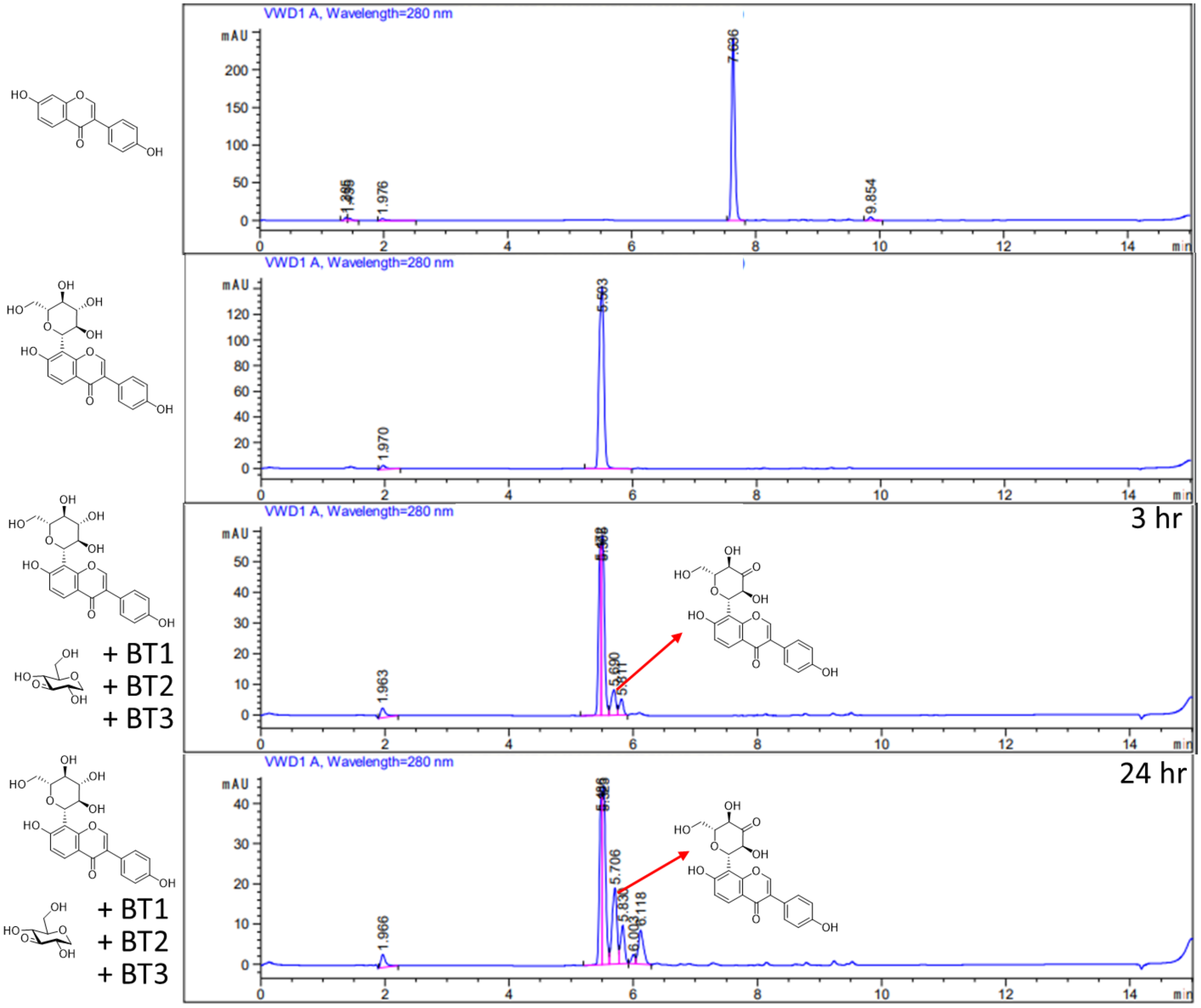

**Figure S58:** LCMS trace for attempted hydrolysis of Puerarin catalyzed by BT123

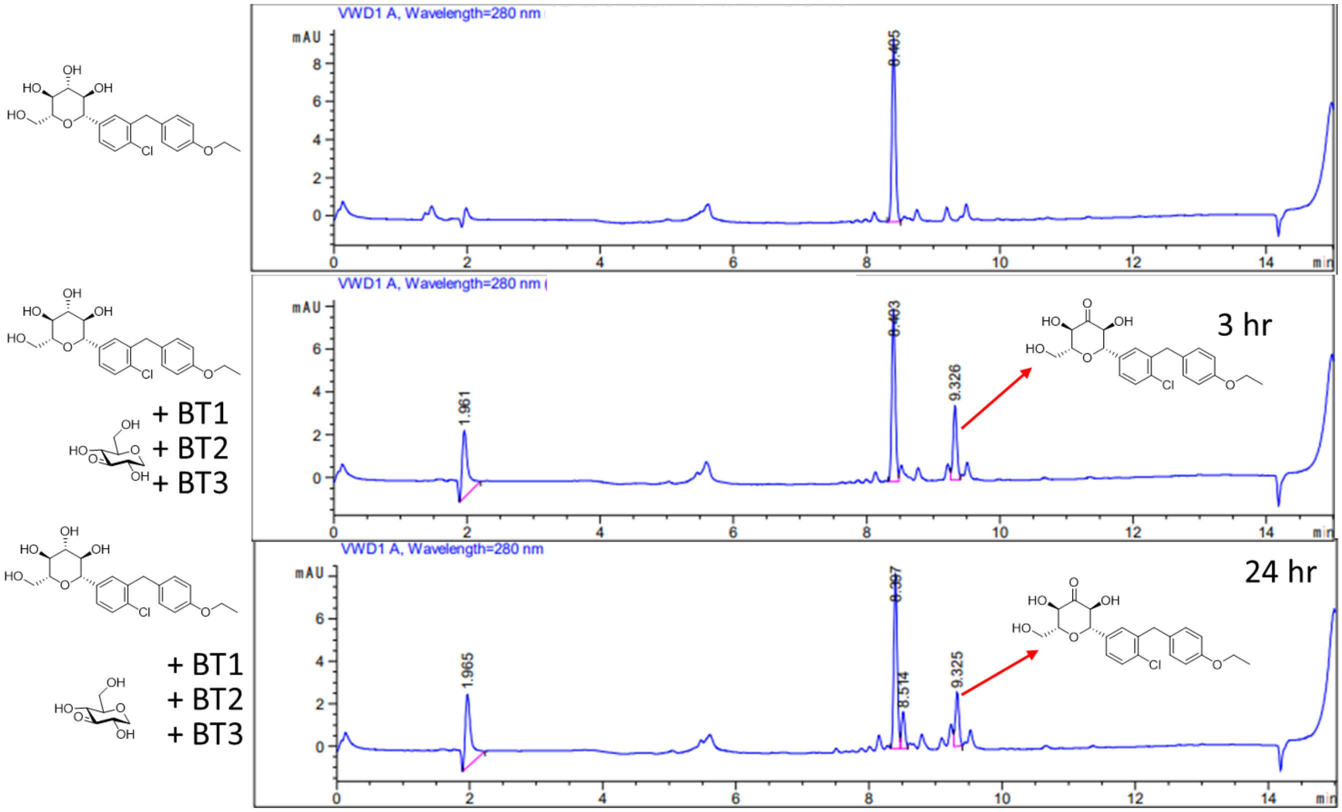

**Figure S59:** LCMS trace for attempted hydrolysis of Dapagliflozin catalyzed by BT123

##### 2.6.1.3.3. Inspection of reaction with amino-glycosides

Treatment of solutions of different deoxy-amino-glycoside substrates (2-amino-Glc-α-pNP, 3-amino-Glc-β-MU, 4-amino-Glc-β-MU, 6-amino-Glc-β-MU, 1 mM each) and 3K-Glc-H (1 mM) with BT1, BT2 and BT3 under the same conditions as the reactions only results in hydrolysis of the glycosidic bond for the 2-amino-α-glucoside. While 3- and 4-OH groups are required by AL/BT3 for binding and catalysis, the 6-amino group is probably required for the binding of BT1. These findings fully explain why amino-glycoside antibiotics are not substrates of these enzymes (see SI section 2.6.2.).

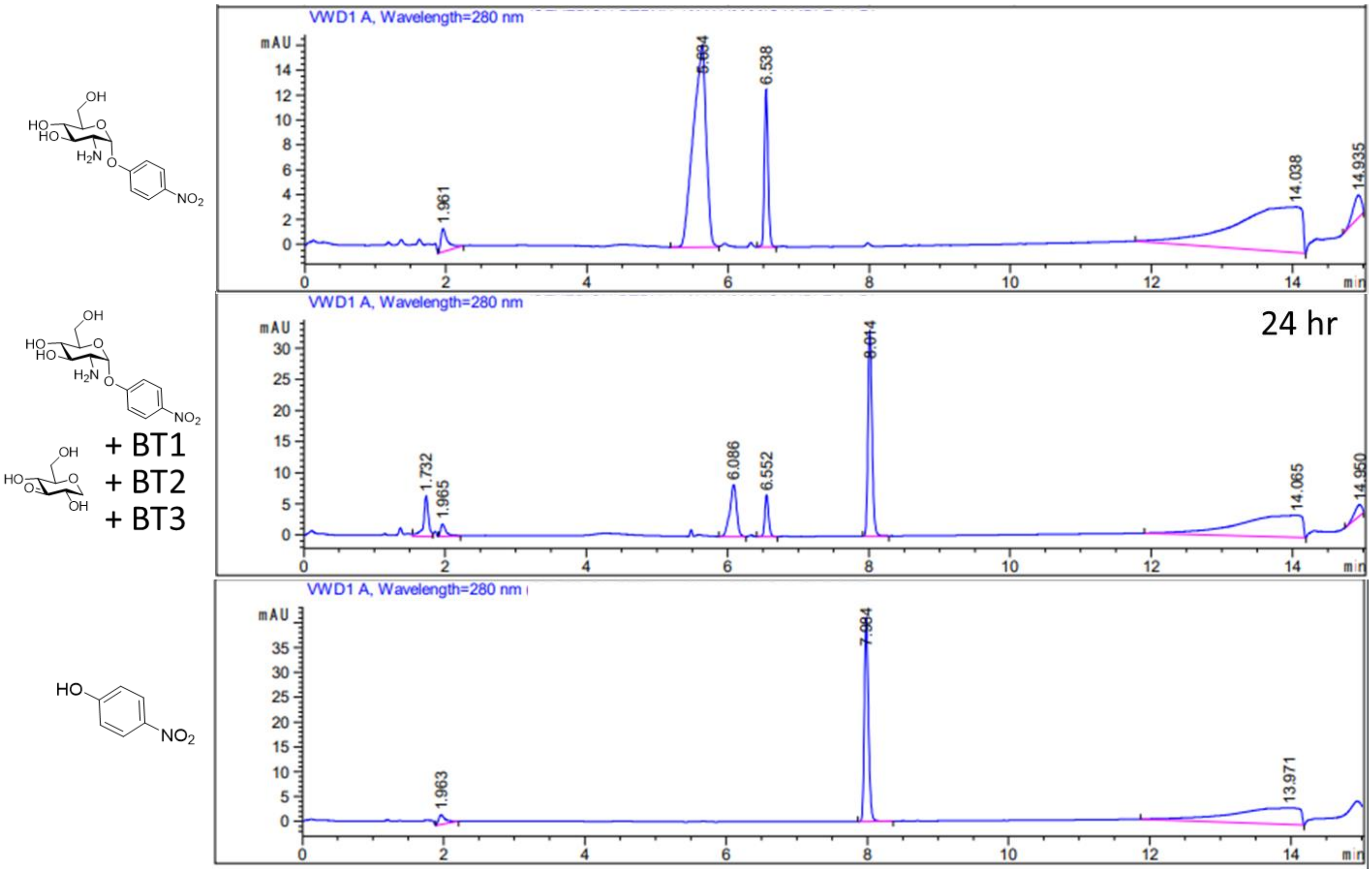

**Figure S60:** LCMS trace for hydrolysis of 2-amino-Glc-α-pNP catalyzed by BT123

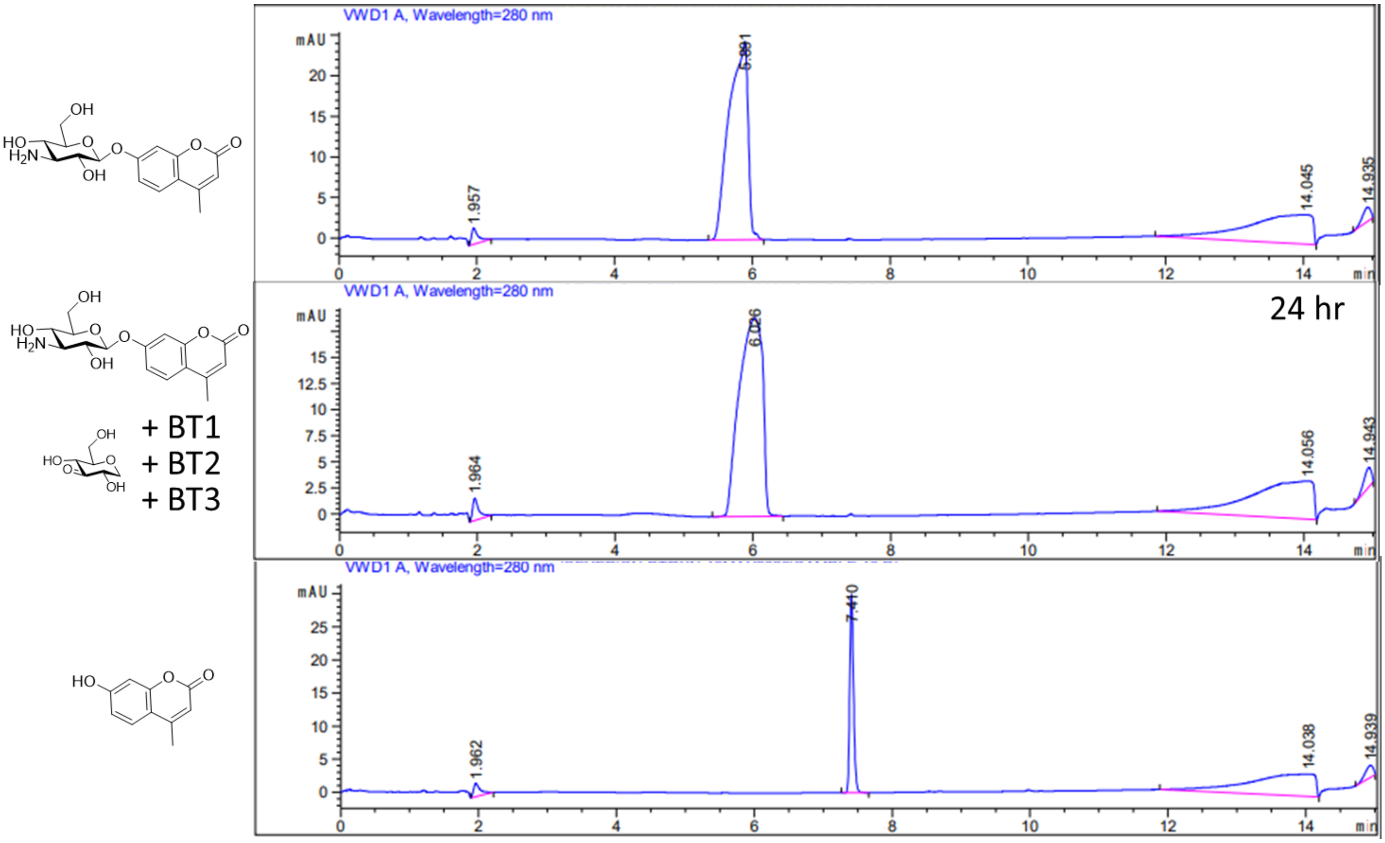

**Figure S61:** LCMS trace for attempted hydrolysis of 3-amino-Glc-β-MU catalyzed by BT123

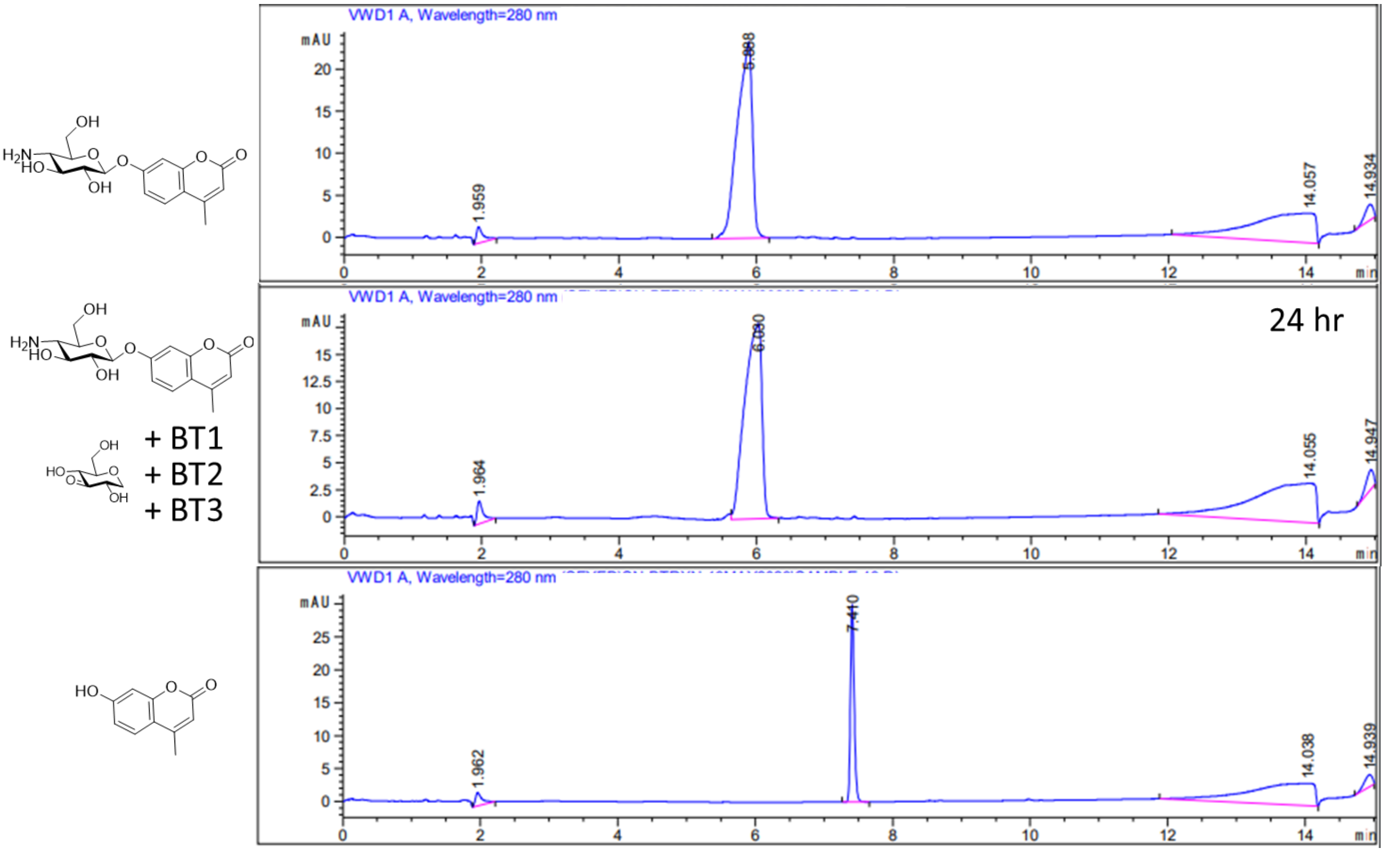

**Figure S62:** LCMS trace for attempted hydrolysis of 4-amino-Glc-β-MU catalyzed by BT123

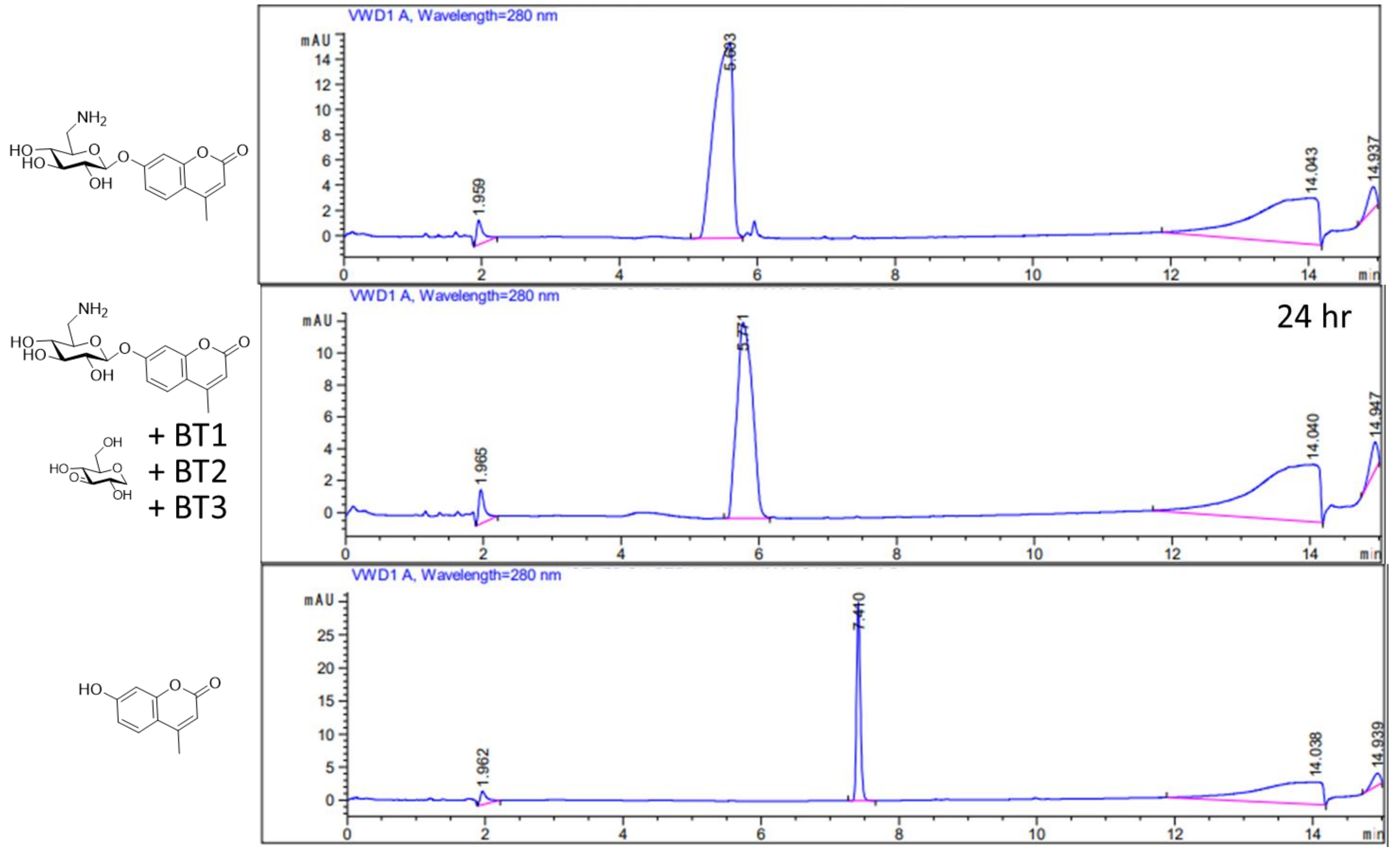

**Figure S63:** LCMS trace for attempted hydrolysis of 6-amino-Glc-β-MU catalyzed by BT123

**2.6.1.3.4. Inspection of reaction with 3- and 4-OMe-glycosides**

Treatment of solutions of different glycoside substrates with blocked 3- and 4- groups (3-OMe-Glc-β-MU and 4-OMe-Glc-β-MU, 1 mM each) and 3K-Glc-H (1 mM) with BT1, BT2 and BT3 under the same conditions as the reactions above does not result in hydrolysis of the glycosidic bond.

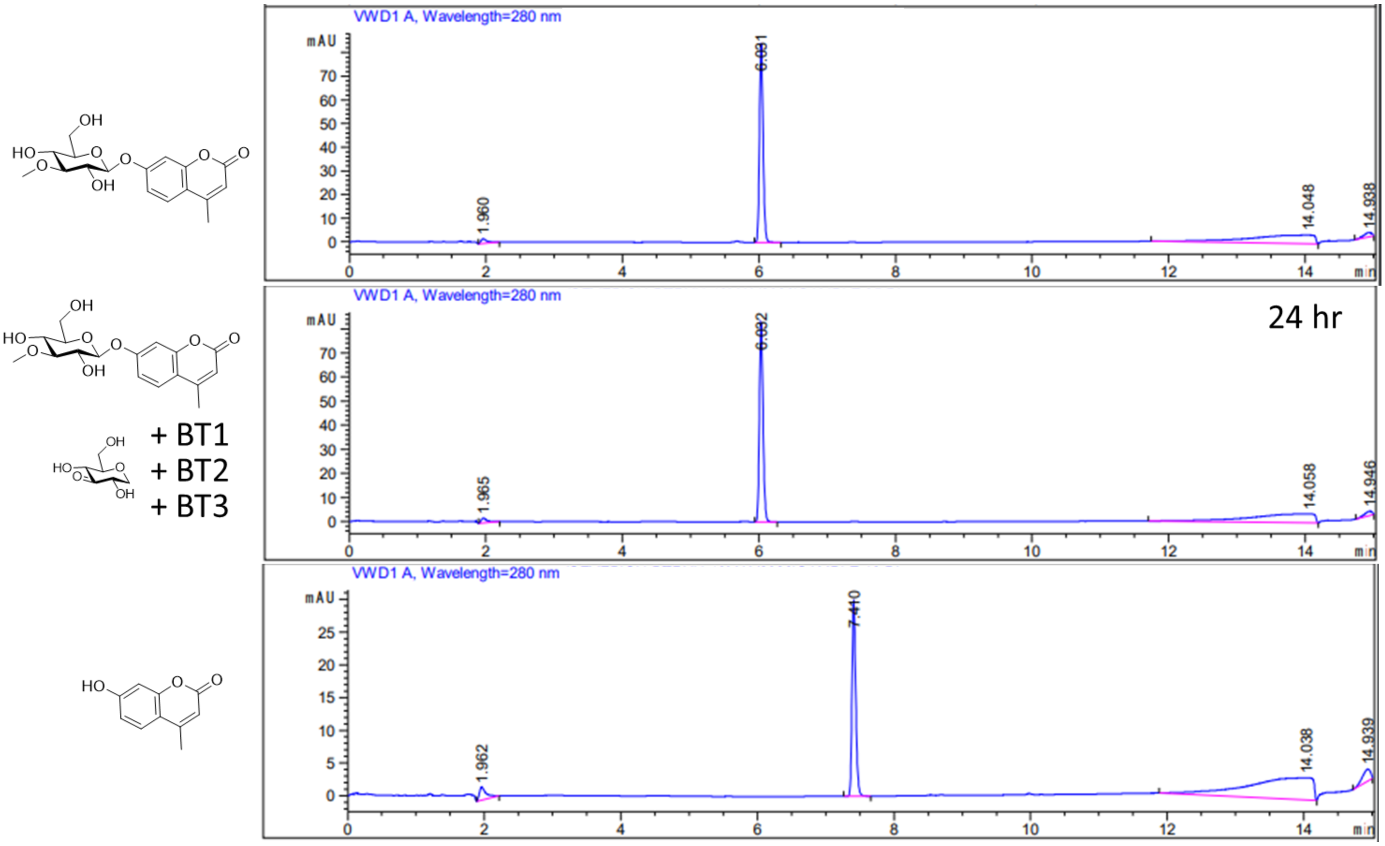

**Figure S64:** LCMS trace for attempted hydrolysis of 3-OMe-Glc-β-MU catalyzed by BT123

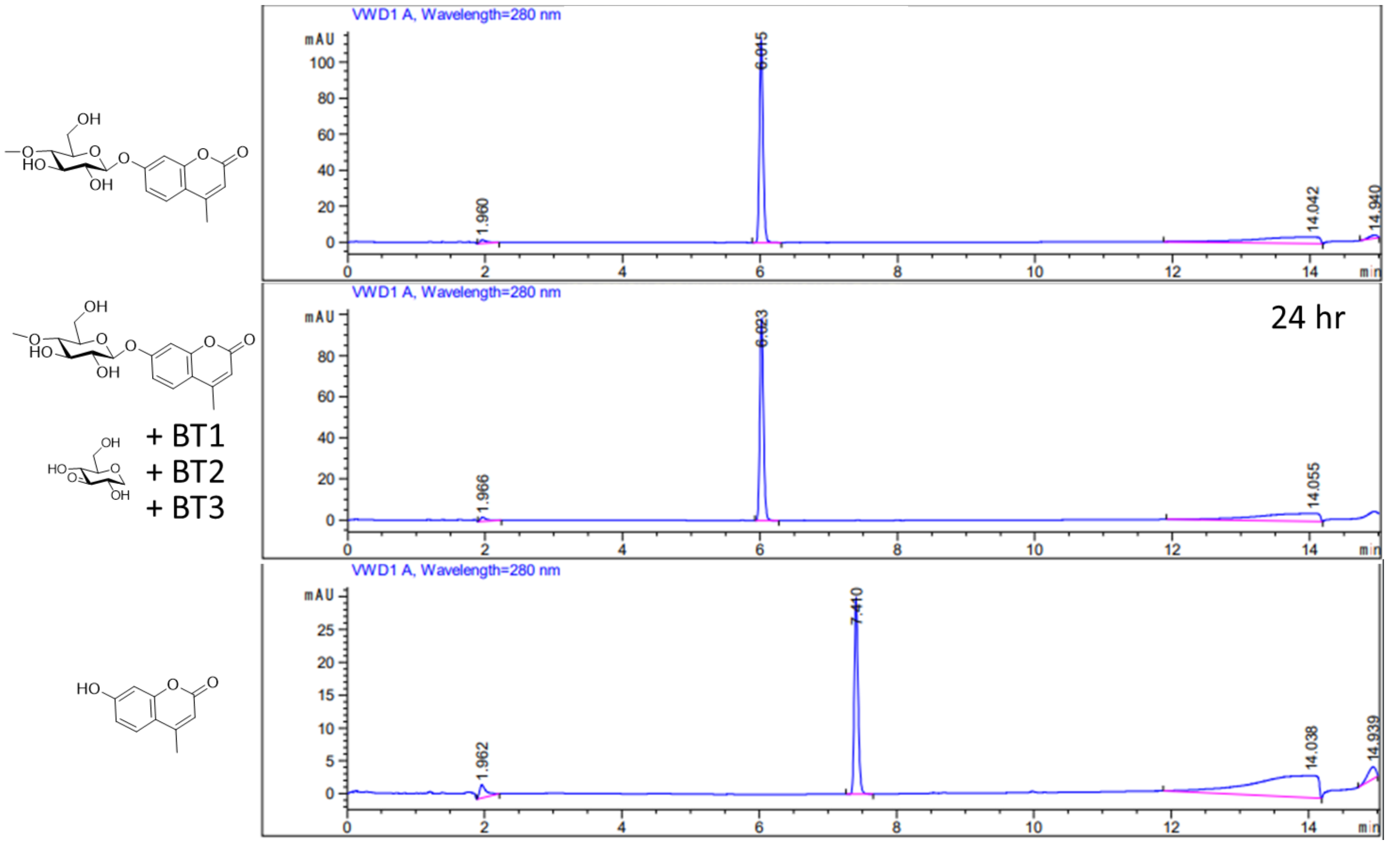

**Figure S65:** LCMS trace for attempted hydrolysis of 4-OMe-Glc-β-MU catalyzed by BT123

##### 2.6.2. Thin layer chromatography studies

All the reactions were performed in the same buffer that was used in kinetic experiments, 25 mM NaPi buffer pH 7.0, with the appropriate metal ions added for each respective system of enzymes. In all the experiments, the reactions were set up using 5 mM of substrates (2 mg/mL for polysaccharides), 4 mM of 3KGlcH and 5 µM of each enzyme. Upon addition of the enzyme, solutions were incubated at 37° C overnight (16 hours). 0.5 µL of the reaction solutions was then spotted on the TLC plates, which were developed using t-Butanol:Ammonium hydroxide:Methanol:Water with the volumetric ratios of 5:4:4:1 as eluent. Controls involved mixtures of the substrates in the assay buffer and the standard for the expected product(s) but no enzyme. The plates were then stained using Hanessian's Stain (Ammonium Molybdate 4.8% w/v, Ceric ammonium molybdate 0.2% w/v, Sulfuric acid 6% v/v in Water). Note that in these conditions 3-keto-glucose and 1,5-anhydroglucitol do not stain (Fig. S67).

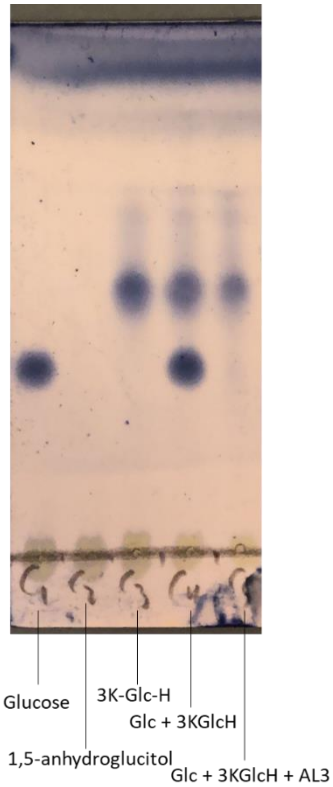

**Figure S66:** TLC reference spots for glucose, 1,5-anhydroglucitol (line 2, doesn't stain), 3KGlcH and 3KGlucose (line 5, doesn't stain)

###### 2.6.2.1. Saccharide substrates of AL enzymes

Analysis of reaction of enzymes AL1-4 with a series of various disaccharides and pseudo-saccharides shows that these enzymes hydrolyse all the substrates tested with the exception of lactose, mellibiose, raffinose, voglibose and puerarin. Substrates that are hydrolysed include cellobiose, ββ-trehalose αα-trehalose, maltose, gentiobiose, isomaltose, turanose, palatinose, sucrose, laminaribiose, sinigrin, salicin. Since sucrose has a similar $R_f$ to that of glucose, its hydrolysis was also monitored by NMR spectroscopy (see section 2.7.). For each pair of spots, the first one corresponds to the control and the other to the reaction in the presence of the enzymes and is indicated by the sign +E.

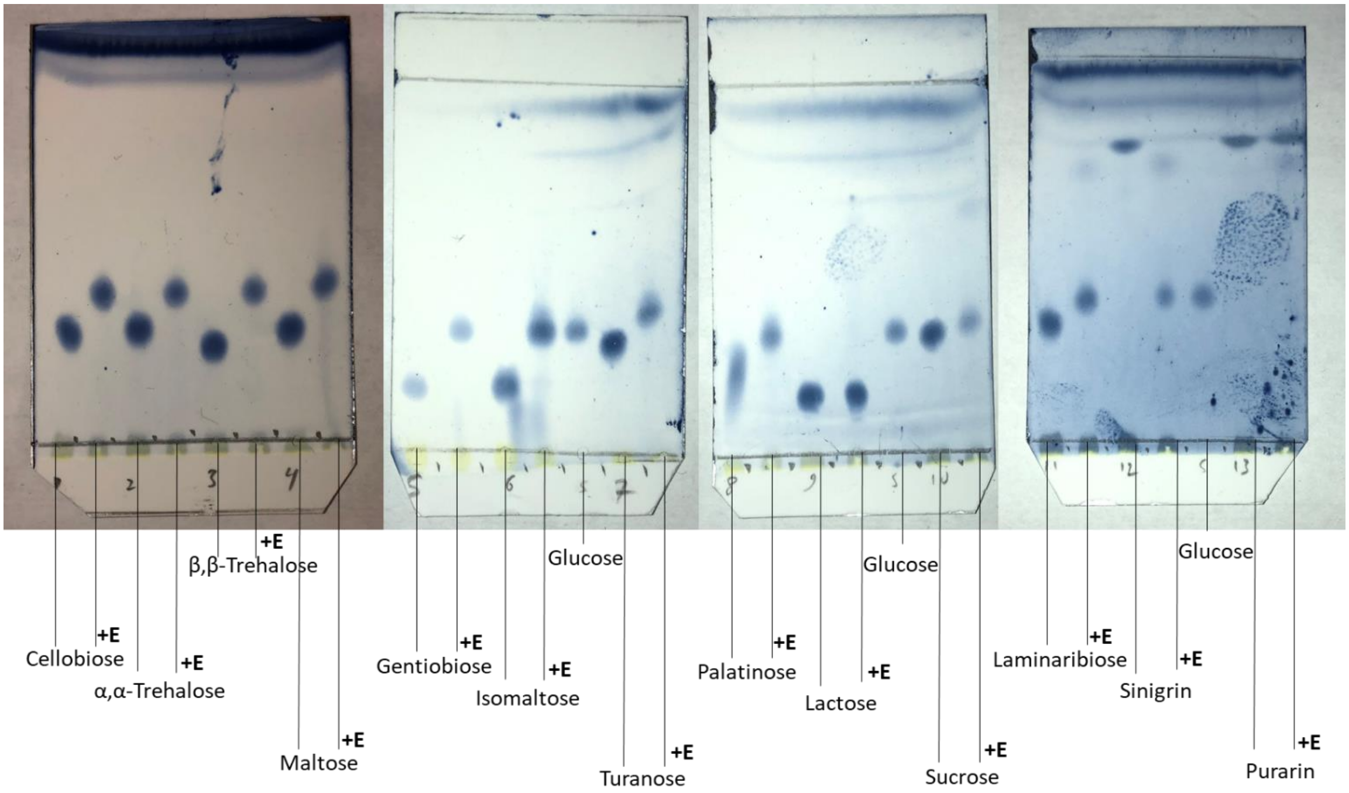

**Figure S67:** TLC traces for hydrolysis of saccharide substrates by AL1-4 - 1

Hydrolysis of Salicin and the fluorogenic substrates can also be visualized using a UV lamp:

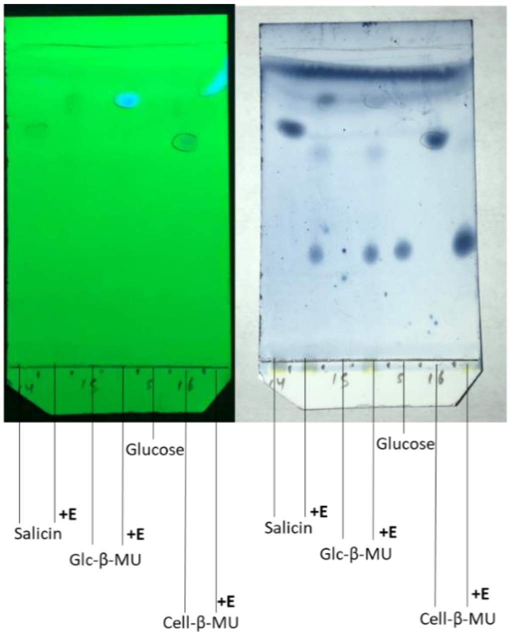

**Figure S68:** TLC traces for hydrolysis of saccharide substrates by AL1-4 - 2

Notably, longer oligosaccharides of various linkage are also accepted as substrates and hydrolysed:

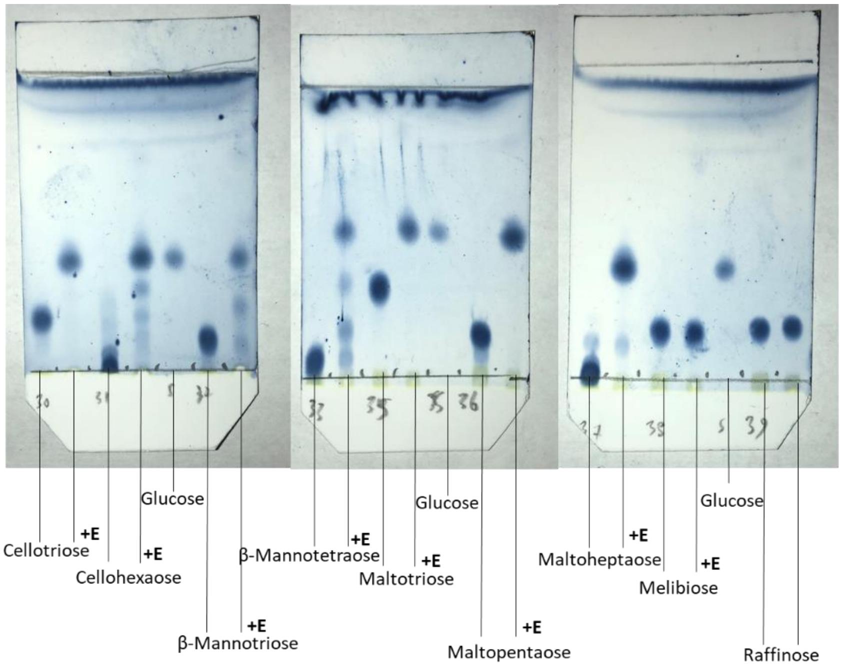

**Figure S69:** TLC traces for hydrolysis of saccharide substrates by AL1-4 - 3

Interestingly, the accepted substrates include acarbose, which has a pseudo-glycosidic bond. Note that here the product of the reaction will be the aromatic compound 4-(Hydroxymethyl)-1,2,3-benzenetriol. This compound was not detected since it is prone to auto-oxidation in the reaction conditions. The sample turns brown in colour (Fig S70), similar to aqueous solutions of pyrogallol[29] (benzene-1,2,3-triol). Also note that the resulting tri-sacharide will not be further hydrolyzed by these enzymes due to presence of the 4-amino group (see SI section 2.6.1.3.3. Fig S62):

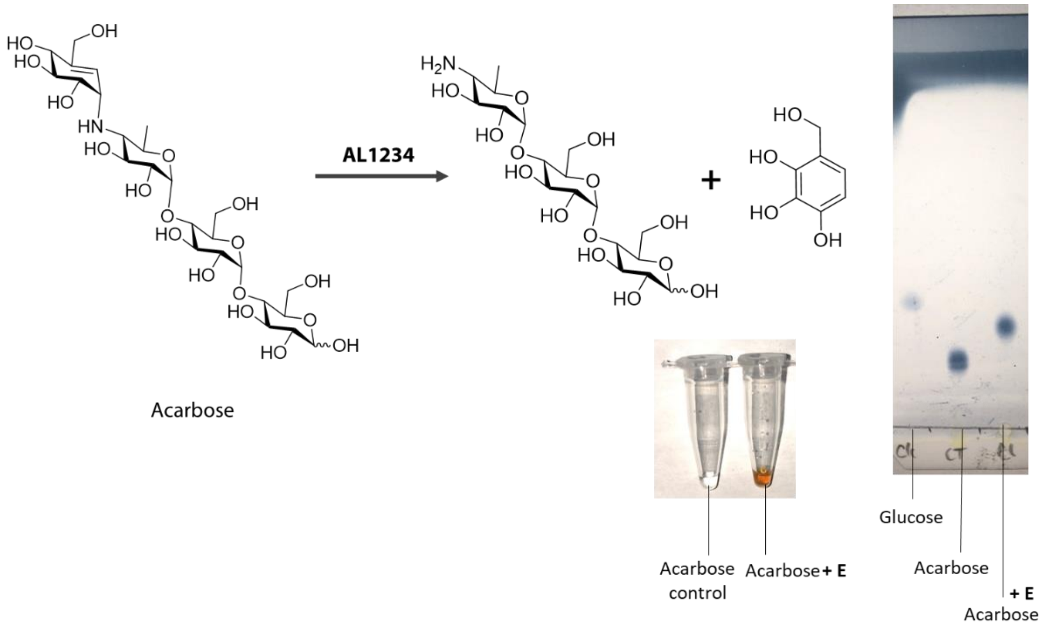

**Figure S70:** TLC trace for breakdown of acarbose by AL1-4 as well as the reaction scheme showing the initial products of reaction

No hydrolysis however is observed for amino-glycoside antibiotics (as expected, see SI section 2.6.1.3.3.), as well as polysaccharides. The only exception is pullulan, which is the most water soluble among the polysaccharides we tested. This, in addition to the location of these enzymes in the periplasm of the cells, indicates that the native substrates of these enzymes are not polysaccharides, but rather oligo- or disaccharides of various linkage types. As well, this shows that these enzymes are exo acting, as expected.

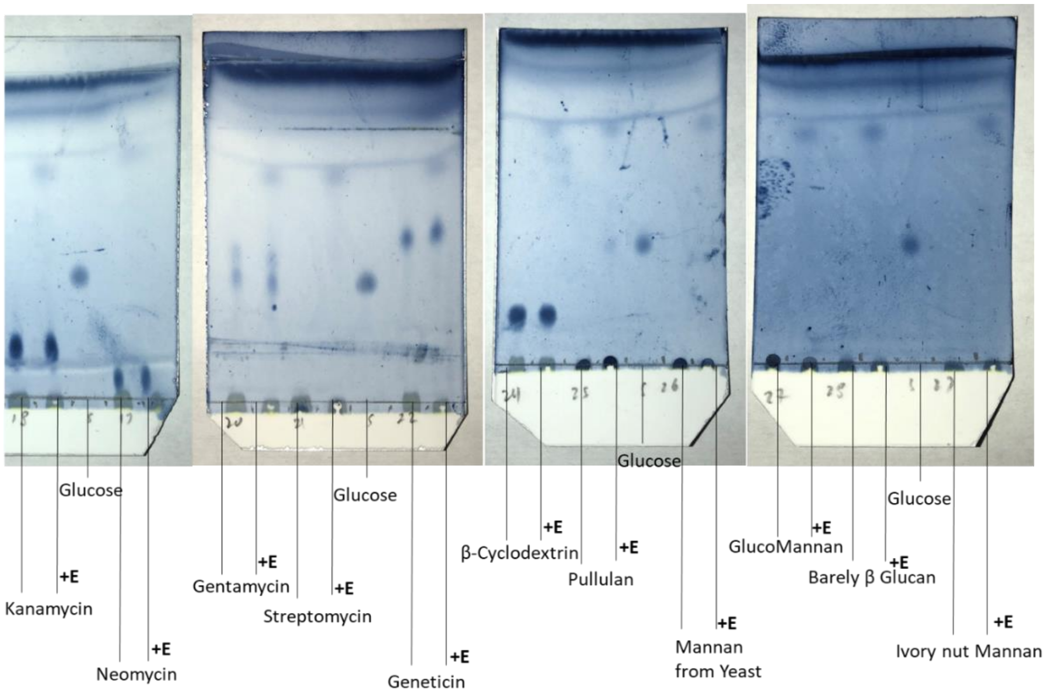

**Figure S71:** TLC traces for hydrolysis of saccharide substrates by AL1-4 - 4

##### 2.6.2.2. Saccharide substrates of BT enzymes

Analysis of reaction of enzymes BT1-4 with the same series of compounds shows very similar results. These enzymes too, hydrolyse all the substrates that are hydrolysed by AL1-4. Hydrolysis of Gentiobiose and acarbose appear to be slower, compared to AL1-4 enzymes.

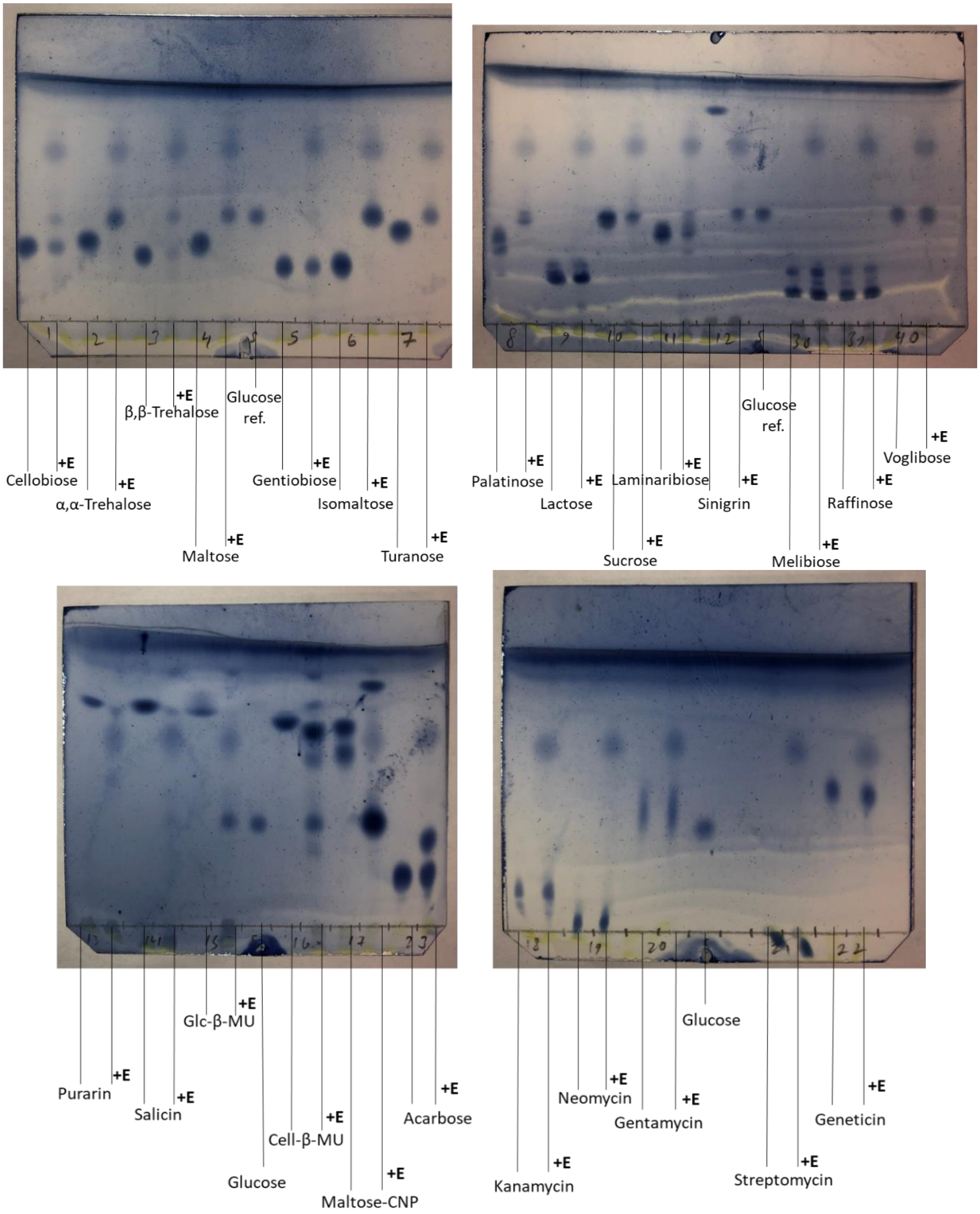

**Figure S72:** TLC traces for hydrolysis of saccharide substrates by BT1-4 – 1

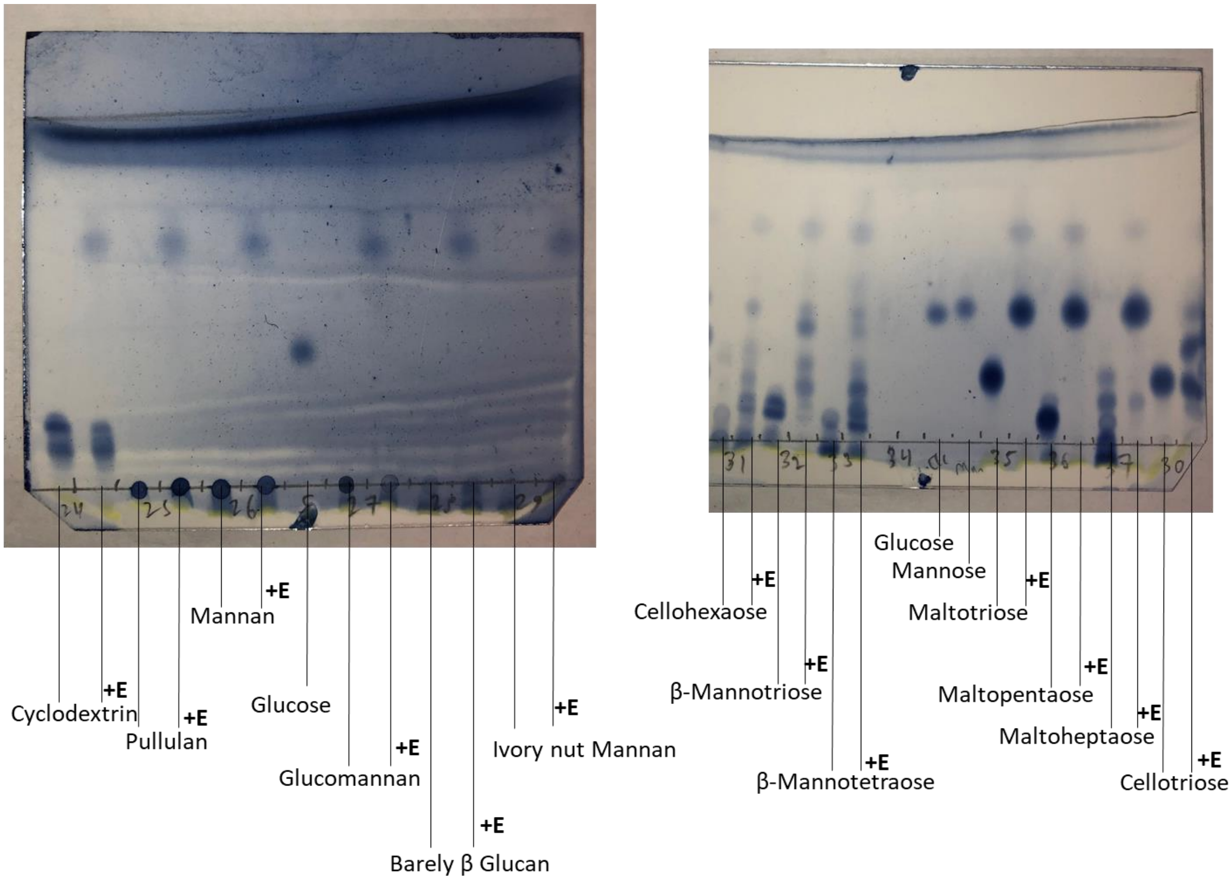

**Figure S73:** TLC traces for hydrolysis of saccharide substrates by BT1-4 - 2

#### 2.7. NMR studies of enzymatic reactions

The NMR experiments were done using a 600 MHz NMR spectrometer equipped with TXI 1.7 mm cryoprobe [Bruker], with solvent suppression using noesygppr1d pulse sequence with a relaxation delay (D1) of 2s, mixing time (D8) of 0.1s, acquisition time of 2s, and with the number of scans ranging from 16 to 64. In a typical experiment, all the reagents except the enzyme(s) were mixed in the buffer with appropriate concentrations (5 mM of each substrate unless otherwise noted) and transferred to the NMR tube and this was used to optimize the NMR acquisition conditions (lock, tuning and matching, shimming). The sample was then removed, mixed with the enzyme(s) (2.5 µM unless otherwise stated) and transferred back into the machine to acquire the spectra. Typically, the first data were collected about 2 minutes after the enzymes were introduced to the sample and the additional data were acquired every 2 to 10 minutes for the rest of the experiment time and also measured after overnight incubation. All NMR assays were performed at the room temperature. The buffer for these experiments was the same as that used in kinetic experiments, 25 mM NaPi buffer pH 7.0, which was lyophilized and reconstituted in $D_2O$, with the appropriate metals (except Mn(II)[30]) added for each respective sample prior to experiments. The details for each experiment can be found below.

Note that for the 3-keto glycoside substrates in aqueous solutions, two separate sets of peaks might be observed corresponding to keto and hydrate isomers of the compounds. The equilibrium is established within a few minutes from when the compound is dissolved in aqueous buffer, but acquiring the spectra quickly allows the transformation of one set of peaks to the other to be observed. Figure S74 below shows one such example, in the downfield section of the spectra for compound 3KGlcβMU where the two set of peaks are most readily observed, as for example the peaks for anomeric proton at around 5.2 ppm. For 3K-α-glycosides however, only one set of peaks corresponding to the keto form is observable.

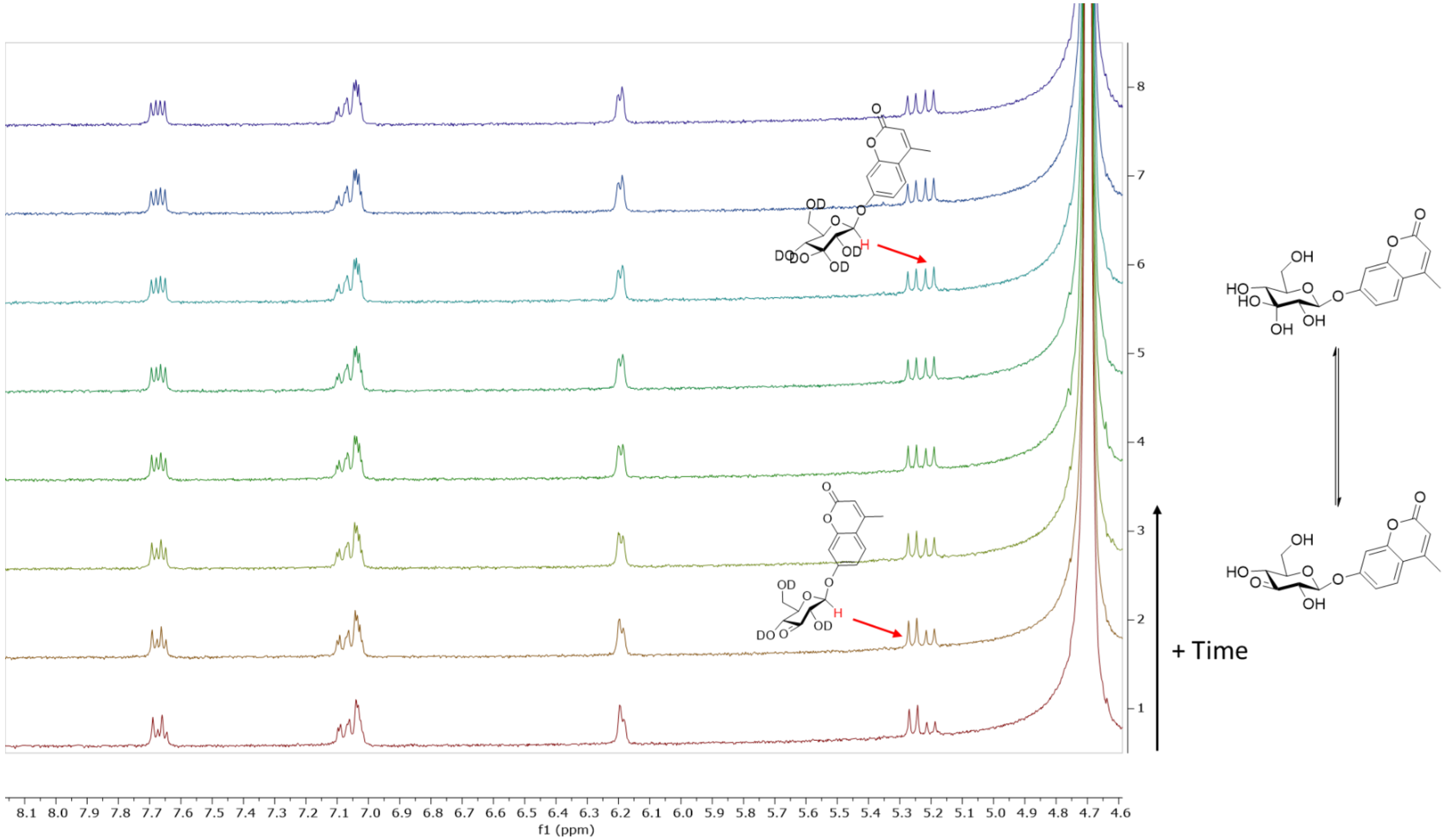

**Figure S74:** H-NMR of keto-hydrate isomerism of 3K-Glc-β-MU

###### 2.7.1. Spontaneous degradation of 3KGlcαpNP

A solution containing 5 mM of 3KGlcαpNP in the reaction buffer was monitored by measuring the H-NMR spectra over the span of a few days (Figure S75). The spontaneous degradation products are the enone, 3-keto-2-hydroxy-glucal and pNP.

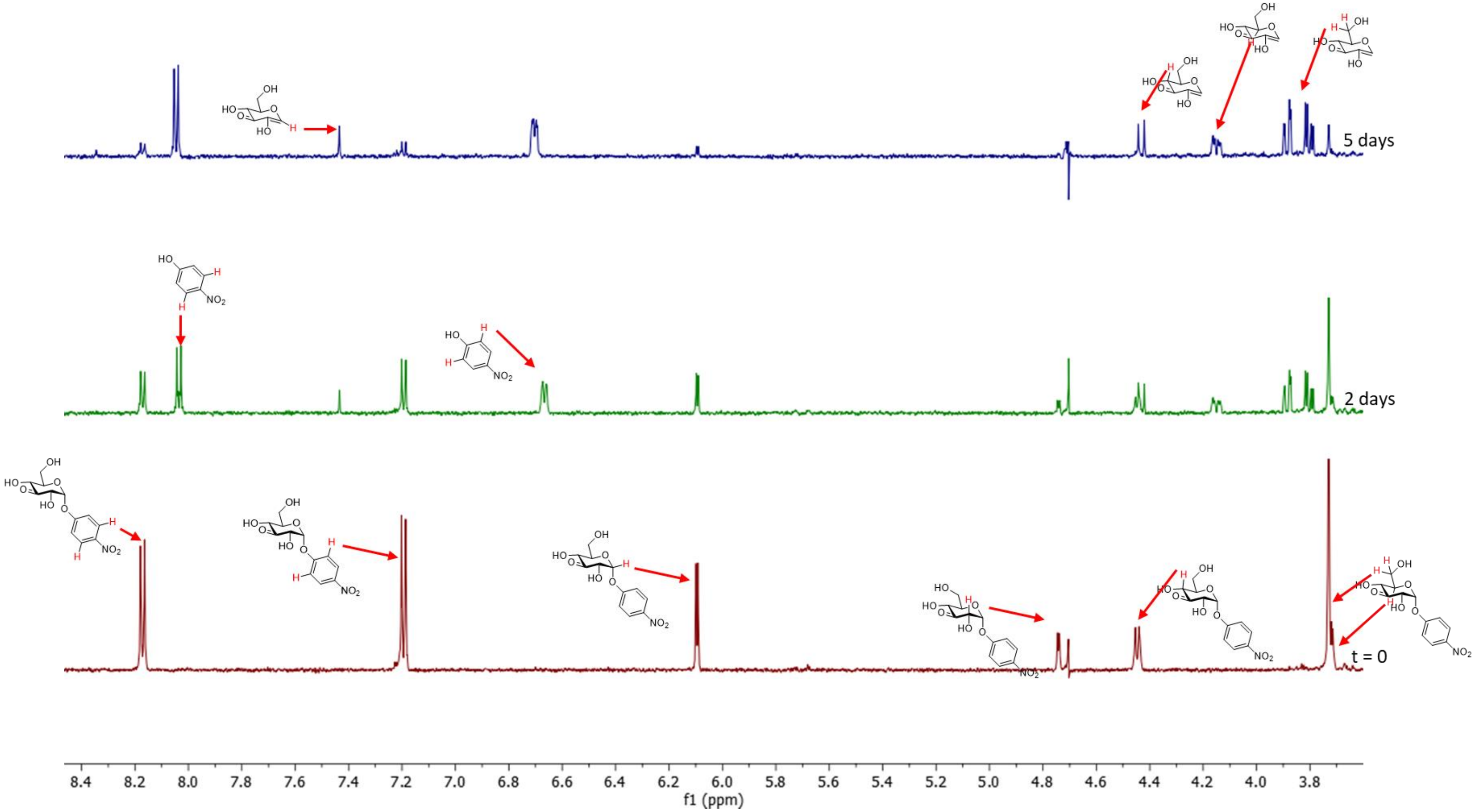

**Figure S75:** H-NMR showing the spontaneous degradation of 3KGlcαpNP

###### 2.7.2. NMR characterization of BT1

To a solution containing 5 mM of 3KGlc-β-OMe in the reaction buffer was added 2.5 µM of BT1 and the reaction was followed by measuring the H-NMR spectra of the reaction mixture. The initial product is 3-keto-2-hydroxy-glucal which is next slowly hydrated by BT1 to form 3-keto-glucose. This product is observable only after incubation for long periods of time. 3-Keto-glucose has been synthesized before and in aqueous solution, it exists as at least ten different isomers, the most prominent of which is the 3-keto-furanose isomer shown below[28]. As well, it slowly decomposes in solution, which in addition to its many isomers is why the signals in the spectra after overnight incubation appear to have lost some intensity. Compared to BT2 (see below), BT1 is much faster in catalysing the elimination from 3-keto-β-glucosides, but significantly slower in catalysing the hydration, which is only observable after overnight incubation.

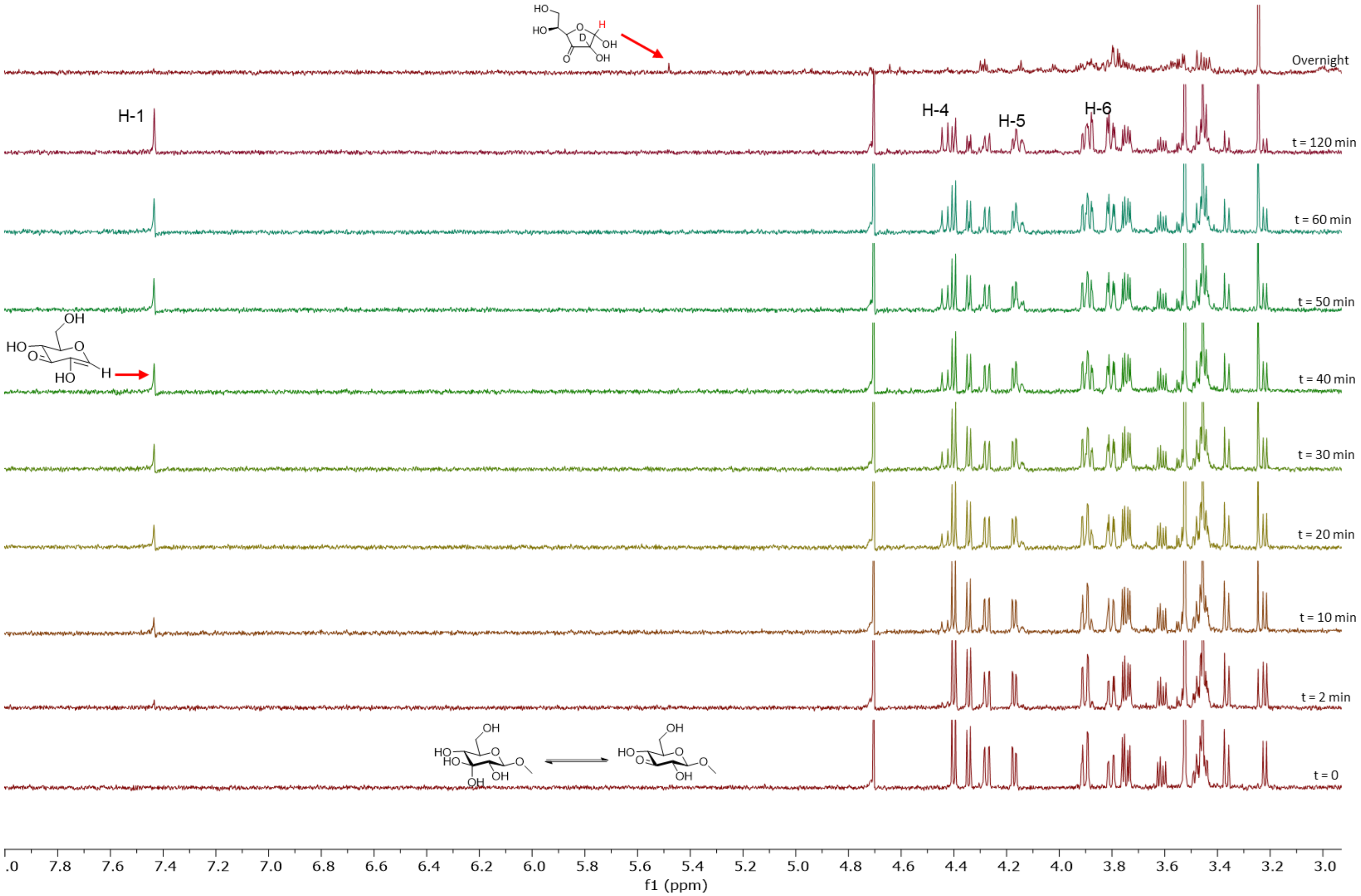

**Figure S76:** H-NMR showing BT1 catalyzed hydrolysis of 3KGlcβOMe

##### 2.7.3. NMR characterization of BT2

To a solution containing 5 mM of 3KGlcαpNP in the reaction buffer was added 2.5 μM of BT2 and the reaction was followed by measuring the H-NMR spectra of the reaction mixture every 2 minutes (Figure S77). The initial product is 3-keto-2-hydroxy-glucal which is next hydrated by BT2 to form 3-keto-glucose, most readily observable as the keto-furanose isomer shown below.

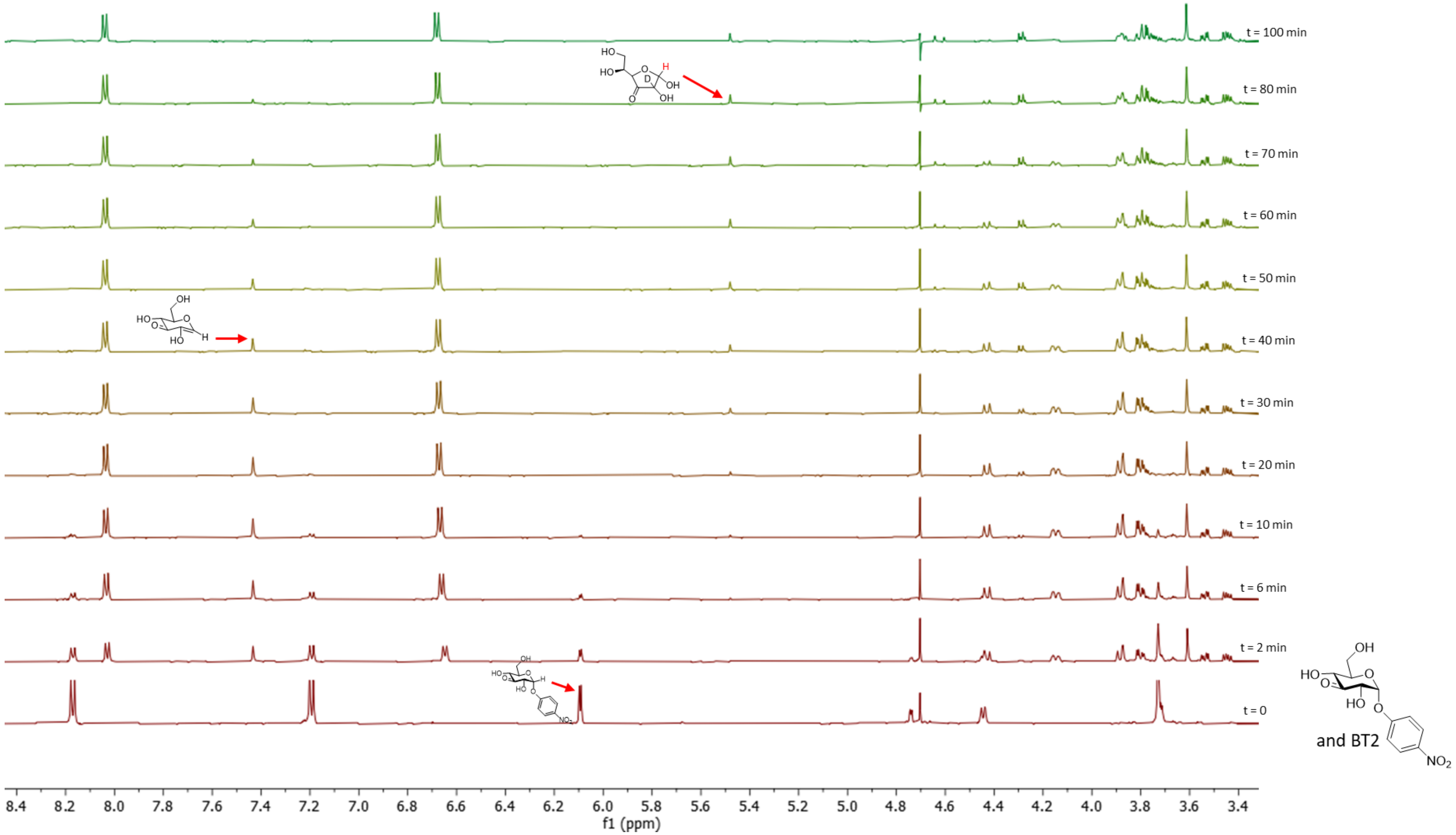

**Figure S77:** H-NMR showing BT2 catalyzed hydrolysis of 3KGlcαpNP

To a solution containing 5 mM of 3KGlc-α-OMe in the reaction buffer was added 2.5 µM of BT2 and the reaction was followed by measuring the H-NMR spectra of the reaction mixture every 2 minutes (Figure S78). The observed product is 3-keto-glucose, since for this substrate with a worse leaving group compared to pNP, the elimination is the rate determining step and the enone intermediate therefore not observable.

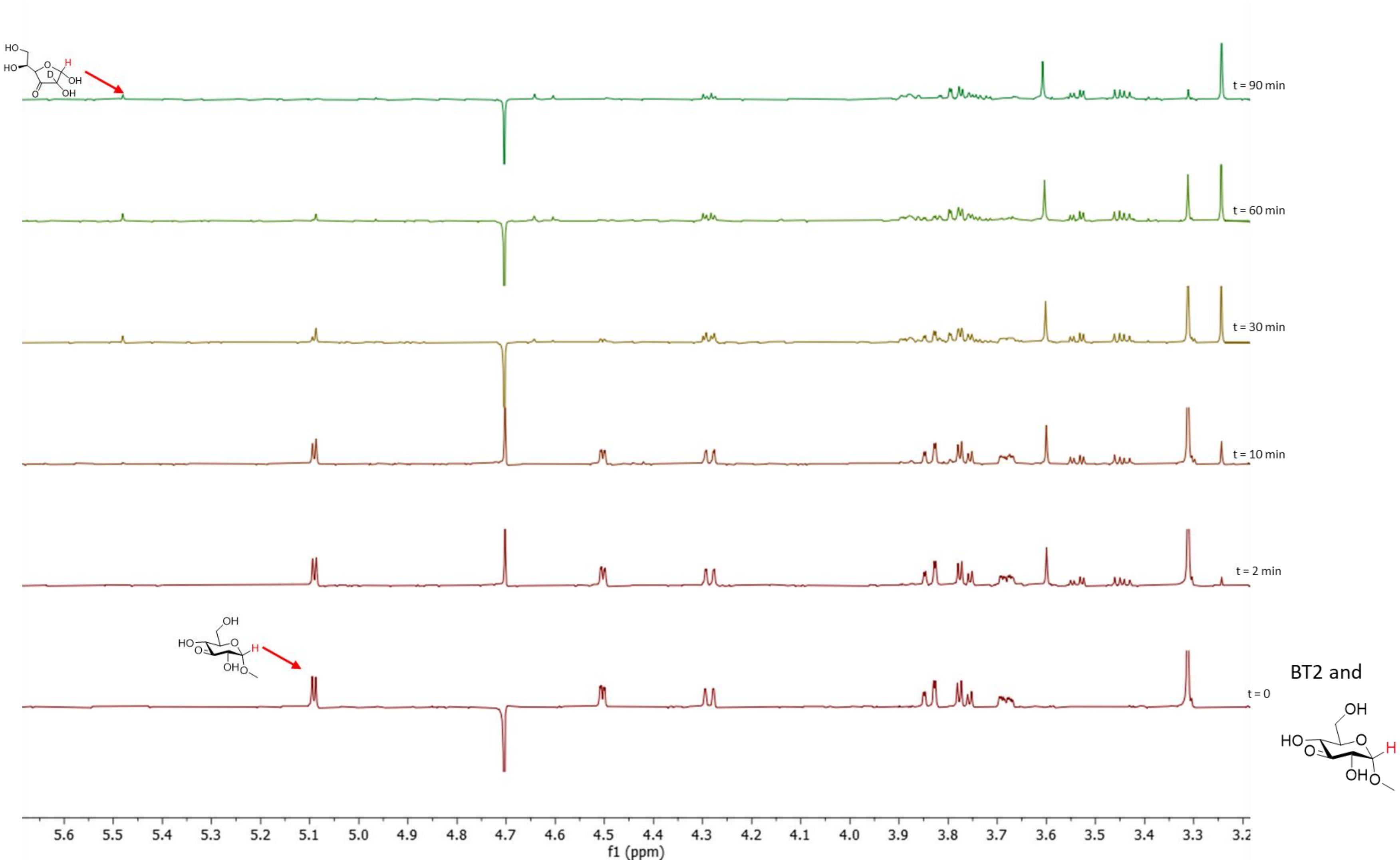

**Figure S78:** H-NMR showing BT2 catalyzed hydrolysis of 3KGlcαOMe

##### 2.7.4. NMR characterization of AL1

To a solution containing 5 mM of 3KGlc-β-OMe in the reaction buffer was added 2.5 µM of AL1 and the reaction was followed by measuring the H-NMR spectra of the reaction mixture every 2 minutes (Figure S79). The product is 3-keto-2-hydroxy-glucal whose H-NMR spectra is reported[31], showing that AL1 catalyses the elimination reaction of H-2 and the aglycone to form the enone.

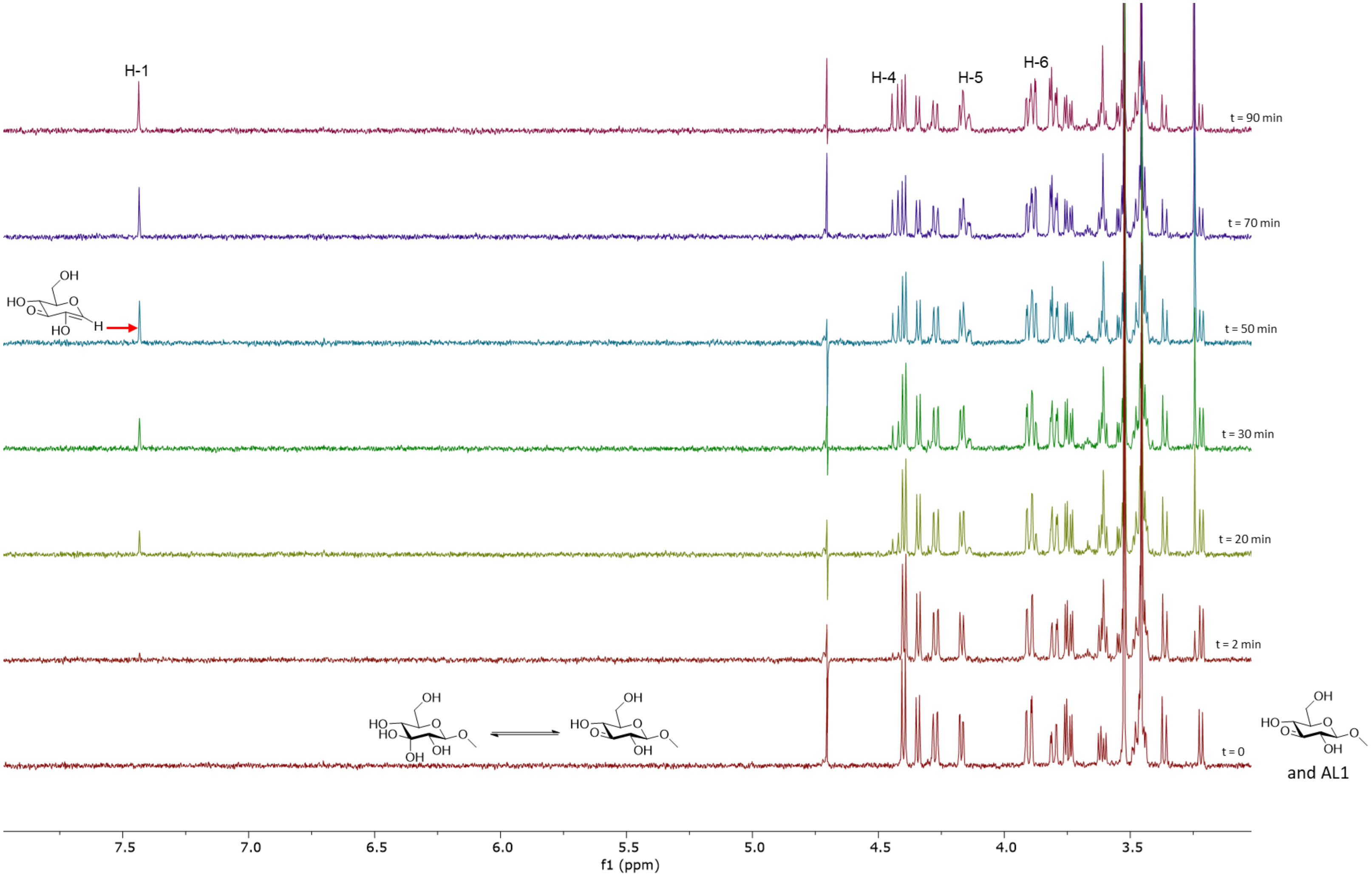

**Figure S79:** H-NMR showing AL1 catalyzed hydrolysis of 3KGlcβOMe

As a control, adding the same amount of enzyme to the solution that contains EDTA (2.5 mM) results in formation of the minimal amount of product even after overnight incubation (Fig S80).

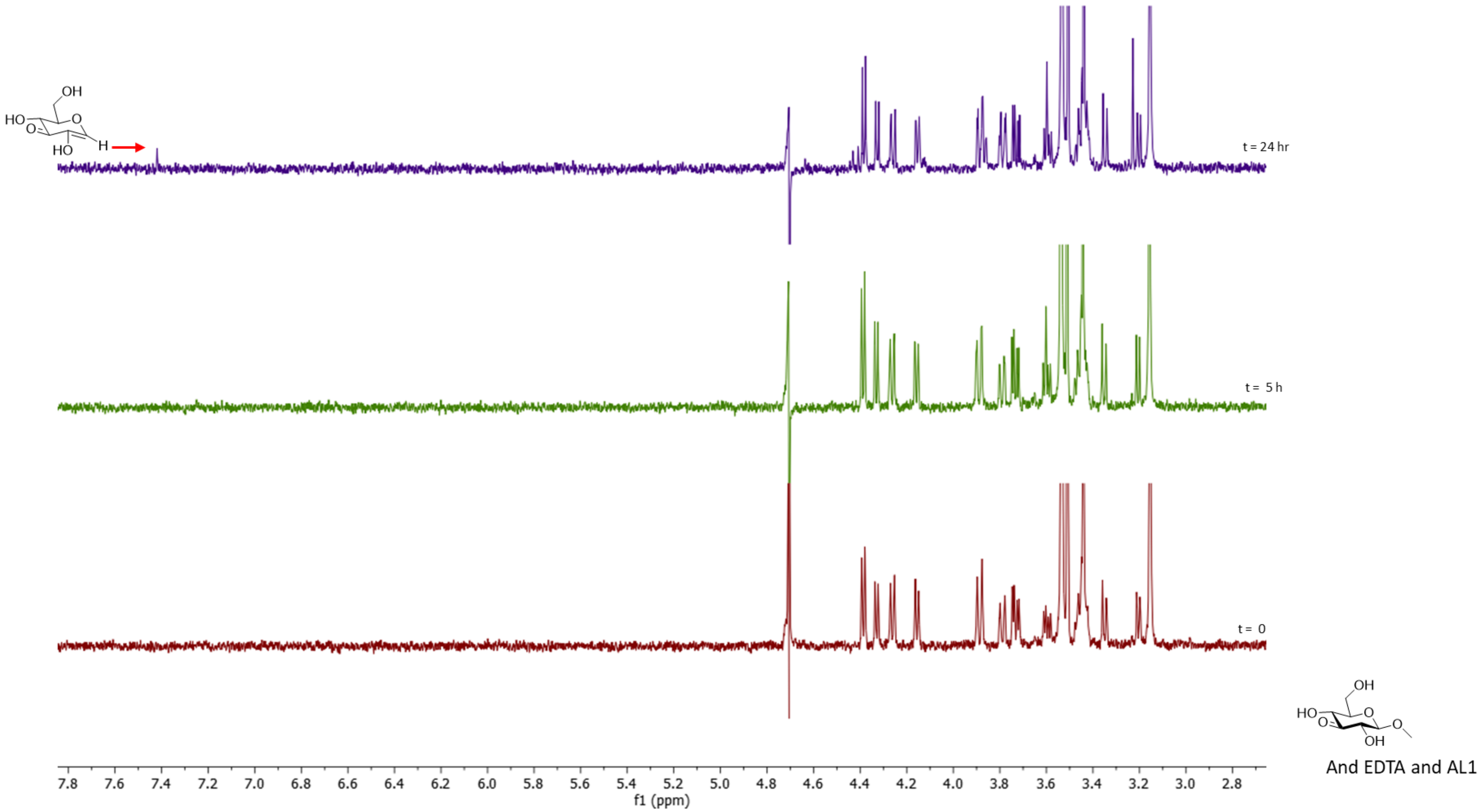

**Figure S80:** H-NMR spectra of the minimal amount of product formed from AL1 catalyzed hydrolysis of 3KGlcβOMe in presence of EDTA

Similar to the case of BT1 (Fig S76), incubation of the 3-keto-2-hydroxy-glucal with AL1 for longer periods of time results in formation of the hydrated product, 3-keto-glucose (Fig S81).

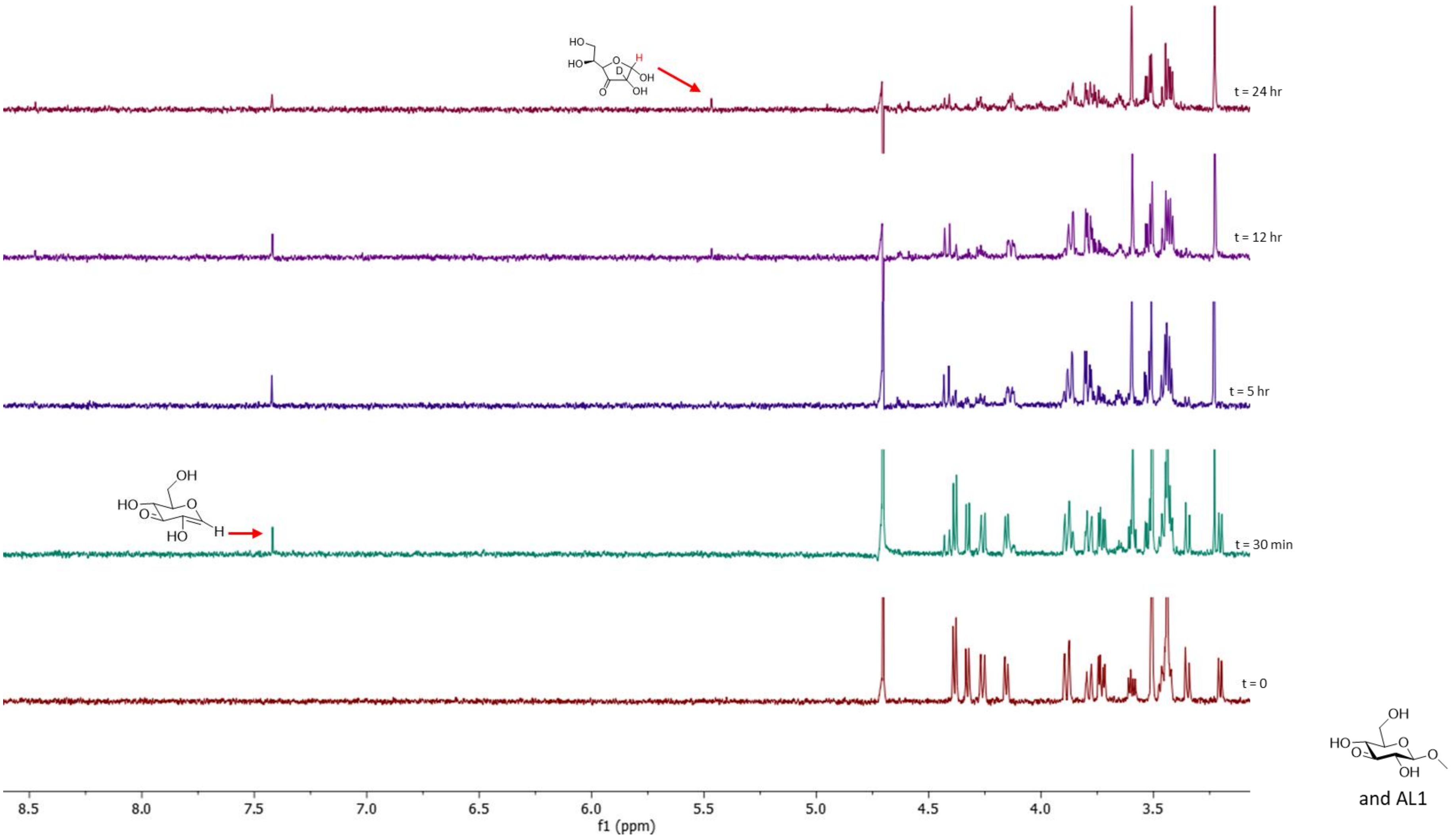

**Figure S81:** H-NMR showing AL1 catalyzed hydrolysis of 3KGlcβOMe and subsequent hydration of this compound upon longer incubation

##### 2.7.5. NMR characterization of AL2

To a solution containing 5 mM of 3KGlcαpNP in the reaction buffer was added 2.5 μM of AL2 and the reaction was followed by measuring the H-NMR spectra of the reaction mixture every 2 minutes (Figure S80). The initial product is 3-keto-2-hydroxy-glucal, showing that AL2 can also catalyse the elimination reaction of H-2 and the aglycone to form the enone with α glycoside substrates. This product however is readily hydrated by AL2 to form 3-keto-glucose. This compound has been synthesized before and in aqueous solution, it exists as at least ten different isomers, the most prominent of which is the keto-furanose compound depicted below[28].

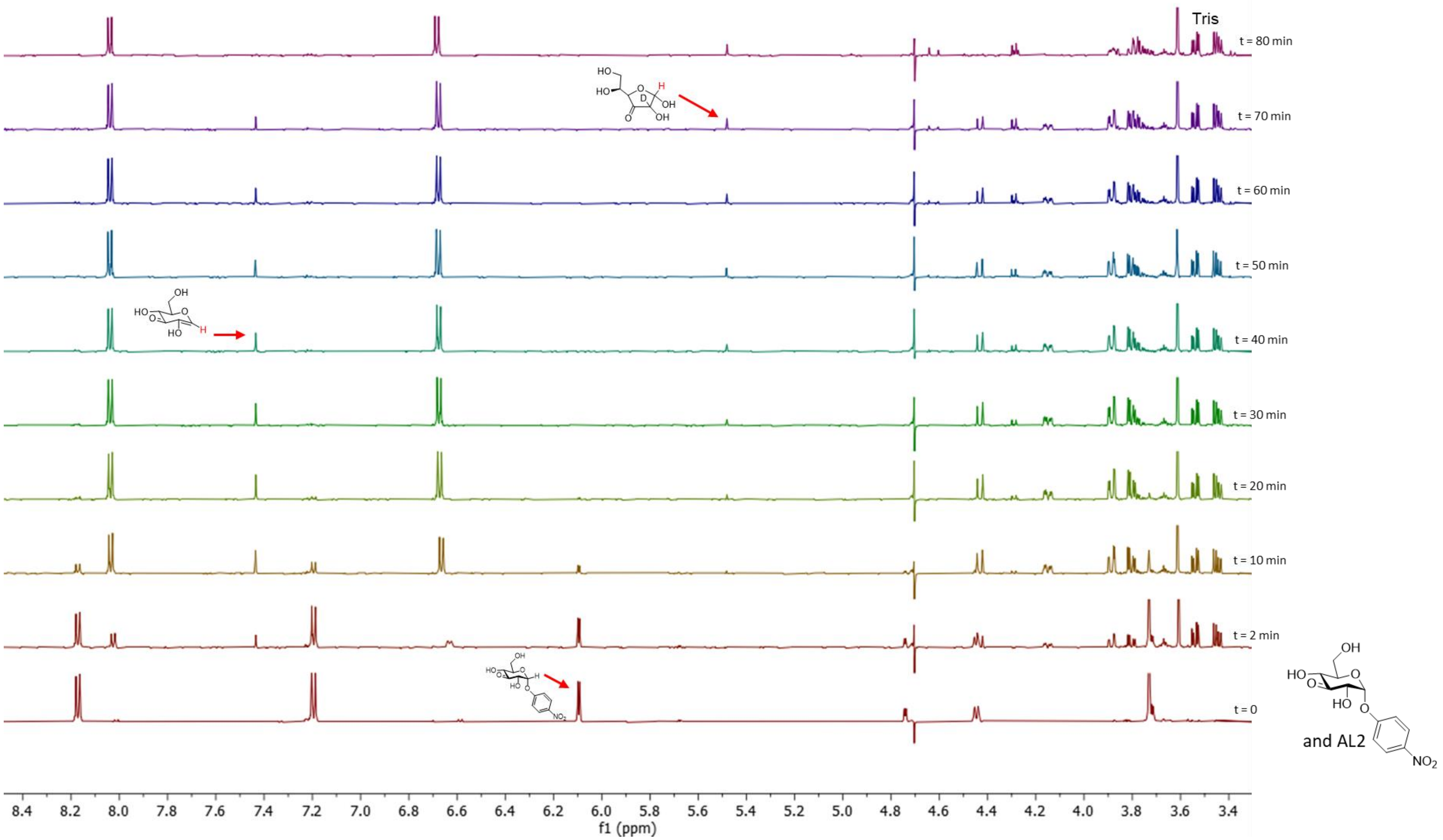

**Figure S82:** H-NMR showing AL2 catalyzed hydrolysis of 3KGlcαpNP

To a solution containing 5 mM of 3KGlc-α-OMe in the reaction buffer was added 2.5 µM of AL2 and the reaction was followed by measuring the H-NMR spectra of the reaction mixture every 2 minutes (Figure S81). The observed product is 3-keto-glucose. Here again, similar to the case for BT2 shown in Fig. S87, since the substrate features a worse leaving group compared to pNP, the elimination is rate determining and the enone intermediate is not observable.

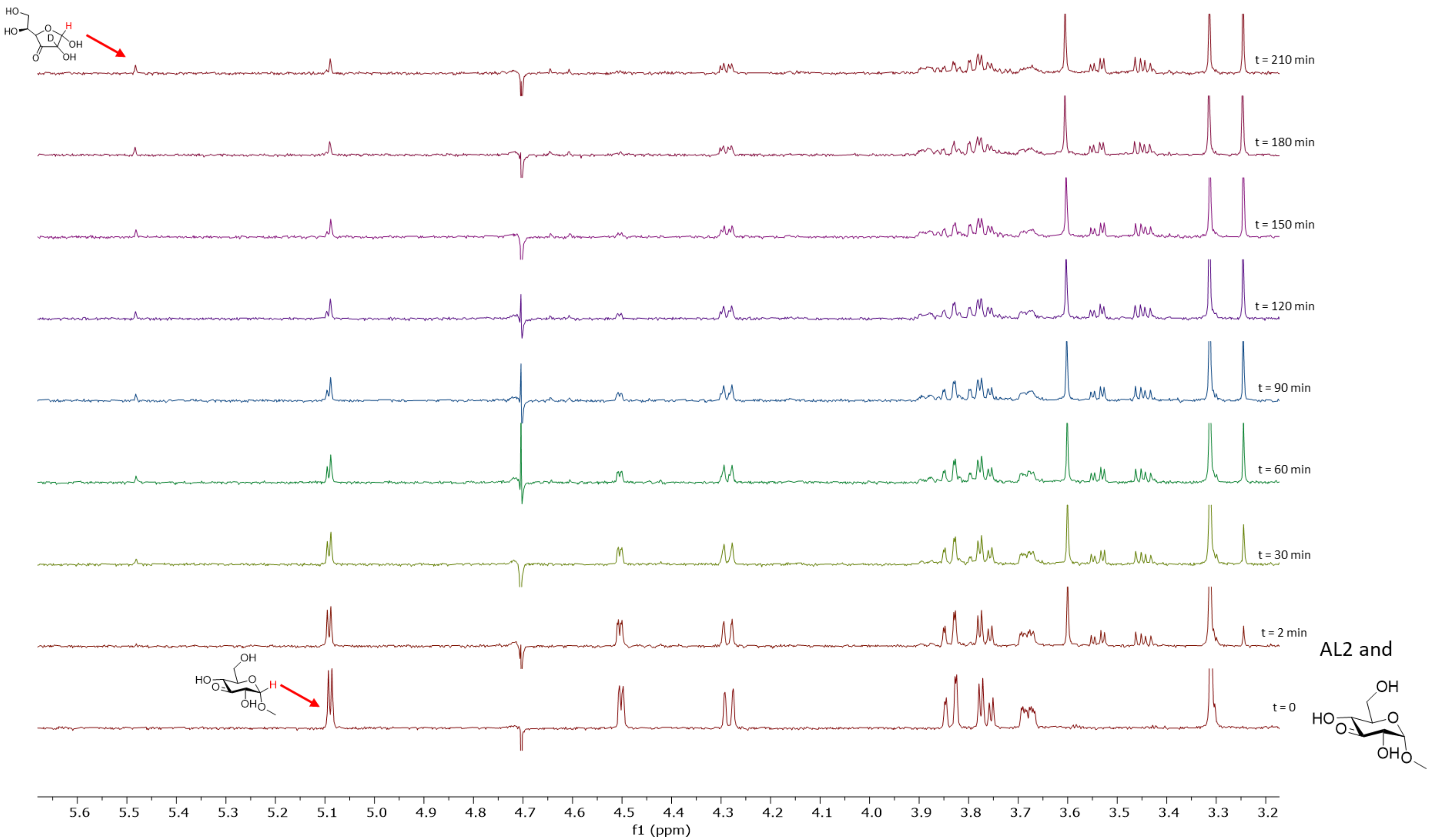

**Figure S83:** H-NMR showing AL2 catalyzed hydrolysis of 3KGlcαOMe

As a control, adding the same amount of enzyme to the solution that contains EDTA (2.5 mM) results in formation of no product even after overnight incubation (Fig S84).

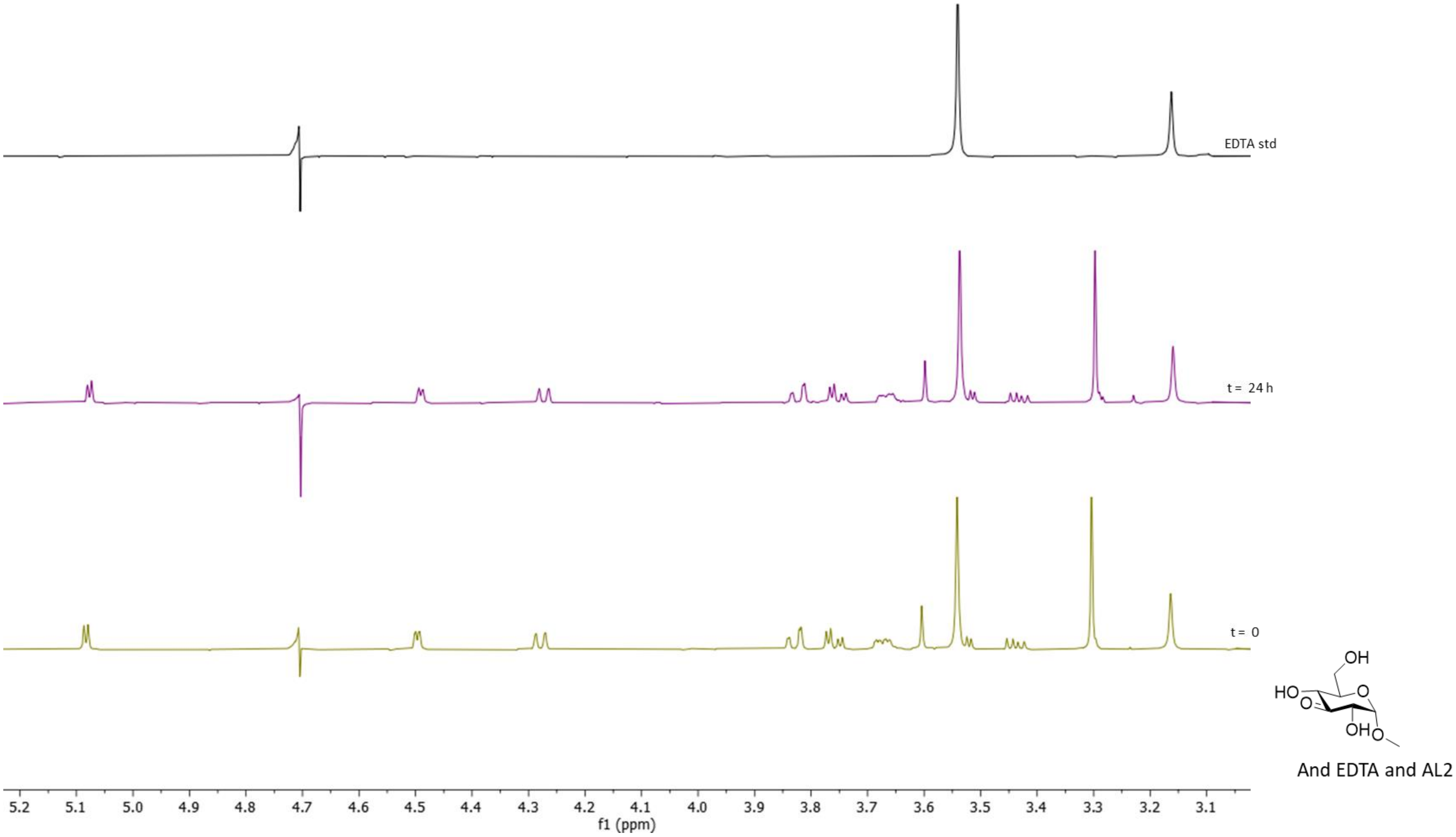

**Figure S84:** H-NMR showing that EDTA reduces the amount of AL2 catalyzed hydrolysis of 3KGlcαOMe beyond the limit of detection

#### 2.7.6. NMR characterization of AL3

To a solution containing 5 mM of 3KGlcH and 5 mM GlcαpNP in the reaction buffer was added 2.5 µM of AL3 and the reaction was followed by measuring the H-NMR spectra of the reaction mixture every 2 minutes (Figure S85). Comparison of the resulting spectra with the control spectra of pure standard compounds shows that AL3 has established an equilibrium between the pair of glycoside and 3-keto glycoside substrates.

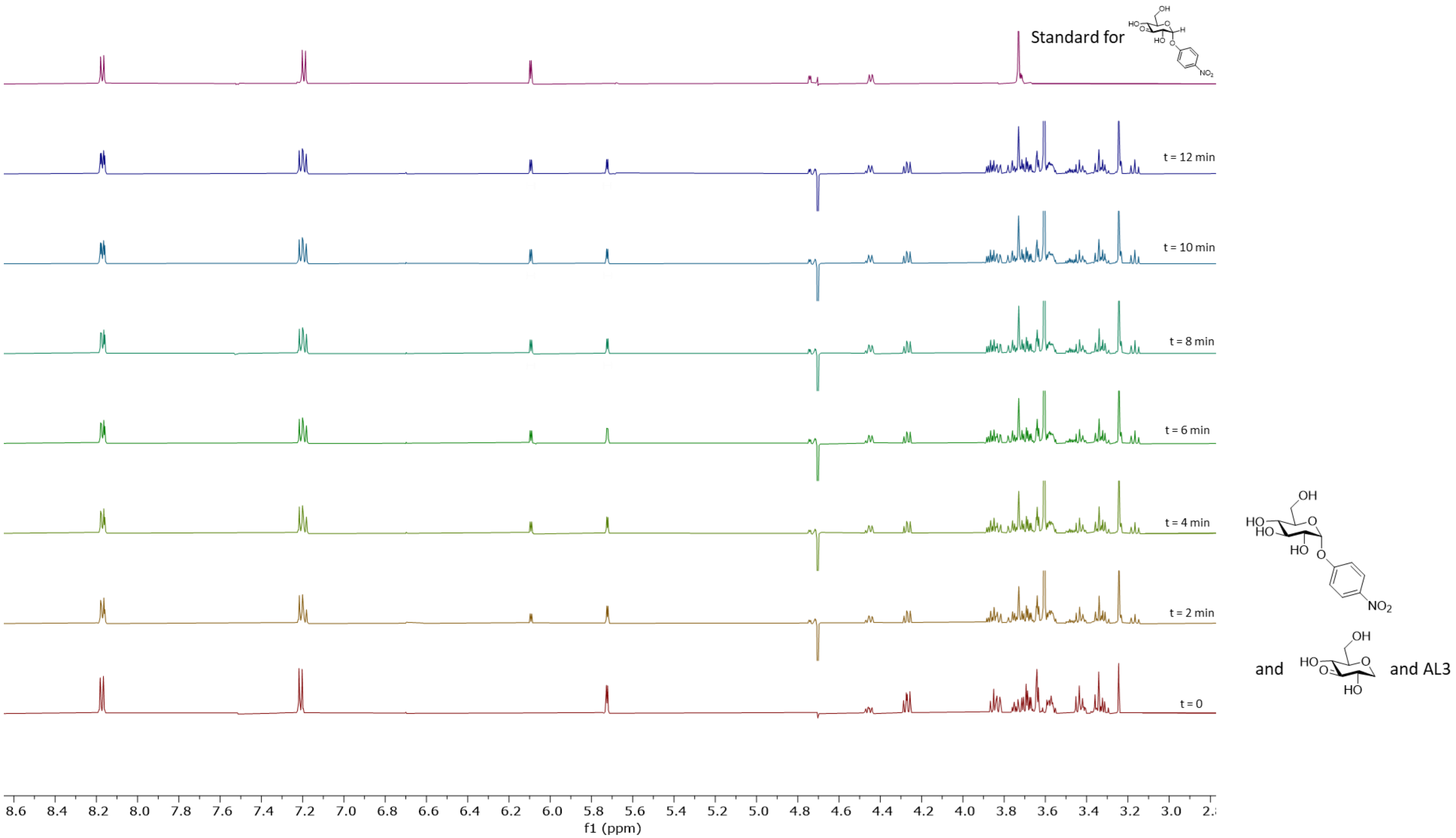

**Figure S85:** H-NMR showing AL3 catalyzed reaction between 3KGlcH and GlcαpNP

Next, to a solution containing 5 mM of 1,5-anhydroglucitol and 5 mM 3KGlcαpNP in the reaction buffer was added 2.5 μM of AL3 and the reaction was followed by measuring the H-NMR spectra of the reaction mixture every 2 minutes (Figure S86). In this case too, comparison of the resulting spectra with the control spectra of pure standard compounds shows that AL3 has established an equilibrium between the pair of glycoside and 3-keto glycoside substrates.

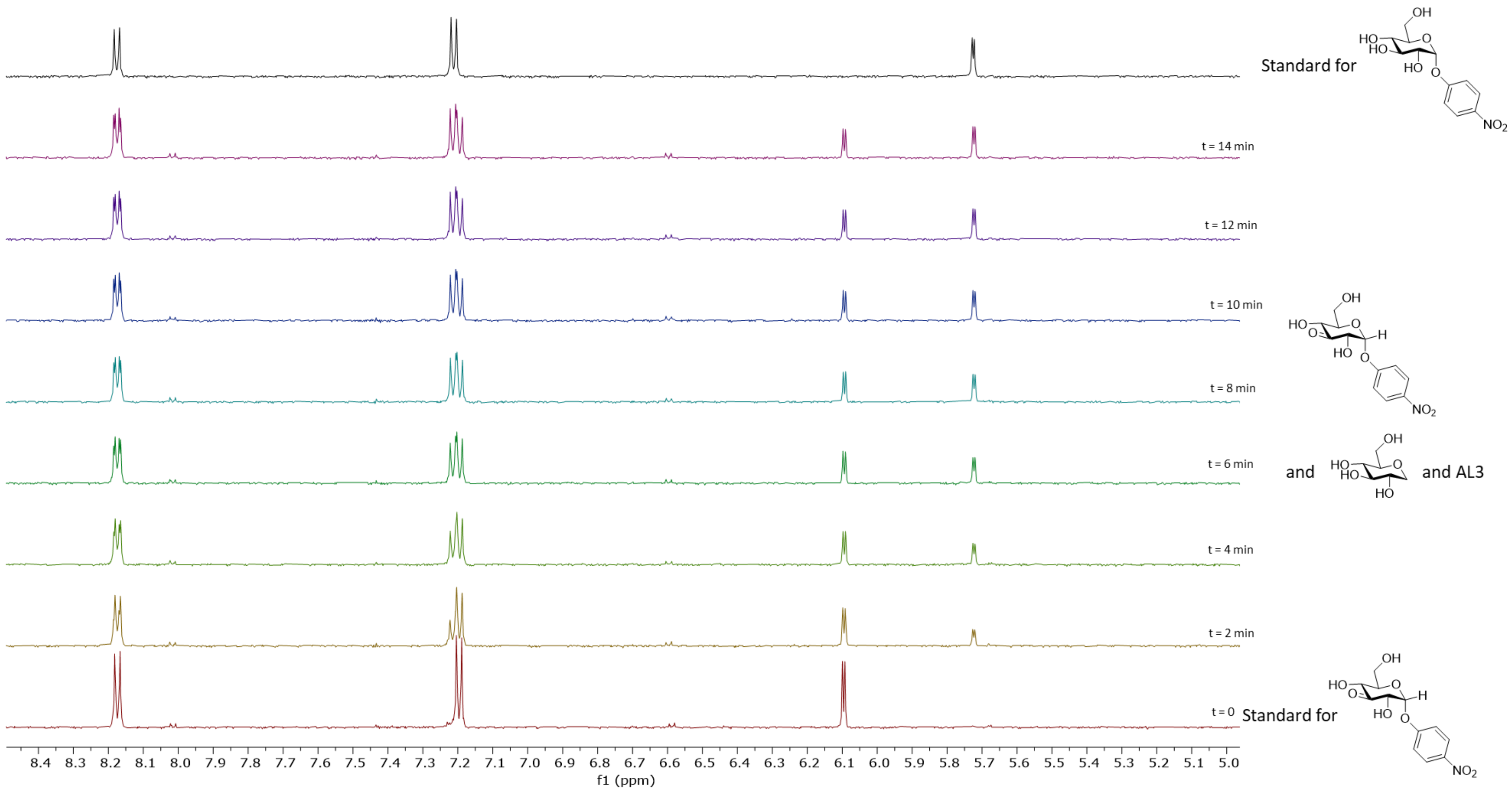

**Figure S86:** H-NMR showing AL3 catalyzed reaction between 1,5-anhydroglucitol and 3KGlcαpNP

Comparing the resulting spectra for both of these reactions shows that as expected, the amount of each compound in the equilibrium is the same.

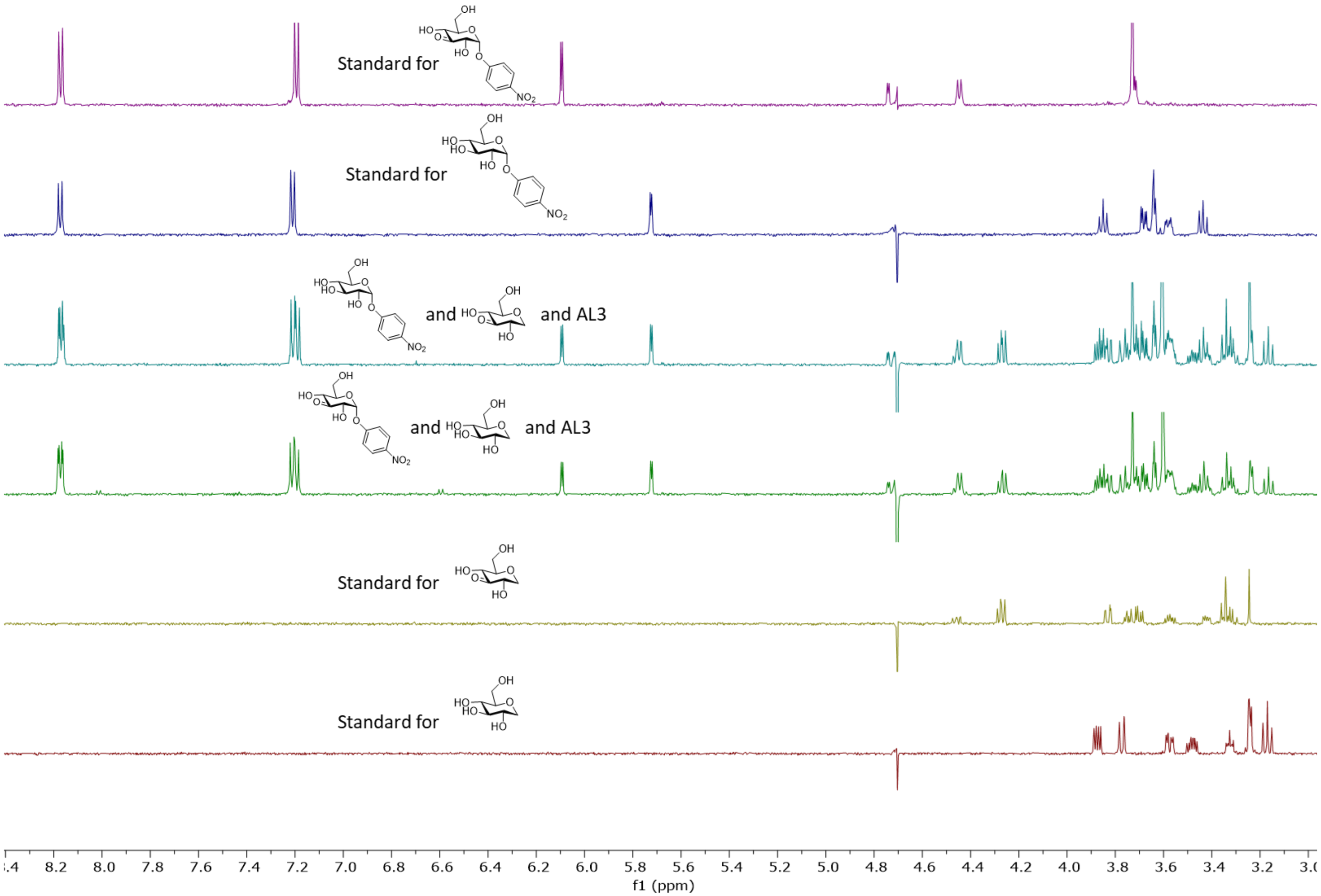

**Figure S87:** H-NMR comparing AL3 catalyzed reaction between 1,5-anhydroglucitol and 3KGlcαpNP and 3KGlcH and GlcαpNP - 1

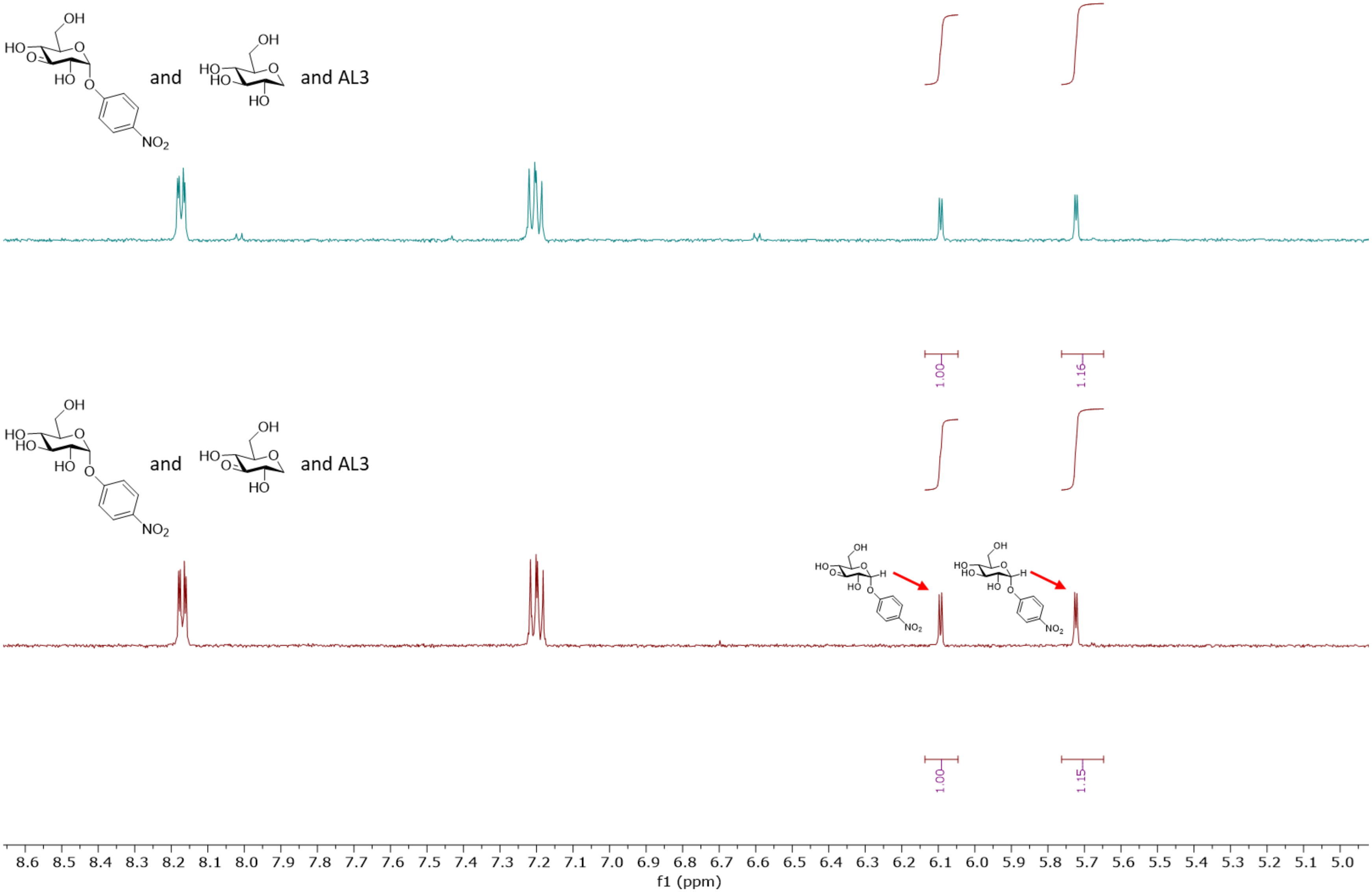

**Figure S88:** H-NMR comparing AL3 catalyzed reaction between 1,5-anhydroglucitol and 3KGlcαpNP and 3KGlcH and GlcαpNP - 2

Next, to a solution containing 5 mM glucose and 5 mM 3KGlcH in the reaction buffer was added 2.5 µM of AL3 and the reaction was followed by measuring the H-NMR spectra of the reaction mixture (Figure S89). In this case, glucose is oxidized to form 3-keto-glucose, which readily isomerises to at least ten different isomers, the most prominent of which is the keto-furanose isomer shown below[28]. Note that in this case, the signal for the anomeric proton of the product is a doublet, unlike the signal for the same compound formed as a result of the reaction catalysed by AL2, where the hydration in $D_2O$ results in deuterium incorporation at the 2 position (See section 2.7.4.).

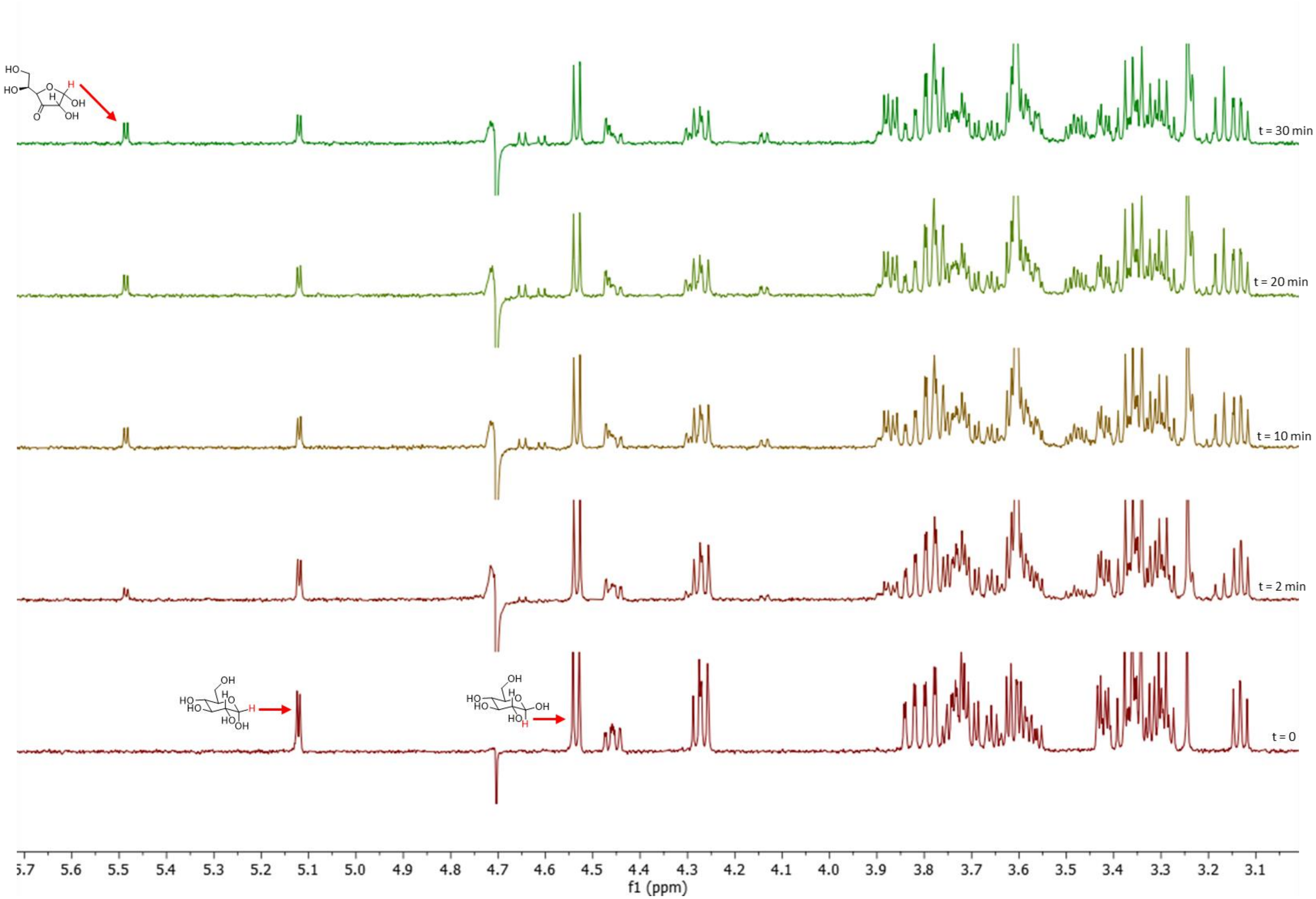

**Figure S89:** H-NMR of AL3 catalyzed reaction between glucose and 3KGlcH

Next, the same reaction was carried out with glucose labelled with C-13 at C1. The C-NMR in this shows formation of the same products, assigned based on the previous report[28] (Figure S90).

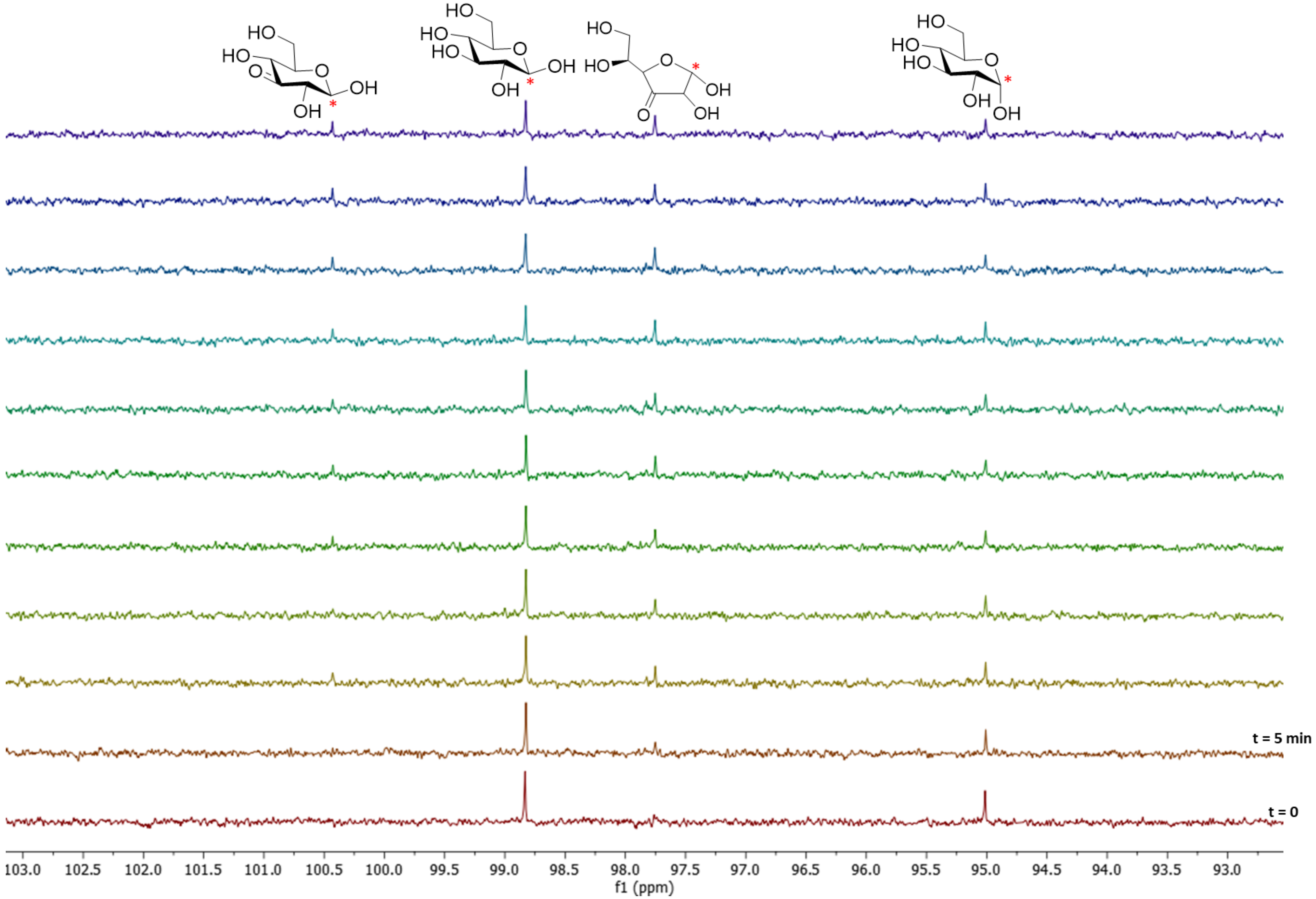

**Figure S90:** C-NMR of AL3 catalyzed reaction between glucose (1-13C) and 3KGlcH

##### 2.7.7. Hydrolysis of GlcαpNP catalysed by AL1, AL2, AL3

To a solution containing 5 mM of GlcαpNP and 5 mM 3KGlcH in the reaction buffer was added 2.5 µM of each of the enzymes AL1, AL2 and AL3 and the reaction was followed by measuring the H-NMR spectra of the reaction mixture (Figure S91).

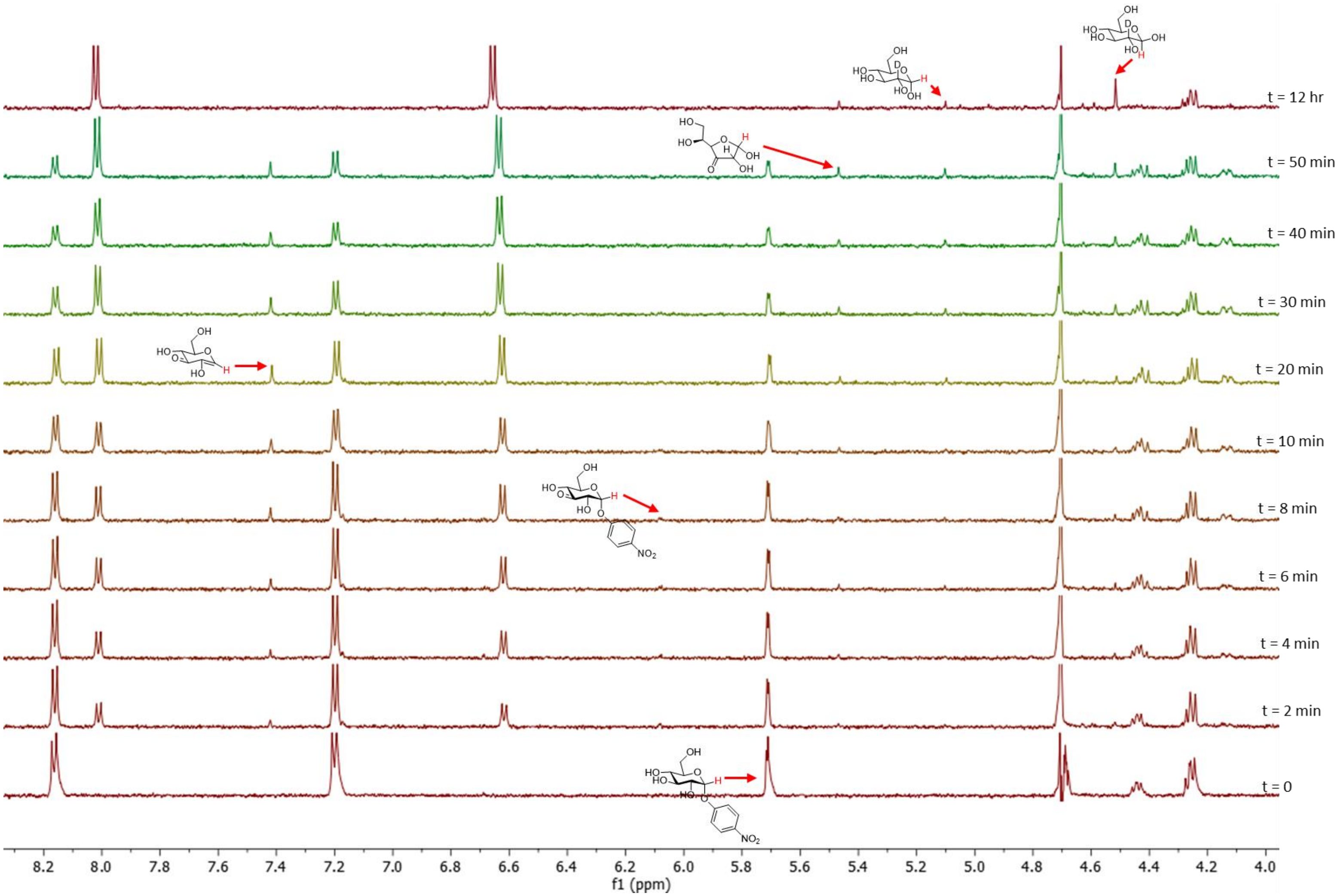

**Figure S91:** H-NMR of AL123 catalyzed hydrolysis of 5 mM of GlcαpNP with 5 mM of 3KGlcH

Next, the same experiment as above was repeated, this time with sub-stoichiometric amounts of the re-oxidant, 2.5 mM 3KGlcH (Figure S92). Note that adding stoichiometric amounts of the re-oxidant is not strictly necessary and the reaction still goes to completion, since the product of this reaction is 3Keto-glucose, which itself can serve as a re-oxidant. However, since most of this compound will occur as the furanose form in solution which is not a useful re-oxidant, only when this reaction is done at high concentrations is there enough of the pyranose form to serve as the re-oxidant. Also note that formation of all intermediates is clearly observed in this case.

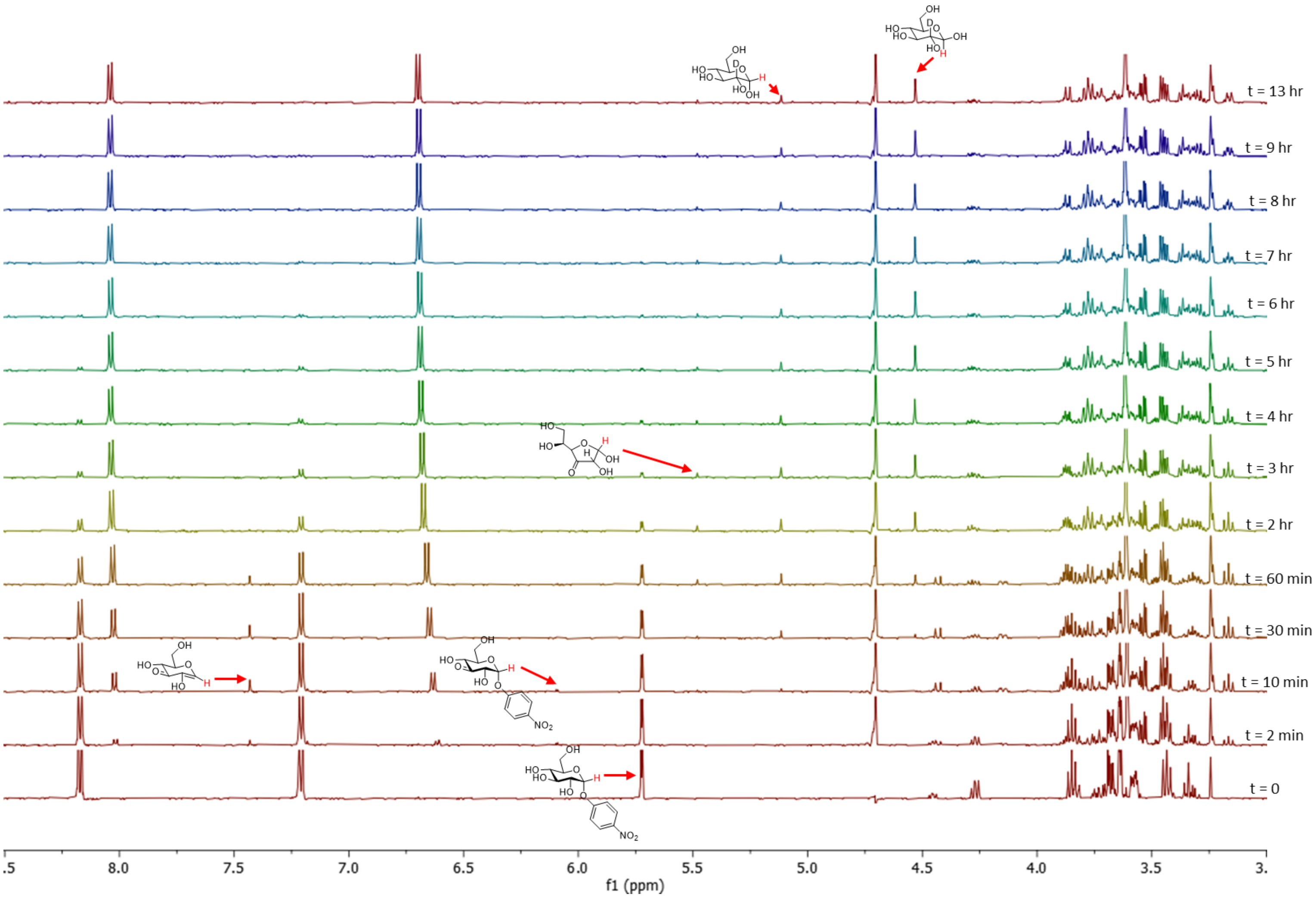

**Figure S92:** H-NMR of AL123 catalyzed hydrolysis of 5 mM of GlcαpNP with 2.5 mM of 3KGlcH

As negative controls, addition of either AL1 or AL2 to the same mixture does not result in any reaction.

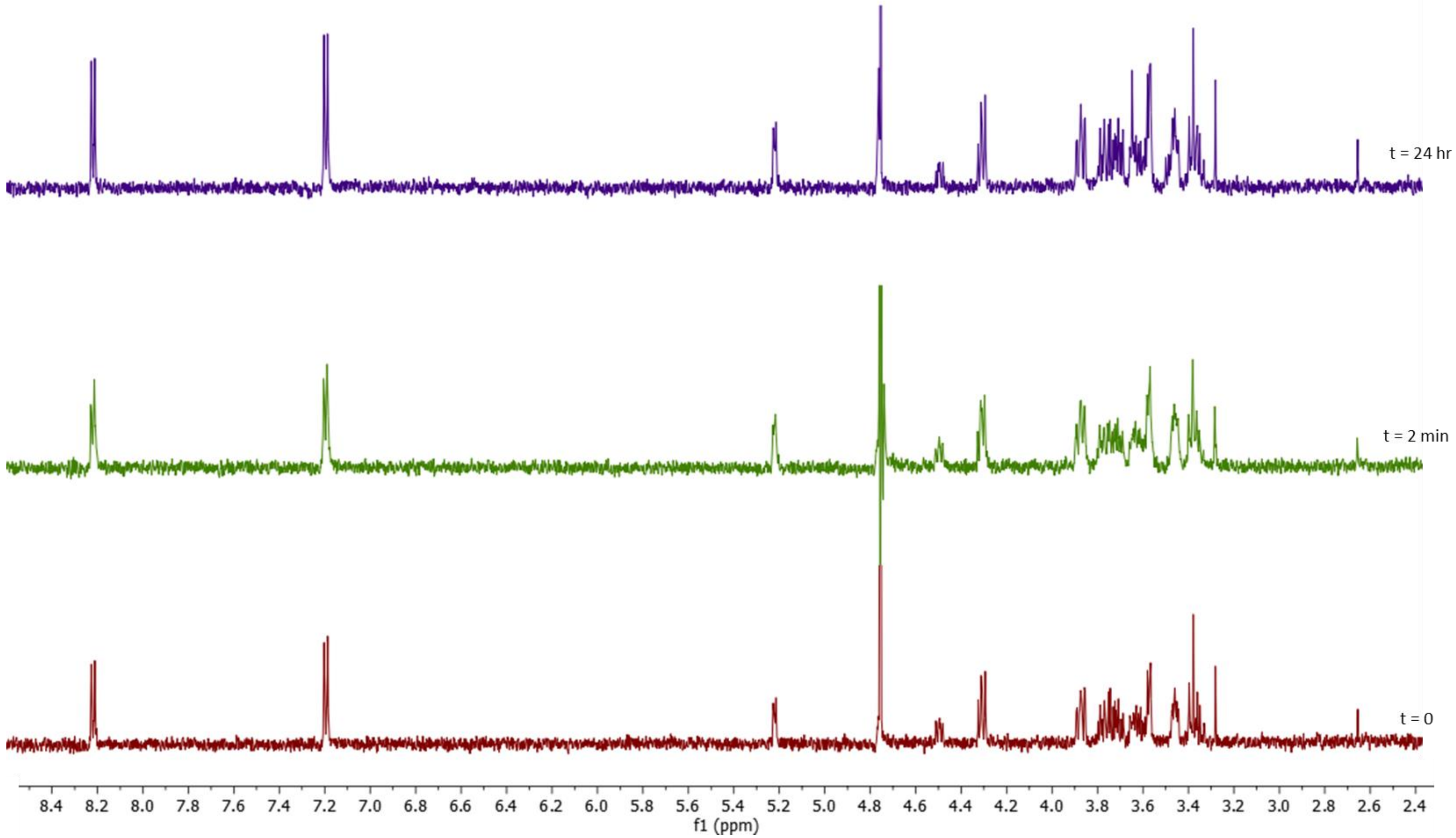

**Figure S93**: Control experiment, showing that AL1 alone does not catalyze the hydrolysis of GlcαpNP

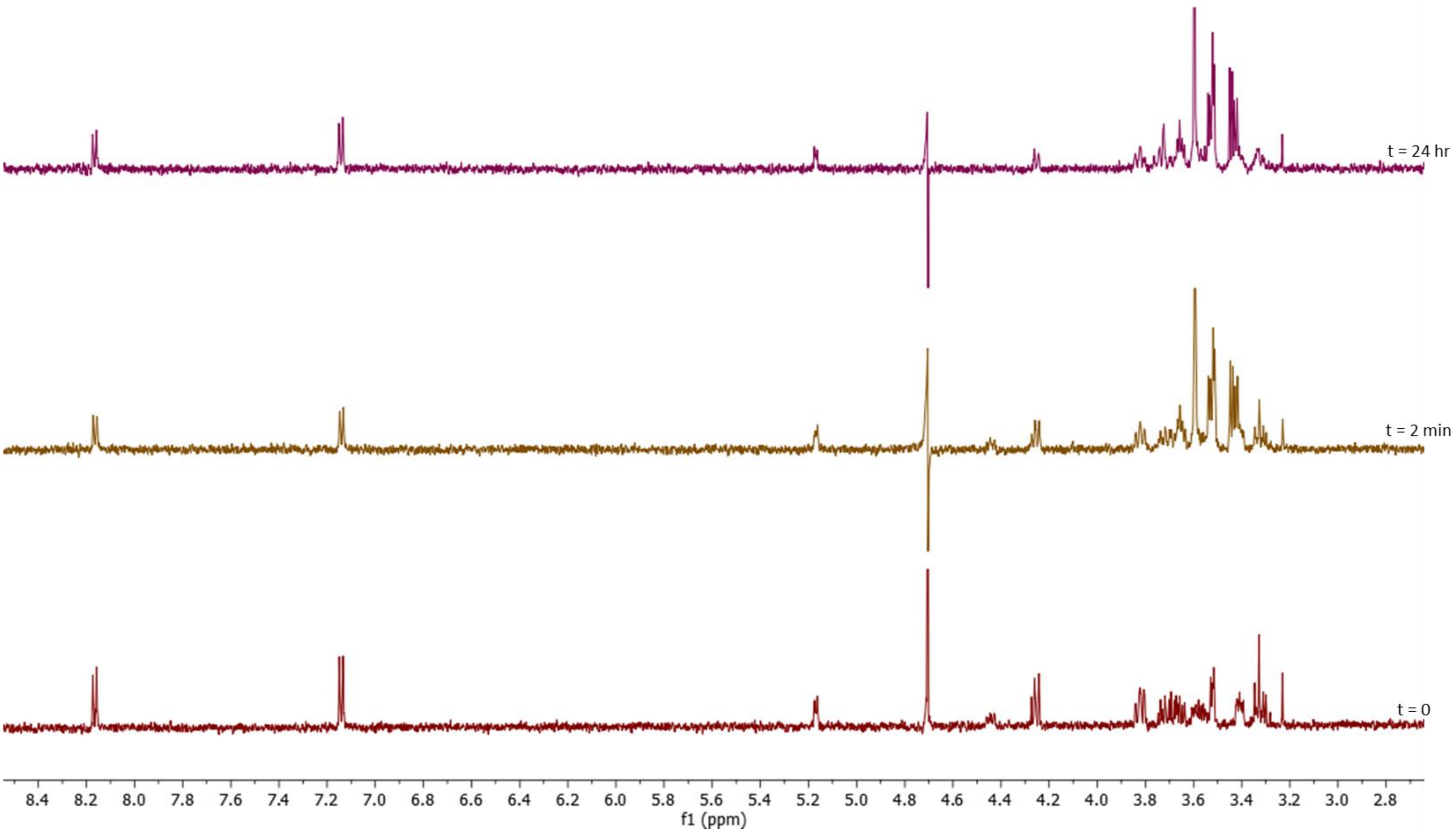

**Figure S94**: Control experiment, showing that AL2 alone does not catalyze the hydrolysis of GlcαpNP

##### 2.7.8. Hydrolysis of GlcβpNP catalysed by AL1, AL2, AL3

To a solution containing 5 mM of GlcαpNP and 5 mM 3KGlcH in the reaction buffer was added 2.5 µM of each of the enzymes AL1, AL2 and AL3 and the reaction was followed by measuring the H-NMR spectra of the reaction mixture (Figure S95).

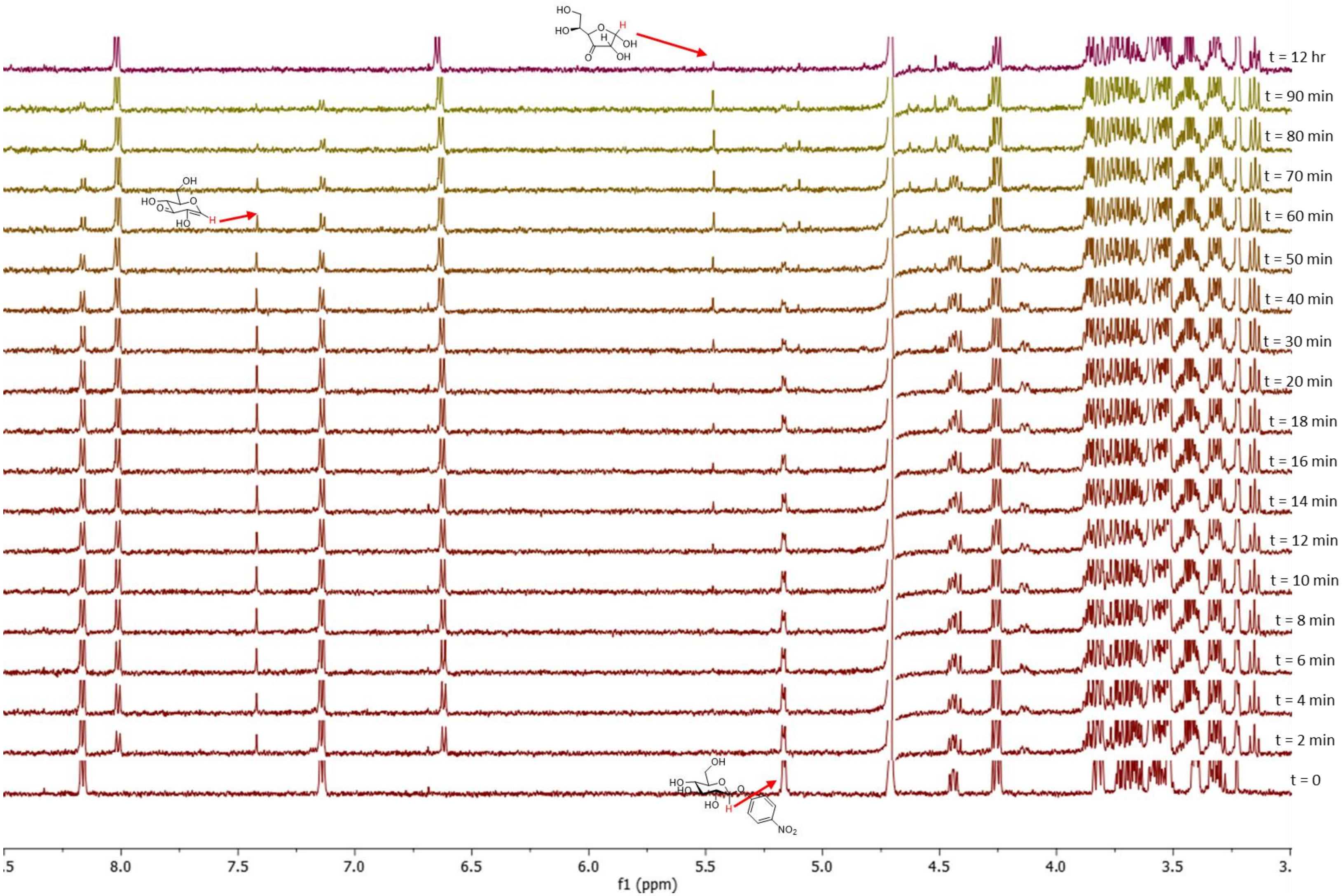

**Figure S95:** H-NMR of AL123 (2.5 µM each) catalyzed hydrolysis of GlcβpNP

Next, performinmg the same reaction with less AL1 (0.25 µM), shows that the formation of the enone intermediate is slower (Figure S96).

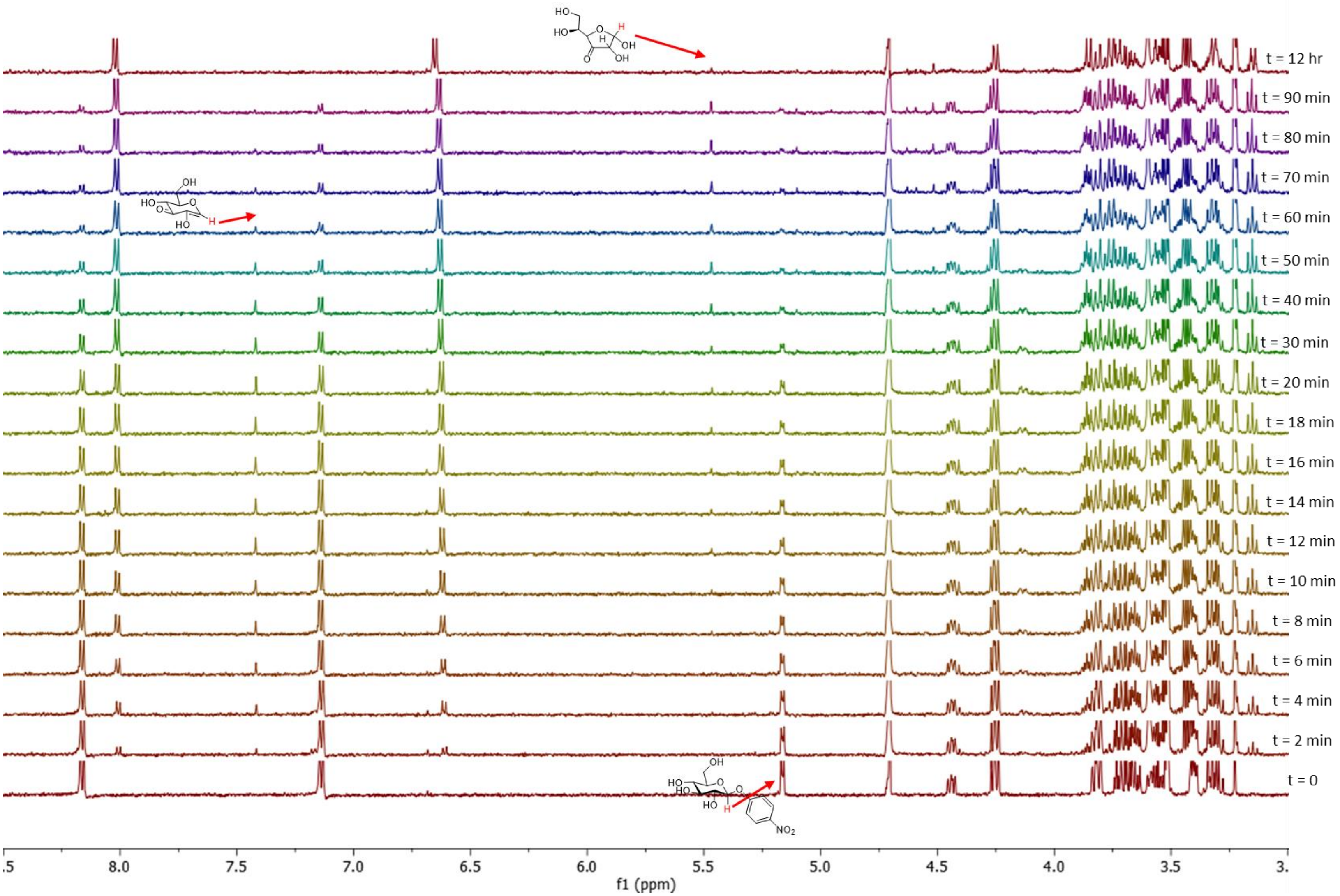

**Figure S96:** H-NMR of AL123 (2.5 µM each of AL3 and AL2, 0.25 µM AL1) catalyzed hydrolysis of GlcβpNP

Next, performinmg the same reaction with less AL2 (0.5 µM), shows accumulation of the enone intermediate and that formation of the final produtc is slower (Figure S97).

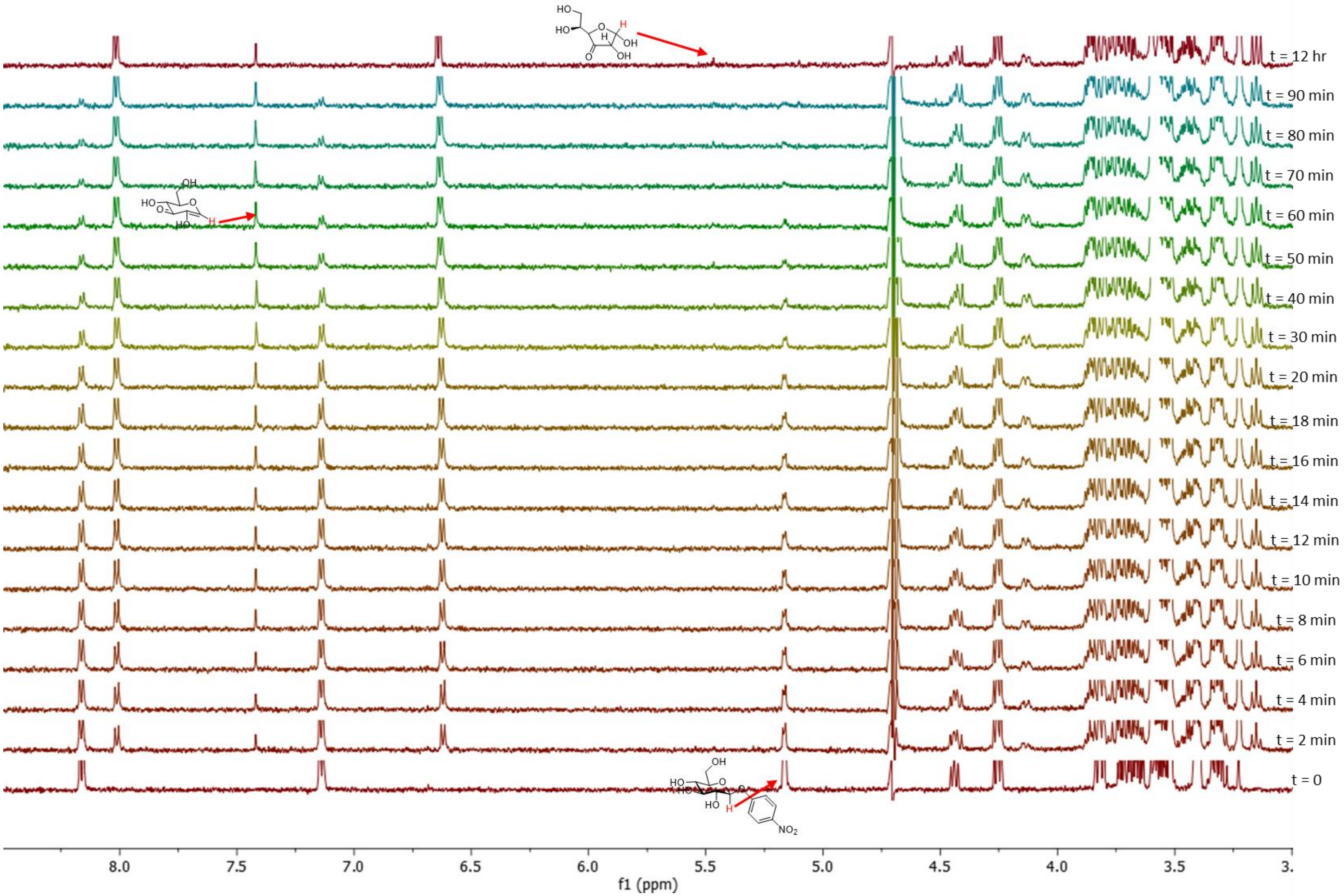

**Figure S97:** H-NMR of AL123 (2.5 µM each of AL3 and AL1, 0.5 µM AL2) catalyzed hydrolysis of GlcβpNP

Finally, performinmg the same reaction with step-wise addition of the enzymes establishes the reactions they catalyze and shows the formation of intermediate and final produtcs (Figure S98).

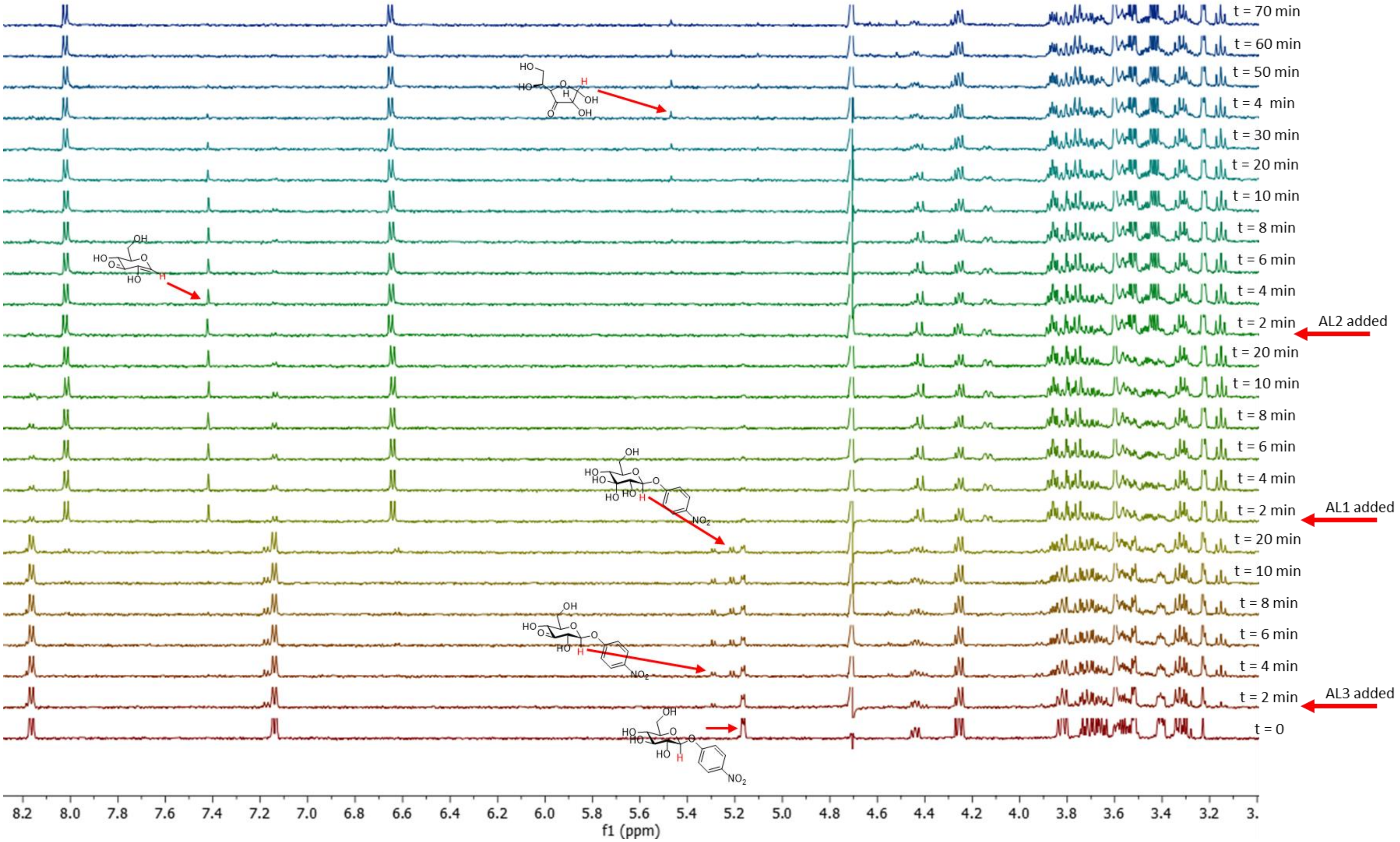

**Figure S98:** H-NMR of AL123 catalyzed hydrolysis of GlcβpNP with step-wise addition of the enzymes

###### 2.7.9. Hydrolysis of Trehalose αα catalysed by AL1, AL2, AL3

To a solution containing 5 mM of Trehalose αα and 5 mM 3KGlcH in the reaction buffer was added 2.5 μM of each of the enzymes AL1, AL2 and AL3 and the reaction was followed by measuring the H-NMR spectra of the reaction mixture (Figure S88).

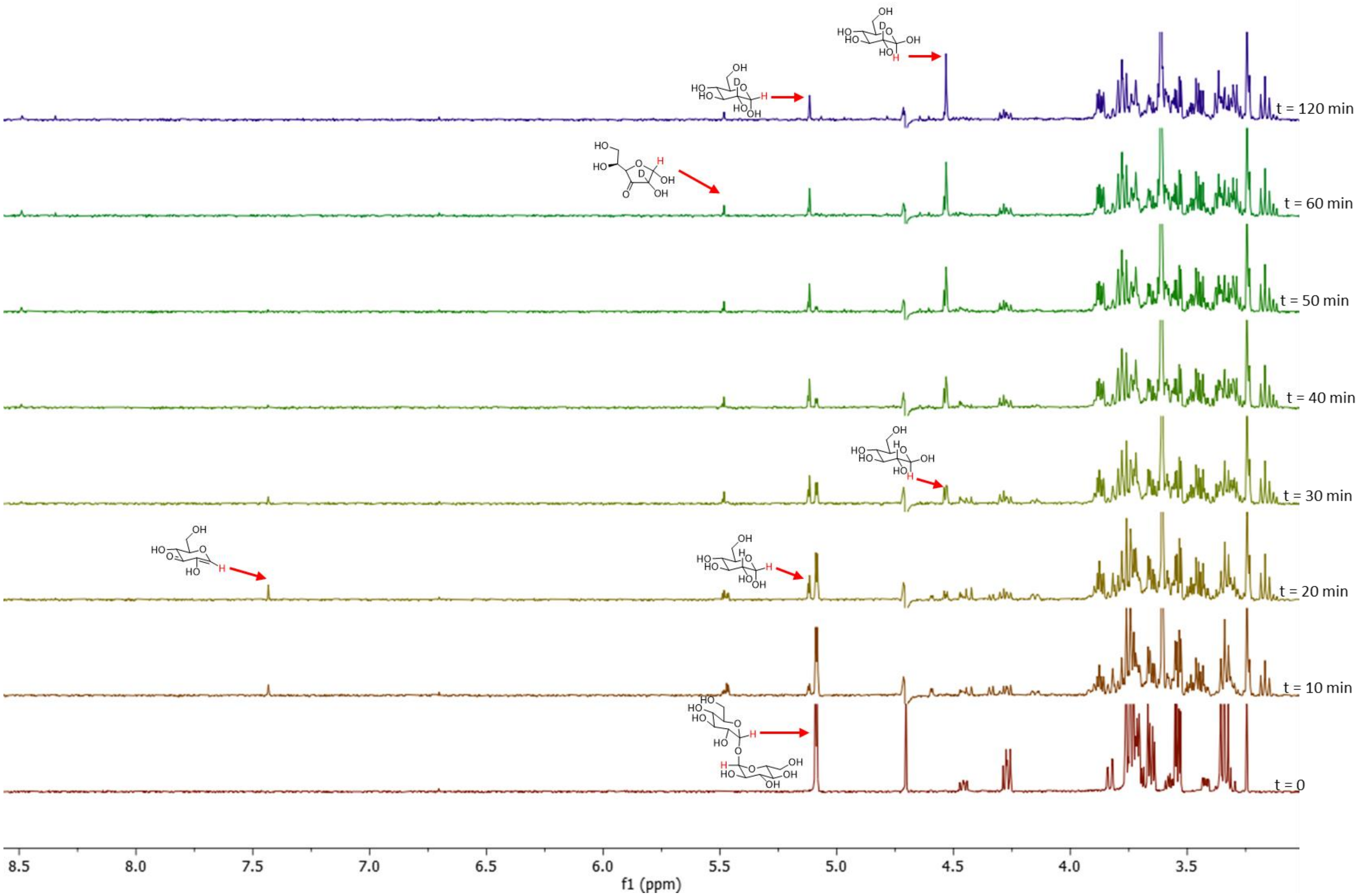

**Figure S99:** H-NMR of AL123 catalyzed hydrolysis of Trehalose αα

##### 2.7.10. Hydrolysis of Sucrose catalysed by AL1, AL2, AL3

To a solution containing 5 mM of sucrose and 5 mM 3KGlcH in the reaction buffer was added 2.5 µM of each of the enzymes AL1, AL2 and AL3 and the reaction was followed by measuring the H-NMR spectra of the reaction mixture (Figure S89).

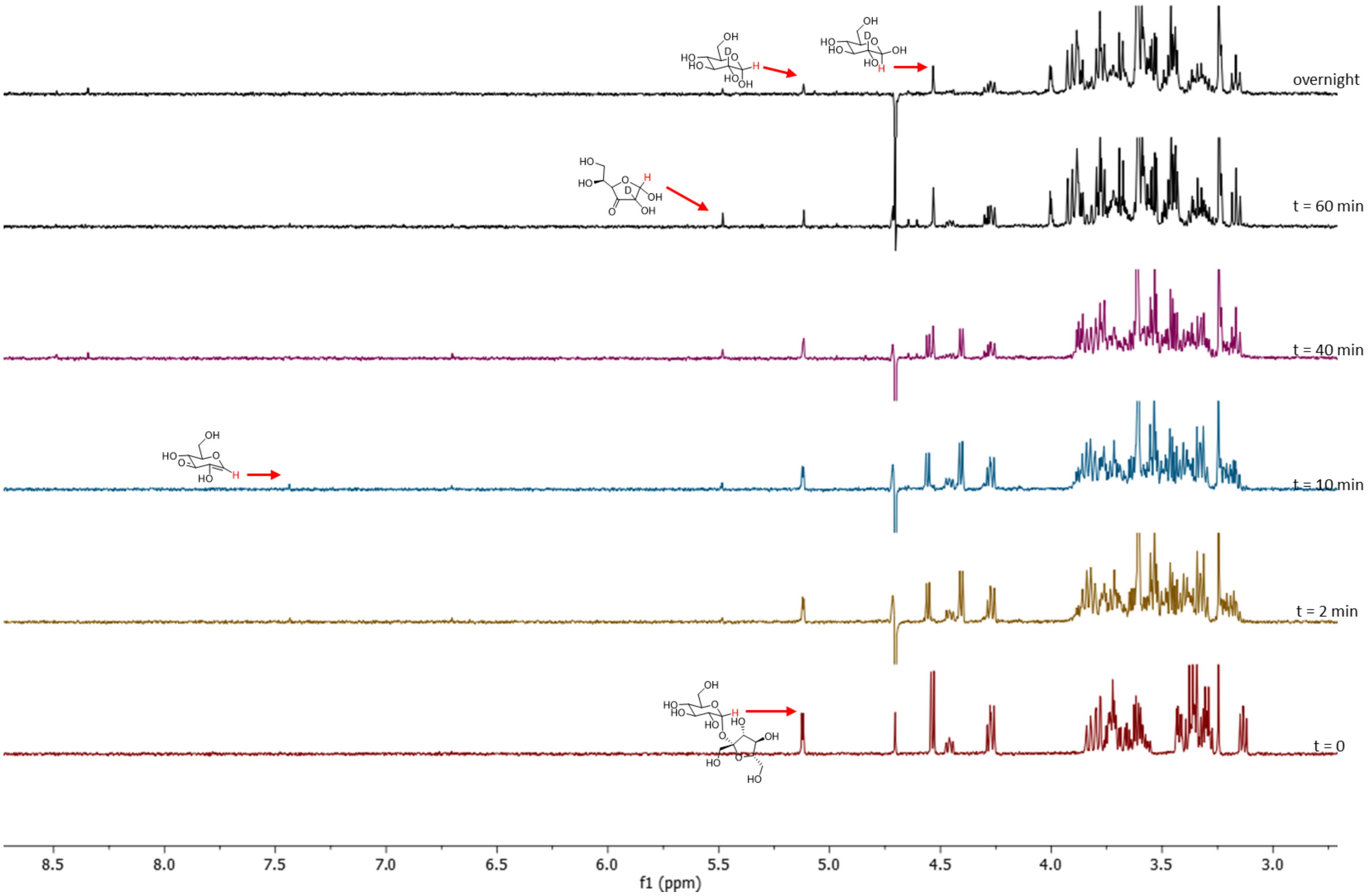

**Figure S100:** H-NMR of AL123 catalyzed hydrolysis of Sucrose

###### 2.7.11. Hydrolysis of GAαMU by enzymes from P2B11

To a solution containing 5 mM of GAαMU and 5 mM 3KGlcH in the reaction buffer was added 2.5 µM of each of the three enzymes from the P2B11 hit, Oxi1, Iso1 and Iso2 and the reaction was followed by measuring the H-NMR spectra of the reaction mixture (Figure S90). The observed products are MU and 2-Deutero-glucuronic acid, thereby indicating that the reaction has gone through a 3-keto intermediate for which incorporation of deuterium is rapid in solution, similar to the case of 3-ketoglucose, discussed above.

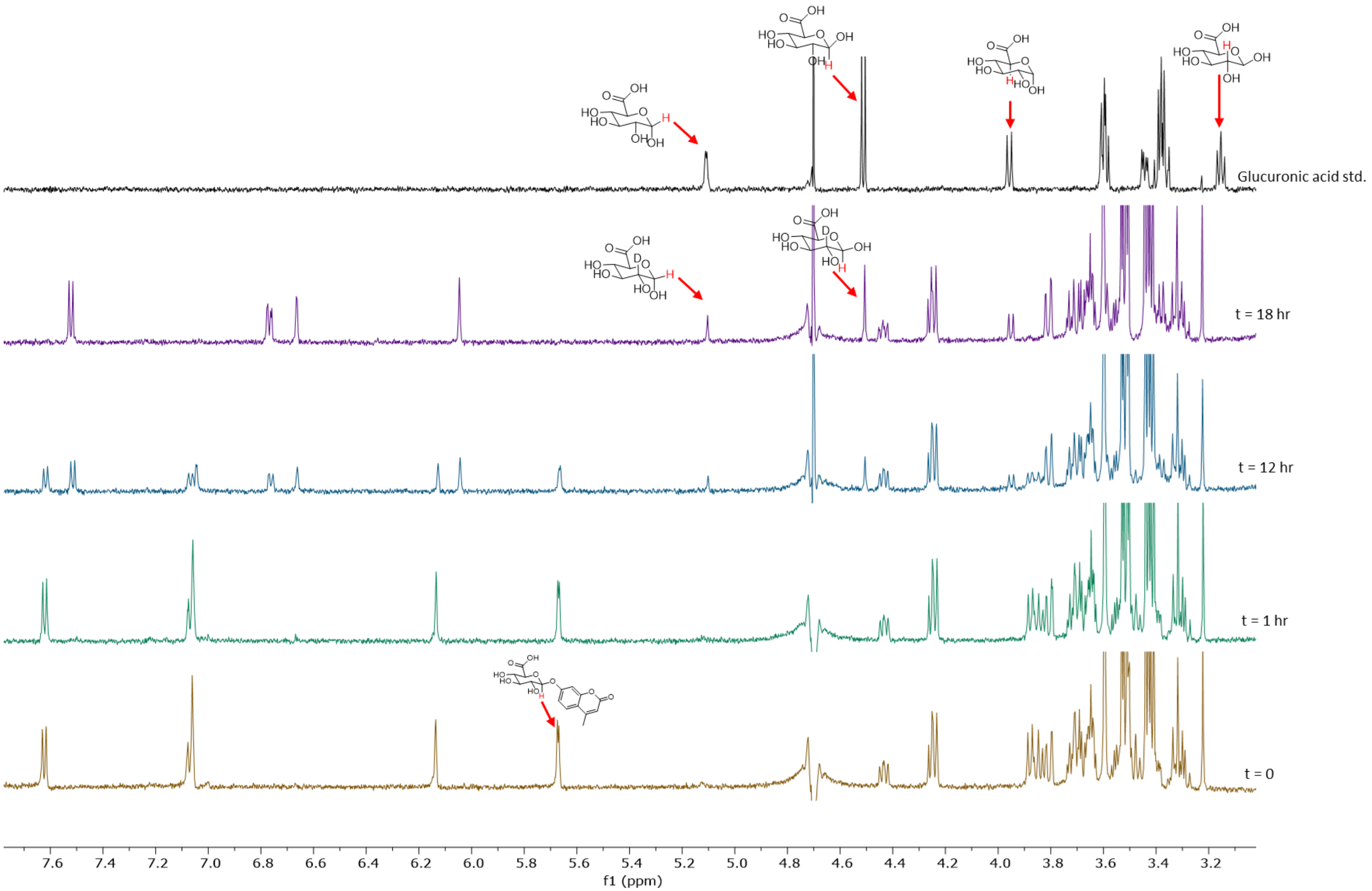

**Figure S101:** H-NMR hydrolysis of GAαMU catalyzed by P2B11 enzymes Oxi1, Iso1 and Iso

##### 2.7.12. Enzymatic kinetics for AL3

The reaction of GlcαpNP with various concentrations and 3KGlcH (10 mM) were initiated with addition of the enzyme AL3 (final conc 0.2 µM) and monitored by H-NMR every 80 seconds at room temperature. Since the relaxation time (T1) for the corresponding peaks of the starting material and the product were measured to be the same, the concentrations of the compounds in solution were calculated based on the integration of the anomeric protons and with respect to the mass balance equation. Initial rates of the reaction were next calculated based on the changes in concentration of the product for the first 5 minutes of the reactions and were used to construct Michaelis-Menten kinetic models for this enzymatic reaction. Note that we have kept the concentration of 3KGlcH constant throughout this experiment and have assumed that at the high concentration of 10 mM, its reduction is faster than the oxidation of GlcαpNP. Also, since the enzyme catalyzes both the oxidation of GlcαpNP and reduction of 3KGlcH at the same rates but we are measuring the rate for only one of these reactions, the total enzyme concentration for calculation of $k_{cat}$ was set to half the total available enzyme, 1 µM.

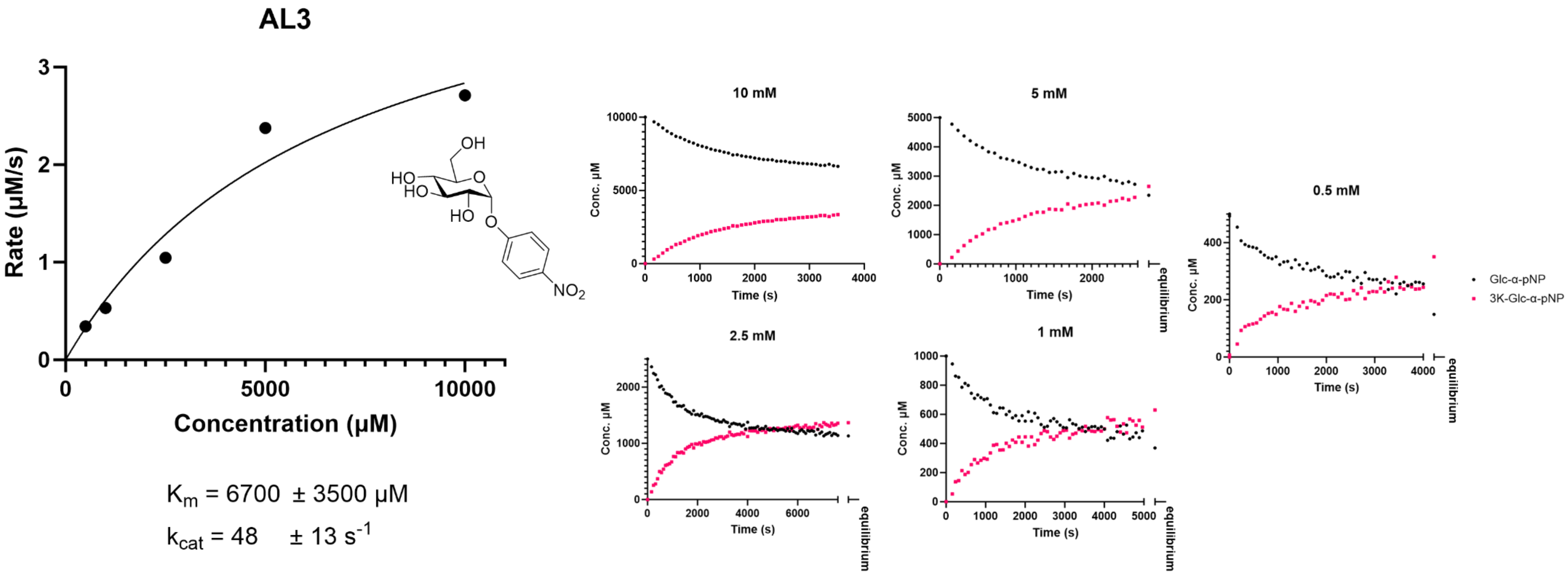

**Figure S102:** Pseudo-Michaelis-Menten kinetics for AL3 – parameters determined by H-NMR monitoring of the area for anomeric peak in different concentrations – reaction coordinates, calculated are shown in the right

#### 2.8. X-ray crystallography

AL, BT, and P2B11 enzymes were screened for crystals using PACT and JCSG+ (Qiagen) crystallographic screens using sitting drop vapor diffusion in INTELLI-PLATE 96 well plates (Art Robbins Instruments) with 0.2 µL of protein mixed with 0.2 µL of mother liquor. Hits were optimized by varying the conditions pH, salt, or precipitant. Hits and final conditions are described in supplemental Table S6. For all crystals, either glucose or trehalose at 25% (w/v), glycerol 30% (w/v), or increased PEG concentration of 30% (w/v) was used as cryoprotectant prior to flash freezing with cryoprotectants for each of the crystals described in Table S6.

Screening and collection of diffraction data (Table S5) was performed at 100 K on beamlines CMCF-BM at the Canadian Light Source using MxDC for data collection, or 23-ID-D at the Advanced Photon Source using JBluIce for data collection. Diffraction data were processed using xia2[32] and XDS[33], with data reduction carried out using Aimless[34] as part of the CCP4 package[35]. Anisotropic datasets, (trehalose bound AL3) were corrected using the STARANISO server[36]. Phasing was carried out using molecular replacement with Alphafold[37] models for all enzymes except BT2, with alphafold models pLDDT scores converted to b-factors using Process Predicted Models in the CCP4 suite, and chain A as the search model in Phaser[38], also part of the CCP4 package. BT2 phasing was completed using PDB ID 3OSD. Sequential rounds of model building, and refinement were carried out using Coot[39] and Refmac[40]. Validation of the final models was carried out using MolProbity[41].

Regions of the structures that were poorly resolved or disordered were omitted from the final models. The final models contain the following chains, built residues, and Ramachandran statistics: AL1 4 chains, residues 30-300, 0.00% outliers, 96.75% favored; AL1 Glucose 4 chains, residues 31-300, 0.00% outliers, 96.92% favored; AL1 Trehalose 4 chains, residues 31-300, 0.00% outliers, 97.02% favored; BT1 3K-GlcH 1 chain, residues 30-304, 0.00% outliers, 97.07% favored; AL2 2 chains, residues 41-100, 108-290 in Chain A, 40-290 in Chain C, 0.00% outliers, 98.36% favored; BT2 2 chains, residues 35-290, 0.00% outliers, 97.05% favored; BT2 Glucose 2 chains, residues 35-290, 0.00% outliers, 97.07% favored; AL3 4 chains, residues 39-490, 0.06% outliers, 97.67% favored; AL3 Trehalose 4 chains, residues 39-490, 0.06% outliers, 95.83% favored; P2B11-Oxido-1 12 chains, residues 11-303, 318-409 in Chain A, 11-35, 43-313, 321-409 in Chain B, 12-36, 45-301, 320-409 in Chain C, 10-303, 320-408 in Chain D, 12-40, 44-303, 320-409 in Chain E, 11-39, 43-303, 317-409 in Chain F, 11-37, 43-310, 320-410 in Chain G, 11-37, 42-303, 320-409 in Chain H, 11-303, 320-409 in Chain I, 11-302, 320-409 in Chain J, 12-35, 45-301, 321-409 in Chain K, 11-35, 46-313, 320-411 in Chain L, 0.00% outliers, 97.56% favored. We note in the refined structure of BT1 in complex with 3-keto-1,5-anhydoglucitol, there is significant residual positive density in the mFo-DFc difference map around the inhibitor and neighbouring active site residue Trp148 (see Fig. S105 D,E). This could indicate an alternative binding conformation of the 3-keto-1,5-anhydoglucitol - for example the glucose bound in the AL1 active site (Fig. S105 A,B) is in a similar but rotated binding orientation (overlaid in Fig. S105 G,H) - or possibly a mixed occupancy with an impurity from the chemical synthesis (see SI section 1.3). Attempts to model this did not improve map interpretation and the selected binding orientation is best supported by an omit map calculated in the absence of the ligand and by comparison with the AL1 active site glucose.

All structure analysis and figure preparation were carried out with ChimeraX[42] and Coot, distributed as part of the CCP4 package. Sugar orientations and density fit were checked and analyzed using Privateer[43], and dimeric interfaces assessed using PISA[44]. Determination of possible catalytic residues was completed by analyzing highly conserved residues in the active site of each enzyme using the ConSurf server[45] (Fig. S92). Metal coordination was checked with the check my metal server[46]. Comparisons of homologous AL, BT, and CT enzymes was carried out using the ChimeraX Matchmaker function. Electron density maps for figures were generated by removing sugars or ligands from the models followed by a subsequent round of TLS refinement resetting B-factors to 30 $Å^2$ and restrained refinement to make omit maps of substrates and co-factors.

**Data availability**
The data that support this study are available from the corresponding authors upon request. Structure factors and atomic coordinates have been deposited with the protein data bank with accession codes 8TCD, 8TCR, 8TCS, 8TCT, 8TDA, 8TDE, 8TDF, 8TDH, 8TDI, 8V31.

**Table S5: Data collection and refinement statistics. Statistics for the highest resolution shell are shown in parentheses.**

| | AL1 | AL1 Glucose | AL1 Trehalose | BT1 3KGlcH | AL2 | BT2 | BT2 Glucose | AL3 | AL3 Trehalose‡ | P2B11-Oxido-1 |
|---|---|---|---|---|---|---|---|---|---|---|
| **Data collection** | | | | | | | | | | |
| Space group | P 21 21 21 | P 21 21 21 | P 21 21 21 | P 32 2 1 | P 31 1 2 | P 21 21 21 | P 21 21 21 | I 1 2 1 | I 1 2 1 | P 1 21 1 |
| Cell dimensions | | | | | | | | | | |
| *a*, *b*, *c* (Å) | 47.57, 112.61, 209.34 | 47.517, 113.68, 208.58 | 47.99, 113.55, 212.44, | 122.29, 122.29, 68.75 | 95.09, 95.09, 159.54 | 75.94, 80.02, 103.15 | 77.04, 81.17, 103.78 | 180.64, 56.75, 220.45 | 186.58, 57.22, 224.96 | 106.26, 185.88, 145.36 |
| α, β, γ (°) | 90, 90, 90 | 90, 90, 90 | 90, 90, 90 | 90, 90, 120 | 90, 90, 120 | 90, 90, 90 | 90, 90, 90 | 90, 108.87, 90 | 90, 106.59, 90 | 90, 111.20, 90 |
| Resolution (Å) | 46.39 – 1.90 (1.97-1.90) | 47.40 – 2.08 (2.15 – 2.08) | 44.30 – 1.50 (1.55 – 1.50) | 45.69 – 1.86 (1.93 – 1.86) | 47.59 – 2.65 (2.72 – 2.65) | 61.15 – 1.46 (1.51 – 1.46) | 43.72 – 1.85 (1.92 – 1.85) | 48.00 – 2.10 (2.18 – 2.10) | 55.31 – 2.95 (3.06 – 2.95) | 48.37 – 2.60 (2.69 – 2.60) |
| $R_{merge}$ | 0.161 (1.562) | 0.135 (0.978) | 0.071 (1.303) | 0.082 (2.707) | 0.145 (2.106) | 0.152 (1.223) | 0.182 (1.051) | 0.181 (1.583) | 0.141 (0.469) | 0.120 (1.721) |
| $I / \sigma I$ | 9.69 (1.26) | 8.93 (1.66) | 11.04 (1.33) | 28.97 (1.16) | 22.90 (1.90) | 13.60 (1.29) | 13.18 (1.81) | 8.68 (1.58) | 3.51 (2.20) | 13.73 (1.11) |
| CC1/2 | 0.996 (0.557) | 0.997 (0.603) | 0.999 (0.575) | 1 (0.619) | 0.999 (0.868) | 0.991 (0.506) | 0.985 (0.615) | 0.992 (0.557) | 0.95 (0.497) | 0.998 (0.507) |
| Completeness (%) | 99.35 (94.72) | 99.68 (98.18) | 99.80 (99.90) | 99.93 (99.86) | 99.12 (97.72) | 99.87 (99.94) | 99.77 (99.96) | 99.66 (99.66) | 89.62 (63.96) | 99.94 (99.98) |
| Redundancy | 6.4 (6.2) | 6.3 (5.3) | 6.4 (6.6) | 19.9 (19.7) | 18.9 (18.5) | 6.4 (5.9) | 6.5 (6.4) | 6.6 (6.8) | 2.7 (2.4) | 6.9 (6.9) |
| **Refinement** | | | | | | | | | | |
| Resolution (Å) | 1.90 | 2.08 | 1.50 | 1.86 | 2.65 | 1.46 | 1.85 | 2.10 | 2.95 | 2.60 |
| No. reflections | 89195 (8375) | 68822 (6689) | 186072 (18379) | 49898 (4903) | 24911 (3236) | 109360 (10835) | 56117 (5543) | 123787 (12234) | 43691 (3079) | 161056 (16054) |
| $R_{work}$ / $R_{free}$ | 0.184/0.226 | 0.177/0.227 | 0.150/0.194 | 0.154/0.164 | 0.197/0.251 | 0.168/0.209 | 0.189/0.236 | 0.199/0.233 | 0.196/0.246 | 0.187/0.232 |
| No. atoms | | | | | | | | | | |
| Protein | 8672 | 8644 | 8685 | 2226 | 7810 | 4075 | 4058 | 14249 | 14220 | 36746 |
| Ligand/ion | 36 | 130 | 100 | 42 | 2 | 2 | 14 | 176 | 705 | 662 |
| Water | 512 | 318 | 870 | 226 | 40 | 553 | 398 | 414 | 140 | 255 |
| *B*-factors ($Å^2$) | | | | | | | | | | |
| Protein | 25.3 | 25.5 | 18.2 | 44.7 | 63.1 | 24.0 | 33.4 | 53.7 | 42.6 | 74.1 |
| Ligand/ion | 35.5 | 41.9 | 27.0 | 86.7 | 43.6 | 21.0 | 52.0 | 44.8 | 46.9 | 74.3 |
| Water | 31.7 | 33.3 | 33.4 | 57.4 | 45.5 | 31.7 | 36.4 | 37.6 | 23.4 | 47.7 |
| R.m.s. deviations | | | | | | | | | | |
| Bond lengths (Å) | 0.015 | 0.016 | 0.015 | 0.012 | 0.012 | 0.014 | 0.015 | 0.016 | 0.012 | 0.013 |
| Bond angles (°) | 2.02 | 2.10 | 1.89 | 1.83 | 2.40 | 1.88 | 2.08 | 2.08 | 1.96 | 2.09 |

‡Anisotropic cut-off applied to merged intensity data.[36]

**Table S6:** Description of protein crystallization and cryoprotection**.**

| **Protein name** | **AL1** | **AL1 Glucose** | **AL1 Trehalose** | **BT1 3K-GlcH** |
|---|---|---|---|---|
| Concentration (mg/mL) | 33 | 33 | 33 | 15 |
| Screen | PACT A8 ‡ | PACT B2 ‡ | PACT A8 ‡ | JCSG C6 ‡ |
| Crystal Condition | 0.27 M Ammonium Chloride | | 0.44 M Ammonium Chloride | |
| | 17% PEG 6000 | 25% PEG 1500 | 21% PEG 6000 | 27% PEG 300 |
| | 0.1 M Sodium Acetate pH 5 | 0.13 M MIB pH 5 | 0.1 M Sodium Acetate pH 5 | 0.11 M Phosphate Citrate pH 4.2 |
| Cryoprotectant | 25% Glycerol | 25% Glucose | 25% Trehalose | 30% PEG 300 |
| Beamline | CLS CMCF BM | CLS CMCF BM | APS 23IDD | CLS CMCF BM |
| Wavelength (Å) | 1.18 | 1.18 | 1.03 | 1.18 |

| **Protein name** | **AL2** | **BT2** | **BT2 Glucose** |
|---|---|---|---|
| Concentration (mg/mL) | 15 | 15 | 15 |
| Screen | PACT H11 ‡ | PACT E11 ‡ | PACT E11 ‡ |
| Crystal Condition | 0.2 M Magnesium Chloride | 0.2 M Potassium Citrate | 0.2 M Potassium Citrate |
| | 25% PEG 3350 | 18% PEG 3350 | 22% PEG 3350 |
| | 0.1 M Bis-Tris methane pH 5.5 | | |
| Cryoprotectant | 30% PEG 3350 | 30% PEG 3350 | 25% Trehalose |
| Beamline | CLS CMCF BM | APS 23IDD | CLS CMCF ID |
| Wavelength (Å) | 1.18 | 1.03 | 1.18 |

| **Protein name** | **AL3** | **AL3 Trehalose** | **P2B11-Oxido1** |
|---|---|---|---|
| Concentration (mg/mL) | 5 | 5 | 12 |
| Screen | PACT E12 ‡ | PACT E12 ‡ | JCSG H9 ‡ |
| Crystal Condition | 0.15 M Sodium Malonate | 0.25 M Sodium Malonate | 0.2 M LiSO4 |
| | 19% PEG 3350 | 17% PEG 3350 | 19% PEG 3350 |
| | | | 0.1 M Bis-Tris methane pH 5.5 |
| Cryoprotectant | 30% PEG 3350 | 25% Trehalose | 30% PEG 3350 |
| Beamline | CLS CMCF ID | APS 23IDD | CLS CMCF BM |
| Wavelength (Å) | 0.95 | 1.03 | 1.18 |

‡ Crystal screen condition optimized.

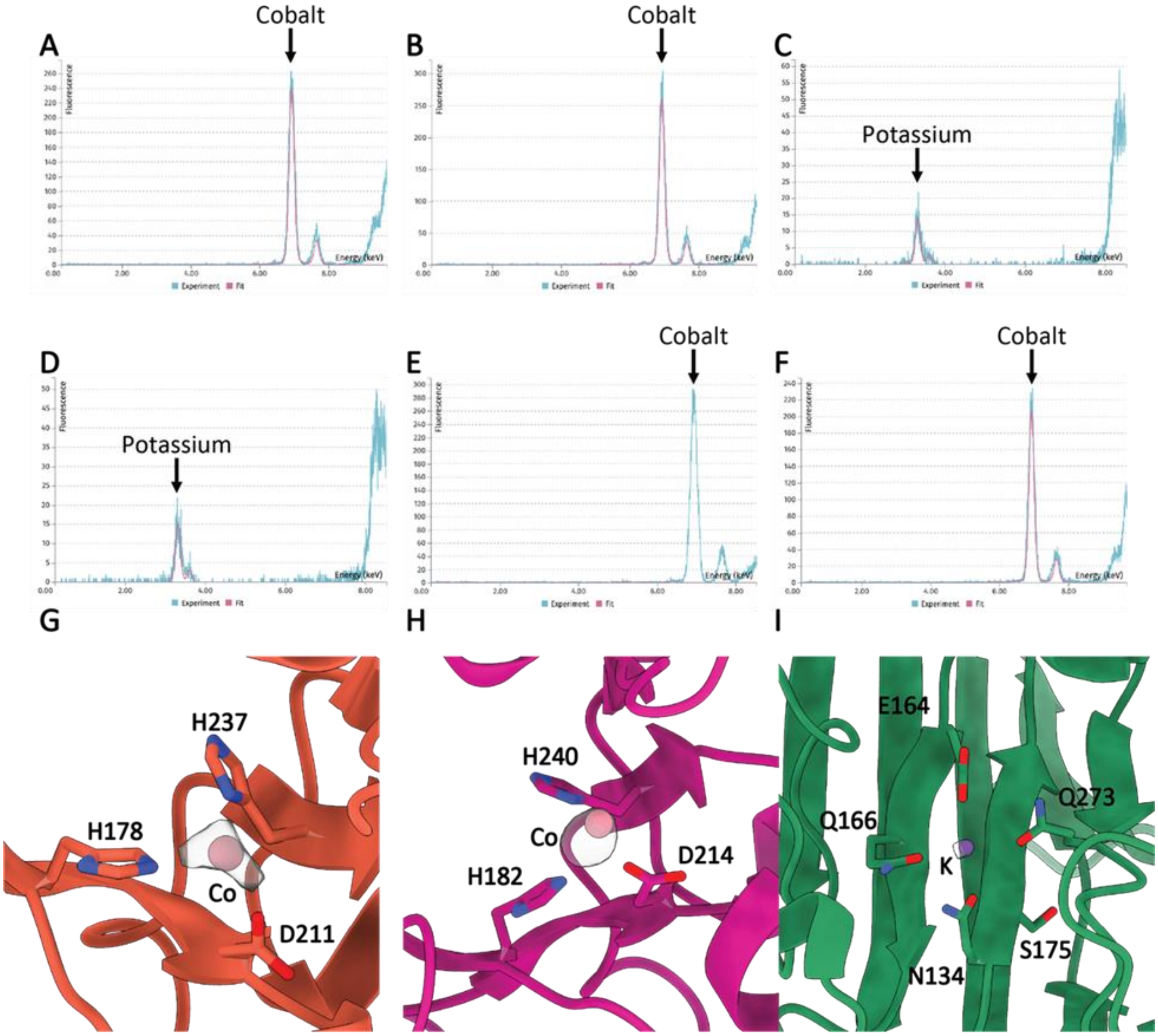

**Fig S103**. X-ray fluorescence scans (XRF) and anomalous density peaks of apo AL1, 3-KGlcH BT1, apo BT2, glucose BT2, apo BT3, and apo P2B11-Oxido-1 crystals. Anomalous density maps generated by CCP4i2[35]. Metal coordinating residues are shown as sticks and anomalous densities as grey surfaces. Metals were checked with the Check My Metal server for proper coordination. (A) XRF scan of apo AL1. (B) XRF scan of 3K-GlcH BT1. (C) XRF scan of apo BT2. (D) XRF scan of glucose BT2. (E) XRF scan of apo AL3. (F) XRF scan of apo P2B11-Oxido-1. (G) Anomalous cobalt peak in apo AL1. (H) Anomalous cobalt peak in 3-KGlcH BT1. (I) Anomalous potassium peak in apo BT2.

.

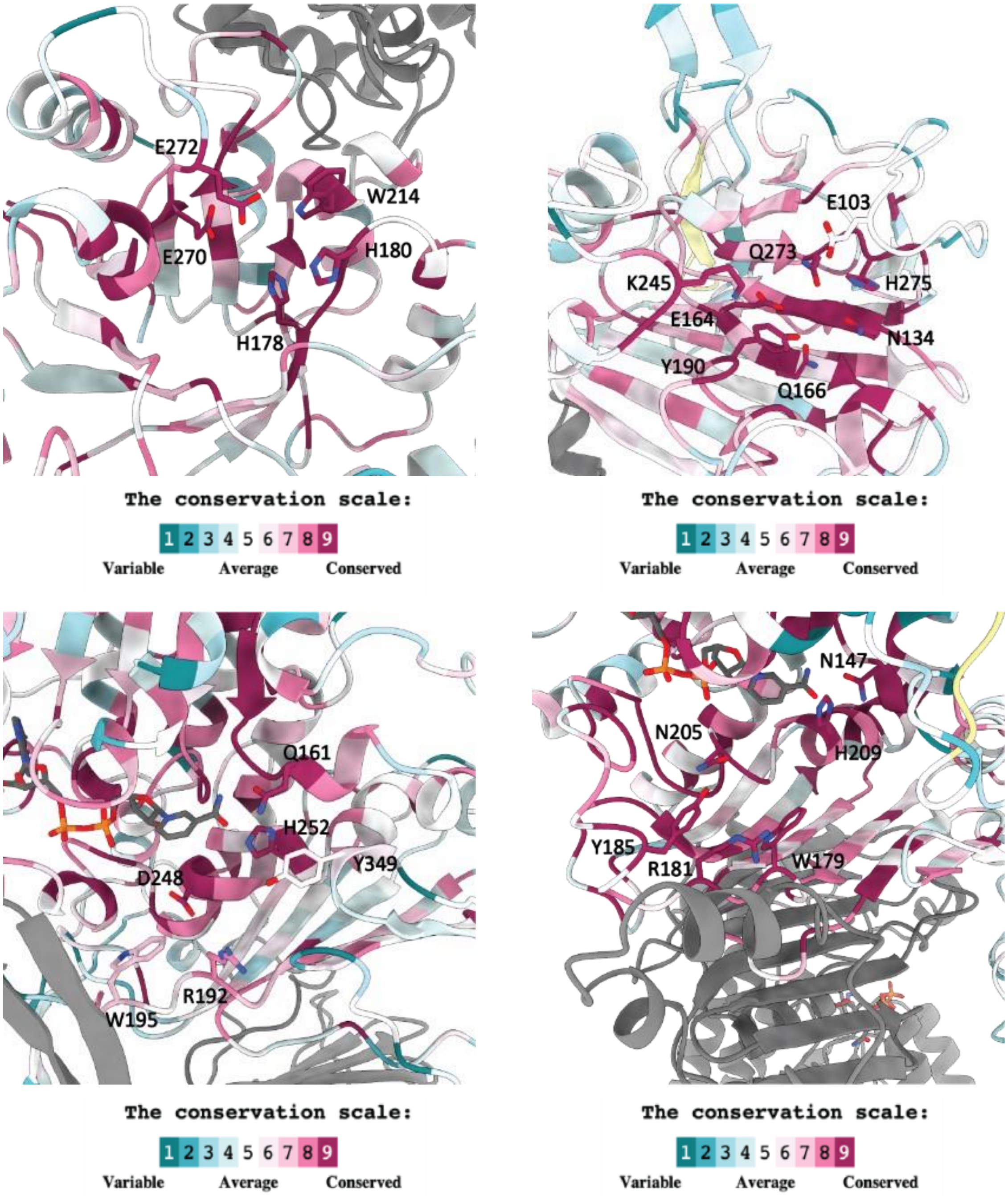

**Fig S104.** Consurf[45] models of apo AL1 (A), apo BT2 (B), AL3 (C), and apo P2B11-Oxido-1 (D) active sites with corresponding conservation scales and conserved residues involved in catalysis or co-factor coordination shown as sticks.

.

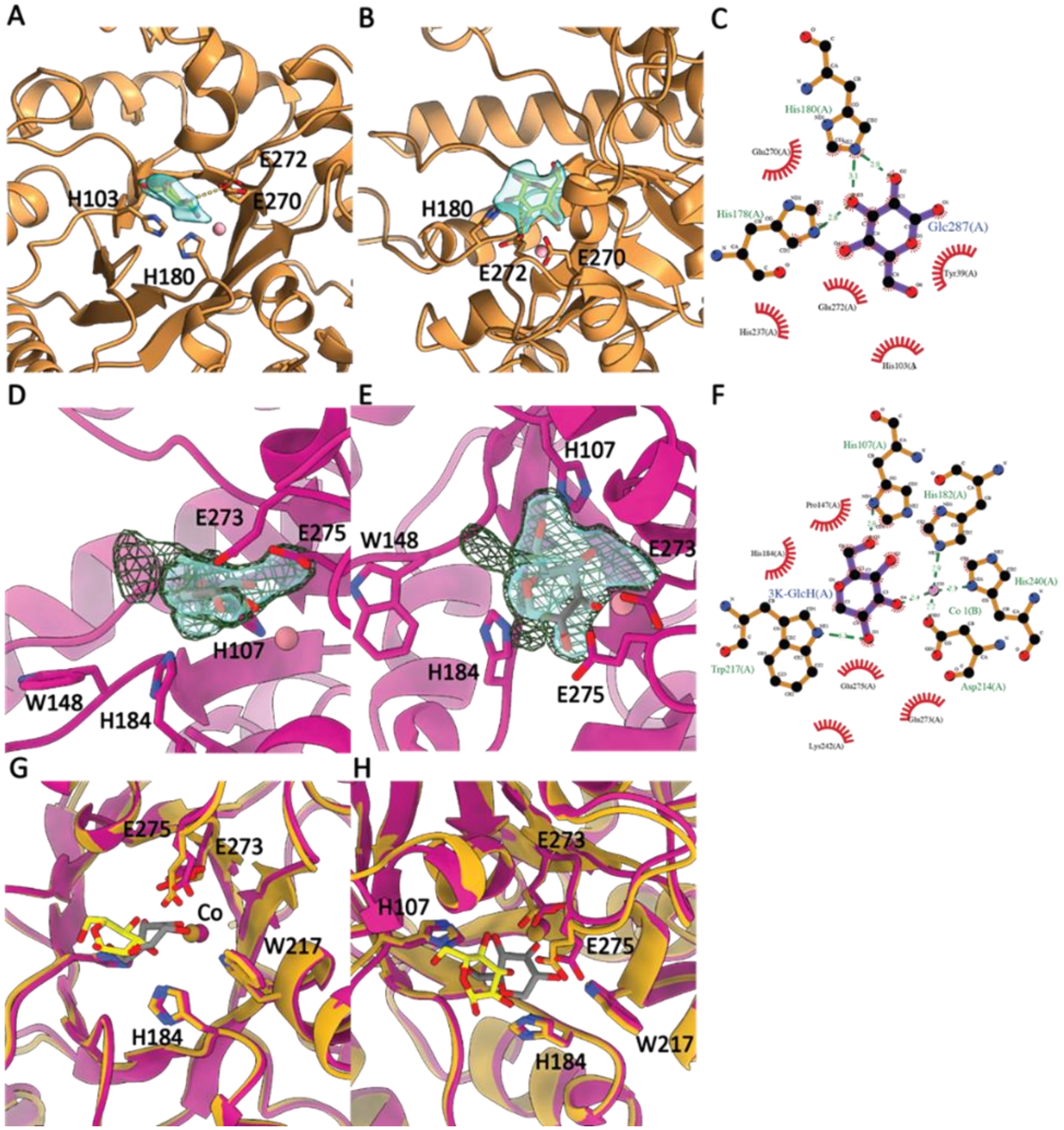

**Figure S105.** 2mFo-DFc omit maps of molecules in AL1 and BT1 active sites. Omit maps were generated in CCP4i2[35] by removing the ligand and completing a round of restrained refinement for 10 cycles followed by 5 TLS cycles after resetting B-factors to 30 Å$^2$ and shown in ChimeraX[42] as blue transparent surfaces at a RMSD of 1.5 in AL1 and BT1 substrate bound models and 0.9 (green mesh) in BT1 3K-GlcH, with interacting residues in active site labelled. (A) Side view of glucose in AL1 active site. (B) Top view of glucose in AL1 active site. (C) LigPlot of glucose and AL1. (D) Side view of 3K-GlcH in BT1 active site. (E) Top view of 3K-GlcH in BT1 active site. (F) LigPlot of 3K-GlcH and BT1. (G) Side view of aligned BT1 3K-GlcH chain A with AL1 glucose chain B. (H) Top view of aligned BT1 3K-GlcH chain A with AL1 glucose chain B. We note in the BT1 3K-GlcH structure (panels D and E), there is significant additional density around 3K-GlcH and neighbouring residue Trp148 as shown in the green mesh contoured at a lower RMSD. This could indicate an alternative binding conformation of the 3K-GlcH (for example the glucose bound in the AL1 active site is in a similar but rotated binding orientation as shown in G and H) or possibly a mixed occupancy with an impurity from the chemical synthesis (see SI section 1.3). Also see SI section 2.8.

.

.

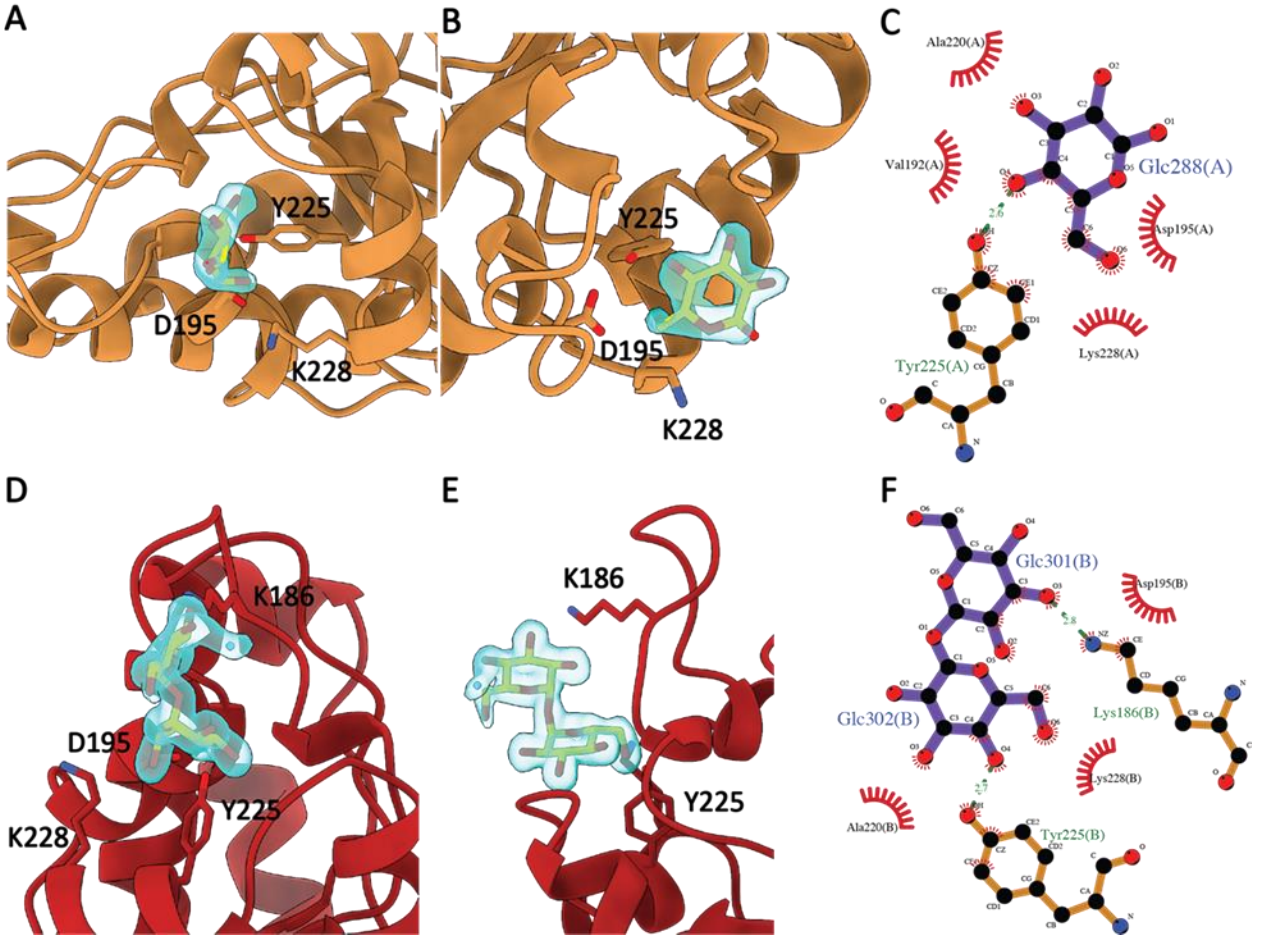

**Figure S106.** 2mFo-DFc omit maps of molecules in AL1 secondary glucose and trehalose binding sites. Omit maps were generated in CCP4i2[35] by removing the ligand and completing a round of restrained refinement for 10 cycles followed by 5 TLS cycles after resetting B-factors to 30 Å$^2$ and shown in ChimeraX[42] as blue transparent surfaces at a RMSD of 1.3 in AL1 glucose and 1.0 in AL1 trehalose with interacting residues labelled. (A) Side view of glucose in AL1 secondary site. (B) Top view of glucose in AL1 secondary site. (C) LigPlot of glucose and AL1 secondary site. (D) Side view of trehalose in AL1 secondary site. (E) Top view of trehalose in AL1 secondary site. (F) LigPlot of trehalose and AL1 secondary site.

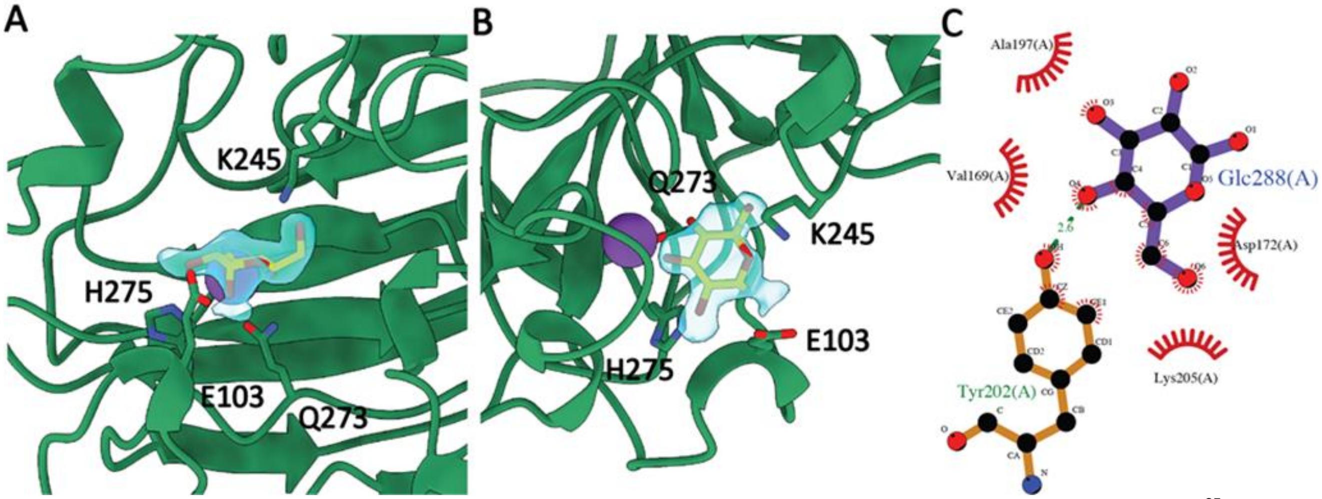

**Figure S107**. 2mFo-DFc omit maps of glucose like molecule in BT2 active site. Omit maps were generated in CCP4i2[35] by removing the ligand and completing a round of restrained refinement for 10 cycles followed by 5 TLS cycles after resetting B-factors to 30 Å$^2$ and shown in ChimeraX[42] as blue transparent surfaces at a RMSD of 1.0 with interacting residues in active site labelled. (A) Side view of glucose in BT2 active site. (B) Top view of glucose in BT2 active site. (C) LigPlot of glucose BT2 interactions.

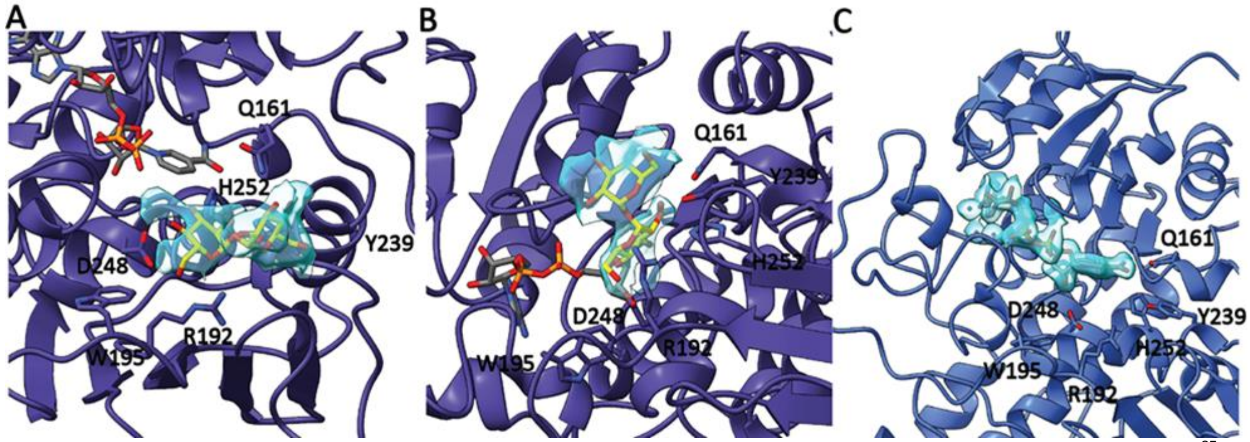

**Figure S108**. 2mFo-DFc omit maps of NAD and active site trehalose in AL3. Omit maps were generated in CCP4i2[35] by removing the ligand and completing a round of r restrained refinement for 10 cycles followed by 5 TLS cycles after resetting B-factors to 30 Å$^2$ and shown in ChimeraX[42] as blue transparent surfaces at a RMSD of 1.0 in AL3 trehalose and 1.5 in AL3 apo with interacting residues in active site labelled. (A) Side view of trehalose in AL3 active site. (B) Top view of trehalose in AL3. (C) Omit map of NAD in AL3 apo

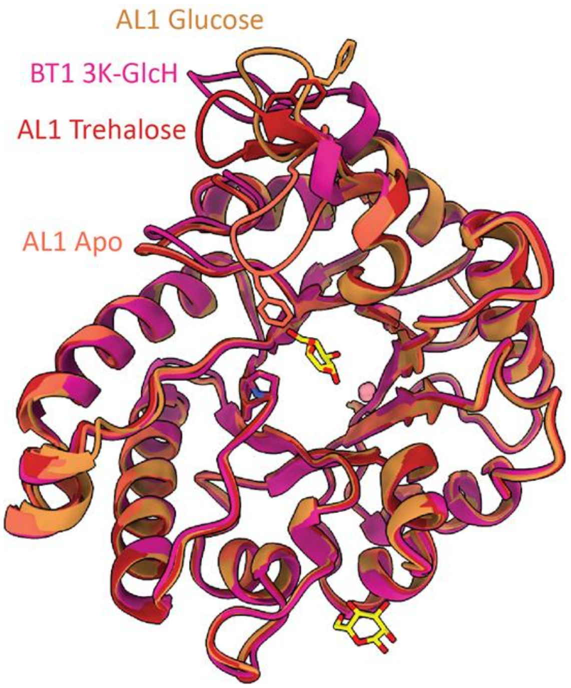

**Figure S109.** Structural alignment of different AL1 and BT1 models using ChimeraX Matchmaker[42] to AL1 apo (tomato), AL1 trehalose (red) RMSD 0.3 Å over 262 Cα pairs, BT1 3K-GlcH (magenta) RMSD 0.6 Å over 257 Cα pairs, and AL1 glucose (orange) RMSD 0.4 Å over 262 Cα pairs

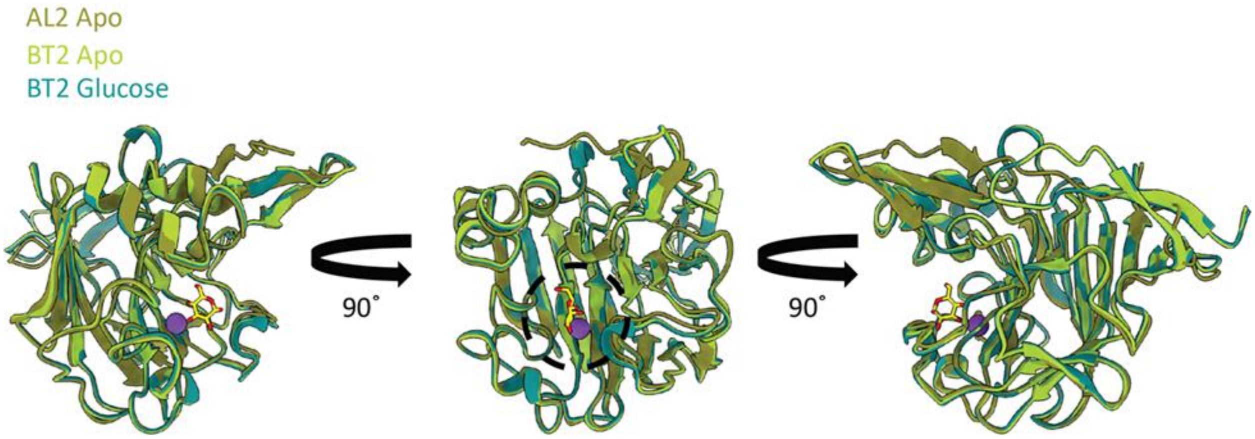

**Figure S110.** Structural alignments of AL2 and BT2 models to chain A apo BT2 (yellow green) using ChimeraX Matchmaker[42]. Chain C AL2 apo (olive) RMSD 0.5 Å over 229 Cα pairs, chain A BT2 glucose (teal) RMSD 0.2 Å over 256 Cα pairs.

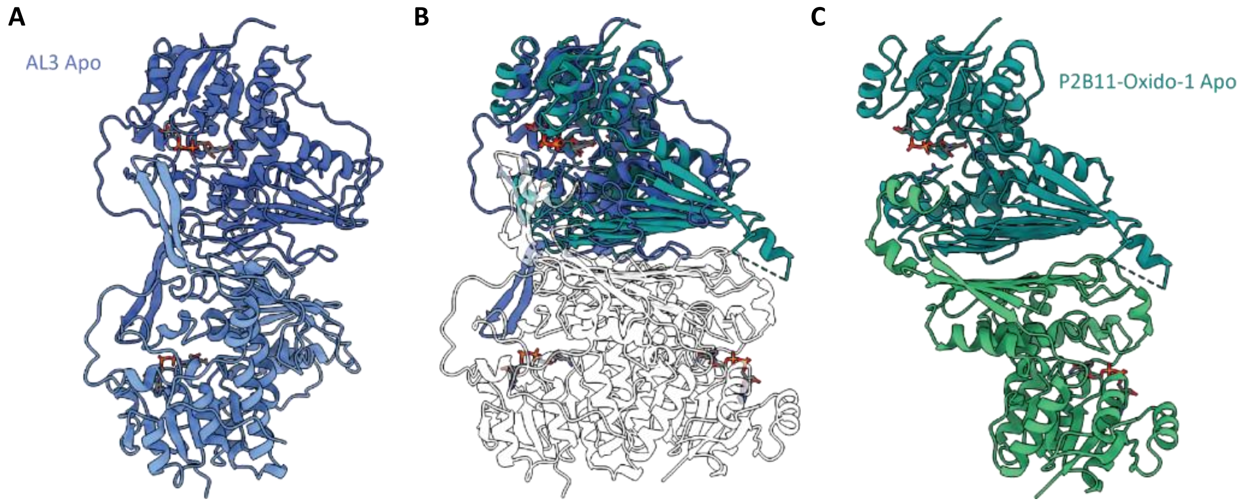

**Figure S111.** Structural alignments of apo AL3 and apo P2B11-Oxido-1 using ChimeraX Matchmaker[42]. (A) Chain A (royal blue) and chain B (cornflower blue) apo AL3 dimer pair. (B) Alignment of apo AL3 chains A (royal blue)/B(transparent) dimer pair with P2B11-Oxido-1 chains J (dark cyan)/L(transparent) RMSD of 1.2 Å across 95 pairs. (C) P2B11-Oxido-1 chain J (dark cyan) chain L (medium sea green) dimer pair.

#### 4. NMR Spectra of synthesized compounds

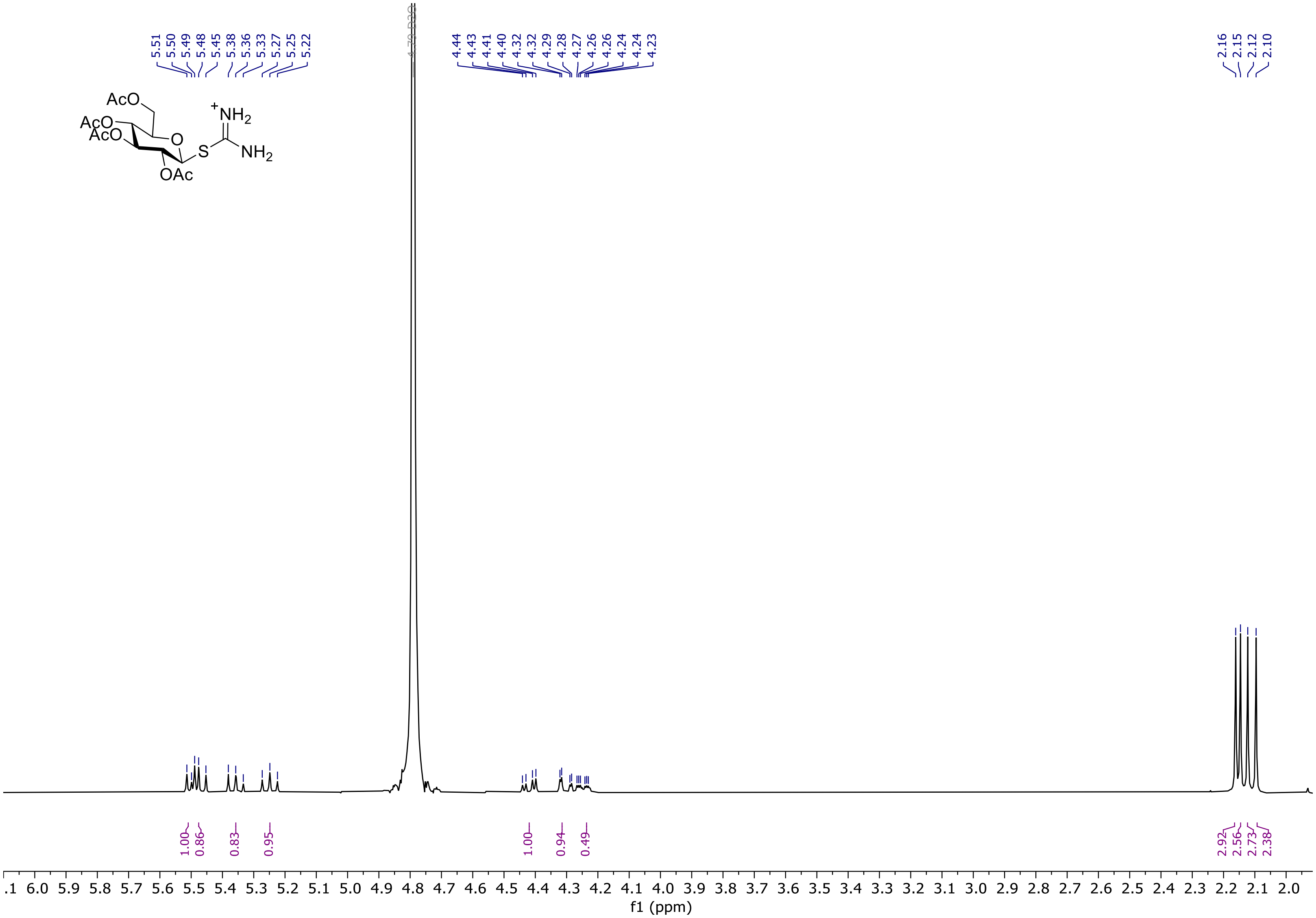

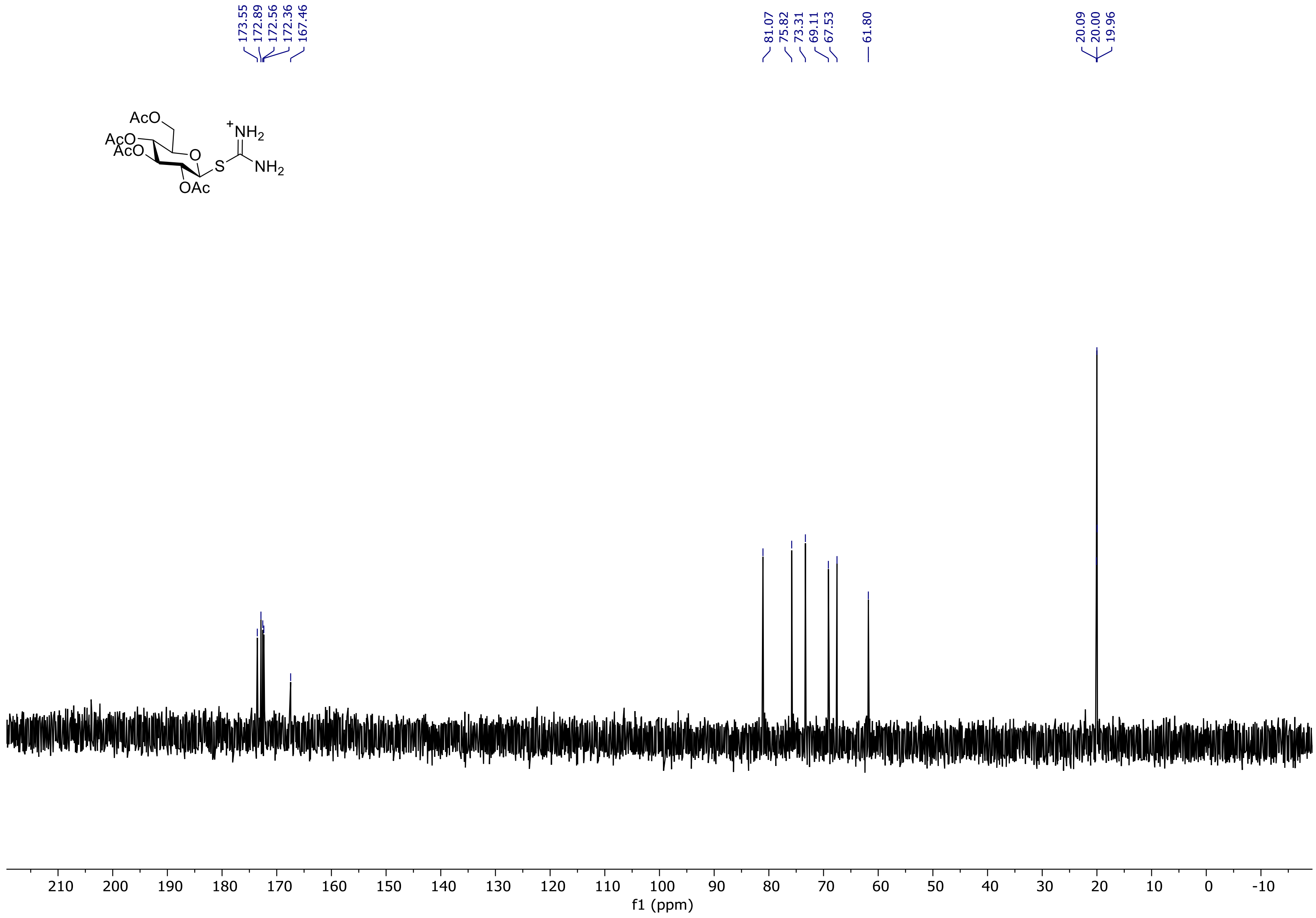
173.55
172.89
172.56
172.36
167.46
81.07
75.82
73.31
69.11
67.53
61.80
20.09
20.00
19.96
AcO
AcO
AcO
O
OAc
S
$^{+}NH_2$
$NH_2$
210 200 190 180 170 160 150 140 130 120 110 100 90 80 70 60 50 40 30 20 10 0 -10
f1 (ppm)

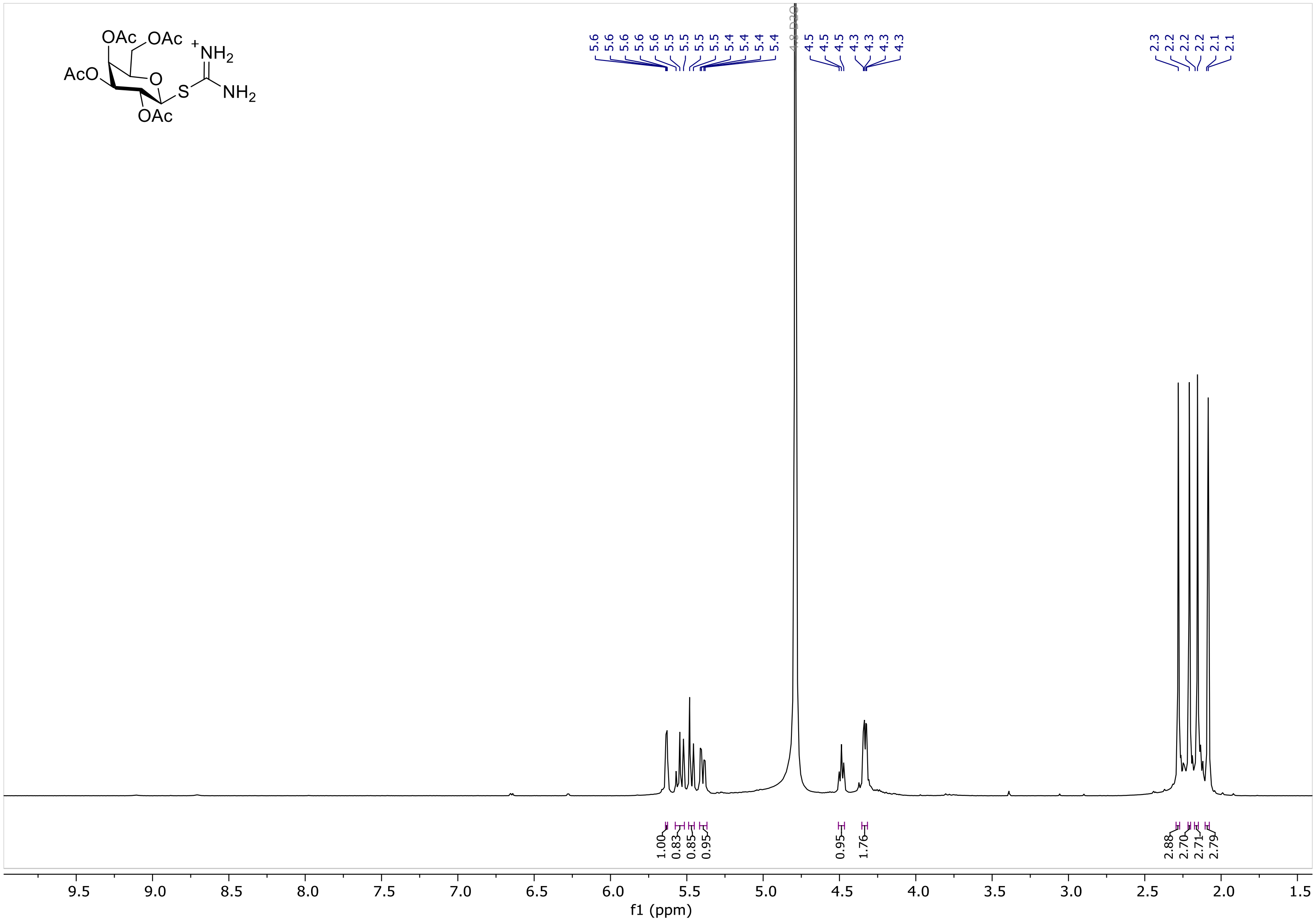
OAc
OAc
AcO
O
S
+NH2
NH2
OAc
5.6 5.6 5.6 5.6 5.5 5.5 5.5 5.5 5.4 5.4 5.4 5.4
4.8 D2O
4.5 4.5 4.5 4.3 4.3 4.3 4.3
2.3 2.2 2.2 2.2 2.1 2.1
1.00 0.83 0.85 0.95
0.95 1.76
2.88 2.70 2.71 2.79
9.5 9.0 8.5 8.0 7.5 7.0 6.5 6.0 5.5 5.0 4.5 4.0 3.5 3.0 2.5 2.0 1.5
f1 (ppm)

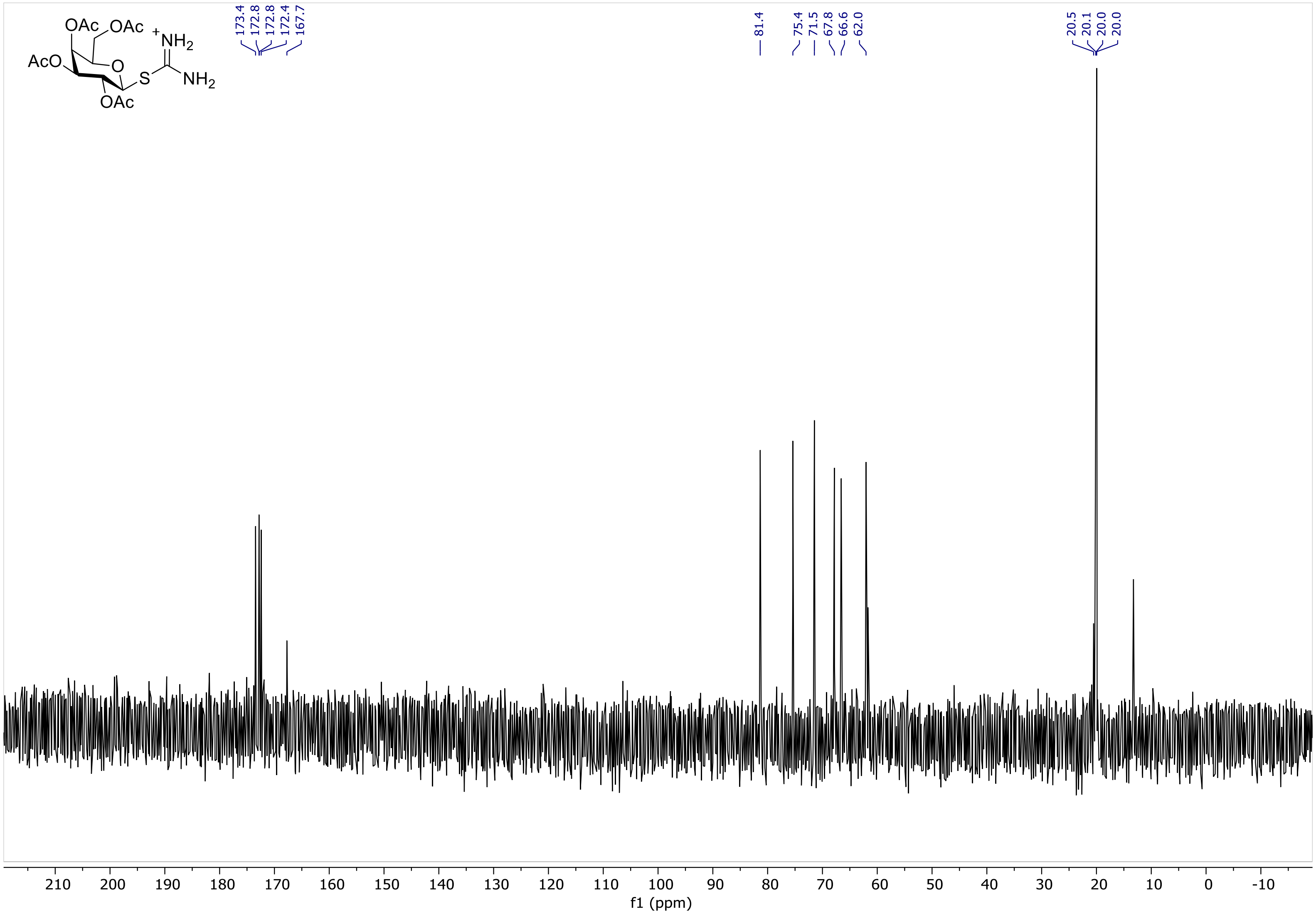
OAc
OAc
AcO
O
S
+NH2
NH2
OAc
173.4
172.8
172.8
172.4
167.7
81.4
75.4
71.5
67.8
66.6
62.0
20.5
20.1
20.0
20.0
210 200 190 180 170 160 150 140 130 120 110 100 90 80 70 60 50 40 30 20 10 0 -10
f1 (ppm)

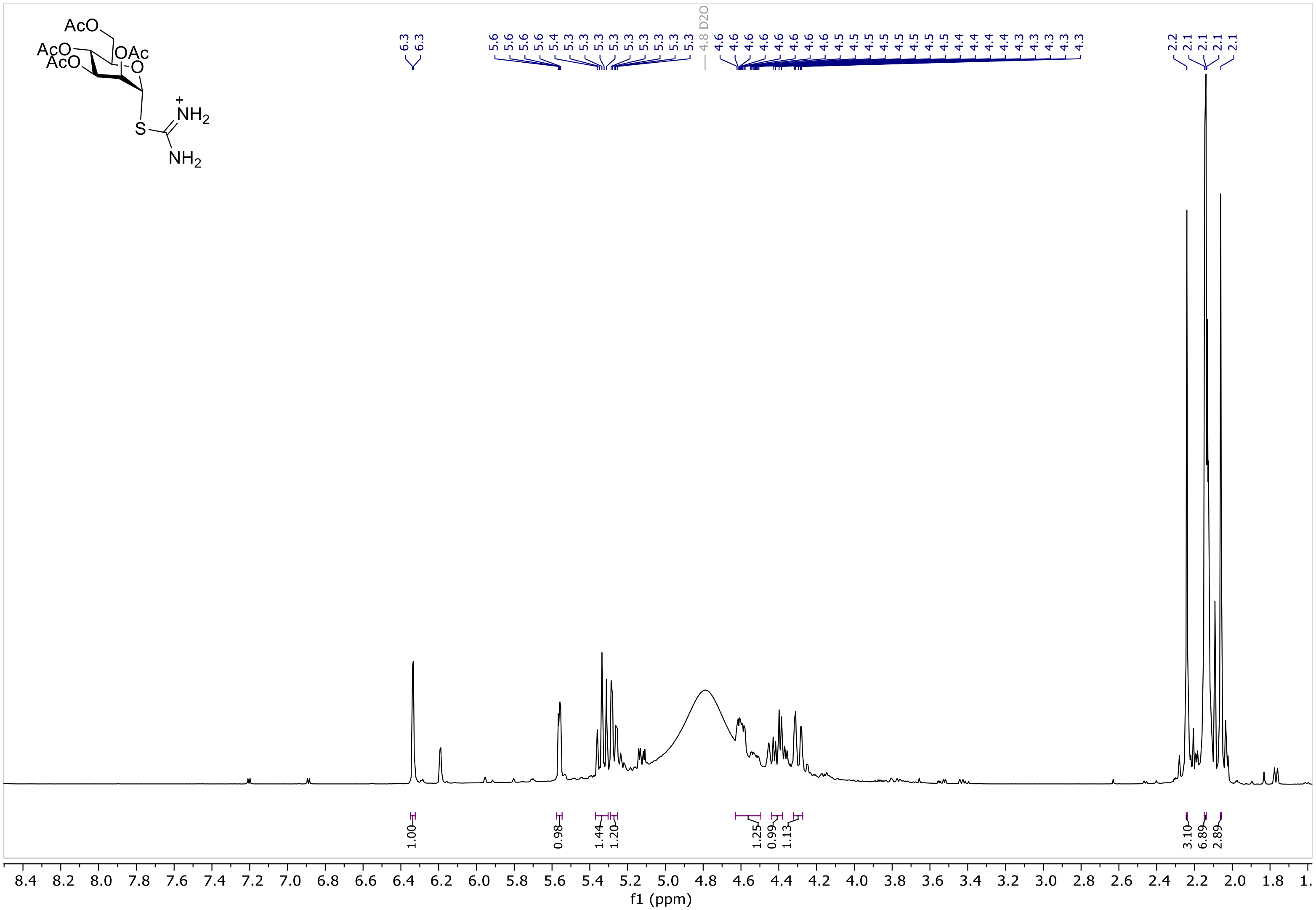
AcO
AcO
AcO
OAc
O
S
NH2
NH2
6.3
6.3
5.6
5.6
5.6
5.6
5.4
5.3
5.3
5.3
5.3
5.3
5.3
5.3
5.3
5.3
4.8 D2O
4.6
4.6
4.6
4.6
4.6
4.6
4.6
4.5
4.5
4.5
4.5
4.5
4.5
4.5
4.5
4.4
4.4
4.4
4.4
4.3
4.3
4.3
4.3
4.3
4.3
2.2
2.1
2.1
2.1
2.1
1.00
0.98
1.44
1.20
1.25
0.99
1.13
3.10
6.89
2.89
8.4 8.2 8.0 7.8 7.6 7.4 7.2 7.0 6.8 6.6 6.4 6.2 6.0 5.8 5.6 5.4 5.2 5.0 4.8 4.6 4.4 4.2 4.0 3.8 3.6 3.4 3.2 3.0 2.8 2.6 2.4 2.2 2.0 1.8 1.
f1 (ppm)

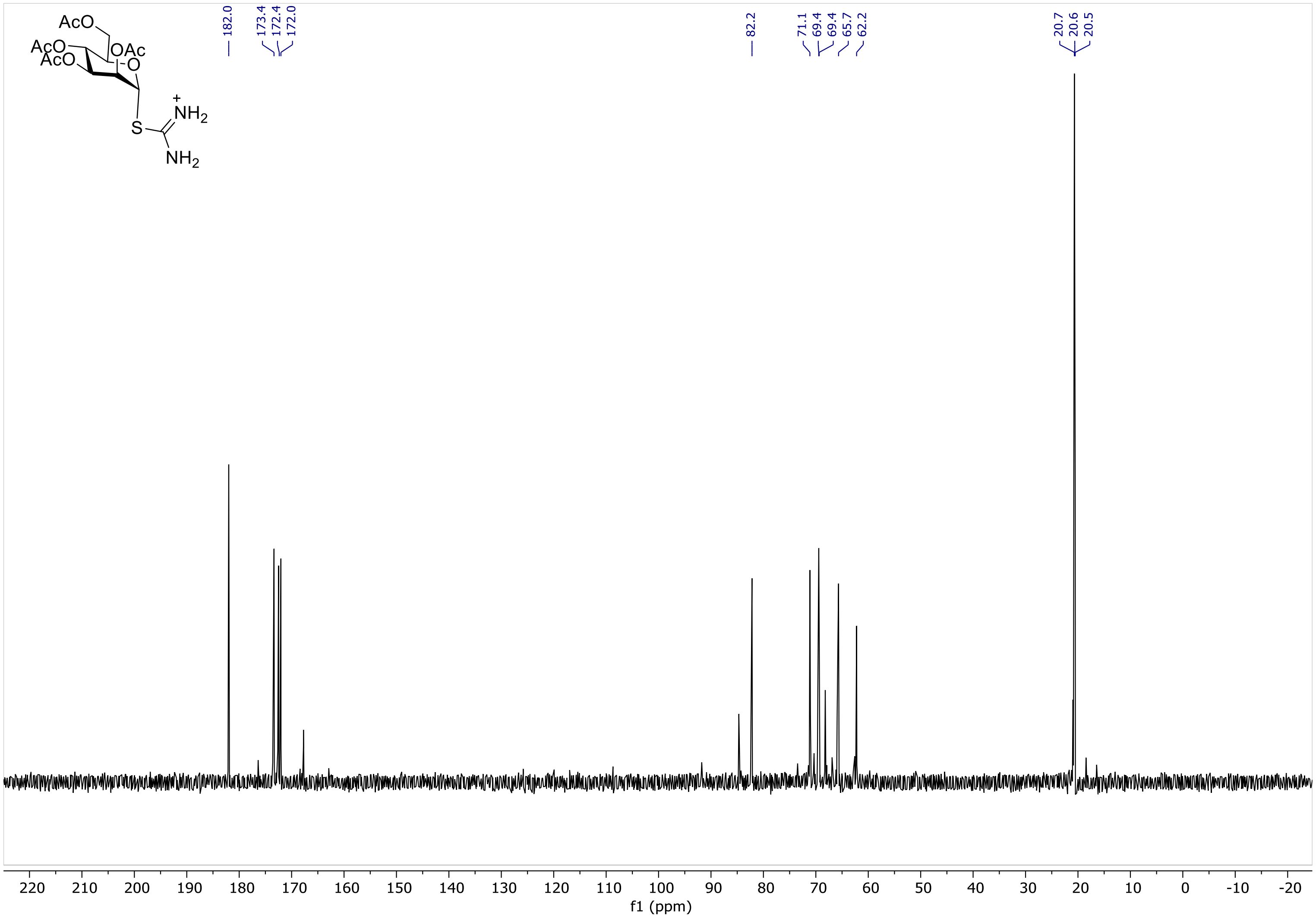
AcO
AcO
AcO
OAc
O
S
NH2
+
NH2
182.0
173.4
172.4
172.0
82.2
71.1
69.4
69.4
65.7
62.2
20.7
20.6
20.5
220 210 200 190 180 170 160 150 140 130 120 110 100 90 80 70 60 50 40 30 20 10 0 -10 -20
f1 (ppm)

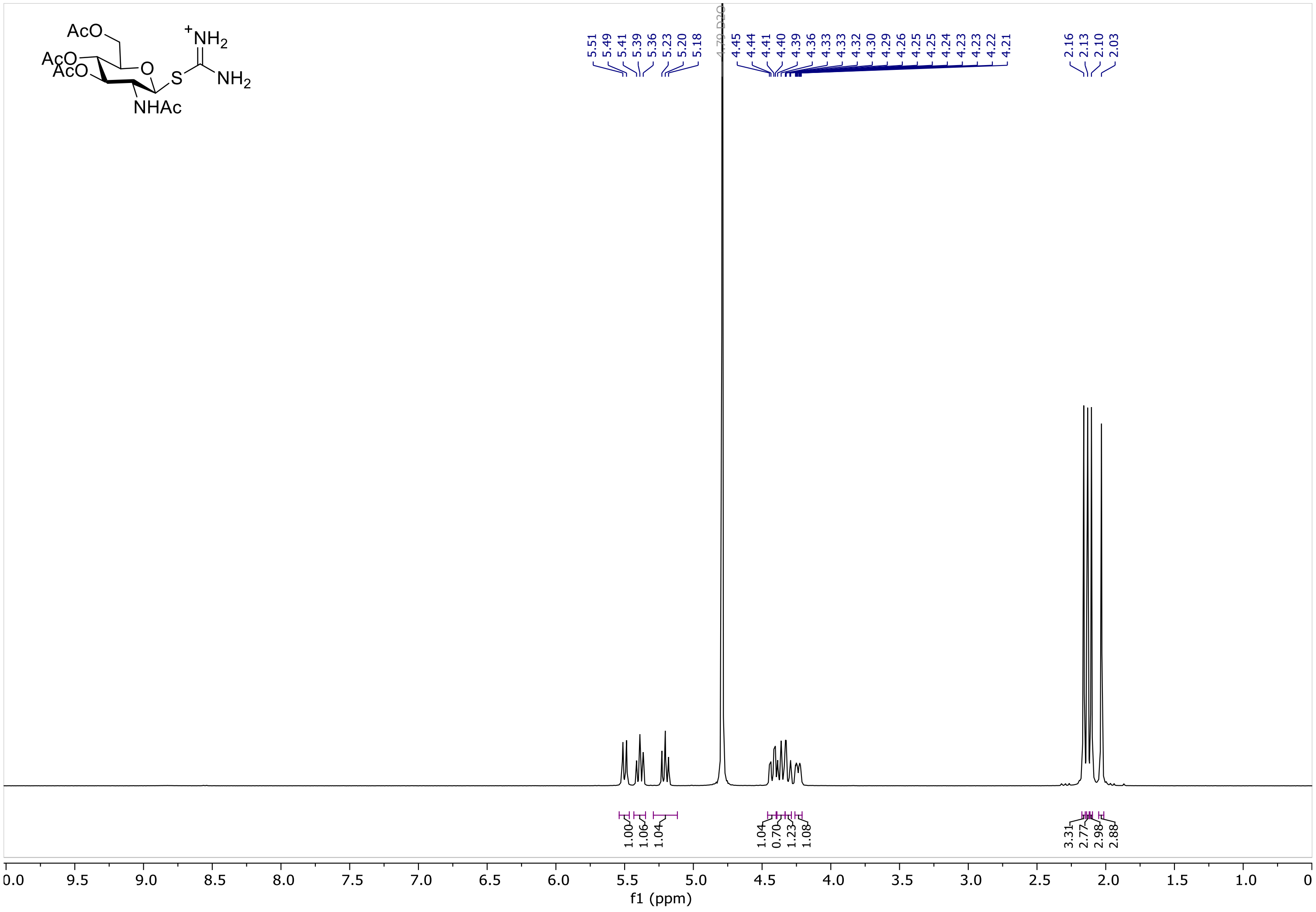

AcO
AcO
AcO
O
S
+NH2
NH2
NHAc
5.51
5.49
5.41
5.39
5.36
5.23
5.20
5.18
4.79 D2O
4.45
4.44
4.41
4.40
4.39
4.36
4.33
4.33
4.32
4.30
4.29
4.26
4.25
4.25
4.24
4.23
4.23
4.22
4.21
2.16
2.13
2.10
2.03
1.00
1.06
1.04
1.04
0.70
1.23
1.08
3.31
2.77
2.98
2.88
f1 (ppm)

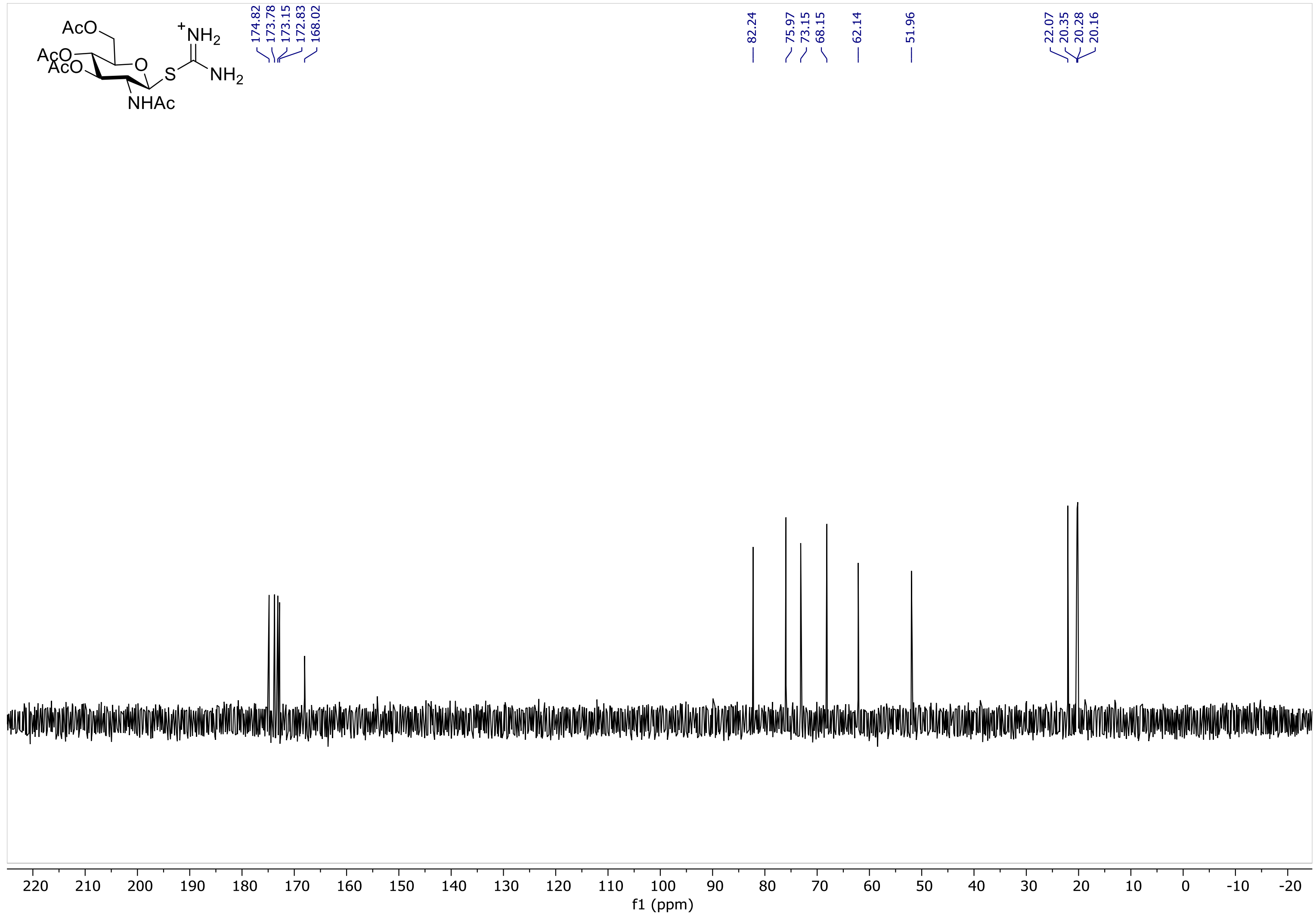

AcO
AcO
AcO
O
S
+NH2
NH2
NHAc
174.82
173.78
173.15
172.83
168.02
82.24
75.97
73.15
68.15
62.14
51.96
22.07
20.35
20.28
20.16
220 210 200 190 180 170 160 150 140 130 120 110 100 90 80 70 60 50 40 30 20 10 0 -10 -20
f1 (ppm)

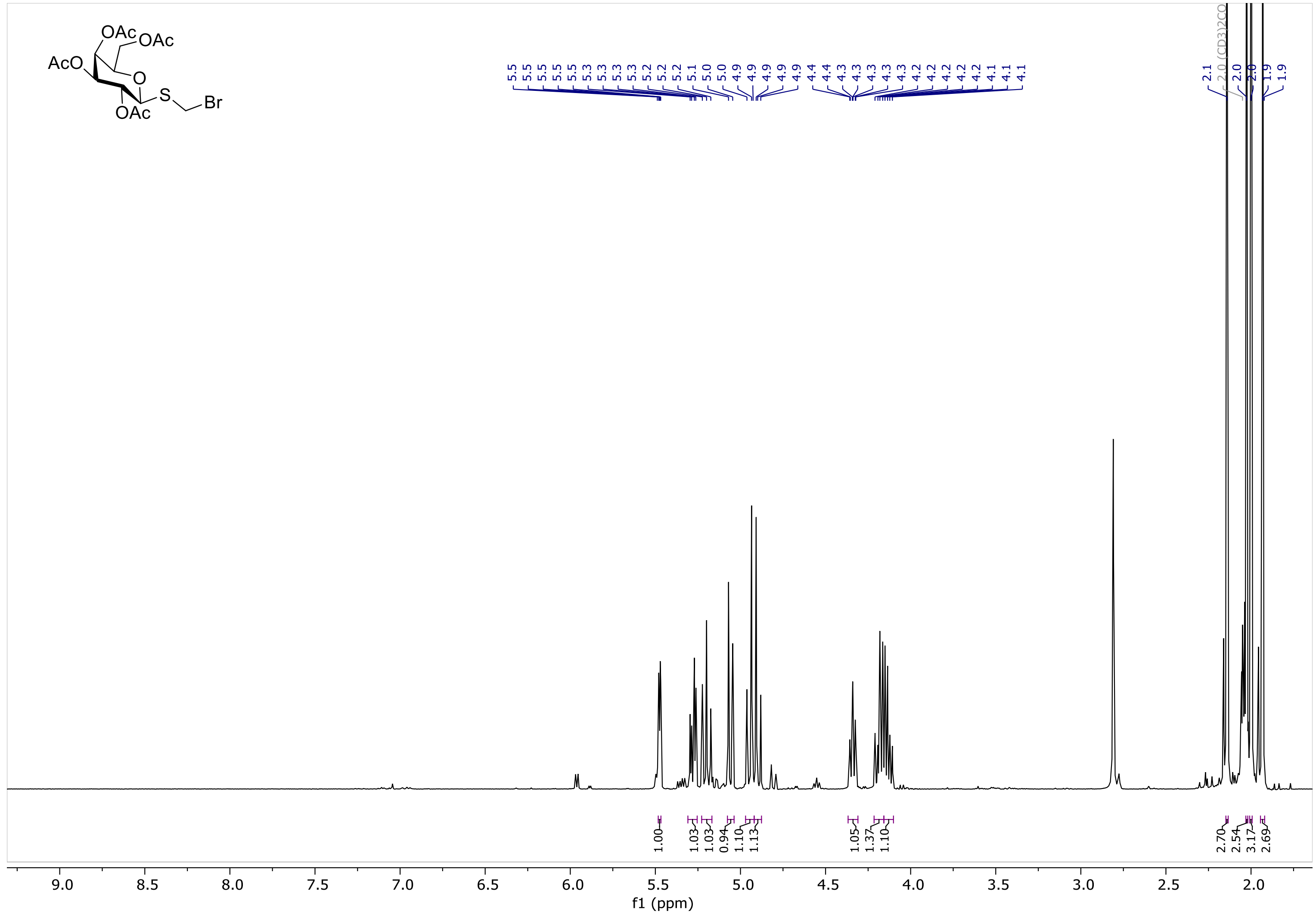
AcO
OAc
OAc
O
S
Br
OAc
f1 (ppm)

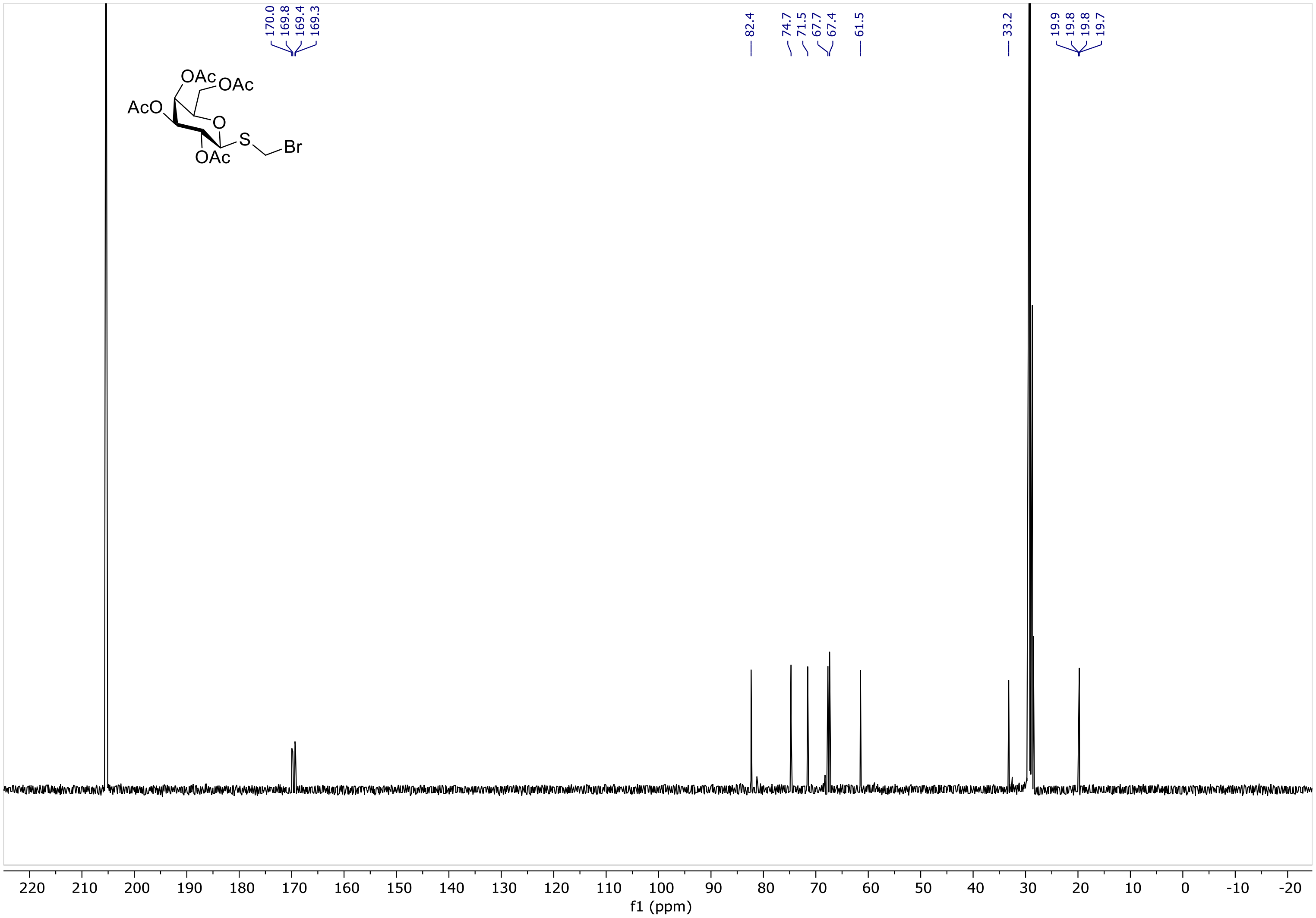

170.0
169.8
169.4
169.3
82.4
74.7
71.5
67.7
67.4
61.5
33.2
19.9
19.8
19.8
19.7
OAc
OAc
AcO
O
S
Br
OAc
220 210 200 190 180 170 160 150 140 130 120 110 100 90 80 70 60 50 40 30 20 10 0 -10 -20
f1 (ppm)

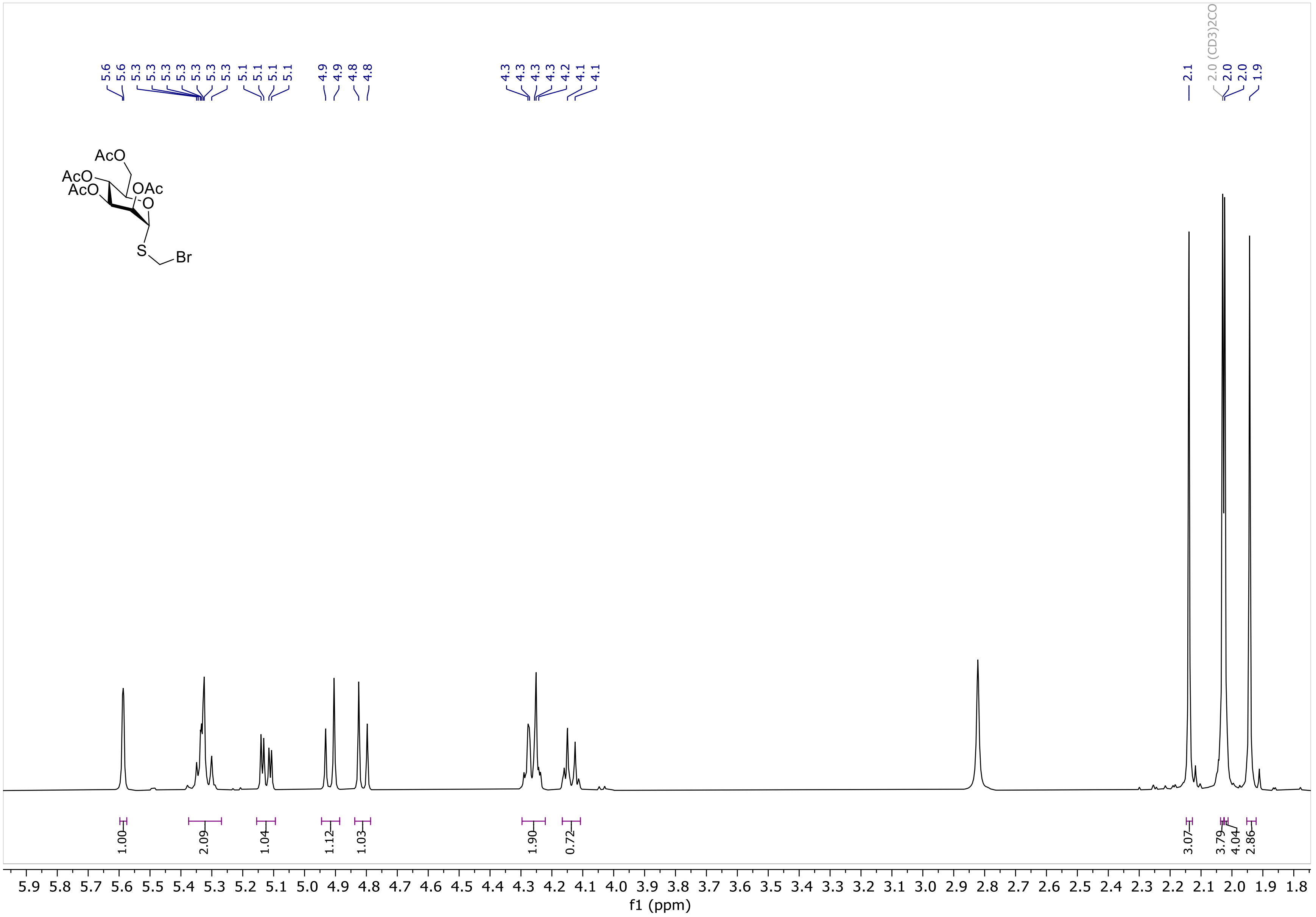
AcO
AcO
AcO
OAc
O
S
Br
2.0 (CD3)2CO
1.00
2.09
1.04
1.12
1.03
1.90
0.72
3.07
3.79
4.04
2.86
f1 (ppm)

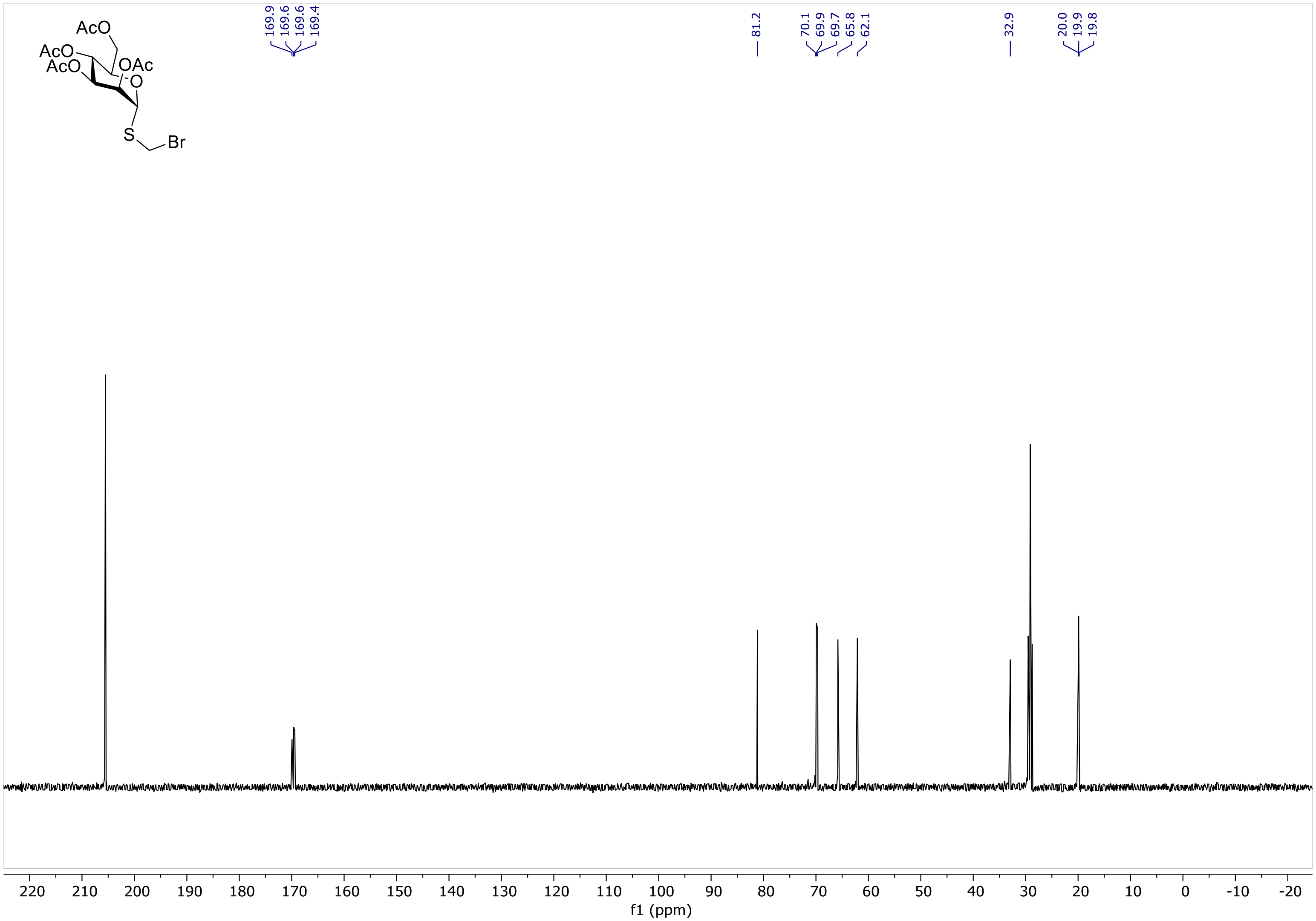
AcO
AcO
AcO
OAc
O
S
Br
169.9
169.6
169.6
169.4
81.2
70.1
69.9
69.7
65.8
62.1
32.9
20.0
19.9
19.8
220 210 200 190 180 170 160 150 140 130 120 110 100 90 80 70 60 50 40 30 20 10 0 -10 -20
f1 (ppm)

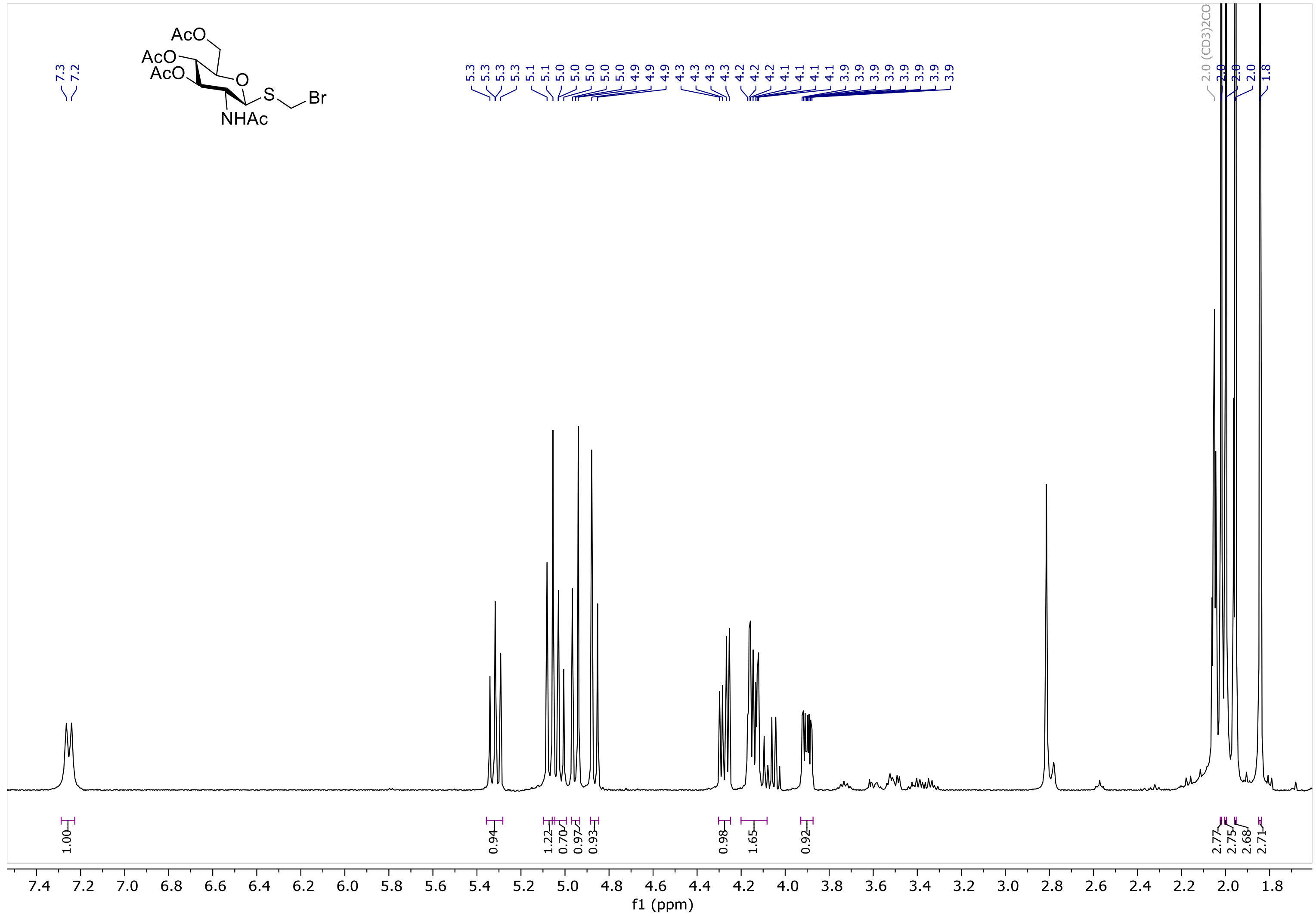
AcO
AcO
AcO
O
S
Br
NHAc
7.3
7.2
5.3
5.3
5.3
5.3
5.1
5.1
5.0
5.0
5.0
5.0
5.0
4.9
4.9
4.9
4.3
4.3
4.3
4.3
4.2
4.2
4.2
4.1
4.1
4.1
3.9
3.9
3.9
3.9
3.9
3.9
3.9
2.0 (CD3)2CO
2.0
2.0
2.0
1.8
1.00
0.94
1.22
0.70
0.97
0.93
0.98
1.65
0.92
2.77
2.75
2.68
2.71
7.4 7.2 7.0 6.8 6.6 6.4 6.2 6.0 5.8 5.6 5.4 5.2 5.0 4.8 4.6 4.4 4.2 4.0 3.8 3.6 3.4 3.2 3.0 2.8 2.6 2.4 2.2 2.0 1.8
f1 (ppm)

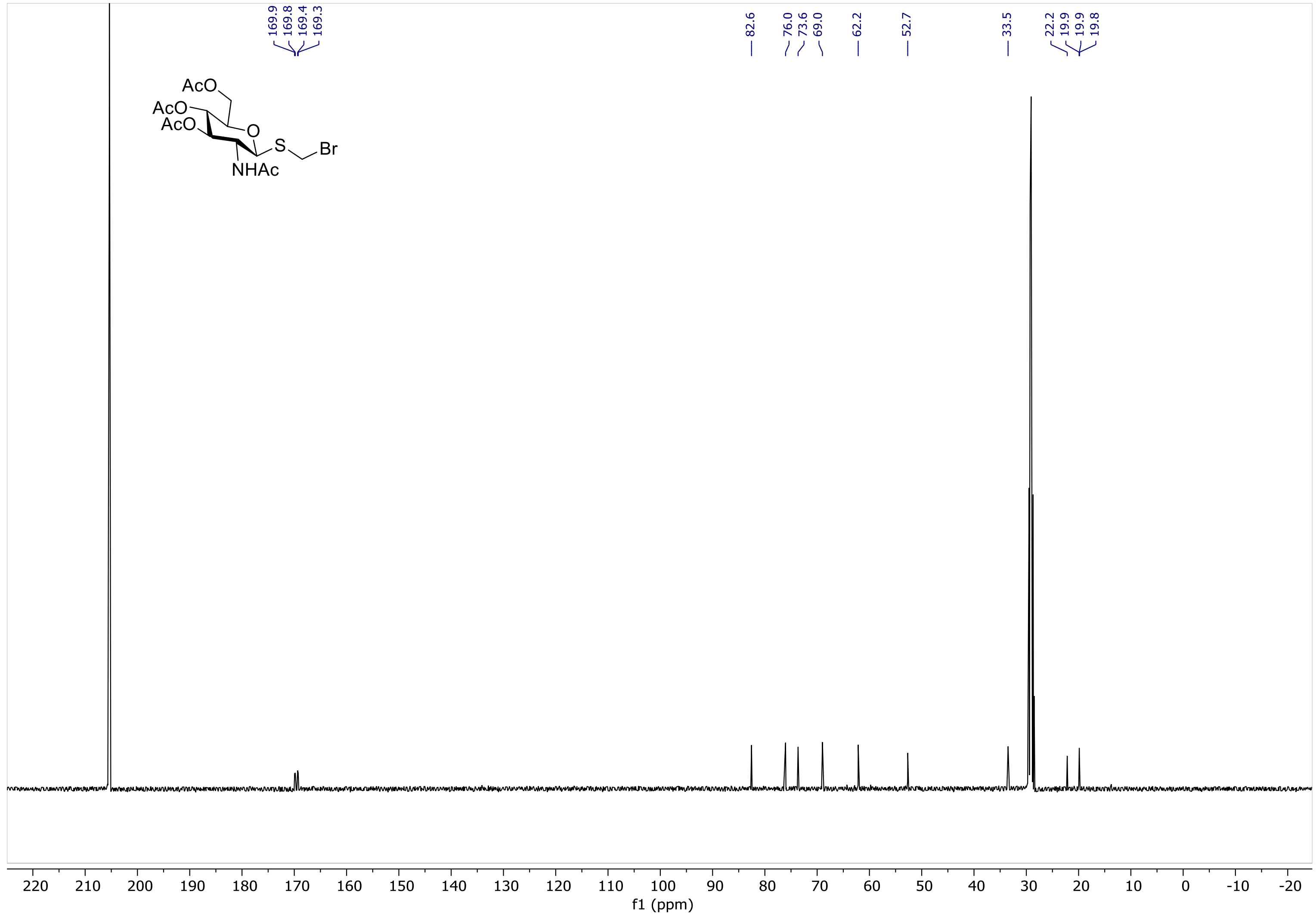

AcO
AcO
AcO
O
S
Br
NHAc
169.9
169.8
169.4
169.3
82.6
76.0
73.6
69.0
62.2
52.7
33.5
22.2
19.9
19.9
19.8
220 210 200 190 180 170 160 150 140 130 120 110 100 90 80 70 60 50 40 30 20 10 0 -10 -20
f1 (ppm)

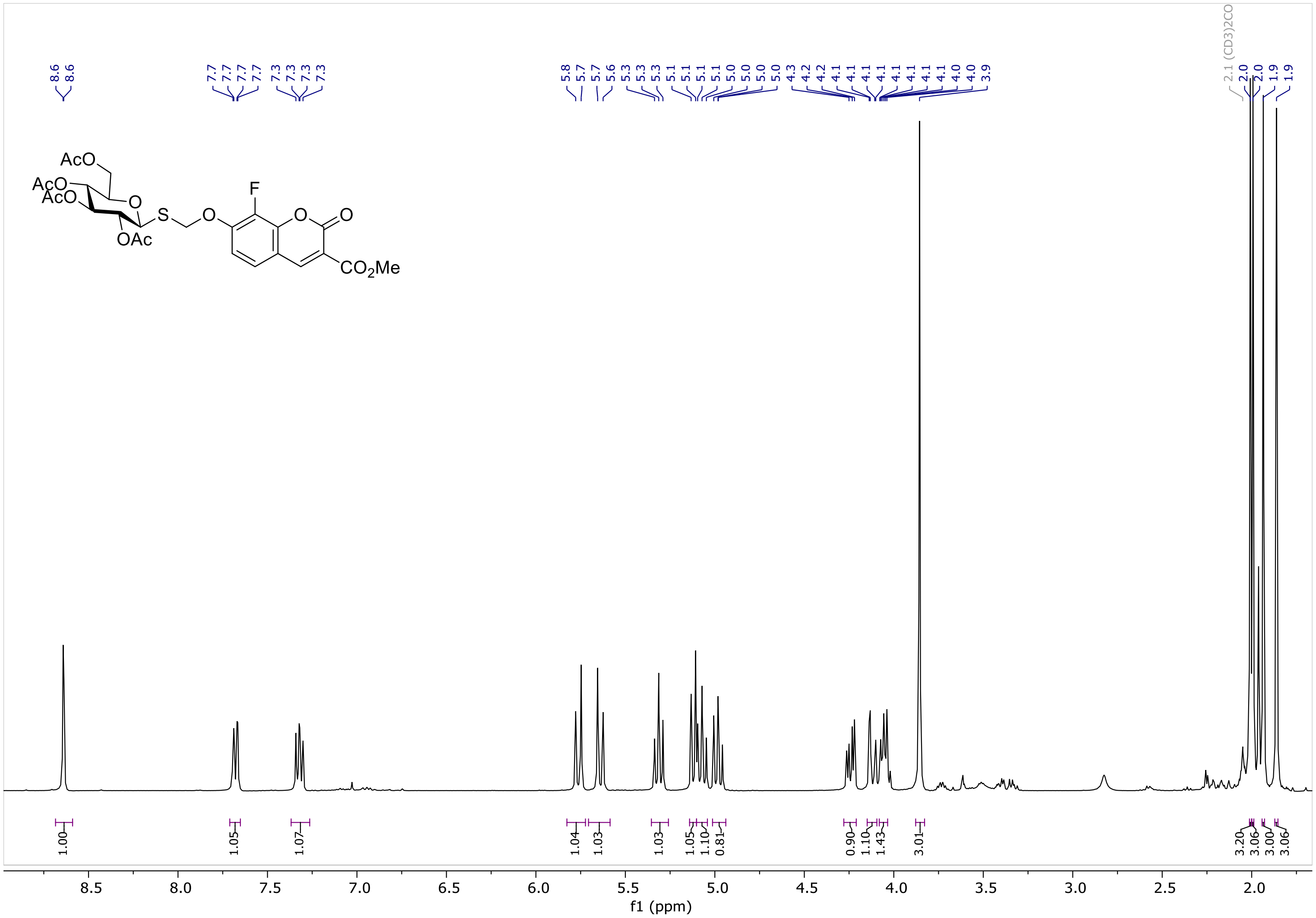

8.6
8.6
7.7
7.7
7.7
7.7
7.3
7.3
7.3
7.3
5.8
5.7
5.7
5.6
5.3
5.3
5.3
5.1
5.1
5.1
5.1
5.0
5.0
5.0
5.0
4.3
4.2
4.2
4.1
4.1
4.1
4.1
4.1
4.1
4.1
4.1
4.0
4.0
3.9
2.1 (CD3)2CO
2.0
2.0
2.0
1.9
1.9
AcO
AcO
AcO
O
OAc
S
O
F
O
O
CO2Me
1.00
1.05
1.07
1.04
1.03
1.03
1.05
1.10
0.81
0.90
1.10
1.43
3.01
3.20
3.06
3.00
3.06
8.5
8.0
7.5
7.0
6.5
6.0
5.5
5.0
4.5
4.0
3.5
3.0
2.5
2.0
f1 (ppm)

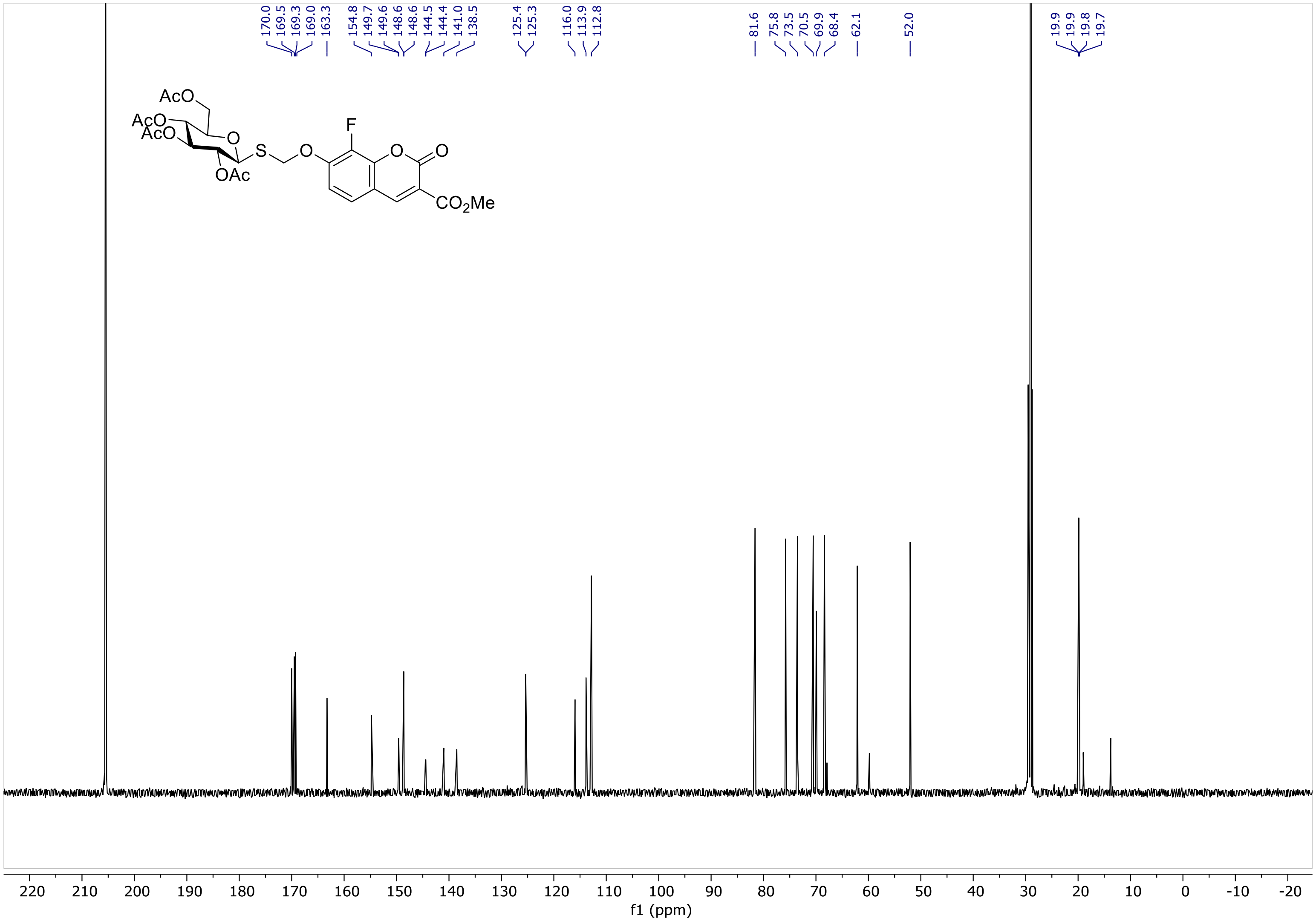
170.0
169.5
169.3
169.0
163.3
154.8
149.7
149.6
148.6
148.6
144.5
144.4
141.0
138.5
125.4
125.3
116.0
113.9
112.8
81.6
75.8
73.5
70.5
69.9
68.4
62.1
52.0
19.9
19.9
19.8
19.7
AcO
AcO
AcO
OAc
O
S
O
F
O
O
CO2Me
f1 (ppm)

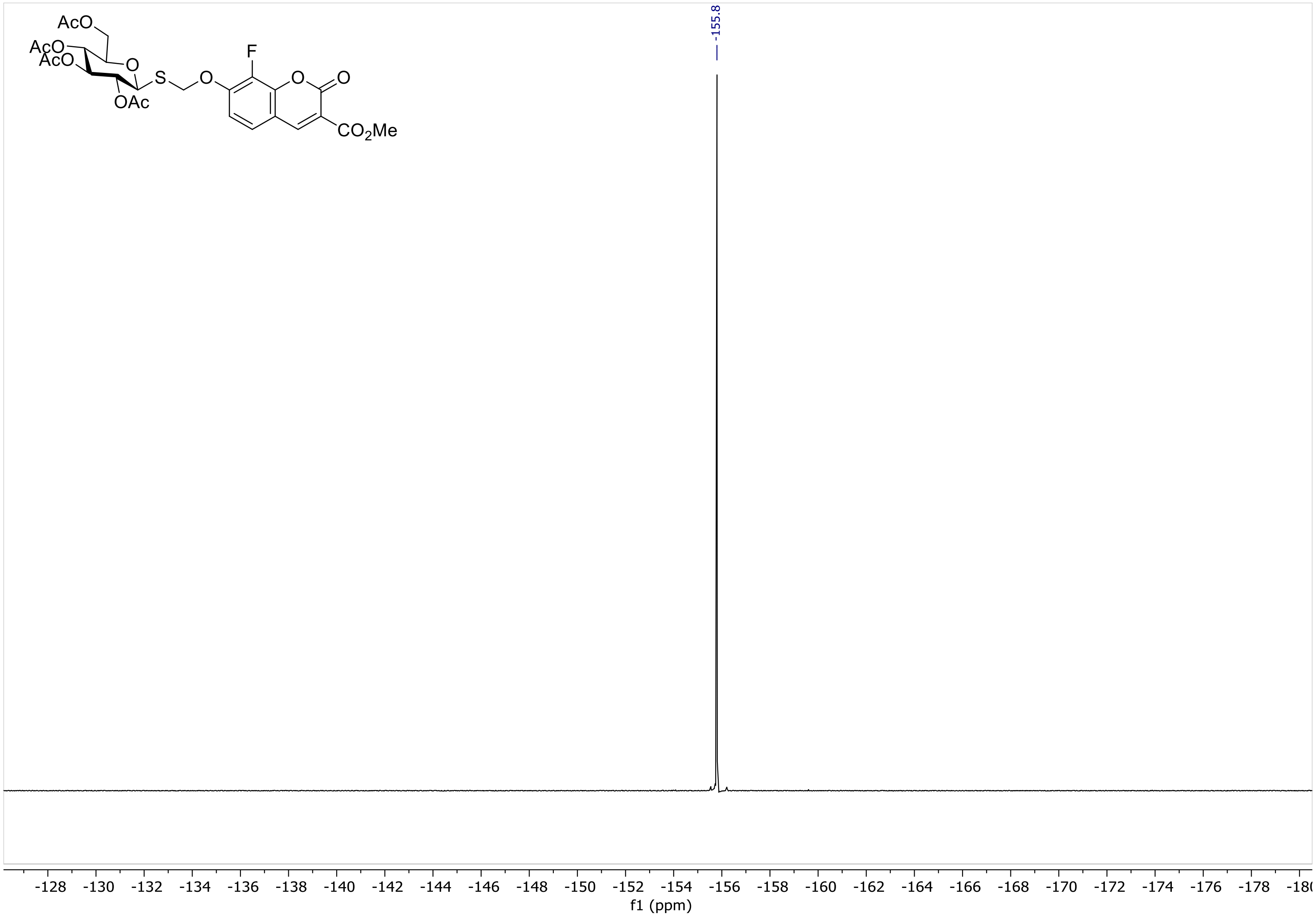
AcO
AcO
AcO
O
S
O
F
O
O
OAc
CO2Me
-155.8
-128
-130
-132
-134
-136
-138
-140
-142
-144
-146
-148
-150
-152
-154
-156
-158
-160
-162
-164
-166
-168
-170
-172
-174
-176
-178
f1 (ppm)

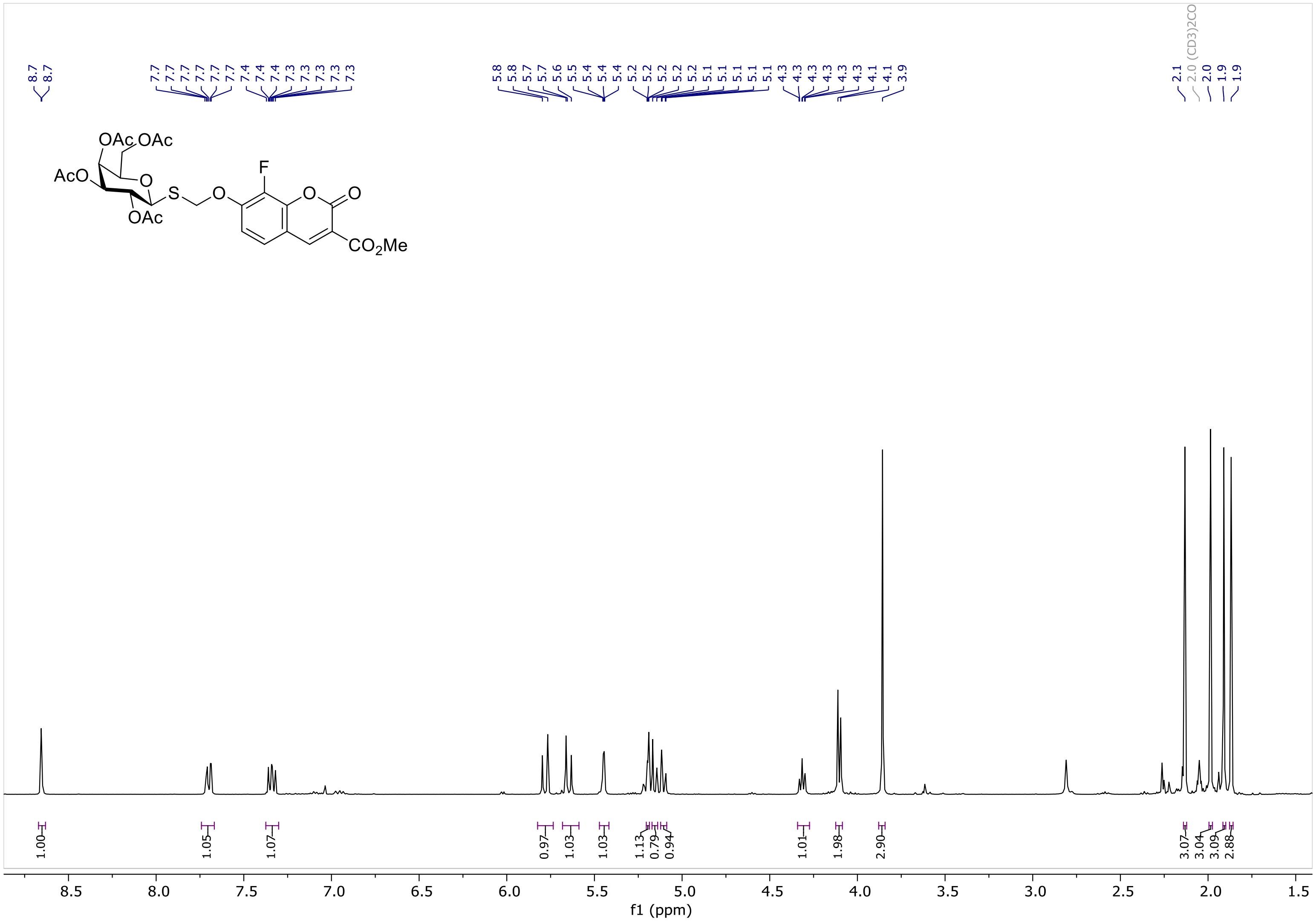

OAc
OAc
AcO
O
S
O
F
O
O
OAc
CO2Me
8.7
8.7
7.7
7.4
7.3
5.8
5.7
5.6
5.5
5.4
5.2
5.1
4.3
4.1
3.9
2.1
2.0 (CD3)2CO
2.0
1.9
1.00
1.05
1.07
0.97
1.03
1.03
1.13
0.79
0.94
1.01
1.98
2.90
3.07
3.04
3.09
2.88
8.5
8.0
7.5
7.0
6.5
6.0
5.5
5.0
4.5
4.0
3.5
3.0
2.5
2.0
1.5
f1 (ppm)

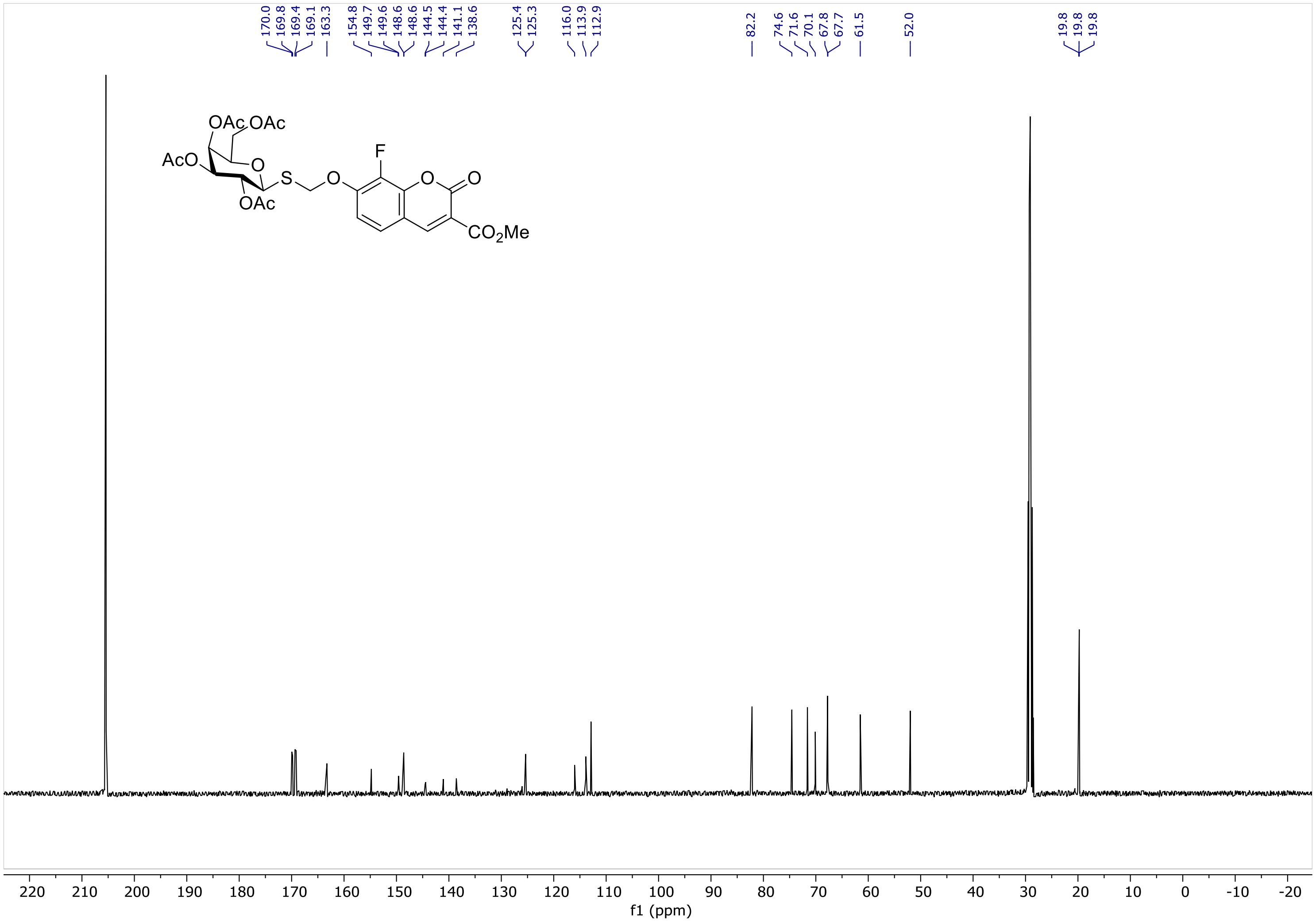
170.0
169.8
169.4
169.1
163.3
154.8
149.7
149.6
148.6
148.6
144.5
144.4
141.1
138.6
125.4
125.3
116.0
113.9
112.9
82.2
74.6
71.6
70.1
67.8
67.7
61.5
52.0
19.8
19.8
19.8
OAc
OAc
AcO
O
S
O
F
O
O
OAc
$CO_2Me$
220
210
200
190
180
170
160
150
140
130
120
110
100
90
80
70
60
50
40
30
20
10
0
-10
-20
f1 (ppm)

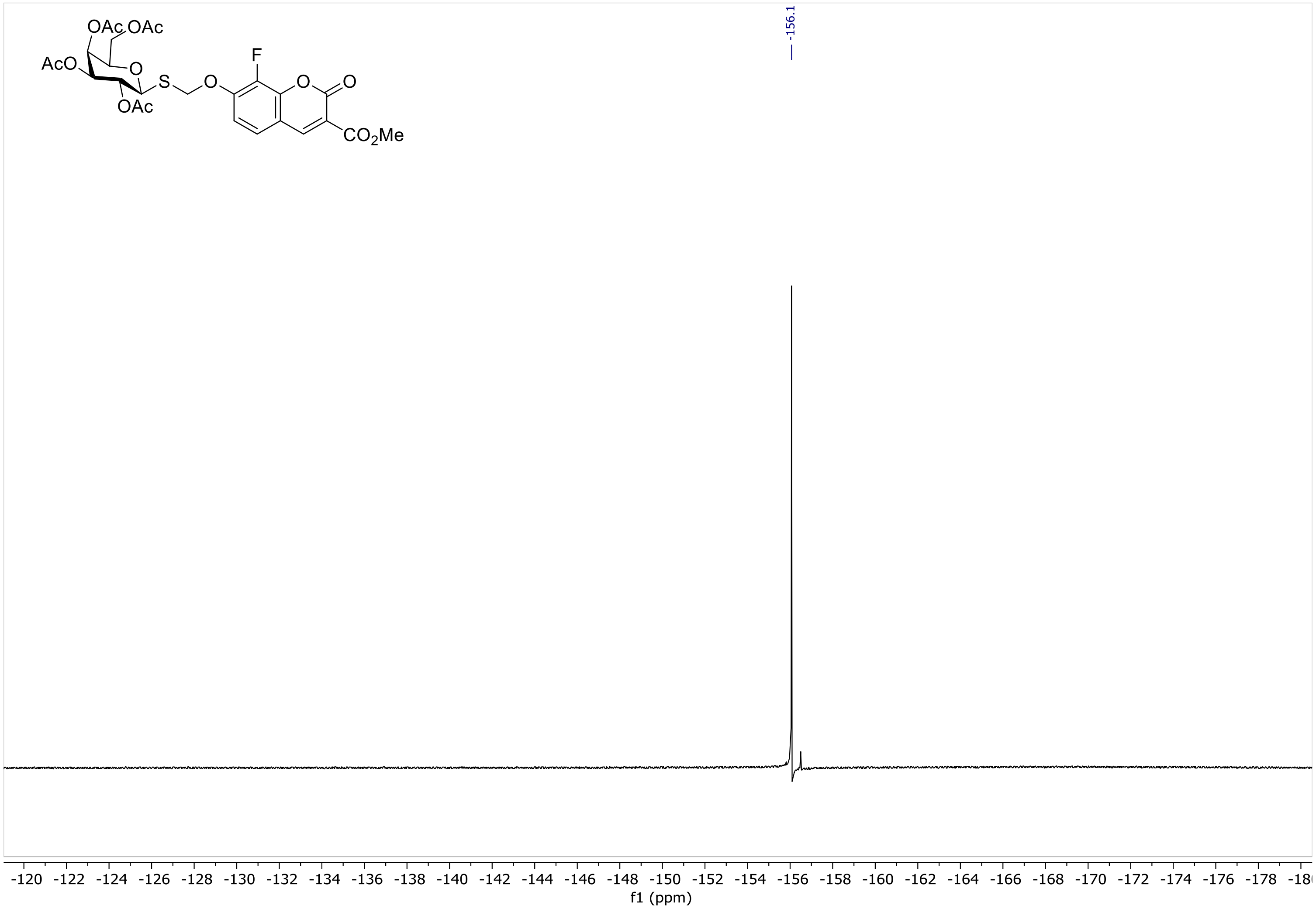

OAc OAc
AcO
O
S
O
F
O
O
OAc
CO2Me
-156.1
-120 -122 -124 -126 -128 -130 -132 -134 -136 -138 -140 -142 -144 -146 -148 -150 -152 -154 -156 -158 -160 -162 -164 -166 -168 -170 -172 -174 -176 -178
f1 (ppm)

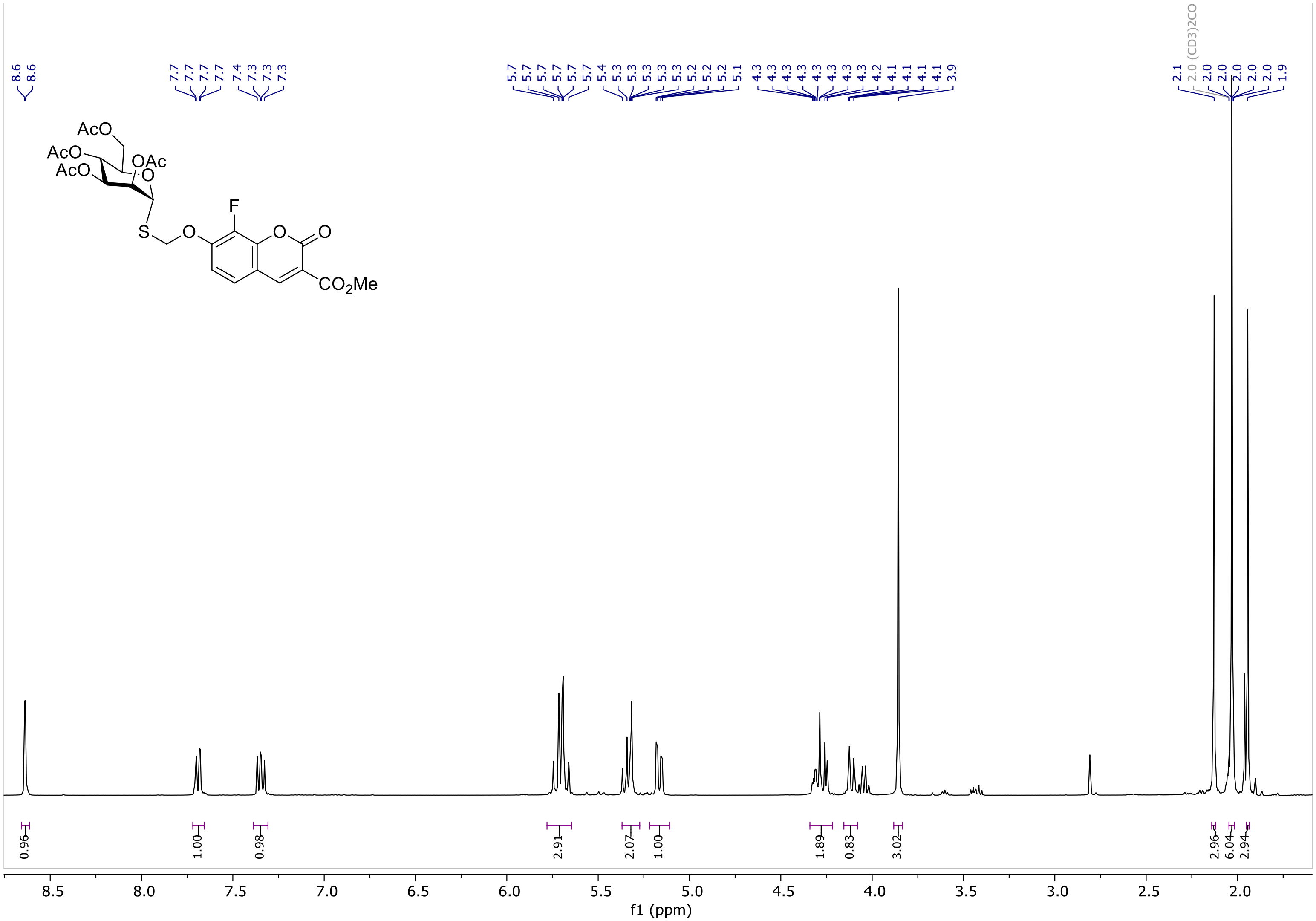
AcO
AcO
AcO
OAc
O
F
S
O
O
O
CO2Me
2.0 (CD3)2CO
0.96
1.00
0.98
2.91
2.07
1.00
1.89
0.83
3.02
2.96
6.04
2.94
8.5
8.0
7.5
7.0
6.5
6.0
5.5
5.0
4.5
4.0
3.5
3.0
2.5
2.0
f1 (ppm)

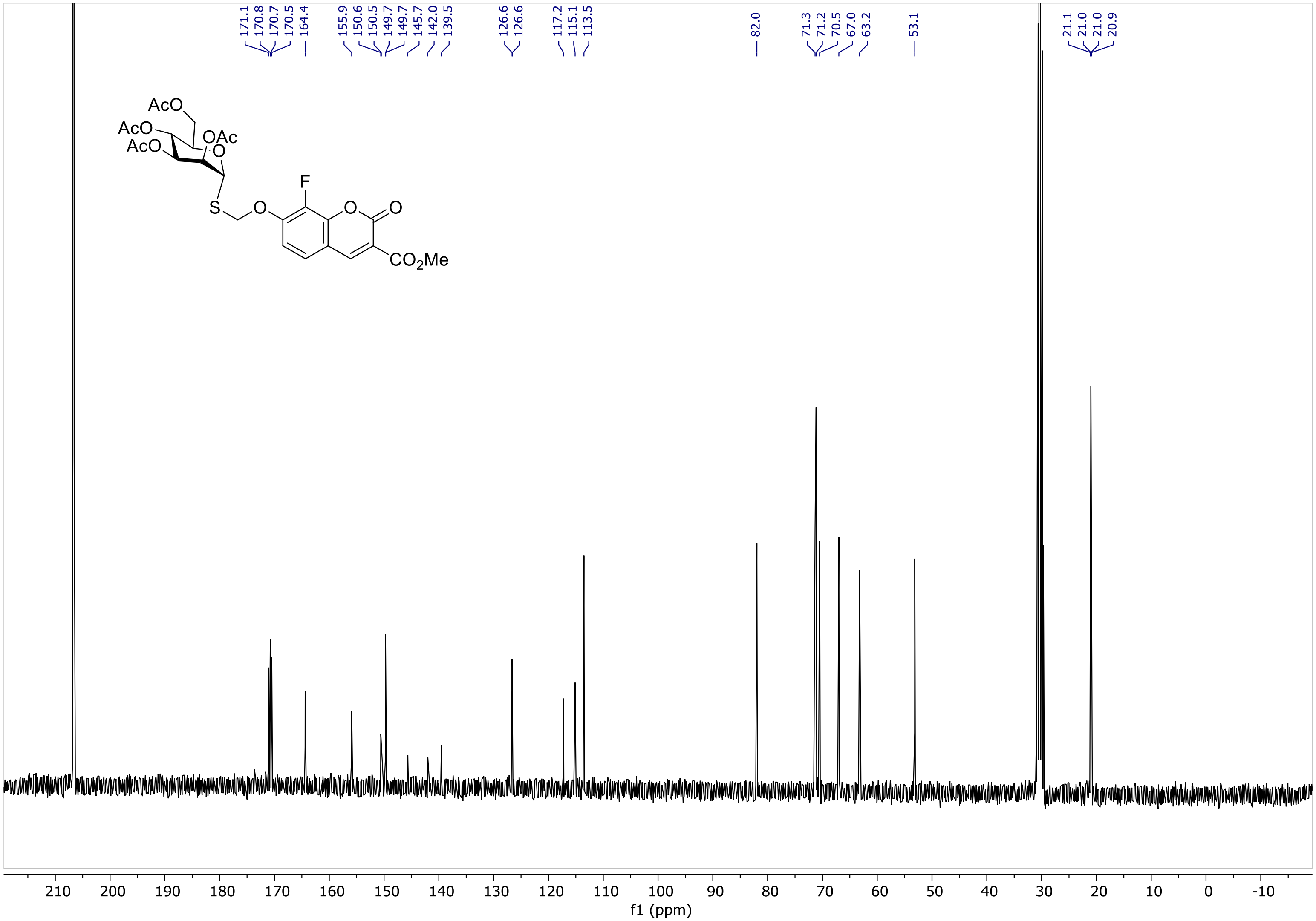

171.1
170.8
170.7
170.5
164.4
155.9
150.6
150.5
149.7
149.7
145.7
142.0
139.5
126.6
126.6
117.2
115.1
113.5
82.0
71.3
71.2
70.5
67.0
63.2
53.1
21.1
21.0
21.0
20.9
AcO
AcO
AcO
OAc
O
F
S
O
O
O
CO2Me
210
200
190
180
170
160
150
140
130
120
110
100
90
80
70
60
50
40
30
20
10
0
-10
f1 (ppm)

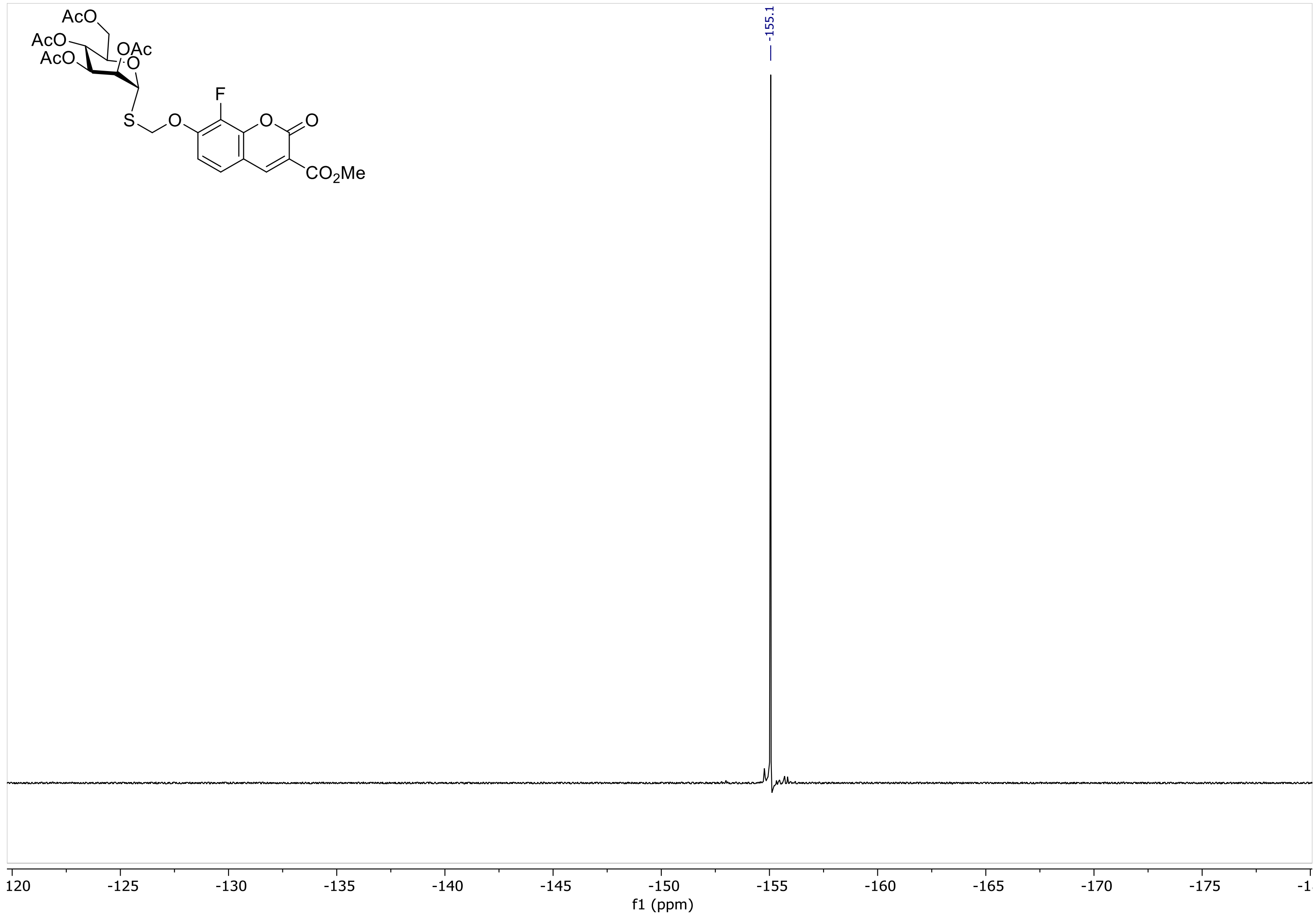

-155.1
120
-125
-130
-135
-140
-145
-150
-155
-160
-165
-170
-175
-1
f1 (ppm)
AcO
AcO
AcO
OAc
O
S
O
F
O
O
CO2Me

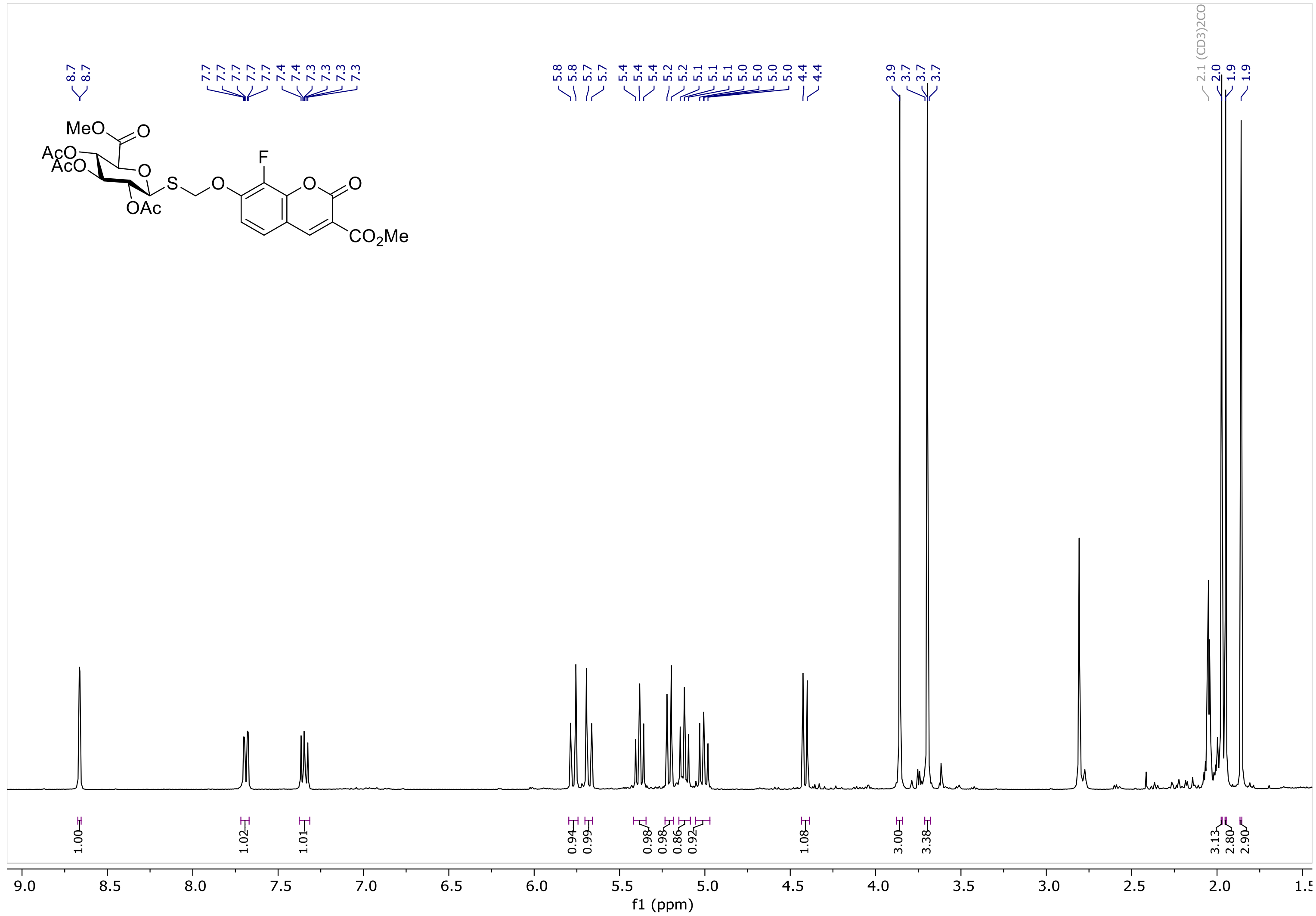

8.7
8.7
7.7
7.7
7.7
7.7
7.7
7.4
7.4
7.3
7.3
7.3
7.3
5.8
5.8
5.7
5.7
5.4
5.4
5.4
5.2
5.2
5.1
5.1
5.1
5.0
5.0
5.0
4.4
4.4
3.9
3.7
3.7
3.7
2.1 (CD3)2CO
2.0
1.9
1.9
MeO
O
AcO
AcO
O
S
O
F
O
O
OAc
CO2Me
1.00
1.02
1.01
0.94
0.99
0.98
0.98
0.86
0.92
1.08
3.00
3.38
3.13
2.80
2.90
9.0
8.5
8.0
7.5
7.0
6.5
6.0
5.5
5.0
4.5
4.0
3.5
3.0
2.5
2.0
1.5
f1 (ppm)

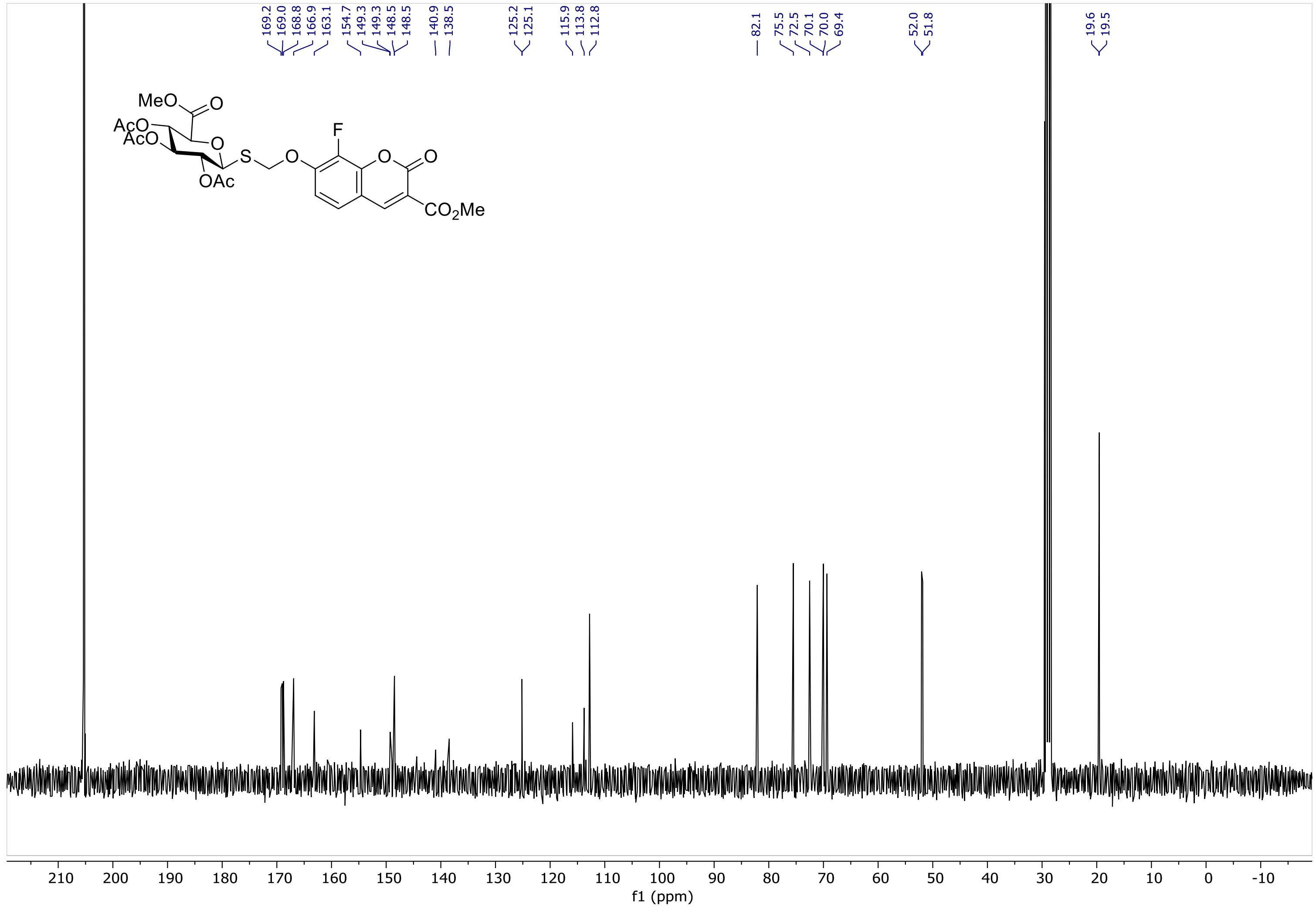

169.2
169.0
168.8
166.9
163.1
154.7
149.3
149.3
148.5
148.5
140.9
138.5
125.2
125.1
115.9
113.8
112.8
82.1
75.5
72.5
70.1
70.0
69.4
52.0
51.8
19.6
19.5
MeO
O
AcO
AcO
O
S
O
F
O
O
OAc
CO2Me
210 200 190 180 170 160 150 140 130 120 110 100 90 80 70 60 50 40 30 20 10 0 -10
f1 (ppm)

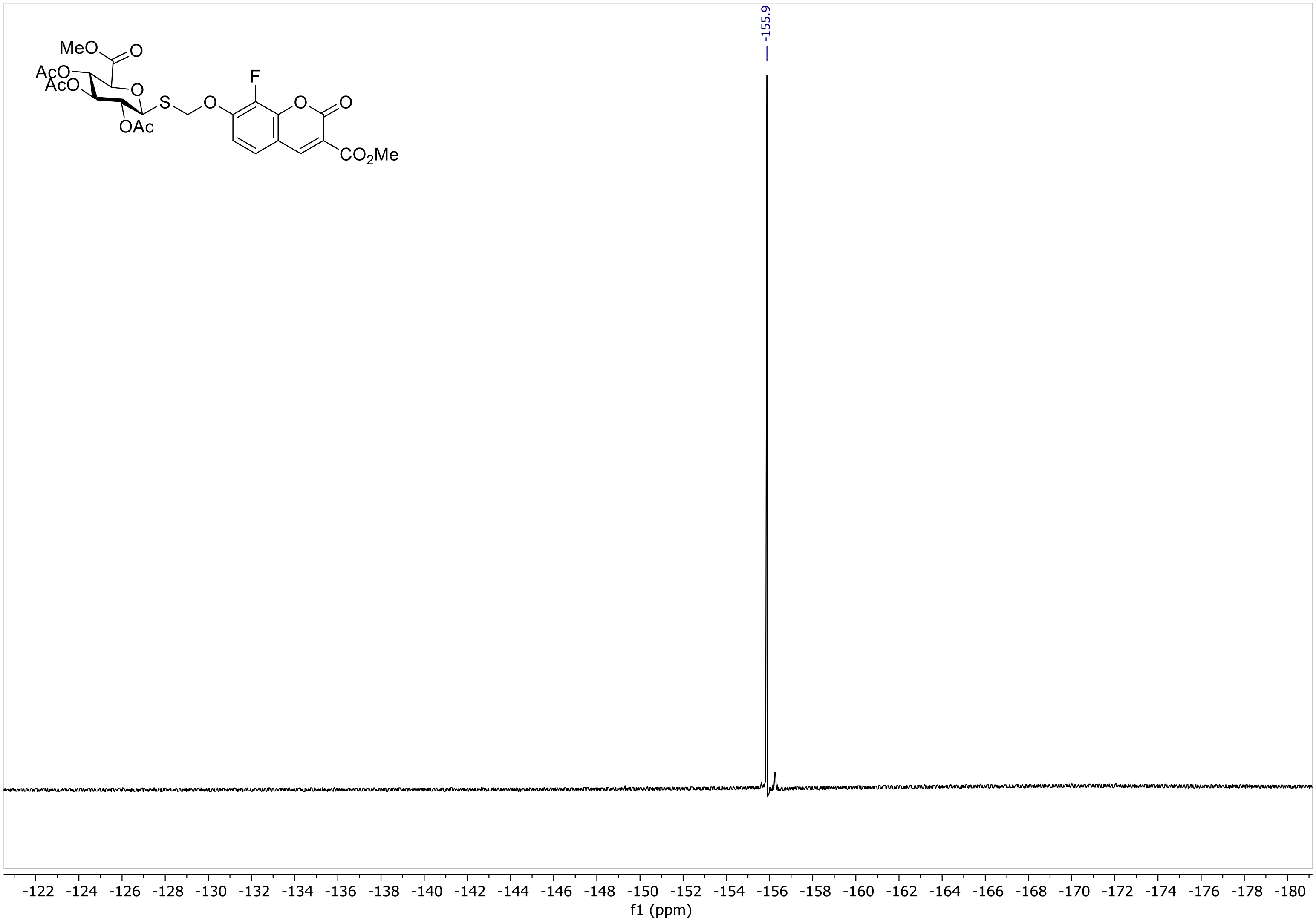

-155.9
MeO
O
AcO
AcO
O
S
O
F
O
O
OAc
CO2Me
-122 -124 -126 -128 -130 -132 -134 -136 -138 -140 -142 -144 -146 -148 -150 -152 -154 -156 -158 -160 -162 -164 -166 -168 -170 -172 -174 -176 -178 -180
f1 (ppm)

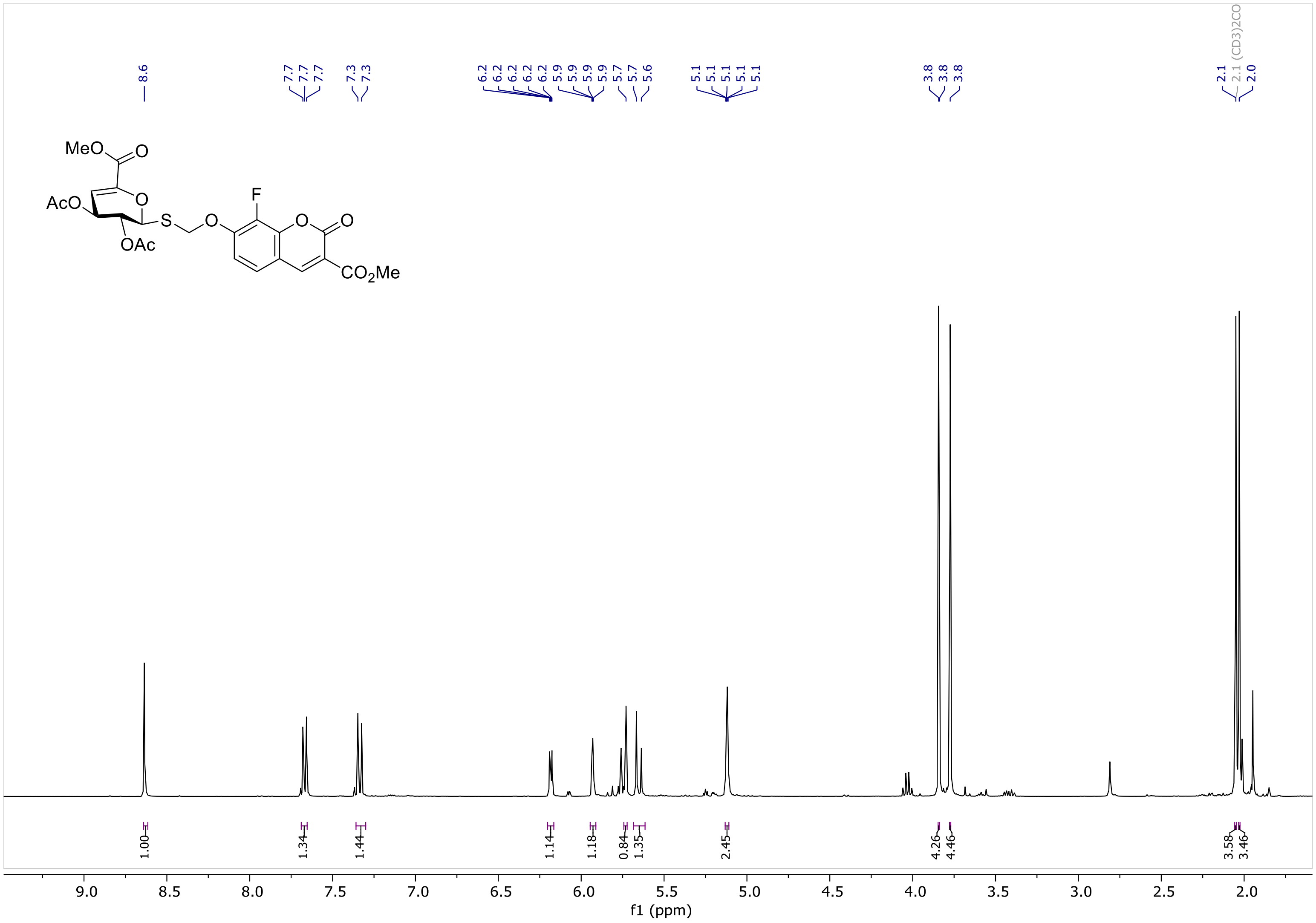
8.6
7.7 7.7 7.7
7.3 7.3
6.2 6.2 6.2 6.2 6.2
5.9 5.9 5.9 5.9
5.7 5.7 5.6
5.1 5.1 5.1 5.1 5.1
3.8 3.8 3.8
2.1
2.1 (CD3)2CO
2.0
MeO
O
AcO
O
F
S
O
O
O
OAc
CO2Me
1.00
1.34
1.44
1.14
1.18
0.84
1.35
2.45
4.26
4.46
3.58
3.46
9.0 8.5 8.0 7.5 7.0 6.5 6.0 5.5 5.0 4.5 4.0 3.5 3.0 2.5 2.0
f1 (ppm)

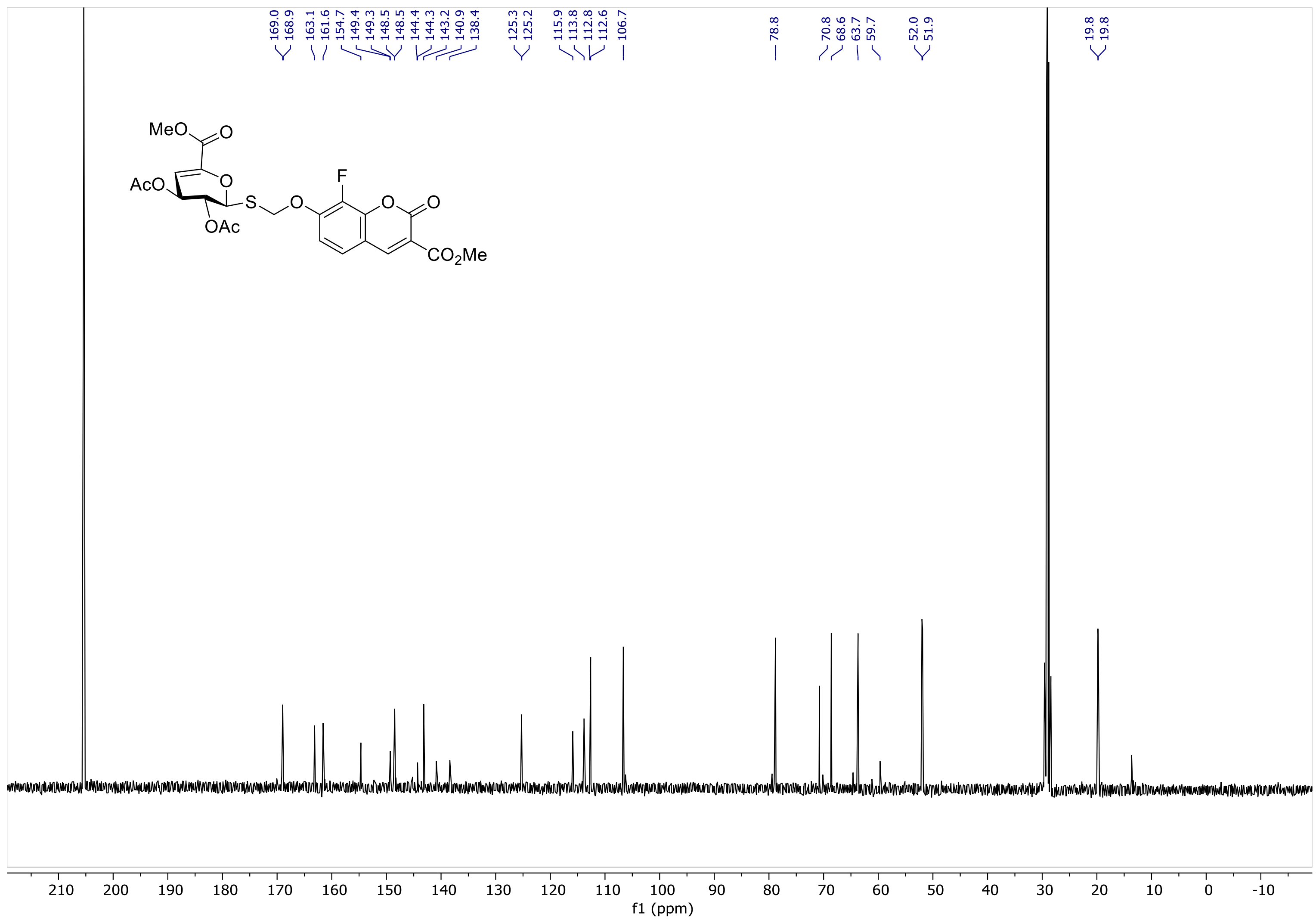

169.0
168.9
163.1
161.6
154.7
149.4
149.3
148.5
148.5
144.4
144.3
143.2
140.9
138.4
125.3
125.2
115.9
113.8
112.8
112.6
106.7
78.8
70.8
68.6
63.7
59.7
52.0
51.9
19.8
19.8
MeO
O
AcO
O
S
O
F
O
O
OAc
CO2Me
210
200
190
180
170
160
150
140
130
120
110
100
90
80
70
60
50
40
30
20
10
0
-10
f1 (ppm)

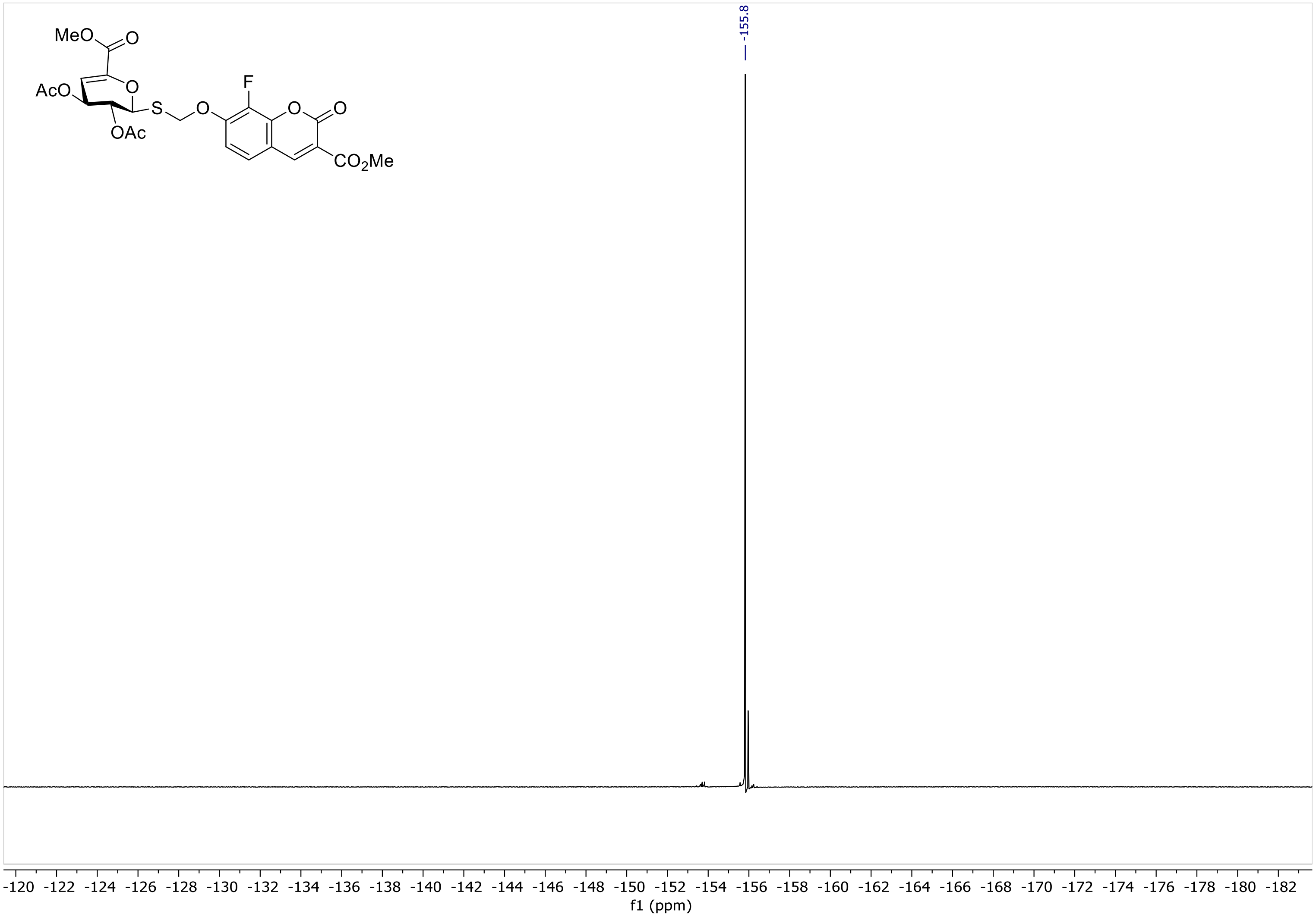

MeO
O
AcO
O
S
O
F
O
O
OAc
CO2Me
-155.8
-120 -122 -124 -126 -128 -130 -132 -134 -136 -138 -140 -142 -144 -146 -148 -150 -152 -154 -156 -158 -160 -162 -164 -166 -168 -170 -172 -174 -176 -178 -180 -182
f1 (ppm)

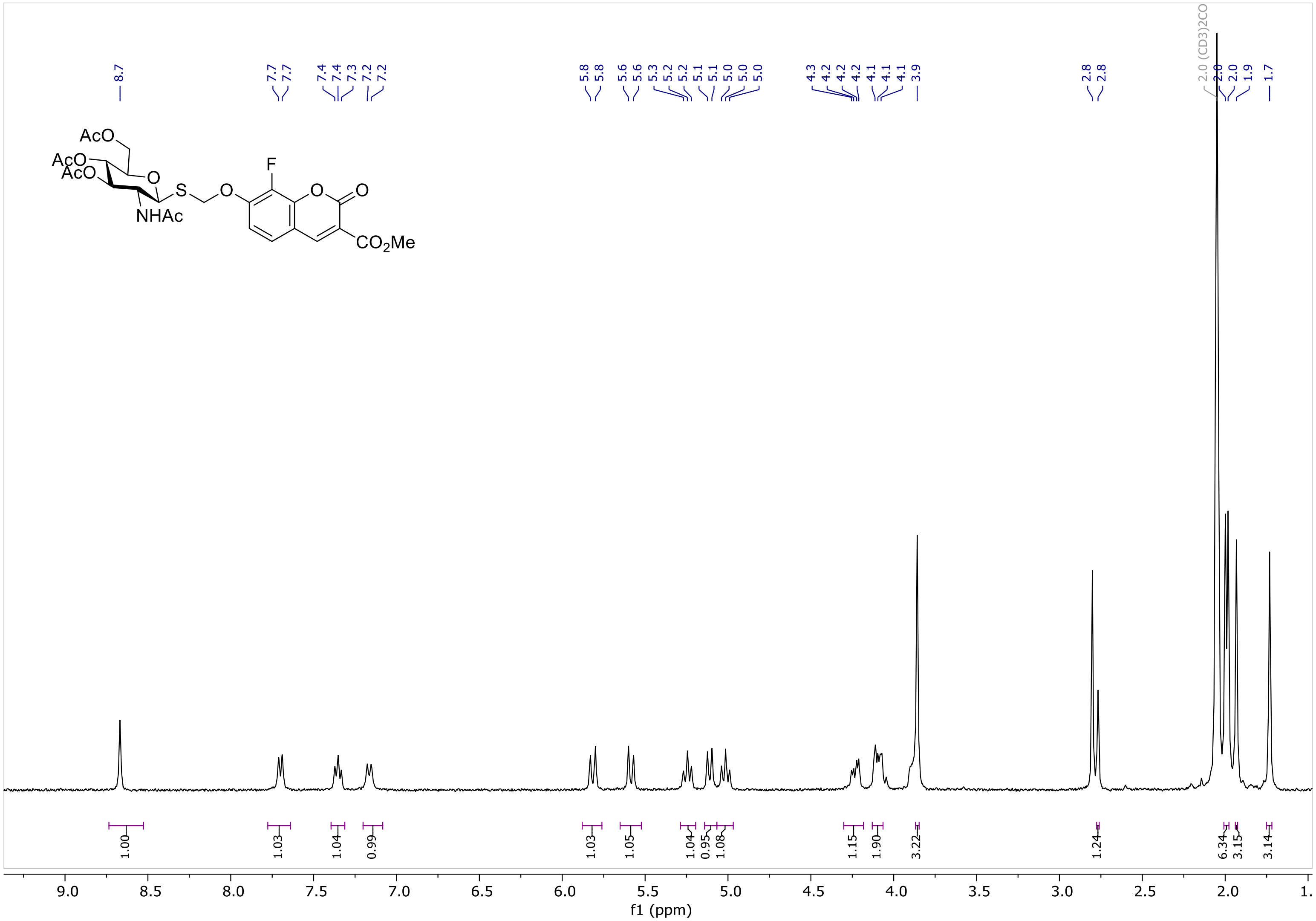

AcO
AcO
AcO
O
S
O
NHAc
F
O
O
CO2Me
8.7
7.7
7.7
7.4
7.4
7.3
7.2
7.2
5.8
5.8
5.6
5.6
5.3
5.2
5.2
5.1
5.1
5.0
5.0
5.0
4.3
4.2
4.2
4.2
4.1
4.1
4.1
3.9
2.8
2.8
2.0 (CD3)2CO
2.0
2.0
1.9
1.7
1.00
1.03
1.04
0.99
1.03
1.05
1.04
0.95
1.08
1.15
1.90
3.22
1.24
6.34
3.15
3.14
9.0
8.5
8.0
7.5
7.0
6.5
6.0
5.5
5.0
4.5
4.0
3.5
3.0
2.5
2.0
1.
f1 (ppm)

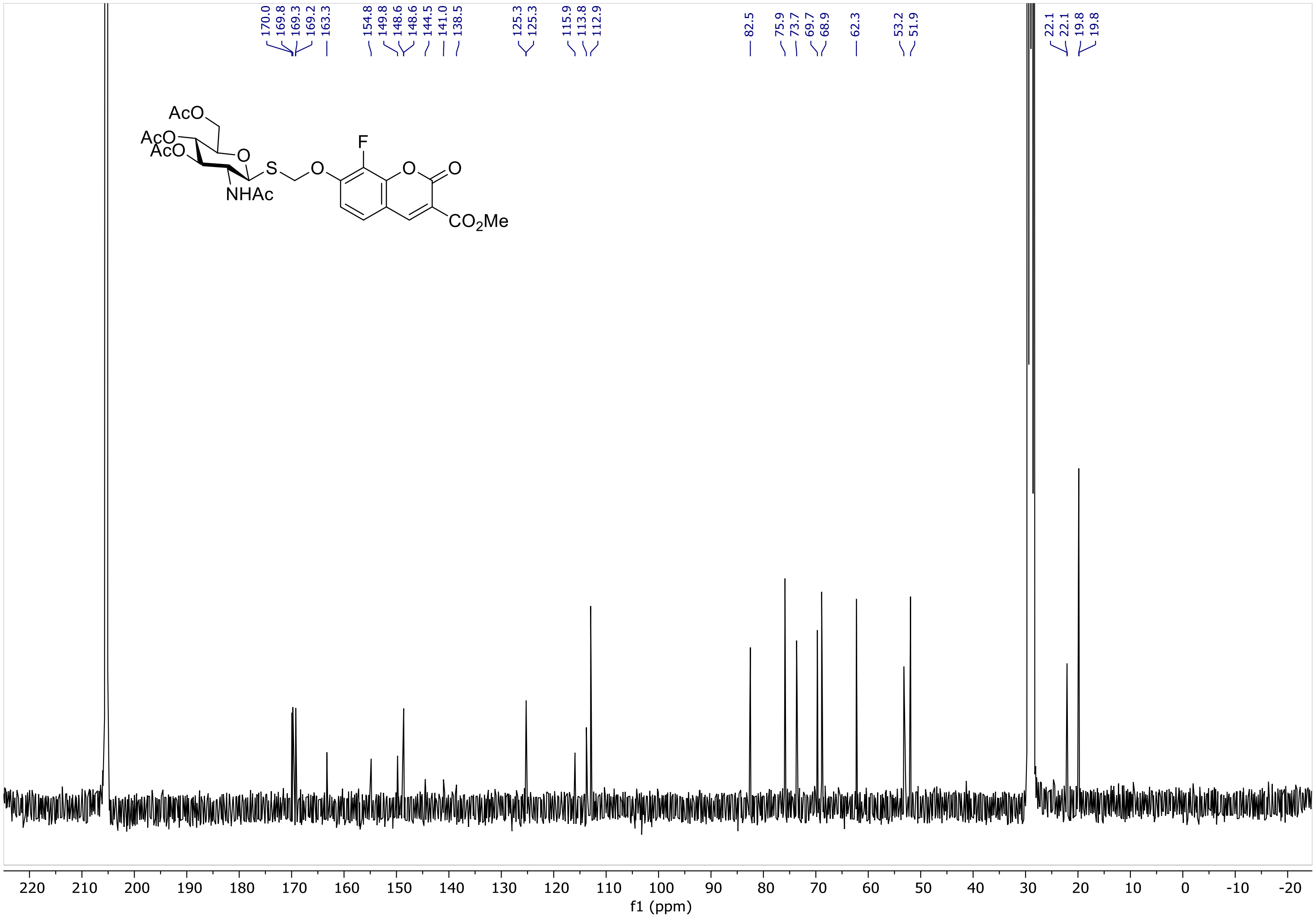
170.0
169.8
169.3
169.2
163.3
154.8
149.8
148.6
148.6
144.5
141.0
138.5
125.3
125.3
115.9
113.8
112.9
82.5
75.9
73.7
69.7
68.9
62.3
53.2
51.9
22.1
22.1
19.8
19.8
AcO
AcO
AcO
O
S
O
F
O
O
NHAc
CO2Me
220 210 200 190 180 170 160 150 140 130 120 110 100 90 80 70 60 50 40 30 20 10 0 -10 -20
f1 (ppm)

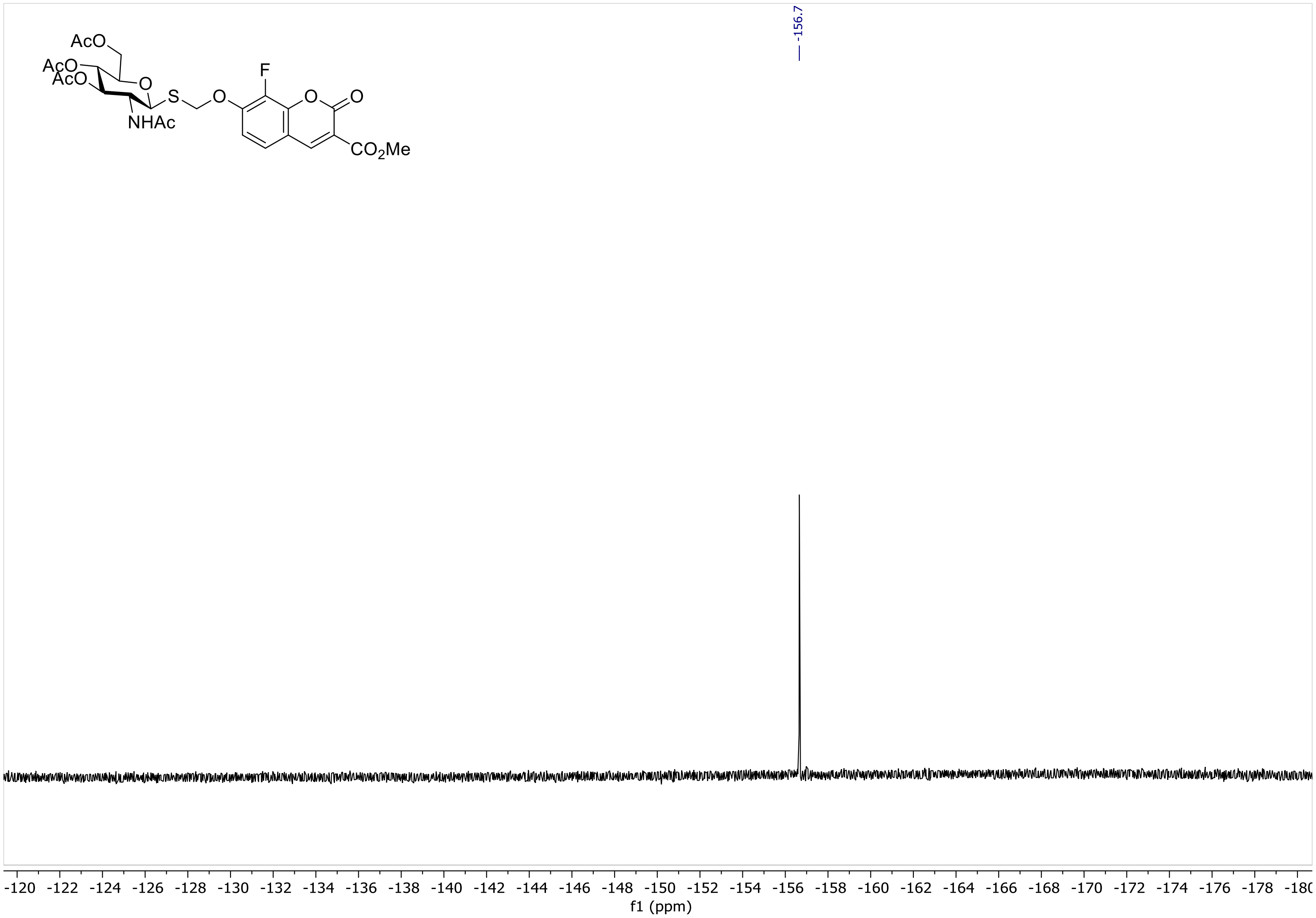

AcO
AcO
AcO
O
S
O
F
O
O
NHAc
CO2Me
-156.7
-120 -122 -124 -126 -128 -130 -132 -134 -136 -138 -140 -142 -144 -146 -148 -150 -152 -154 -156 -158 -160 -162 -164 -166 -168 -170 -172 -174 -176 -178 -180
f1 (ppm)

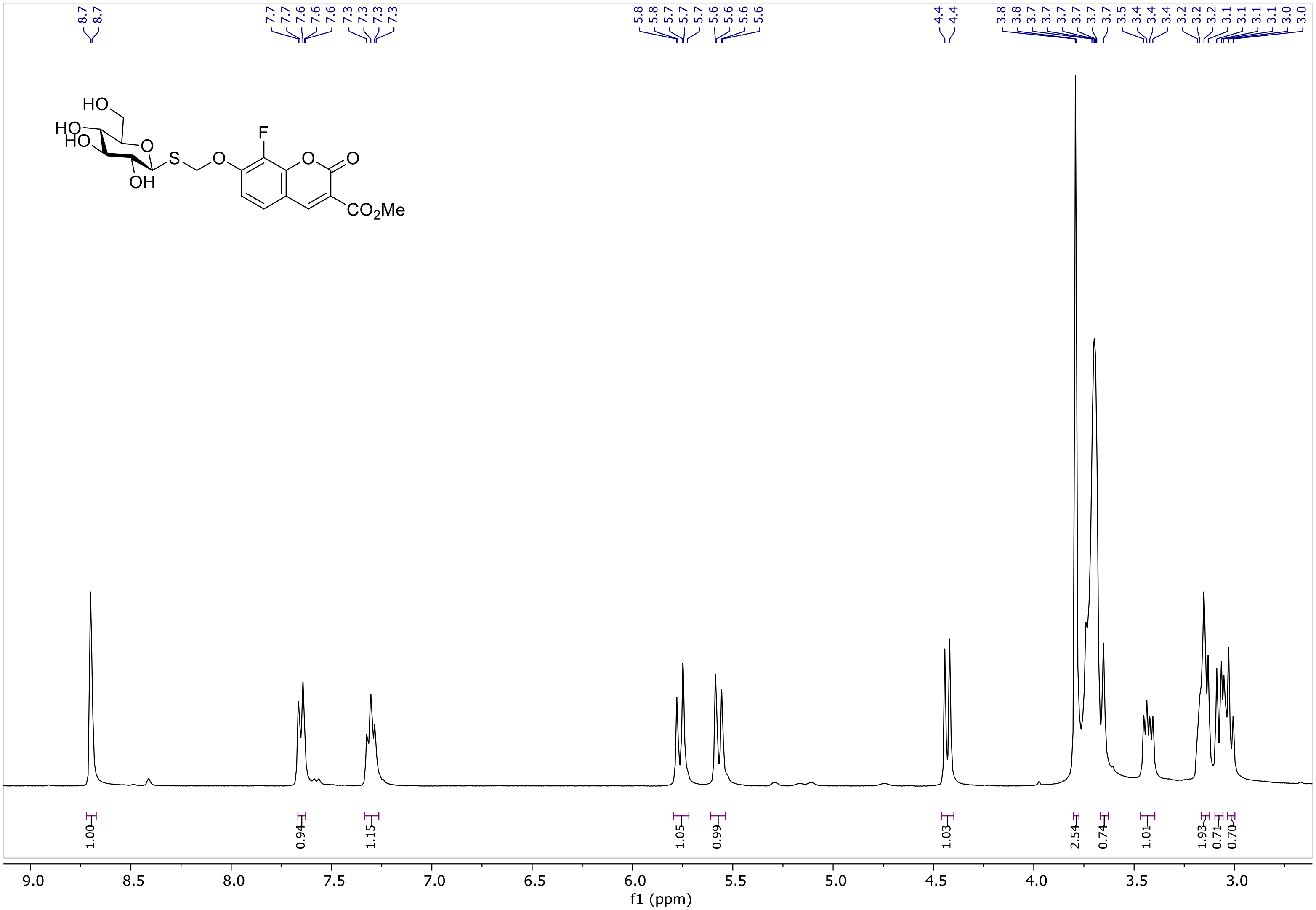
8.7
8.7
7.7
7.7
7.6
7.6
7.6
7.3
7.3
7.3
7.3
5.8
5.8
5.7
5.7
5.7
5.6
5.6
5.6
5.6
4.4
4.4
3.8
3.8
3.7
3.7
3.7
3.7
3.7
3.7
3.5
3.4
3.4
3.4
3.2
3.2
3.2
3.1
3.1
3.1
3.1
3.0
3.0
HO
HO
HO
O
OH
S
O
F
O
O
CO2Me
1.00
0.94
1.15
1.05
0.99
1.03
2.54
0.74
1.01
1.93
0.71
0.70
9.0
8.5
8.0
7.5
7.0
6.5
6.0
5.5
5.0
4.5
4.0
3.5
3.0
f1 (ppm)

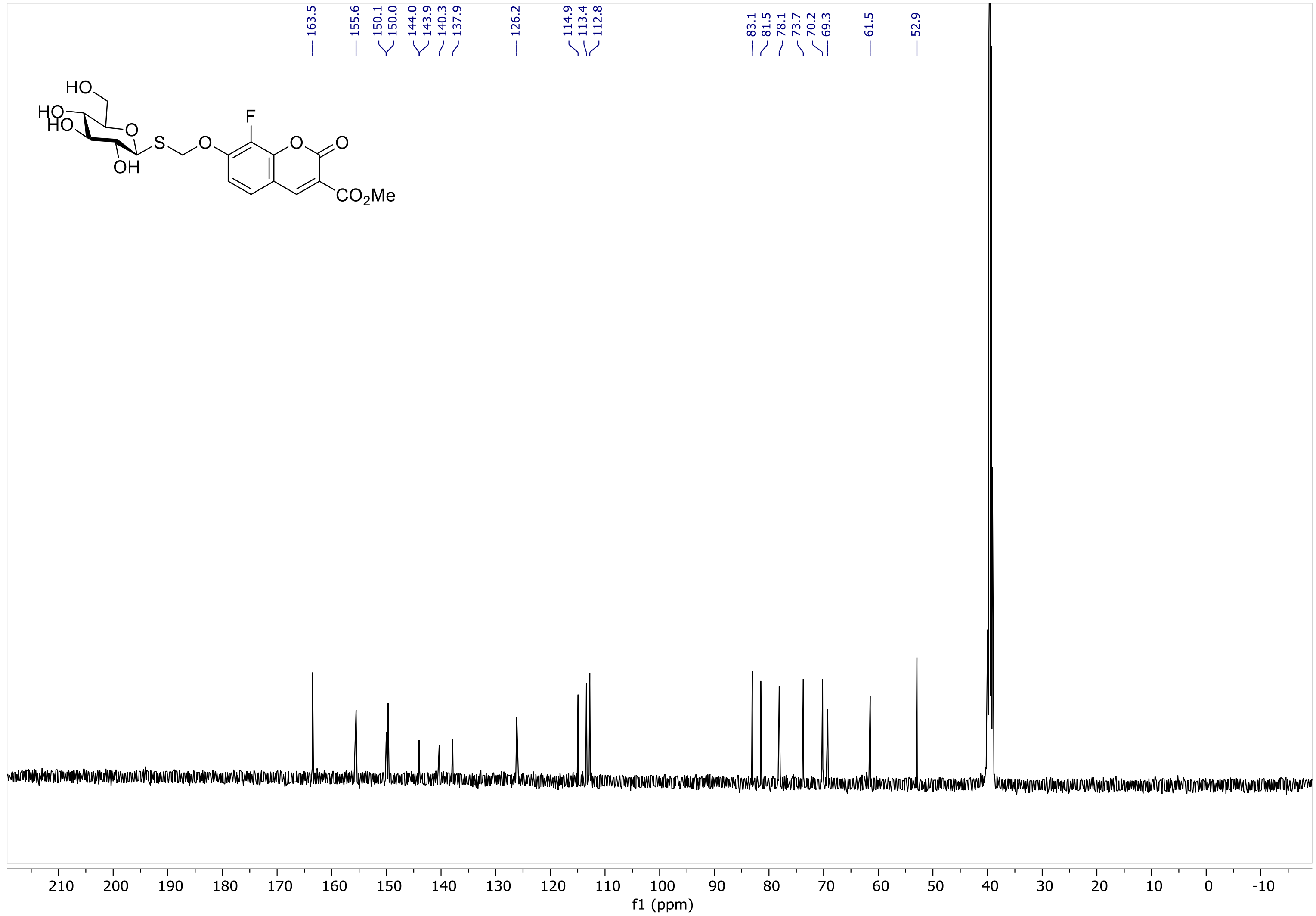

163.5
155.6
150.1
150.0
144.0
143.9
140.3
137.9
126.2
114.9
113.4
112.8
83.1
81.5
78.1
73.7
70.2
69.3
61.5
52.9
HO
HO
HO
O
S
O
F
O
O
OH
CO2Me
210 200 190 180 170 160 150 140 130 120 110 100 90 80 70 60 50 40 30 20 10 0 -10
f1 (ppm)

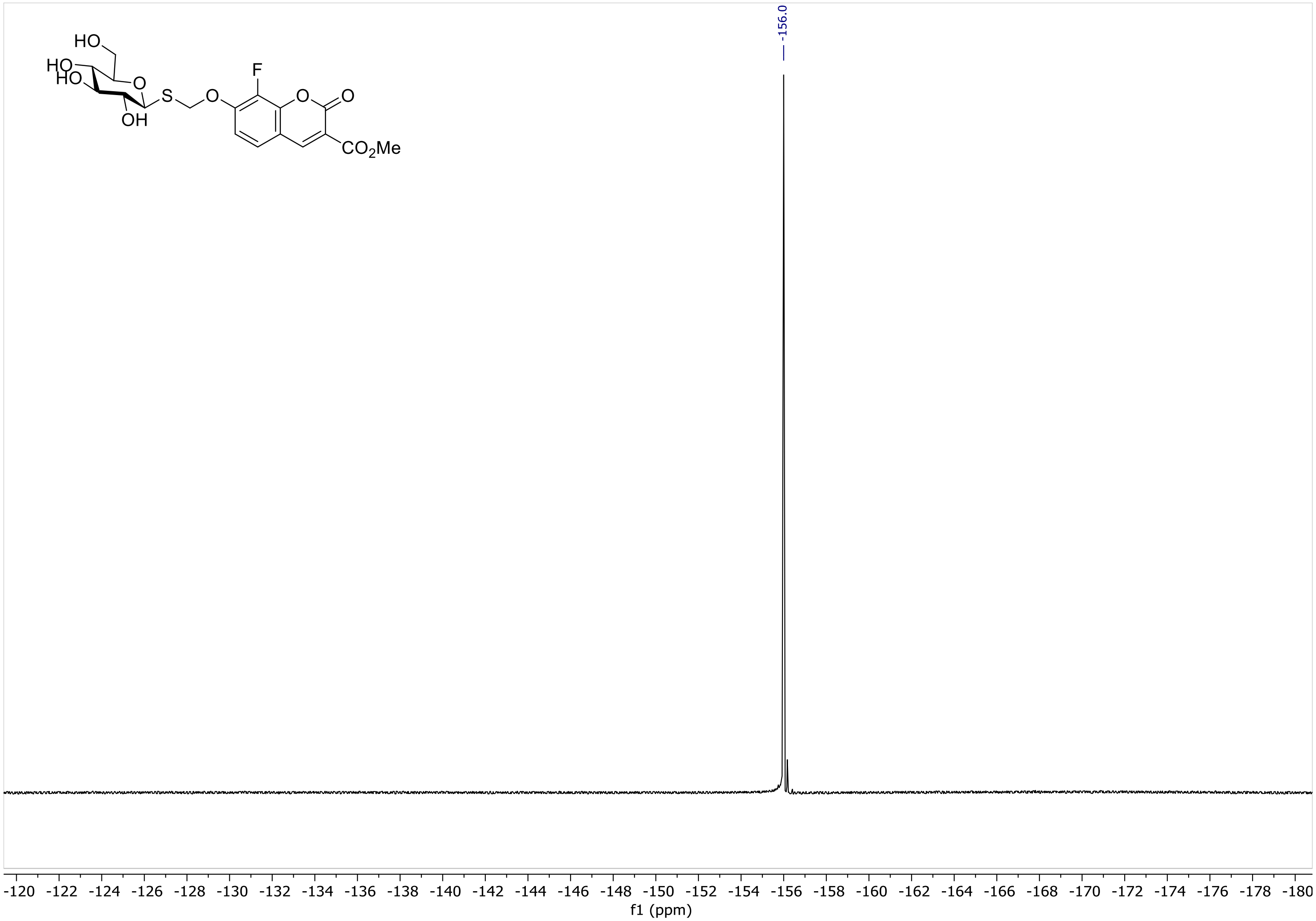
-156.0
HO
HO
HO
O
S
O
F
O
O
OH
CO2Me
-120 -122 -124 -126 -128 -130 -132 -134 -136 -138 -140 -142 -144 -146 -148 -150 -152 -154 -156 -158 -160 -162 -164 -166 -168 -170 -172 -174 -176 -178 -180
f1 (ppm)

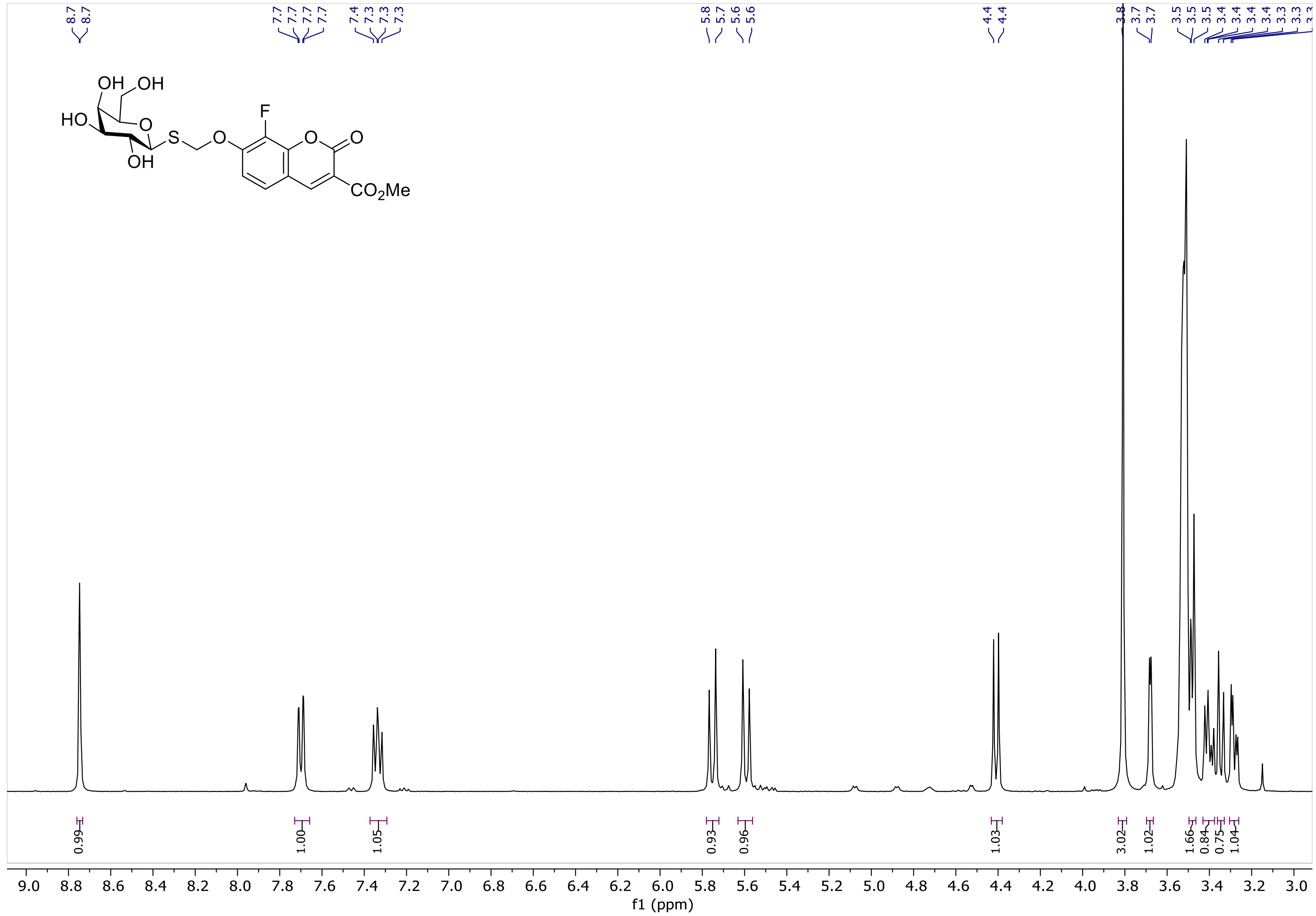
8.7
8.7
7.7
7.7
7.7
7.7
7.4
7.3
7.3
7.3
5.8
5.7
5.6
5.6
4.4
4.4
3.8
3.7
3.7
3.5
3.5
3.5
3.4
3.4
3.4
3.4
3.3
3.3
OH
OH
HO
O
OH
S
O
F
O
O
CO2Me
0.99
1.00
1.05
0.93
0.96
1.03
3.02
1.02
1.66
0.84
0.75
1.04
9.0
8.8
8.6
8.4
8.2
8.0
7.8
7.6
7.4
7.2
7.0
6.8
6.6
6.4
6.2
6.0
5.8
5.6
5.4
5.2
5.0
4.8
4.6
4.4
4.2
4.0
3.8
3.6
3.4
3.2
3.0
f1 (ppm)

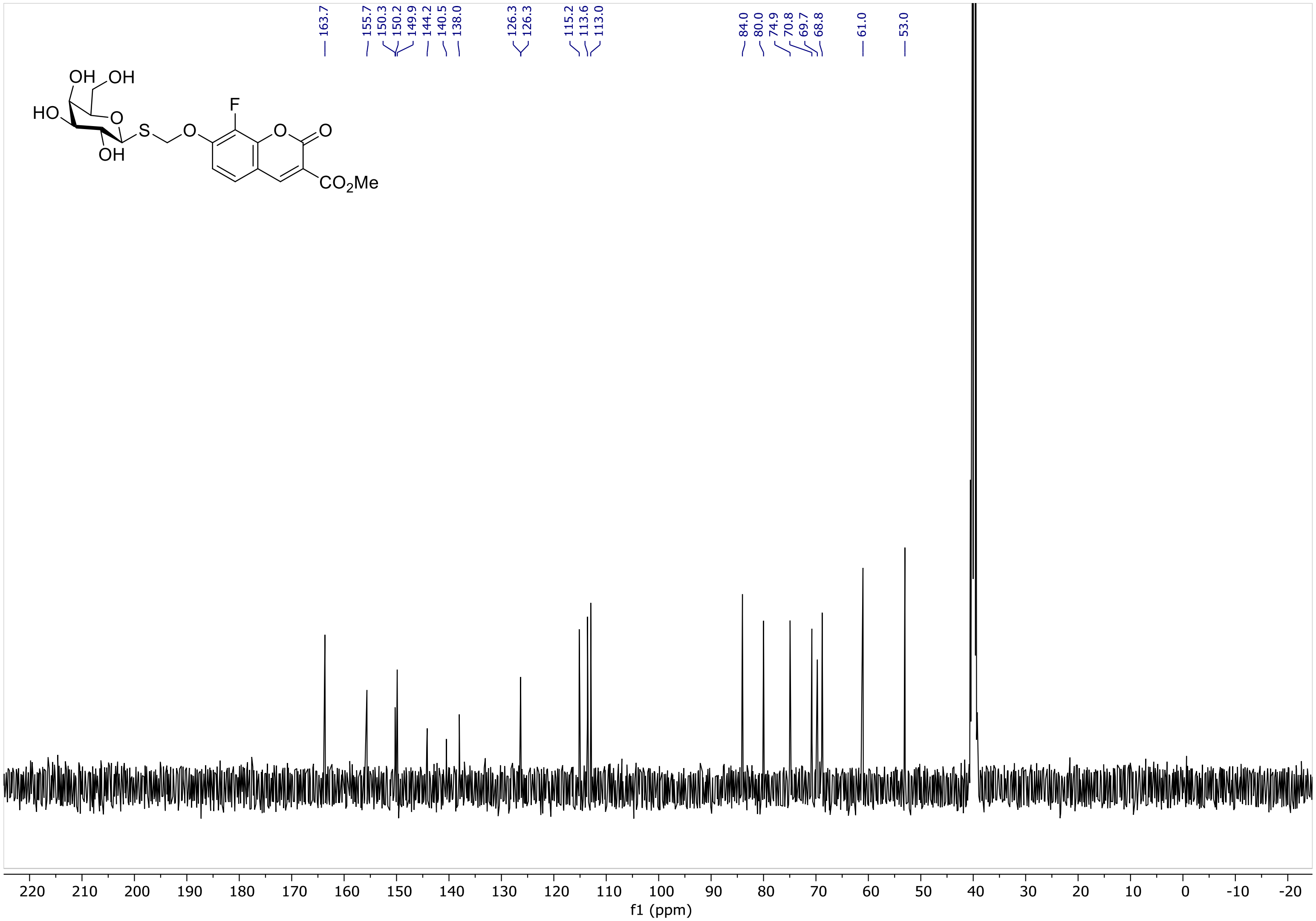

163.7
155.7
150.3
150.2
149.9
144.2
140.5
138.0
126.3
126.3
115.2
113.6
113.0
84.0
80.0
74.9
70.8
69.7
68.8
61.0
53.0
OH
OH
HO
O
OH
S
O
F
O
O
CO2Me
220 210 200 190 180 170 160 150 140 130 120 110 100 90 80 70 60 50 40 30 20 10 0 -10 -20
f1 (ppm)

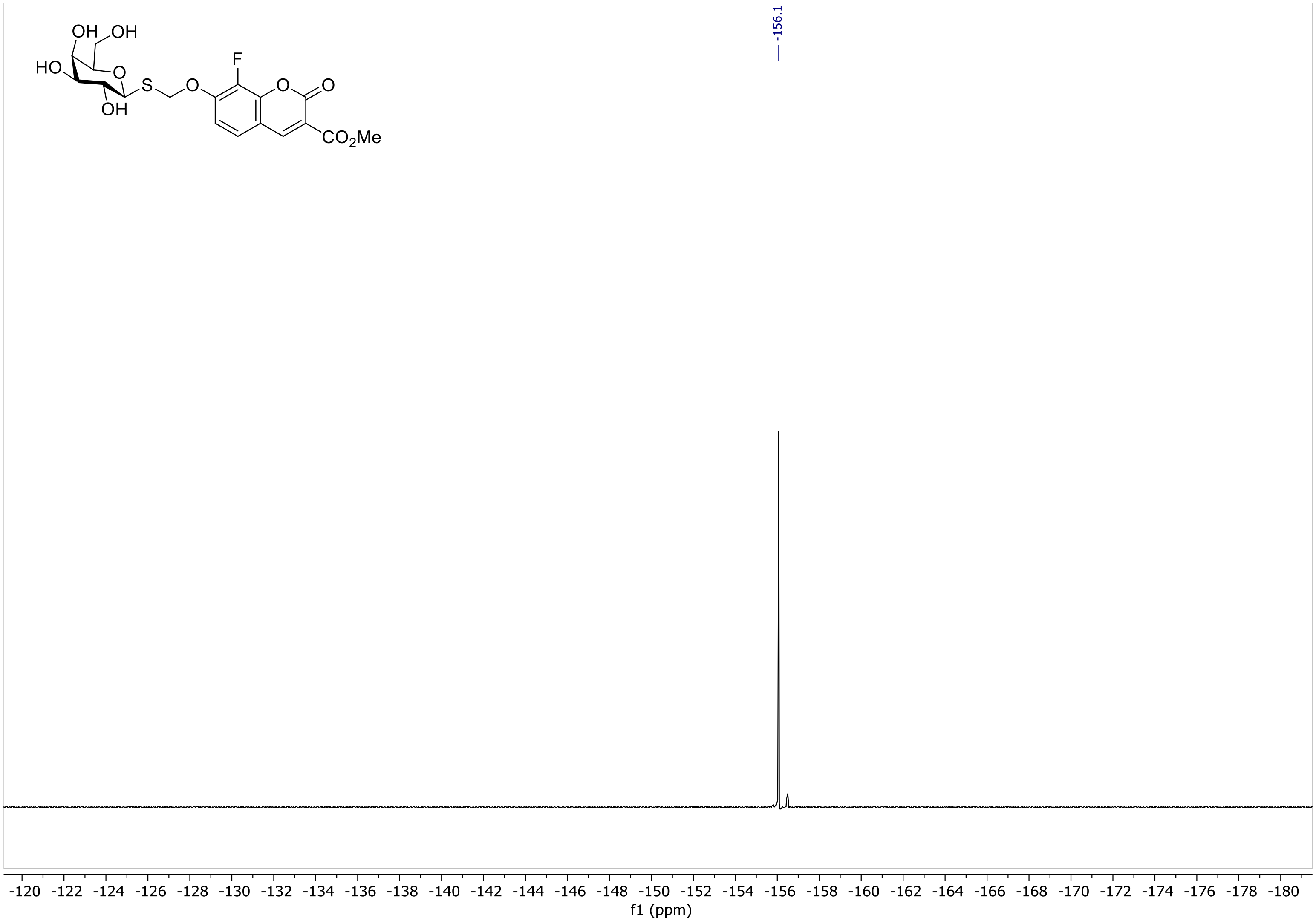
OH
OH
HO
O
S
O
F
O
O
OH
CO2Me
-156.1
-120
-122
-124
-126
-128
-130
-132
-134
-136
-138
-140
-142
-144
-146
-148
-150
-152
-154
-156
-158
-160
-162
-164
-166
-168
-170
-172
-174
-176
-178
-180
f1 (ppm)

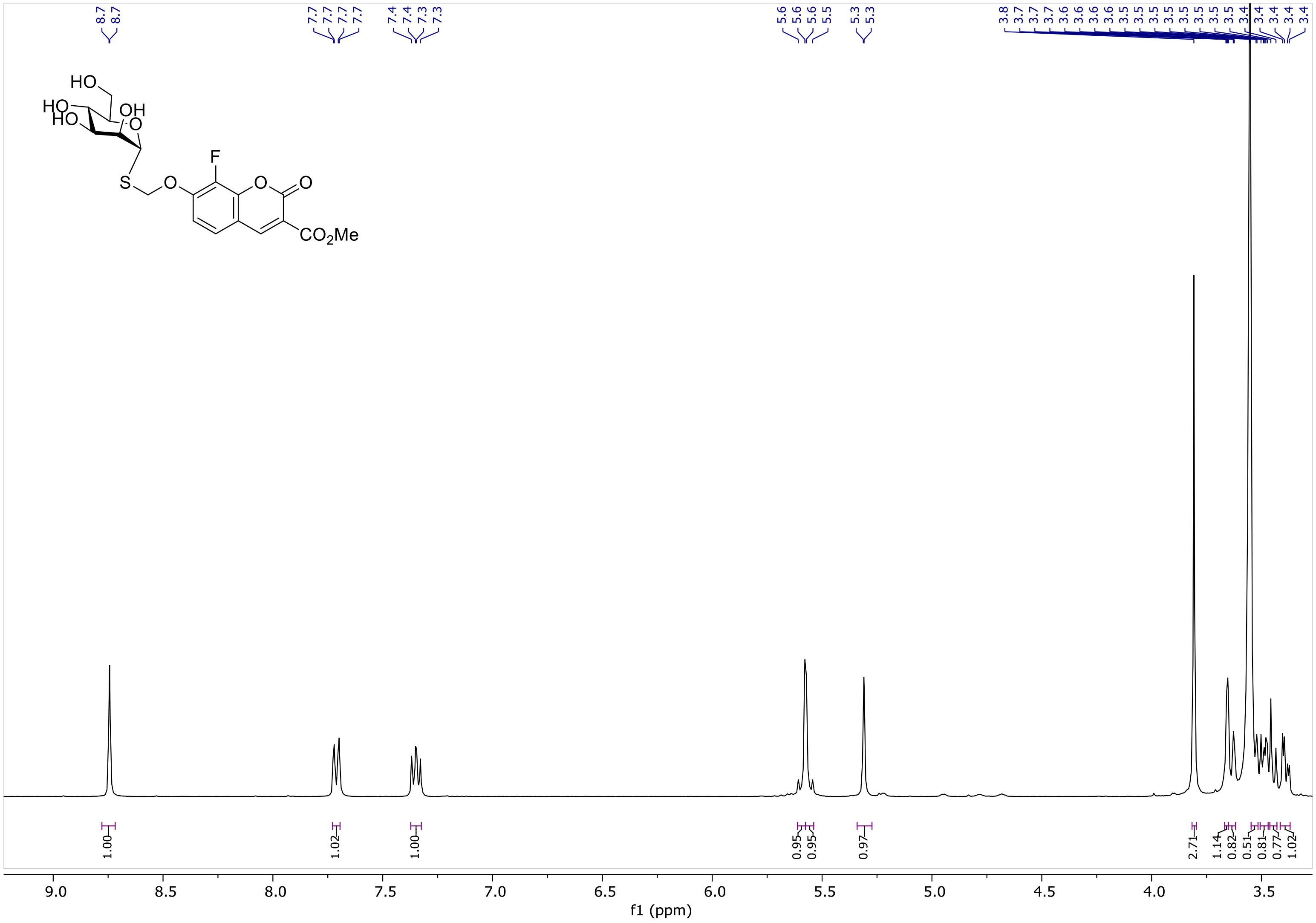
8.7
8.7
7.7
7.7
7.7
7.7
7.4
7.4
7.3
7.3
5.6
5.6
5.6
5.5
5.3
5.3
3.8
3.7
3.7
3.7
3.6
3.6
3.6
3.5
3.5
3.5
3.5
3.5
3.5
3.5
3.5
3.4
3.4
3.4
3.4
3.4
HO
HO
HO
OH
O
F
S
O
O
O
CO2Me
1.00
1.02
1.00
0.95
0.95
0.97
2.71
1.14
0.82
0.51
0.81
0.77
1.02
9.0
8.5
8.0
7.5
7.0
6.5
6.0
5.5
5.0
4.5
4.0
3.5
f1 (ppm)

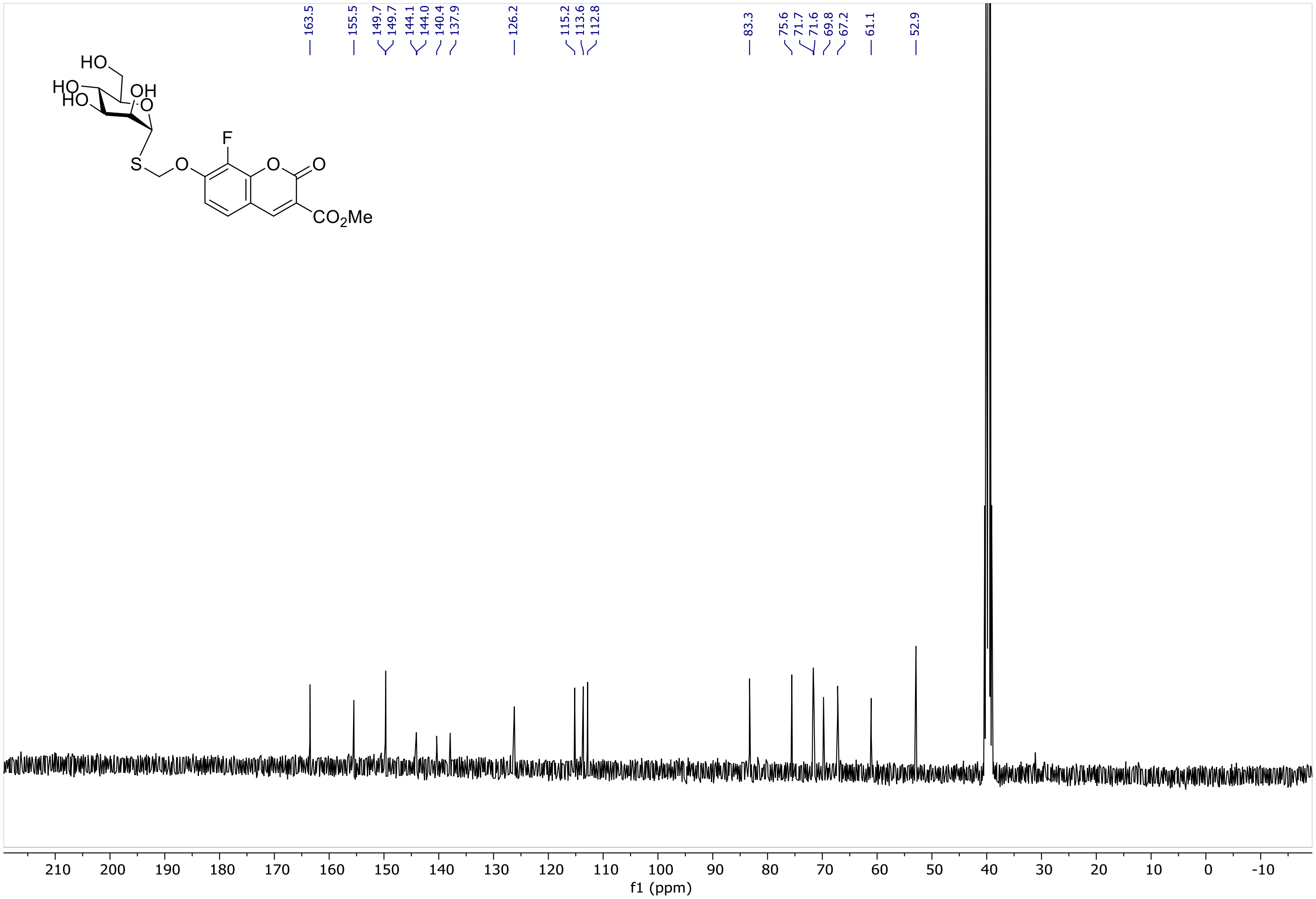
163.5
155.5
149.7
149.7
144.1
144.0
140.4
137.9
126.2
115.2
113.6
112.8
83.3
75.6
71.7
71.6
69.8
67.2
61.1
52.9
HO
HO
HO
OH
O
S
O
F
O
O
CO2Me
210 200 190 180 170 160 150 140 130 120 110 100 90 80 70 60 50 40 30 20 10 0 -10
f1 (ppm)

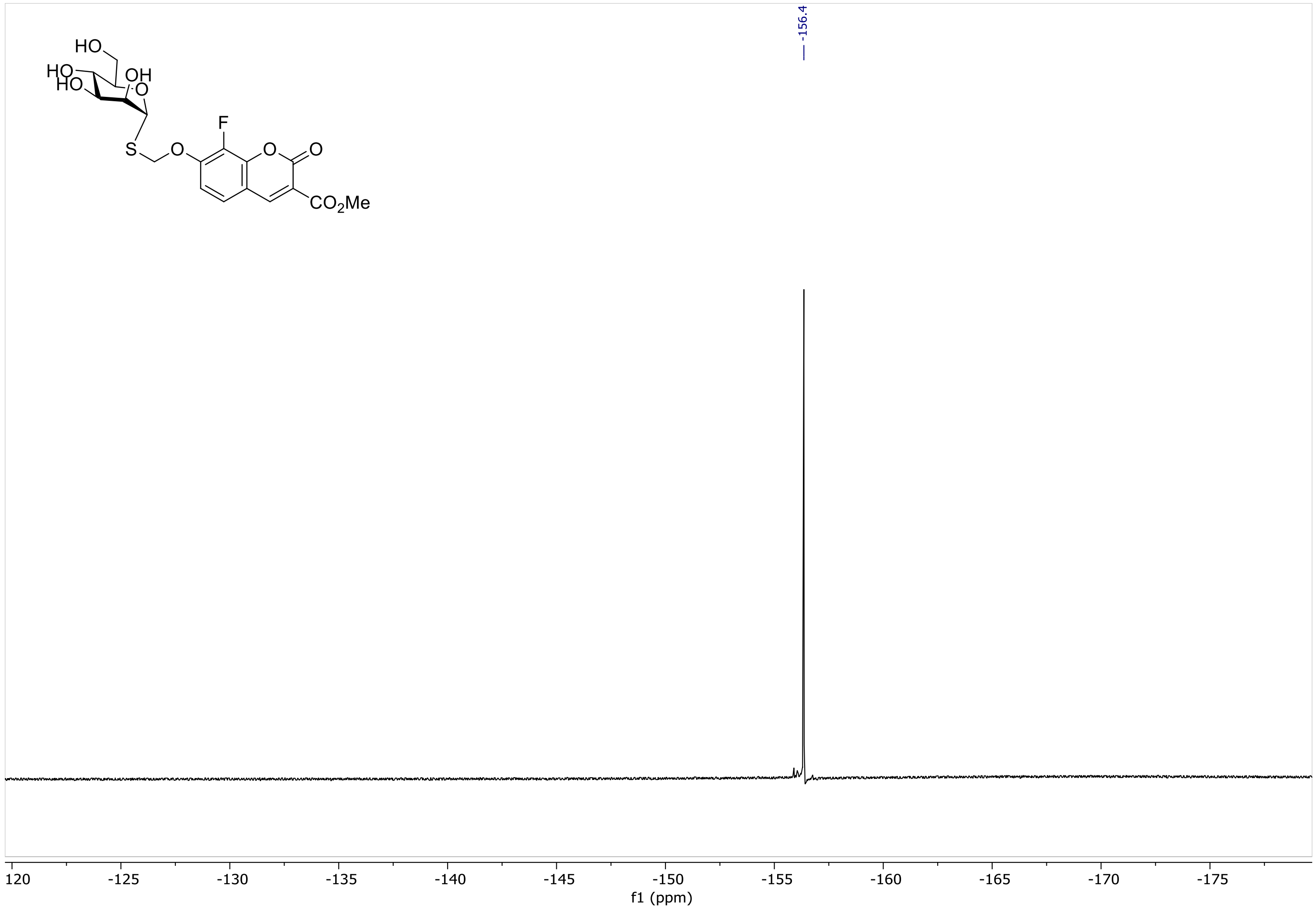

-156.4
HO
HO
HO
OH
O
S
O
F
O
O
CO2Me
120
-125
-130
-135
-140
-145
-150
-155
-160
-165
-170
-175
f1 (ppm)

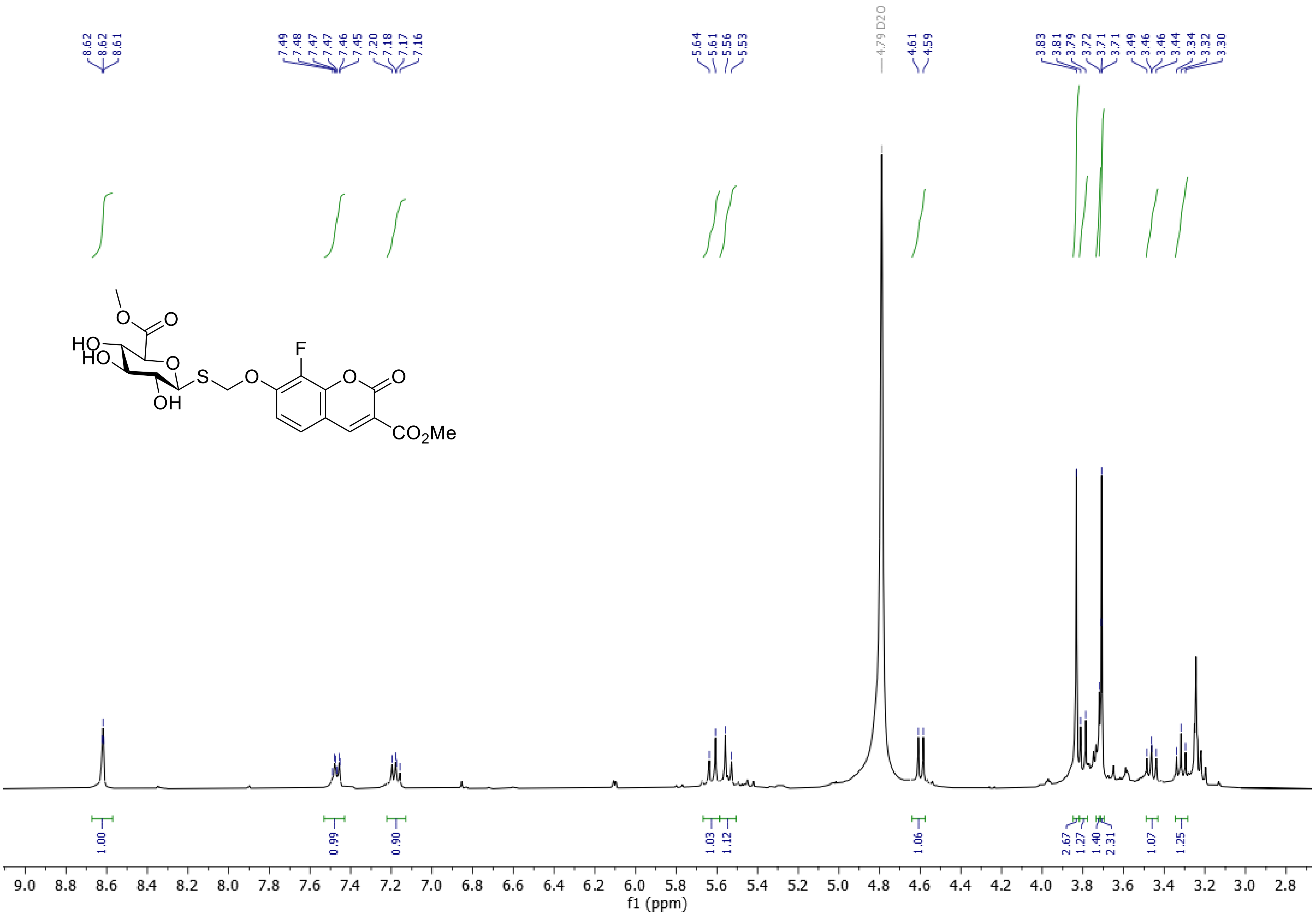
8.62
8.62
8.61
7.49
7.48
7.47
7.47
7.46
7.45
7.20
7.18
7.17
7.16
5.64
5.61
5.56
5.53
4.79 D2O
4.61
4.59
3.83
3.81
3.79
3.72
3.71
3.71
3.49
3.46
3.46
3.44
3.34
3.32
3.30
HO
HO
O
O
O
OH
S
O
F
O
O
CO2Me
1.00
0.99
0.90
1.03
1.12
1.06
2.67
1.27
1.40
2.31
1.07
1.25
9.0 8.8 8.6 8.4 8.2 8.0 7.8 7.6 7.4 7.2 7.0 6.8 6.6 6.4 6.2 6.0 5.8 5.6 5.4 5.2 5.0 4.8 4.6 4.4 4.2 4.0 3.8 3.6 3.4 3.2 3.0 2.8
f1 (ppm)

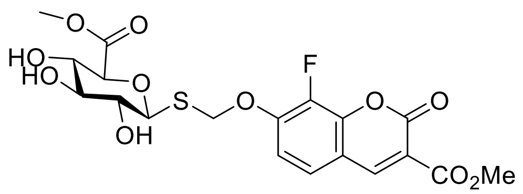

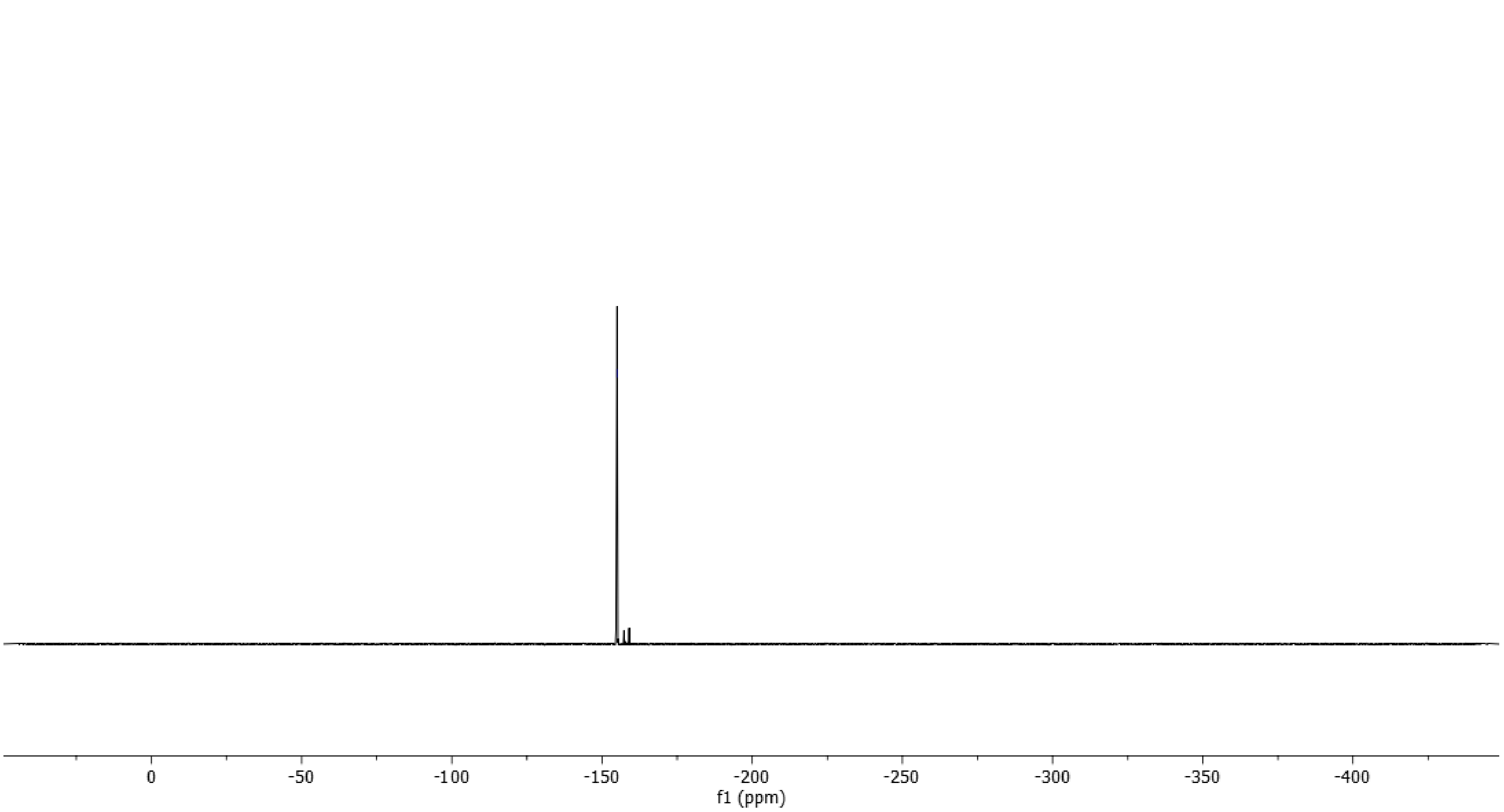

-155.00
0
-50
-100
-150
-200
-250
-300
-350
-400
f1 (ppm)

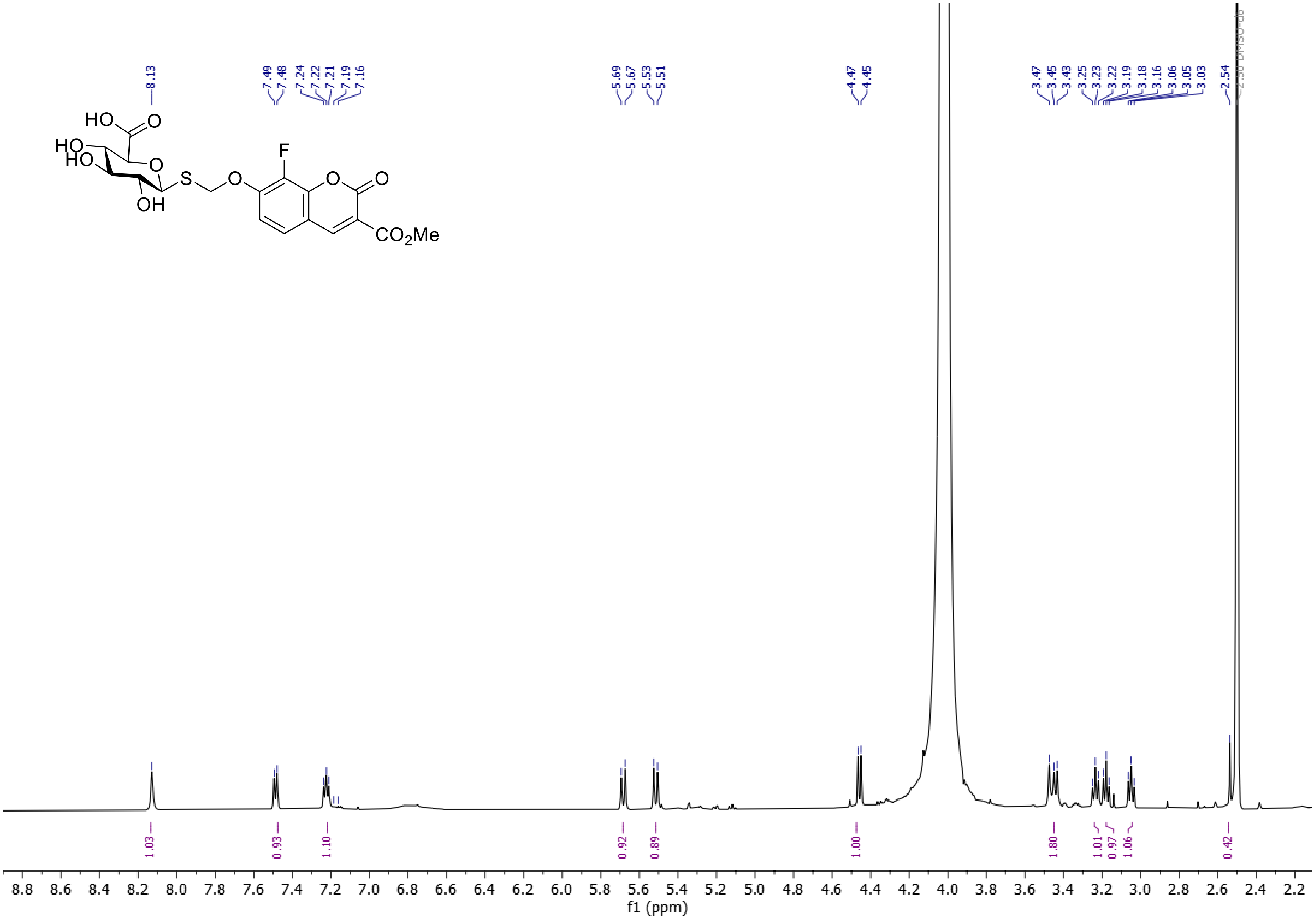

HO
O
HO
HO
O
S
O
F
O
O
OH
CO2Me
8.13
7.49
7.48
7.24
7.22
7.21
7.19
7.16
5.69
5.67
5.53
5.51
4.47
4.45
3.47
3.45
3.43
3.25
3.23
3.22
3.19
3.18
3.16
3.06
3.05
3.03
2.54
1.03
0.93
1.10
0.92
0.89
1.00
1.80
1.01
0.97
1.06
0.42
f1 (ppm)

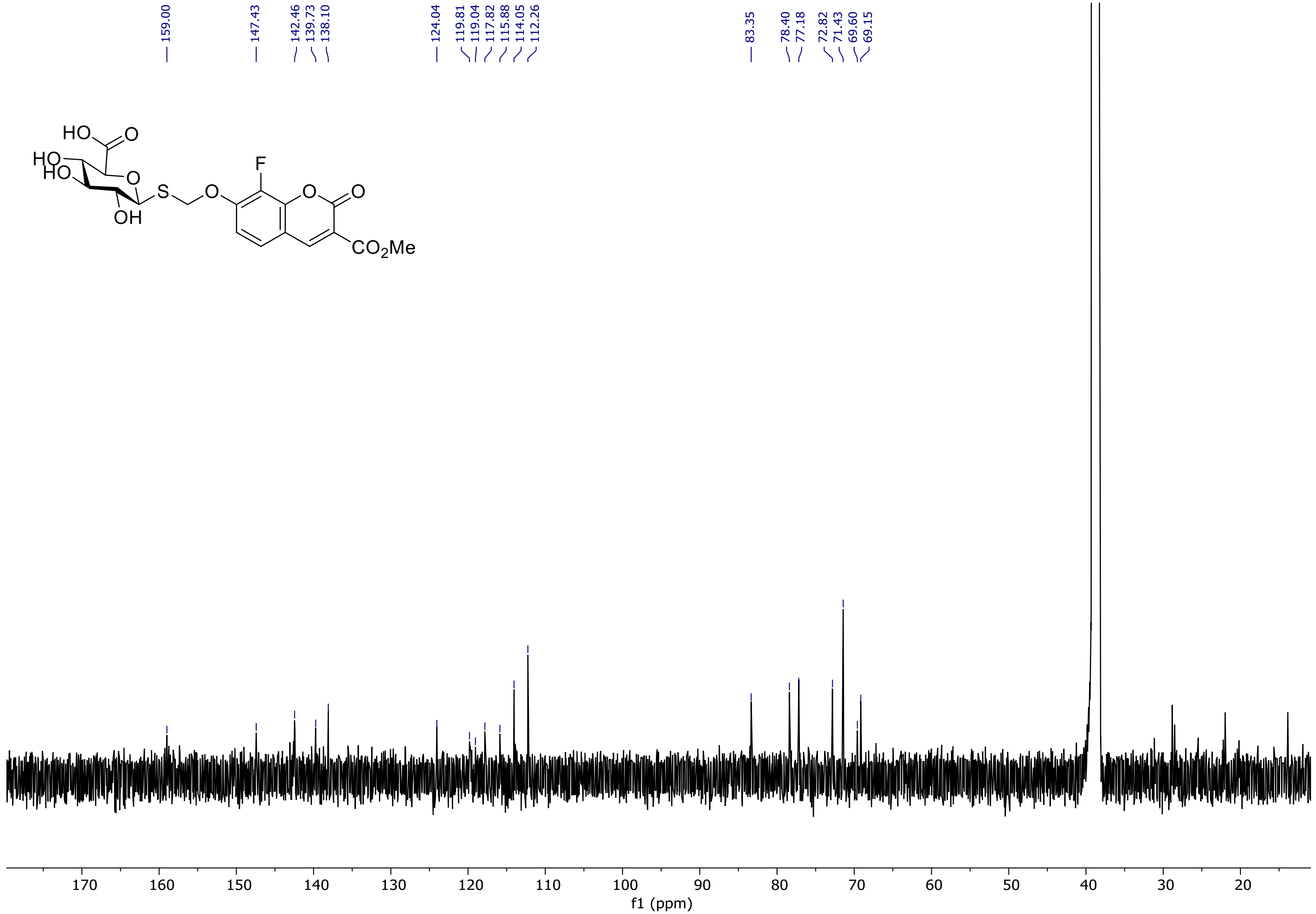
159.00
147.43
142.46
139.73
138.10
124.04
119.81
119.04
117.82
115.88
114.05
112.26
83.35
78.40
77.18
72.82
71.43
69.60
69.15
HO
O
HO
HO
O
S
O
F
O
O
OH
$CO_2Me$
170
160
150
140
130
120
110
100
90
80
70
60
50
40
30
20
f1 (ppm)

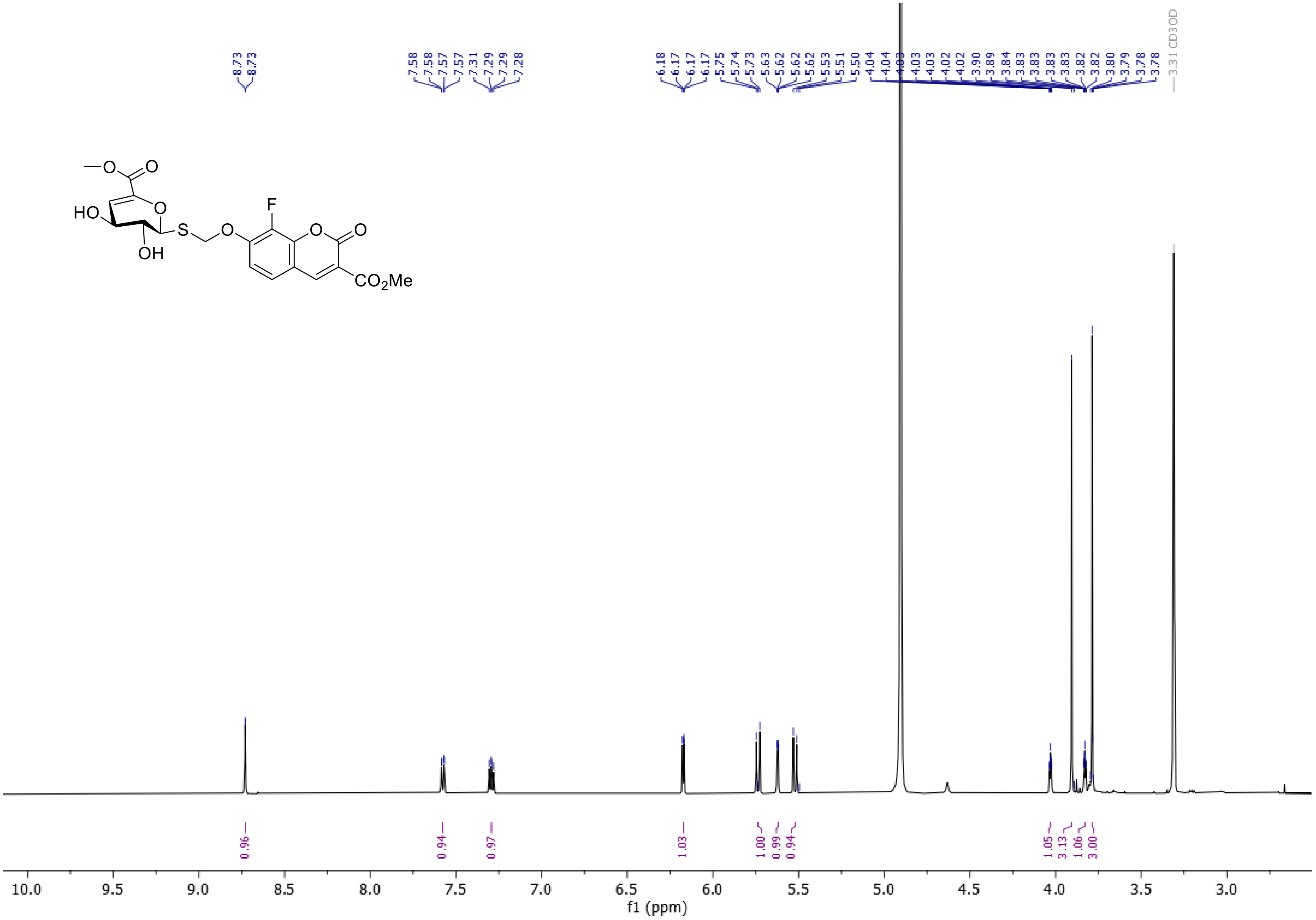
8.73
8.73
7.58
7.58
7.57
7.57
7.31
7.29
7.29
7.28
6.18
6.17
6.17
6.17
5.75
5.74
5.73
5.63
5.62
5.62
5.53
5.51
5.50
4.04
4.04
4.03
4.03
4.03
4.02
4.02
3.90
3.89
3.84
3.83
3.83
3.83
3.83
3.82
3.82
3.80
3.79
3.78
3.78
3.31 CD3OD
0.96
0.94
0.97
1.03
1.00
0.99
0.94
1.05
3.13
1.06
3.00
10.0
9.5
9.0
8.5
8.0
7.5
7.0
6.5
6.0
5.5
5.0
4.5
4.0
3.5
3.0
f1 (ppm)
HO
OH
S
O
F
CO2Me

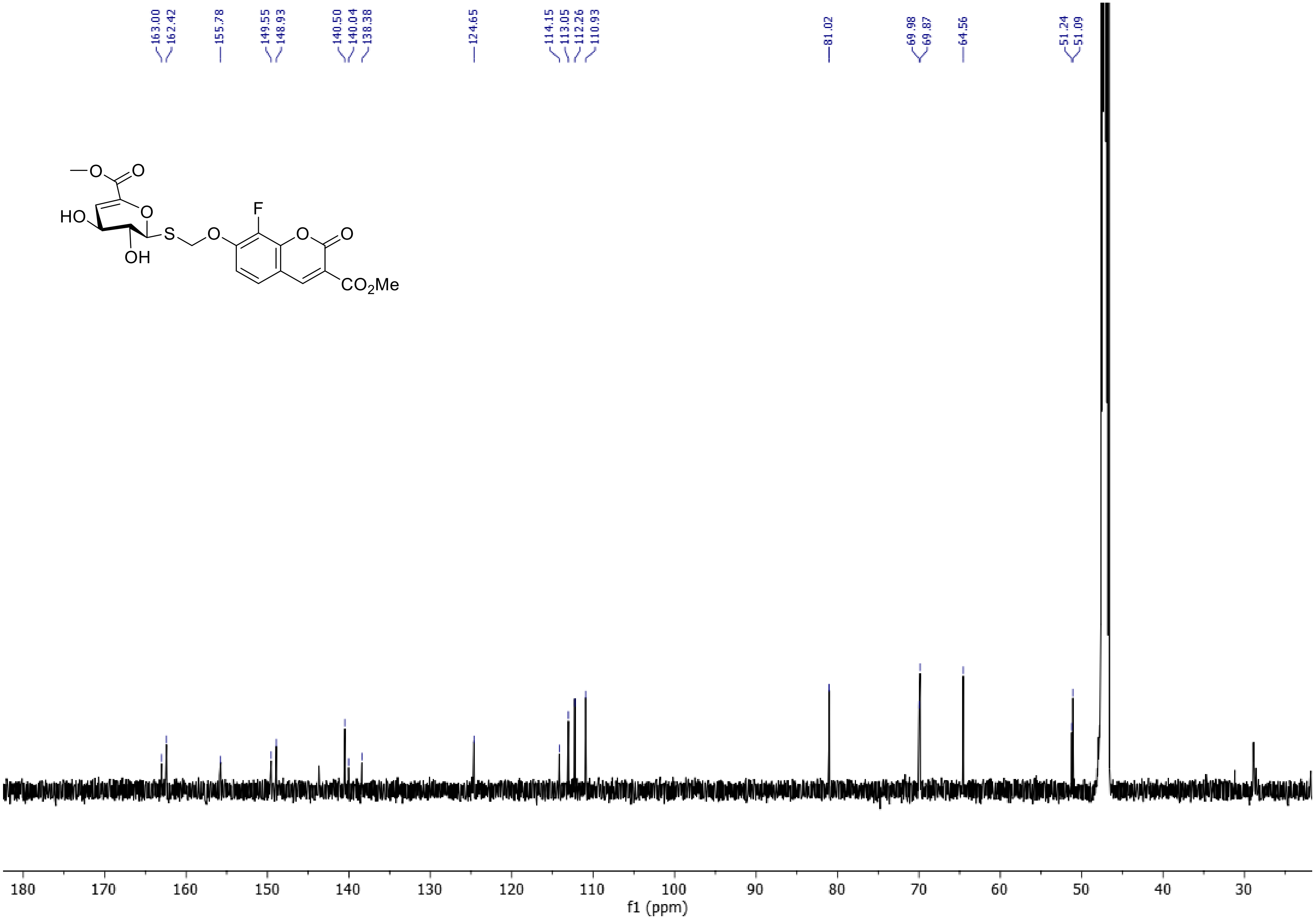
163.00
162.42
155.78
149.55
148.93
140.50
140.04
138.38
124.65
114.15
113.05
112.26
110.93
81.02
69.98
69.87
64.56
51.24
51.09
O
O
HO
O
S
O
F
O
O
OH
CO2Me
180 170 160 150 140 130 120 110 100 90 80 70 60 50 40 30
f1 (ppm)

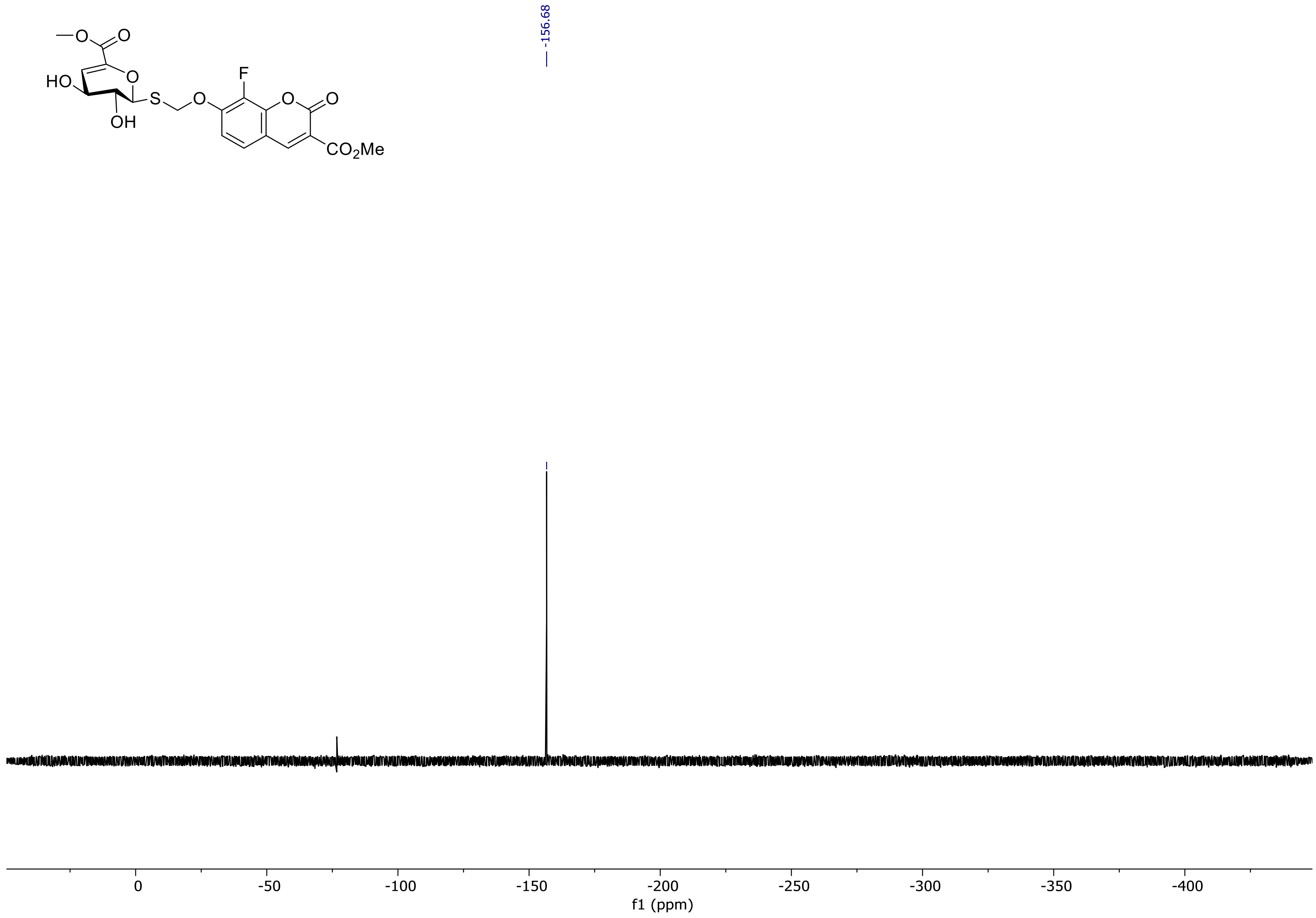

-156.68
HO
OH
S
F
CO2Me
0
-50
-100
-150
-200
-250
-300
-350
-400
f1 (ppm)

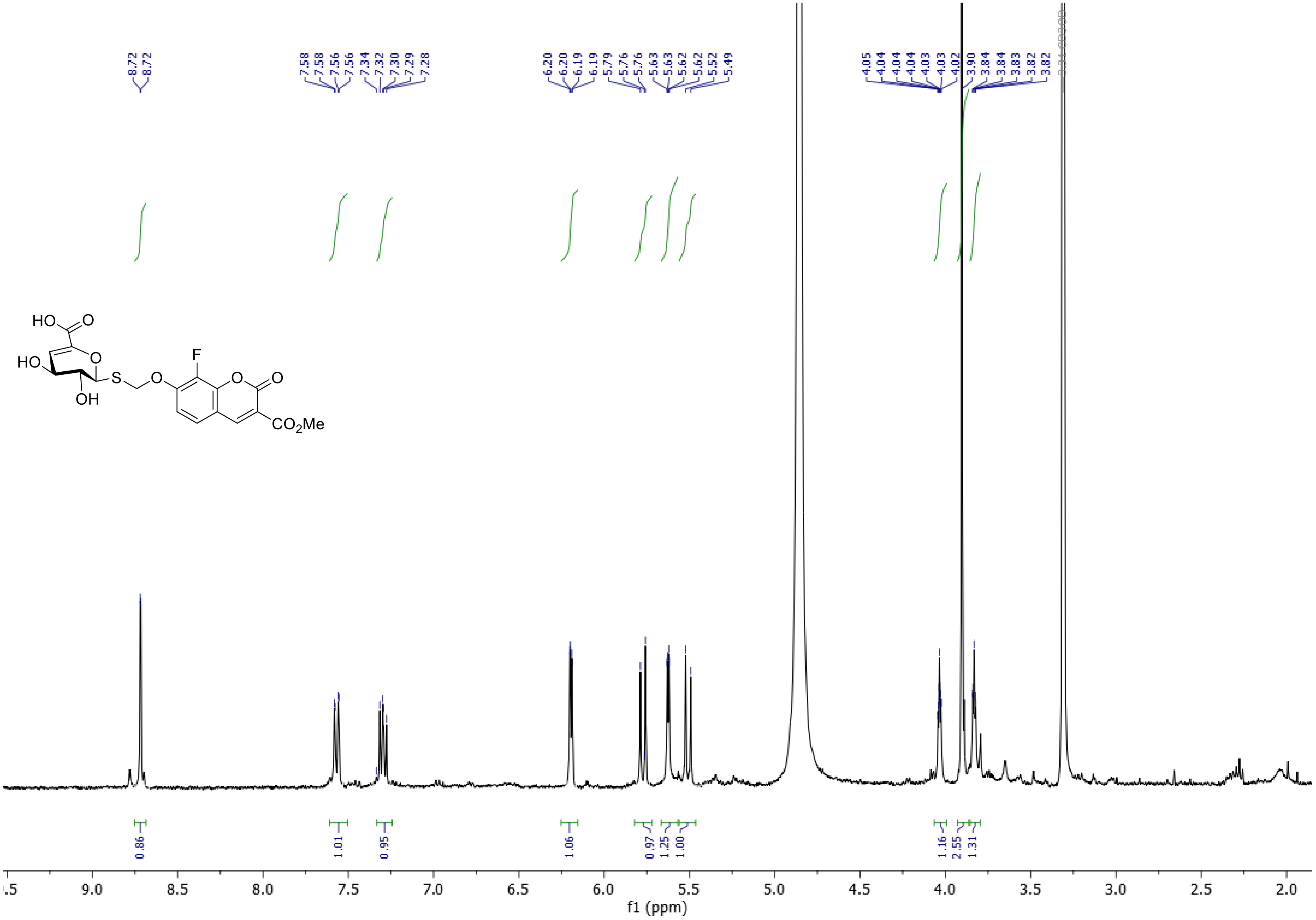

HO
O
HO
O
S
O
F
O
O
OH
CO2Me
f1 (ppm)

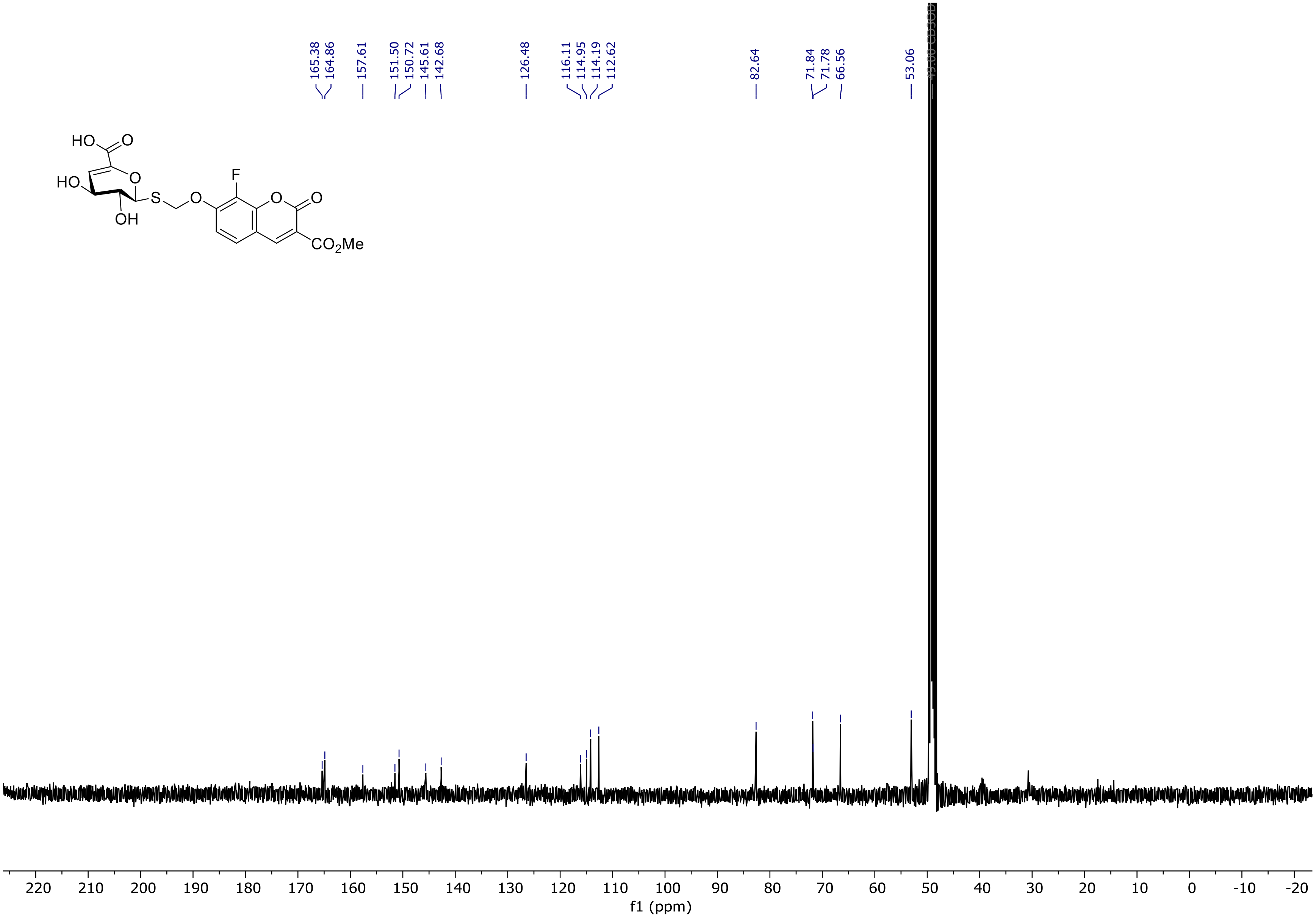
165.38
164.86
157.61
151.50
150.72
145.61
142.68
126.48
116.11
114.95
114.19
112.62
82.64
71.84
71.78
66.56
53.06
HO
O
HO
O
S
O
F
O
O
OH
CO2Me
220 210 200 190 180 170 160 150 140 130 120 110 100 90 80 70 60 50 40 30 20 10 0 -10 -20
f1 (ppm)

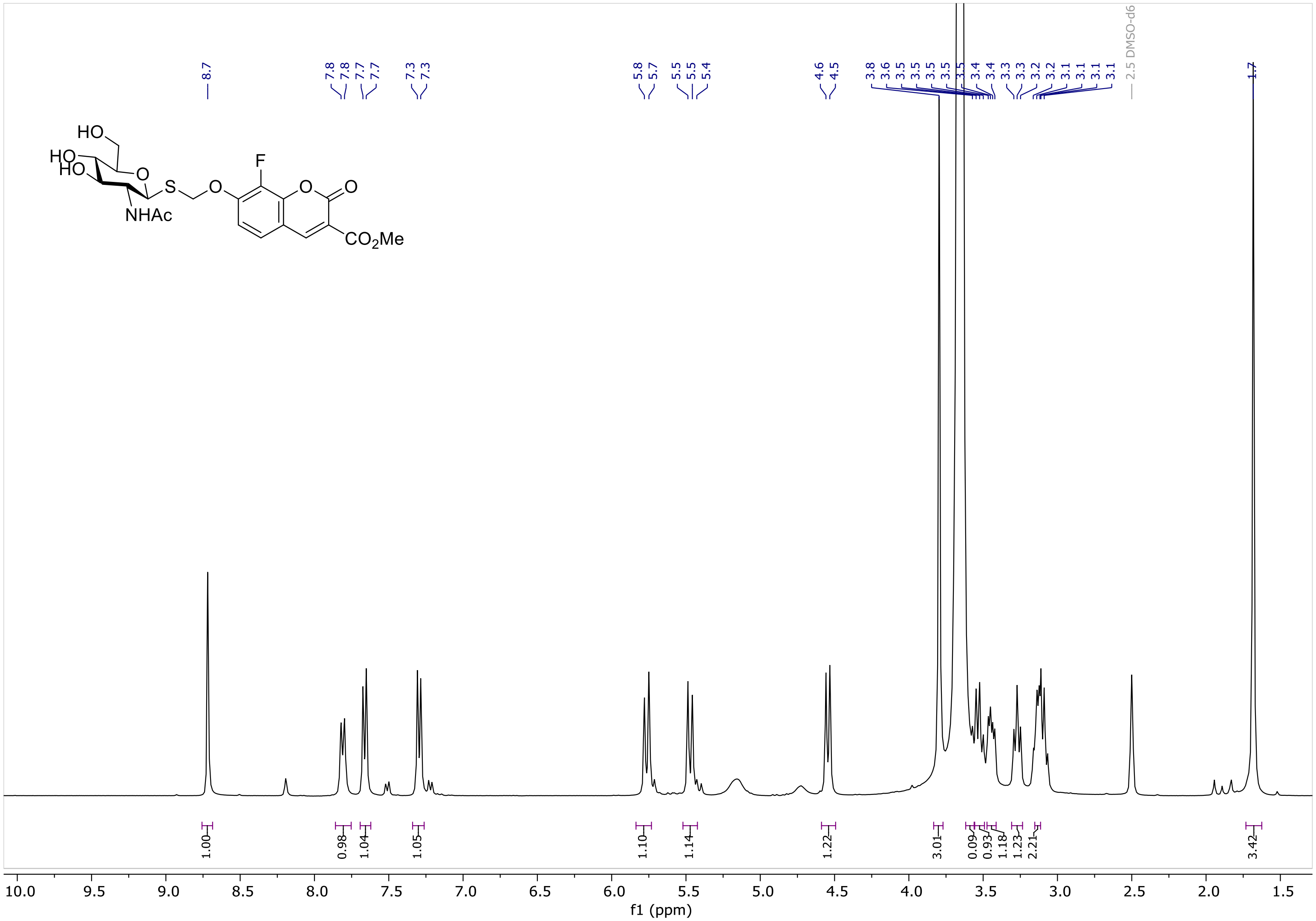
HO
HO
HO
O
S
O
F
O
O
NHAc
CO2Me
8.7
7.8
7.8
7.7
7.7
7.3
7.3
5.8
5.7
5.5
5.5
5.4
4.6
4.5
3.8
3.6
3.5
3.5
3.5
3.5
3.5
3.4
3.4
3.3
3.3
3.2
3.2
3.1
3.1
3.1
3.1
2.5 DMSO-d6
1.7
1.00
0.98
1.04
1.05
1.10
1.14
1.22
3.01
0.09
0.93
1.18
1.23
2.21
3.42
10.0
9.5
9.0
8.5
8.0
7.5
7.0
6.5
6.0
5.5
5.0
4.5
4.0
3.5
3.0
2.5
2.0
1.5
f1 (ppm)

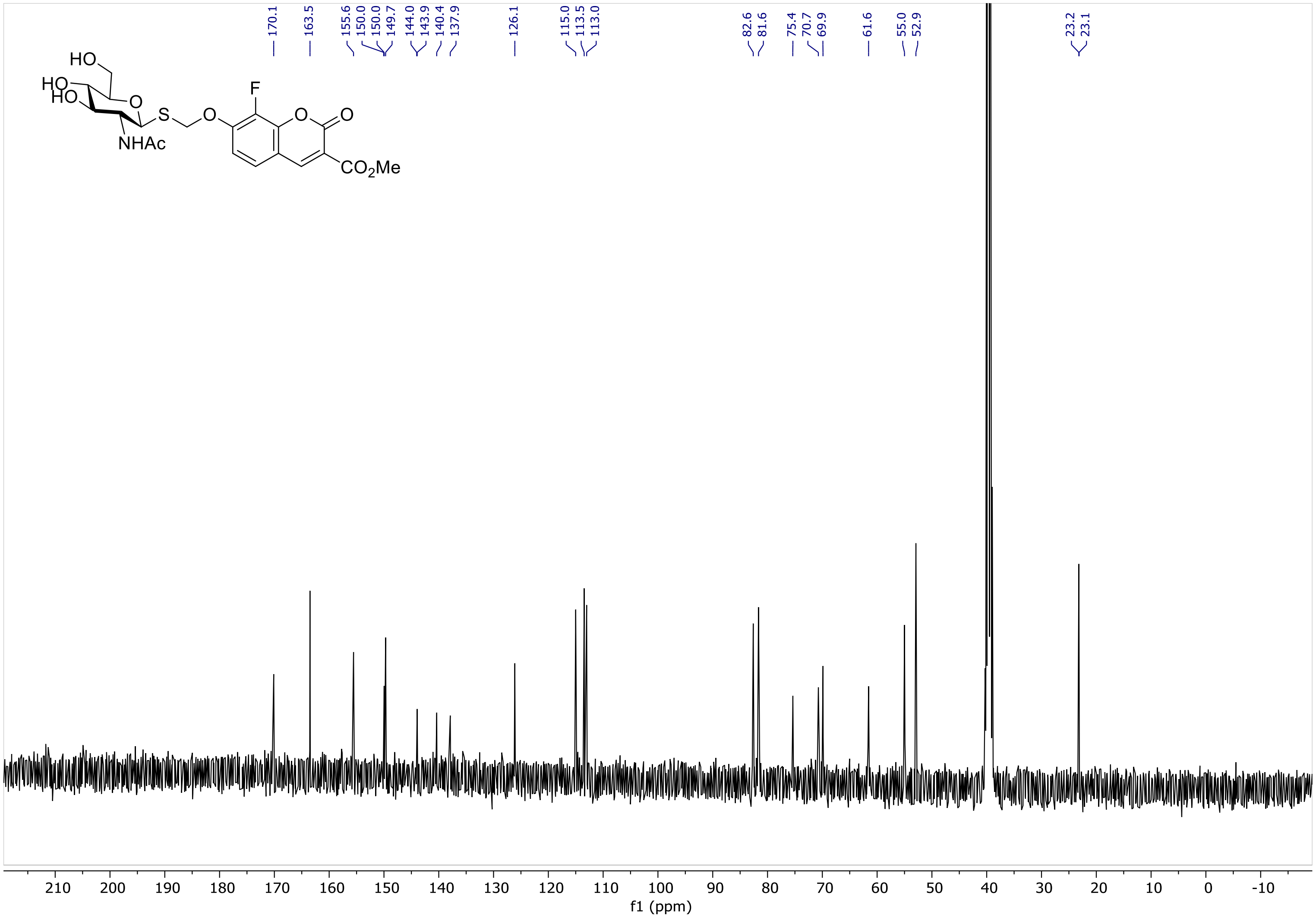

170.1
163.5
155.6
150.0
150.0
149.7
144.0
143.9
140.4
137.9
126.1
115.0
113.5
113.0
82.6
81.6
75.4
70.7
69.9
61.6
55.0
52.9
23.2
23.1
HO
HO
HO
O
S
O
F
O
O
NHAc
CO2Me
210
200
190
180
170
160
150
140
130
120
110
100
90
80
70
60
50
40
30
20
10
0
-10
f1 (ppm)

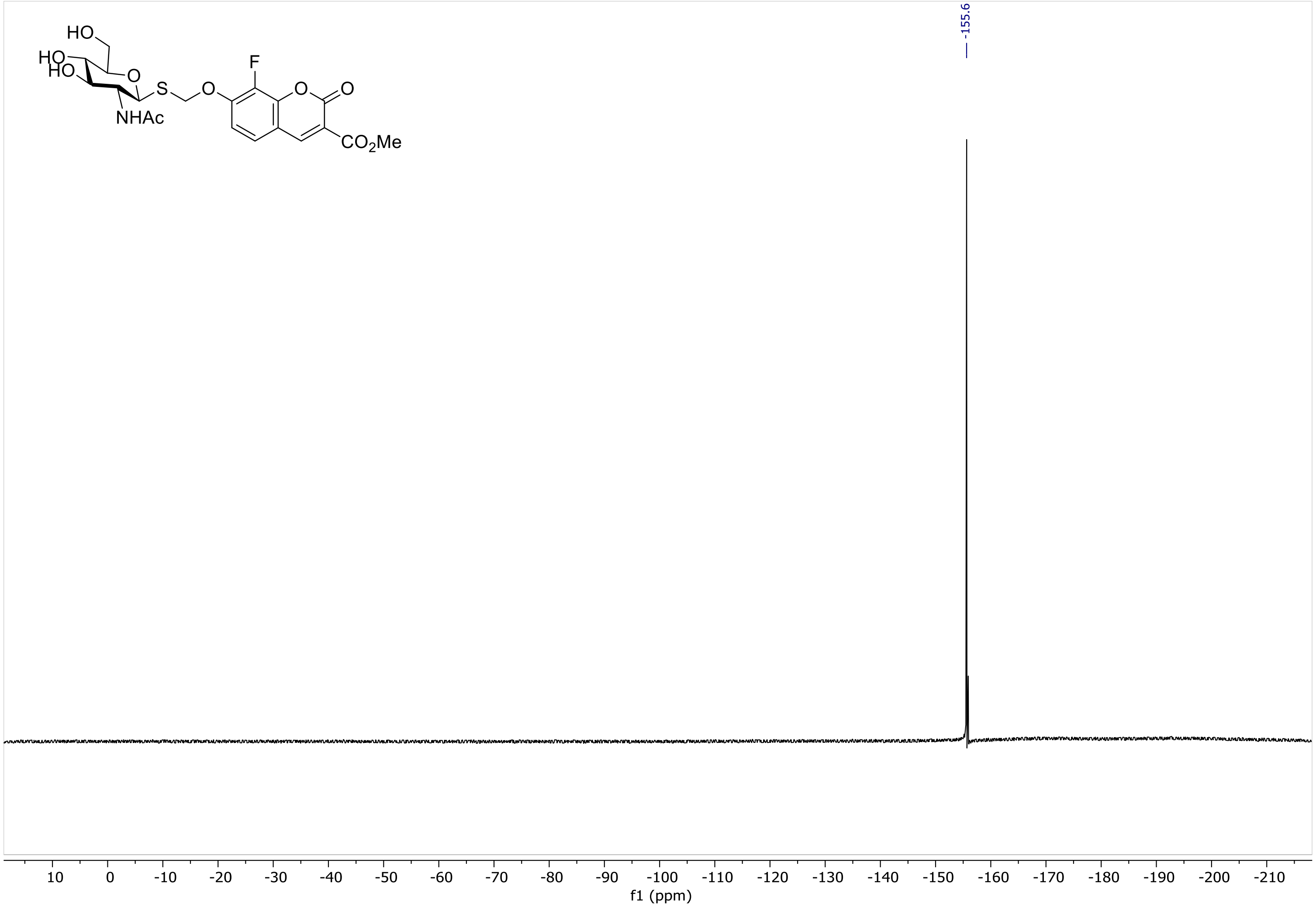
-155.6
HO
HO
HO
O
S
O
F
O
O
NHAc
CO2Me
10 0 -10 -20 -30 -40 -50 -60 -70 -80 -90 -100 -110 -120 -130 -140 -150 -160 -170 -180 -190 -200 -210
f1 (ppm)

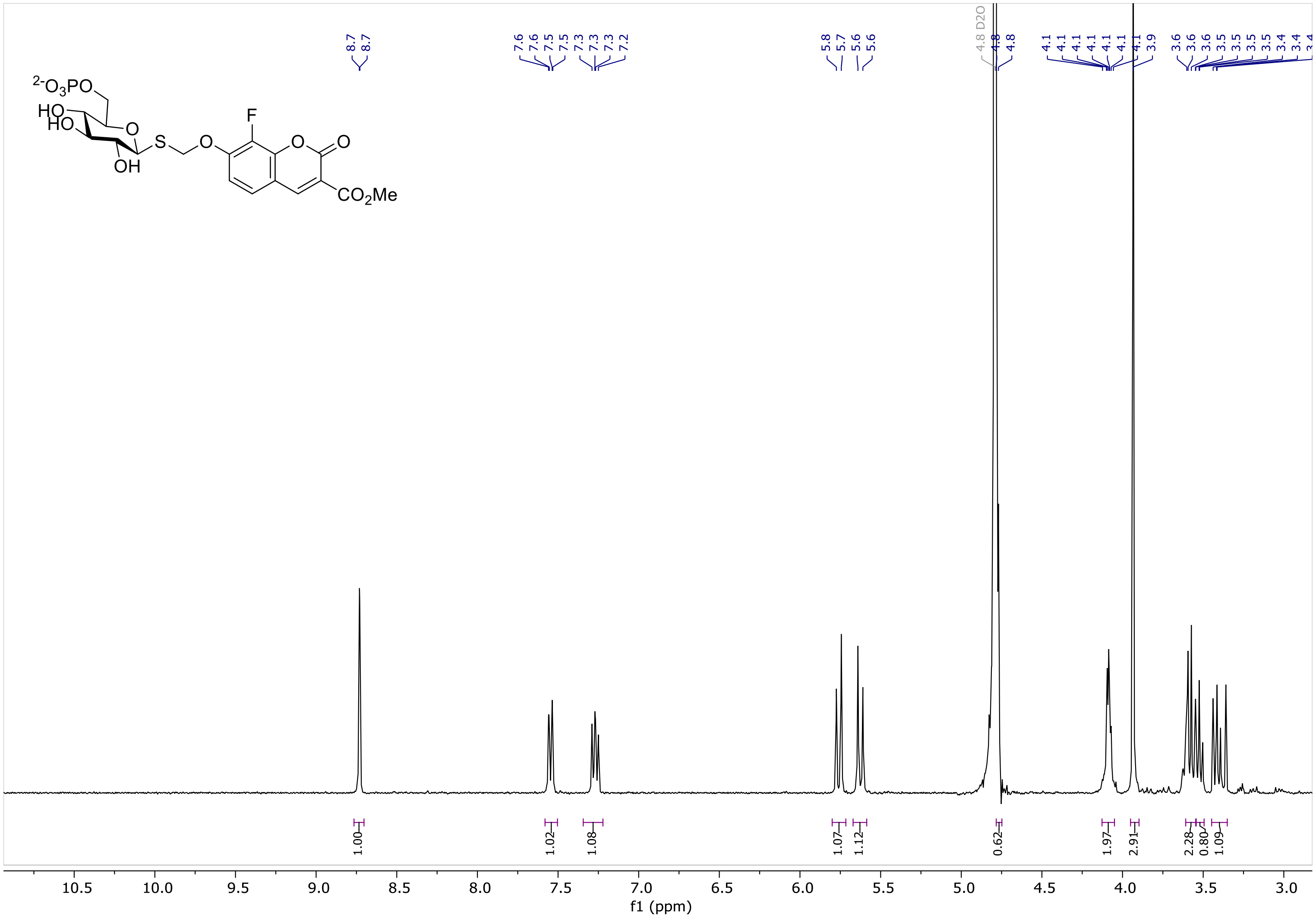
$^{2-}O_3PO$
HO
HO
O
S
O
F
O
O
OH
$CO_2Me$
8.7
8.7
7.6
7.6
7.5
7.5
7.3
7.3
7.3
7.2
5.8
5.7
5.6
5.6
4.8 D2O
4.8
4.8
4.1
4.1
4.1
4.1
4.1
4.1
4.1
3.9
3.6
3.6
3.6
3.5
3.5
3.5
3.5
3.4
3.4
1.00
1.02
1.08
1.07
1.12
0.62
1.97
2.91
2.28
0.80
1.09
10.5
10.0
9.5
9.0
8.5
8.0
7.5
7.0
6.5
6.0
5.5
5.0
4.5
4.0
3.5
3.0
f1 (ppm)

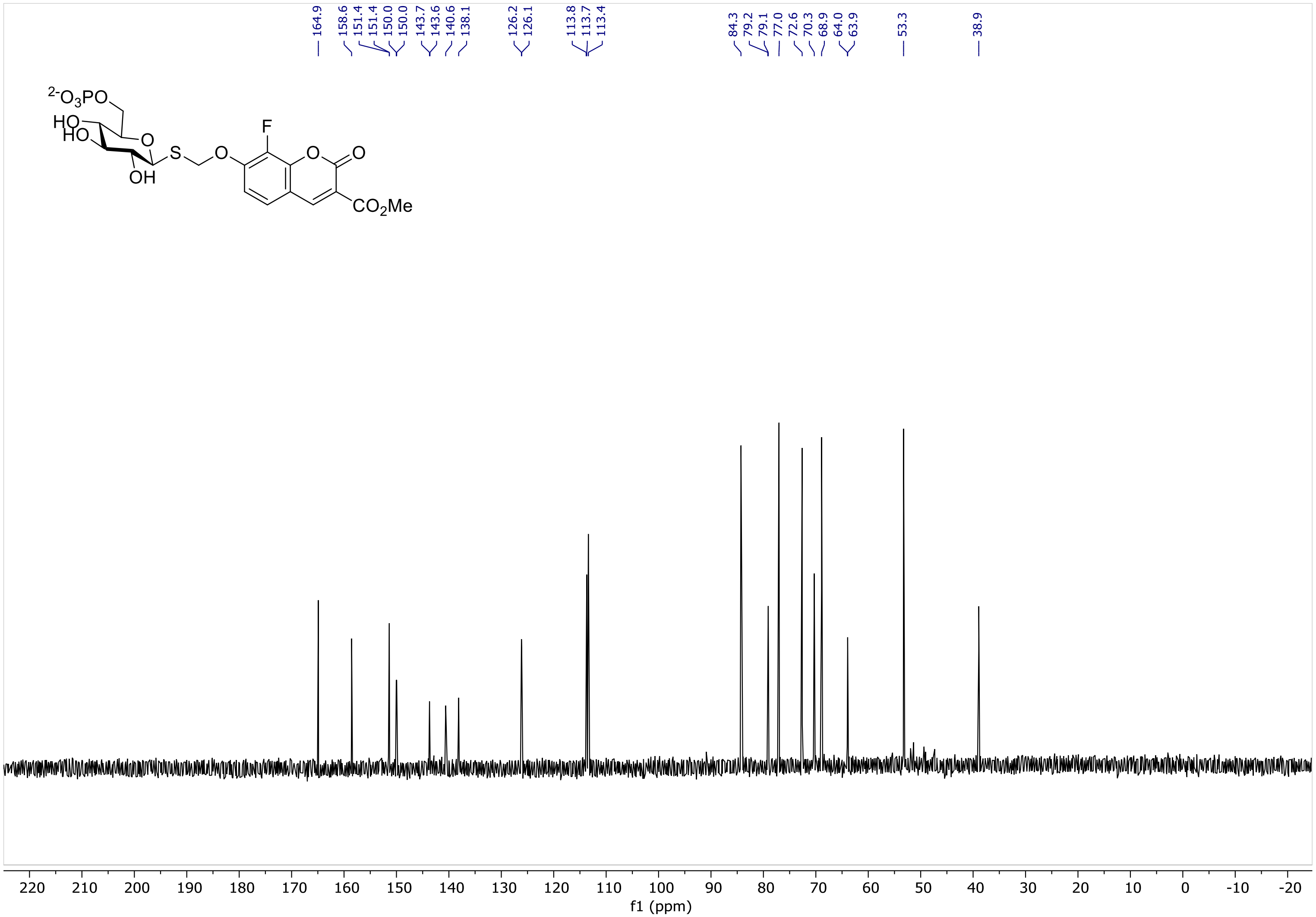

164.9
158.6
151.4
151.4
150.0
150.0
143.7
143.6
140.6
138.1
126.2
126.1
113.8
113.7
113.4
84.3
79.2
79.1
77.0
72.6
70.3
68.9
64.0
63.9
53.3
38.9
220 210 200 190 180 170 160 150 140 130 120 110 100 90 80 70 60 50 40 30 20 10 0 -10 -20
f1 (ppm)

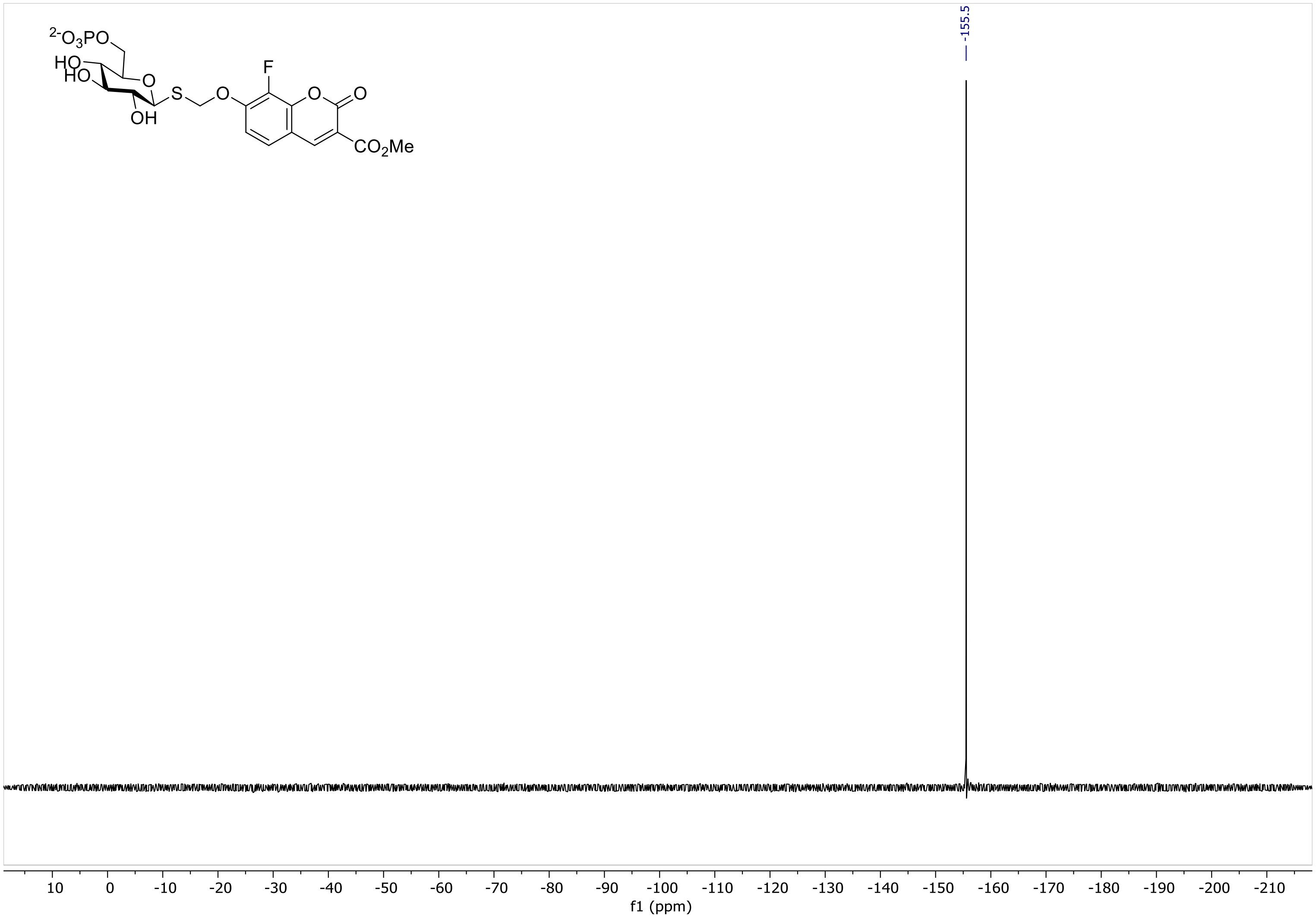

-155.5
F
O
O
O
S
OH
HO
HO
$^{2-}O_3PO$
$CO_2Me$
10
0
-10
-20
-30
-40
-50
-60
-70
-80
-90
-100
-110
-120
-130
-140
-150
-160
-170
-180
-190
-200
-210
f1 (ppm)

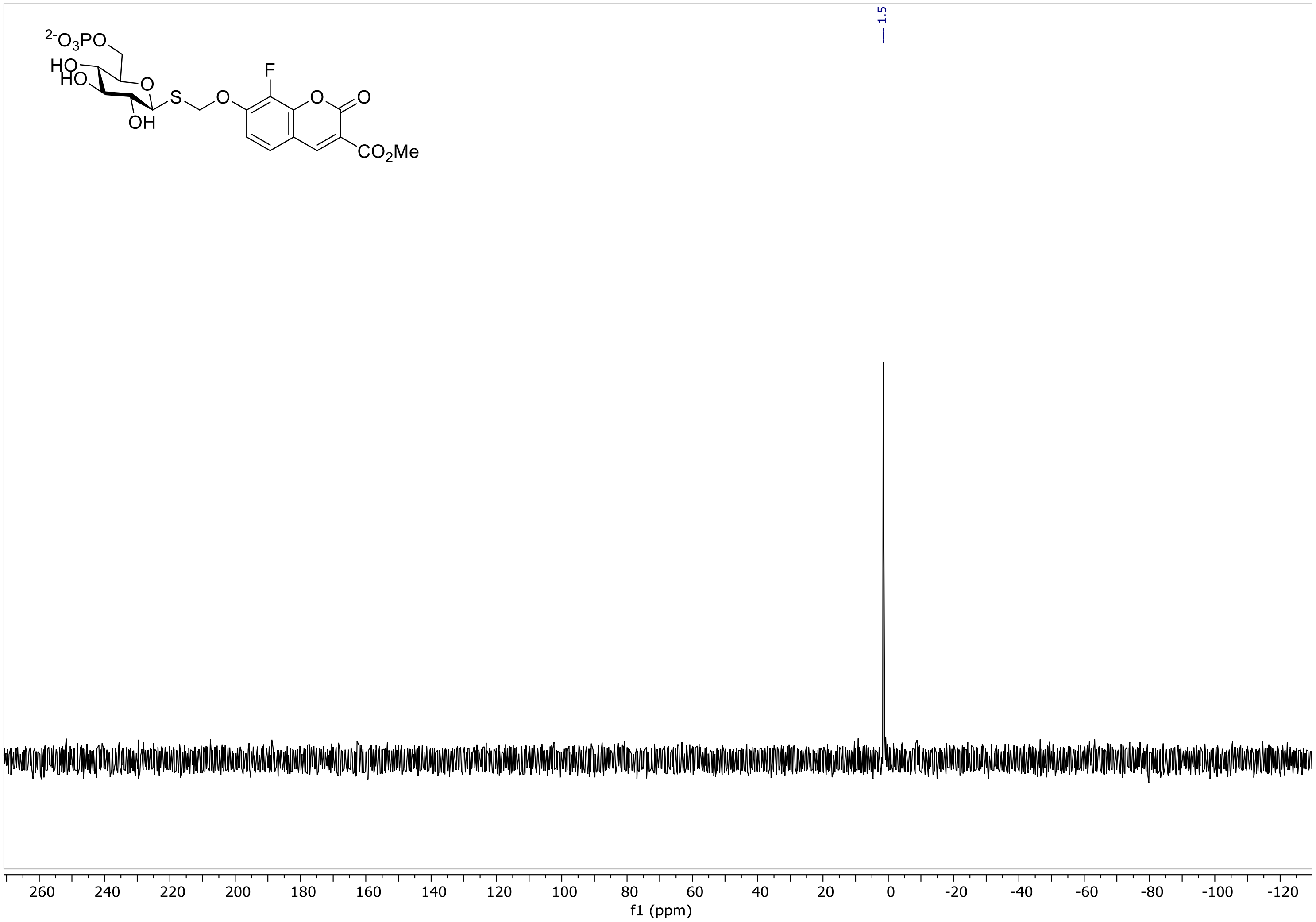

1.5
f1 (ppm)

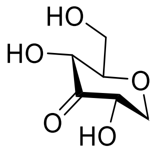
HO
HO
O
O
HO

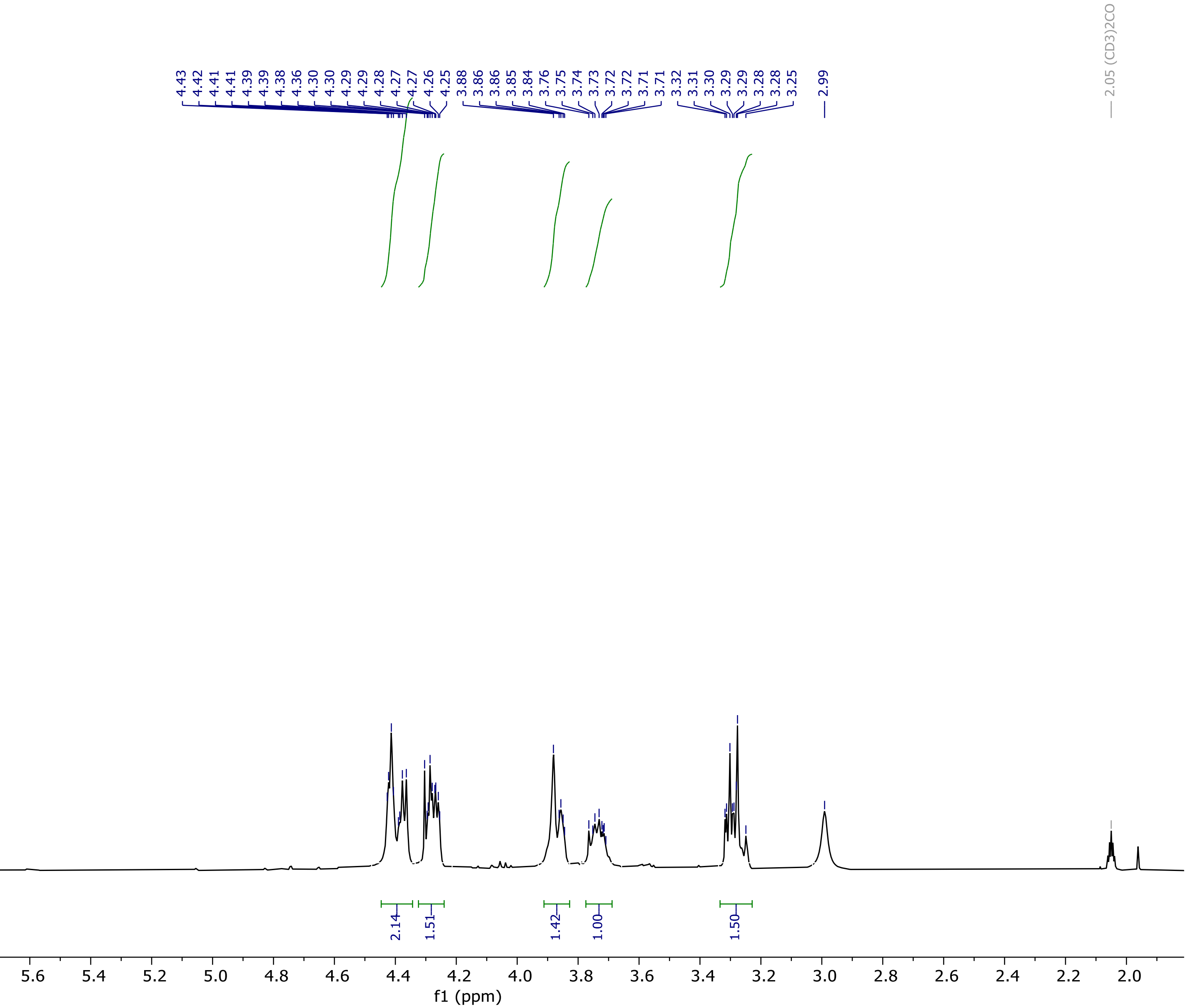
4.43
4.42
4.41
4.41
4.39
4.39
4.38
4.36
4.30
4.30
4.29
4.29
4.28
4.27
4.27
4.26
4.25
3.88
3.86
3.86
3.85
3.84
3.76
3.75
3.74
3.73
3.72
3.72
3.71
3.71
3.32
3.31
3.30
3.29
3.29
3.29
3.28
3.25
2.99
2.05 (CD3)2CO
2.14
1.51
1.42
1.00
1.50
6.4
6.2
6.0
5.8
5.6
5.4
5.2
5.0
4.8
4.6
4.4
4.2
4.0
3.8
3.6
3.4
3.2
3.0
2.8
2.6
2.4
2.2
2.0
f1 (ppm)

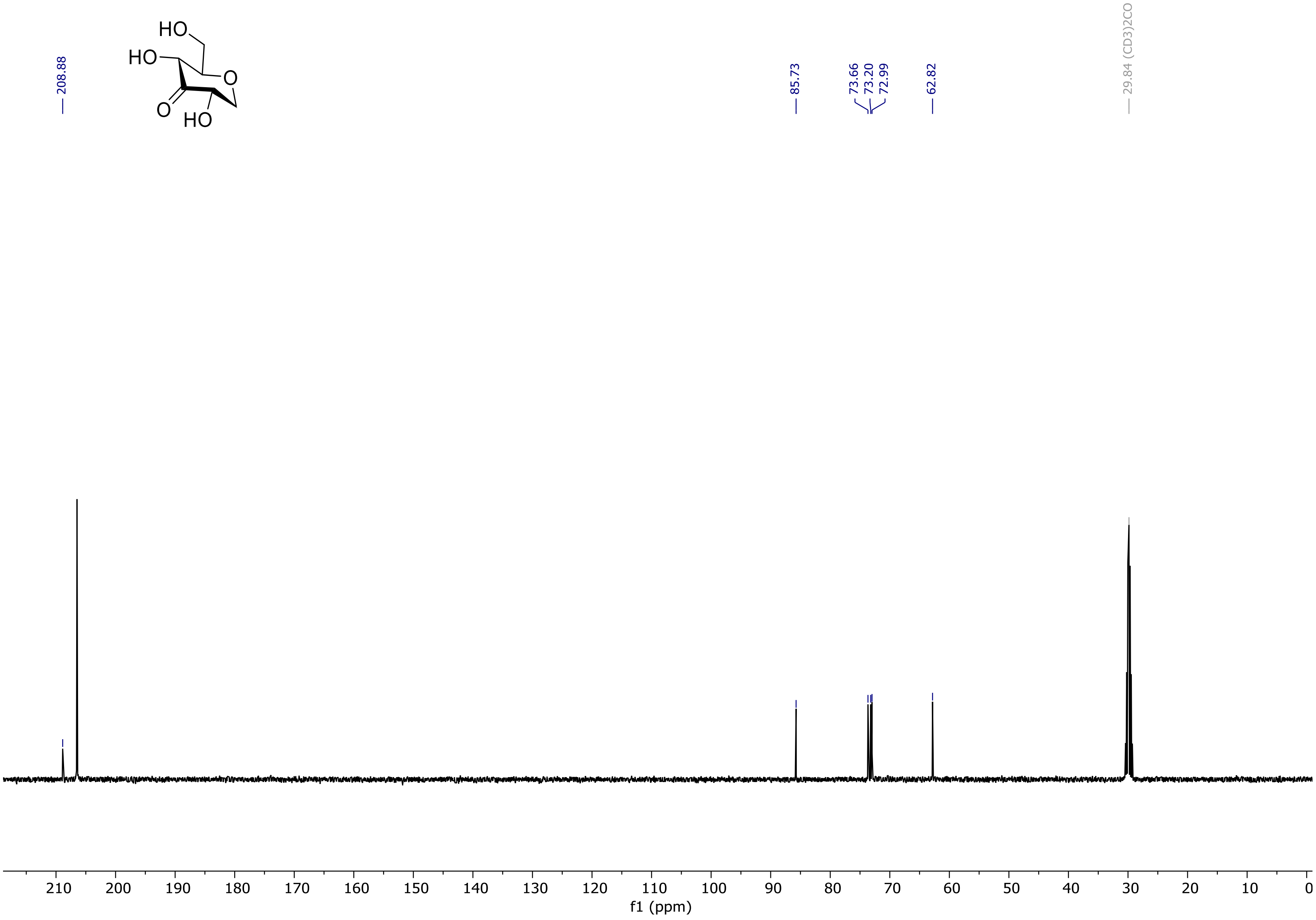

208.88
85.73
73.66
73.20
72.99
62.82
29.84 (CD3)2CO
HO
HO
O
O
HO
210 200 190 180 170 160 150 140 130 120 110 100 90 80 70 60 50 40 30 20 10 0
f1 (ppm)

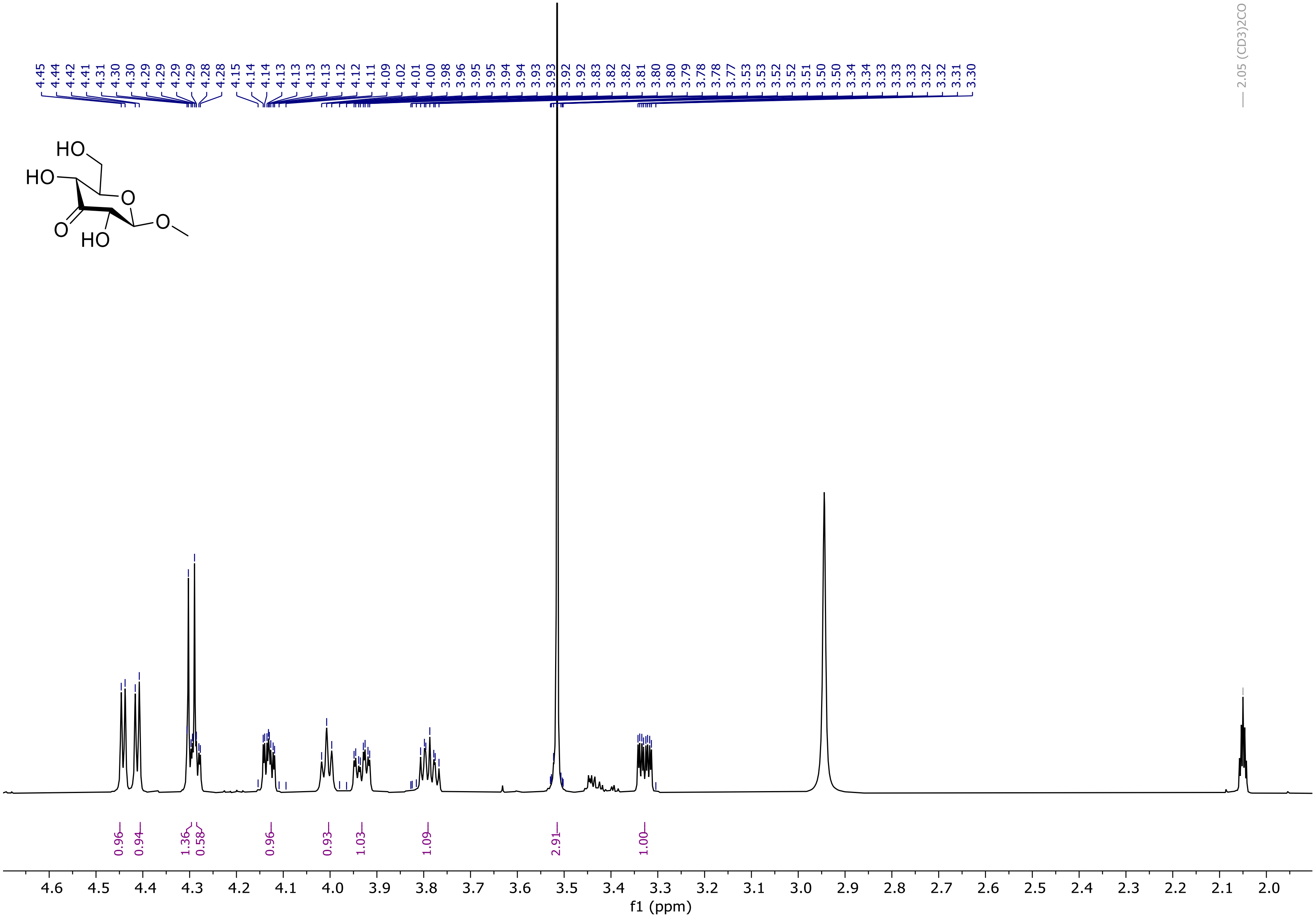
HO
HO
O
O
HO
O
f1 (ppm)
2.05 (CD3)2CO

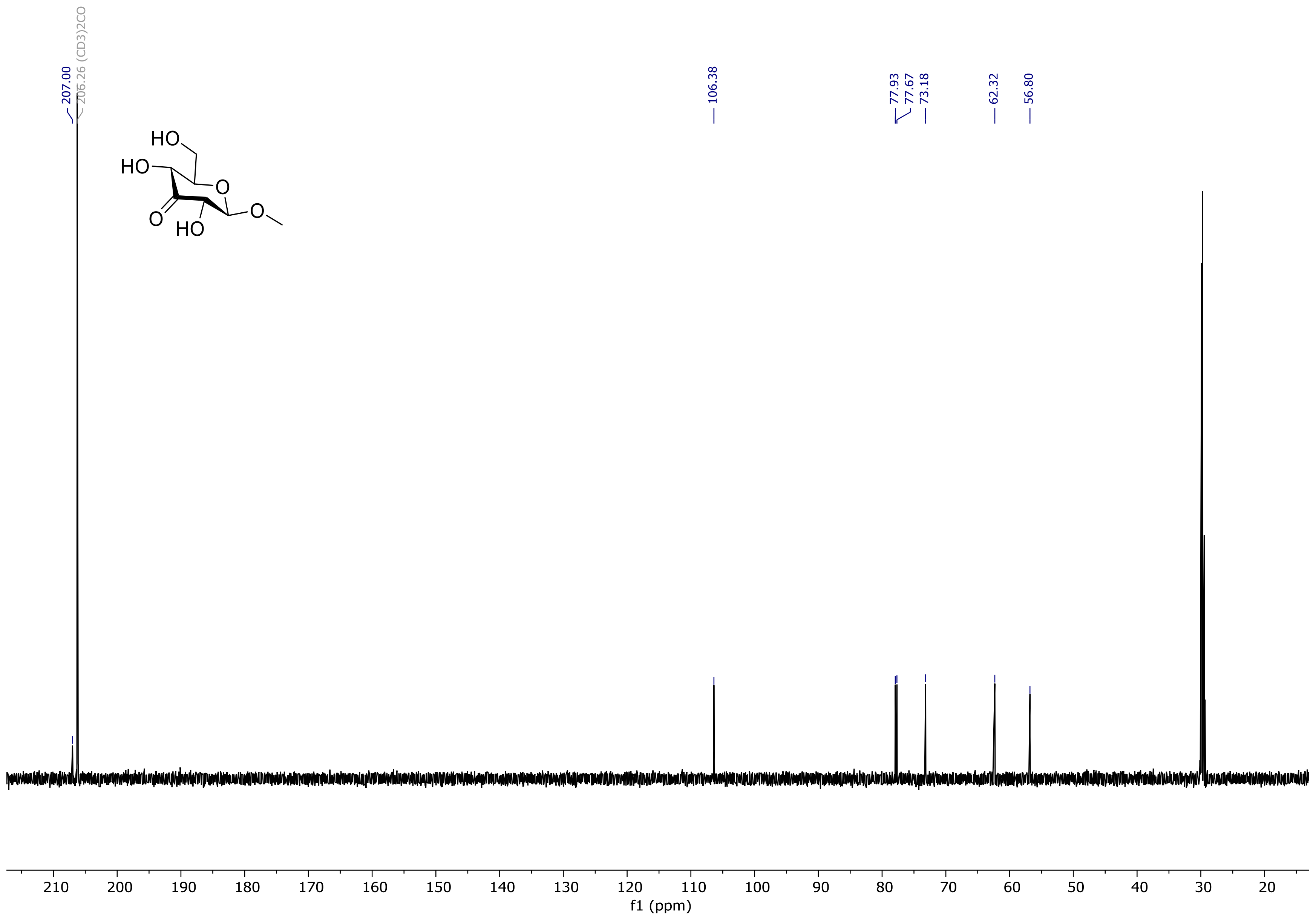

207.00
206.26 (CD3)2CO
106.38
77.93
77.67
73.18
62.32
56.80
HO
HO
O
O
HO
O
210
200
190
180
170
160
150
140
130
120
110
100
90
80
70
60
50
40
30
20
f1 (ppm)

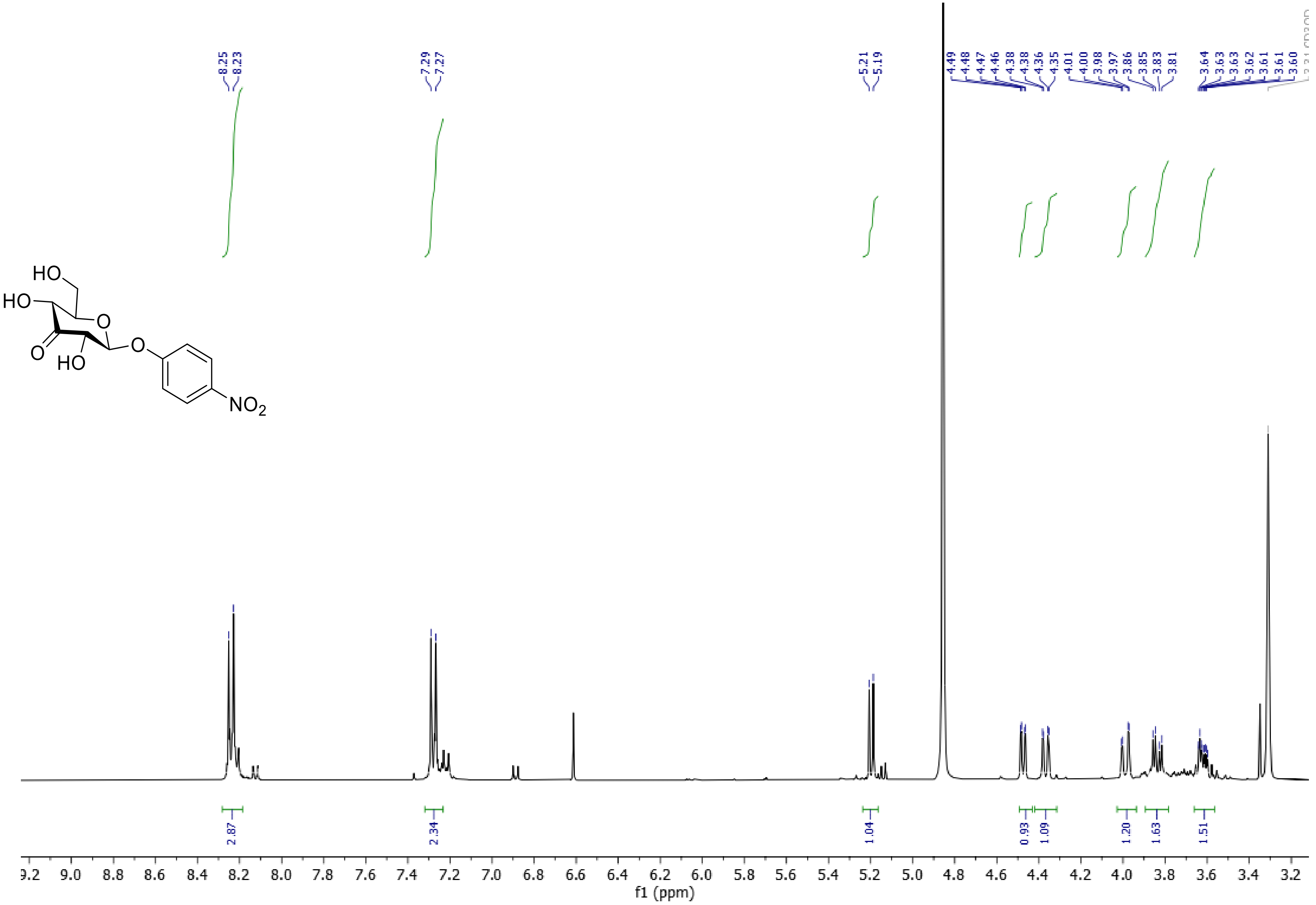
8.25
8.23
7.29
7.27
5.21
5.19
4.49
4.48
4.47
4.46
4.38
4.38
4.36
4.35
4.01
4.00
3.98
3.97
3.86
3.85
3.83
3.81
3.64
3.63
3.63
3.62
3.61
3.61
3.60
HO
HO
O
O
O
HO
$NO_2$
2.87
2.34
1.04
0.93
1.09
1.20
1.63
1.51
9.2 9.0 8.8 8.6 8.4 8.2 8.0 7.8 7.6 7.4 7.2 7.0 6.8 6.6 6.4 6.2 6.0 5.8 5.6 5.4 5.2 5.0 4.8 4.6 4.4 4.2 4.0 3.8 3.6 3.4 3.2
f1 (ppm)

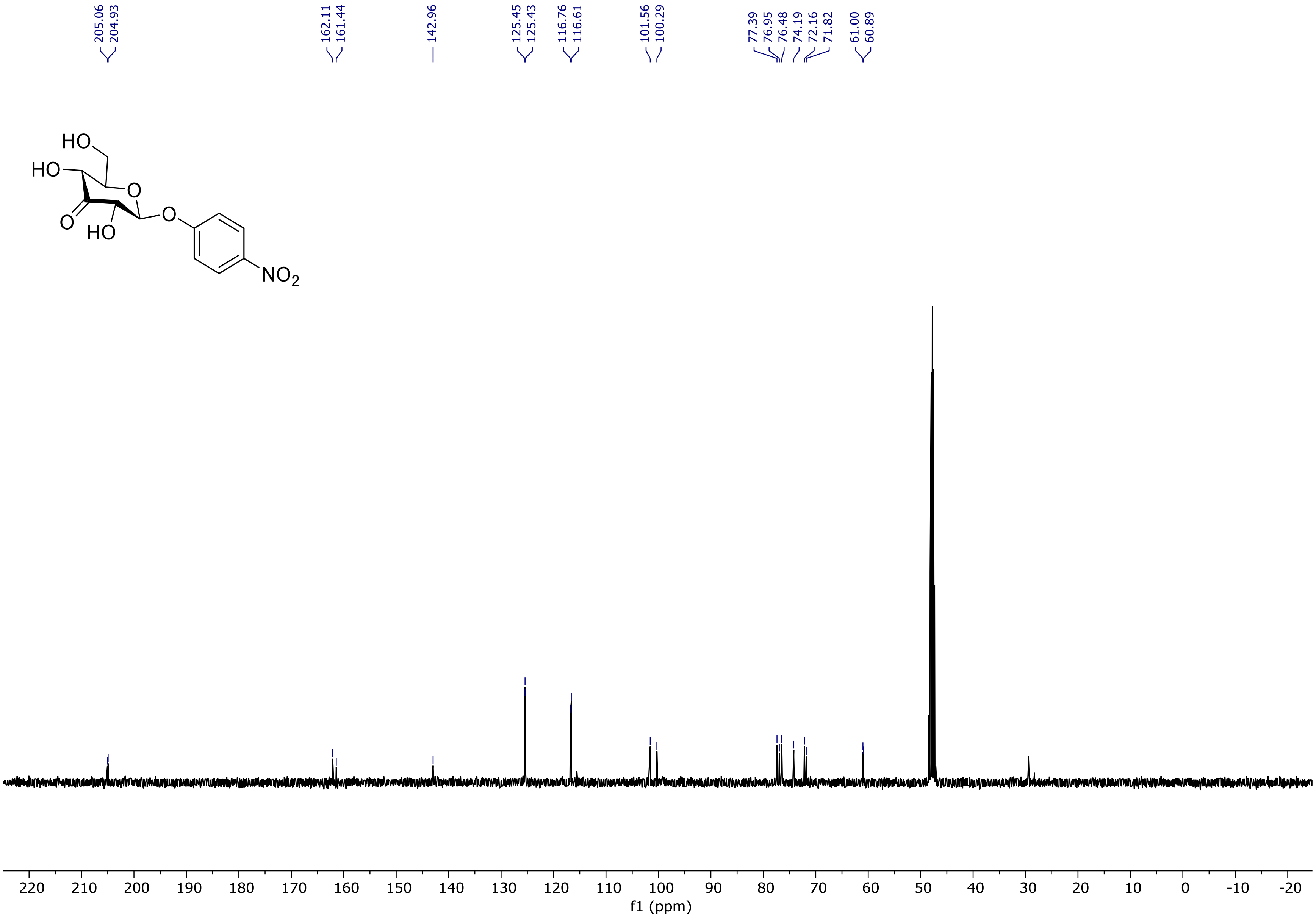
HO
HO
O
O
HO
O
NO2
205.06
204.93
162.11
161.44
142.96
125.45
125.43
116.76
116.61
101.56
100.29
77.39
76.95
76.48
74.19
72.16
71.82
61.00
60.89
220 210 200 190 180 170 160 150 140 130 120 110 100 90 80 70 60 50 40 30 20 10 0 -10 -20
f1 (ppm)

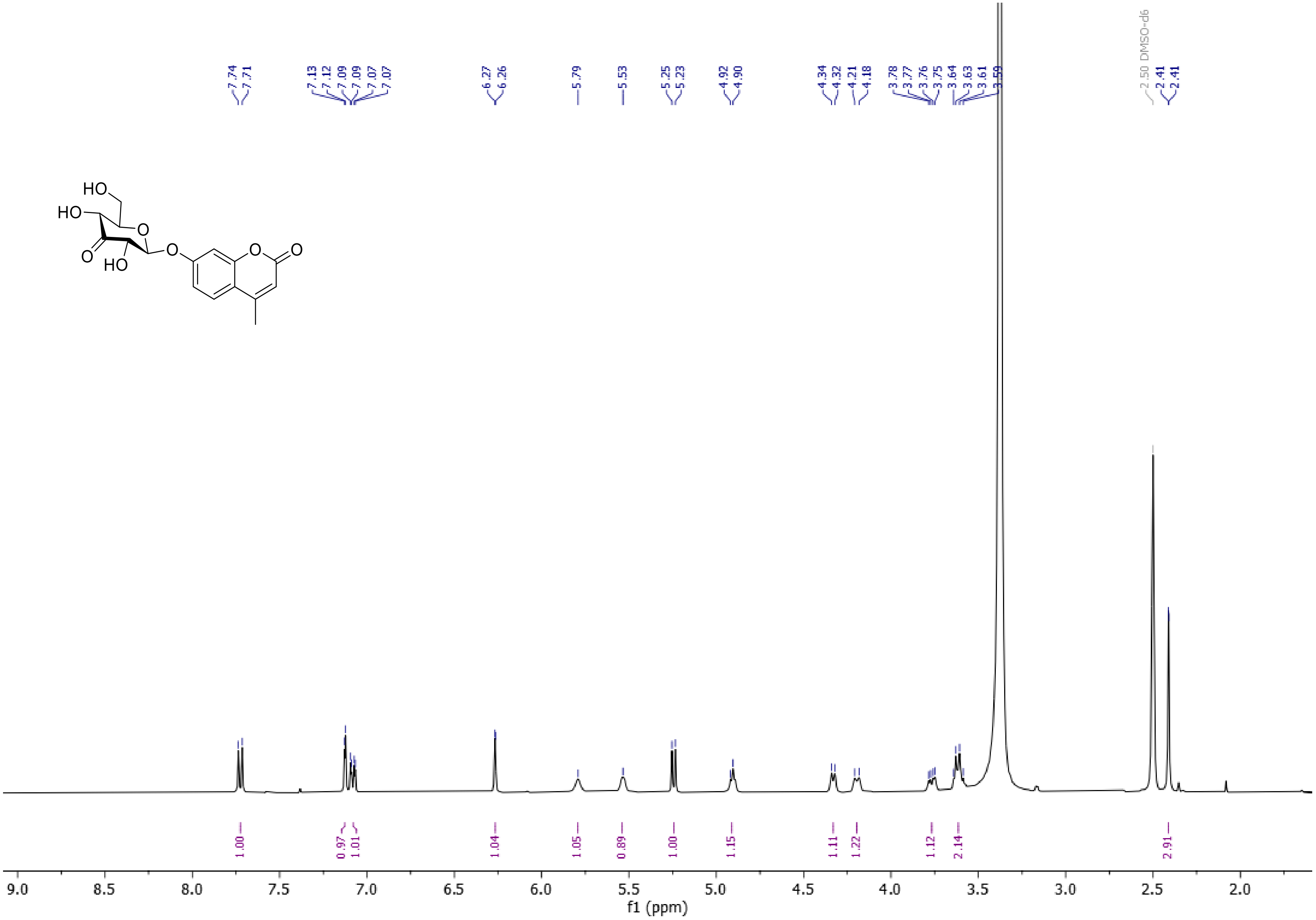
7.74
7.71
7.13
7.12
7.09
7.09
7.07
7.07
6.27
6.26
5.79
5.53
5.25
5.23
4.92
4.90
4.34
4.32
4.21
4.18
3.78
3.77
3.76
3.75
3.64
3.63
3.61
3.59
2.50 DMSO-d6
2.41
2.41
1.00
0.97
1.01
1.04
1.05
0.89
1.00
1.15
1.11
1.22
1.12
2.14
2.91
f1 (ppm)

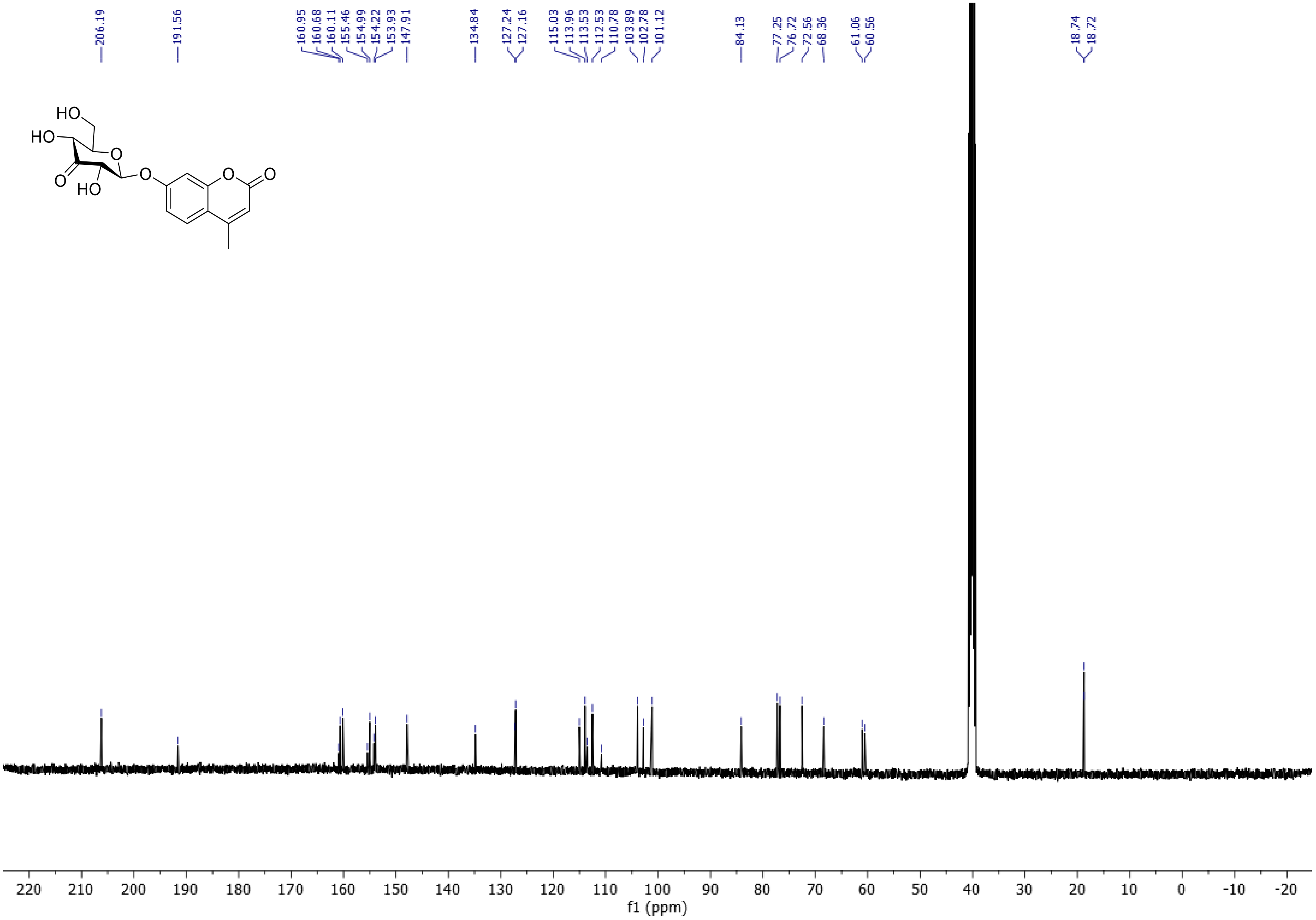

206.19
191.56
160.95
160.68
160.11
155.46
154.99
154.22
153.93
147.91
134.84
127.24
127.16
115.03
113.96
113.53
112.53
110.78
103.89
102.78
101.12
84.13
77.25
76.72
72.56
68.36
61.06
60.56
18.74
18.72
HO
HO
O
O
HO
O
O
O
220 210 200 190 180 170 160 150 140 130 120 110 100 90 80 70 60 50 40 30 20 10 0 -10 -20
f1 (ppm)

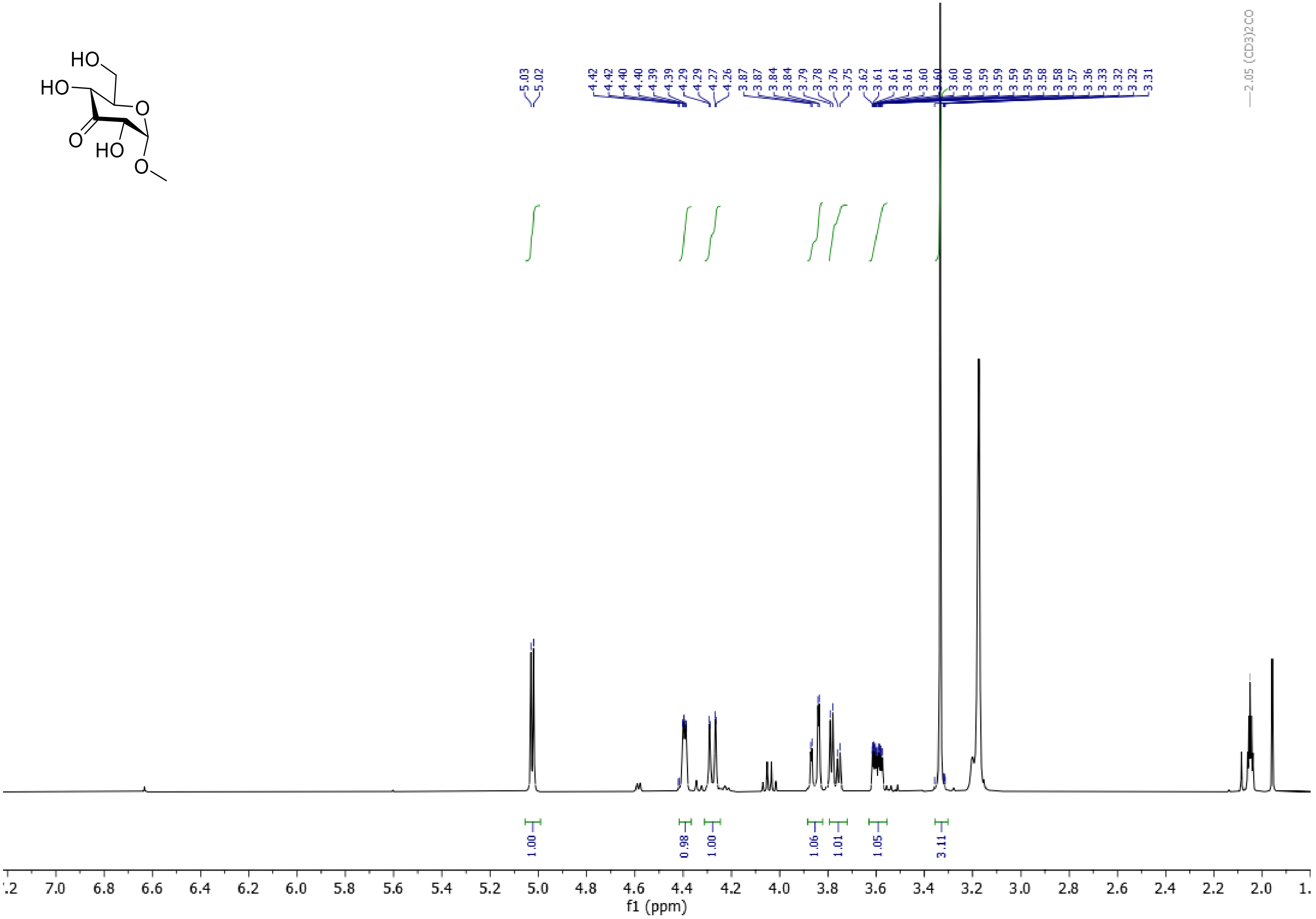
HO
HO
O
O
HO
O
2.05 (CD3)2CO
1.00
0.98
1.00
1.06
1.01
1.05
3.11
f1 (ppm)

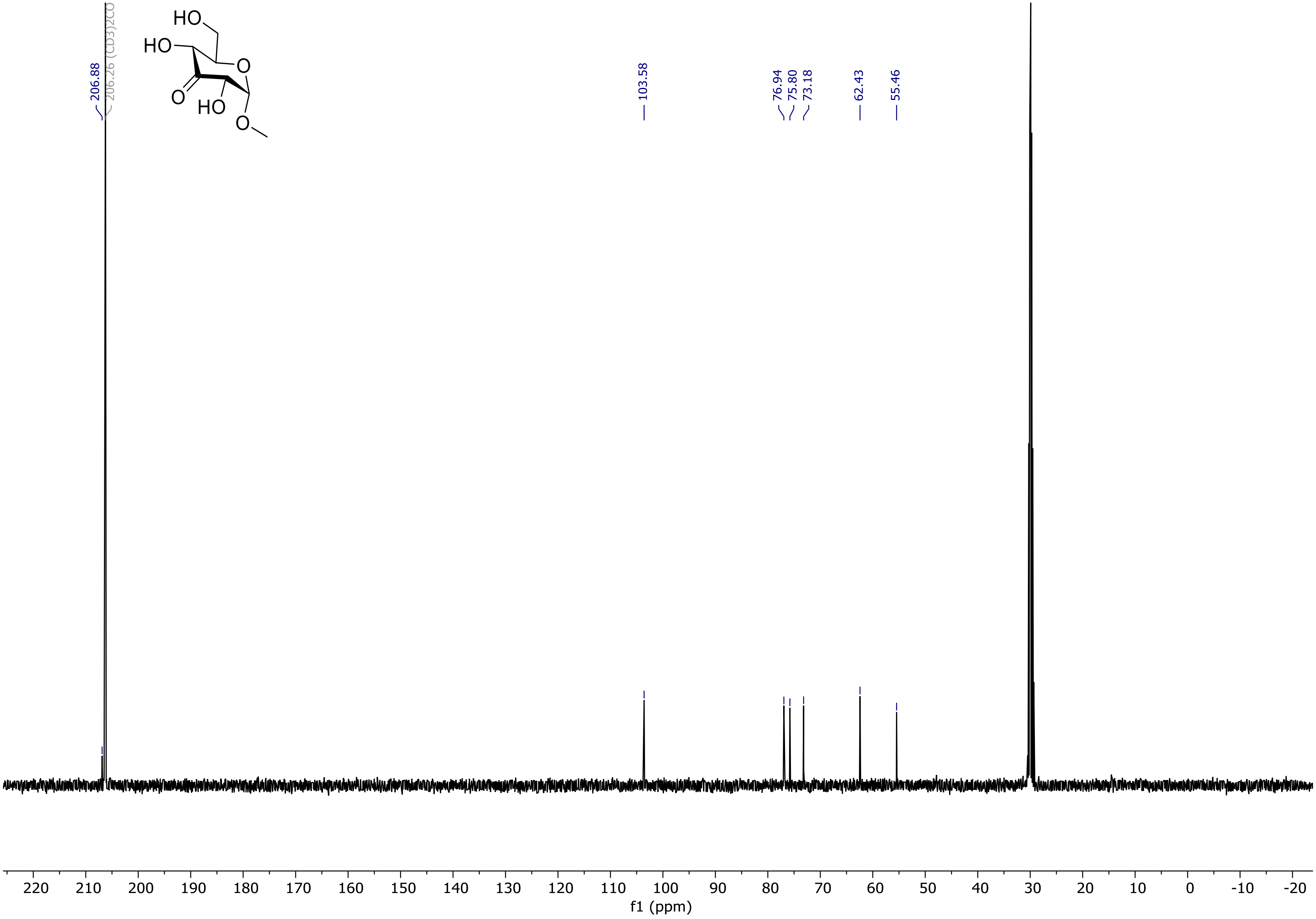

206.88
206.26 (CD3)2CO
HO
HO
O
O
HO
O
103.58
76.94
75.80
73.18
62.43
55.46
220 210 200 190 180 170 160 150 140 130 120 110 100 90 80 70 60 50 40 30 20 10 0 -10 -20
f1 (ppm)

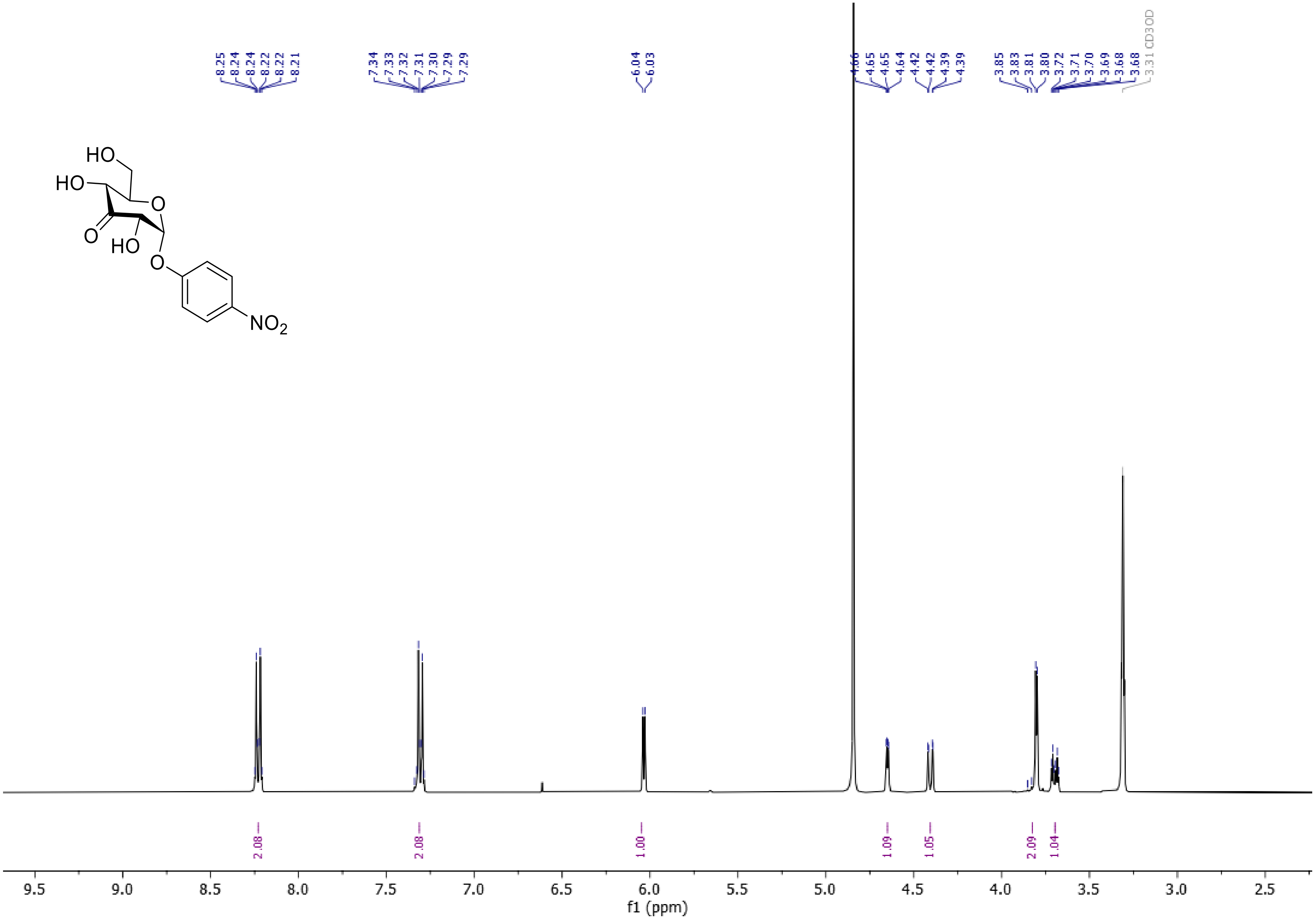
HO
HO
O
O
HO
O
NO2
8.25
8.24
8.24
8.22
8.22
8.21
7.34
7.33
7.32
7.31
7.30
7.29
7.29
6.04
6.03
4.66
4.65
4.65
4.64
4.42
4.42
4.39
4.39
3.85
3.83
3.81
3.80
3.72
3.71
3.70
3.69
3.68
3.68
3.31 CD3OD
2.08
2.08
1.00
1.09
1.05
2.09
1.04
9.5
9.0
8.5
8.0
7.5
7.0
6.5
6.0
5.5
5.0
4.5
4.0
3.5
3.0
2.5
f1 (ppm)

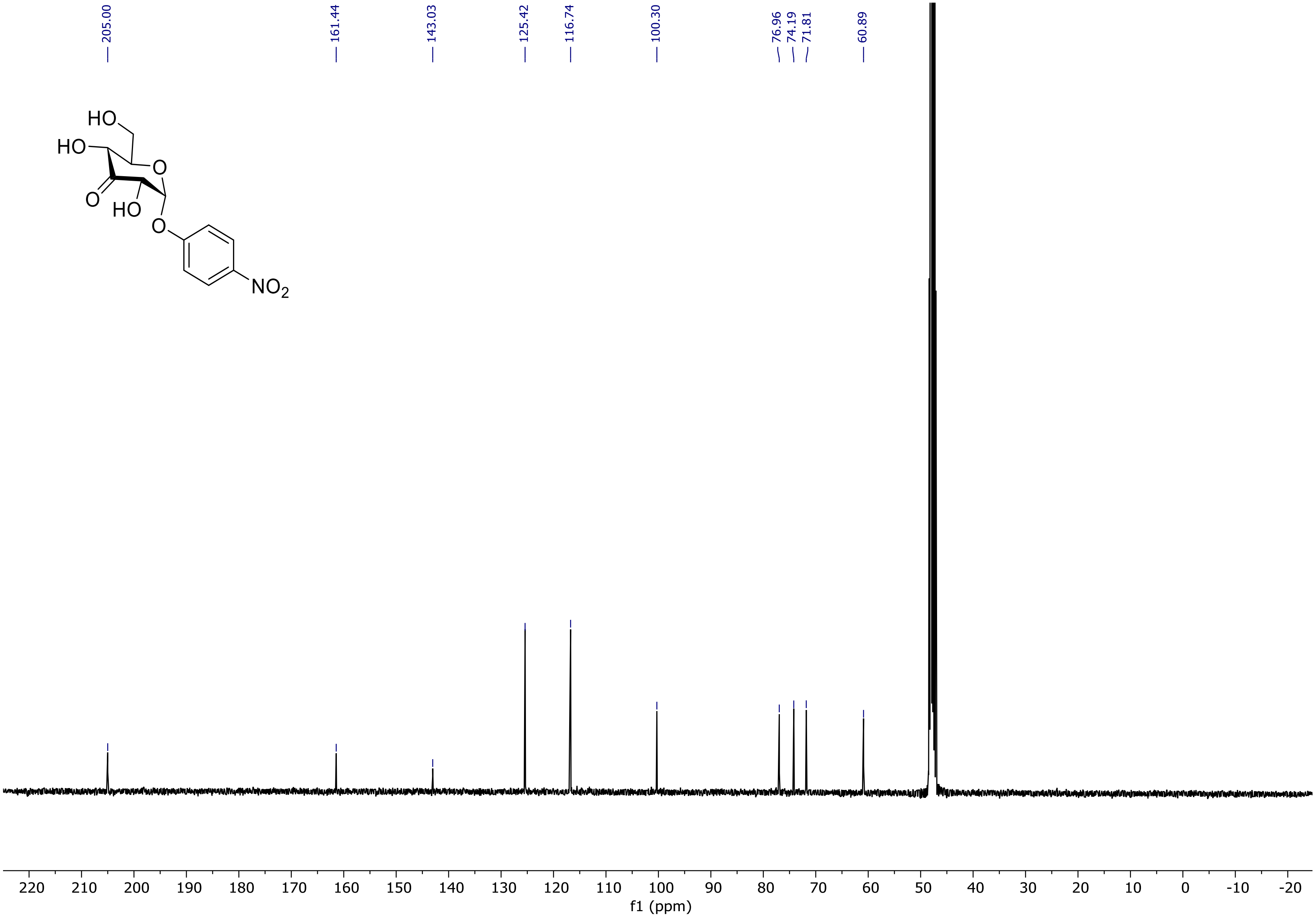
205.00
161.44
143.03
125.42
116.74
100.30
76.96
74.19
71.81
60.89
HO
HO
O
O
HO
O
NO2
220 210 200 190 180 170 160 150 140 130 120 110 100 90 80 70 60 50 40 30 20 10 0 -10 -20
f1 (ppm)

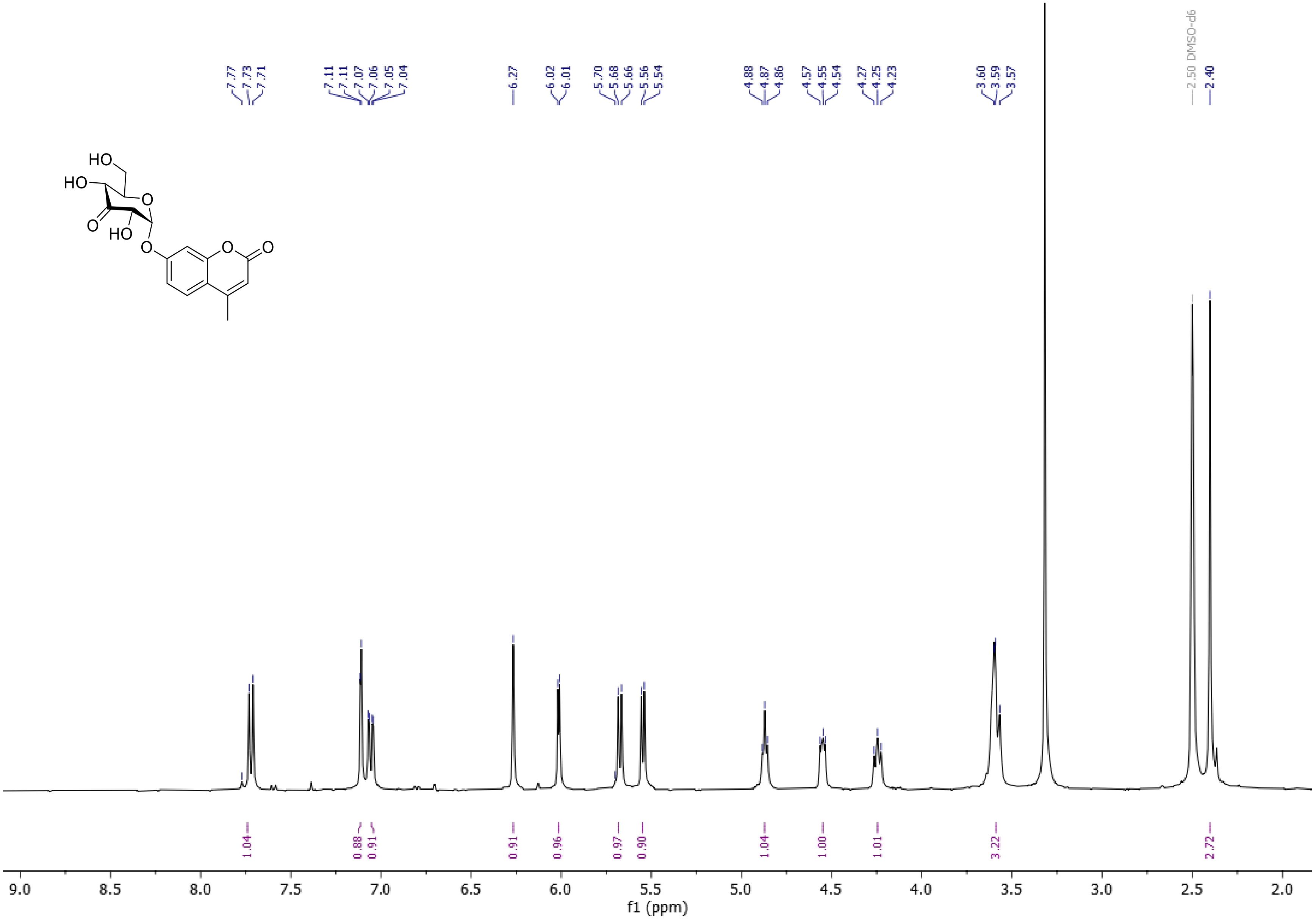
7.77
7.73
7.71
7.11
7.11
7.07
7.06
7.05
7.04
6.27
6.02
6.01
5.70
5.68
5.66
5.56
5.54
4.88
4.87
4.86
4.57
4.55
4.54
4.27
4.25
4.23
3.60
3.59
3.57
2.50 DMSO-d6
2.40
HO
HO
O
O
HO
O
O
O
1.04
0.88
0.91
0.91
0.96
0.97
0.90
1.04
1.00
1.01
3.22
2.72
9.0
8.5
8.0
7.5
7.0
6.5
6.0
5.5
5.0
4.5
4.0
3.5
3.0
2.5
2.0
f1 (ppm)

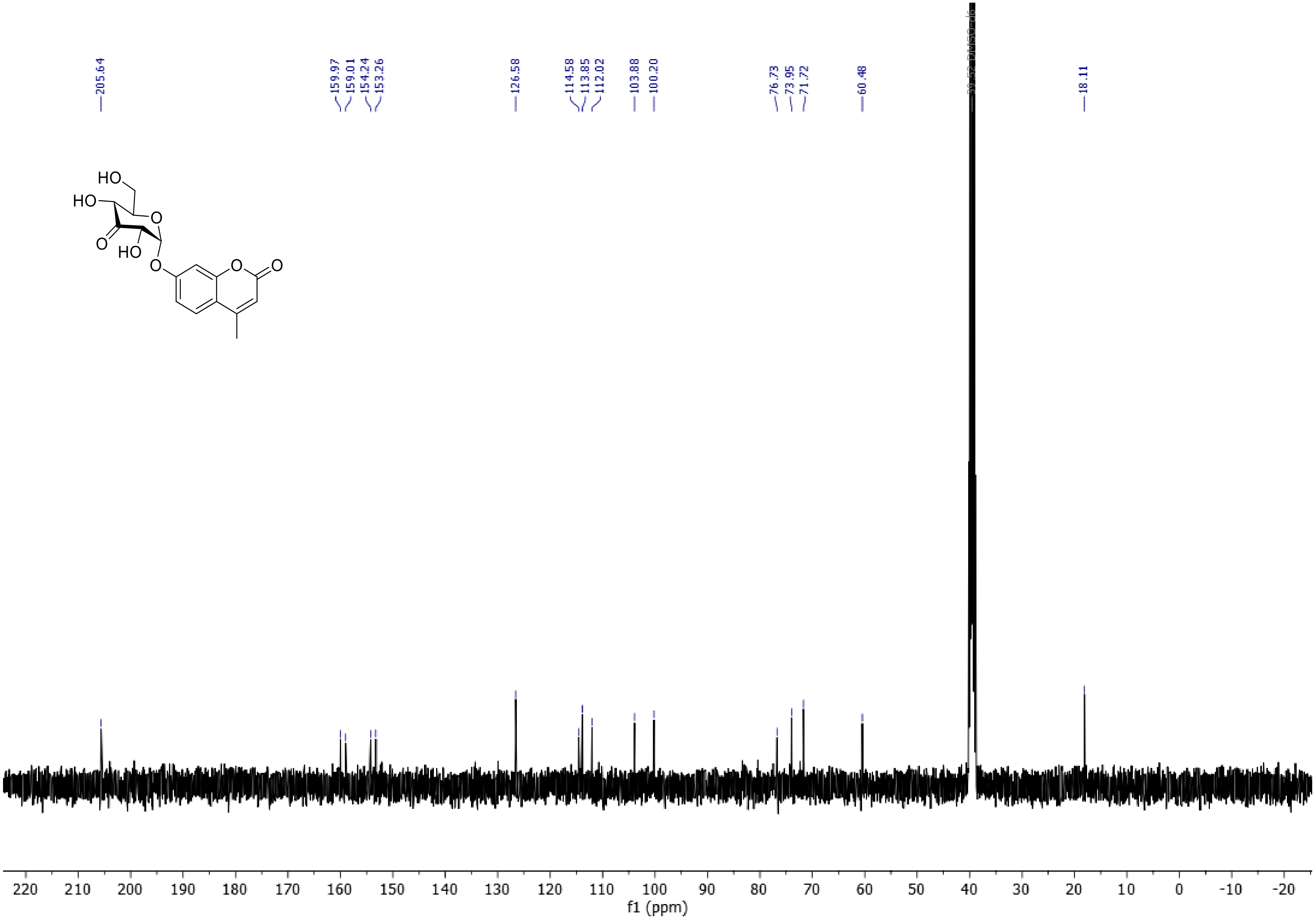

205.64
159.97
159.01
154.24
153.26
126.58
114.58
113.85
112.02
103.88
100.20
76.73
73.95
71.72
60.48
18.11
HO
HO
O
O
HO
O
O
O
220 210 200 190 180 170 160 150 140 130 120 110 100 90 80 70 60 50 40 30 20 10 0 -10 -20
f1 (ppm)

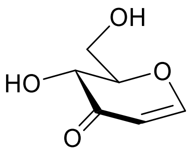

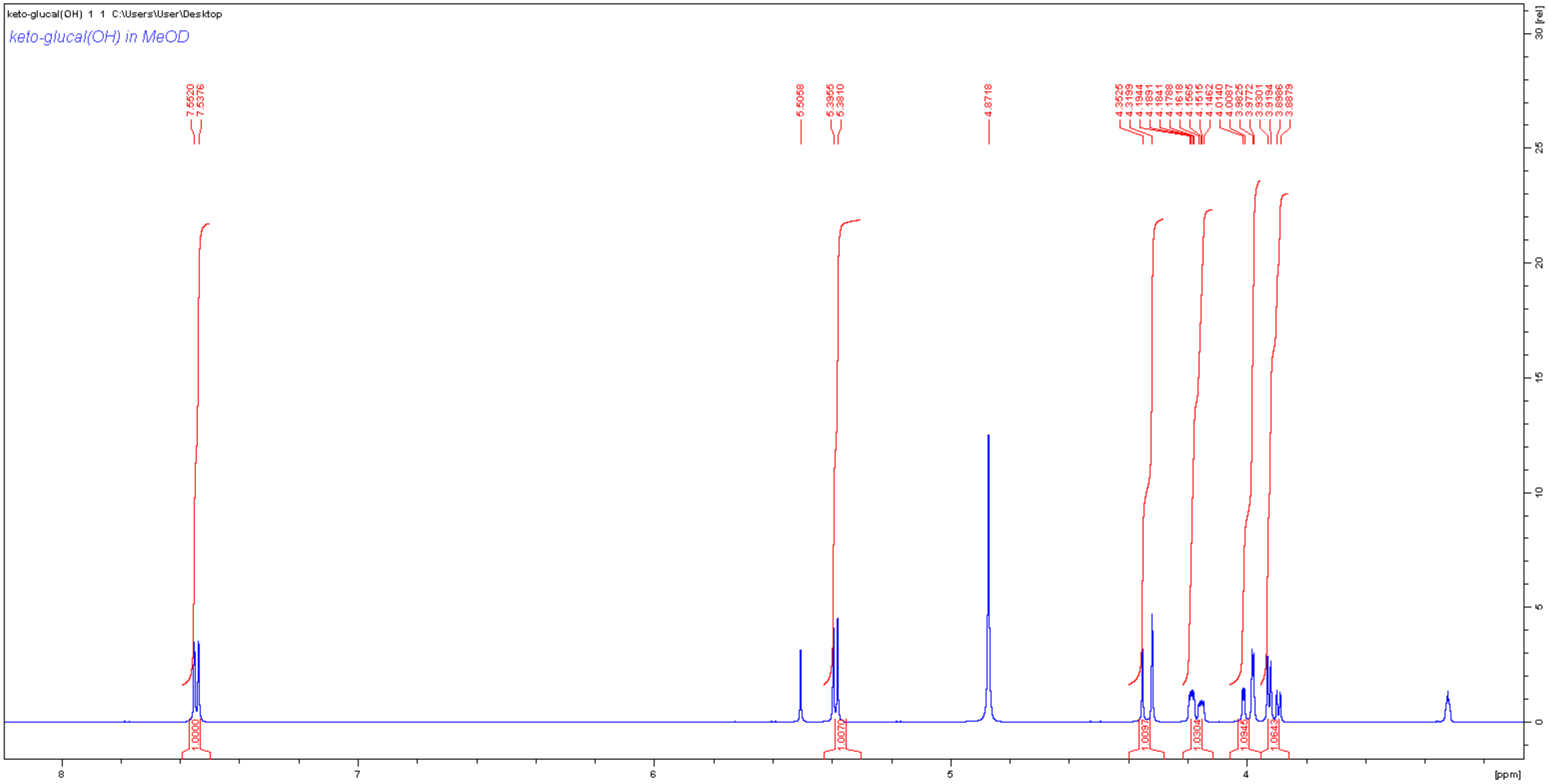
keto-glucal(OH) 1 1 C:\Users\User\Desktop
keto-glucal(OH) in MeOD
[ppm]
[rel]

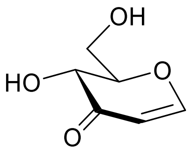
OH
HO
O
O

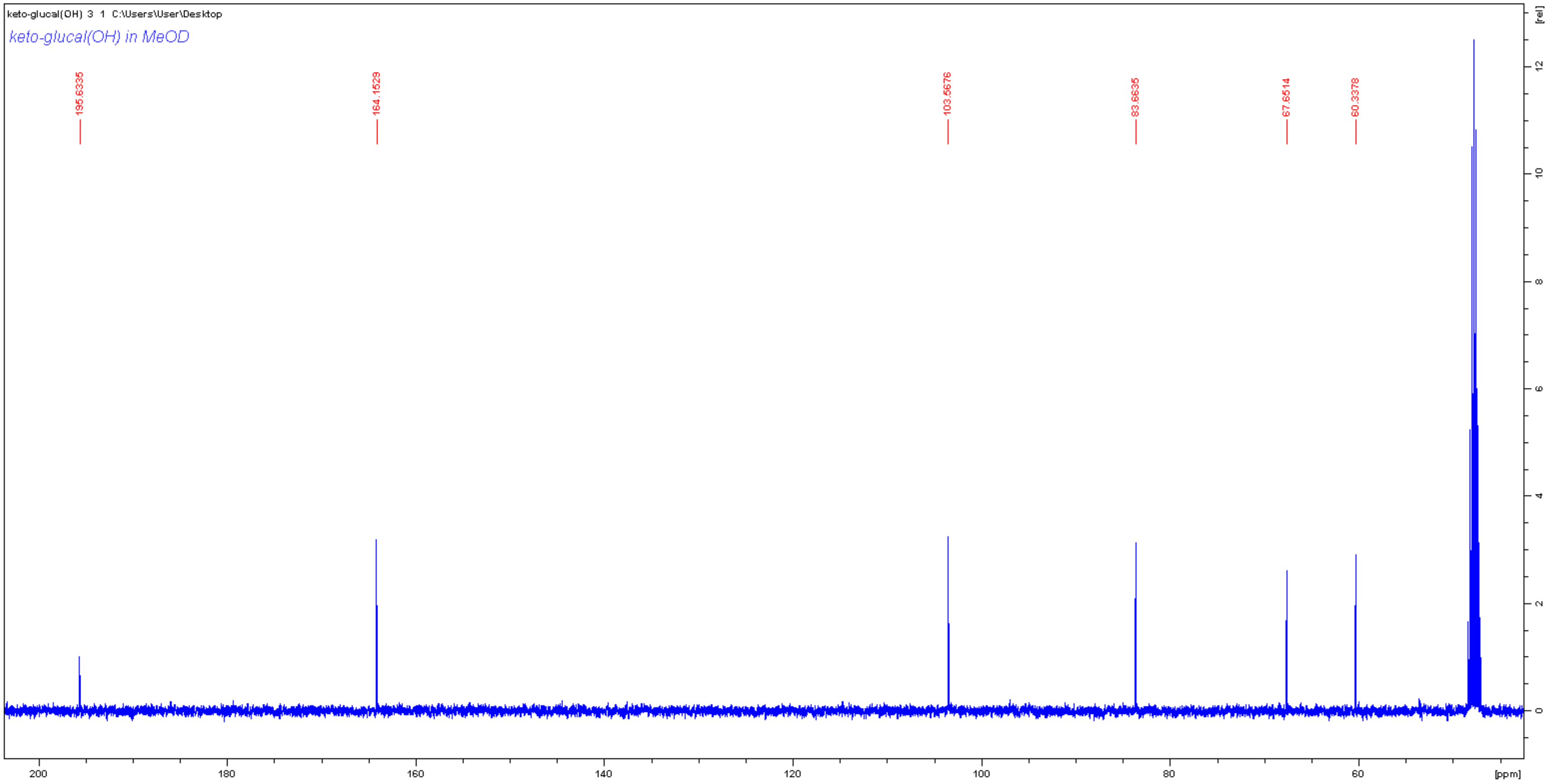
keto-glucal(OH) 3 1 C:\Users\User\Desktop
keto-glucal(OH) in MeOD
195.6335
164.1529
103.5676
83.6635
67.6514
60.3378
200
180
160
140
120
100
80
60
[ppm]
[rel]
12
10
8
6
4
2
0

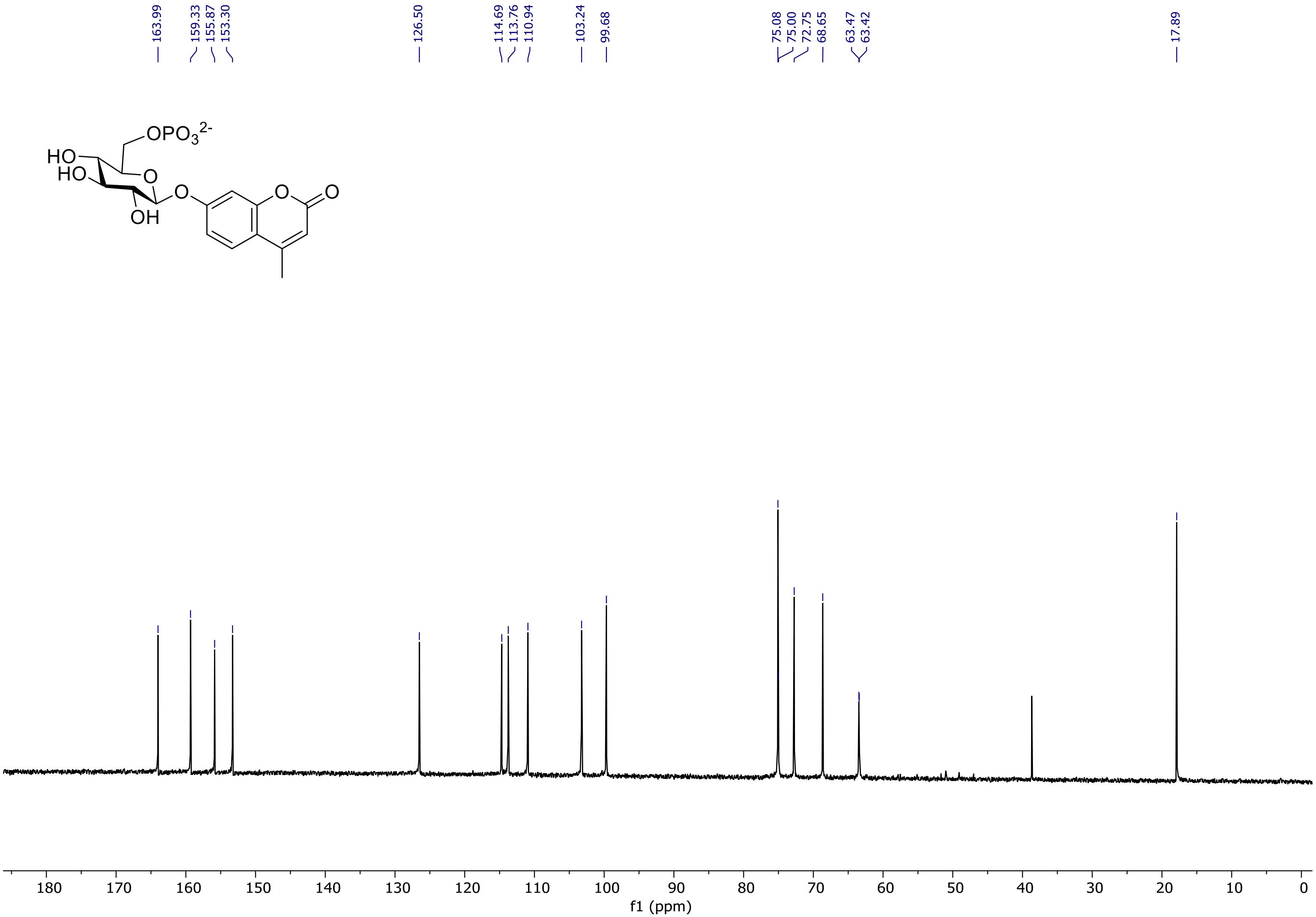
163.99
159.33
155.87
153.30
126.50
114.69
113.76
110.94
103.24
99.68
75.08
75.00
72.75
68.65
63.47
63.42
17.89
OPO3 2-
HO
HO
O
OH
O
O
O
180 170 160 150 140 130 120 110 100 90 80 70 60 50 40 30 20 10 0
f1 (ppm)

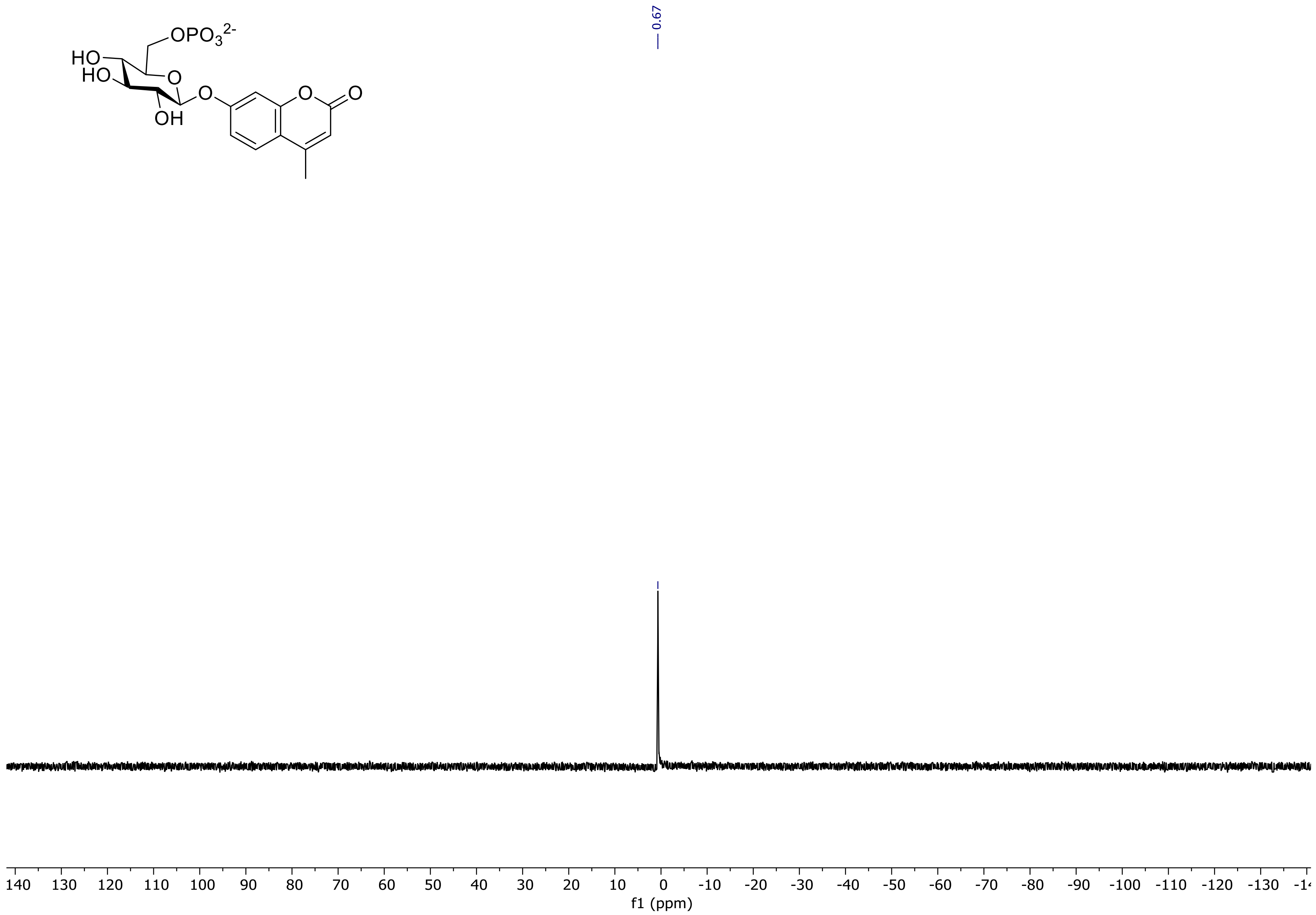

OPO3 2-
HO
HO
O
OH
O
O
O
0.67
140 130 120 110 100 90 80 70 60 50 40 30 20 10 0 -10 -20 -30 -40 -50 -60 -70 -80 -90 -100 -110 -120 -130
f1 (ppm)

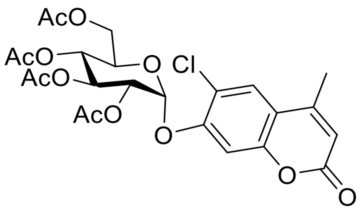
AcO
AcO
AcO
AcO
O
Cl
O
O
O

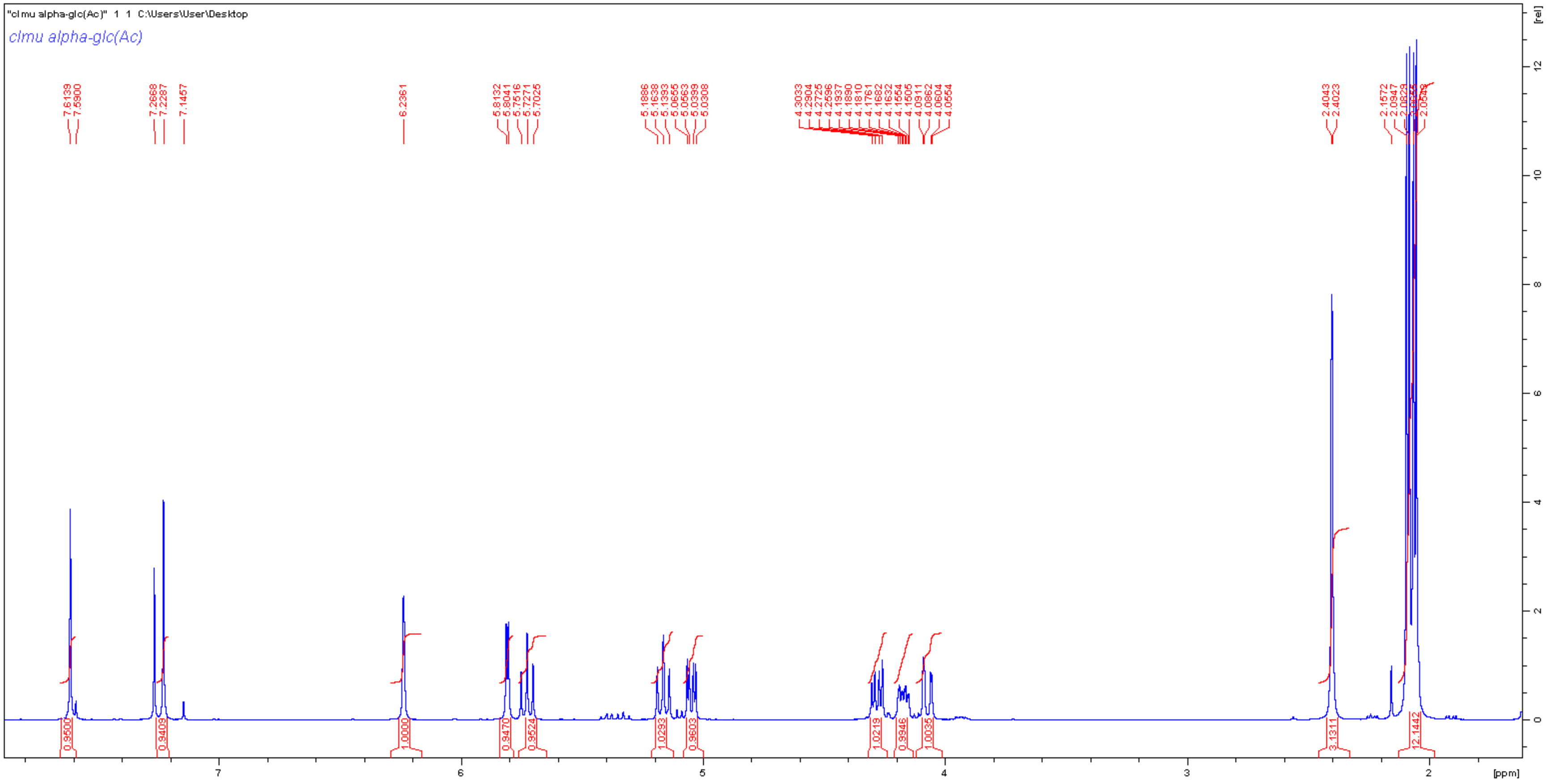
"clmu alpha-glc(Ac)" 1 1 C:\Users\User\Desktop
clmu alpha-glc(Ac)
[rel]
[ppm]

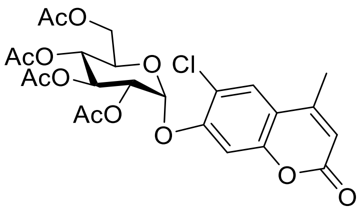

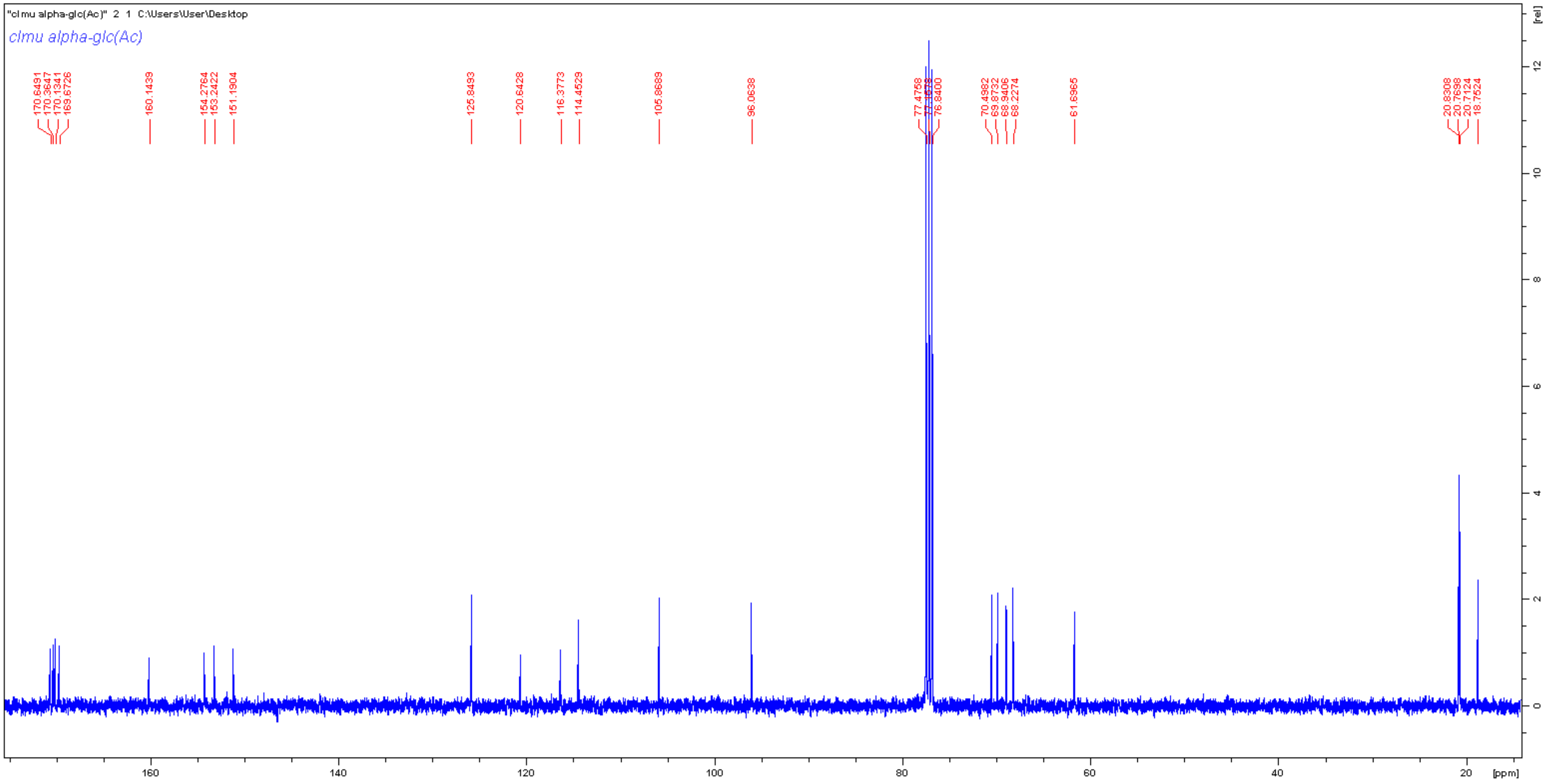

"clmu alpha-glc(Ac)" 2 1 C:\Users\User\Desktop
clmu alpha-glc(Ac)
170.6491
170.3547
170.1341
169.6726
160.1439
154.2764
153.2422
151.1904
125.8493
120.6428
116.3773
114.4529
105.8689
96.0638
77.4758
76.8400
70.4982
69.8732
68.9406
68.2274
61.6965
20.8308
20.7698
20.7124
18.7524
[ppm]
[rel]

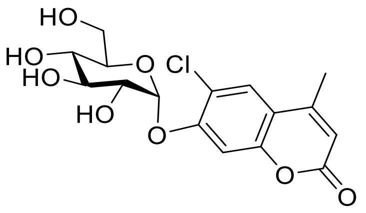

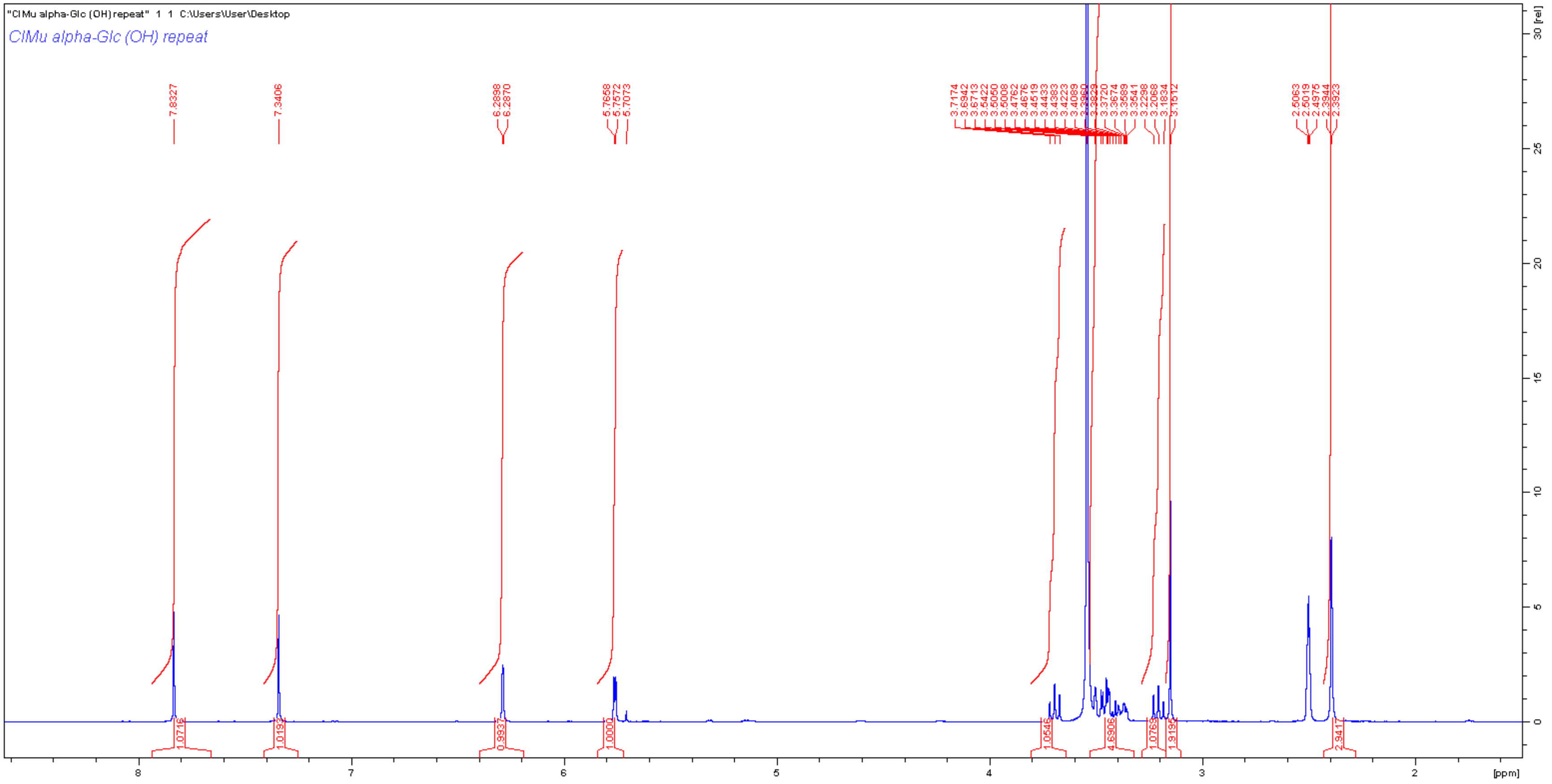

"ClMu alpha-Glc (OH) repeat" 1 1 C:\Users\User\Desktop
ClMu alpha-Glc (OH) repeat
[rel]
[ppm]

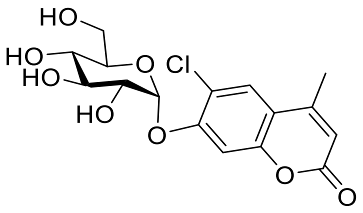

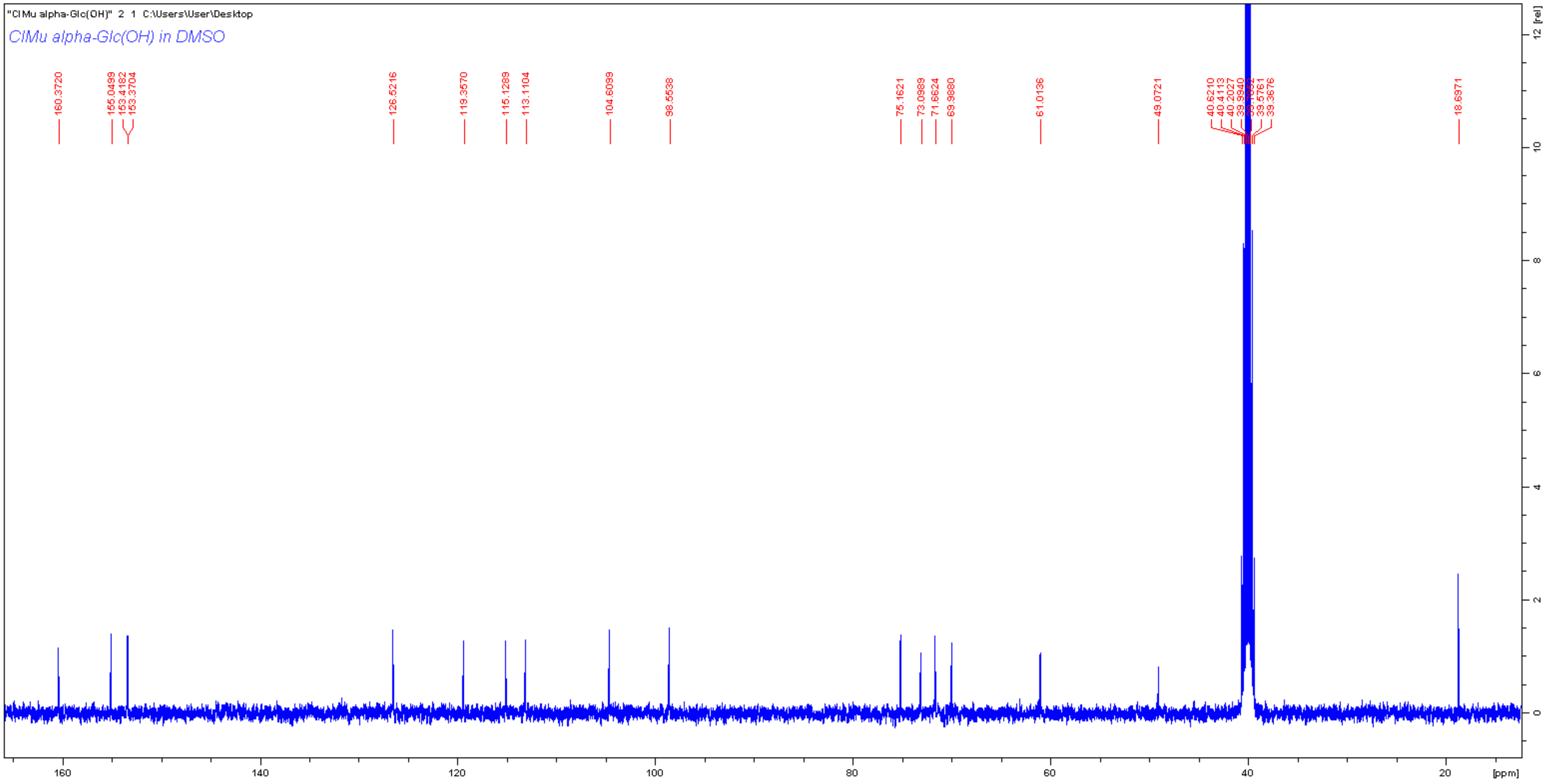

"ClMu alpha-Glc(OH)" 2 1 C:\Users\User\Desktop
ClMu alpha-Glc(OH) in DMSO
160.3720
155.0499
153.4182
153.3704
126.5216
119.3570
115.1289
113.1104
104.6099
98.5538
75.1621
73.0989
71.6624
69.9880
61.0136
49.0721
40.6210
40.4113
40.2027
39.9940
39.5761
39.3676
18.6971
160
140
120
100
80
60
40
20
[ppm]
12 [rel]
10
8
6
4
2
0

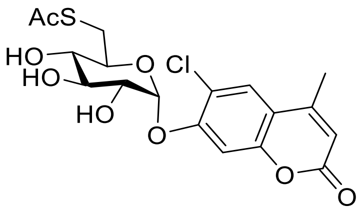
AcS
HO
HO
HO
O
Cl
O
O
O

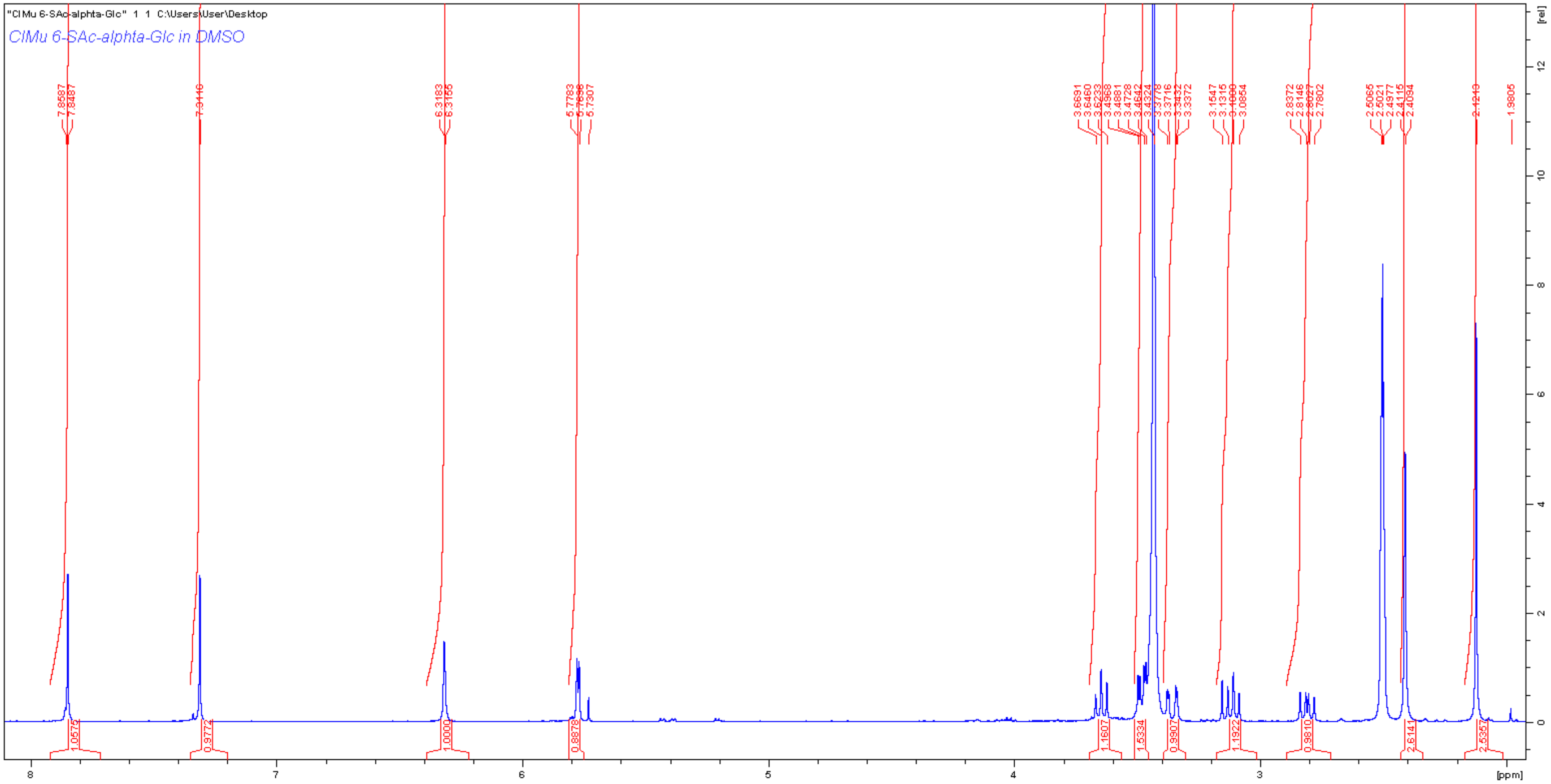
"ClMu 6-SAc-alphta-Glc" 1 1 C:\Users\User\Desktop
ClMu 6-SAc-alphta-Glc in DMSO
[ppm]
[rel]

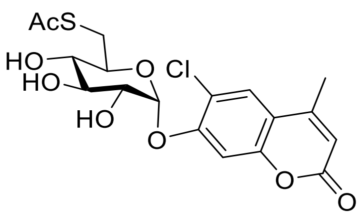
AcS
HO
HO
HO
O
Cl
O
O
O

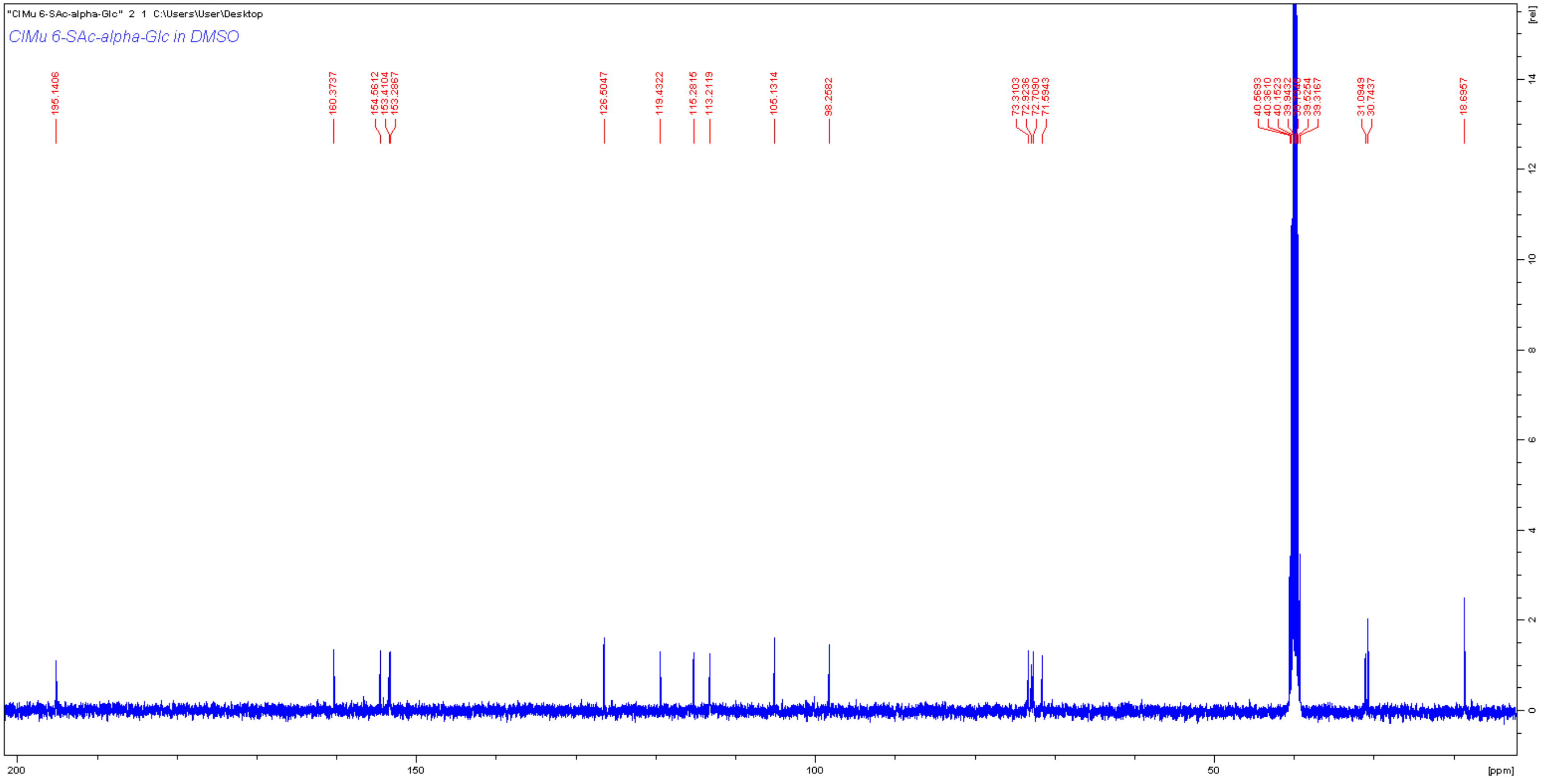
"ClMu 6-SAc-alpha-Glc" 2 1 C:\Users\User\Desktop
ClMu 6-SAc-alpha-Glc in DMSO
195.1406
160.3737
154.5612
153.4104
153.2867
126.5047
119.4322
115.2815
113.2119
105.1314
98.2582
73.3103
72.9236
72.7090
71.5943
40.5693
40.3610
40.1523
39.9432
39.7346
39.5254
39.3167
31.0949
30.7437
18.6957
[rel]
14
12
10
8
6
4
2
0
200
150
100
50
[ppm]

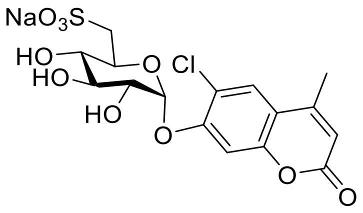

NaO3S
HO
HO
HO
O
O
Cl
O
O

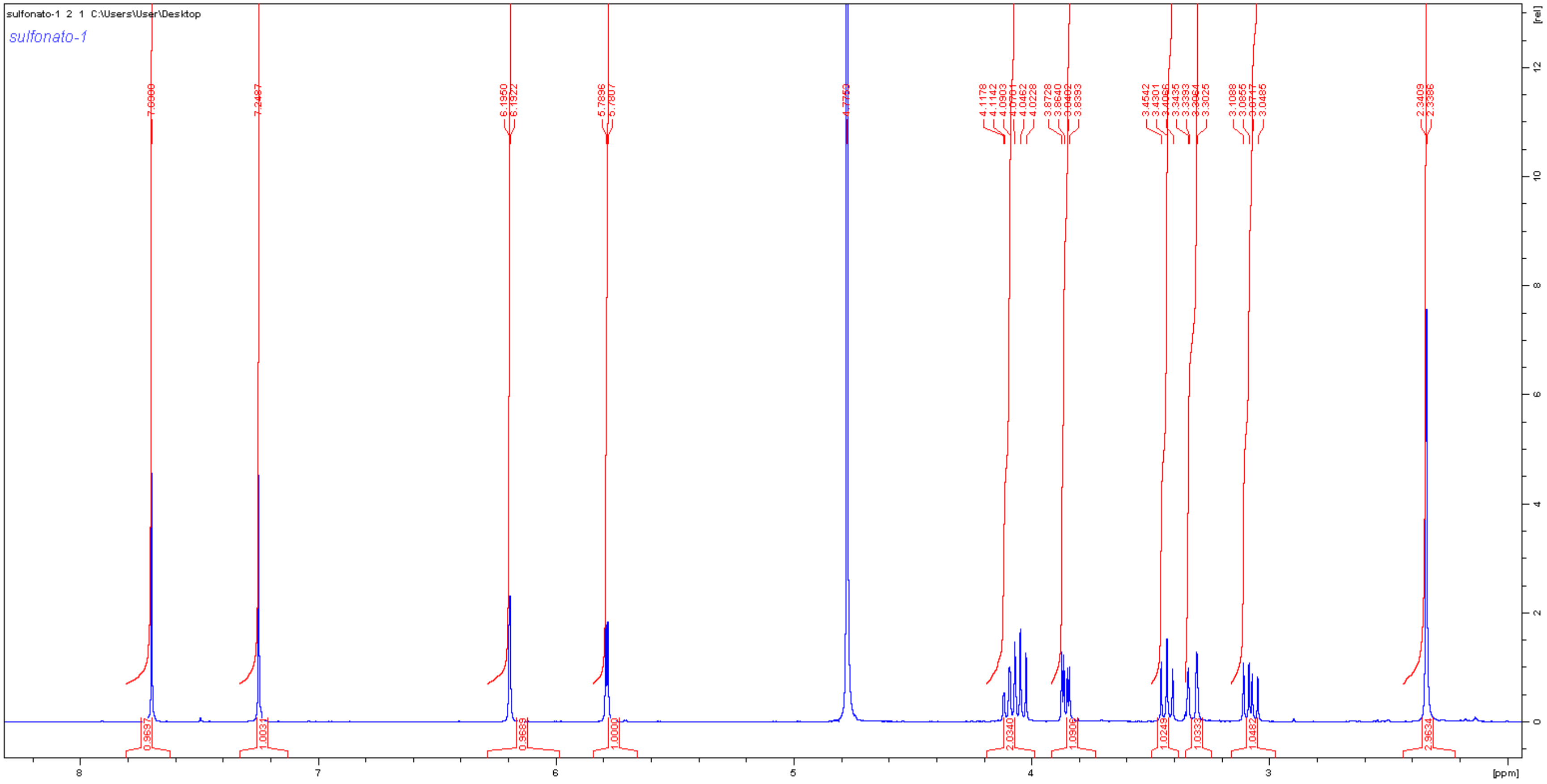

sulfonato-1 2 1 C:\Users\User\Desktop
sulfonato-1
7.2487
6.1950
6.1922
5.7896
5.7807
4.1178
4.1142
4.0903
4.0462
4.0228
3.8728
3.8640
3.8393
3.4542
3.4301
3.3435
3.3393
3.3025
3.1088
3.0855
3.0485
2.3409
2.3386
0.9697
1.0031
0.9689
1.0000
2.0340
1.0906
1.0249
1.0333
1.0482
2.9534
[rel]
[ppm]

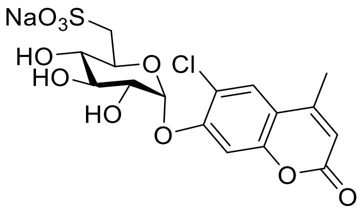
NaO3S
HO
HO
HO
O
Cl
O
O
O

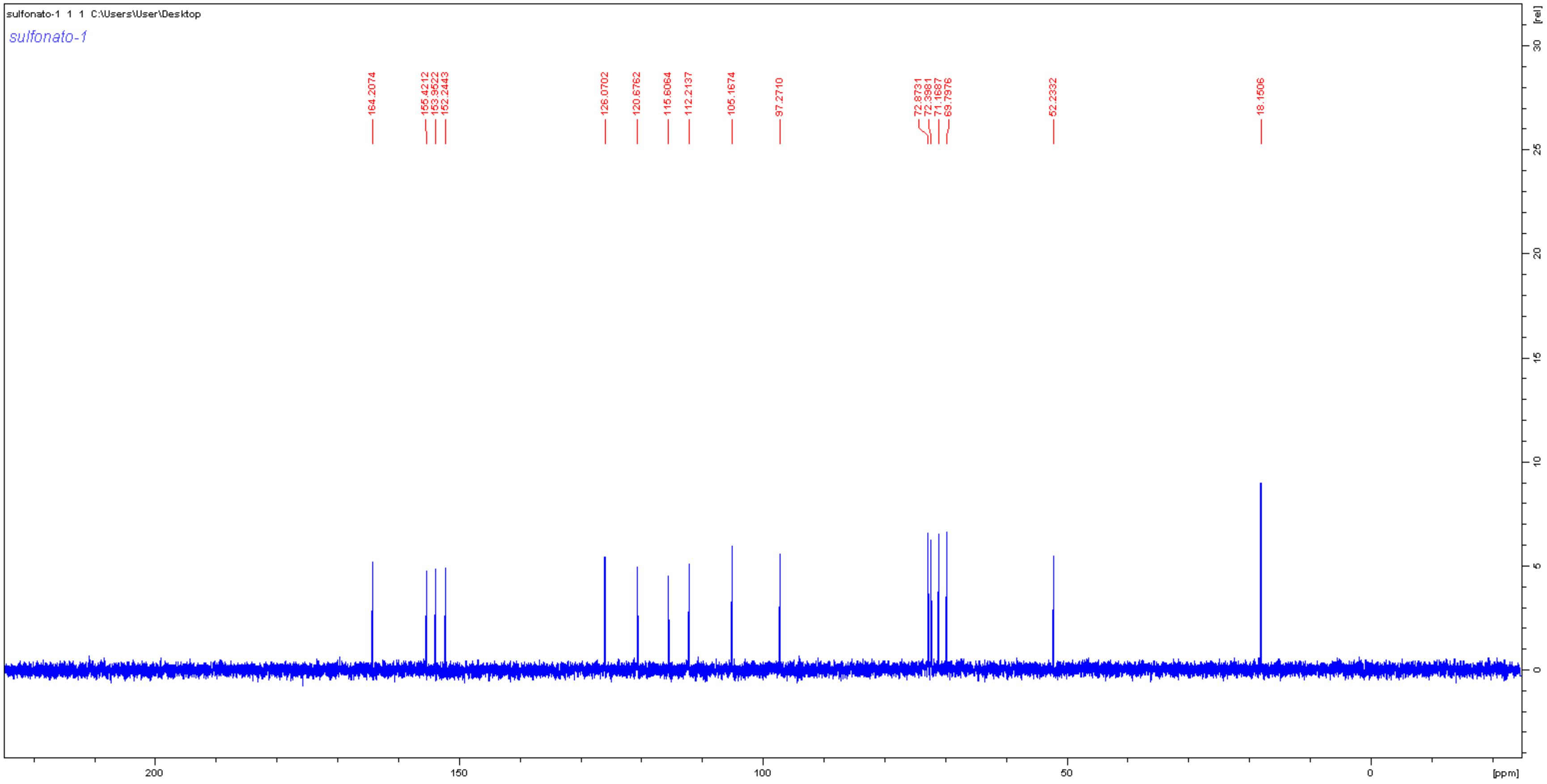
sulfonato-1 1 1 C:\Users\User\Desktop
sulfonato-1
164.2074
155.4212
153.9522
152.2443
126.0702
120.6762
115.6064
112.2137
105.1674
97.2710
72.8731
72.3981
71.1687
69.7976
52.2332
18.1506
[rel]
[ppm]

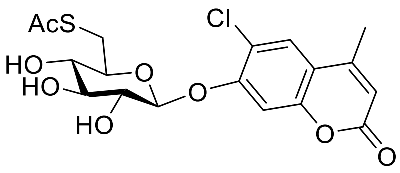
AcS
Cl
HO
HO
HO
O
O
O
O

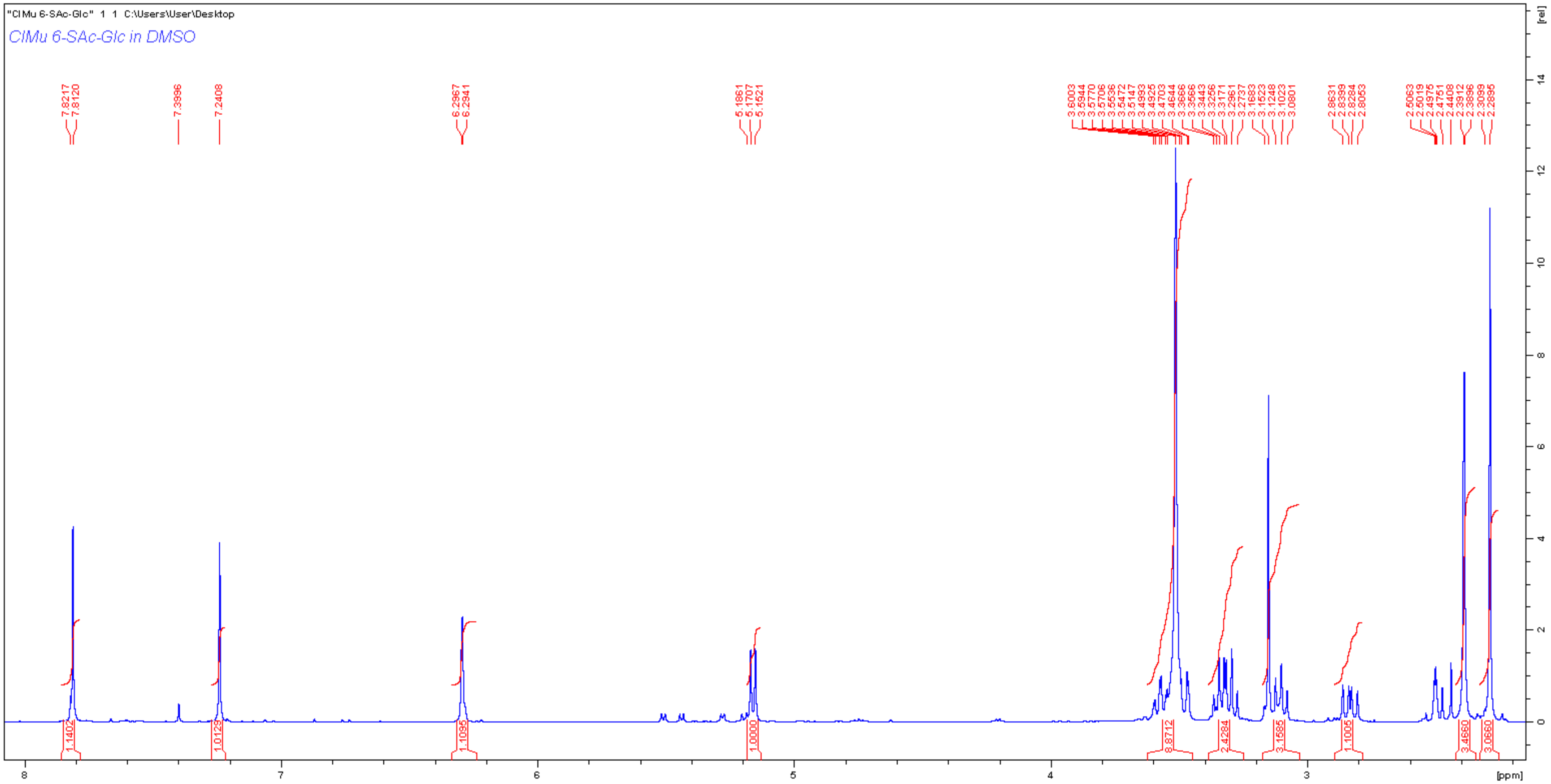
"ClMu 6-SAc-Glc" 1 1 C:\Users\User\Desktop
ClMu 6-SAc-Glc in DMSO
7.8217
7.8120
7.3996
7.2408
6.2967
6.2941
5.1861
5.1707
5.1521
3.6003
3.5944
3.5770
3.5706
3.5536
3.5472
3.5147
3.4993
3.4925
3.4703
3.4644
3.3666
3.3566
3.3443
3.3256
3.3171
3.2961
3.2737
3.1683
3.1523
3.1248
3.1023
3.0801
2.8631
2.8399
2.8284
2.8053
2.5063
2.5019
2.4976
2.4751
2.4408
2.3912
2.3896
2.3099
2.2895
1.1402
1.0129
1.1095
1.0000
8.8712
2.4284
3.1585
1.1005
3.4660
3.0660
[rel]
[ppm]

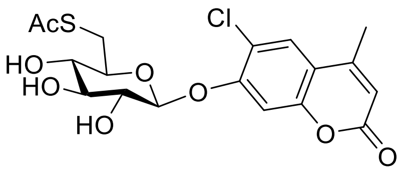
AcS
Cl
HO
HO
O
O
HO
O
O

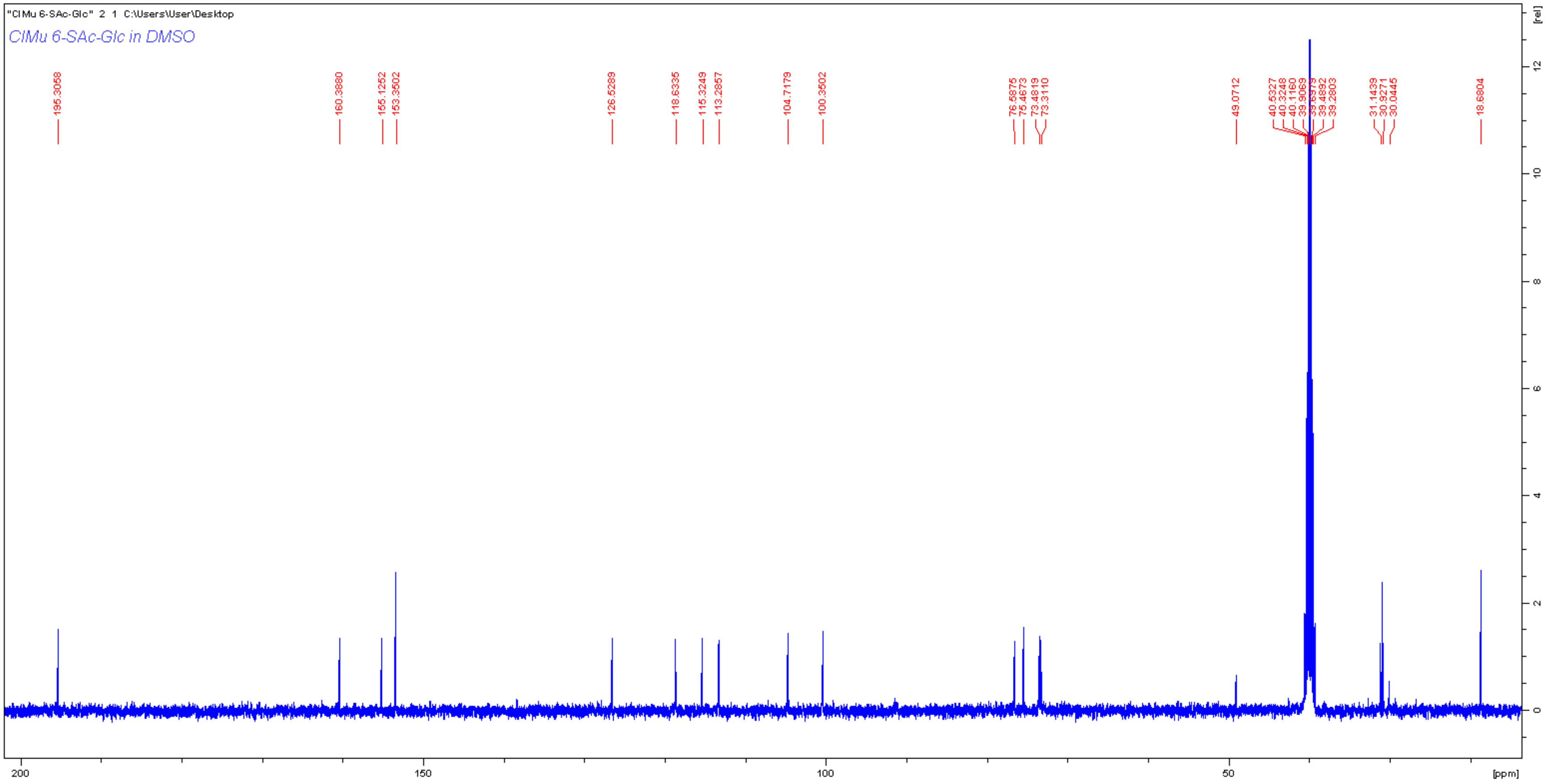
"ClMu 6-SAc-Glc" 2 1 C:\Users\User\Desktop
ClMu 6-SAc-Glc in DMSO
195.3058
160.3880
155.1252
153.3502
126.5289
118.6335
115.3249
113.2857
104.7179
100.3502
76.5875
75.4673
73.4819
73.3110
49.0712
40.5327
40.3248
40.1160
39.9069
39.6979
39.4892
39.2803
31.1439
30.9271
30.0445
18.6804
[rel]
200
150
100
50
[ppm]

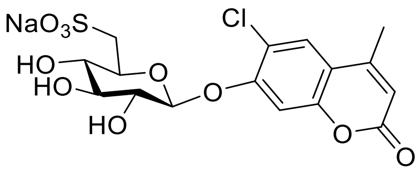

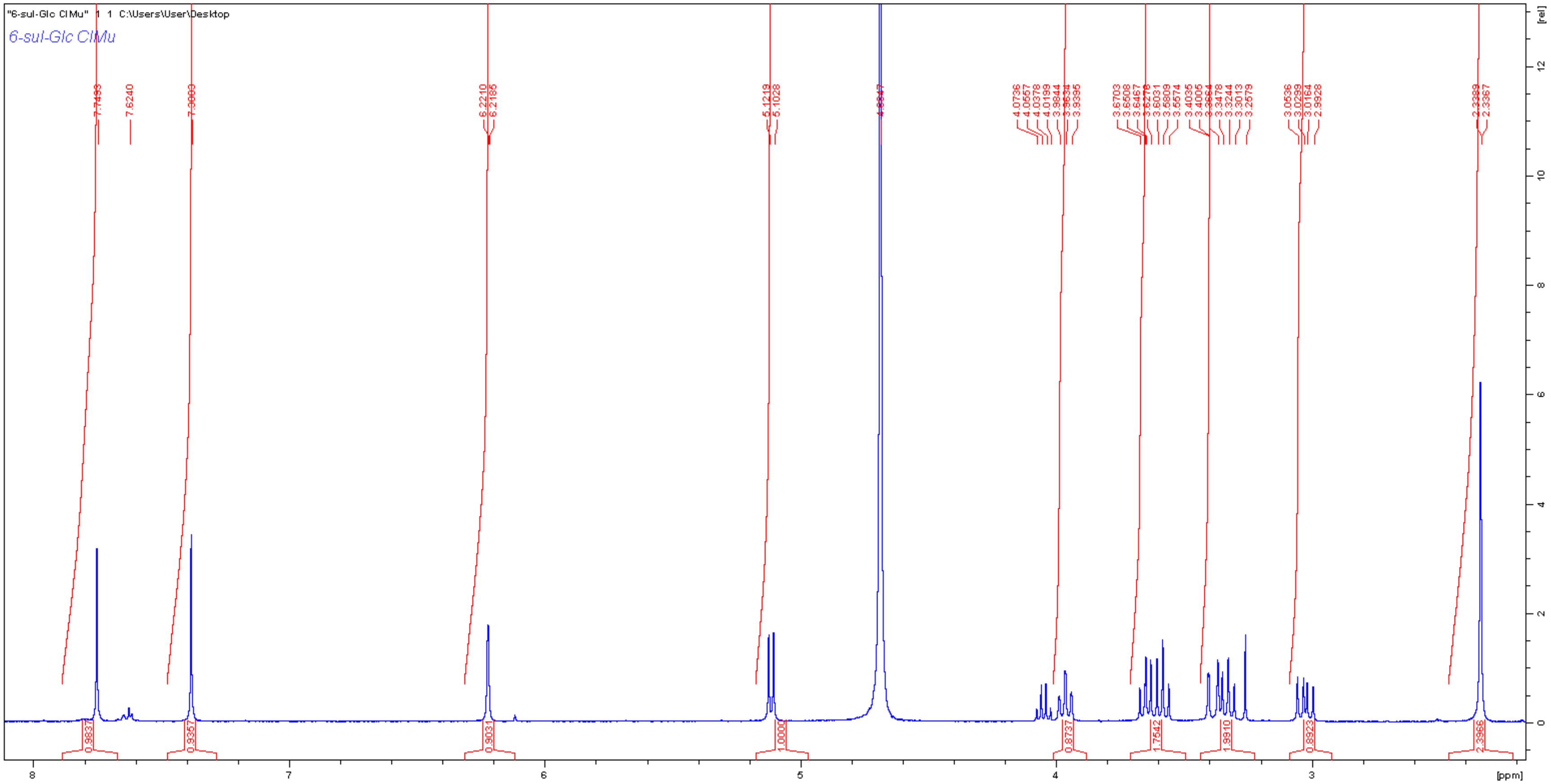
"6-sul-Glc ClMu" 1 1 C:\Users\User\Desktop
6-sul-Glc ClMu
[ppm]
[rel]

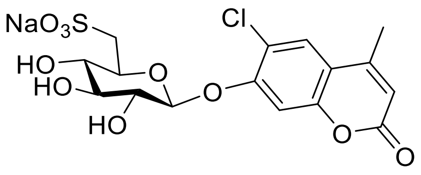

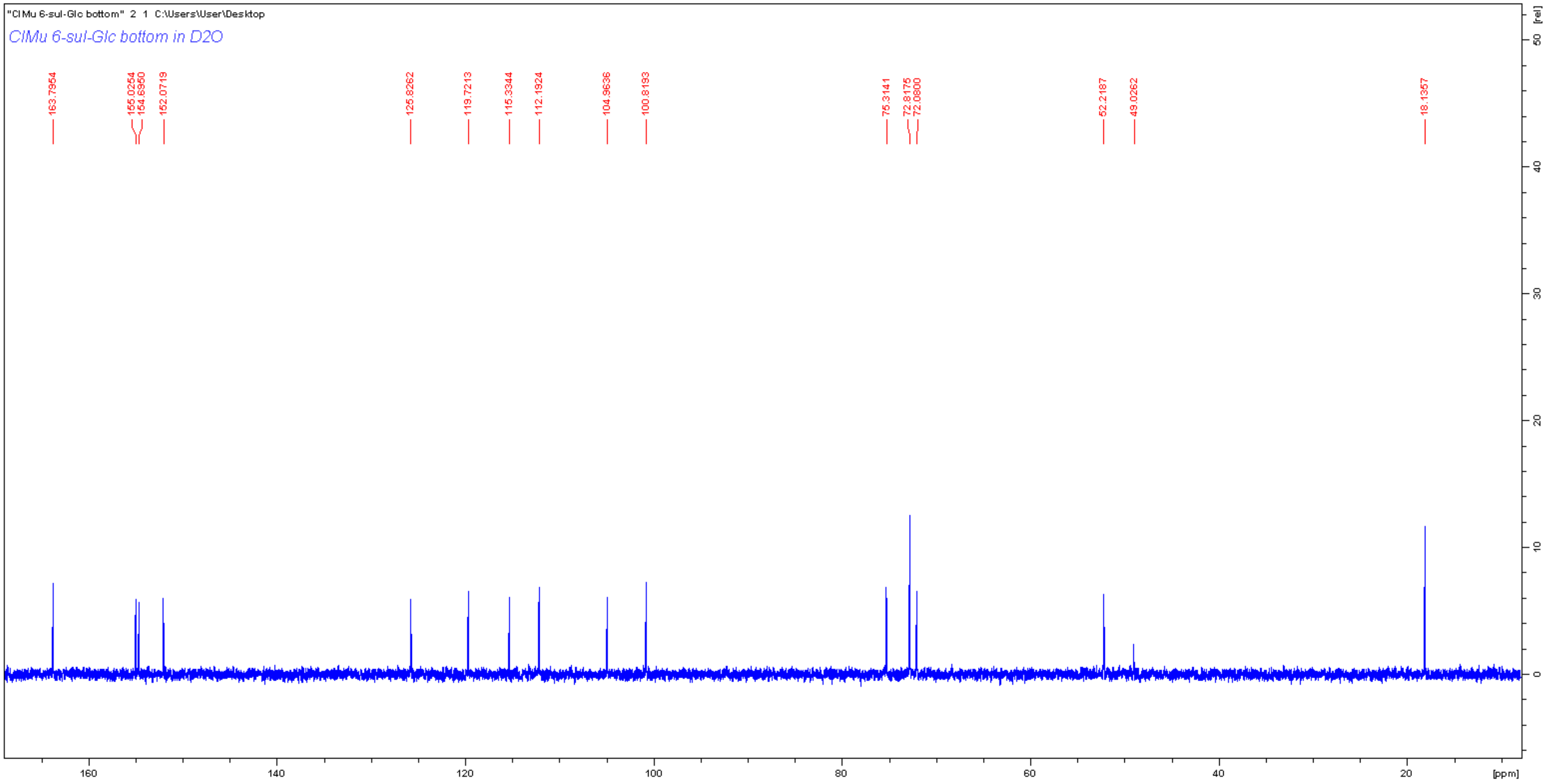
"ClMu 6-sul-Glc bottom" 2 1 C:\Users\User\Desktop
ClMu 6-sul-Glc bottom in D2O
163.7954
155.0254
154.6950
152.0719
125.8262
119.7213
115.3344
112.1924
104.9636
100.8193
75.3141
72.8175
72.0800
52.2187
49.0262
18.1357
[ppm]
[rel]

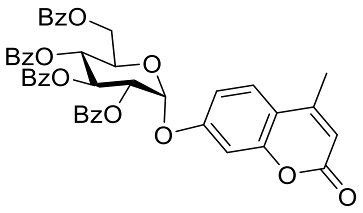
BzO
BzO
BzO
BzO
BzO
O
O
O
O
O

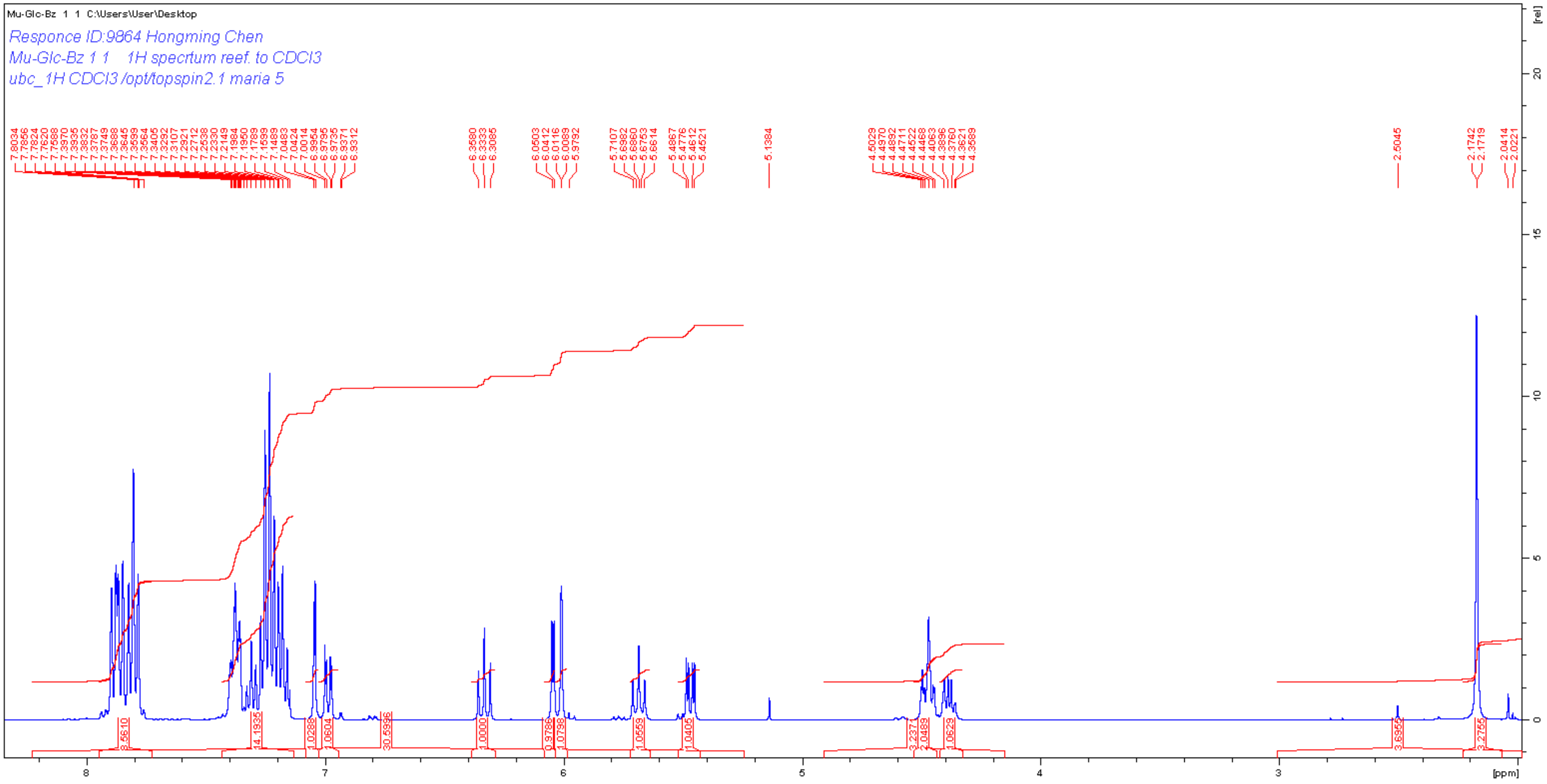
Mu-Glc-Bz 1 1 C:\Users\User\Desktop
Responce ID:9864 Hongming Chen
Mu-Glc-Bz 1 1 1H specrtum reef. to CDCl3
ubc_1H CDCl3 /opt/topspin2.1 maria 5
[ppm]
[rel]

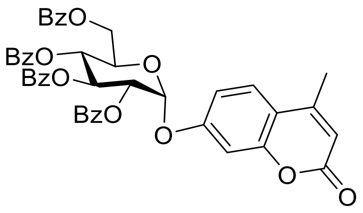
BzO
BzO
BzO
BzO
O
O
O
O
O

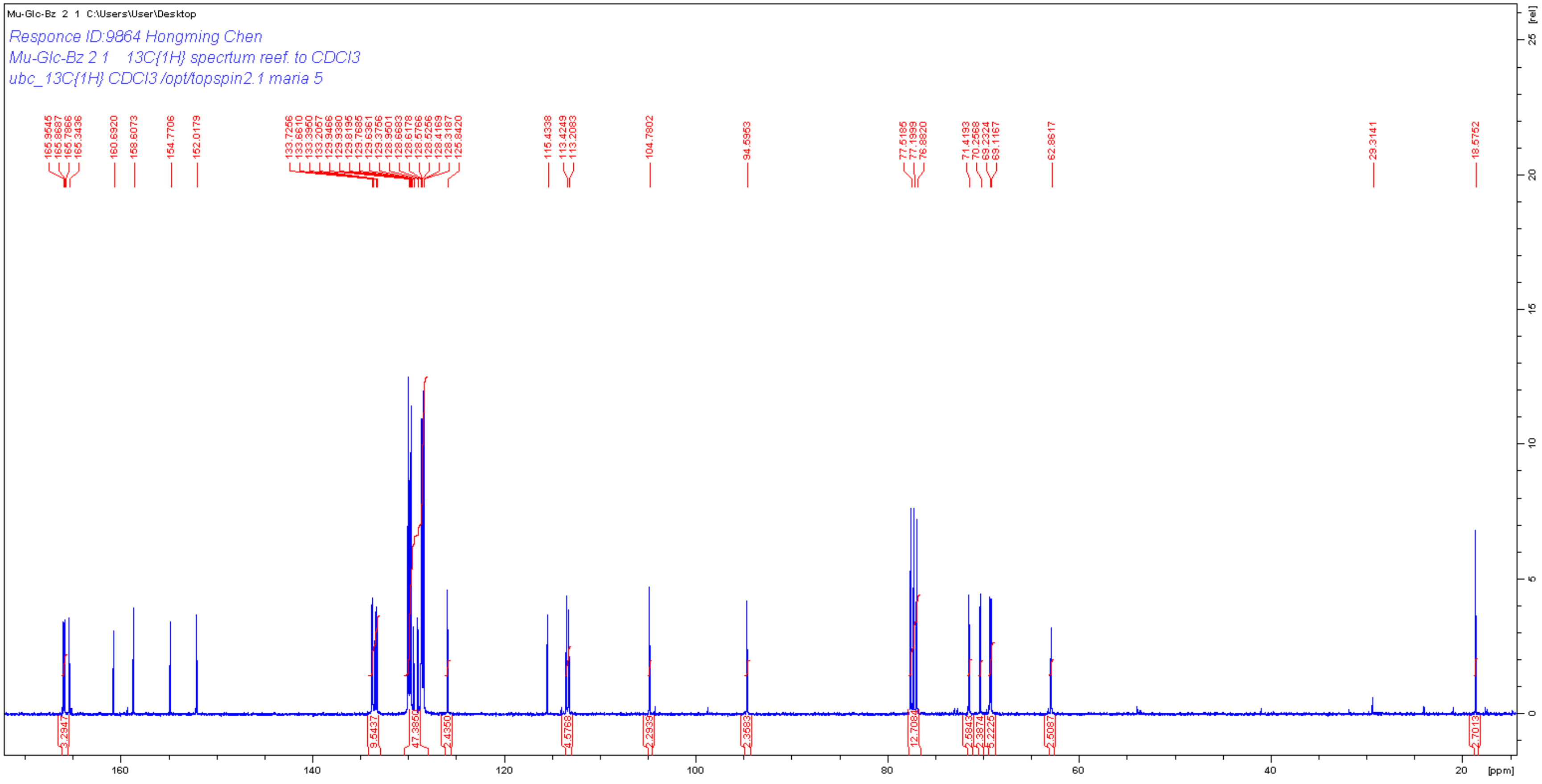
Mu-Glc-Bz 2 1 C:\Users\User\Desktop
Responce ID:9864 Hongming Chen
Mu-Glc-Bz 2 1 13C{1H} specrtum reef. to CDCl3
ubc_13C{1H} CDCl3 /opt/topspin2.1 maria 5
165.9545
165.8687
165.7866
165.3436
160.6920
158.6073
154.7706
152.0179
133.7256
133.6610
133.3950
133.2057
129.9466
129.9380
129.8195
129.7685
129.6361
129.3756
128.9601
128.6983
128.6178
128.5766
128.5256
128.4169
128.3187
125.8420
115.4338
113.4249
113.2083
104.7802
94.5953
77.5185
77.1999
76.8820
71.4193
70.2568
69.2324
69.1167
62.8617
29.3141
18.5752
3.2947
9.5437
47.3850
2.4350
4.5768
2.2939
2.3583
12.7084
2.5843
2.3874
5.2225
2.5087
2.7013
160
140
120
100
80
60
40
20
[ppm]
[rel]
0
5
10
15
20
25

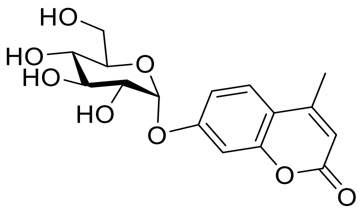

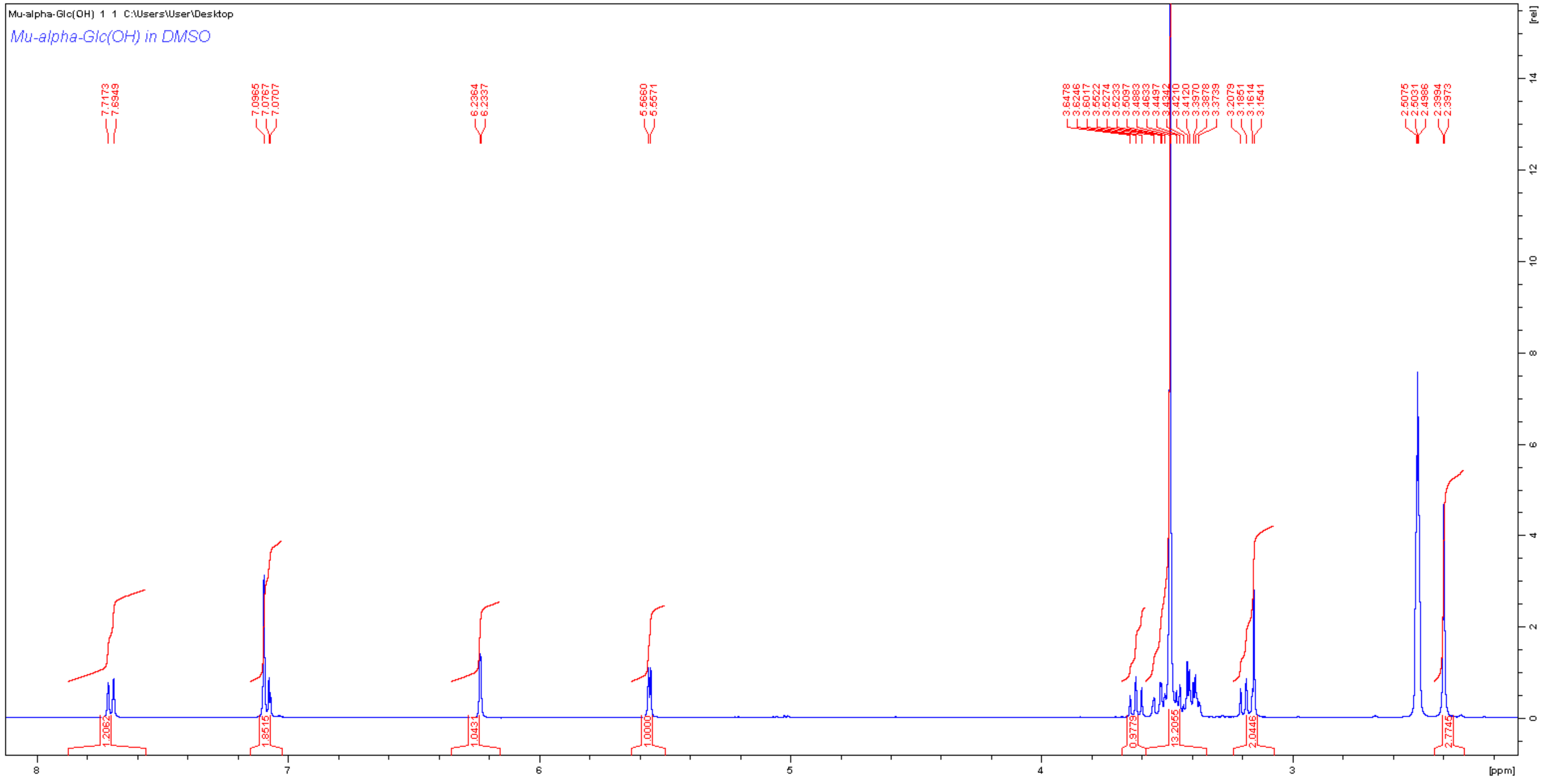
Mu-alpha-Glc(OH) 1 1 C:\Users\User\Desktop
Mu-alpha-Glc(OH) in DMSO
[rel]
[ppm]

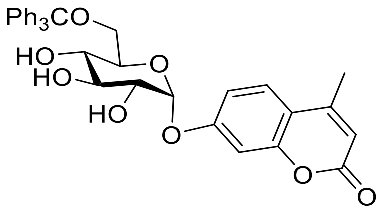

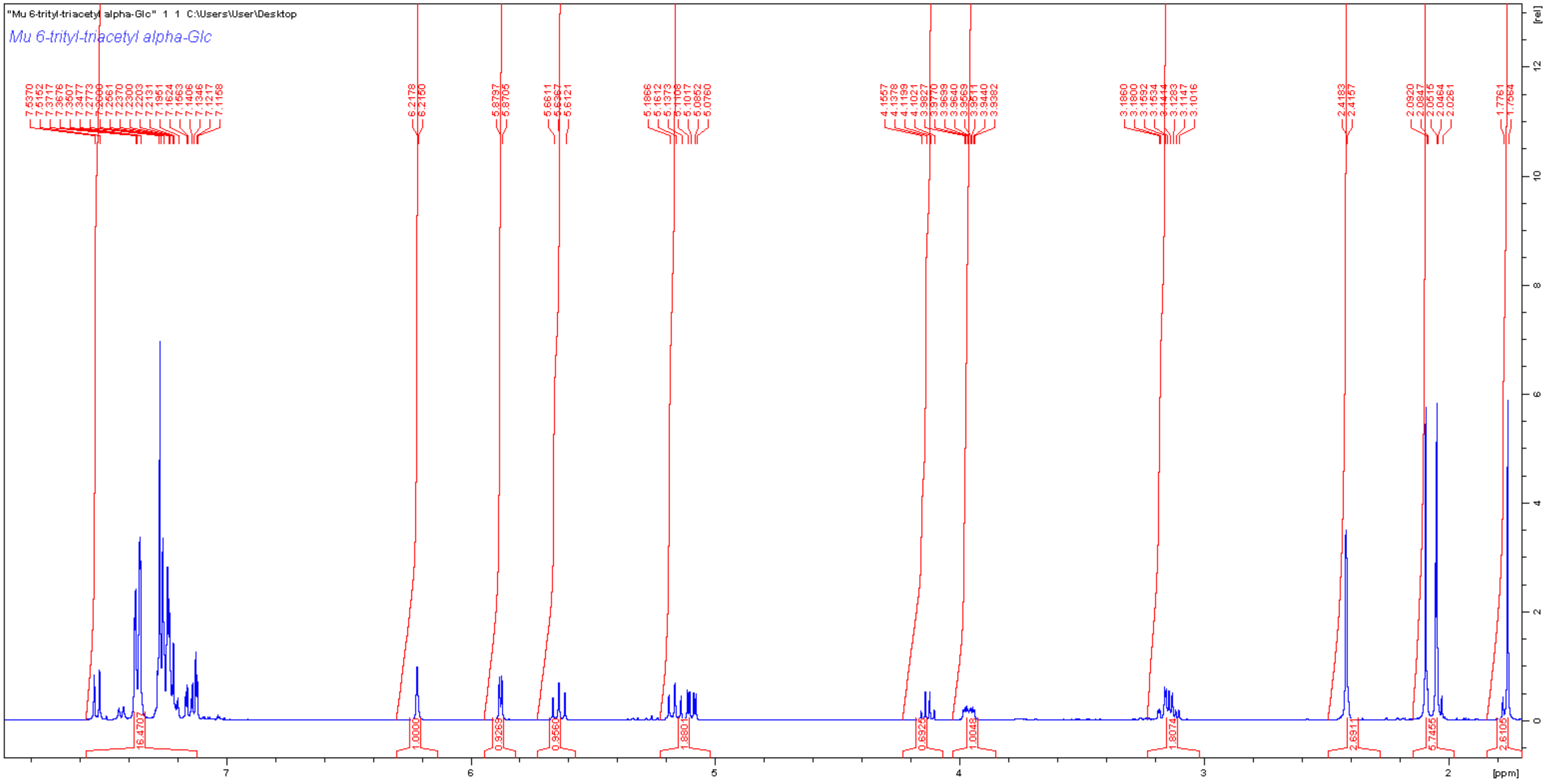
"Mu 6-trityl-triacetyl alpha-Glc" 1 1 C:\Users\User\Desktop
Mu 6-trityl-triacetyl alpha-Glc
[ppm]
[rel]

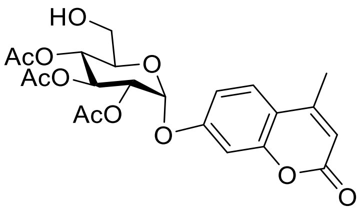
HO
AcO
AcO
AcO
O
O
O
O

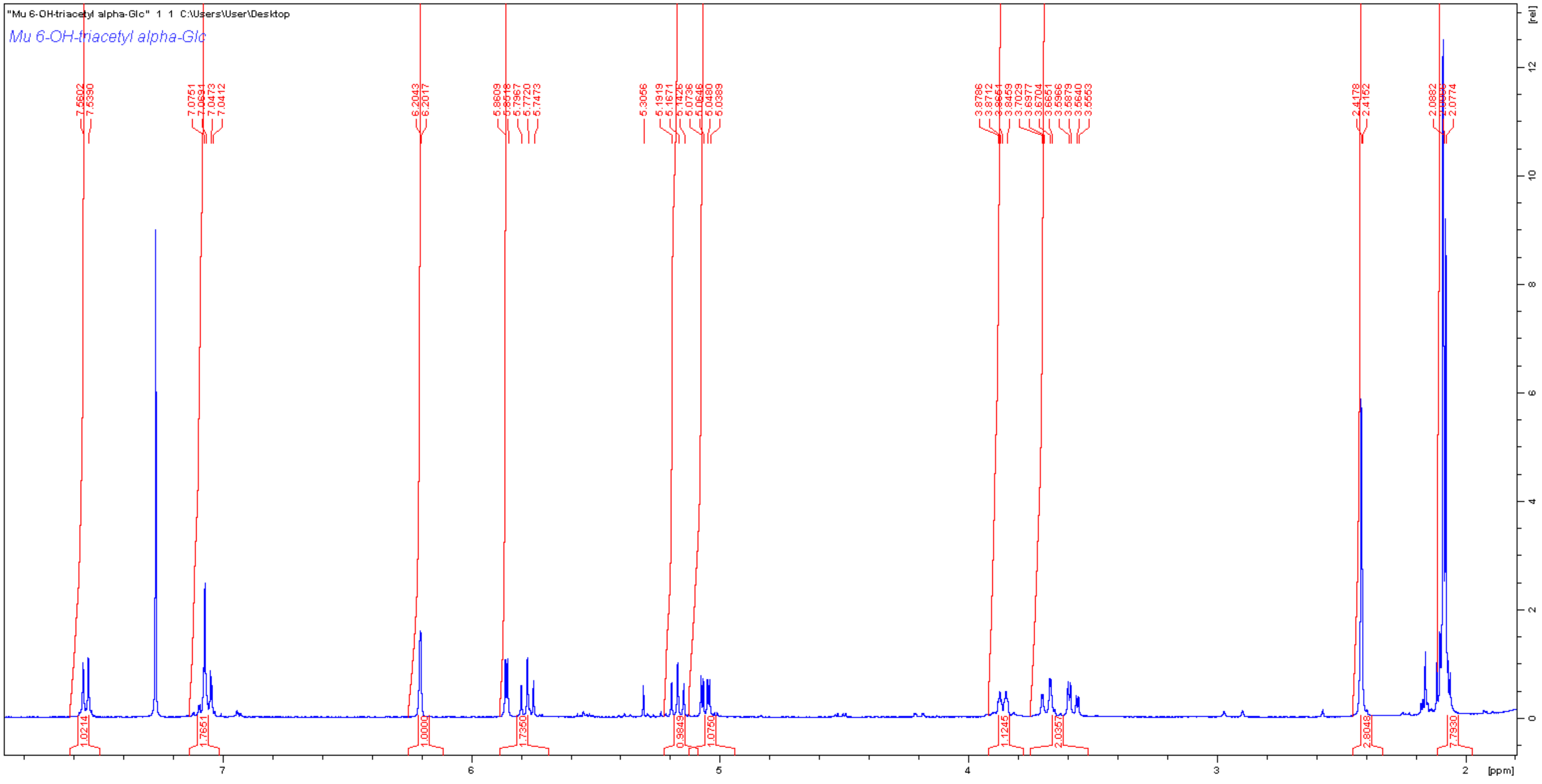
"Mu 6-OH-triacetyl alpha-Glc" 1 1 C:\Users\User\Desktop
Mu 6-OH-triacetyl alpha-Glc
[rel]
[ppm]

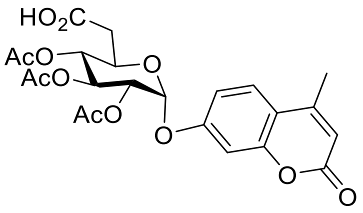

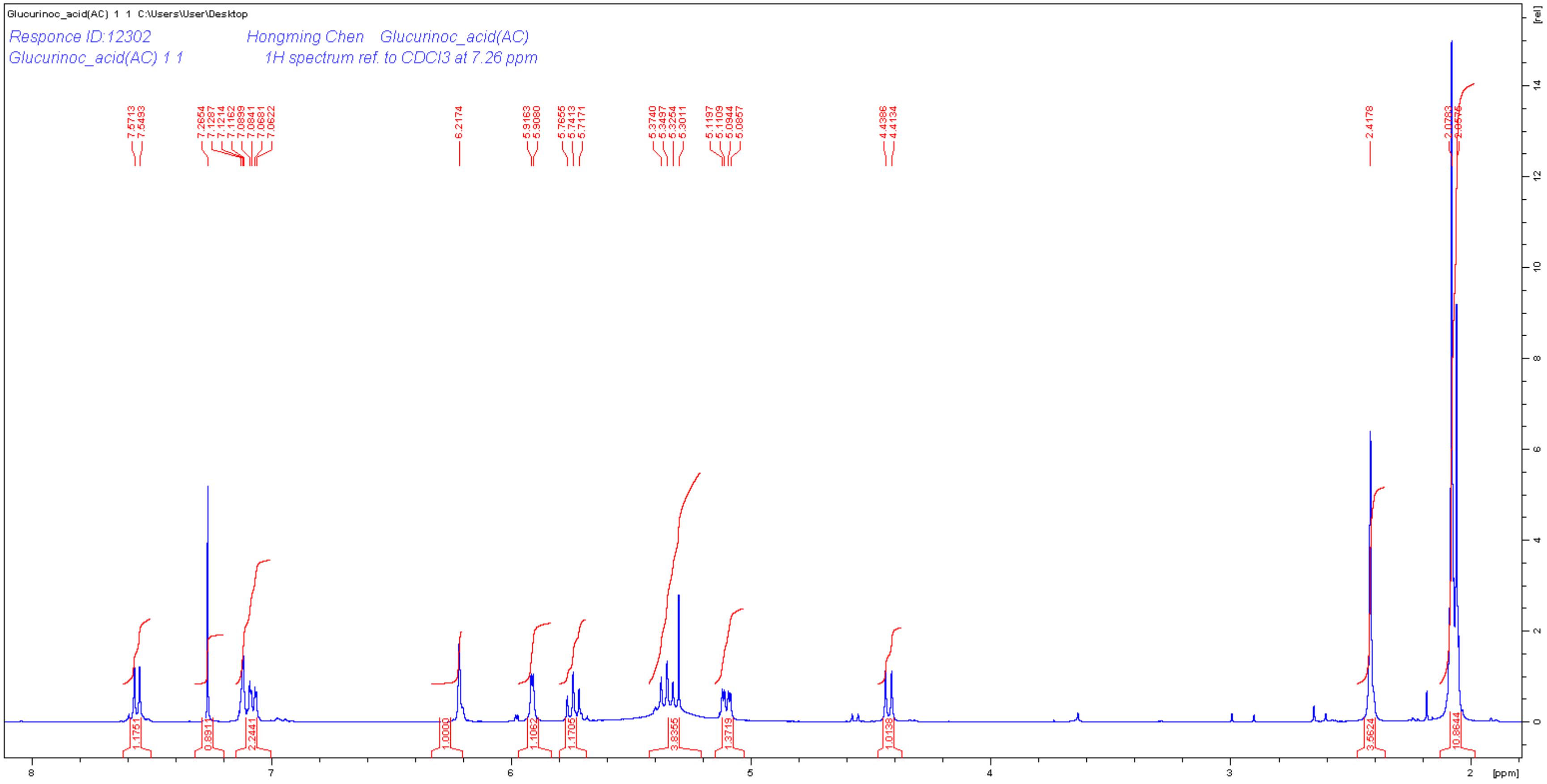

Glucurinoc_acid(AC) 1 1 C:\Users\User\Desktop
Responce ID:12302
Glucurinoc_acid(AC) 1 1
Hongming Chen Glucurinoc_acid(AC)
1H spectrum ref. to CDCl3 at 7.26 ppm
[ppm]
[rel]

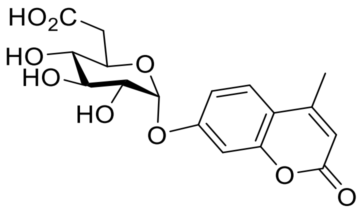

HO2C
HO
HO
HO
O
O
O
O
O

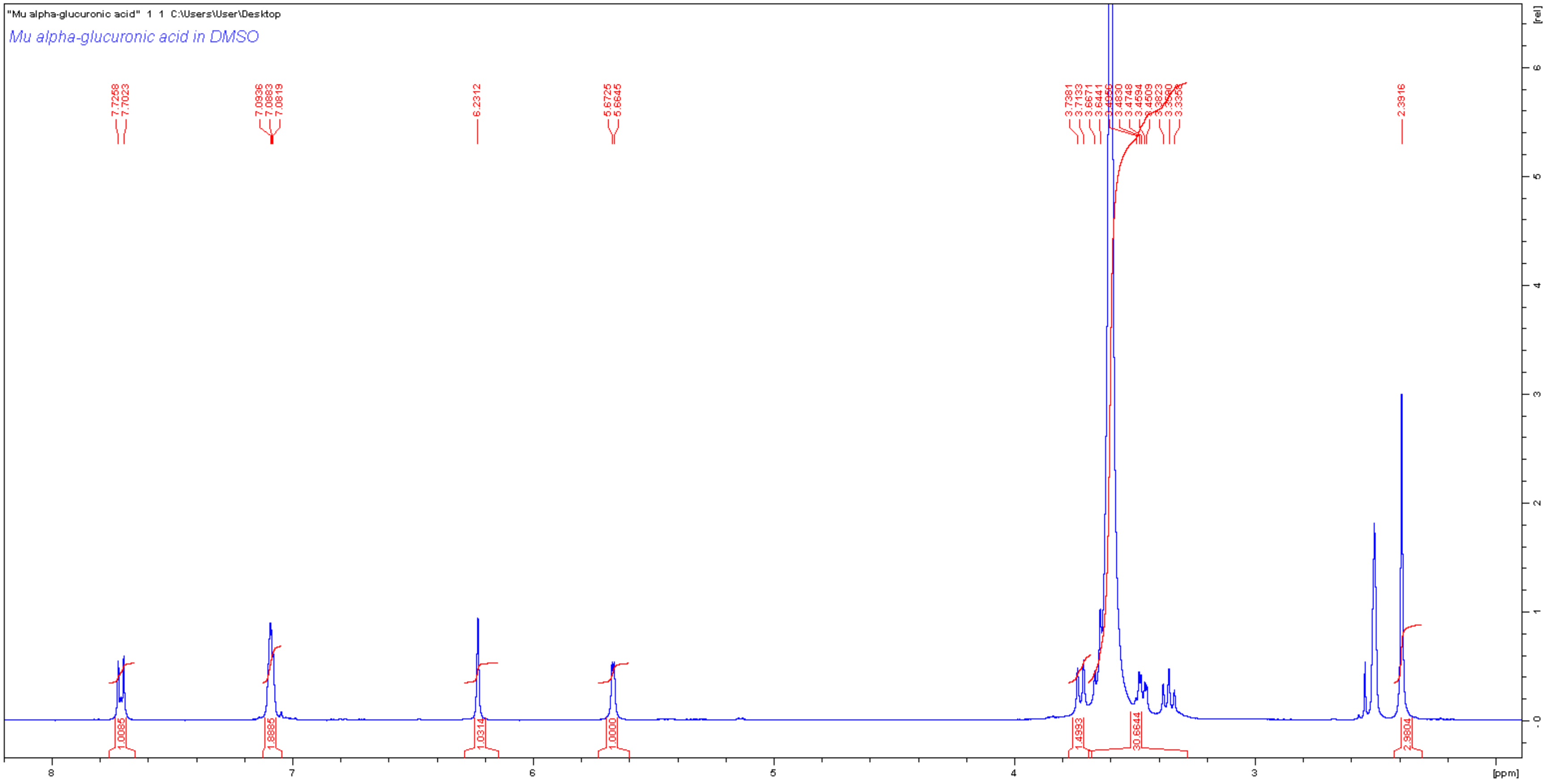

"Mu alpha-glucuronic acid" 1 1 C:\Users\User\Desktop
Mu alpha-glucuronic acid in DMSO
[ppm]
[rel]

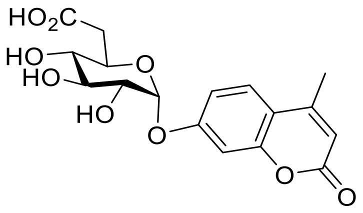

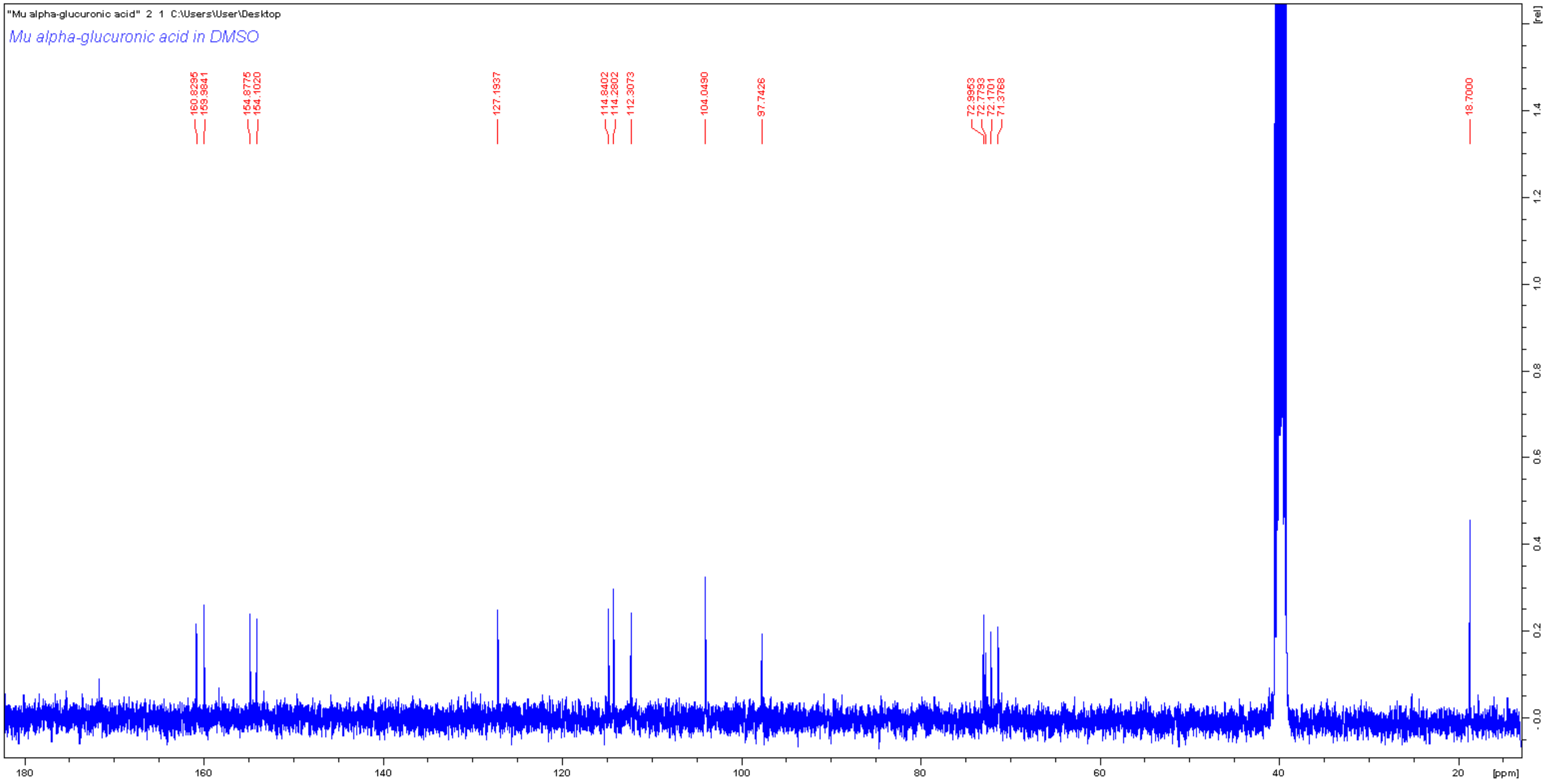

"Mu alpha-glucuronic acid" 2 1 C:\Users\User\Desktop
Mu alpha-glucuronic acid in DMSO
160.8295
159.9841
154.8775
154.1020
127.1937
114.8402
114.2802
112.3073
104.0490
97.7426
72.9953
72.7793
72.1701
71.3768
18.7000
[rel]
[ppm]

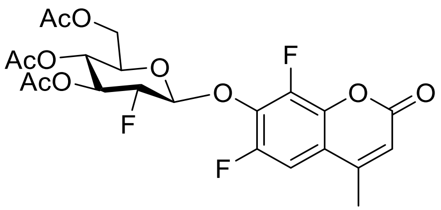
AcO
AcO
AcO
O
F
O
F
O
O
O
F

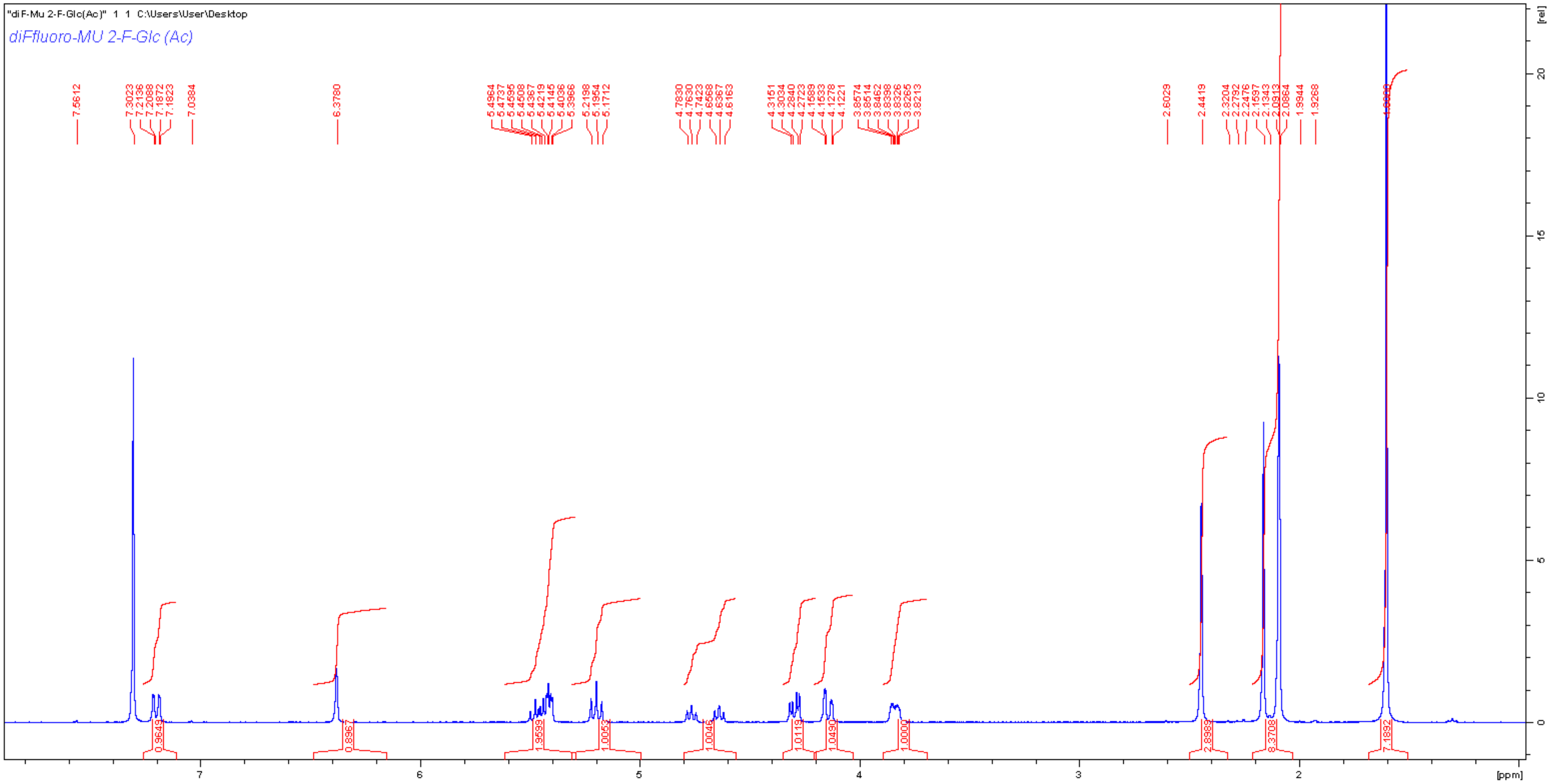
"diF-Mu 2-F-Glc(Ac)" 1 1 C:\Users\User\Desktop
diFfluoro-MU 2-F-Glc (Ac)
[ppm]
[rel]

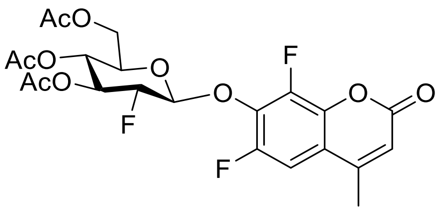
AcO
AcO
AcO
O
F
O
F
O
O
F

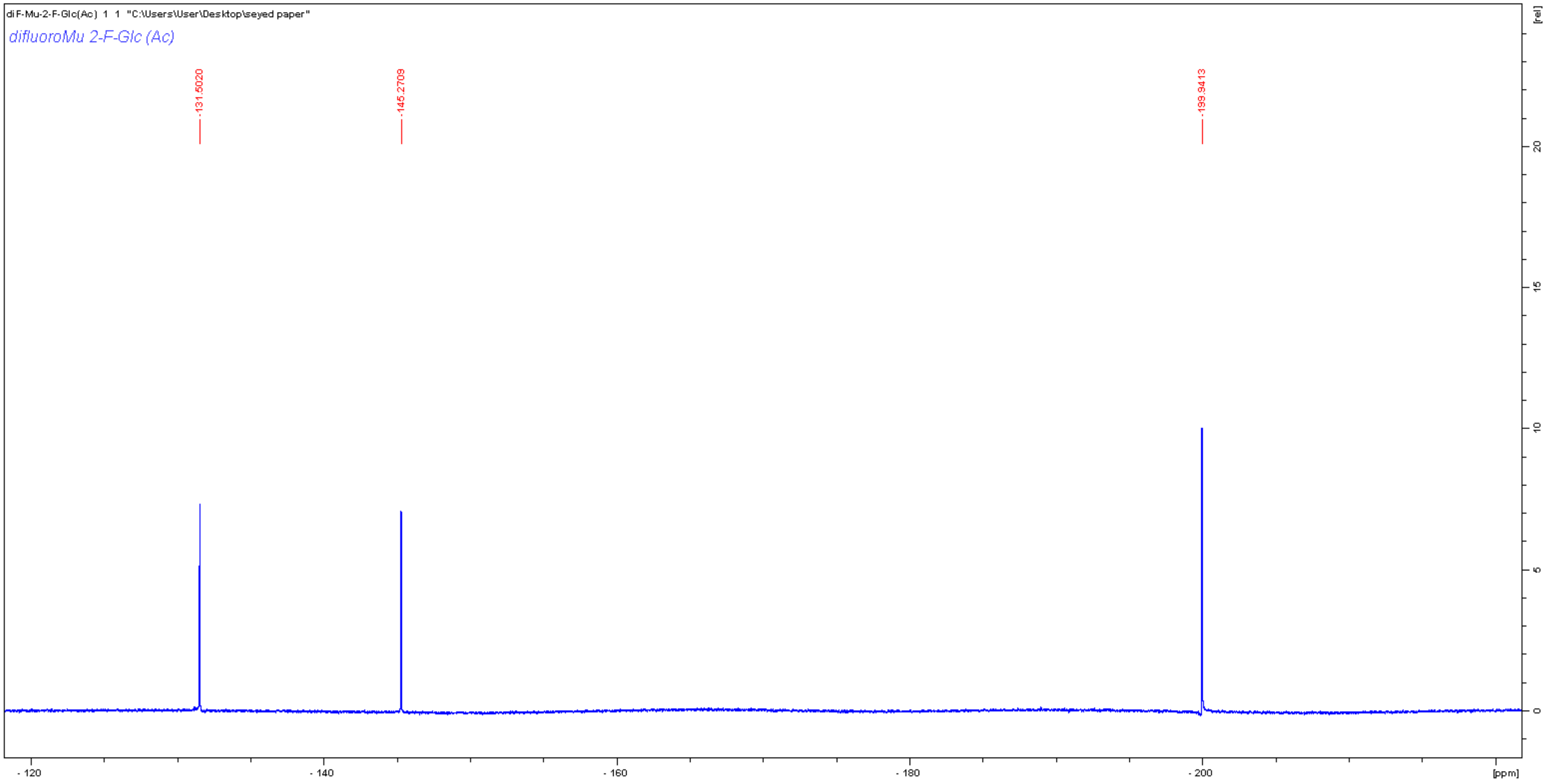
diF-Mu-2-F-Glc(Ac) 1 1 "C:\Users\User\Desktop\seyed paper"
difluoroMu 2-F-Glc (Ac)
-131.5020
-145.2709
-199.9413
[rel]
20
15
10
5
0
-120
-140
-160
-180
-200
[ppm]

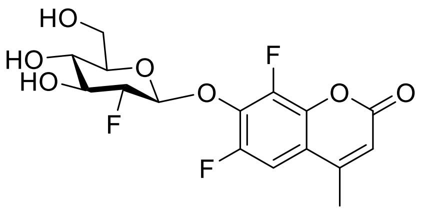
HO
HO
HO
O
F
O
F
O
O
F

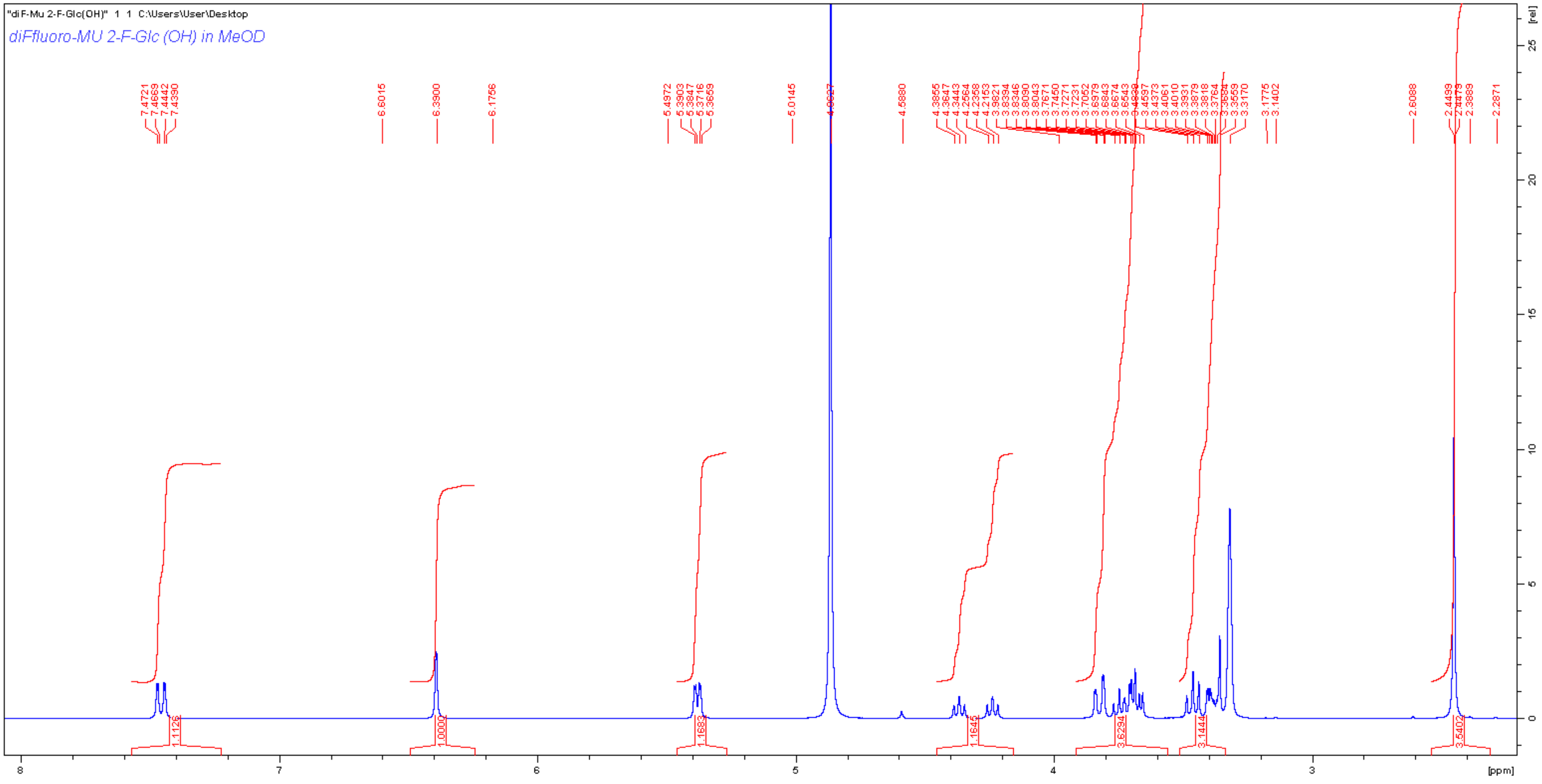
"diF-Mu 2-F-Glc(OH)" 1 1 C:\Users\User\Desktop
diFluoro-MU 2-F-Glc (OH) in MeOD
[ppm]
[rel]

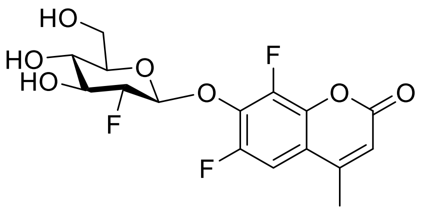
HO
HO
HO
O
F
O
F
O
O
F

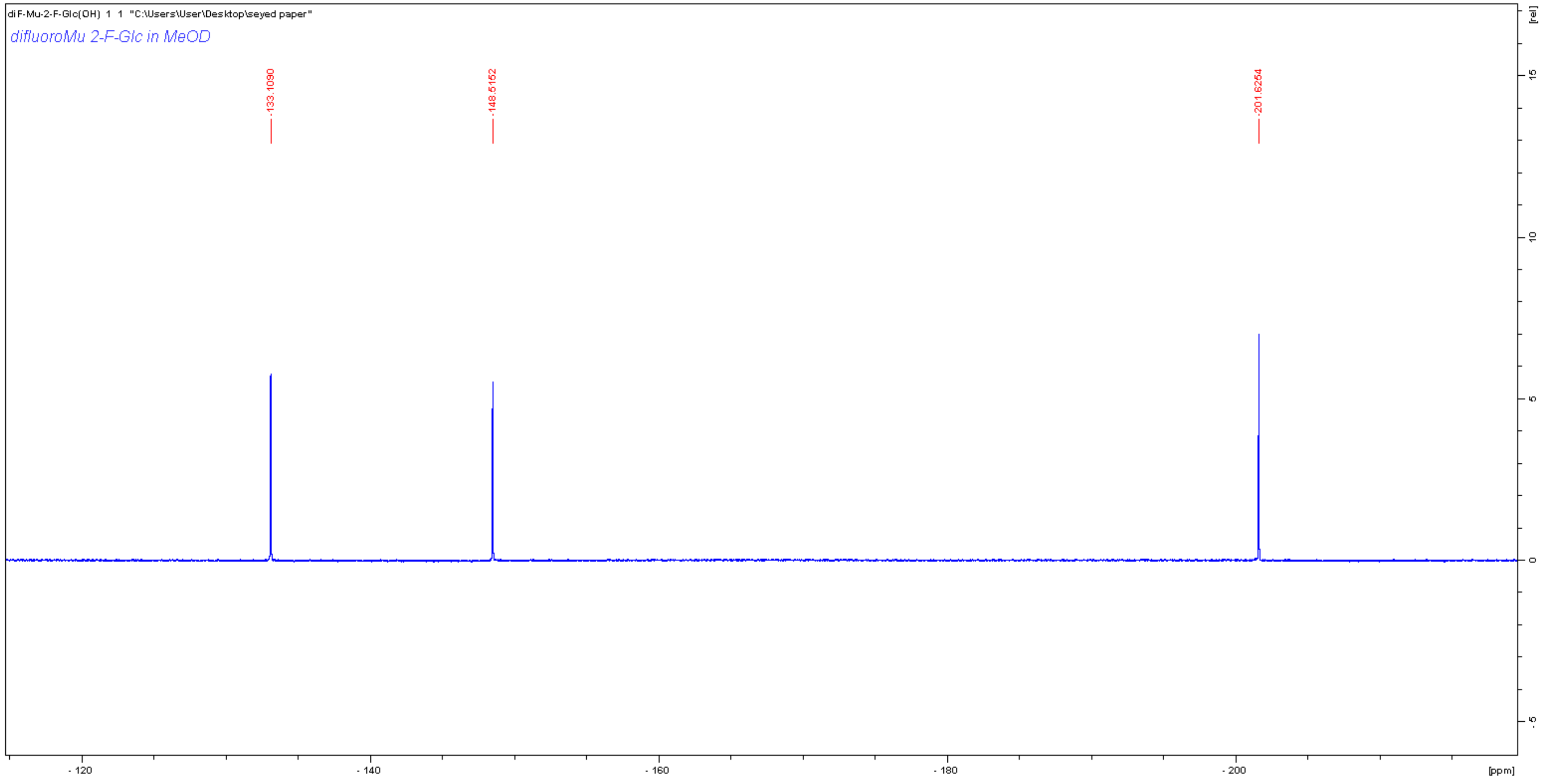
diF-Mu-2-F-Glc(OH) 1 1 "C:\Users\User\Desktop\seyed paper"
difluoroMu 2-F-Glc in MeOD
-133.1090
-148.5152
-201.6254
[rel]
15
10
5
0
-5
-120
-140
-160
-180
-200
[ppm]

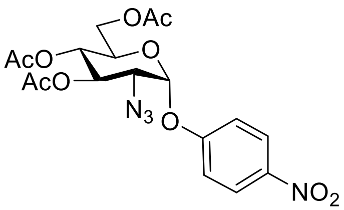

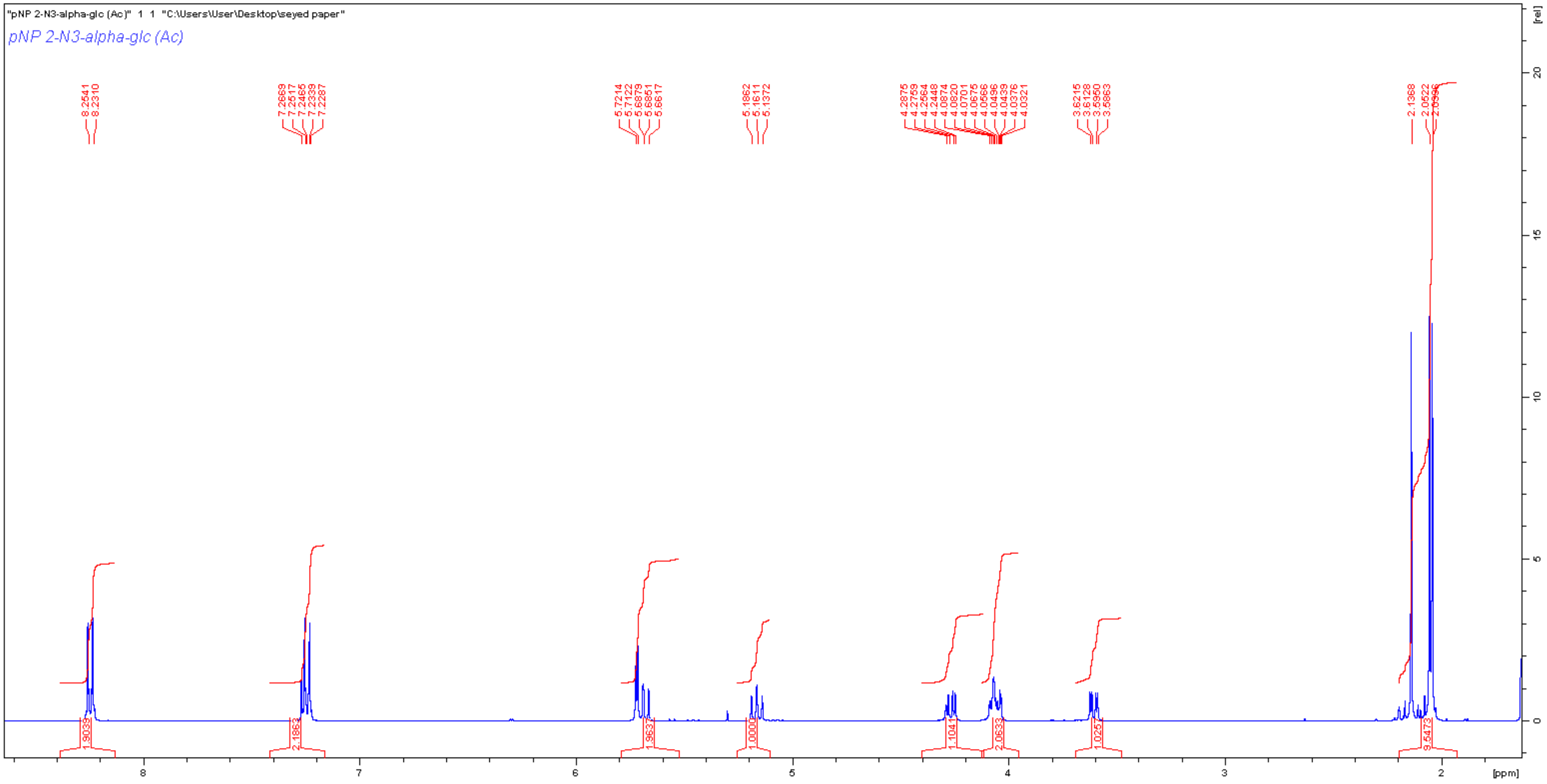
"pNP 2-N3-alpha-glc (Ac)" 1 1 "C:\Users\User\Desktop\seyed paper"
pNP 2-N3-alpha-glc (Ac)
[ppm]
[rel]

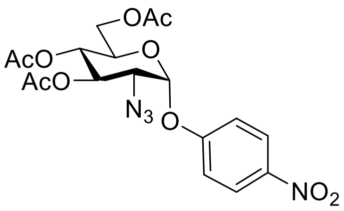

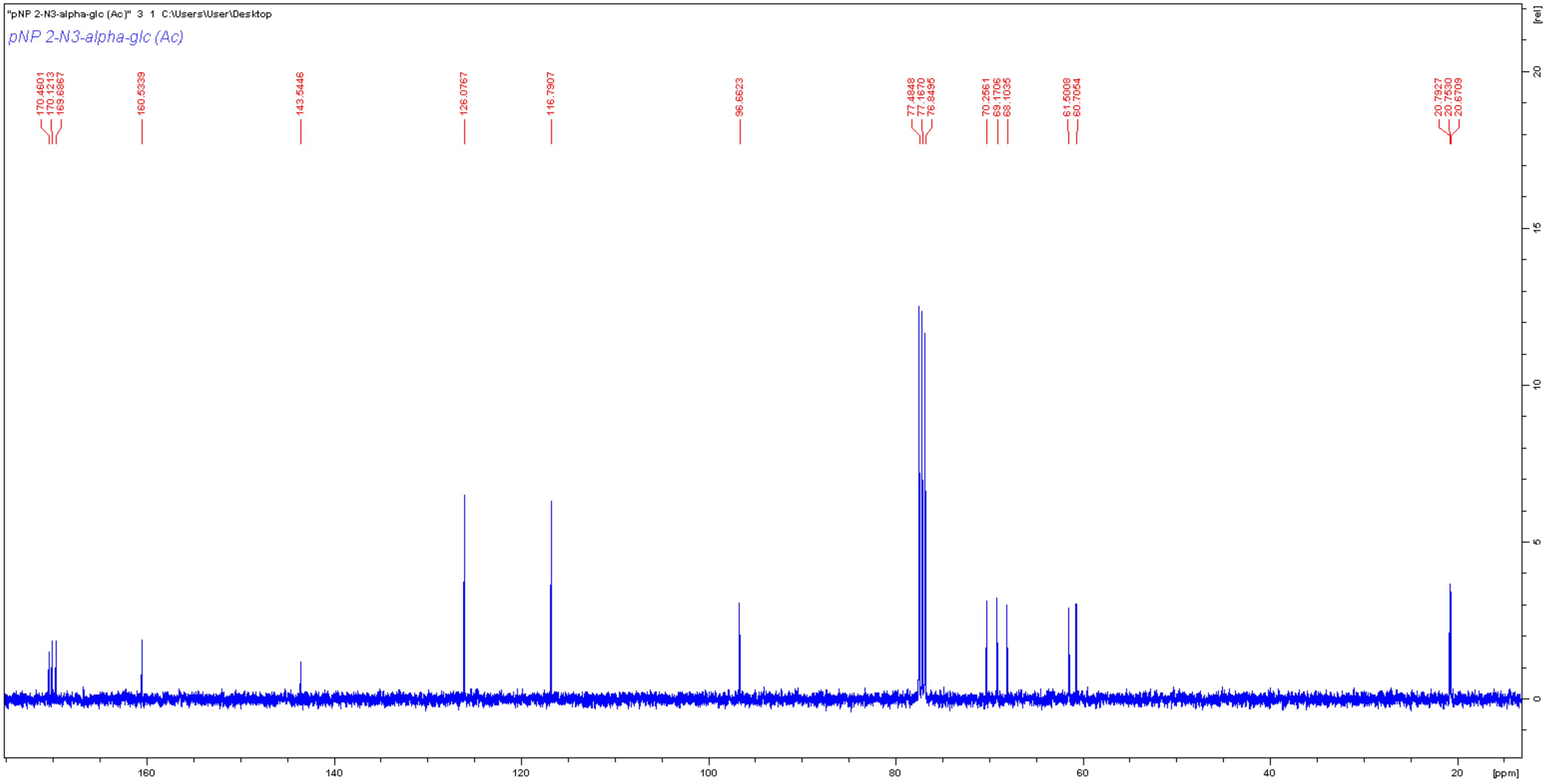

"pNP 2-N3-alpha-glc (Ac)" 3 1 C:\Users\User\Desktop
pNP 2-N3-alpha-glc (Ac)
170.4601
170.1213
169.6867
160.5339
143.5446
126.0767
116.7907
96.6623
77.4848
77.1670
76.8495
70.2561
69.1706
68.1035
61.5008
60.7054
20.7927
20.7530
20.6709
[rel]
[ppm]

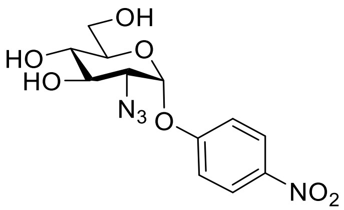

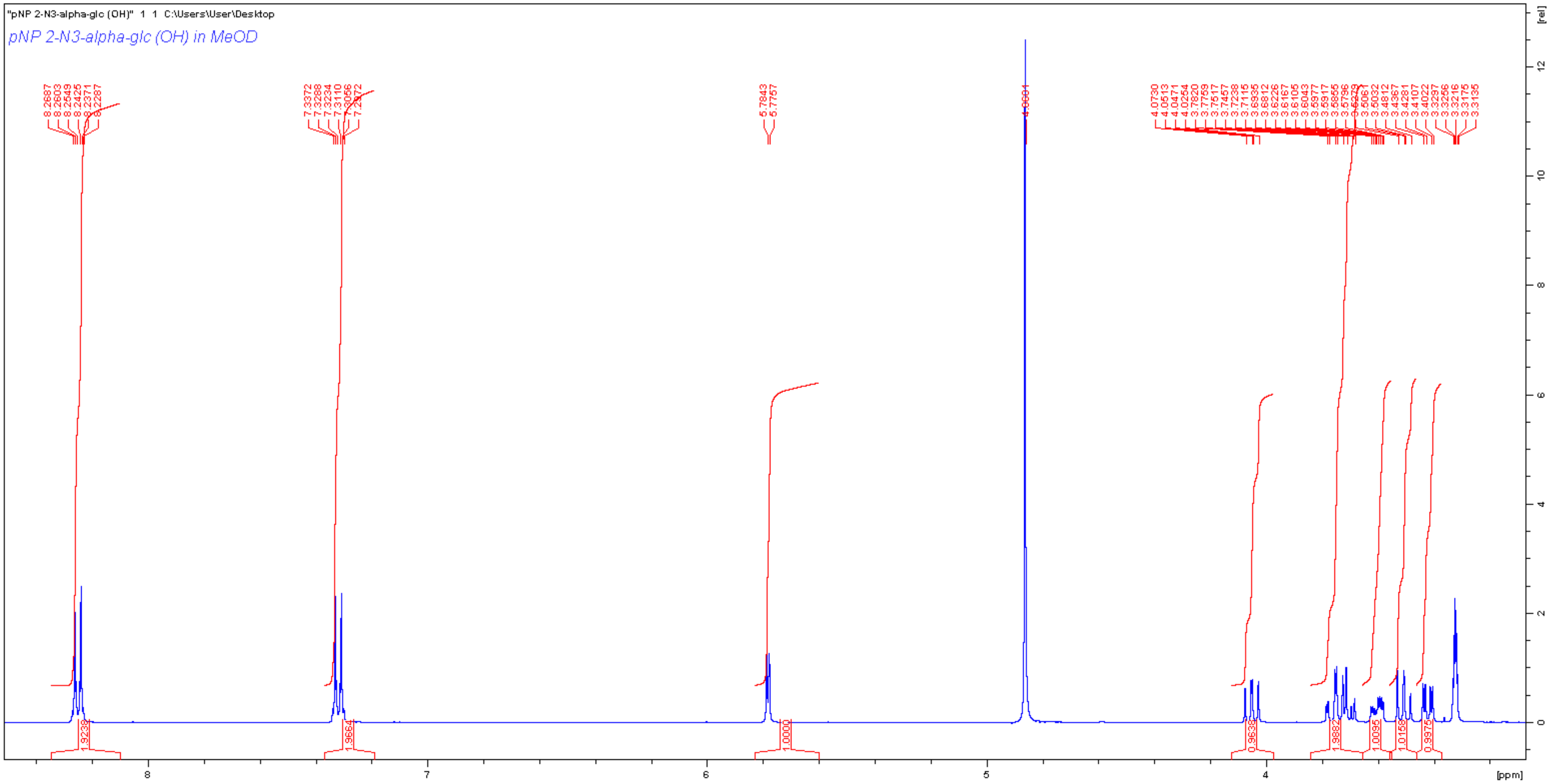
"pNP 2-N3-alpha-glc (OH)" 1 1 C:\Users\User\Desktop
pNP 2-N3-alpha-glc (OH) in MeOD
[ppm]
[rel]

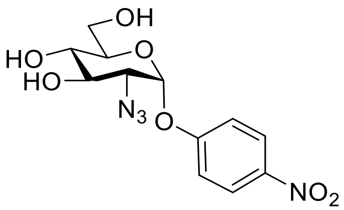

OH
HO
HO
O
N3
O
NO2

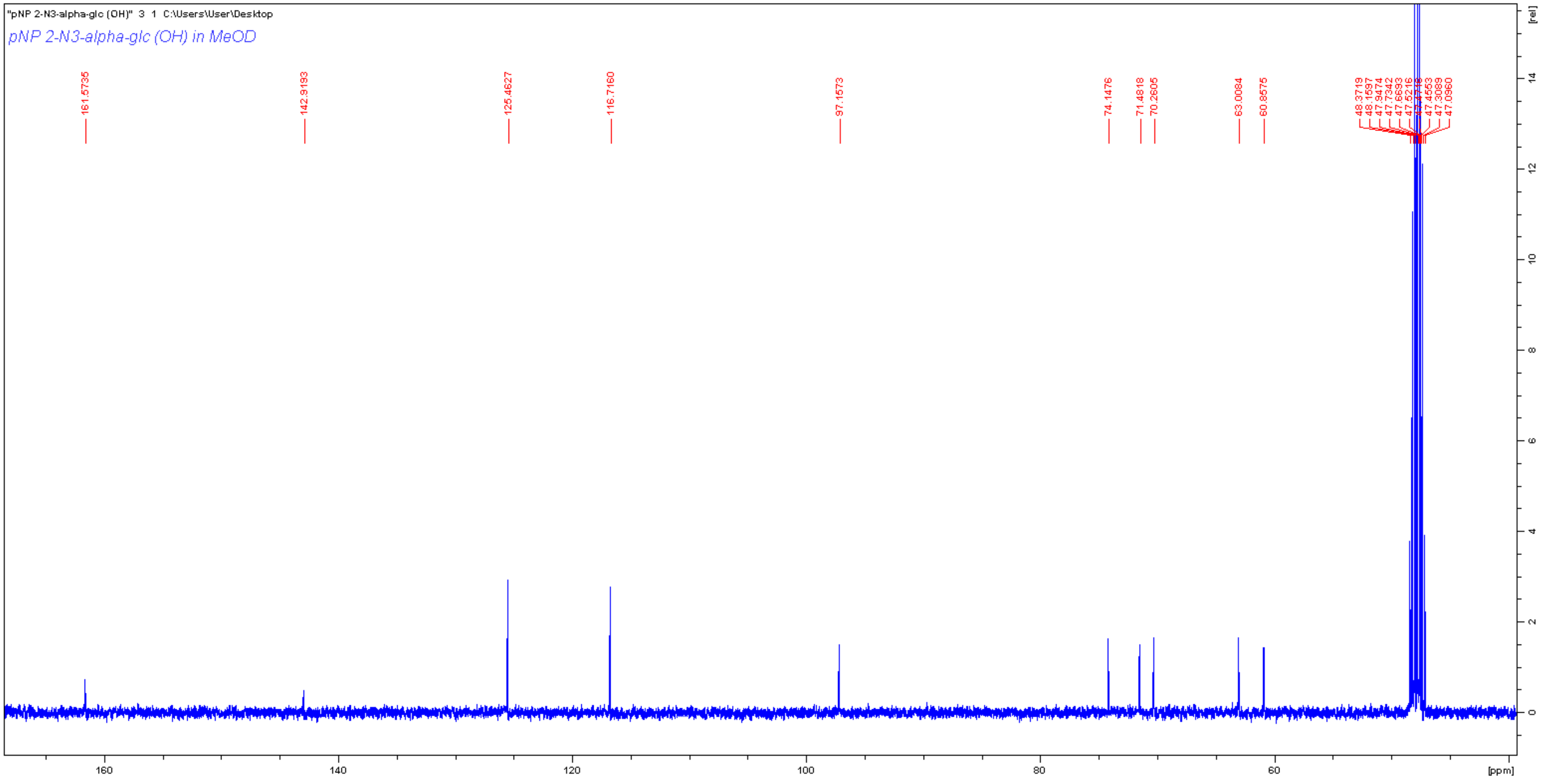

"pNP 2-N3-alpha-glc (OH)" 3 1 C:\Users\User\Desktop
pNP 2-N3-alpha-glc (OH) in MeOD
161.5735
142.9193
125.4627
116.7160
97.1573
74.1476
71.4818
70.2605
63.0084
60.8575
48.3719
48.1597
47.9474
47.7342
47.6693
47.5216
47.4553
47.3089
47.0960
[rel]
[ppm]
160
140
120
100
80
60

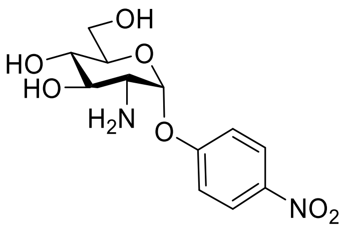
HO
HO
OH
O
H2N
O
NO2

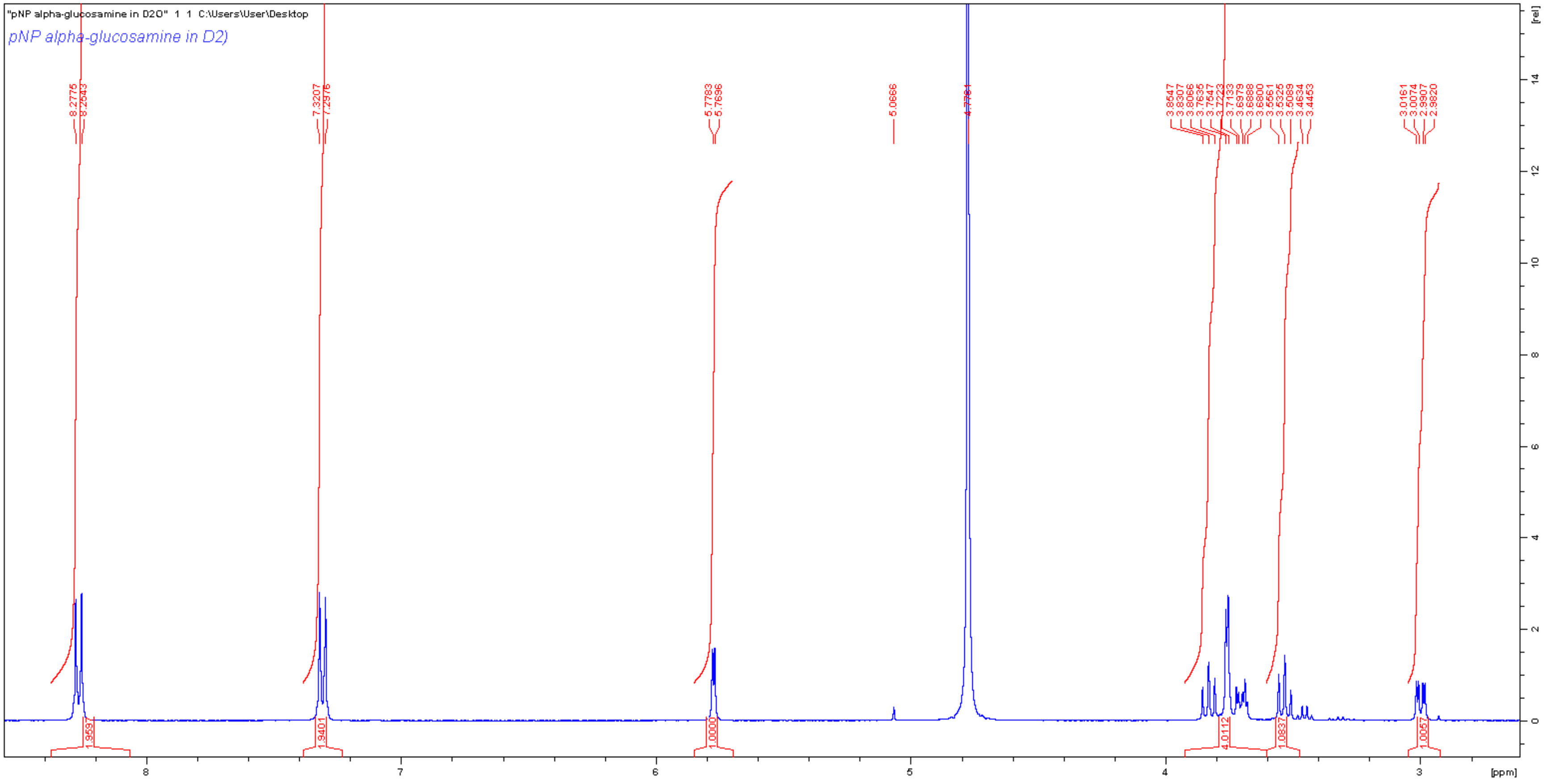
"pNP alpha-glucosamine in D2O" 1 1 C:\Users\User\Desktop
pNP alpha-glucosamine in D2)
8.2775
8.2543
7.3207
7.2976
5.7783
5.7696
5.0666
3.8547
3.8307
3.8066
3.7635
3.7547
3.7223
3.7133
3.6979
3.6888
3.6800
3.5561
3.5325
3.5089
3.4634
3.4453
3.0161
3.0074
2.9907
2.9820
1.9597
1.9401
1.0000
4.0112
1.0057
8
7
6
5
4
3
[ppm]
[rel]

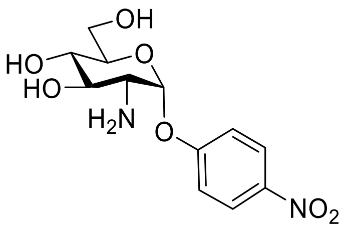

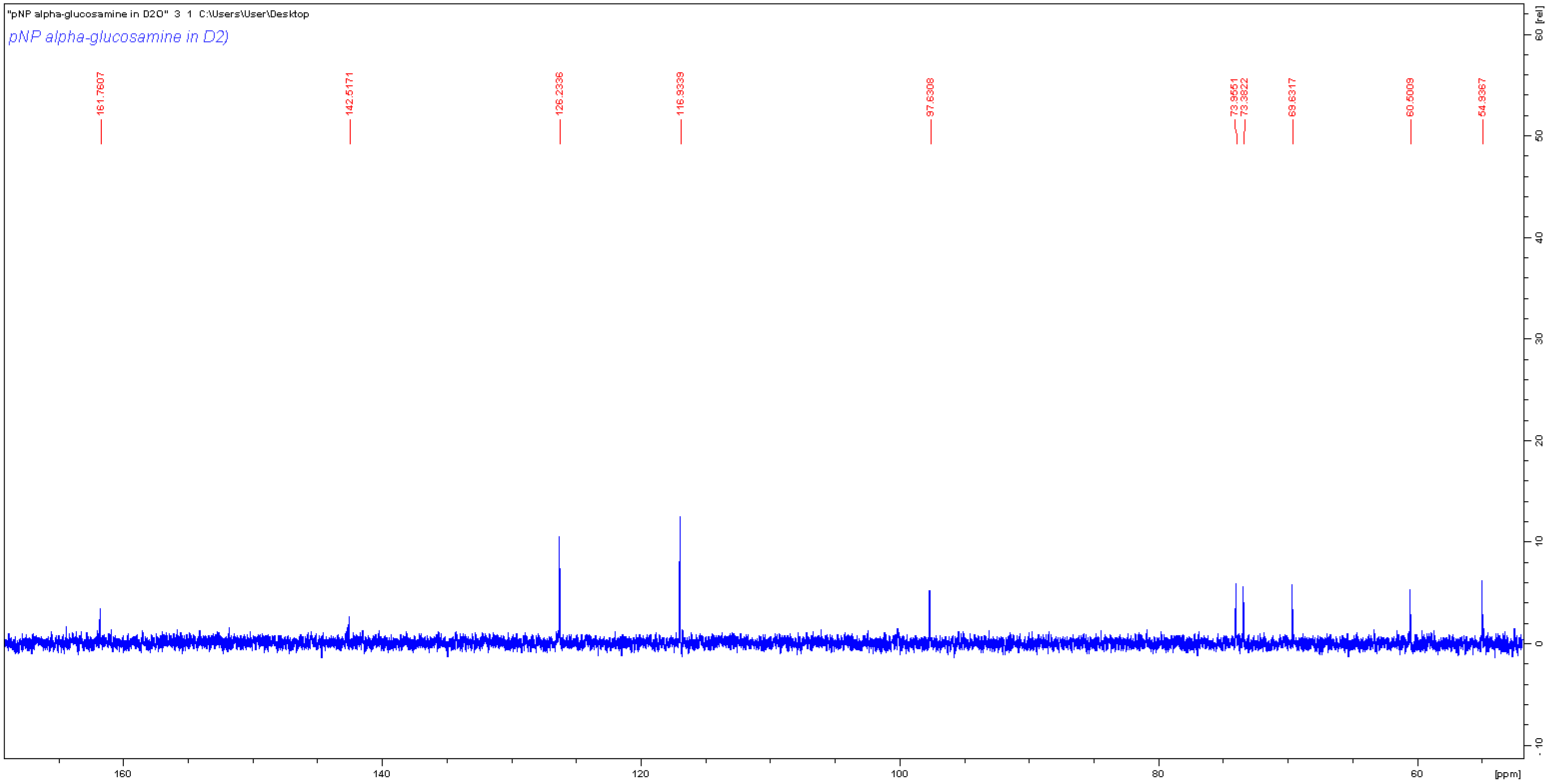
"pNP alpha-glucosamine in D2O" 3 1 C:\Users\User\Desktop
pNP alpha-glucosamine in D2)
161.7607
142.5171
126.2336
116.9339
97.6308
73.9551
73.3822
69.6317
60.5009
54.9367
[ppm]
[rel]

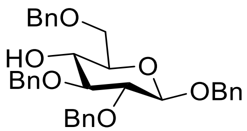
BnO
HO
BnO
O
OBn
BnO

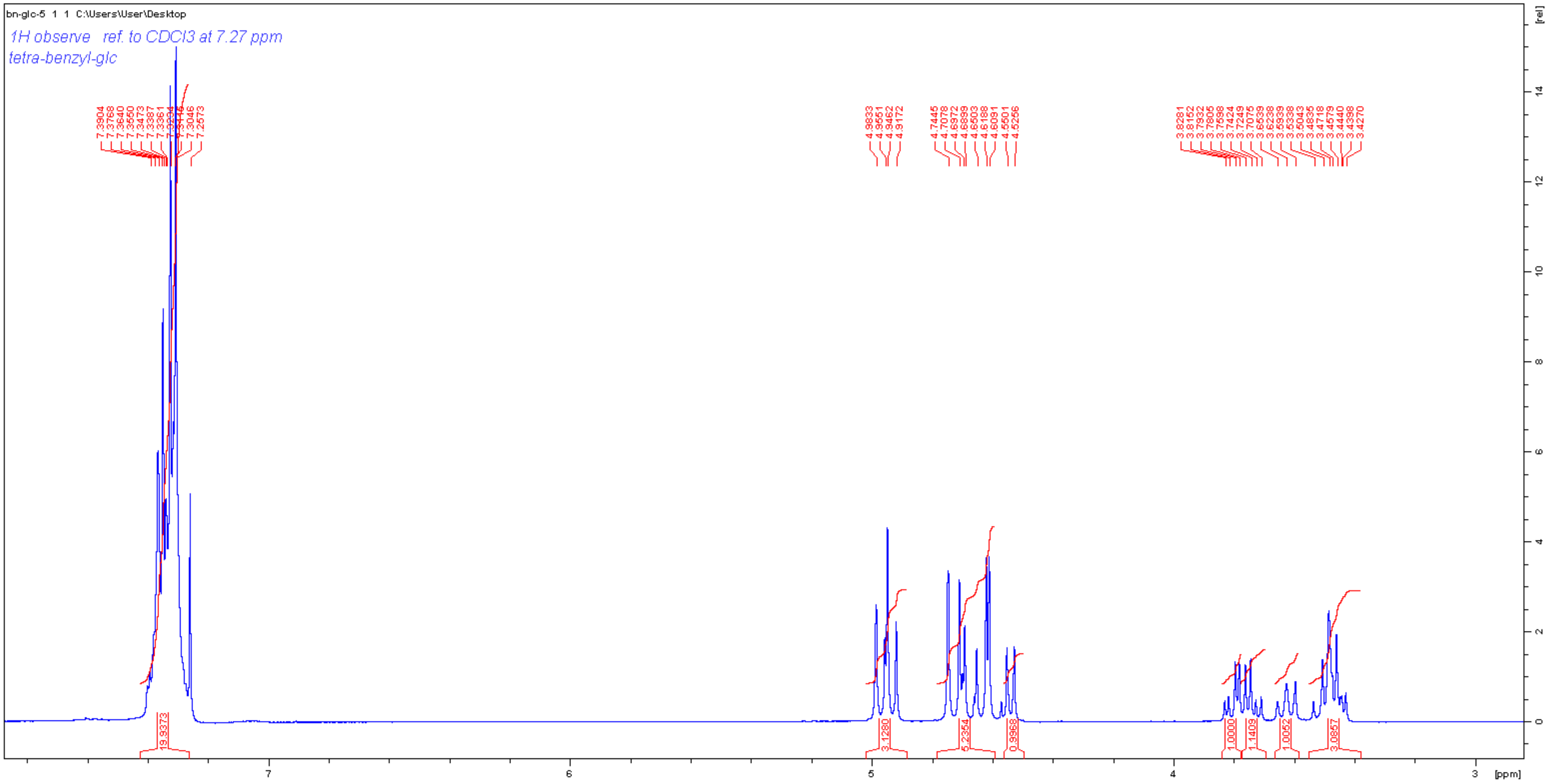
bn-glc-5 1 1 C:\Users\User\Desktop
1H observe ref. to CDCl3 at 7.27 ppm
tetra-benzyl-glc
7.3904
7.3768
7.3640
7.3550
7.3473
7.3387
7.3361
7.3046
7.2573
4.9833
4.9551
4.9462
4.9172
4.7445
4.7078
4.6972
4.6899
4.6503
4.6188
4.6091
4.5501
4.5256
3.8281
3.8152
3.7932
3.7805
3.7598
3.7424
3.7249
3.7075
3.6639
3.6238
3.5939
3.5338
3.5043
3.4835
3.4718
3.4579
3.4440
3.4398
3.4270
19.9373
3.1280
5.2354
0.9968
1.0000
1.1409
1.0052
3.0857
[rel]
[ppm]
7
6
5
4
3

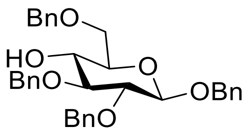
BnO
HO
BnO
O
OBn
BnO

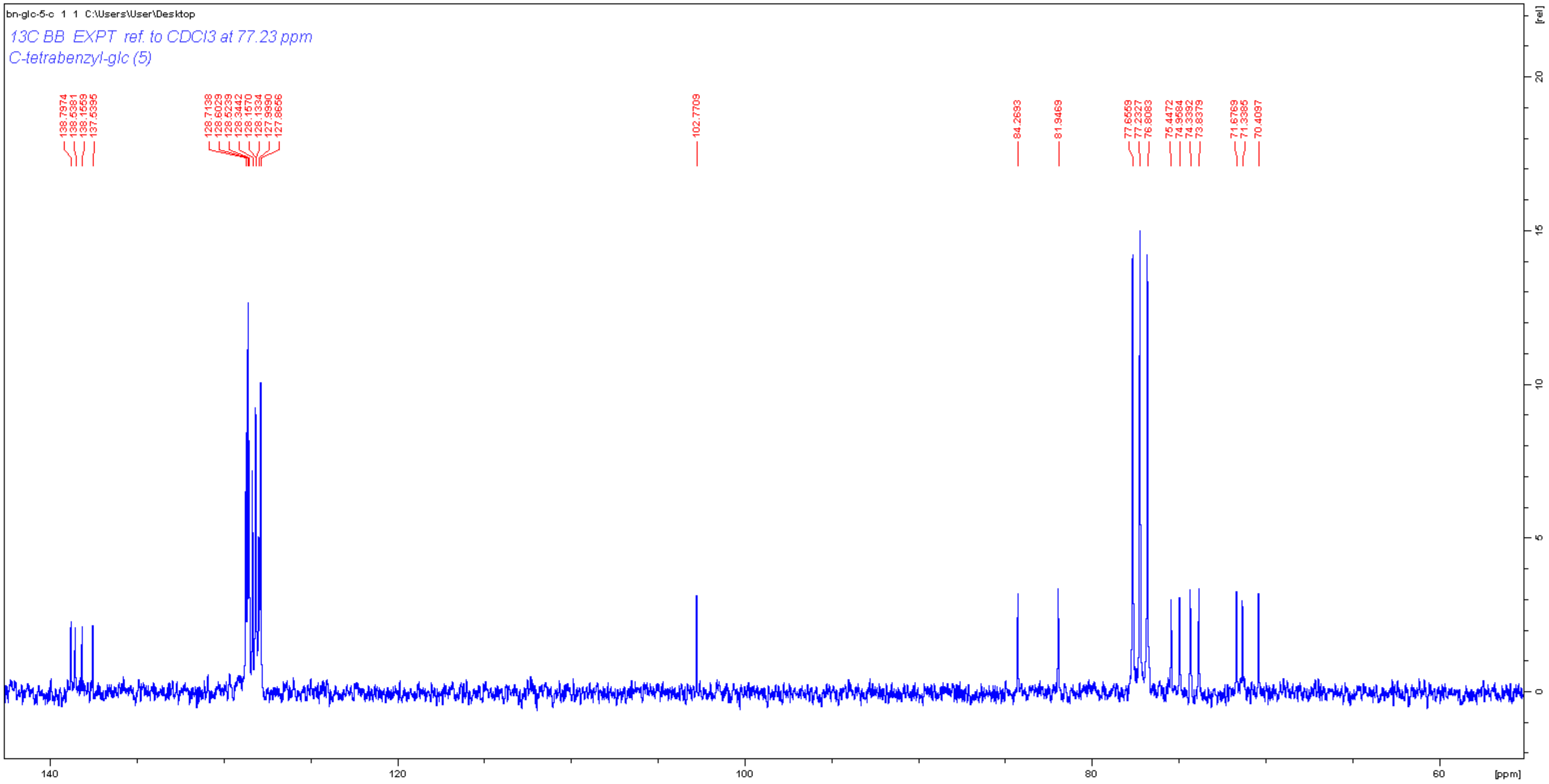
bn-glc-5-c 1 1 C:\Users\User\Desktop
13C BB EXPT ref. to CDCl3 at 77.23 ppm
C-tetrabenzyl-glc (5)
138.7974
138.5381
138.1559
137.5395
128.7138
128.6029
128.5239
128.3442
128.1570
128.1334
127.9990
127.8656
102.7709
84.2693
81.9469
77.6559
77.2327
76.8083
75.4472
74.9584
74.3392
73.8379
71.6769
71.3385
70.4097
[rel]
20
15
10
5
0
140
120
100
80
60
[ppm]

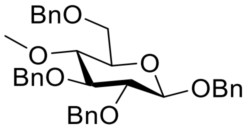
BnO
O
BnO
O
OBn
BnO

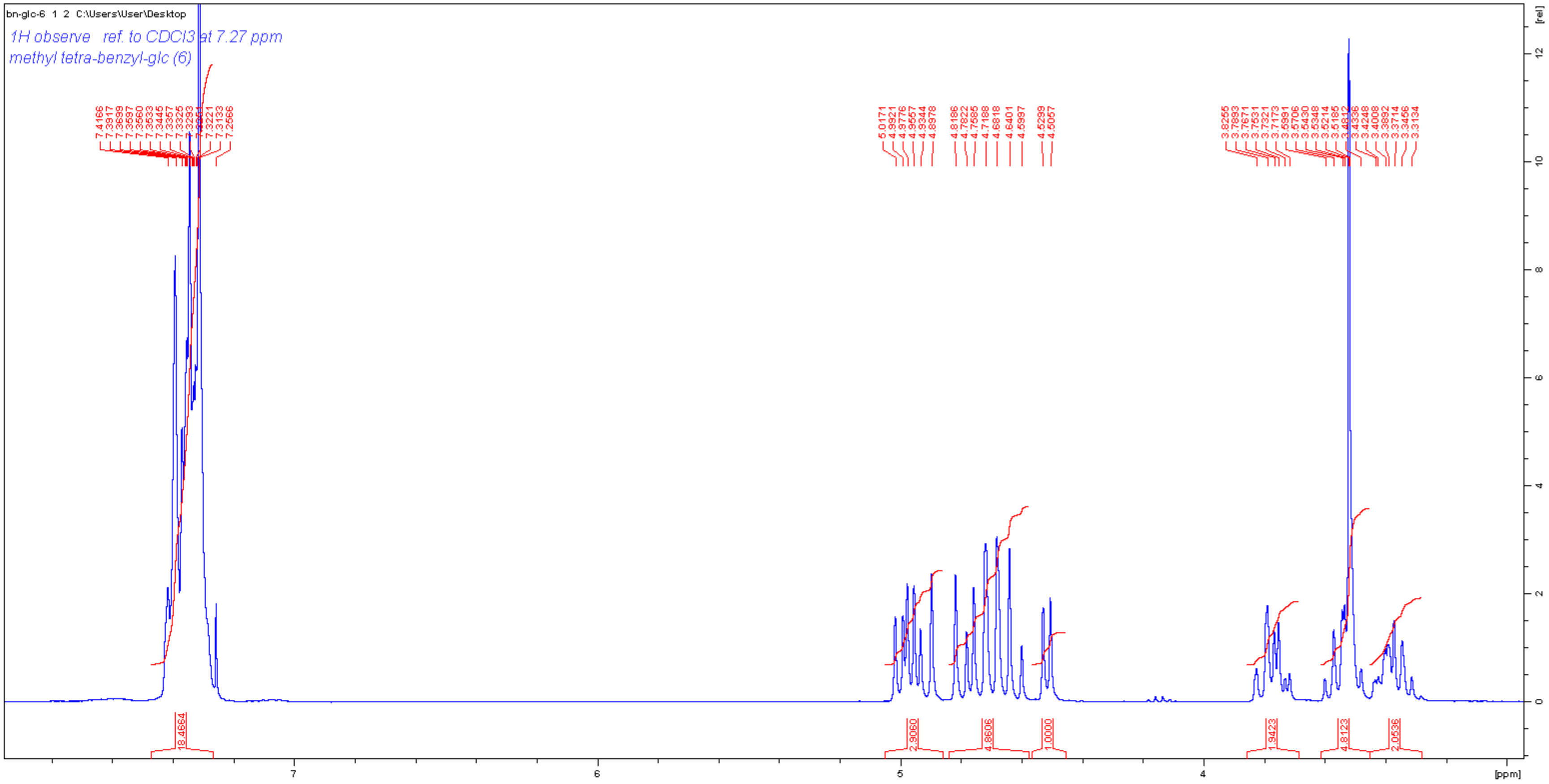
bn-glc-6 1 2 C:\Users\User\Desktop
1H observe ref. to CDCl3 at 7.27 ppm
methyl tetra-benzyl-glc (6)
7.4166
7.3917
7.3899
7.3597
7.3560
7.3533
7.3445
7.3357
7.3325
7.3293
7.3261
7.3221
7.3133
7.2566
5.0171
4.9921
4.9776
4.9557
4.9344
4.8978
4.8186
4.7822
4.7585
4.7188
4.6818
4.6401
4.5997
4.5299
4.5057
3.8255
3.7893
3.7671
3.7531
3.7321
3.7173
3.5891
3.5706
3.5430
3.5348
3.5214
3.5185
3.4812
3.4336
3.4248
3.4008
3.3892
3.3714
3.3456
3.3134
18.4664
2.9060
4.8606
1.0000
1.9423
4.8123
2.0536
[rel]
[ppm]

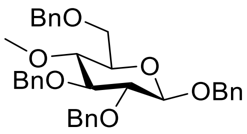

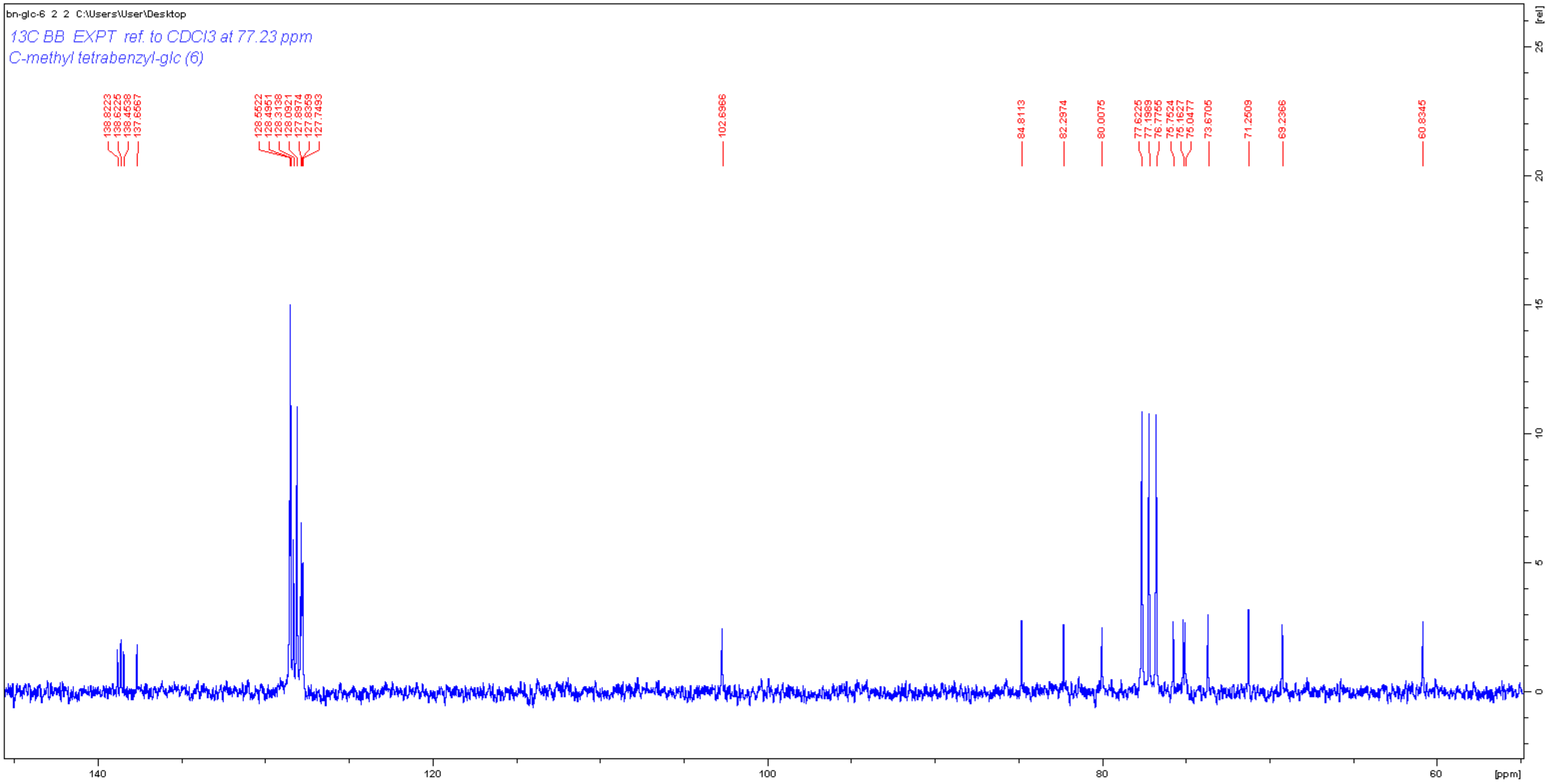

bn-glc-6 2 2 C:\Users\User\Desktop
13C BB EXPT ref. to CDCl3 at 77.23 ppm
C-methyl tetrabenzyl-glc (6)
138.8223
138.6225
138.4538
137.6567
128.5522
128.4951
128.3138
128.0921
127.8974
127.8359
127.7493
102.6966
84.8113
82.2974
80.0075
77.6225
77.1989
76.7755
75.7524
75.1627
75.0477
73.6705
71.2509
69.2366
60.8345
[rel]
[ppm]

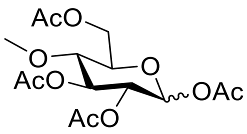

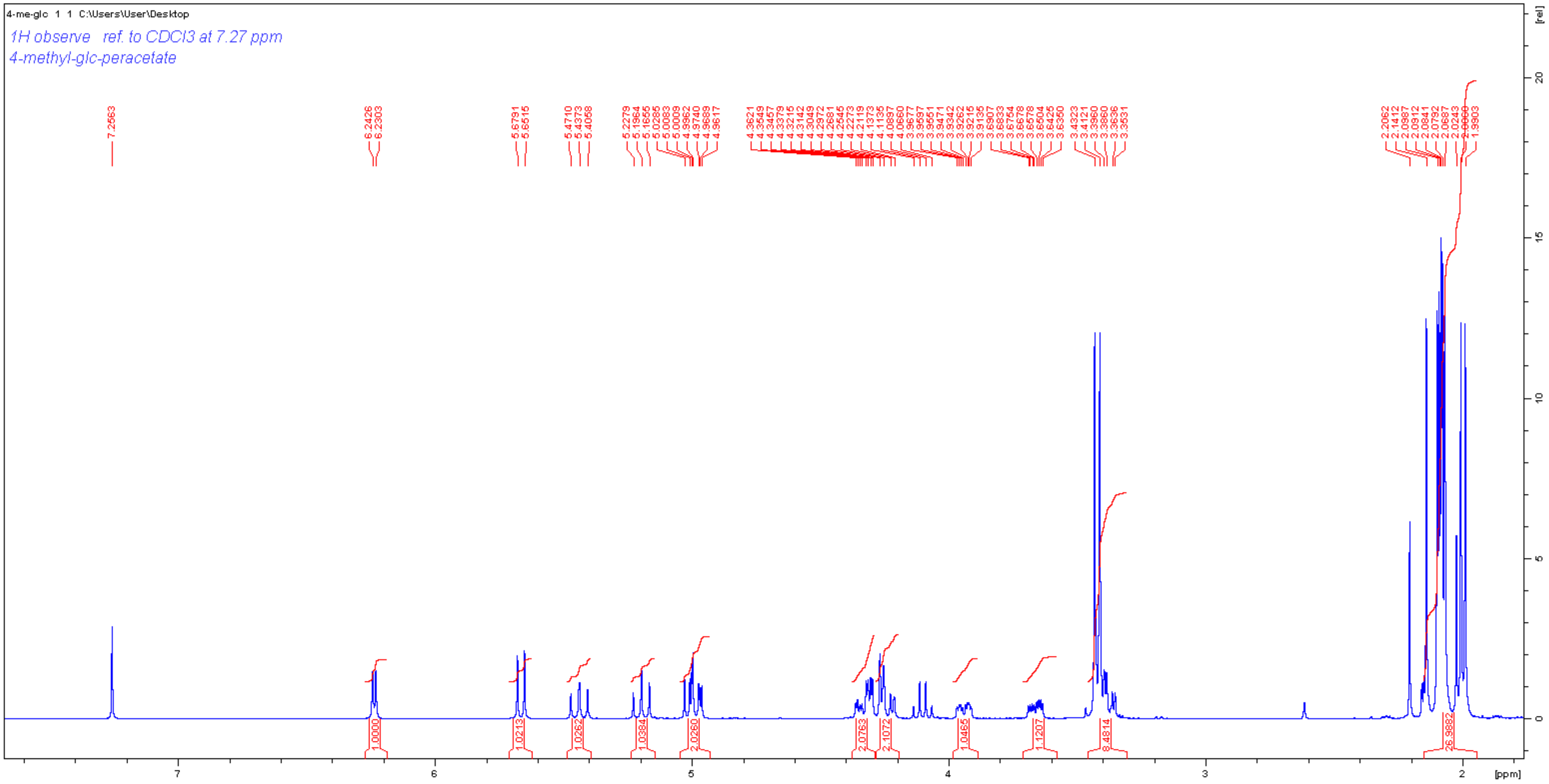
4-me-glc 1 1 C:\Users\User\Desktop
1H observe ref. to CDCl3 at 7.27 ppm
4-methyl-glc-peracetate
[ppm]
[rel]

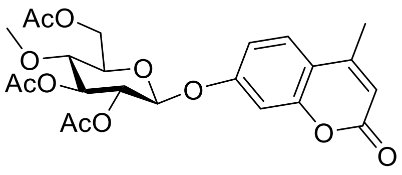
AcO
O
AcO
AcO
O
O
O
O

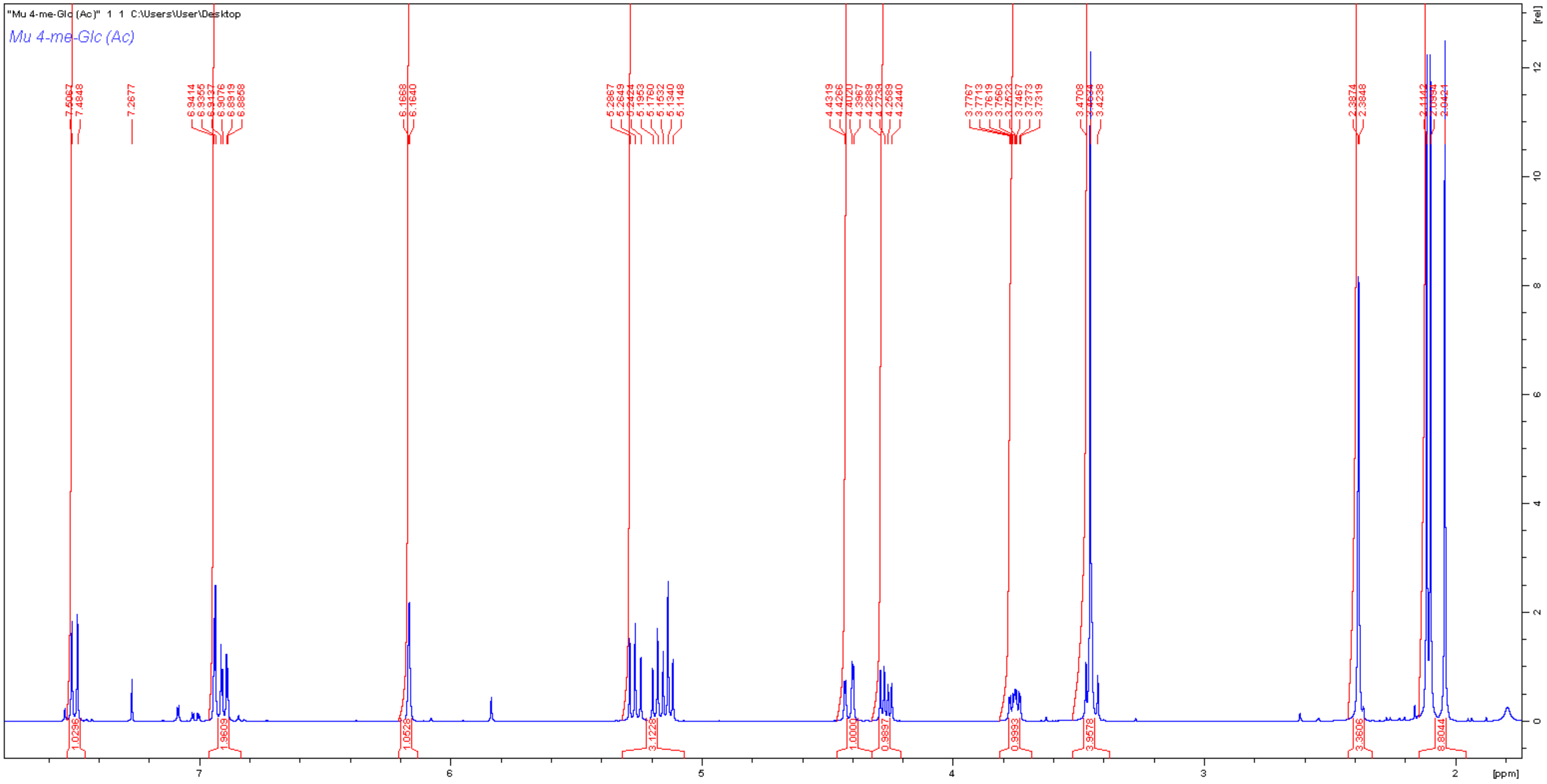
"Mu 4-me-Glc (Ac)" 1 1 C:\Users\User\Desktop
Mu 4-me-Glc (Ac)
7.5067
7.4848
7.2677
6.9414
6.9355
6.9137
6.9076
6.8919
6.8858
6.1668
6.1640
5.2867
5.2649
5.2424
5.1953
5.1760
5.1532
5.1340
5.1148
4.4319
4.4266
4.4020
4.3967
4.2889
4.2739
4.2589
4.2440
3.7767
3.7713
3.7619
3.7560
3.7523
3.7467
3.7373
3.7319
3.4708
3.4238
2.3874
2.3848
2.1142
2.0994
2.0421
1.0296
1.9609
1.0528
3.1228
1.0000
0.9897
0.9993
3.9578
3.3606
8.8044
[ppm]
[rel]

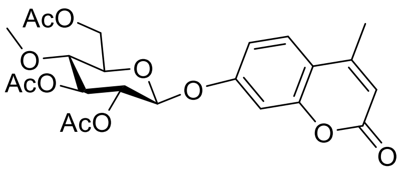

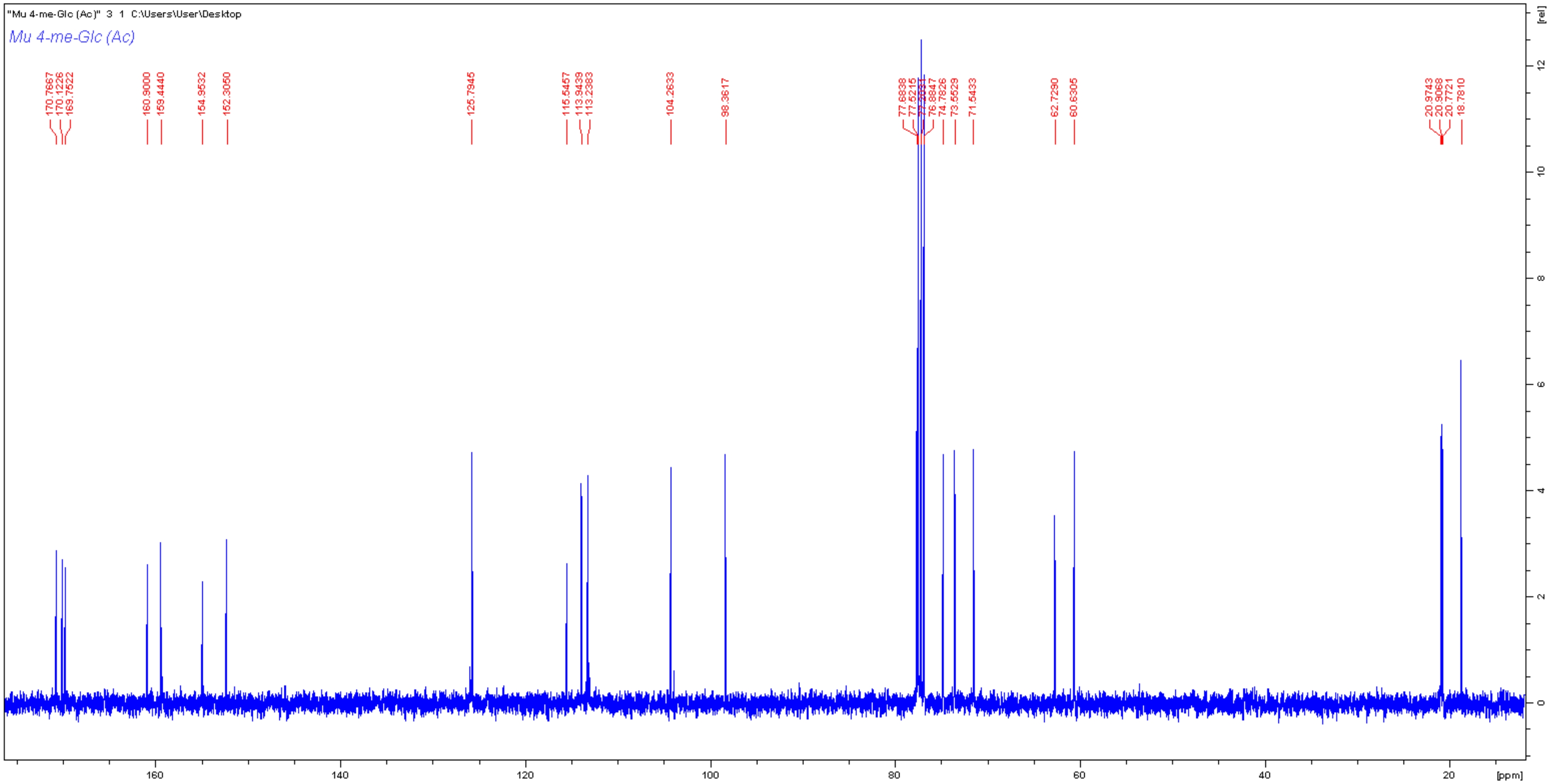
"Mu 4-me-Glc (Ac)" 3 1 C:\Users\User\Desktop
Mu 4-me-Glc (Ac)
170.7667
170.1226
169.7522
160.9000
159.4440
154.9532
152.3050
125.7945
115.5457
113.9439
113.2383
104.2633
98.3617
77.6838
76.8847
74.7826
73.5529
71.5433
62.7290
60.6305
20.9743
20.9068
20.7721
18.7810
[ppm]
[rel]

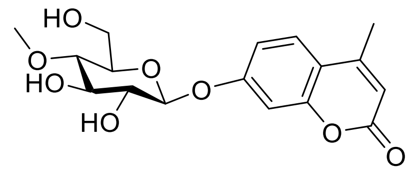

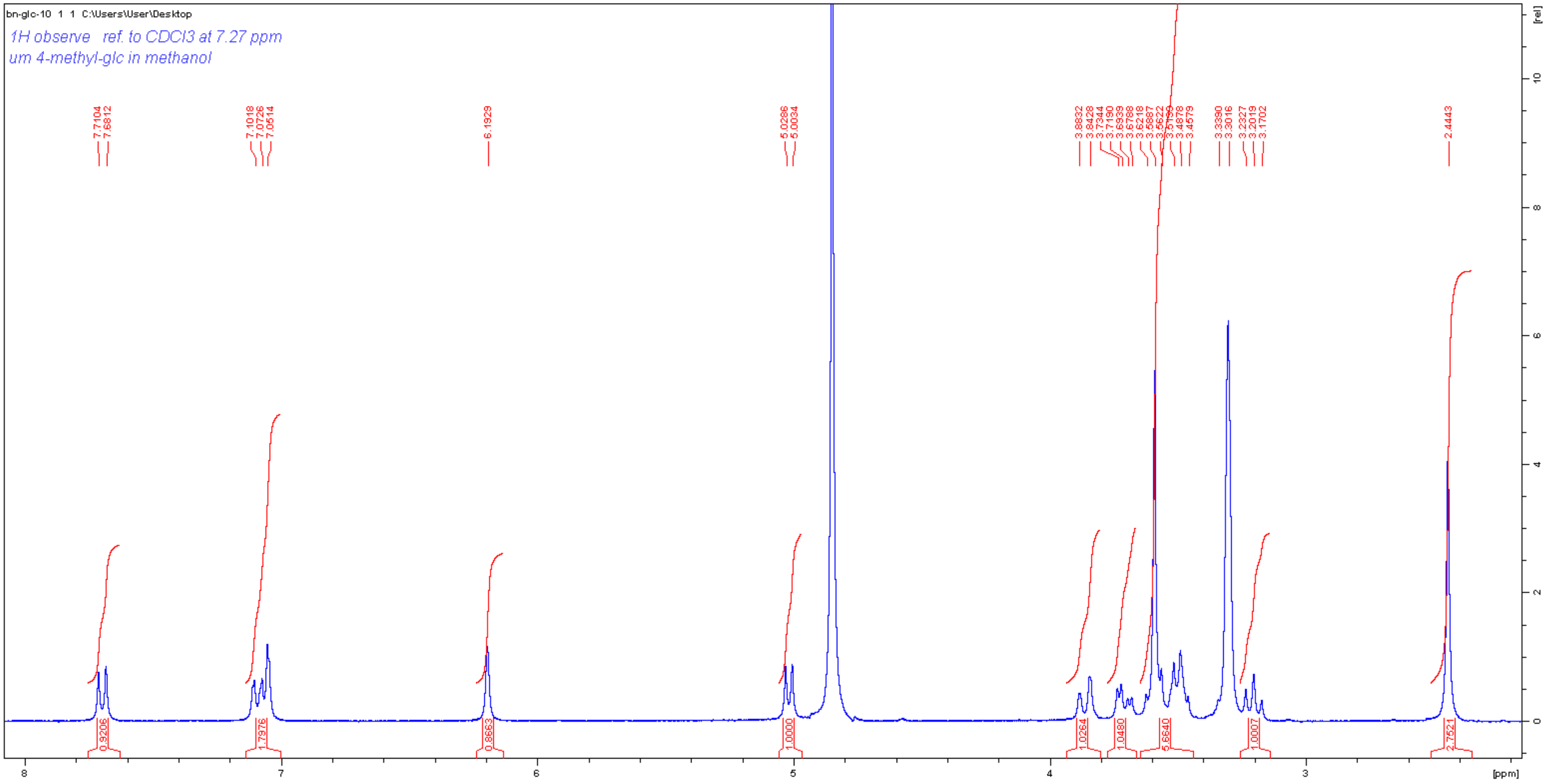
bn-glc-10 1 1 C:\Users\User\Desktop
1H observe ref. to CDCl3 at 7.27 ppm
um 4-methyl-glc in methanol
[ppm]
[rel]

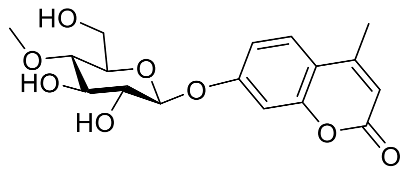

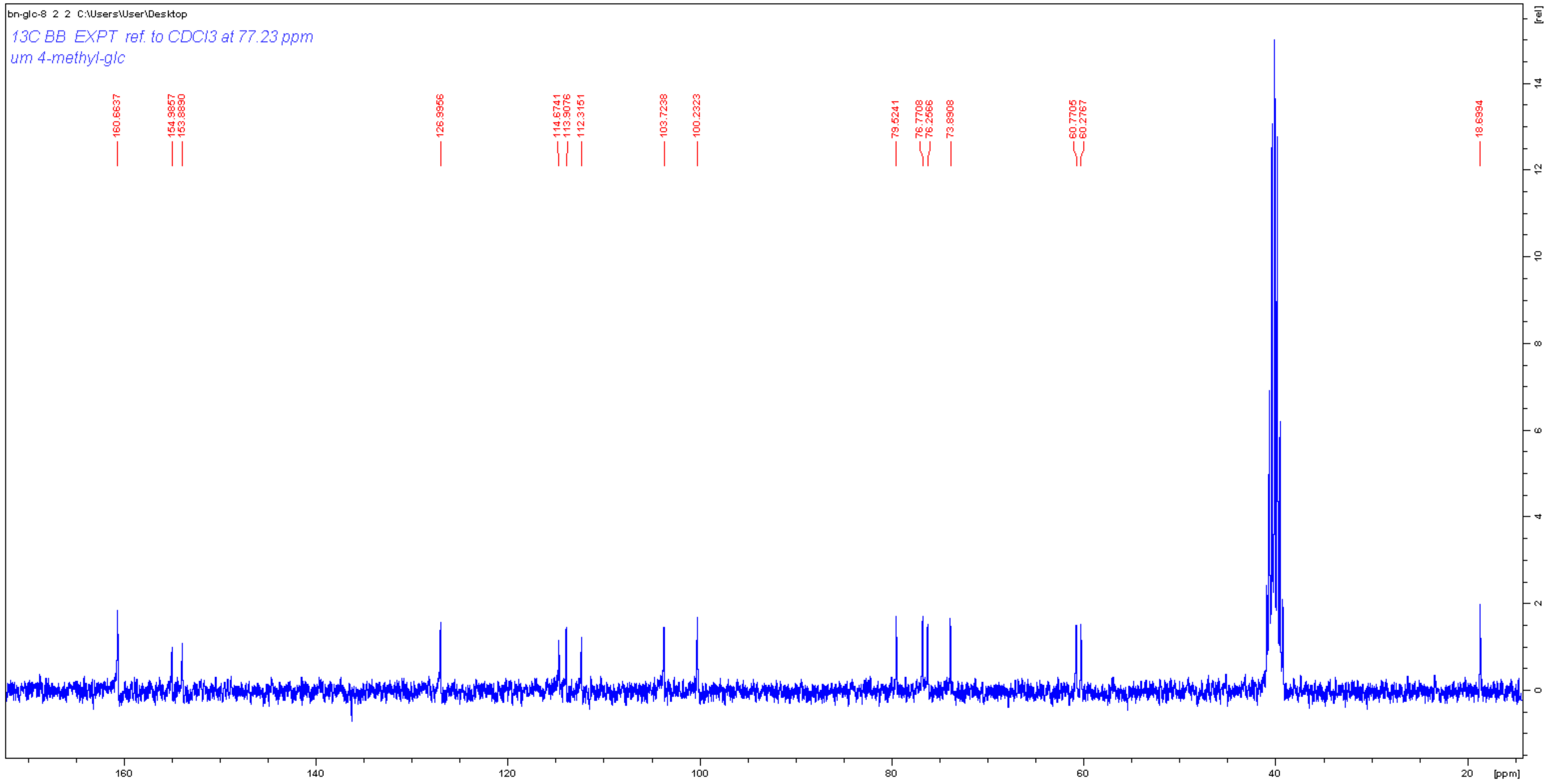

bn-glc-8 2 2 C:\Users\User\Desktop
13C BB EXPT ref. to CDCl3 at 77.23 ppm
um 4-methyl-glc
160.6637
154.9857
153.8890
126.9956
114.6741
113.9076
112.3151
103.7238
100.2323
79.5241
76.7708
76.2566
73.8908
60.7705
60.2767
18.6994
[rel]
[ppm]

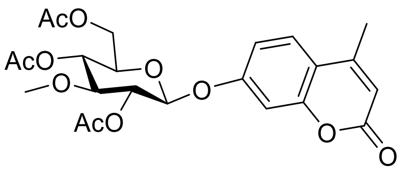
AcO
AcO
O
O
AcO
O
O
O

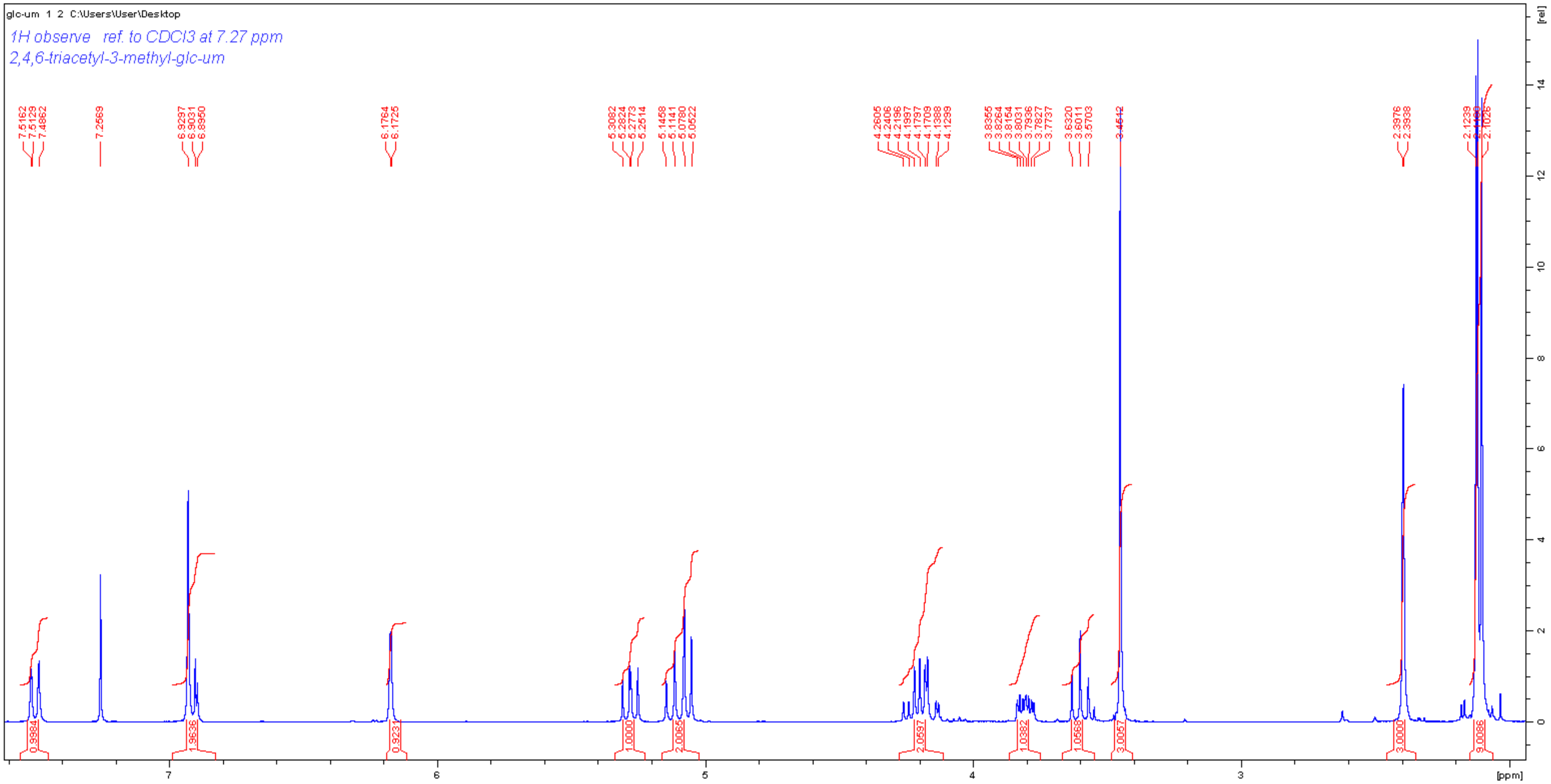
glc-um 1 2 C:\Users\User\Desktop
1H observe ref. to CDCl3 at 7.27 ppm
2,4,6-triacetyl-3-methyl-glc-um
[ppm]
[rel]

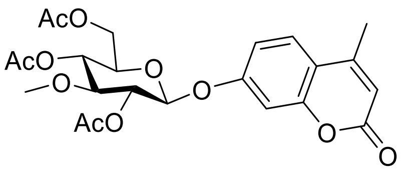
AcO
AcO
O
O
O
AcO
O
O
O

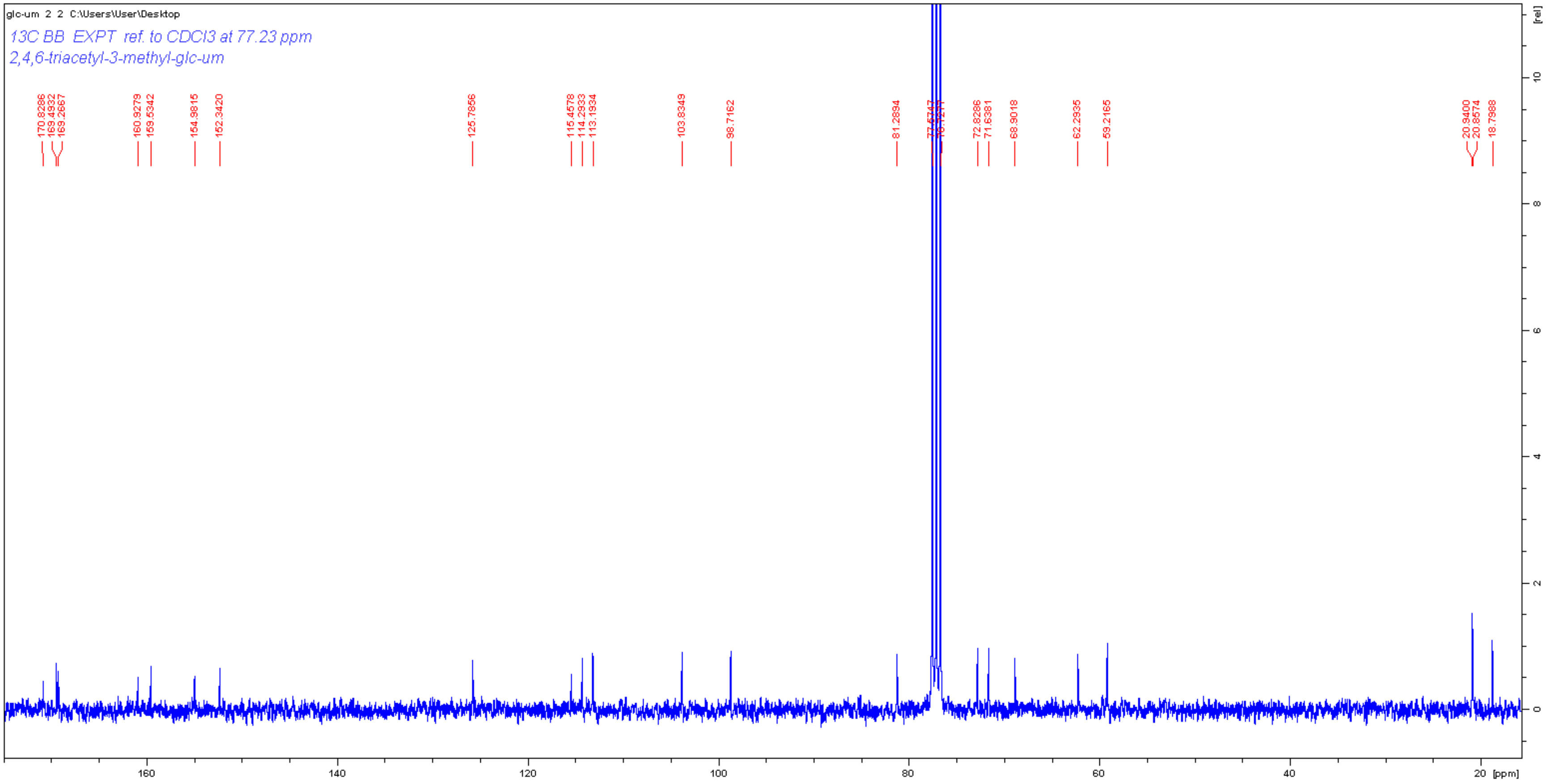
glc-um 2 2 C:\Users\User\Desktop
13C BB EXPT ref. to CDCl3 at 77.23 ppm
2,4,6-triacetyl-3-methyl-glc-um
170.8286
169.4932
169.2667
160.9279
159.5342
154.9815
152.3420
125.7856
115.4578
114.2933
113.1934
103.8349
98.7162
81.2894
72.8286
71.6381
68.9018
62.2935
59.2165
20.9400
20.8574
18.7988
[rel]
10
8
6
4
2
0
160
140
120
100
80
60
40
20
[ppm]

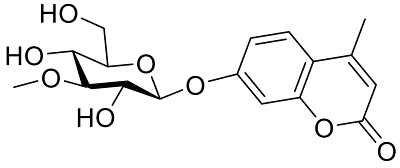
HO
HO
O
O
HO
O
O
O

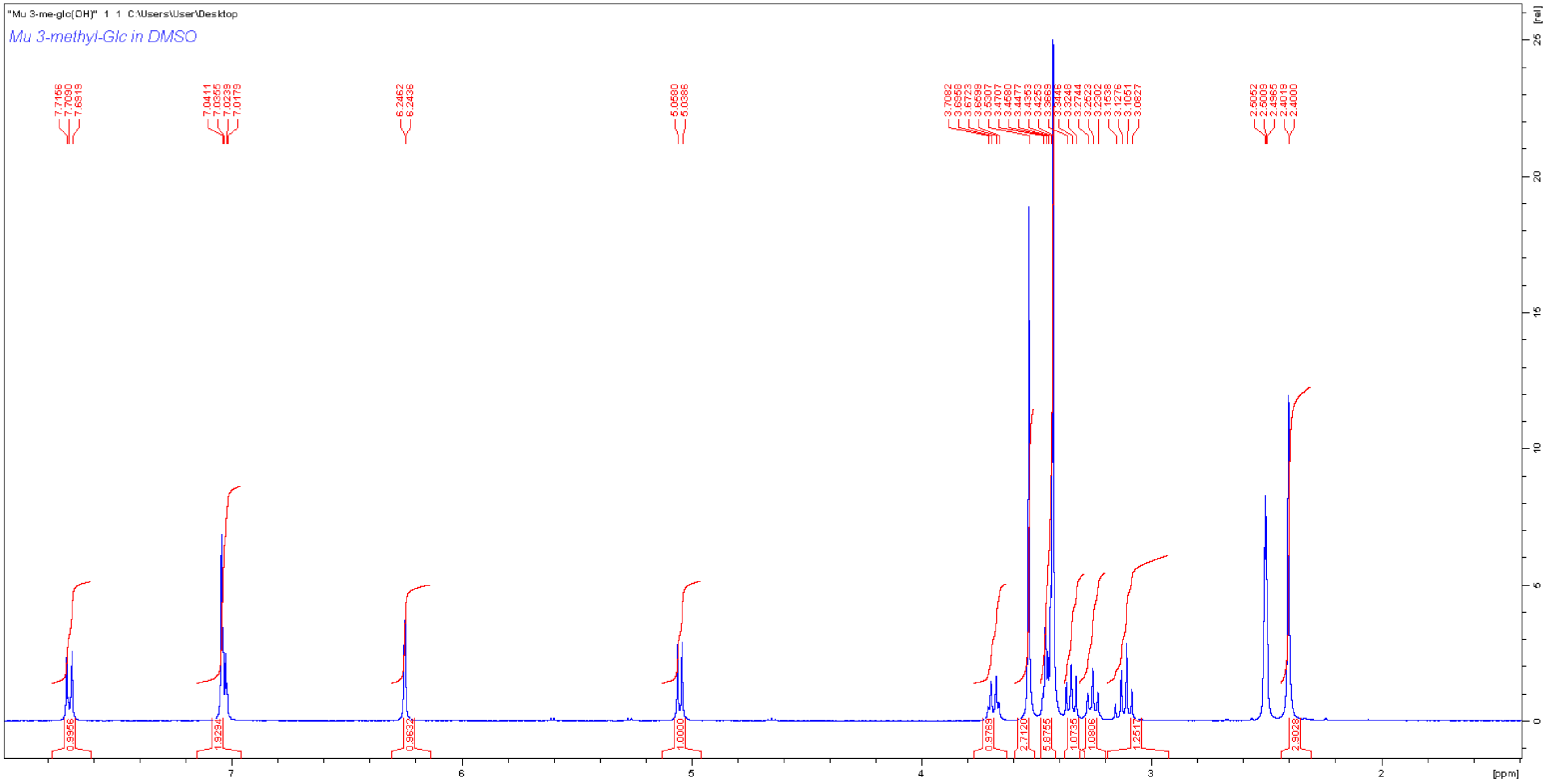
"Mu 3-me-glc(OH)" 1 1 C:\Users\User\Desktop
Mu 3-methyl-Glc in DMSO
7.7156
7.7090
7.6919
7.0411
7.0355
7.0239
7.0179
6.2462
6.2436
5.0580
5.0386
2.5052
2.5009
2.4965
2.4019
2.4000
0.9956
1.9294
0.9632
1.0000
0.9769
2.7120
5.8755
1.0735
1.0806
1.2517
2.9029
[ppm]
[rel]

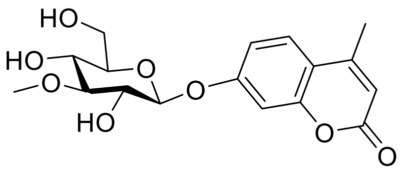

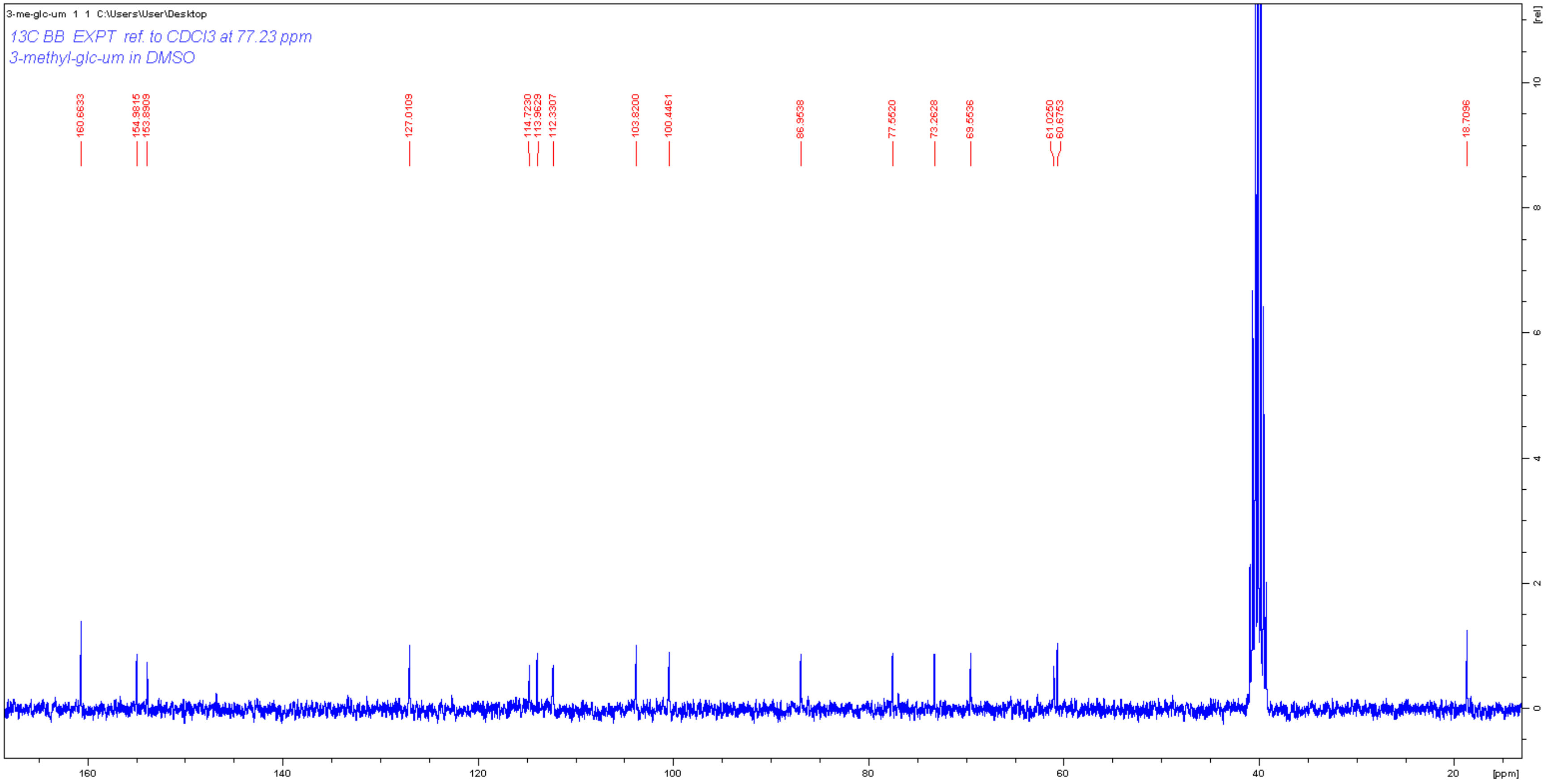

3-me-glc-um 1 1 C:\Users\User\Desktop
13C BB EXPT ref. to CDCl3 at 77.23 ppm
3-methyl-glc-um in DMSO
160.6633
154.9815
153.8909
127.0109
114.7230
113.9629
112.3307
103.8200
100.4461
86.9538
77.5520
73.2628
69.5536
61.0250
60.6753
18.7096
[ppm]
[rel]

#### 5. Nucleotide and amino acid sequences

##### P1C11 Fosmid sequence

Highlighted yellow: AL1, Green:AL2, Turquoise: AL3, Pink: AL4, Teal: AL5

CGGTCTGATTATTAGTCTGGGACCACGGTCCCACTCGTATCGTCGGTCTGATAATCAGACTGGGACCACGGTCCCACTCGTATCGTCGGTCTGATTATTAGTC
TGGGACCATGGTCCCACTCGTATCGTCGGTCTGATTATTAGTCTGGGACCACGGTCCCACTCGTATCGTCGGTCTGATTATTAGTCTGGAACCACGGTCCCAC
TCGTATCGTCGGTCTGATTATTAGTCTGGGACCACGGTCCCACTCGTATCGTCGGTCTGATTATTAGTCTGGGACCACGATCCCACTCGTGTTGTCGGTCTGA
TTATCGGTCTGGGACCACGGTCCCACTTGTATTGTCGATCAGACTATCAGCGTGAGACTACGATTCCATCAATGCCTGTCAAGGGCAAGTATTGACATGTCGT
CGTAACCTGTAGAACGGAGTAACCTCGGTGTGCGGTTGTATGCCTGCTGTGGATTGCTGCTGTGTCCTGCTTATCCACAACATTTTGCGCACGGTTATGTGGA
CAAAATACCTGGTTACCCAGGCCGTGCCGGCACGTTAACCGGGCTGCATCCGATGCAAGTGTGTCGCTGTCGACGAGCTCGCGAGCTCGGACATGAGGTTG
CCCCGTATTCAGTGTCGCTGATTTGTATTGTCTGAAGTTGCTTTTACGTTAAGTTGATGCAGATCAATTAATACGATACCTGCGTCATAATTGATTATTTGACGT
GGTTTGATGGCCTCCACGCACGTTGTGATATGTAGATGATAATCATTATCACTTTACGGGTCCTTTCCGGTGATCCGACAGGTTACGGGGCGGCGACCTCGC
GGGTTTTCGCTATTTATGAAAATTTTCCGGTTTAAGGCGTTTCCGTTCTTCTTCGTCATAACTTAATGTTTTTATTTAAAATACCCTCTGAAAAGAAAGGAAACGA
CAGGTGCTGAAAGCGAGCTTTTTGGCCTCTGTCGTTTCCTTTCTCTGTTTTTGTCCGTGGAATGAACAATGGAAGTCCGAGCTCATCGCTAATAACTTCGTATA
GCATACATTATACGAAGTTATATTCGATGCGGCCGCAAGGGGTTCGCGTCAGCGGGTGTTGGCGGGTGTCGGGGCTGGCTTAACTATGCGGCATCAGAGCA
GATTGTACTGAGAGTGCACCATATGCGGTGTGAAATACCGCACAGATGCGTAAGGAGAAAATACCGCATCAGGCGCCATTCGCCATTCAGCTGCGCAACTGT
TGGGAAGGGCGATCGGTGCGGGCCTCTTCGCTATTACGCCAGCTGGCGAAAGGGGGATGTGCTGCAAGGCGATTAAGTTGGGTAACGCCAGGGTTTTCCCA
GTCACGACGTTGTAAAACGACGGCCAGTGAATTGTAATACGACTCACTATAGGGCGAATTCGAGCTCGGTACCCGGGGATCCCACGTGCGAAGAGAGCACCT
TCATGCCTACGGCCTCCACGTCGCGCCTGAACTCCTCGGGGGTGTTGCCGTAGAATTTACCGTCGTTGTAACTGGCGGCCTCGATGCTGGTGTAACCCATGT
CGGCGAGTGCCTTCAGGACGGGTTTGTAGTCGTGCTGGTTTCTTCCGAAACTCCCGATCAGATCCCTGACCGAGTAGAGCTGGATACCGATATCCTTCTTTG
CGTCCTTCTTCGCTTTGGCAAATGCCCCGCAGGGAGCAATTGCCGAAGCGAGCAGGATGCAAACCGCTGTAAACAGTTTACGTTTCATACGTTACCGTTGTTA
ACGGAATTAGTCGAGCACCTTGATGCGGATGTTGCGGAACCATACATCGTCGCCGTGATCCTGGAAGCCGATGTAACCTTCGCGCTGTGCTCCGCCGCAGTT
GTTCAGCAGTTCGAATGCCAGGGGCCACTTCTCGGGGCTGAACTTGCTCTTTTCGAGCATCTCCGTCCACTGGGGAGTCCAGAGGTGGTACTCGAGCACGTT
CTGGTCGTTCTGGCCGTGTACCACGGTGCCTTTGTAAACCATGATCTTCGCCTTGTTCCACTCGCCATAGGGCTTGGCGTTCTGGGGAACGGCCGGGATCAT
GTCGTAAAGCGAAGCCGACTTGCGGTTGCCGTCGACGCCCAGTTTGGCATCGGGATGGTTCTCGTTGTCGAGTACCTGGTACTCGGGGCAGGAGATGTAGA
TGGGCTCCATCCGCTGTTTGCCGTCCTTGGTCGTGGCTACCTCCTGGCCGAGGTAGAAGATACCCGAGTTGCTGCCTTTGGCAACTTTCCACTCCACTTCGA
GCTCGAAGTTCTTGAACTTGTGCGTGAAGATCAGGTCGCCGCCGTCGCCGTCCTGGGCTTCACCGCCGCCCGAACCGTTGAATTTGATCGCACCGTTGTCGA
TCGTCCATTTCGAGGGCACGTGGTCTTTGCCGTAGCCGCGCCAGCCGTTGAACGTCTTGCCGTCGAAGATCACGTAGTAGCCGTCCTTGTCCTGCGGGAAAC
CTGCGAGGTCTACCGTCGCATTGTTGATGACCGTGTATTCGGGTACGGCTTTGGTTTCTGCCGCTGCCGTGTCGGCTGCCGCTTTGCCTTTGGCCCCCGGGC
CGCCGCATGCCGCCATCGTTGCGGCTGCTGCGATGCATGTCGAGAATAAAATGACTTTTTTCATATTCTTTCCGTTGTTTTAATTTGAGTTGATAATCTTGTCCG
TAGATTAACGCGGCATGTCGGGCAGTTTCCAACCGTCGCGGTAGGTGTGCTTGATGAGCTCCTGTGCGAATTGCTGTGCGTTGACCGGATCCGTCCAGGTCT
TGTCGAAGGTCGGATGGCCGTCCTTGATGTGGAACCCGTCCTTGATGACGGCACTGATCGTGGCGTCATCGGGGATGTTCGTGAAACGCATGTTCGGGCCG
TCCCAGAGCAACTCGCGGTTCAGGTTCTGCAGGCGGACTGCCAGCACGCCCATAACCACCATTTCATTGAACGGGCCGGCTTCGCTGAAGTCCGATGCGCT
CGGCACGCGGTCGTCGGCATCCTCCTTGCAGGCACGTACCCAGTCCATCTGGTGCGACTCCTTGATCTCGCGCAGTACCTTCGGTGCGTTGGGCACGCGAC
CCGAAACCAGGTAGGGGTTCACGCCGTAGCAGCCGCAGATCAGCGTATCCTTCGTACCGTAGAAAATAACGCCGCCGCCGGCCATGTTCAGGTCTTTGCCT
GCGGGCAGCCCTTCGGGGCGCGCCGGTTTCAGGCCGCCGTCGTACCAGTATACCTCTACCTCGGGCATGGCTACCTTCGGCATGTTGTCGCGTGCCGGGA
AGACGAACTTCACCGTCTGCGCCATCGGTGCGGATTCGTTGAGCAGCAGCGTCGAGCTACCCTGAACCTTGGTCGGATAGCCGAGCTTCAGGCCCTTGAATA
CCGGGTGGAGGATGTGGCAGGCCATATCGCCCAGTGCGCCGGTACCGAAATCCCACCAGCCGCGGAAGTTCCACGGGGTGTAGATCGCGTTGTAGGGACG
GTAGGGAGCCGGGCCGATGAAGGCGTCCCAGTTCAGCGTCTTGGGGATGCGCTGGTCGTCCTCGGGACGTGAGAGCCCCTGCGGCCAGATAGGACGGTC
GGTGAAGGTCTCCACCTTGCGGACTTCGCCGATCTCTCCGTTCCAGATCCATTCGCAGACCTTGCGGACACCCTCGTCCGACGCACCCTGGTTACCCATCTG
GGTGGCTACTTTATATTTGTCGGCGAGCTTGGTGAGCAGACGCGATTCGTATACGGTATGCGTCAGGGGCTTCTCGACGTAGACGTGCTTGCCGGCCGTCAT
CGCATCGGCTGCGATGATCGCGTGCGTGTGGTCGGCCGTAGCCACCATCACGGCGTCTGCCGACTTGAGCATTTCGTCGAACATCTTGCGGTAGTCGTTGT
ACTTCTTGGCAGCCGGGTAGCGCTTGAACACGTGGTCGGCATATTTCCAGTCGACGTCGCACAGGCCGATGATGTTCTGGCTTTCGAGGCCGCGGAGCACC
GACGAGCCGCGGCCGCCGATACCGACACCCAAGATGTTGAGTTTGTCGCTCGGAGCCGTGTGCCCGAACTTCTTACCTAAAACCGTACCCGGAACGACCGC
TAAGCCGATCGCGGCAGCAGCTCCTTTTTTCAGGAACTTTCTCCTTGACATTTCGCTTGACATTGTTTCTTGATTTAAGTTAGTTGTTTTTGAGGGACTATTTATT
TTCGTTGTTGGATTTTTCCGTACCCGAGTTTTCGATGTCGTTCTCGATGTTGTTCATGCCTTCCTTGAAGCTGCGCACACCCTTGCCGAGACCCTTCATCAGTT
CCGGGATTTTTTTGCCGCCGAAGAGCAGCAGTACGATCAGGGCTATGACGATGATTTCGCCTGCACCCAGGTTGCCGATAAACAGTAAGTGGTTCATACCGA
TTGTTTTTAAGGTTTATAAAGTTTGTTTTAGTATCAGATTGCGGCTTATGCGCGCTTCCTCAGCAACCCGTAGCCATTCGACATCGTGCGGCGGCGTTCGCGAA
TCTCCTGCAGCGTTTCCAGATTTCCGATTGCCGGCAGGTCGTGCTGCCGCGAATCCTGGAGGAACCGCCATGCGTATTCCGATGCCGTGGCGGCGTCGCGG
AACGAGGGGAGCGACGGATTGCGTTTCCCGCTTTCGATCGACTCGGCGAACAGTTTGCACAGCACGTCGATGTTTTTGCCTCCGTAGGGCTGTATGAGCCG
GTGCGTCTGTGTCACTCCGTGCAGTTCGACGATCGCCGTCTTGAAATCGTGCGTCATGCGCGCGATGCCTTTCGTGCCGATGATGTCGATGTATGAATTATGT
GTCTGTTTTTCGGCCAGTTGTCCGTATACATGTCCCTGGGTAATGTCGAATACGATGCCGTTCTCGAACGTGCCGTGGCACTGGAGCCACCACGGATCCTTGT
AATCCCACATGCGGATGGCCTGTGCGTTCCAGGTCTTGAATTCCGATTCGGCATACCAGCGTGCGATGTCCACGTAGTGCATGCCGCAGTCATGGAACGACG
GGCCCTCGAATTCGTGTCCTTCGCCCGGGGCCAGCCCCGGCGTCATGTGGCATACGCGGATGATCGCCAGTTCGCCGATTTCGCCCTGTGCGATAAAATCC
TTGATGGTATTGTGGTACCACGAATTGCGCAGGTAAAGGTTTACCGTCGAGAAGATTTTGGCCTTCTCGGCCAGCGCGACCGTCTGCCACTCCTTTTCCACCG
TGTCGGCGATCGGTTTCTCGGCTATGACGTGCTTCCCGCTCGCAATGGCTTTCTGAATCTGCTCCGGGCGGGAATCCGCCAGGGCGAACAGCCCGACCGCC
TGAATCTCGGGGTCGTTGAATATGATTTGTTCGTCGGATATTACTTTCGACCGGGGGGAGAGCCGCTGGGCCAGTTCCCGTGATGCGGGAGATACGTCACAG
ATGTAAACGATATCCCAGTTACCGCTTTTTTGCATCTCTTCGAGGTAAAAACCTCCCATTCTGCCAAAACCGATCAGACCGACTTTAATTGGATTCATTTCGTAT
AAAATAGGTTTAAGTAGGTGTTTACAATCCGATGCAATCCGGCTGCCCGCTGCGGGAGAGGGTGGCTGGACGCTGTGTTTACAAAATTAATCCTATTATATAC
CATATTTATAGGTTGGGGCGGAAATTGTAGAACCTGTAAAGTATTATATGTTTGTAAATAAATATTTTATCAAAAAAATATACAAAAAAAATGTAGTTGCCGTGGA
ATCGGTCACGGCCTGTTTTCGGGCGGCACACCGATTCGCATGACCATTTTTTCGAAGTCCTCGCGCGATATTGCGGCCTTGTTCCGGGCTTTGGTGCGGTAG
TTATATACGGTGCTCAGCGAGCATCGCAGGAACGAGGCGATCTTGACGCTGTCGGTGATGCCCAGCCGCAGCAGGGCGTAAATCCGGAGTTCCTTGTTCAG
CAGGTCTTCCGATTTGAGCACGATCCGTTCGTCCTCGACCAGCAGGGCGTTGAAATCGGCGACGAACGTCGGATAGAGGTTCAGGAATATGGCGTCGAAATT
CTTATACAGTTCATCCAGTTCGTTCTCCATCATCGACGTGGATTTCAACTGTTTGTATAACTCGTCGATCTGTCGGTTCTGCGCCAGTTTGTTCAGCGATTTCCG
GTAGTCCTCCAGTTTGTCGATATACATCGAACAGAGGTCGAAGAACCTAGCTATGTACTGTTCCTTGATGATGTTCGAATCGGTGAGCTGCGTGTTTTTCTCAT
GCAGTTCGTCGTTGAGCTGCATCAGGCGCCCGTTCGTCCGCGACAGCTCCTCCTTGATGCGGTACAGCTTGTGTATCTGCTGGTAAACGTAGATGAACAGCA
GCACGAGGAACAGCGACAGGACGCTGATCAGGATCAGGTAGTGCTGCAACTTGGATTTGGACTTGGCCTCCTTGGCCTGGTAGATCGAGTTGATGATCGAAT
AGAACTTTGACATCTGGGTCGTGCGGAACTGCACGTTGCTGAATACCGCATCTTCGATGGCCGCCTGCGTGTACTTGAAAGCCCGTGAAAGGTCGCCCTGGT
CGTAATACATCAGAGCCAGCGTGTGGAACGAGGCGTTCTCCTTCACCGCACTCTTCGCATCGGCGATCGCGGAGAGCATGAAATATTTCTTTCCAGCCTCCG
GGTCTTTGTTCAGCCGGTAGAAATAACCGTAGGCATAGGCCGTCTCCGCGTAGAGCGGCGTGTCCTTGCCGACCTCTTCCAGCAACTCGTCGAGGATCGCC
CTGGCTTCGTCCATCTTACCGTTTCCCATAAGTTTGTAGCTCAGGTGTATCCTGTACCGGAACGATCCCGGGTCGGAAACCATCATCAGCGAGTCCCGGTAGA
TATTCATTTCGTGCAGGTGGCTGCGCTGGCTGCTCAGCGTGGCGTAGTGCTGGAAATAGCGGTTGTTGGCCTCGTAGTAGACGGGAAGCAGGTCGCGCGGG
AGTTCGGACGATCGGATGGAGCGGAGTATCTGGTTCGCCTCGACGCACATGCCCGAAAAAGAGTAGAGCTGGGCGAGGTACATGTTCGACAGGTAGGTCAG
CCTTTTGTTGTCCAGCTGCCTGGCGATTTCGAGGTTCCGTTCGACGTAGCGGATGGCCGAATCCAGCTTGTATTTCCTGAATTCTTCGTACAGCTTCGTATTGA
TCTCGTACTCCTGTTCCGGGGTCAGGTTGCTTACGCCCAGCAGGTTCCTTAGCCTGCTGATCCGCTCTTCCTTTTCGGCTTCGAAAATATGTTTGTTCTCGATT
ACGTAGTCCAGATATTCCGAGAGGGATTGCGTCTCGTCCTGCCGGCACGACGCTAATGCCAGCAAGCCGGAGAGCAGCAGGAGTAGCCTTATCGGACGTAT
GGAATTTCGTAGTTTCATGGCCTCTTCGTATGCAGCCTTCCGTAGCAAAAACGGCAGGTTGCGGGCTGCCGTAAAAAAATGATATTTTATGCTACGAAGTTAAG
GCTATTTATAGGATAAATATTGTTTCTTTTTATGTATTTATTGTCTGATATTCGTATTTTTTAGGGTGTCTTTGGCGATTGCTGCGCTTCGGGAGCCGGGGATATT
CCGGAGGCTTCTTTTCTGGGACATAATGCCGGATTTGTGAATCAGGCATCTGTAATCCGTTGTGTCATTGTCTGAAACATTTTTTTATGGACTTCCAAATATGGT
AACATGCCACAGGTGTTTCCCTCTTCCGCTTCCATTGCAGGCATCTGGCTGTGGGACACGGTATTGTCCGGGATTTTCCGCCGATGGGGCCGGCCCGTAACA
ACGATCCGATATTACGGAAACGTTTTTTACACCGGCAAAGTGTGTCCCCGCGGACTGATTTGCGCGAAAGCTGTTTCGCCCCTCCCCTCTCTTTCGGACGGGA
ATGGCGCAAAAACGATAAATTTTGCTACCTTTGTCGCTTGTATGAAGGCATCGCTCAAAAAGACTTATGCTTCGCTCGGCGCCGACGGCCTGGTAGCTTTGTG
GCTCGGCGTGTGGTGGGTGTGCAACCTCCTGCAGGCCGGGTTTTCCGAACTGGCCAACGACGAAGCGTATTACCATATGTTCGCTGAAAACCTCGCATGGG
GATATTTCGACCATCCGCCGATGACCGCCTTGCTGGTCTGGCTCGGCGAGCACCTTTTCGGGGGCGAACTGGGCGTGCGGTTTTTCTTTACGGTGCTGCAAC
CGCTCTACCTTTATATACTATGGCGGATAATCCGCCCCGCGGATGCCAACCGGCGCGACGCGGCGCTTTTTGTCGTCCTCTCGGCCGCGACGCTGATGTTGC
AGCTCTACGGCTTCATCGCCGTGCCCGACGGCCCGCTGATGATGACCACCGCGCTGTTCCTGCAGACATTCAAGTGGTTCTCCGAGAACCGGCGTTGTGCA

TGGCTCTGGATGGGGGTTGCCATGGCGCTGATGGCTTACAGCAAATACCACGGGGCGCTGGTCGTACTCTTCGCGCTTGCGGCGACCCCCCCCCGTGTGTT
CCTGCGTCCGACCCTATATCTCAGCGGCGCGGTAGCGCTGTTGCTGCTCGTGCCGCATTTCGTGTGGCAGTACGAGCACGACTGGGCCTCGCTGGCCTATC
ATCTGGCGGGGCGTAATTCGGTGTTCCGGCCCAATTATGTCGCCGAATACCTGCTCAACCTGCTGGTGGTCTTCAATCCCTTCTTCGTGCCGTTGTACGTGAA
GGCCTGGATCGCCGTGAAGCCGCAGAACGCCGTCGAACGTGCGCTGAAGTTTATCCCCGTGGCGTTCATCGTTTTTTTCCTGCTCTCGACATTCCGCGGTTA
CGTGCAACCCCAATGGGTGATCGTCGCCGTTTTCGGCCTGATCTATACGCTTTTCACCTATGCCCGCCGGCATTCCTGCACGCGGCGTTACCTGATGTGCGC
CGGGCTGGCGACCGTTTTGCTGGTCGGTCTGGTACGTCTCGAAATGATCTTCAATCCGCTGGGCCTGCGTTTCGAGGTCTTCGACAATCCTGCGAGCTACGG
CCGGATCGCCGGGATCGCCGACGGCCGCCCCGTCGTTTTCCGCCATGGCTATGCCGTCGCGGCCAAATACGCCTTCTATACCGGGGGCGAGGCCTACTGC
CAGCCCAATATCCGCTACCGTACGCACCAGTGGCAGTTCCGCGATGACGATACGCGCTTCGCGGGACACGAGGTGCTGGTCGAAACGCCTTACGACGGGG
CGGATACCACGGGCCGCGTCCGGAGCGTCGTGTTGCCCAACGGCAAACGTTTTTCGTGGTTCGTCGACCCGCATTTTCATCCGACGCGCCTGGTGGATATT
GCCTTTACGGGGCTGCCCGAACAGGTCGTTGCGGGCGATACGCTGCGTTTGACGCTGCGCATCTCGAACCCCTATCCCTATGCGATCGGGGTCTGCGGGGA
TCGGTCGCTCGTGATGCTCTGGAAGCACGGCCGCTTCCGCGTCGAGGAATTCGATACGGGGGAGCATTTCAATATCCCGGCCGGGGATACCATCACGCGCG
AGGTGCGGTTCGTCGTCCCGGAGCAACTCTCCGGCGCCGGTTTCGATGCCGGTTTCGCCCTGCGCCGCGATGGGTATACGAATTGGTTTAACGGAAACCT
CTCCGCACGGAGGTCGTCGCACCATGATCCAGCAGTTCATCAAATTCTGTGTCGTCGGCGGTTCGGGGGTCTTCGTTGATTTCGGGATCACTTACCTTTGCAA
GGAGTGGCTGCGCCTGAACAAGTACCTGGCCAATTCGCTGGGGTTCCTCTGTGCGTCGACCTCGAACTATGTCCTGAACCGCATCTGGACGTTCCATAATGA
AAACCCCGACATTGCCGGGCAATACCTCCGTTTCCTCGGTATCGCAGCCGTCGGGCTCGTGATCAACAACCTCACGATCTATCTGCTGCACGAACGCTGGAA
GCTGAATTTCTACCTGGCTAAATTATTTGCCATCGGGGTAGTCACCTTCTGGAATTTCTTCATGAACTATTTCTTTACGTTTTGATTTCCGGGGTGCTCGGATTC
CGCTGTAAAGTATGTATGCATAAATGCCAGCCTCCCGGGAGGCTGGCGTCCGTATTCCTGCCGGATGGCTGTCACTGCCTTTTCGCGCTTCTGATCGAACGG
AACGGGAGTCCCAGCACGCCGAAGAACGCTTCGCGGAAGATGCCGCCGCTCATTTTCGAGGTGCCTTCGCGGCGTTCGACGAAGATGATCGACACCTCCTT
GAGTTTGAGTCCCAGCCGCCATGCCGAGTATTTCATCTCGATCTGGAACCCGTATCCCTTCATCTTCACGGCGTCGAGGTCGATCGTCTCCAGCGCGTGGCG
CGAGTAGCAGACGAACCCCGCCGTTGCGTCGCGCAGGGGCATGCGGGTCACGATGCGTACGTACATCGAGGCAAAGTAGGACATCAGCAGGCGCGACATC
GGCCAGTTCACGACGTTCACACCCTGCACGTAACGCGAACCTACCACTACGTCGTTCCCTTCGGCCGCAGCCTGGTACAGCCGTAGCAGATCGTCGGGATT
GTGCGAGAAGTCGCAGTCCATTTCGCAGATGTAGTCGAACCCGTTGTCCAGTCCCCAGCGGAACCCCGTCAGATAGGCCGTGCCCAGCCCCTGCTTGCCGC
TGCGTTCCAGCAGGTGCAGCGTCGCGGGGAATTCCTGCTGGCGCTCTTTGACGATTGCTGCGGTGCCGTCGGGCGAACCGTCGTCGATGACGAGCATTTCG
AAAGCCTCGGGCAGCGAGAAGACCTTGTCGATCATCGCCGAAATATTCTCTTTCTCGTTGTATGTCGGGATAATTACCAGTTTGCGCATAGGGGCTCGTTTTG
GAATTTCTTACACAAAGGTAACATTATTTTGCAAATCTGTGCCGTGCGCCCTGTTTTTCTGCCGTGCGCCCTGTTTTTCTGCCGGGCGCCGTCGCCGGGCTGC
CGTTTTACCGTCTGTCGGTTGTTTTGCGGCCTGTCCAGGCCGTTTTTCCGCCTGCCCGGCCGTTTCGGTGCCGGGGCGCGGGTTCACGGCGGCCCGTGAAG
TTGCCTGTTTGCGCCGTTTCATGGCGCGGATCGCCTGTTTGTCCGTGTCGCTCACGCGCAACGATTCGACGATTTTTTGCAGCGATTTGTTGAATGTCCAGTC
GTCGAGCGAGCAGCTGCCGAGCCACTCGTGTGTCCGCTGCGGGAATTTTACATAGCAGACCGACACCGCCCATGCCACGGCCATACGGGCGTAGTAGGCTT
CGTGGCGGACGCCGTCGAGCAGTTGCAGCAGTGCCTCGAGGTGCTCCCCGTCGACGAAATTCCCCAGCCCCGTCACCACGGCGAAACGCACGTCGTATTC
GGCTTCGGATCGGAAATAGGGCTGTATGAATTCCCACATCGGCTGCCGTTCGGCCGCCTTCAGGCGCCAGCAGAAACAGTCGCAGACCGCCCAGTTGTCGA
TTTTGGGCACGAAGCGCGCCACATGCTCCAGCTTCTCGGCCGGCGTGCAGCGGGCGCAGCCGATCACCAGCCCCTGCAACATCCGCTCCTCGAAATAGAAA
TCCTCGGCTTCGGCCAGGTATGTGCGCCAGTCGCCTGCGGCGATCTGGCGGGCTATTTTGCGCAGTACCGGCAGGCGTATGCCCAGTACGTTTTCCACGCC
CGGCATCAGTGCCGCGGTGAATTTCATGTATTTCGGCTCCTGGAGTTCGAGCAGTTGGTCGCGGAGGGTCATCGTTCCCGGAATTTGGAATTTTGCCGCCCC
GAATAGGGCGGATTGCCGTGTGTGGTTATTGTTTTTTCGTTACGGTGCGGTAGCAGCCGATGCCGCTCACCTCGGTCGATACCTCGCCGATCAGGGGCGCTT
TGGCATTCACCCCTGTCTGCACCATGATGAAGTCGACCTGTGCGAGGTTGGCCGGGGCGCCATCGGCCGTTACGGCGTCGCTGATCCGCAAGCGGTTGGTC
ATGCCGGTGCGGTCGGAGGTCGAATAGTTGTCGGCGTAACCCCATGCAAAAGGCTCCATCGTCCATTTTTCCCCGGCTTTTCCGATATTGTCCGGCAGCCGT
GTCCCTTCGAATGTGAATCGCTCCCCGAGCCATTCGGGGTAGTAGGCCTGCGTATGTTCTTTCACACGGTCGATCGTGCCGCTGCCGCCCGCGTTGTCACTC
CAGGCTACCGGGGCGTCCGCCTCCGCGGGGCGGGTATAGGTGATCTTGTAGCCGCGGATCGTTTCGGCGTTGTCGTATTCGCTGCCCCGCAGTTCGTACCA
CGTATCGTTGGGCGCTCCGTCCCCGTTGGCGTCCTGCGACACCCACACGATGCCCGGTTCGGACGACGTGTCGAACTGGTTGCCTTTGACGTAGAGGTCGT
ATCCGCCCGTGTTGGGCACGGGTTCCGCGAAGGCCGCCACGATGCGGCCTCCCCAGCCGCCGAGCGAAACCCAGTTGTTCGAGGCGAGGCGGTCGGCGG
CATAGCGCAGGGCGTCTGCCTCCGATGCCACATGGTCGAACCCGGCCATCGTGCCGTTGATGAACTGGCCCGGTGCCGGGGTGTATTCGAGGATCGTGGC
GACGGGCTCCTGGATCTGCGTGGTCGTGGTCTCTTCGCCGGAATTGCAGGCGGCGAAAAGAAGCGCCGCGAGCGGGAATGCATATTTATTCATGTTTTCCAT
AATTTTTACGGTTTCGGCCGGGATTCGGCCCGGGTATTCCCCGGGCTTGCCGGCGCCCGGGCTCCTGCCCCCCCCCCTTCCCGGGCGTTTTCTCCGCGCCG
GAGGGGGGCCGTCCGCAACTGCACCGGTCGCCCGGCGTACGATCCGCCGCAGGGTGGCACGGGGCCGCGTCGGGGTGTCGGTTTGCCTGCCTGTCCCGA
ACGGGACAAAAATACTCTTTTTTCGGCATATGGCAGCCGGGATACGGGCAAAAAGCCGGGGTTTCGAAAAAAAGATTAACTTTGTGAAAAATATCCGGGAAGT
ATGGCGGTCGAAATAGAGTGTGTGACAGAGGTATCCGATGAACTCGCCGAGGCGTTCGCAAGGCTCATGCCTCAGCTTTCGCCCCGGCTGGGGGCACTCTC
CCGCGAGGCTATGGAACGTGTTGTGTGCAGCGATGGCGCGGCGCTGTTCGCCGCCCGGCTGGATGGCCGGATCGCCGGGGTGCTGACGCTGGTGTGGTA
CGACGTACCCTCGGGCCGGAAGGCATGGGTCGAGGATGTGGTGGTCGATGCCGCAGCCCGCGGCTGCGGTGCCGGTGAGGCGCTGGTGAAGGCTGCCAC
GGAGCATGCCGCCCGCATCGGTGCTGCGAAGGTGATGCTCACCTCGAACCCTGCGCGCGAAGCTGCCCGGGGCCTTTACCGTAAAATCGGATTCGAAGAG
GCGGGGACTACGGTTTTCGTCTTTAAAACAGATAGGAAATGAAAAAAATCGTGAAGATTGTGGCCATTGTCGTGGCCGTTGTTCTGGTGGCGGCTTTTGTCGC
CCCGATGTTGCTGAGGGGGAAGATCGCCCAGATCGTGAAGCGCGAAGCCAATGAGATGCTCGACGCACGGGTCGATTTCGAAAAACTCGACATCAGCCTGC
TGCGCCATTTCCCGCGCGCGTCGCTCGACCTGAAAGGGCTGACCGTCGTGGGTGCGGGGCCGTTCGAAGGCGATACGATCGTTGCCGCGAACCGCATATC
GGTCGTCGTGAACCCGATGTCGCTGTTCGGCGACGACGGGTTCGAAGTGACGAAGGTCATCCTCGCCGAGCCTGCGCTGCACGCGCATAAACTCGCCGAC
GGCAAGGTCAACTGGGACATCGTGAAGGCTTCGGGGGAGGAACCCGCCGCCGGGGAAGTGTCTTCGGAGGAAGGTTCTTCGTCGTTCAGCCTCTCGATTCG
CGATTTCAGGATCAGCGACGCGACGATCCGCTATGAAGACGATTCGACGGCATGCGTTTTTCGACGGCACCCCTCTCCCTTCGCCTGCGGGGCGACATGT
CGGCCGACCAGACCGACCTCGACCTGCGGCTTACGGCCGGTGCCGTGCGGCTGGTTTCGGGCGGAATTCCGCTGCTGAGCGACGCCGAGGCGGAGCTCG
ACGCCGTGATCGCGGCCGACCTGAAGAACAACCGTTTTACCTTCTCCCGGAACCGTCTGCGCCTCAATGCGATCGAGCTTACGCTCGACGGCTGGGCCGAA
ATGAAGGACGATGCCGTGGCTATGGATATTACGGCCGGATGCGACAAGGTGCAGTTCAAGGACGTGCTTTCGCTTGTCCCGGCATTCTATACCCGCGATTTC
CGCAACCTGTCCGCCTCCGGCGAACTTTCGATGGCGCTGTGGGCCCGTGGCGAGATGCGCGGGGCGGCGCTCCCGGCCTTCGAACTCAAGACCGAGGTG
CGCAACGGCAGTTTCCAGTATTCGTCTTTGCCCAAGGCGGTCGACGGAATCAATATCGCCGCACGGATAGCCAACCCCGGCGGCGTCATGGACAAGACCGA
GGTAGACCTTTCGAAATTCACGCTCCGCATGGCGGAAACACGCTTTCGGCATCGCTCTATGCGACGAACCTCGCCAGCGATCCGGTATTCCGCGCTGCGG
CCGACGGACGGGTAGACCTGGGGGCGGTGAAAGAGGTTTACCCGCTCGGTGAGGATGTAGCCCTGAGCGGACTGATTAGCGCCGACGTGAAGGTTTCGGG
GCGCATGTCCGATGTCGAGAAGGCCCGTTACGGGCAGATCGGCGCCTCGGGGACGTTCGTCGTCGAGAAACTCGGCCTCTCCATGCCCGGCCTGCCCGCG
GTGCATATCCGCCGTGCCGCGGCTACGATCACCCCGGCGTCGATGACGCTGGGCGAGTTCGGCGTGACCGTGGGTAAGAGCGACCTTGCGGCCAACGGCC
AGCTTACGGGGTATATCGGCTACCTGCTGCGGGGCGACAAGCTCTCCGGGCGCCTTTACGTGAAATCCGACCTGCTCGACCTGAACGAGATCATGAACGCC
ATGCCGGCCGACGAAGAAACCGCCGGCGGGGAAGCTGCTGCCCAGGCGCCCGCCGAGTCCCCTGCTCCCGCGCAGGCGCTCGAAGTCCCGCGCAACCTC
GACCTTTCGCTCAATACCGAGTTGCAGAAGGTGTTGTTCCAGAAGATGACCATCGGCGGCATCACCGGCGAAATGCGCATGGCGGACGGAACTTTGTCGCTC
AGCCGCCTGCGTATGCAGCTCTTCGGAGGTACGGCTACCGCATCGGGCAGCTACTCGACGGCCTCCGATCCGCAGCGCCCGGCGCTGAAACTCTCGTTGG
GGCTTTCGGGGGCATCGTTCTCCAAGACCTTCGACGAGTTGGAGATGGTGCAGAAACTCGTCCCGGTCTTTGCCAAGACGGGCGGCGACTATTCGCTTTCG
CTCGATATGTCCGCGACGCTCGATGCGCAGATGTCGCCCGACCTGCAGTCGGTCAACGCCACGGGCGAGATCAAGTCGGCCAATATCCGCATCCAGAACAT
TGAGGCGTTCGATGCACTGGCCAAGGCGCTCAACAACGACAACCTGCGTAAGATCGAGGCCAAGGACGTGGCTATCCGCTTCGCTATCCGCGACGGACGTA
TCGCTACGGAGCCTTTCGACCTGAAGATGGGCGACATCAGGATCAACATGTCGGGTTCGACGGGGCTCGACCAGACCATCGACTATACGGCCAGGGTAGCC
CTCCCGGCCGGTTCGACGGGCGGTATCCTGCAATCGGTCAACGTCGGTATCGGAGGCACGTTCACCTCGCCCAAGATCACGCTCGGTGTGAAGGAGGCTG
CCGAGCAAGCCGTGAAGAACGTCGTCGACCAGCAGATCCAGAAGCTTACGGGCAGCGAGTCGCTCGGCGAGGAGATCCGCAAACAGGCCGACAACCTCCG
GGCCGAGGCCAGGAAGGCGGGTGAGAAACTCGTCGAGGCCGCCCAGGCGCAACGGACGAAGCTTATCGACGGTGCGAAGGAGAAAGGTGCGCTGGCCAA
GTTGGCTGCCGAGAAGGCCGGAGACAAACTGGTCGAGGAAGCCCGCAAACAGGCCGACAAACTTGCGGCCGAGGCCGATAAACAGATCGAAAAACTCACG
GCGAAGCAGGAATAAACGATGCTTGTCATTTTTGCCGTCATCATCGGAGGAATCATTGCGGGGCGCCTGCTGTCGAGCTGGCGCCTCGTATTCGTCTCCCGT
ATGATTACCGTGATTATCTGGCTCCTGCTGTTCCTGCTGGGACTGGAAGTCGGTTCCGACCCTGCCGTCGTGGGTGGTATGGCGACCCTCGGACGCACGGC
CTTCGTCATCTTCGCATGCTCGGTCGCGGGCAGCATCTGTATGTCGTGGCTGTTGTGGCGCTGCGTCCGGCGCCGGGACGCTGTGCCCGACGGAGGCGGT
ACGGAATCGCCGGGACTGCCTGCCGCGGCTGCGGATTTCCCGCGAGGGGCGGCAGGCGGCCAGGCCGCAGCTGGCGGGGACACCCGACCCCTTTCCGTA
TGGGGCGCCTTGCGGGGGAGCCTCGTGATCGTCGCCTTTTTCGTCGCAGGGTGTCTGGCGGGGTTGTTCTCCGCCTTGCCGTTCGACGTGGCGGGGTCGC
GCCTCAGCACCTATGTGCTCTATGCCCTGATGTTTTGCGTGGGGATCACCCTCGGCAGCGATACGGCCCTTGCGGGGCGTGTGCGGCGGCTCGATCCGCG
GCTGGCGCTGCTGCCCGTGATGACGGCCGTCGGTACGCTGGCCGGCGCTGCCTTGTCGACACTGCTCTTGCCGCCGCTTACGGCCGGGGACACGATGGCC
GTCGGCGCTGGGTTCGGCTACTACTCGCTTTCGAGTATCTTCATCGCCGATTTCCGGGGCGCGGAGCTGGGAACGGTGGCATTGCTCTGCAATGTCATGCG
CGAGCTGTTCACACTGTTGGCCGCACCCCTTGTCGCCCGGTGGTTCGGGCCGCTGGCCGCCGTCTCGATCGGCGGAGCCACGACTTTCGATACCACCCTGC
CGATCATCACGCAGTCCGCAGGGAAACCCTATGCCGTGGTGTCGATCTTCCACGGCTGCGTGCTCGATTTCAGCGTCCCGTTCCTGGTGACGTTCTTCTGCA
CGATGTAAATTCGGATACGGATGCCCTTCCGGAGCGCTTTTGCCGCTGTCACGGCAAAAAGTCCCCGCCACAGCGGGCTGGGATGGACGGCAATAAACACG
GCGGAACGCCGTGGACGTTTTGTGGTTATCTTTTTATGTAGGTCTGCAATGCCGCGACGTATTCGGCGAAGGCATCGCGCCCGGTGTCCGTGATGCGGCAG
ACCGTGCAGGGCATCTTCCCGCGGAAGGTCTTTTCGATTTCGATATAATCCGCCTTCGCCAGCTTGTCCAGTTGTACCGAAAGGTTGCCCGCCGTGGCCCCT
GTCTGCTGCCGGATAAAGACGAAGTCGGCGCTCTCGACCCCGATGAGGATCGACATCACGGCCAGTCTCAGTTCCGAATGCAACAGGGGATTCAGTTCTTTG

AACATTGCTTGCGGGTTATGTGGCGGAGGATAAAACCGGGTAGCAGTACGAGCAATATGCCATGGGCGGAGAGTTCGAGCATCCACGTTCCGCCCTGGGCG
TCGAACGGCATGCGGAACGTATAGTTGAACGCCATGGTGAGCTCGAAGACCGAGAGCATGAGCAGCGCCGCGAACGACCCCGATTGCTGGGAACCTTTGCC
CCGGTAGAATTCGGCGGTGATGGCCAGTGCCGTGCCGAGCAGCAGTTGCATGACGCCTACGGGGTTTATCGTGTTTCCGGGGATGGTTACGGCATAGACGA
TCGCGGCGGCCGAGATCATCGCCGGCATGCCCCATATCAGGATCAGCAGGTCGTCGAACCCCGTGCGGACGAGGGCCTTCCGCCGCGAATTGCGAACCGT
CCAAATATAGCCGCAGACCAGCGGAACCAGCCAGCTCCACAGCACGCGCACGTCGCCTGTCCAGCGGTAAAGCCCGTATTCGGCCAGTGCCGTGAGGCTG
AGCAGCGCTCCCCAGCAGATGAACAATTCGCCCCGTACGTAATTCATGCGGCGCATATTGCGGTCGAAGACGCGGGCGATGATGTCCAGCGCTTCCGTTTC
GGATATTTTCTGCCTGTCCATTGTGTTATTTATATCGTATGGTTCGACTTGTAGTTGAGGATATGCCCCGGGATAATCATCATCACGACGAATCCGGCGGCGAA
GAGGGCCGGCATCCAGTAACCGCTCACGATGAGCGAAACGGGAGCGAGGACGATAGCCGCCGTACCGCCGGCGGTGGCCGGGCGGAACCGGATGATCAG
CCCCGTGATCGTGGTTCCCATGCCCATCGTCAGCAGGATGATGAACAGGATCGGCAGCCGTATCTGGCTGAAAAGCGAGAGCATGCTGACGAACCACGCCG
TGAGGCCCATTACCAGCCATACGTTGTTGATGACGCGGTCGACGAACGTGCTGACTTTGCCTTCGGTATGCTTGTCGCGTGTCAGCCACATCAGCAATGCGC
CCAGTACGGGGAGCAGCATCCACAGGATGTTCCATTTCGGATCCTGGAAATAATACACCGCACCCATGACCAGGAGCGTCGTGAAGACGGTCACGTAACCCC
AGACCAGGAACGGCCGTCCCGAGTTGCGCAGCATACGGTTGCGGGTGTTGTCGATCATGCGGCTGATGAGCGCGATGCTCTCGGCGGCATTCAGTTGCTTG
TCTTCCATAGCGATGTGTTTTTGCGTCCGTTTCATGTCGTTTCAGTGGGCGCCGGGTGTTTTGCCGCCACCCTTGCCGGCACGGTACTGCCCGGAATGTTTAT
AACGGTAATGGAGCAGGTGCCCCGGGATGATCAGCATCAGGACGAATCCCGCCGTGAATAATACGAGGTGCCAGTGGTCGGTAACCGCGAGCGAAACCGAT
CCCAGCACGATCCCCGCCGTGCCTCCGGCGATGCCGGGCGTGAACCGCAGAATCAGGCTGGTGATGGCGATTCCCATGCCGATCATCAGCAGGATGGTGA
ACGTTACCGGAATCCGGATGATGAGCAGCACCGCCAGCGAGCCGACCAGCAGGGCGGCGAACCCGATGACCTTCCAGATTATGTCTATCACCTGGTCGACG
TAAGTGCGGGAGCAGGATCCGGTCGTTTCGGATCCGAGCAGGCGGGTTATGAGGTAGCCGGCTACCGGCAGCATGAACCAAAGGTAGTTCCAGTGTGTGTT
GTCGAAATAGCTTGCCAGCCCCCAGACGGCGAGTGTCGTGGCGACCGTAACATAACCCCATGTCAGCAGGTGGAGCCCGGCGCTGCCGGCCATGCGGTTG
CGTGTATTTTCGATCATGCGGCCGATGAGTGCCAGACTCTCGGCGGCGTTCAGTTTCTTATCTTCAGTCATGCGCGTCGTTTGATTTCGCCACTTTGTCCGCA
TCGCACCTTACCTGCCTTCTGAGCGCGTAGCCGGCACCCGTGAGCCCGATGAGAACCGCTGCGGCGAATTGCAGCAGGTTGTAGTCGGGGTCGGCATTGAG
CCAAGAGTCCCGGGTAATACCGAAGCCGATCGGTATTACGGCGCATGATGCGTAATACATGTACTTGTTGCGGATTACCTGCCCCGAGATGAAGAGTCCGGC
GCCGATCAGCAGCAGCGCAGGGAAGAGCGGGTCGACCGCCCACGGGTCGTACCCGCAAATCAGGATCGCCGACAGGACGACCAGTGAGGCTATGCAGCT
CCATATCTGTTCCAGCAGCCGGTCGGTGTAGCTTTTGACCTGCGGGGCGCGGAACCTGTTATGCCACCAAGTCACGGCGCCCCCTGTTGCCGCGGCTGGCA
GCCACAACCATAGCCAGGTGCGGCAGTCGGTGGTTTTCAGCAGGACGAAGATCACGATGGCGGTCGCGGCCAGCAGGTATCCCCACAGGAGGAACAGGTT
GCTGCTGCCGAATGAGAGCCGTTGGCGGGTGCCGCGGATCATCTGCTGGATGAGTTCGATGCGCTGGTCTGCGTCGAGGTTTTTTTCTTCCATGGCGGATG
GTGGGTTACTGGGGATAAAGTTACGAATTATTTTTTATTGTTCCGTGCTTTTCACGAGGCGCTTGCGCTGGGTTTCCGCCGCGCTTTCGACGGTGCGGGCGC
GGAATTTGTTGCCGGTGCGGAAATTCCAGGCCGCGCCGACTTTCCACAAGCGCCCCGACCAGGCTTGCCGTGTGTCGATCGTGCGCCGCATGCCGTCGGTC
AGGGAGGCGAACTCCCAGTCGCAGCCGGTGATGTTCGACAGGGTGCAGAAAAGCGTCAGGCGTTTGCCGAAGAGCTGTTTTTTGACGGTCGCCTGGAGCGT
GTGGCGCGGCCCGACCTCGCTGTTGCCCGAGTAGAGGCGGCTTTGGGCACGGTAGGCGGCTTCGGCGAAGAACCCGGCCGGCAGCGTGAACGAGCAGGT
GGCGTCGGCGAACATCAGGTAGTGCGTGGCCGGGCTGTCGGTGCTGTTGAGCCGGATGCGCTGCATCACCCCGACGGCATTGACCGAGAGGTTCCACCAG
CGGGTGATTTTCGCCGGGACGCTCGCCGCCACGAACCAGTGGTTCTCCGTGTAATGGTTTTCGGGACGGATATACACGGCGTCCGGGTCGGTTGCGTCCCT
GTGTGCGATTTCGCGAATCAGGTCGTGGTGCAGGTTGCCGCCCACGGTGAGCGTGTAACGGTATTTCCATACGGCCGTGAGCGAAAAACGGTGTATGTAGG
TCGGCCGCAGCGAGGGGTTGCCCGACTGCCAGCTGTAATCCGATATGCGGATCCGTGCGGGGTTCAGCGCCGGGAATGCCGGGCGTTCGATGTTGCGGCT
CCACTGCGCCACGAGCAGCCATGTGCGCAGGGCGTTGAGCGCGCGGCTGACCGAAACGTTTGGGAACAACCCGAAATAGGAGCGGCCGAAGGCCTGCGAT
GCGACGGAGGTGTATTCACCGCGCAGCCCGGCCGAAAGCTCCCACCGCCCGGCGCTGTACTCCAGCGAGGCATAGGCCGCGCCGATCTGCTCGGTGTAGT
GCTGGTCGTAGCCGTATTCGGGCAGTGTCTGCCATGTGCCCTGCGTCTGTGCCGCGTAGCGGCTTTCGTCCGCCATGCCGTTGCGGGTGTAGCGCAGCCCC
GCGCGCAGTGTCAGGCCGTGCGGCAGTTTGCGCGTCAGCCCGGCGTCGGCCGTCAGGATGTCGTACACGGAGCCCGTGGAGGATTCGTAGAGCGAGTCG
CGTACGGTTCCGGGTGCGGAGGTCGTCGTGCGATAGCTGTTGTCGCCGTCGGAGGTGTAACGCGTATAGTCGGCGATCAGTTTGAACTGCGAACCCAGCGT
GTCGATCTTCCAGATGTAATTGGCGGTCGCCGTCAGGGTATTCCCGCCCATGTGCTGGCGGTAGCGGCTGTCGCTCTGAGTTTGTCCGAGTGTGGTGCGCG
ACAGTGTCGGCATGTCGGTACTGCGCTCTGTGTATTCGACGTCGAAGCCTGCGGTGTGCTTCGGCGTCGGGTCGAAAACCGCGGAGAAACGTCCGCGCCCG
TAGTTTGTACGGCTCTTCGAATCGCTCCTGCCGGCGAAGGGCAGCCGTTGCCTGGCCTGTTTGCGGGTCTCCGTGAAGCGGCTTTCGGCAACCGGAGTGAA
GCTTCCCGACCCCGAGGCGGTGAGTGTCCAGCGCCCCGTGCGGATGCCGATGCGCCCCGTGGGGGTATAGCCTGCTATCCGGTTGCTTTGCACCGTGGAG
AATTGCAGGTTGCCGTCCATGCCGCCCTCGCGTCTGCGCCGCAGTGTGATGAGGATCACGCCGCCCCGGCTGTCGGCGGCGAACTCGGCGCCCGCCAGCG
GTACCACTTCGATGCGGGCGATGTCGGCTGCGGTCAGGCTGCGCAGGTAGGTGACGGTTTCCTCCGCCGAACCTTTCAGCTCGCGGCCGTCCACGTAGACT
TTCGTACCGCCGGCGCCGTTGATCGAGATGCCGTCCTCACCCAGCCATACGCCCGGCGCCCGTGCCAGCAGTTCGGTGCCGTCCTGTCCCGCGAGTGCGG
GCGTATTCCCGATGCTGACCACGAACCTGTCGGCCTGCCGCGTCACCGTCTCGGCGGTCACCGTCACCTCGCTGATTCCGAGAGGTGCAAAGGGTAAATCA
CCGAGGTCGGTATCCGCTGCGGCGACGCATACTTCCTGTTCGAAGGGATGGTAGGCGACGTGGCGCAGTACGAGCGTGTAATGGCCCGTATCTGCCGTGA
GCACGAAGCTCCCTGCGTCGTCGGTCGTCGTACCCGCTGCCTGGCGTCCGTCGCGCAGCAGGACTGCCGTGGCGTAGCCGACCGGGCTCTCGCCCTCCAG
GACGCGCCCCCGGAGGGTATGTTGAGCTGCGGCCGGGATCGTCGCGGCGAGCAGCAGTGCGGTGAGCAGCGGGCGGTACATGCGCCTATTCGTTTTCGGT
GGCAGGATAGTAGAGCTTCAGCGTAGCCGTGTTGTGCTCGGCGGGCAGCTTGCATTTTCGGGTGGCGTAGCCTTCGGCAGGGCCGCGCTCCCCCATCTTCA
TTTTGTAATCGCCGTCTTCGTCGTGGAAAAGCGAAAGCTCGGCCGCATCGGCTTCGATGCCTTCGAGCGTCACGACGACTTTGCCGGCCCTTGCGGCTGCCA
TGCCGTAGACGGGCTGTTCCTGCCCGGCGACCTTGGCCATGACGAGGATGTTGCCCTTGTCGTTGCGAATGTCCGATACGGAGACTTCGAGCGTTTTGGCC
GAGACGGCCGTGGAGGCCAGCATGAAGCTGGCGGCCAGTAGCAGGTGTAAGGTTTTCATAACCGGGATTTTTATGGGTTTAACGGTAATCTTTGTCCTCGTC
GCAGCCGGTCAGGTACCAGACCAGCGAGAGGATCAGGTTCAGCCCCCAGCCGGCCATGACCCATACCACCCACCAGTAGTGGGGCGAGGTGTACCAGTTG
ACGAAGGTGAGGAACGCGCATACCGCGACGTAGGTGGTGATGCGGCGGGCGAACCAGCGGCCGAAAAGGCGGCGTTTGTCGGTGGTGTTTTCCAGAAGTT
CCATAGGTTTGCGGTGTGAAATGACTCTCTTGGCTTATAAATCTGTTGTAAATATAAAGTAATTTATAAAACAAACCAAATTTCCGGACGGGAATTTTCGAAAAA
ATTCGATACCTATATTAACAGCCGAGTAGTTTTTATGCGCGCGGGGTATTCCCGGTTGCCGGAAAAAATGTAAATTTGCGGGAAATAAATTGATACCGCACCGA
TATGAAGAAACTCATTTTGTCCCTTATTTGCCTTGCCGCCGCATGGGGTGCCGCGGCCAAGGTCGAACTGCCCGAGATCATCGGCAGCGACATGGTGCTCCA
GCAAAATGCCCAATGCCGCCTCTGGGGCTGGGCCGATCCCCGGGCCACGGTGCGGATCGAAACCTCGTGGGGTGCCGAATGCACCGTGCGAAGCGATGCC
GACGGCCGTTGGTCTGCGCGGGTGAACACCCCGGCCGGCAGTTATGAGCCGCAGCAGGTGACGATCGTAAGCGGCGATAGGGTCGTGCTCGACAACGTAC
TGATCGGCGAAGTCTGGTTCGCCGGCGGGCAGAGCAATATGGAGATGCCGCTCCGCGGGTTCAATAACTGTCCCATCGCCGGGGCCAACGAGATCATAGCC
CGTGCGGGGAGTATGCGCGACAGGCTCCGCTACGTGAAAATACCCCTCACGGCGGCCTATACGCCCCAGGATCGGGTTCCGGGGAAATGGAGGCCATGTA
CGGTGGAGACCGCGCCCGATTTCACCGCCACGGGTTATTTCTTTGCCGAACTGCTCAGCGACGTGCTGGACATGCCTGTCGGCATTGTCGACTGCACGTGG
GGCGGGACGCGTGTCGAGGGTTGGACGAACCGCGAGATACTGGAAACCTATCCCGATATCGACCTGACGGAAAAAGGCATCGAGGCTATGACCGACTGGCT
GCGCCCGATGGTGATGTACAATGCCATGTTGCACCCGGTTGCGGGGTATACCGTCCGTGGCTTCCTGTGGTATCAGGGTGAATCGAACGTAAACCAGTATAA
GGACTATGCCGTCCGCCTTTCCAACATGGTAGGGCTGTGGCGTTCGCTCTGGAAGCAGGGCGATATCCCGTTCTATTACGTCGAGGTCGCCCCGTTCGCCTA
TGGCAACGGCTGCGGCGCCTATCTGCGCGAAGCGCAGTGGAAGGCGCAGGAACTCATCCCCAACAGCGGCGTGATTTGCACGAACGACCTGGTGGAACCC
TACGAGGCAACCAATATCCACCCCAAGAATAAACGGGAGGTAGGGCACCGGCTGGCCTACATGGCACTCAACAAGACCTACGGTTTTTCAGACATCGCCTGC
GAAAGCCCGCGGTTCAGGTCGATCGGGATCCGCGACGGAAAGGCTTACGTTGCCTTGTCCCATACCGAAGAGGGGTTCAGCCGTATGGAGGATATCCGCGG
GTTCGAGGTGTGCGGCCCCGACCGGGTATTCCATCCGGCCGATGTGGAGGTCGACAGGCGGGAGCTGTTGCTGGTCGTCTCGTCGAAGGAGGTTGCCAGT
CCGGTTGCCGTACGGTACGGGTTCCGCGACTTTATGCCCGGCAACCTGGCCAACATCCGGGGGCTGCCGGTCGTGCCGTTCCGCACGGACGATTTTTGAAT
GCCGTGATACCGTAACAGGAAACCCGCCGGGAGGCGGGTTTCCTGTTTATTTGGCCGGCATTTCTTTCAGTACCACGTCCACCCATTCGGGATGGCGGCGG
ATATATTGGGCGATGAAGGGGCACTGGGGGATCATTTGTAGTTTCTTGGCGCGCAGATCCTCCAGTACGTCGTGCACCAGCTCGGCGCCGATGCCCTGCCC
TTCGTATTTTTCAGGGACTTCGGTGTGTGTCAGGATAATGAATCCCTGGGTTTTGATGTATTCGATGATGGCGATGTCGTCGCCCAGGTCGAATTCATAGCGC
TTTTCGTCGGCATTGTCGACCAGCGTGTGTTTTTTTCTTTCCATAACCTTGCGTTTTAGATAGTGGCTTTACCTCTTCAAATATAGTTCCTAATCGGAAAAATGT
TATTTTTGCATCCGAAAGATGGAGCCTGTTTCCGACATACTGCTATCCTGGTTCGCCCGCGAAGGCCGTGACTTGCCCTGGCGTCGCACGCGTGACCCTTAC
CGGATATGGCTTTCGGAGGTTATCCTGCAACAGACGCGCGTGGCGCAGGGATTGGAGTACTACCTGCGCTTTACGGAGCGGTTCCCCGACATCGCCGCACT
GGCAGCCGCGCCCGAGGACGAGGTGCTCAAACTGTGGCAGGGGTTGGGCTATTACAGCCGTGCCCGCAACCTCCATGCTGCCGCGAGGCAGGTCATGTCT
CGGTTCGGCGGGGTGTTCCCTGCCACGTACGGGGAGGTGCGCGCGCTGCCCGGCGTGGGCGACTATACTGCGGCGGCCGTCTGCTCGATCGTTTACGAC
GCCCCGTGTGCGGTGCTCGACGGGAATGTATACCGGGTGCTGGCACGGCTTTTCGATATCGGCATCCCGATAGATACGACGGCCGGGAAGAGGACGTTCG
CCGAGTTGGCACAGTCGCAGCTGGATACTTCCCGCCCGGGACTTTACAACCAGGCGATCATGGATTTCGGAGCCCTTCAATGTACCCCCGCGCAGCCCCGG
TGCGGGGATTGCCCCCTGGCTGGGCGGTGCCTCGCCCTGGCTGCCGGCACGGTGGACGTGCGTCCCGTAAAACAGGGGCGTGCGAAGGTGCGCGACCGC
TGGTTCAACTACCTGCACGTGACATGCGGCGGGCAGACGCTGCTGCATCGTCGCGGCGGAGGCGATATCTGGCAGGGGTTGTATGAATTTCCGCTTATCGA
AACCGACAAGGCCGTGGATTTCGCGGAGCTGGCTGCAACGGCGGCGTTCGGGGAGCTGCTCGGCAACGGCAGGTGGCACCTGCTTCGCAGCGTCACGATG
CCCGGGCACCGGCTTTCGCACCAGCTTTTGCATGCGGTTTTCCACCGCATCGAATGCGATGCCATGCCCGCTTTCCCGTCCGTGGAAACCGCCGCGCTTGG
CGGCTATGCCCTGCCTCGGCTGGTCGACCGCTACCTGATCAAATACCAGATATAAGCGATGTAAATCGCCAGGAAGATAGCGCCTTCCCACCGGTCGAGCTT
TTTGCTGCGGAACGTGAAGGCTGCCATGAACAGCAGCAGCGAACTGAGCACGACCACGAGGATGTCCGTCAGCGTGATCCCGCCCATCGTCAGAGGGTGGA
TCGTGGCGCTGATGCCCAGGATCAGCAGGATGTTGGCGATGTTCGATCCGATGACGTTGCCGAGCGCCATTTCGGCCTTGCCTTTGAGCAGCGATACCACC
GACGAAGCCAGTTCCGGCAGCGACGTGCCGCCCGCCACGAGCGTGATGGCGATGACCGATTCGCTGATTCCGAGCGTGCGGGCGATCGCCGTAGCGCTTT
TCAGGAACATCTCGCCGCCGAAGATCAGTCCCGCGAGCCCTCCTGCGATCATCACCCCCGCCAGCCACAACGCCATCGGTTTTTTTGTGCCTTCGGCGTGTT

GCGCCCCGTTTTTCAGGGCAGCTCCGGATACGGGAACGGTCATCCCGGTTGCCGCACCGGGCATGCTGCGTTGGCGTTTGGTCGTGCGGATCATGTACCAC
ATCAGCCCGATGTAGAGCGCGAGCATGACGATGCCGTCCGGGCGGCCGATCCTGTCCGTTGCCCCGGCGCATACCAGGCGGTCGGAAGTGACCGCCAGCA
GCACGAGCGACGCGGCCATGCCGAACGGGATATCGCGCCGGATGTTCTCCTTCGTAAGCACGAGCGGCCGGATCAGCGCGCAGATTCCCAGGATTACGAA
GACGTTGAAAATGTTCGAGCCGACGACGTTGCCGATCGCCACGTCGCTGTTGCCCGCCGCAGCCGAGAGCACCGAGACCACCAGTTCGGGCATCGAGGTG
CCGACGGCGACGATCGTCAGGCCGATGATGAATTCCGGGACGCGGAAACGCTGGGCTACCGCCGCAGAGCCGTCCGTCAGGAAATTTGCGCCGCCGAGGA
TCAGCCCCAGGCCGACGAGTAGTAACAGGATGTCCATTGTTTTATCCGAGTATGATAAGACTTACGGCCATGAGTGCCATGCCGGCCACCACGCCGTAGATC
GAGATGTGCGCTTCGCCGTATTCGCGGGCCGCAGGCAGCAGTTCGTCGATCGAAATGAATACCATGATTCCCGCCACGCCCCGCGAGGATGCAGCCCATCAG
TGTCGGCGTCATGAAGGGCATCAGCACCAGGTAGGCCAGCACCGCCCCGACAGGTTCCGCGAGCCCCGACAACAACGAGAGTTTGAAGGCTTTTTTGCGGT
CGCCCGTGGCGTAGTAAACCGGGATCGAGACGGCAATGCCTTCGGGGATGTTGTGGATGGCGATCGCCGCGGCGATGGCAACGCCCAGAGCCATGTTGTC
CACGGCCGAGGTAAACGTGGCGATACCTTCCGGAAAATTGTGTATGCCGATTGCCAGTGCGGTCATTACGCCCATGCGCATGAGTTTCGGGTTGCGGGGCT
GCTTGTCCATCTCCTCGACCATGTGTGCCTCGTGGGGGTTTTCGAACGAGGGGACGAGCCGGTCGATGATGCCGATGAGCAGTATCCCGGCGAAGAAGCTG
GCGACGGTGACGATGATCCCCGTCTGTGCGCCCCATTCGTCCGAGAGGGTGGCGTGCGCCTGCTGGAACAGCTCGACGAACGATACGTAGATCATCACGCC
GCCCGACAGGCCCAGCGAGAACGACAGCAGCCGCTTGTTGGTGCGGCGGGCAAAGAAAGCGATGGCGCTGCCGATGCCGGTGGCCAGTCCGGCGCCCAG
CGTCAGCAGCAAGGGTATGAGGATATTGTGTTCCATAGGGTTTACCGTTCGTTTATCTGGCTCTATCGCCGCGATGATGGTGCAAACGGGCGGCAAGGGAGC
TTGCTCTCAAATGCCCGGGTGCAACTTATCTTCTGCAAAGATAGTGCTTTTCCGGTGAACACCGTAAAAAACCGGGCCCTTGCGAGCCCGGTTTTTCTGATAG
GAGTATAACCTCCGCCTATTCGGAGTGCTGTTTCCGACAGGCATTTGTGCGCCCAAGCGCTGATATGCCGATGTGGCGGTGCGGTGCTGCCGGTCTGCCGG
ACGGCATGCTGCGTCCGGTTATTTGCCGTGTGCTTCGCGGATCAGCGCGCCCAGCACTTCGATGCTGTCGGCCGTGATGTGCGGCTGCGGGTCGGCCTTGT
CCGCCACGTCGAGCGTCGTTTCGCCCGAGAGCACCAGCACGCCCATGGCCTTGGCATTGTGCGCCATCGCCACGTCGGTGTAGATGCGGTCGCCGACCAT
CGCGATCTGGTCGGGTTGCAGGCCGTGGCGCGAGAGTATGCCCGAGAGCATGTTGGGATCGGGTTTGCCGAGCGTGATGTCGGGCCGCCGTCCCGTGGCA
TGTTCGATGCAAGCGCAAATGGAACCGCAGTCGACCAGTATCGTCGGCTGGTCGGTGGGGCATACCCGGTCGGGGTTGGTGGCGATGTAGGGCACGCCCT
GCGACACCCACCATGCGGCGCGGCACAGGCGGTCATACTGCAGCGTCATGTCGAACGCCGCGACCACCACGTCCGGCACGTCGTCCGCCGAATCTGCGGC
CGATTCGAAACCTGCGGCCTCGAATTCCGAGATCATGCTCGGCGTGCCGAGCAGGAAAAGCCGTTTGGCCGCAGGGTAATGCTCCCTGATGTAGTCGATCG
TCGCCAGTGCGGTGGTGTACATCTCGTCGCGTGTCGCCCGTATGCCCAGCACGTCGAGTTTGTGCAGGTAGTCGGCGATCGACTTCGACGGGTTGTTGGTC
AGGAACGAATAGCCGATGCCCATCTCCTTGAGCCCCGCGAGGAACGGCTTGGTGTAGGGAAAGAGCGACATGCCCATGTAGATCGTGCCGTCCATGTCGAG
CGCCACGTGCCGGATGCGGCGCAGGTGTTCCATCAGTTCGTCGTGCGTGTAGACCGAGCGATATTTTTCGAATGCAGCCATTTGTTTACTTGCGGTTGAGAT
TAAGGTTCTGGAATAAGTAGGGGTAGTCGGTTTCGATGACGTCGCAACCCGTCGGGATTTCGAGTTCATAGCCCTGTATCTTCTGCGCGTCGGTGGCCCTGC
GGTCGTGCGTCGGGGCGAGCGAGATCATGCACATCACGCCGTTGCGGTGCAGGTTGTCGTAAAGCTCCTGCTGGTCGGGCAGCATCATCGTGCCGACGTA
GGCCATCACCTGCGACCAGGGAATCCGCTCCCCGACGTATTCGTTGAACTCCTTCATGTTCTTGCACCAGCACGAGAACATCGCCTGCGGGTCGCGGTCGA
GATACGTGCGCGCCTGTTTGGCGTTGCGCACGGTATACATCACGTTGGCCGGTGCCAGGCGGTTGATGAATTGCGACATCACTTCGAGCGGGACGTCCTTG
ATGTCGAGGTTGAGGATCGTCTTGCCCTGGCTCCATGCGATGCACTCCTCGAGGGTGGGGATCTTGTAGGTCGTGACGTTGCCTTCGCGATCCTTGAGGAAG
AACTGCTGCAACTCCGCGTAGGTGTAGTCCGAAACCTTGCCCGTGCCGGTCGTGGTACGGTCGATCGTGGCGTCGTGCATCAGCACGATCACGCTGTCTTT
GGTCATGCGCGGGTCGATCTCGAAGAACGAGGGCATCATCGTGAGCGTCTTCTCGAACGACTCGATGCAGTTTTCGGGGAATCCCGTCACCATGCCGCCGC
GGTGGCCGCTGATGACGACCGGCCGCTCGGGCGAATAGCGGAACCACTCGTGCAGCTGTGTGCGGTCTTTGATGTCGATGTAGTAGGGCCCTTCGGCCTTT
TGCCCGCCCGGGGCGCATCCGAGCAGCAGGGCGGCGGTGGCGAAGGCGGTCAGTAGTCTGGATGTTTTCATAGTGTGTTTTATTGGTTCTCTGCGGTTTGT
TCCAAGAGCAAAGGCGCCCGGAAGCGCCTTTGTCCCTGTTCTGTTTCCTTTCAGGCGGTTACTTGTCGATCCGGTGGCGTTTGATGACCTCGCCGCCCGTGT
TCACGACGTCGATGTCGATTCCCCTGCGGTCGGCCTTCACCACGAGCTTGTCCCGGTTCGAATTTACGAGGATCGGGAAGTTCGTGCCCCGGCTCCCGTCG
TCTATCCACTGGTAGCGGTGGTAGTGCCCGCATAGCATGATGTCGACCGAAGCTTCGTTGAGGATCGGTACGAACAATCTGCGGATCTCCTGTTCGCCGTGC
CAGCTGCTTTTCCCGCCCGGGATCATGTGCAGCACGACGATGTGCAGGGGTGCCGCGCGGTATTCCTCGCTTTCGACCACGCCCTTCAGCCAGCGGGCTTC
CTGTTCGCGGTAGGCGTCGGTGGTCGACAACCCGTAATAGCGGATGTCGCTGTCGGGTTTGTCCTCGCCGCAGTCCAGCACGACGAAATAGGCCGGCCCCT
GCCGGAACGTGTAGTATACCTCGCCCGTGGAGGTCGGGAAGTAGTCGAGGAAACTCGGGGAGAAAGAGCCGCGGTTCTCGTGGTTGCCGCGCGTGGCGAA
GATCGGGATTCCCGCGGGCGTCAGCAATTCGGCGGCCGAATTGAGATACCCCTCGAAAAGCTGCTTCTCCGACTCCATCGAAGTGAGCATGTCGCCGTTGA
AGCAGACGAAATCGGGTTTGGCCTTTTCGGCACCTTCCAGCAGCAACCGGAAGATCGAATCCTGGGCGTGAATGTCGTTCACAACCCAGAATTCGGTTTGAT
CCTTCTTTTCGTCGAGCGTAGTGGCCGTATAGGGCTTGTGTTTGAGGATGTCGCTCCCGTAACCTTCGCCCAGGACTACCCGCTTGTTGCCTTCGTCGCAGA
GCACGCCCTGCTGCATGATGCGGTAACGGTAGGTCGTCCCCGGAGCCAGCCCCTCCACGCGTACGCGGTGCAGGCGGCCTATGCGGCGTTTGCCGATGTG
CGAATCGTAATATTTCGGCCGCTCGGCGGCGTAGAAATGCGATCCGTCGTCAGGAGCGACTTCGACCCACGATGCGGCATTGATGGTCGTCGTCCACACGA
CGGTGAAACCGTCGTCGGTGACCTGTTGCAGGTAGGGCCCGTGGGCAATGCGTACCGAGGGCTGTCCGATGGCAATGCCCCAGGCCGATAGGGCGACGAG
TATCGAGATCAGTCGTTTCATTTTGTTTTTGCTTTTGAATCGCAATTCACAAAGTTACGAATTTTTCCGTAATAACCTACTCGCCGACCGGGAAATCCTCGGCAA
GGCCCGTGGTGAATTTCGTTTGCGTGAAGGTCGTGCCGAACCGATCTGTGGCACGCACTTCGACCGACGCTGTCGGTGACTGAAGCGTGTAGTAGTACAGG
TGGCGGACGTTGCGCTGGTAGTAGCTCGTGGGTTTCTTATTGCTGTCCAGTTCGGCCTTGCCTACTACGCCGATGTGGTAACCCAGTGCCCAGGCATCCCAG
GACTTGTCCTTTTCATTGCCTGTGTAGTCGAAATGCGTCATTTCCCCTGTTTTTATGCCATTCTCGTATACTTCGATTTTCCAGTCCTTGTCGGCGTTCCAGACA
TTGGCTACGATCTGGTCGTCGCCCTTGTAATAGAACTGGTAATTTTTCGAGTAATCACGCATGAAGATGTCGTTGGCGCGGTAGAGCCGTATCTGCTGCTGGG
CATCGAGCCCCGTGCCCTTGTATACCCAGTCGGTAATCGTGGCGCCCTTGATGTTGTAGATGGCGTATCCGTTGGGCGTGCCGTCGACGTTGATCGTGGAGT
TCCACCATGCGCCGCAGGCCGCGCCATGCACGTGTTCGTAGTGGCTGTCGTAGACGTTGTTCTCGGCGTAGTGCGTGTGCCCCGACATGACGTGCACCTCG
GCGAACGGCTTGAGCAGTTCGAATACCGCTTGTACGTTCTGGTTGCCGCTGGCGCGCATGGGGATGTGTACGCAGAGGATCACCATTTTGTCTTTCGAAACA
TAACTCAGATCCTGTCGGAGCCATTCGACCTGGTCGTCGCGGAATCCGAGCGAATAATCCTTGTGGCTCGGGAAAATGATGTTGTCCATGGCGACGATGTGC
ACGTTGCCGCGGTCGAACGAATAGTTTACGGGGCCGGTGGGATCCTCTAGAGTCGACCTGCAGGCATGCAAGCTTGAGTATTCTATAGTCTCACCTAAATAG
CTTGGCGTAATCATGGTCATAGCTGTTTCCTGTGTGAAATTGTTATCCGCTCACAATTCCACACAACATACGAGCCGGAAGCATAAAGTGTAAAGCCTGGGGT
GCCTAATGAGTGAGCTAACTCACATTAATTGCGTTGCGCTCACTGCCCGCTTTCCAGTCGGGAAACCTGTCGTGCCAGCTGCATTAATGAATCGGCCAACGC
GAACCCCTTGCGGCCGCCCGGGCCGTCGACCAATTCTCATGTTTGACAGCTTATCATCGAATTTCTGCCATTCATCCGCTTATTATCACTTATTCAGGCGTAGC
AACCAGGCGTTTAAGGGCACCAATAACTGCCTTAAAAAAATTACGCCCCGCCCTGCCACTCATCGCAGTACTGTTGTAATTCATTAAGCATTCTGCCGACATG
GAAGCCATCACAAACGGCATGATGAACCTGAATCGCCAGCGGCATCAGCACCTTGTCGCCT

**P2B11 fosmid sequence**

Highlighted yellow: P2B11-Iso1, Green:P2B11-Oxi1, Turquoise: P2B11-Oxi2, Pink: P2B11-Iso2

GATAGTGTTCACCCTTGTTACACCGTTTTCCATGAGCAAACTGAAACGTTTTCATCGCTCTGGAGTGAATACCACGACGATTTCCGGCAGTTTCTACACATATA
TTCGCAAGATGTGGCGTGTTACGGTGAAAACCTGGCCTATTTCCCTAAAGGGTTTATTGAGAATATGTTTTTCGTCTCAGCCAATCCCTGGGTGAGTTTCACCA
GTTTTGATTTAAACGTGGCCAATATGGACAACTTCTTCGCCCCCGTTTTCACCATGGGCAAATATTATACGCAAGGCGACAAGGTGCTGATGCCGCTGGCGAT
TCAGGTTCATCATGCCGTTTGTGATGGCTTCCATGTCGGCAGAATGCTTAATGAATTACAACAGTACTGCGATGAGTGGCAGGGCGGGGCGTAATTTTTTTAA
GGCAGTTATTGGTGCCCTTAAACGCCTGGTTGCTACGCCTGAATAAGTGATAATAAGCGGATGAATGGCAGAAATTCGATGATAAGCTGTCAAACATGAGAAT
TGGTCGACGGCCCGGGCGGCCGCAAGGGGTTCGCGTTGGCCGATTCATTAATGCAGCTGGCACGACAGGTTTCCCGACTGGAAAGCGGGCAGTGAGCGCA
ACGCAATTAATGTGAGTTAGCTCACTCATTAGGCACCCCAGGCTTTACACTTTATGCTTCCGGCTCGTATGTTGTGTGGAATTGTGAGCGGATAACAATTTCAC
ACAGGAAACAGCTATGACCATGATTACGCCAAGCTATTTAGGTGAGACTATAGAATACTCAAGCTTGCATGCCTGCAGGTCGACTCTAGAGGATCCCACCTCG
ATGGCACGGCGTATTTCGACCGCATCGTCAAGCACGATACGCCGCTTGCGGATACGTATTACATCGTCGAGAAAGATGTGCCCAAGGGGTATGCGGGCGCC
GTCGACCCCATTCGCGTCATCGTCGATGCTGACGGAGTGCACGCCGACGCGGGCGTGGCAGGTGACGACGTCACGGTTTCCGTCGGCATCGGCTCGCTCA
TCGATTCGATGAACCACTACGGCAGCAACGACGGTGTCGAGGTGACGCTCCACGATGTGGTGGCGCAGCTTCGCGTCGGTACGGTGAATGACAGCAAGACG
CCCGGGTTCTATTGGATCGGCGGTTGGGAAAAACCCGCGCCTGAGGCCGAGCAGCAAGATCTGTACTTGCGCTATTCCGACGAGAGCGCCACGCACGAATA
CGTCTGGAATAATAAGCATAAGCCCAAGAGCGGCACGCGATTCACGACCGATGAGGGCTTCATCCGCGCTGCGGTCAAGCAGGACGCCGAGTCGTGCGATC
AAAATCACGGTATCGGCACCTGGGATAACTTGGGGACCAAGGACATCACCAATCTCTTCACCGGCTCGACTGTGATCCACGTGACGAACCATCGTGTCGCCT
CGCTTGAGGTGGAAAAGCAGGTCGTGATTCCCGAGGGGCTCACCGGCCCCGTCGATTGGAAGGACAAGGCCTTCACGTTTGTCTTCGAGTTCAAGGACAAG
GATGGCAAGCCGCTCGCCGGGGAATTTGACGCACGCGTGTTTGGCGCCGACGGCGTGCAGCGCGGCGACGACTTCAAGATCGCGAGCCGCGGCACCCATA
CGCTCAAGCACGGTGAGGTCATGCGCGTGTACGGCTTGCCGGATGGGGCGACCTATACGGTGAACGAGTCCGTGGAGCAGATGCCCGAGGGCTTTACCCA
GACGCTGCCGCACGATGGGGCGGGCAAACCCACAACCGCCACGGGTGCGATCAAGCTGGGCGATACATCCCGCGTGCTGTTCGAGAACACGTACACGCCT
GCGCCCGCCATCCTCAACACCGCGGCCACCCTGCAGGTGAAGAAGGAGTTCGTCGACGCCGAGGACAAGCCTGCGTGGGATCTCGAGTTCAACGTGAAGC
CGACGTTCAGCTTCGTGTTGGACACCGACCCGGGCACGCCGATGCCTGATGGCGCGACTCTCAATGAGAATAACCGTCTCCAGTCGATCATCGTGATCGATC
GCGAGGATGCGCAGGCGGGGTACGCCCGGAGCTTCGGCACCATCCATTACGAGCATCCGGGCACCTACCTGTATGCGCTTGCCGAGAAGACGCCGGATGC
CGCCGACCGCATCCCTGGCGTGTCGTATTCCGACGCCGCGTATCGCGTGGTGGTGACTGTCTCTGATGATAAGGCGGGCAAACTCGTCGCAGACGTTCGGC
TGCGCAAGCTCGCTGACGATGCCGGCGTTCCACAAGGCGATGACGAAGCCGGGGTACCCGTTAAGCCTCAGAACGGCGTTCACACCGCGTTCTTTAAGAAC
GTTTTCGCCCTGAATAAGGTTCAGATCGGCCCGTTGGCGACGAAGGTTCTCGATGGACGTGGTTTCACCGTCGACGGCTCGGGCGCGGGCGAGTTCACGTT
CCGCATGCGGCCCGCCGGCGACCATGCGGCTGACGCTCCCATGCCCGAGGGCACTGCGGGCGAAGGTGCATCTCGATACATCGAGGTGCACAACAAGGG
CTCGATGCTCTCCTTCGGACAACCCACGTTCACGTTCGAGCATCTAAACGGTGCTGATAGGCGGGGTCGCTTCGACTACGAGCTGTTCGAGGTCATCCCAGA
TGATGCCGTCAACGCTAATAACACAAAATGGAGCGACGCGTCGCCCGCGGAGAAGCTTGCCGGCGGCTTCACCAAACAAGGTATGACATATGACTCGACGC
GCTTTGTCGCGCGGGTGAACCTCGCGCTCCAGGATTCCCCGCTCAAGCGCACTGGTGGCACAGAGGCGCTCGTTGCCACGATTACCTACTACCAGGTCACG
GGGCTGAACGAGCAGGGCGTTTGGACGGGTTCTGACAACGATCTGACTCCGGTTGAGCCTACGCCCGGCGGTGCCGACCGCGTCGAGTTCCTTAACACGTA
TCGTGCCGCTGCCTCAACGGCGGGCATCAGACTCACTAAGACGCTTGAGGGCCGCGATATGAAGGACGGGGAGTTCTCGTTCACGATCGAGGGCGCCGAT
GAGAAATCTCGCAAGCTGCTGGCTGAGACGGACAAGAGCTTCAGCGCTCCTGCGGCGATGAGCGGCGAGAAGGCCGTTATGGACGGCAAGCTCCAGCTTAA
GTTTGACCAGTCGCATGCGGGCCTCACGTATACGTACTACATCAAGGAGGTCACGCCCCTATGGGGTGGTGAGCACATACCCGGCCTCGATGGCGGTTGGG
ACAATTCGATTTACCGTCTTGAATACCGCGTGATCGACAACCTCGATGGCACGCTCAAGGTGGTCCCAAGGCTCTATCGTACGTACGATCGCTCGGGTGCCA
TCGTCAACATTCCTCTCGACCTTGGCACGCACACTGATTCCAGCGGCGTGATCACGCTCGATTTCGTGAACCGTTACAGCGCGAGCACGACCTTCGCCGGCA
TCGAGGTCGCCAAGACCATGTATGGCAAGGACCTGGAGGCCAAGCAATTCGGCTTCACCTTGGAACCGGTCGACCCAGCCTCGGCGCAGAAGATCCTCGAC
CGCGGCCTTCCCGCCGATGAGGAGCAGCTCGAGCGCATCTTCGCCGCCGGGGAGGCTCCGGAGGGCGTCCCGCACCTGATGGAGAAGCTCAAGGGCCTG
AAGTTCACGCAGGCGGATACAGGGAAGACGTATTCGTATCGGGTTAAGGAGAATGTCGATCCCGCATGGGATGAGGACGGCGAGGCGCCCGGCGTTCAGG
TCGATGGCGTGACCTATGACCGCTCGCAGTTCCTGGTCTCCATCAAGCCCGCGGACAAGGGCGACGGAACCATGAGGACCGTGACCACGGTCAAGCGCAC
GCATGACGCGCTCACGGGCGATCAGCTGGCAGAGCCCGAGCCGATCGGCAAGTGGGATAGCGACAACCCCGCAGAGGGCCAGGCCGAAAAACCGCAGGC
GCCCTTTGTGAACAAATATAAGCCAAGGCCCGGCGCCCTCGAGTTCGATCGCTTCACCAAGCGTCTCGTCGGCCGCGATTGGTCCGCTGCGGACTCCTTTGA
GTTCACCTGCGAGCAGATGTCCTTCAGCTCTGGTGGACATGACTACAAGCCGGGTGATGAGGGCTTCGTCCTCGTTCCGCTCGTTGTCGCGCCGGTGAATG
CCGACTCGCCTCAGAACGAGAACGGCTTCAAGCTGTTTGGGATTTCGACGAGCCCGGACGTCCTGTACCAGCAGAGCGGACGGTACGACTACCGCCTTCGT
GAGATAGTTCCCGACGACGCGGTCAACGCGGCAGGCGTGCGCTATGAGGACGCGACGCCCGAGCAGAAGGCAGCCGGTGGCTTTGTCAAGGATGGCGTTA
TTTACAGTAATCGCGAGGTTCGCTTTACGGTTTCCGCATTCGACCTCGGCACGGGCGACACTTCTCGTGATTGGGCCCATCTTGCAGGATGGCCCGCAGGAGG
ATGGCGCGTACTTCACCAACAAGGTGCGTCGGGCGGATACGGCCCTCATGGCCAAGAAGGTGCTAGAGGGACGCCCGAATCCGCTCACCGAGGGCGAGTT
CTCCTTTGTGCTCGCACCGGTCGACGGCGGTCAGCCGCTCGTCTCGAGCAATGACGCCCAGGGTAACGTGACGTTCGCCGGCTTGACGTTCGACGAGCCAG
GCGTGCATCAGTACACATTGCGCGAGGTGCATGAGGCCCCGCAGGCGAACGGCGTGACGTTCGATGATGCGGTCTACGATGTGACGGTGAACGTCACTGAG
GCCGCGGGCGGTGAGCTCGTTCCTGAGGTGTCGGTGTTCAAGGACGGCAAGCCGGTTGAGGGCGTCCCCGTGTTCACGAATCGCTACGAGTCCACGGTCG
ACTACTCCGCGCTGGGCGGCCTCAAGATCACCAAGGTCCTCAACGGCCGCGATATGGCAGACGATCAGTTCGCCTTCACCGTGAAGGCGACAGGCGACGAC
GCCGAGGCGATTGCCGCCGCTGCGAGCAAGCTCGGTTTTGCCGAGGGGCAGACGGAGCGCGTCTTTTTGTCCCACACCGCGAAGGTGGGGCAGTCGACAC
CATAGATATTCTCGCCGGTCTTCCCGGTGGACTTGTGTTCACCCAGGCTGACGCGGGGAAGACCTTCGCCTACGAGGTGCGCGAGAGCACGGGTGGCGGT
GCCGGATACACCAACGACCAGACGGTCTACGGCGTTTCGATTTCCATCGCGGATGACGGTGCGGGCACGCTGACGGTCACCACGACCGTCACGGGTGGCG
CAGAGGGCGAGAAGACCTACGTGTATACGAGCGGCCAGACGGCCGAGGAGGTCGAGCCCGTGGTGCTGCCCTTCGTGAACAGCTACGCGGCTCAGGGTGA
CGTCTCGATCGCGACGAGCAAGGTGCTCCACGGACGCCCACTCGAAGCAGGGGAGTTTTCCTTTGAGCTGAGGACGGGCGACGGCGCGGCTGGCAGCAGC
GAGGTCATCGCGCGTGCTGAAAACGCTACCGACGGAACCGTGGACTTCGGCAAGATCGAGTTCATGCTCGATACGTTGGCTGAGGCCGTCGATAAGGGGTA
CGCCACGCGCGATGCCCAGGGCATCTGGCATGTGCGCTACACCGCCGTCGAGATCGCTGACGGTCTGCCCGACGGCGTCTCCGCGACCACCCAGAGCCTG
AGCTTTGAGGTGAGCGTCGAGGACAAGGGCGACGGCACGCTCGCGGCGACGGTGCAGGGGTTGGACGGCCACTCGTTTGAGAACACCTACGTCACGAACG
GTGGCGAGCCGGTGTCCATGGCACTGGTGGGCCGCAAGGTGCTGACGTACGCCGAGGGCCTGGCGCCTGCGGACATCACCGGCAAGTTCACCTTCACCGT
TTCGGGCGAGGACGGCGCGCCGATGCCCGAGCGCACCGAGGTCACCAATGCCAAGGACGGCTCGGTCGACTTCGGTATGATCAAGTTCACCCTCGACGAC
CTCAACCGCAAATGGGCTACGGAGCATCCCGAGGACACCGGCATGGAGGCGCTTGCCGACGGCGAGGCGGCTGCGCGCAGCGGGAAGCCGCGCACGGTG
TCCTTCACCTACACCATCACGGAGTCCGGCGAGGTGCCGGGCGTGACTAACGATCAGAACGCATCGCGCACGGTGACCTTCACCGTGACGGACGACGGCTC
GGGCGCCCTCACCGTGACGCGTGATCCCACCGAGGGCGCGAGCTTCACCTTCACCAATGCTTACACGGTGGACGAGCTTTCCTCGAGCGTGACTGACCAGA
TTGCGATTTCCAAGGACCTCACGGGCAAGCCGCTTGCGCAGGACGCCTTCACCTTCGTGCTCACCGAGCTCGTCGATGACGGAGCGGGCGCGGACTTGCAG
GTCCTGACCGCGACGAACGGCGCGGACGGCAGGATCGCTTTTCCGGCGCTCACCTATACGGAGCCGGGCGTCCACACGTACCGCCTTGCCGAGGTCGTCG
ATGAGAACGTTGGCGGCATCGAGTTCGATAAGGCTGTGTATACGGTGCGCACGACGGTCGTCGACAACGGCGACGGCACGCTTTCCGCCACGCACGAGGTT
CTCGACGAGAGCGGGAGCAAGGTCGATGCGGTGGTGTTCGCGAACGCCTACGATCCCGTGGGCGTCACCGTGCCGCTTATGGCATCCAAGATGCTGACCG
GTCGCATGCTAGAGGATGGTGAATTCTCCTTCGAGTTGGTGGATGCGGACGGTACGGTCCTGCAGACGGTCTCGAACGACGAGCGCGGCAACGTCGCGATTC
ACACCCCTGCGTTATACGGCTGAGGACCTGGCGGATGCCGCTTGGGTCGATCAGGACGGGGTCATCTCGCGTGCGCGTGAGTTTGCATACGCCGTGCGCG
AGGTCGTTCCCGCGGATGCCGTGAACGCGGATGGCGTGCGCTGGTCCGATGCGAGCGCCGATGATCGCGCCGCGGGCGGGTTTGCCAAGGACGGCGTGA
CTTACGACGGGCGCGTGTTCGCAGCGACGGTCACGGTCTCCGACGACGCCACCGGCTCGCTGGCGGCTACGGTGACGTGGGATGCCACCCCGGCGTTCG
AGAACGTCTACACCGACCCCGAGGAGCCCGACAAGCCGGATAAGCCGGATAAACCGGATAAACCTGACAAGCCCGAGGACCCCGAAAAGCCGGACAAGCC
GGAGAACCCGGACAAACCGGACAAGCCGGATAAGCCGGAGGACCCGAACGACCCCGAATTGCCTGATGGGCCGCAAGACCCCGGTACGACTGACGCAAACCC
CAGGCCCCCGGCGGATCGAGCGCCGGTAAGCTGCCCGCAACGGGCGATACCTCCCTCGTGCTGACGGTCGGCGTCGCGATCGCGGGAGTCGCTGTTGCG
GGCCTGGGAATCTGGCTGGCGCGTCGCAAGCGGTAGCTCAGAGGCAGCTGGTCCATCTTGCGCGAGAACGCGCATCTATATGACGGCGAGGCGCCCTTGG
GGATTTCCAAGGGCGCCTTTTCAATGCCGCTTGGACGGCTGAATGGCGGTACGCGCCTGACTCGTCATGACACGGTGCCACCGGGGGAGAAGTGTATACAT
TGATTAAGCGCAACCGCGGCGTGGGCAATGAGGAGATCAGCGCCCGCCCTCACGGCATCTTCGAGTTCTTCATTGATTCTGCGTCCCAGCTCTTGGAAGCGC
GCGAATTGGCCCTTGCTCAGGTTGAGCATCTTGCGGTTGGCGCCGTCCGCCGCCGCGTCACCCAGCGTTCTCGCACCTCCTGCCCGTACGTGCGCTCATTG
TCTTCAACGAGCCCTCGTTTCATGCCCTCGAATTTCTAACGAAATTGGGTTCACCAACATTGCGGCTGATGAGGTCACGGCGACGCATGGCAACGAGGTCTCT
TTTGAGACGATTGTCGAGAAGGATCCCGAGTACATCTTTGTGCTCAACCGCGACGCCGCGATTGGTTCCGACGGCGCGGACCAGGCCCAGGAGGTGGTGGA
GAACGACCTTACCCGTCAAACTCAGGCGTATCGCAACGGCTATATTGTCTACCTGAGCAACCCGGCTGTTTGGTATACGGCCGAGGGCGGAATCGAGGCTCT
GAACATCATGCTCGAGGATCTTGAGGTGTGCTGCTGTAGGGCGGCGTAAGGCCTGCATATGGCCTTGTTTAAGCGGGCGCAATCATTCGGTGCGCGGTAT
CGGATTGGTTGTGCCCGCGTTGCCATGTGCTAGGGGAAATCCCTGGCTGCGCCCATGGCTGTTTTGCGGGACGTTTTCAGGACATAATCAGGACACTATCGG
GACATGTCCTGATTATGTCTCGATTCGCGGCAGGATGAGGCGGGCGTTTTCGAGGGGATAGCGTGTCGGAAAACTCGTCCGCGTTGTTCCATACGGTGAGGT
TAAGAGATGCTCTGCGTAAAGACGTCGACAAACTCCTCAAAGGTATCCGTCTGCATAAGGCGGCTTGTCTTGTCGATGGGGGAGTAGAGGCCGATGATGATG
TCCGGGGCATCGCGGAAGTAGGCCATCTCGTCCTTAGCGATGCCGATGAGAAGGACAAAGTGCACCTTGTGCCTTCCCCAGGGTATGGCGGTATCGTTTTTG
ATATCCCTTCTAGACATTTTTCCGTGCGGTATTCGTGGCCTGCCATAGCTTGTTTAAAGCCCCCGCATCTTAAACATTTCGAATAGGTGGAAGAAGTGGCTGGG
GCCTTATGCAAATAACGGCCATCTATACGTGGCAAATCGTGGGTTAGGCGCACGGACCATACCACATACAAGGTCCCCAGCCAAGGAAATCATACCTTATGG

CGCTGGAGGGCTTTCGGCTCAAAGGCCCCAACGGTAGATGCAGACACCGCAAGCGATGCTCAGGGGAAGGCCGGCGGTCTGCGCGGGACCATTCTCGGCG
CGCTGGGGCTGGAAAACGCCAGCTCGCGCTGACTCCTGCAGGCCGCATCGAATCGGGATGGTCAATGAGTGCCACGAGGTGGGTATCTCAGTTTTATACAT
TCGCGTAGGGCGCTGATTTGCGTATGTTTTGCTCTAAGTTGGGAGACGGCTGTAGAGTATAGTTAGCCAGAGGGGCGCACGGAGAGCGCATGGAACGGAAT
GCGGAGCGGCGTGGAGGATTATTGGGGATCCGCACGCCGCATTCGGACCCCTTGAGCGCCCAGCGGGAGCCGTACCTTACCCGCCTCGCGGCCCTCATT
GACCACGCTGATCGACCACGGTTTCTCATGGCTCAAATCGGATGGATAAGGCTACTCGTTTTCGCAGTTGCCGGGCAAGCTGTCTGCATATTGTTTTCCGCGT
AAGCGTGGAATGAAAATGTCCCATTGGTGCGAGGCTAGCGCAAGGCGCTGGGCCCCGTAAAGCGGCCTGCGACCCGCAAATAAGGTCAATGGCCTGCTGC
GAGTATCCCCAACTAGGTTGTTCATCATGCAACGACAGATTGTGCGAATAAGCATCGTTGCCCTTCATGTTGCAGAAGTCAACGAACACTCTGAAATGGAAATC
GTTGAACGCTTTCGGCTCACCTTTAACTTTGAGAACGATGCCCTCCTTCTTGAGCTTTCGGCGTATGAAATCGGCGCCTTTCTTTGTCCGATATATGTACTTGT
TGATTGGATCGGTGAGCGAGTGAAGGATGGAGACGGCGGCGTCAGCGTCGCTTGCGATTCGCACGGGAATGCCCTCACCTTTCTTCTTGGTGAGGCGCAGC
TCGGCGGCATAGCCGATGCCGGTGCCATCGGAACGACTCGCCACGATTGAGGAGCCAAGTTCGAGCATCTTCTCGACAACCTCGCGTGGATAGCGCGCCTT
GATGACGTCGTAATCAACATCACCGCAACGGACGGAGAGCGCCAGGTAGTTCTCGGGGATGCGGTCGCTTATTTCTATGCCATGAAGCTCGCGAAGCTTCTC
CTCGTAGAACTTCACGCATTCCTGTAGGATGGGGCCGTAGGACAGTTCGTACTCGTCGGTGACAAAGTGGGAGCCCGTGTTGCGGAACTCGCCCACTACTTC
CATGTTCAGGCGTAGGGGGTCGTTTTCGTTGGTAACGACCTTCTTGAGGCAGTCAGAAAGCGACAGCGTGCGGTCGCTGTCCGGATAGTAAACGGCATCGTA
GCCAACCTCGCGGATATGGCGGTGTATTGGGCAAACGCGCGCGGAATCGTTCACGGGTATGAGGCGTCCAATGGGTTTCGATACTACAGGCCGTGGGTAGG
TGTAACCAGAGAGCAAGTTGCCGCTATGCTTATGGCGCATGTCGAGGGAGAGCCGGGGTCGGTCAGCGGTCTTAAGGGTTTTTCTGATTGGTCTTCTACTTC
AAGCTAGAGGGTTGTTCCACGGCAGCATTGTCATCACTTCATTGTGATTAATACTGGTATTTAGGCAACATTATATTATTAAGAGGTTGTGCCAGTGCTCATTTG
ATCTTATTTGCCTGAATGAGGTCTCATTTCGGCGCGGACAGTTGAGGGCGAAGCCCCATGGTTCAAAACTATAATATCAGCGTTTGATATATGCCTTAGAGGAT
GCTGGGGGCGAATGGTGAAAGAGTCGACAACGGATCTTACAACGGGAAGCCCTCTAGTAAGGTTGCTCGTCTTTTCCATTCCCCTTGTGCTTGGCTCGTTTGT
CCAGCAACTGTACAGTTTTGCCGATACCGTTGTGGTTGGTCGTCTTATAGGGGCGGATGCCCTTGGAGCTGTCGGCGCCACGTATGCCCTTTATTTTTTGGTG
CTTGGGTTTGTCCAAGGTGCCGGTATTGGGTTTGGTATTCCAGTGGCACAAAGATTTGGCGTGGGGGATAAAGAGGGCTTCCTTCGATACCTGTGGAACGGC
GCCTGGGTCTGCGCGCTGGTTTCTTTAGTGCTTACGCTGGCGACGGTTTTTGCAACCAGGCCTTTGCTGGTCGCAATGGGAACCCCAAAGGAACTCTTTGAT
ATGGCGGCAAGCTATATTGGGGTTGTGTTTGCCGGGATACCCGCAACGCTGTTCTATAACTACACCGCTTCTGTTTTGCGTGCGGTGGGAGATTCGCGGCGC
CCGTTTATCTTCCTTATGGTGTCCTGCGCCGTAAACGTTGCGTCCGATTTGTTTTTCATCCTTGTGTGTGGTTTGGGTGTTGTGGGCTCGGCTCTGGGAAACAT
TATAGGAACGGTACTTTCTGTCGTTCTTAATTGCTGGTGGGCCTTTTCTCGTGTGCCTCAGATTCGCTTTGACCAATCGGCTGCAGCATGGTCTACCAAACACG
CGCAAGACTTGTGCGCGGTTGGGCTGCCAATGGGGTTTGAATACTCCGTGTCGGCGCTTGGAGCCCTTGCGATGCAAGGGGCGATTAACGCGCTTGGGTCT
GCAGCGGTCACGGCGCAAACGGCGGGCGAAAAGATCCGGCAGCTCTTTACTATGCCCATGGAGAGCGTGGGCATGGCAATCGCCTCGTATGTGGGGCAAA
ACTATGGGGCGGGGCGCTTTGATCGAATACGGCAGGGTATATTGAGCGCGGCGGCGATTCAGCTTTTGTATTGCGCTGCTTGCTGGGTGGTCATCTTTGTCT
TCAAAGAGCAGCTCATTGGGTTTGTGCTGGGCGAGGAAGTCGATGCTGATATTGTTTCCGGGGCCGTCGACTATCTCTTTTTCGTAAGCTGCTTCTTTATGCT
CCACGGGCTTCTTATGGTGTTTAGAAACACCTTGCAAGGCATGGGCCATAGCGTGCAGGCGGTGGCTTCGGGCGTGAGCGAGTTTGCGGGTCGCGCTGCG
GGCAGTCTTGCAGCTGTGGGCGGCATGGGATTCGCCGGCATCTGCTTGGCAAACCCGCTTGCGTGGGGGTGCGCGCTTGTTTATTGCCTTGTTATGGTTGG
CCGCTTGCTTGCCGGTAAGAACGGTAAGAAAAAGGGGCGCGCAGCGGCTAGATAGCCGCTGCGCGCCCTTGGAAACTGGGCGTTTATGGGGTGATTGCGCT
AAAGGCTAGGCCTTATAGCCAAAATCCGGAATCTCGTGCCAGCGCTGGCGGAACCAGTGGGCAGCGAGCTTGGGCTTGCGATCGCGGGTGAAGAGGCCCTT
CTTGTTGCCCTCGACGCGCAGCAGGCTCTGGCTGGTCGCGAAGTCGGCGAAGTTCCACACCTGCTCGCCCACCACGCAGGGGAACTCGTCAAATGCCTTGA
GGTTCTCCTTGTAGTAATCGACCTGGTACTCCTCCGTGTACATTACGGGCGTGGTATTGTGCATGCCGGCGACCGTGTCGGCTCCAAATTCGGTAAACATAAC
CGGCTTGCCCACGCTCGTCCAAAACTGAAGCTCGCGGCGAAGCAGCTGGCAGGCGTTCTCGATATCCGGTCCGCCAAGATACCAACCGTAATAACGGTTGA
GGCAGATGACGTCGGAGAGCTCGGCGGTTACATCGTTTTCGGGGTGCACCGTCTGAACGCTGATCAGCGTGCAGGGGCGGTTTTGCGGGTCAAGTTCATGC
GCCAGGTCAAAGAGCGGCTTGAAGTAGTCGTACGCGCCGTTGTTGCGGTCGGAGTCGGGTTCGTTGGCGATGCTCCACATGACAACGCAGGCGTGGTTCTT
ATCGCGTGCGATAAGGTCGCGGATAACGTCCTTGTGATGCTCCTGCGTTTGCACGCCGTTCTGGGGGTCGAACGTCTTGATGTTCTCGCCGCCAAAGTTCGC
GCCGCCGCCAAAGCCAAGGTTGACGCCCACCGCCGTGGTCTCGTCCACCACAACAATGCCCTCGGCATCGCAAAGACGCATCATTTCCTCGCTGTAGGGAT
AGTGGCTTGTTCGGAAGCTGTTTGCCCCCTGCCACTTCATGAGCGAGATGTCTTTCACGTTCATGGGCAGGTTCTCGCCGCGGCCTGCCGGGAAGGTGTCCT
CATGCTTGCCGTAACCCTTGAAGTAGAAGGGCTTCTCGTTAATAAGGAACTTTGCGCCCTCGACGCGGACCGTGCGGATGCCGTAGGGCAGCTCGTAGACGT
CTTCGTCATAGGTTGCCTTGACGGTATAGAGGTAGGGTGCGCCGGGGTTCCATAGGTGCGCGTCTTCAACAACGAGCGTGCCTGTGGCGCCGGCGGATTCG
GCCACCTTGTTGCCGGCGGCATCAAGGAGCTCGACGGTGCAGACGGCATCCTCGCCCGCGGCCTCAACCTCATAGGTAAGGGTTGCGGTGCTGCCGTTTAC
CGCAGGCACAACCGTGATGTCTTCGATATGCGTCTGCGGCGTGGTGTAGATATATACCGGTCGCGTGATGCCGGCGTAGTTGAAGAAGTCAAAGTTGGGGTC
GTTGCGCTTCTTGGGCTCCACGTTGTCGCCGCCCATGACGCTGCCCATAAGATCGCCCATCAAACCGCCCTTGCTGCCAACGGGCAGGGTGGTGTAGTCGA
TGACGTTGGAAACCGCGACGGTGAGTAGGTTGTCGCCATCGCGAAGGACGTCGTTCAGCTGGACCTCAAAGGGCAGGAAGCCGCCCTTGTGCTCGCAGATG
AGCTCGCCGTTCAGATAGACCTTGGCGTGGTGGGTGACGGCGGAGAAGCGCAGCATCACGCGCTGGGCGCGCACAAAGGCGGGAACCGCGATGGAGCGC
TGATAGATCACCCAGCCGTAGTGGTCGCGAAAGTCGATGCCCTCCTTCAGGTCGTTATAGGAGGCGGGGACGGCATGGTGGTGGCGTCTTCAAGCGGGC
GGTCCTGCCAGCCGGCGTCAAGGCCCGTGCCGTCGTCCAGCTTAAAGTTCCAGATGCCGCTTAGGTTTTCGACAAAACGCGATGGCGTAGCGGATGGGTAC
AGCATGGATGCATCCTTTCGTAGAAGAACATGACGGAGCTGTTGTACCCATAAGTAACATATCAGATTCTTGAAGTCATCGATGTGATATGGCGGGAATGGGG
CGTACCTTCGAGTGGTTGTCTATTTACCGCGAGCGGTATACATGGTGTAAGGAATGCAGGTATTACCCATAAAAAACAAATAGCAATCCGCTCGCAGTGAATA
CGCGACCTTCGACTGGATTATTGACAGAAAACAAAAGATGTCTGAATCGAGTGGAACTAGCATATTAGCCATGAGAATCGCGATGCGTCGCGCATTCGCATGT
CAAATTTGTCCAAAAAGAAAGGCGACTGTTATGACAGAGCATGTTCAGCTCAAGGCGGGCGGATATCAAAAGATTCGCGATATCGATCCGCGACTTGTTTCTT
ATAACGTGGAGATGACCGAGGTCACCGGCGGCACGTTTTGGAAGCCCTATACCGCTGGGCAGATTGCCGGAACGGAAGAGTTTCCAAAAATGAAGGACTTTT
CTGATCTTGGCACCTGATGGCGATGTTTCCTCCCGCCGACCTGTCCAATAGTCGCCTGCGCGCCCTTGCAAAGGACCTGGGGCCCGCATGGGTCCGCGTA
AGCGGTTCTTGGGCAACCAAGACGTACTATGATTTCGACGGCCATACAAATGGCGTTGTGCCCGAGGGCTTTGAATCTGTGCTGACTCGCGAGCAGTGGGAG
GGCGTTCTCGATTTTGTAAAGGCTATCGGCGCCAAGCTGCTGGTGTCCGTTGCCAACTGTCCGGGCGTTCATGATGGCATTGGTCCGTGGCAGCCGGATCA
GGCGCGATTGCTCTTTGACTTTAGCCGTGACTACGGAGTTCCCGTCGATGCGGCGGAGTTTATGAACGAGCCCAATATCTTGACGGGCACCGGCTACGATCT
CCAAGACGGCTACACGCTCAAAGAGTTTGCGCGCGACCAGGACCTCTTCTTTAGCTTTATGCGCGAGAACTATCCGGAGATAGAGCTTGTCGGCCCCTGTGT
TGTGGGCGATGCGATGGCGACCATTTTGGGTGAGGACGAGGGCGGAATCCTTGGCGCTGACAAGGCGGACCTTATGGCGGCGATGGGCAAGGCGTTCTTC
TTGACCAACGACCTGCTCGACATTGTCCAGGAGCGTCCCGGCGTGTTCAGCTACCACTATTACAACGGAATCTCCGAGCGCGGAGCGGCGATGGGCGGCCA
CTGGGATGCGTCTCAGGCTCTGTCCGAGCAGTACCTGTCGGTCGCCGGGCAGTCCGCGCTCCTTTATGCACCCATGCGCGATTCTTATACGCCCGGAGCTC
CCATGTGGGTTACCGAATCCGGCGACGCGGGCTGCGGCGGCAATACCTGGGCCTCCACGTATCTCGACGTGCCTCGCACGGTGAATGAGCTGGGACTGTT
CGCCACCATTACCGACGGTGTGATTTTCCATAACACGCTATGCTCTTCTGACTACGGATATCTTGATCGCGAGACGCATGAGCCTCGTCCCAATTACTGGGCG
GTGCTTCTGTGGAATCGCATCATGGGCGTCGAGGTATTTGATTCTGACGAGGAAATTCGAGAGGGCGCCCACGTGTACGTGCATTCGCGTCGCGATGGGAA
GCCGGGTTACGCCGCCGCCATCATCAACAACTCCACAACGGACGCCACCGTGGTCGACCTGCCCAAGGATGCGGAGGTCTACGCGCTATCTTCCGATTCTT
TGCGTTCAAGCACGATCATGCTAAACGGGGCGCCGCTTGCTTTGGGCGAGGATGACTCCTTGCCGGAGCTTGCAGGCAGACACGTTGAAGCGGGTGCTTTT
GAAGTGGCGCCGGCGTCGGTGGTCTACGTGGTCATGTAGCTGGCCGCTTCGGGAAGAAGCCAAGGGAGCCGGGCGGATACTGCATTGCGGTGTCCGCCCG
GCCTTTCCCTGCAGGTGGTTTTTCGCAGCGTGCCTAGCTCGCAAAGTAATCCGGAAAGGCGGACGTGACAATTTCTTGCTCTGCATTCATAAGGTGCAGGTG
GAGACCGGAGATAAAGCGCGCCTCCTCTGGCCTGCGGTTGCAGATGGCGTCAAGGAGAAGCTTGTGCTGATCCATGATGGTTTCCCATTCAAGGCCGCTTAC
CTGGGTTCTCAGCCAGCGAAAGCGCTCAAGATCGGTGTTGTTCGAGTCGATCCAGCGCCAGATCTCATGCCTGTCGGCAATGATGAAGAGTCGCTCGTGAAG
AAGGTTGTCCAGATGGAAGAAGGCGCGGGCGTCGCGCTTCGCAACGGCTTCTTGCTGCTGGGCAAATAGCGAGGTGAGCTTTGCCTTGGCTGCATCGTCCA
CATGCGCGCAGCATTCAACCGTGATACTTTTTTCTAGATGTTCGCGCACGTAGCGAGCGTTTTCCGCCTCGCGCAGGCTGATGCGCGAAACGTAGGTGCCGC
TTTGCGGAACGGTCTGCACAAGGCCCTGTTGGCTCAGGCGTACCAACGCCTCGCGAACGGGGGTGCGCCCAATGCCCAGATTGTCTTCCATGGTTTTGACG
GATATCTTTTGCCCGGGGATGAGGGTGCAAAAGATAATGTCCTGCCTAATCTGGTCATAGGCGGACGTCTGAAGGTTGGGTGCGGAATTGCTATTAATCATGG
TGTAGCTCTCTGCAGGTCGATCCAGCTGGTCACAACGAACATCTTACACTTGTTATGCGGACGGCGCCGTTAAAGGCGGCAAAGCTGCCGTTTGCAGACGAG
GCGATCCGGTGGGGTCGAATGTCTGCTCGTGAAAGAAATACATCTGATATATTAGTTGTGCGGCGAGCTTGATGCAAAGGCGAGATCCCTTGCGGTCTTTCC
GCGCTATCCACTACAGAGGAGGAACGATGGTTTCACTTAAGAACTGGCAGGCCTCGAAGGACGAGCTGGAGGCATGGGCGTGCATCTCCCCAAGTTTGAC
GTGACGGCGGTGAAGGAAGCGGGCGCTCGCCAGCCGGTTTGGATCCACATGGGCGGCGGCAACCTGTATCGCGCGTTCCACGCGGAGGTCGCTCAGCGT
CTGGCAGATGCCGGCGAGCTTGATCGCGGCGTTGTGGTAATTGAGACGTTCAGCCCGTTCACCGTTGAAAACGTATACAAGCCTTATGGTTGCGACTCCCTT
GAGGTTGTCATGAACGCCGACGGCTCCCTTGATGAGCGCGTGCTGGCATCTACTGCGGACGCGCTGTTCGCCAACCCTAGCTGTCCTGAGGACTACAATCG
TACAAAGGAGTATTTCGCCAGCCCCGAGCTGCAGCTTGTGACCCTTACCATTACCGAGAAGGGCTACGGCCTGCGCGACGGCGCGGGCAACCTGCTGCCTT
ATGTCGAAACCGAGATCGAGCAGGGTTCCGATCATCCCGCCTCGACTATGGGTATTGTCGCCGCCCTGCTACTAGAGCGCTTCAACGCCGGCGCGGCTCCC
GTCGCCATGGTCTCCACGGACAACTTCTCGCGCAACGGCGAGCGTTTCCGCGCGGCCGTGCTCACCATCGCCGAGGGATGGCAGGCTGCGGGCCACGTGT
CCAAGGAGTTCGTCGAGTATGTTGCGGACGAGTCCCGCGTGAGCTTCCCCTGGTCCATGATCGACCGCATCACCCCCAATCCCTCGCCCGAGACCGAGGCC
GCGCTAAAGAGCCAGGGCTGGGAGGACCTTGATATCCGCACGTCTGAAAACGGTCCGGCATTTGCCGGGTTCGCCAACACGGAGGTCGCGCACTACCTGGT
TATCGAGGACAGCTTCCCCAACGGCCGCCCTGCGCTGGACGCCGGCGGCGTTATCCTCACCGACCGCGAGACCGCGGAGAAGGCCGACACCATGAAGGTG
ACCGCGTGCCTGAATCCCCTGCATACCTGCCTTGCCGTGTTTGGCTGCCTGCTGGGCTACAACCGCATTTGGCGTGAAATGGAGAACGAGGACCTGGTCGC
GCTTATCAAGCGCCTTGGCTATGAGGAGGCGCTGCCCGTTGTGGTCGACCCCAAGGTTATCGACCCCAAGGAGTTTATCGACGAGCTGCTGACCCTTCGCCT
TCCCAACAAAGCGCTGCCCGACGCCCCGCAGCGTATCGCAAGTGATACCTCGCAGAAGATTCCGGTTCGCTTTGGCCACACGCTGGCGGCATACGTTTCCG

ACCCGGAGAAGGATCCCGCGCAGCTTACCTGCATTCCGCTCACCATCGCCGGATGGCTGCGCTACCTGCTTGCCGTCGATGACAAGGGTGAGGCCTTCGCG
CCCAGCCCCGATCCGCTTCTTGCCGACCTGCAGTTCCAGCTCAAGGGCATCGCGCTTGGGGAGACCGACGCCGCCAAGGTGCACGACGCCGTCGCCCCCA
TCCTTTCGAATAAGGAAATCTTTGTGGTCGACCTGTATGAGGTTGGCTTGGGCGAGAAGGTGGAGGGCTTCTTCATGGAGTTCCTCGCCGGACCCGGAGCCG
TTGCGGCAACCATCAAGAAGCACGTGTAGCGACATCGTCCGCATGGGCCTTGAAACATCGCAAATAAGATGGAGAGGAAACTGATATGGAACTTACCTTCCG
TTGGTATGGACCCAATGAGGAGAAGATCACCCTGGAGCAGATCGGCCAAATCCCCGGCTGCCGCGGCATCGTGGGCACGCTGTTCGATATCCCCGCCGGTG
AGGTCTGGCCGCGCGAGCGCATCCACGAGCTGCGCCAGATGGTCGAGGACGCGGGCCTTACCCTCAAGGTCATCGAGAGCGTCAACGTTTCGGACGATAT
CAAAATCGGCACCCCGCTGCGCGACCAGCACATCGAGGCGTACAAAGAGACCATTCGTAATCTTGCTGCAGAGGGCGTCAAGGTTATCTGCTACAACTTCAT
GCCCGTGTTCGACTGGGTCAAGAGCGACCTGGACTATAAGCTGCCGGATGGCTCGTCCACCCTGGCGTTCATCCGCAAGAATATCCCGGATGACCCGCAGG
AGATCATCGACAACGTGGCCAACGGCTCGGGCGAGTTCACCCTGCCCGGCTGGGAGCCCGAGCGCCTGTCCCAGGTGCGCGCCCTGCTGGACGCCTACAA
AGATGTGGACGAGGACAAGCTCCGCGAGAACCTGGTCTACTTCCTCAACGCCATCATTCCCACCTGCGAGGAGTGCGACGTCAAGATGGCGATCCATCCGG
ATGACCCGCCGTATTCCATGTTCGGTCTGCCGCGCATCATCAAGAACCGCGAGGACCTGGACTGGCTCTGCAACGCCGTGGACAGCCCGTACAATGGCATC
ACCCTGTGCTGCGGTTCCATCGCGGAGGAGCCCGGCAACGACATCTACTCCATCCTGGCCGAGTTCACCCGCCGCGGCCGCATCCACTTTGTGCACATGCG
CAACATCAAGTACATCAGCCCGGAGGAGCCCGGCAACAAGGACTTCTACGAGGCCCCGCATCCCAGCGAGTGCGGCTCCCTGGATATGGTTAAGATGATGC
GCGCCCTGCACGACAACGGCTTACCGGCTATATCCGTCCCGATCACGGCCGCATGATCTGGGGCGAGACGGGTCGTCCCGGTTACGGCCTGTACGATCGC
GCGCTGGGCATCGCCTACATCAACGGCGTGTGGGAAGCCCTTGAGAAGCAGGATCGCTAAAGCTTAGAAAACGCGAGCGGTAATCCAAGAAATAGGCCCT
CCCAGAAACACGATGTGTCGGTTTTTGGGAGGCTTTCTTCAAAAATCAAAACTAACATACCAGTTATGGAGTAACGGGAACCGCCCACCGACAGGGATGGCAA
GGGCGCTCCCGTAGTTTTGCTCTAAAAGTAATATATAAGTTGGAAGGTAAACGATTTCGCATTCAATCATGGCGGTAATTAGTTTAGGAGGCTCCATGGGTAGC
GTGCGCATTTCCGGATTTGCCGACGAGATCAGCTCTGATTTTGACAAGCAGCTCGATGTGGTGAAAAAGCTTGGTATGAGCTATATTTGCCTGCGCTCTGCGG
GGACCAAGGGCGTGGCGGACTATTCGCCTGCTGATTTTGCGGACGAGCTGTGGCCCAAGATGCAGGCGGCTGGGATCGGTCTTTCTTCAATCGGCTCGCCG
ATTGGCAAGGTTGGCATCAACGATGAAGAGGGCTTCCAGAAGCAGCTTGTCAGCCTGGAGGGCCTGTGCCAGATCTGCGAGATGACGGGCTGCCGCTACAT
CCGCGTCTTCAGCTTCTTCATTCCCGCGGGCGAGGATCCCGACGCTTACTACGACAAGGTCATCGAAAAGGTCAAGCGCTTTGTTGAGATCGCCGAGCGCCA
CGATGTCATCTTGATTCACGAGAACGAGAAGGACATCTTTGGCGACATCGCGCGCCGTTGCCAAGAGCTCTTCGACGCCATCAAGAGCGATCATTTCAAGGC
GGCATTCGACTTTGCAAACTTTGTCCAGGTTGGCCAAGACCCGGTCGCCGCCTGGGATTTACTGCATGAGCATGTGGTCTACATCCACATCAAAGACGCGGT
CCATGGCAGCAATGAAAACGTCGTCGCCGGCACCGGCGATGGACATATAGAAGAGATTCTCAAGCGCGCCATTGTGGACGAGGATTACGAGGGCTTCCTCA
CGCTCGAGCCGCACCTGGTCATTTTCGATTCGCTTAAGATGCTTGAAACCAAGGACGTGAGCGACATCATCCGCGGCGATAAGGCTAAAGACGGTGAAGAGG
GCTACACCATGCAGTACAACGCGCTGGTGGAGATTCTCGGCCATATTGGCGCGACCGCGTCGTAAGAGCGCGCACGATATATCGGCTATAAAGTGAACGGC
CGCCGGAATGAAGCCTGGCGGAGAAAAGGAGCAAGAAATGGAAAAGGTGCGTTACGGCATCATCGGCGTCGGTAATCAGGGCGGTGCCTACGCTGGCTTTC
TGACCGGGACGGGCAACGTCCCCGGCATGCCGGCGGCCCCCGTGCCCCCCGCATTGCCGCGCTGGGCGCCCTGTGCGACATCGATCCCCAGAAGGAAGAGA
TGTGCAAGGAGAAGTATCCTGATGTCCCGTTCTACAAGGACTGGAAGGATATGGTCGCCTCTGGCGACGTCGACGCCGTCATCACCACGGTGCCCCACTACC
TGCACACCGAGATCGCCATCTACTGCCTCGAGCATGGTATGAACGTTCTGGTCGAGAAGCCCGCCGGTGTGTACGCCAAGTCCGTCCGCGAGATGAACGAG
TGCGCCGCTGCCCACCCCGAGGTGACCTTTGGCATCATGTTCAACCAGCGCACCAACAAGCTGTATCAGAAGATTCGCGAGATCGTCGCTTCCGGTGAGCTT
GGCGAGATCCGTCGTTCCAACTGGATCATCAACAACTGGTATCGTCCCGATAGCTACTATCGCCTGAGCGACTGGCGCGCCACCTGGGGCGGCGAGGGCG
GCGGCGTCCTGGTGAACCAGGCTCCCCACCAGCTCGACCTGTGGCAGTGGATTTGCGGTATCCCCACCACTGTTTACGCCAACTGCATCAATGGCAGCCAC
CGCGACATCGCCGTCGAGAACGACGTTACCGTTCTGACTGAGTACGAGAACGGCGCCACCGGCTCTTTCATCACCTGCACCCACGATCTGCTTGGTACCGAC
CGCTTTGAAATCGACTTGGACGGCGGCAAGATTGTTGTCGAGGACAGCAAGAAGGCCTATATCTACCGCTTCAAGGAGACCGAGACCGCCGTCAACGCCAG
GGATATGGACTGGATGCAGATCGCTATGCTGACGTCTTCCAACGGCAACTCTGATGACAAGATGTTCGAGGTCGAGGAGTTTGAGAACACCGATGGCTGGGG
CTACCAGCACACCACGGTTATGGAGAACTTCGCCCAGCACATCATCGACGGCACCCCGCTTCTGGCCCCCGGTTCCGACGGCATCAACGGCGCGCCTGG
CAAACGCCATCCAGCTCTCCGGCTGGACGGGCGAGAAGGTCGCTAACCCGTTGACGAGGACAAGTACCTTGCTGAGCTGAACAAGCGCATCGAGGCTGAG
GGCAAGTTCCCCGTTCGCGAGTAACACGAATATGGGCATGAGGAGGCCGGAACAATGAAACGGGCGGCGATTGTGGGTATGGGAGCGATCTATCCCATTCA
TTCTGCAGCGATTCAGGCATTAGACGGTATAGAGCTTGTTGGCGTTTGCGATATCGATGCCAAGAAGCGCGCAGCCGCGCCCGAAGGCGTTCCGGCCTTCG
AGAATGTACGCGAGATGGTCGAGCAGACGCATCCGGACTGCGTGCACGTTTGCCTGCCTCATTACCTGCACTATCCCATTTCAAAGCAGGTTGTCGAGATGG
GCGTGAACGTGCTGTGCGAAAAGCCGGTCGCCCTGAACGGCCGCGAGGCGCTTGAGTTTAGGCGCCTTGAGCAGGAGCACCCCGAGGTCAAGATCGCGGT
GTCGCTCCAAAATCGTCTCAACGAGACGACCGAAGAGCTGGTTCGCATCATCGGCAGCGGAGAGCACGGCAAGGTCATGGGTATTCGCGCCGAGGTGCCAT
GGTATCGTCCCCTTGCGTATTACCAGGCCGGGCCGTGGCGCGGCAGTTGGGACCAGGCGGGCAGCGGCGTCATGATGAACCAGGCGATCCACACCATCGA
CCTCATGTACCTGCTGGGTGGCCCCGTCCAGCGCATCAAGGCCTCTGTAGAGCAGATCCTTGATTACGGTATTGAGGTCGAGGACACCGTGAGCGCTCGCTT
CCAATATGAAAACGGCGCTATAGGCCATCTGTACGCTACCAATGCAAACTTCAAGAACGAGGGCGTCAACATCTCCGTCGATCTGGAGTGCGCGAGCTTCCG
CATGCGCGACAACGTTCTCTATGAGGTTGGCGAGGGCAACGTCGAGACCAAGCTGGTCGAGGACCAAAAGCTGCCGGGTAGCAAGTTCTACTACGGCGCGA
GCCACAAGAAGCTCATCCGCGGTTTCTATGATTGCCTTGAGGATGGCTCGGACGAGTATATCCACGTGCGCGATGCCTACATGAGCGTGCATCTAATCGATG
TCATCAAAGACTCCGGTCTTACCGGCTCATGGGCGAATGTATAACAAATGCTTTATAGGATCCTGCAGCGATTTGCGGGGTCGAGAAAGGAAGAGCTATGGCT
GAAAAGGGATTGATTGGCGTTCAAATGAGCACCATCGCGCCTGCGAAGATGCCTAAATTTGACGCATACGAGGCTATGGGCAAGCTGAGCGACATCGGCTAC
CACTGCGTTGAGATCTCCCAGGTTCCCATGACCAAGGAGAACGTGGATGGTTTCCGTCGCGCCATCGATGAGCTTGGCTTGAATGTTTCTTCCTGTACTGCAT
CGGTTGGCCCGCTTATGCCCGGCGTTCCGGGTGAGACCCTGAGCGATCCCGATGATTTCAAGAAGATCGTCGAGGATTGCCATGCTCTCGACTGCGATATG
CTGCGCATTGGCATGCTGCCCATTAGCTGCATGGGAAGCTTCGAGAAGGCCATGGACTTTGCCGCGCAGGCAGAAGAGTGCGCTGCGAAGCTGAAAGAAGA
GGGAATCGACCTGTACTACCACAACCATCACGTGGAGTTTGTGCGCTACAACGGCGAGTACCTGCTGGACATCATCCGCGACCACGCTCCGCACGTGGGCT
TTGAGCTGGATACCCACTGGATTCACCGTGGTGGCGAGGACCCCGTGTCGTTCATCAAGAAGTATGCGGGTCGTATTCGTCTGCTTCATCTCAAGGATTATCG
CGTTGTGGAGCCCAAGTTCCCCGAGGGCAATTTCGATCCGGCTGGGTTCATGCAGGCGTTCACGAGCAATATCGAATTCGCAGAGGTTGGCGAAGGCACGC
TCGATATTAAGGGCTGCATCGAGGCGGGTCTTGCCGGAGGCGGCGAGTACTTCTTGGTGGAGCAGGACGACACCTATGGTCGCGATCCCTTCGAGAGCCTT
AAGATCAGCCATGACAACCTGGTCAAGCTTGGCTATGAGGATTGGTTCTAATCGAGCTAGACGTTCATCGGCATCAGGATGAATCAATAAGACAGCATGCAAA
GGCGTGGCCTCAACGGTCGCGCCTTTTTGCGTTTGGGCGCCCGGGCCCCGTGATGTATGACAAAAACTCCGGCCCGCATTGGTGGTGCGGGCCGGGGCGG
AGAAAGGGGGGCGAATTATCGATGCGGATATGGACTAGTACGTGAGGCCAAATCCAGCGCGGTAATAGTTGCCGTCGGCGTCTTGGTGGGCATAGCTGCCA
CCGGGCACGTTAGCTAGGATTTCGGGGTCGATGTAGCGATCCTTATCGATGCCGGGGACGTCGTTGGGCGACGCGCAGACCTCACGGACATGCCCGTCGAT
CTCGCGCTCGCTGATAGCGATGACGTCAGGCGAGGAAACCATGGTGAGGGGCATCTTGCCAGTGGGTGCAAACGAACCCGTAAGTACGTCGAGCTGGGCG
TTGAGCGCGTTATCCATCGCGACGCCGCTGACGGTGTACTGCATGGTGAGCGCGTCAACGTAGGGCTCAAGCGTGTCGAGCAGCCAGGGGTTATTAACAAC
AAGCGTGGCTATTACCTTGCCGCCGCTCTCGTGAACGGTATGGGCAATGCTGGCGATGCGTTCCACGCCGTGCAGCGTGCTCACCTTCAAGATCTCGCCGG
TCTTCTGCTGGCTCTTGTTGGCTTCGCGCGCTTCGACCTCGATATTCTCTCCCATCTCGCAGGACGGCATCGCGGTTTGGTTGAACACGATGTTGTTGAGGAT
GGGCCAAACGTGGAGGTATGCGAAATCGGCCTCTTCGGGCGTGTCGACCACGGTGAAGCCACGCTGCTCAAACAGCGCTTCGATCTTATTCTGAAACTCGTC
CACGTCTCCACCGCCTGCTGCGGCGCCCATGGCCTGAGCCAGGGCAAGGCCAGGGTCATCGCCCTTCAGGCAGCACACATAAACTTTTGCTCCAGCAGCGA
GGGGGAGGGTTTGGTTGTGGTTCTTAGCCAAGACAACGGCCTTTTGATTGGCAGCGTAAGCAGAGCGCTTGGCGTTCTCCAGGTTGTCATGACGAGCTTGGT
CGGCCTTATGGATGTCCAGGTAGGGATTGTCCACGCGTTTGGTGCGGAACATGGAGAGCAGGCGACGGTAGTTGGCGCGATCAAGGTCTTTCTTGGGCAGC
AAGCCCTCCTCGACCGCCTTTACGATGTTCTCGGGGTCGATGGGTCCGCCGATAACATCTGTGCCGGCGCTGATGGCTTTGGCGTAGCGCGCGGGGACCGA
CTCGTCCTCAAGACCGTAGTTCTGAACGTAGATGATGCCTGAGTCCGAGTTCACGTATCCATCGAAGCCCATGACGTTGCGGGCGAGCTCAGTGATGAGCCC
CTTGTTGTAGGTCGAGGGCACCTCCTCGGTCGAGGTGACATGGCCGTGGTAGGTCTGCTCGACGCTGCGGCCGTCCTTGGTGGGTCGGGAGTAGTCGGGC
ATGATAGCGGACGCCTTGGCGTCAAATGCTGCCTGGAACGGAGAAAGGTGATACTTCTCAAGCGATCCTTCGGTGGGATACAGGCGCCATTGCCCAATGGG
GACATGCGGCTCAAAGCCGTTCTCGCTTGGGCCGTCGCCGGGGAAGTGCTTCACGGTGAGCACGACGGACTCCTCGTTGAGGCCGTCTCCGCCGTGCTGG
TAGCCGCGGATAAGGGCATCCGTGATGGCGGACGTGGCGTCGACGCACTCTCCGTAGGTTCCGCTAATGCGGGGCCAACGGGGGTCGGTCGCAACATCCA
CCTGCGGGCCGTACATAATATCAAGGCCGGCAGCCTTCATCATCTTGCGATCGGTGTCCGCCATGTTGAAGACAAGGTCGGCGGAACCATCACCCAGAACG
GCGGCGCCAATTCCCAGCTGGTCGGGATAGCCTATGTTGATGGGGTTGGTATGCAGCGAGAAGGGGATCGCCACGCCGCCCTGGCATGCCTCGTGCTCAA
GCATCTGGGTCGCCACGTTGGTACAGCGCGACCATGGAGGCGTCGGCTGCCATGTCCAGGCGGTAGACGCCTGCGGTCAGCTTCTTTTCCAGGATATTG
CTGTCGTCGATGCTGGACTCCATTCCAAGGAAGGAGGGGGCGTTGAGCGCCTCAGGAGAACCCTTTGCCGGCATGACTTTACCAACGATAACGTTTCCGTCT
TCATCCGTTGCCTCGCCAAGGTGGCGGACAACCGGGTTGTTCCACAGGTTGTTCAGCACAAGTCCCGCCTGCTGGTCGAGGCGAAGGTGGGAAACCATGTC
CGCCGCTCGCTCTTCCGCGCTGAGGCGCCAATCCTCATAGGGGGAGAGGACGCCGTCGTTGTCCATATCCTTGAAATACAGGCCATCTTGAACAATGACGCC
GCAGGTGGTGACTCCAATGTCGGGGCCGCTATCGTTGTGGTAGATCACGGGCTCAAGGTAGGCGTCGCGGATGCAATCGCCTTCTTCGAGCTCATAGGGCG
CTGGTATATAGCCATCTTTTAAGGTCGGTTCAAATTCTTTCTAGCATGTCAGTCCTTTCGATTGAATGCTCAAAGGTTGCTGTTGATTATTGCCGCA
GCTATTGAATAGGCATATCTACTCTAAAGTGATATATCAAGCTACTTATAATTGCAGAAAATCAGCGAATGGTAGGGTTTTCTCTGTGAGCGAATGACAAAGCG
GTGTAAATATGATACATCGGCAAACCAGTGGCTACCACGAGTCAGACTACCTAATCTAGCTGCTGAAAGATAGAGCTACATATCTTGTTGATTTGAGCTTAATC
GTACGCTGGCAGACAAATTTCCACCGCCTACGGAACAACTGGCATATCTCAAACATACCAGTTTCAGTAATTAGTATATTAGTTTTGCTCTTCAAGTTTCTCGTT
GACGCACATTCCTCGCTCCGCGCGGTTGGGCGAATCGGTGTCAAGGGGAGCAAGCAATCGTTCTGCTCATGCGATGTACGCGTAGCGCATGGGCGAAGATG
CGGGAATTAACAGAAGGGGATGGAAAATGGCTAAAGTTGAAAAGGTCGATAGTAACGGCCTTCTTAAATTCGGATTCAAAGATGTTTTTGCCTACGGATTGGG
CGACTTTGGATGCAACATGAGCTTTGCTCTCAAGAGCTTCCTCATTGTCTATTGGACCCAGTTCATGGGTATCGATAGCATACTTATGGCATCGCTGCTGCTGA
TCGTTCAGGTTTGGGATGCTATCAACGACCCCATCATTGGTGCCGTGGTAGACGCTGATACCCACAAGTACCGTCGTAACAAGTTCCTCACATATGTATGGGC

GGGTTCCATTGGCCTGCTGGTTGCTGGTGCCCTGTGCTACATTCCGTGGCCCTCGGCTCCCTACGTGGTCAAGTGTATCCTGTTCGTGGCGGGATACATGCT
GTGGGATGCGTTCTATACGGTTGCAAACGTTCCCTACGGCTCGCTGCTTTCTCTTATCTCTAACGATCCCGTTGAGCGCACCAAGCTCTCTACCGGCCGCAGC
GTGGGCAGCATGGTAGGCGCCCTTACCACCGCTATTCTGCTGCCGTTCTTCATCTATGATGCGAGCAACAACCTCCGCGGTGAGGTCATGTTTGGCATCGCC
TTGGTCATGGGCGTCCTGGGGTTCATCGCTTTCCAGTACATGATTCGCAATACCGTCGTTCGAATACCGTCGTTCGCGTTCAGTCCGCTGCCGCCCCTTCGA
CCGGGGGCAAGAAGGCCACCAAGAACCCCCTCGTGGGCATTGGTCACTTCCTGCGTAACCGTGCGGCAATCGGTGCAACTATTGCCCCCGTCGGTACGTTT
ATCTCTATGTACGGCTCTCAGACCGCAAGCACCATTTTGTTCCAGTCCTATTTCCAGAATGCTCAGATTTCCGGTCTGGTTAGCATGCTGTCCTATGTTGGCCT
GTTCGCATTCATTCCGTTTGCGTCGAAGATCGTCGATAAGTGGGGAAAGAAAGAGTCCGTTACCGTTGGCATCCTGTTCTCTACCGTGGCATATGCCGCTATG
TTCGTACTTCCCATCACTCCCGATTCGAACGGTCTGATGATCTATATCGCCTGCCAGCTTGTCGCTGCCGTTGGTCTCGGCCTCAGCCAGTGTGTTTCCTGGT
CCTTCATGGCTGACGCTATGGACTACGAGGAGTGGAAGTTCGGCACCCGTAACGAGGGCACTACCTACGCCATGCACTCCTTCTTCCGTAAGCTTGCCCAGG
GCGTTGGTCCTTCGCTCGGTTTGGTTTTGGCTACGATGTTTGGCTACAACGAGGCCCTTGGCGCCGCGCAGACCGCTGCTACGGCATTCAACATGCGTATGC
TCGCGCCTGCGATGCTGCTGGTAGGCAACATCATTATGCTTATCGGCTATGGATTGGTGTATAACCTGGACAAGAAGACCGTTGCACAGATGGAGCGTGACC
TTGGCAAGAAGCACGAGTCTGCCGTTAACCTTAACGCCGCTTTGGGCGAAGACTAGTTAGTGGTTACTTTGCTGCGGTCGCATTGACATATCGCATAAAGACG
GGGTGGTGCGCTTTTGGCGGGCCACCCCGGTTGAAGTTAGAGGCCTTGTGCGTAGGGGCAAGGCGTAGGAGAAAGGGCAACCCCATGCTGCTTAACGATG
ACTTCCTGCTTACAACCGAATGGGCAAAAAAGCTCTACCACGCCCATGCGGAGAAGATGCCCATTATCGATTACCACTGTCATCTTGATCCCCAGCAGATCTA
CGAGGACAAGCATTTTGAGAACCTCACCGAGGTATGGCTGTTCGATAACGGTGCGGGCGATCACTACAAGTGGCGCTTGCTGCGTGCGAACGGTTCGGACG
AGAACTCGATTACCGGCGATGCGGATCCGTACGCCAAGTATCTCGAGTTTGTAAAGGCCATCGAGCGCGCAATCGGCAACCCCATCTACGAGTGGAGCCAC
CTCGAGCTGCGCCGCGTGTTCGGTATCGACCTCACTATCAACCAAAAGAACGCGGAAGAGATCTGGAATCGCGCCAATGAGATCATCGTTTCCCCTGAGTTC
TCGGCCAAGGCGCTTATTCACAAGTTCGATGTGAAGTGCCTTTGCACAACGGATAACCCCGCGAGCGACCTTGCGTACCATAAGCTTCTGGCCCAGCAGGAG
GACGAGAACGGCTTCAAGGTTCTTCCCACCTTCCGTCCCGACGACTTGATGGCCGTGGACGTTCCGGGCTTCGCCGCCTACGCCACCAAGCTTGGCGAGCT
CGCCGGCGTAGATGTTTGCGATTGGGAGAGCCTCAAGCAGGCCGTGGCACAGCGTGTGGACTATTTCCATGAGGTTGGAGGCCGCTTTGCCGATCACGGCG
CCAATTCCTTCTTCTTCCATAAAGCGGACGAGGACGAGGTTGCCGCGATTGTTGCCAAGGGCCTTGCGGGCGAGGCTCTGAGCGATGAAGAGGTTGGTGCC
TATCAAACGGAGCTAACGCTTTTCTTGATGGGCGAGTACGCGCGCAATGGCTGGACGCTCCAGATTCATGCCAACTGCCTGCGCAATGATTCGACGCTTGGT
CTTGAGCGCTGCGGCAAGGACGCGGGCTTTGACTCGGTGGGCGACCAGGCGGGGCTTGCCACCCAGCTCCGCAGCCTGTTTGATGTCGCCGAGCAAGCCG
GTAGCCTTCCCAAGGTCATCCTGTACAGCTTGAATGAGAGCGATTTCCTGTCGCTCGCTACGCTGTGCGGATCTTTCCAGGGCGGCGGCATTCGCCAGCGCA
TGATGCTCGGCTGCGCATGGTGGTTCAATGACCAGTTCGACGGTATGAAGAAGCAGTTGACCATCTGCGCGCAGCAGGGTCTGCTCGGCAACTTCACCGGC
ATGCTTACCGATTCTCGTAGCTTCCTGAGCTATCCTCGTCATGAGTACTTCCGCCGCGTTCTGTGCCACACGCTTGGCGAGTGGGTTGAGCAGGGCCGCGTT
CCGGAGGACGAGGAGTACCTGGGAGCGATTGTCGAGGACATCTGCTACAACAACGCTCATGATCAGTTTGGCTTCTTTGAGTAGGGCGCAGCTATCGGCTCA
AGAACCGTCGATTCATTCATAACCCGGAGGTCGCAGGTTTAAATCCTACTCCTTCGACCAAGAATTCGCAGCTCGGAGATCCAACGGTCTCCGAGCTTTTCGC
ATTCGTGCTACGCGCGGGGCCATATCCCTTTCAATGACTTTTTCAACAGATGAGGCGCTCTCAAGAAGGGTGTTGCTATTAGAAGAATGTGGCCATGAAATTA
CAAAGAGCGATGCGTAATGGCCGGAGCCTCATGCAAATAGCGGCCATCTATACGTGGCAAAGCGCGGGTTGGGTGTGCGAGCCACGCCGCAACGAGGCCC
CCGGCCAAGGTCATCATGCCTCGCGGTTCGGGAGGGCTTGCGGTTGGGAGGCCCCGTCGGTCGCCGCGTAGATCGCGTCGTGCAGAAACCGCGCGGCAC
CCTCTCCGCAAGGTCCGTCATAGTAGGCGGTAAAGCGCTCGTCCGCTATGTACATATCTGCCATGTTGCGGTGCGCCCGCGCGGAGTAGATGCCGTCCGGC
CAAAACGCGCGCAGCCAGGTTGCGTGCATCTGCGCCAGGCACCGCGCCGCGGCATCGGCTGGGTCGCCCTGCTCCATCGCGCGAGAGAGCTGCTTTAAAA
TTTCCTCGCCCAGCTCATTGACGTCATCCCATTCCTGCTGATCCATCTGCAGCAGCTTCTCATTGGCGGCATTCACGGTTTTGTCGCCGTATTTAGCACGAAC
CTCCTGGCCGTACACGCGCTCGTTCTCGTCCACGGCTTGCTGCTTCAAGGCCTCAAACTTCTGCCTGTCGCTCTGCTGCTTGCCCAACTTGTTCTGGTTCTTC
ATGTCGCGGTGCCCTTCTAAGCGGTCGATGGTATTTTCCACGGTGAGGATAAGCTGGTCTAACTCATGGCGCCTGCGGTGCAGGGCGGAGAGGTGCTCGCG
CATAGCGGACGCCTGATCGCGGCCATCCGCATCCAGAAGCGTCGCGATGTTCGCCAGCTCAATGCCAAGGCCTCGGAAGATCAGGATCTGCTGCAGGCGCT
CGACGTCTGCAGGGCGATATACGCGGTAGCCGTTTTCACGCCGCGCCGGCACTAGCAGGCCAATCTGGTCGTAGTGGTGGAGCGCTCGCACGCTCACTCC
GCTGATGCGCGCCAGCTGCGCGGGCGTGTAGCCGCCGCCGTGAAGCTCGCCGCCCCCGCCAACGTTGCTGCCGTTATGGCGAGCGGCGTCGTAATCCTCA
AAGTTCCCAATCATGACGTCACCTCCTTGATACGGCAGTATGCGCTATGACGCGGCGAAAGGGTCAAGGCTTGCGCAAAAGTTTGTCCGCACGAATAGCTAC
CATATAAATAAGTTATATCCGCACGCGCCACATTGGTGCGGGCAAGCGCGTTCAAGAGGCTTCAAGGGGGGGCGTCTTATGGGTTTGGGGGACGATATTTCC
AAGCGCACGTTCGACAACTACTTTTACGAGGAGCAAGAGCGCGATACGCACGCGGCGGGCGGCAGCAACAGCGATCTTGGCTGCGGCGGCGTTCTCATGG
CGACGGTGCTGATCATAGGTGCCATCGTGATTGGGTTCATGCTGATCATCTGGTTCCTGGAGAGCAGGGGGTAGGGGCTCCTTGCGCCCAACAGCCCCCAT
CCTTCCACGCATAAGTCCATCCGTGGTTCCAACTGCGACCTTGTCCACGTGCCCGTCCGCGTGCCCGCCCGCGATTGCGTCTGCGGCCTGGAGACCGCCAG
CGGCAAATGCGTTGCAATAAACCTCTATTCCAGAGGATAAAATGAAGCAAGTGTGTATACGCCGCGGCGGATTCTCCGCTGTGGACTTTGGGAAGGAATCAA
CATGAAGCGCGTTTTTGGTGTCGACCTGGGCGGCACAACCATCAAGTTCTCCATCATTGACCAGGCTGGCAATTTCATCGACCGTTGGGCCATCCCTACCAAT
ACGGAGAATGGCGGCGCGAACATTCCGGGCGACATCATCGAGTCCCTTAAGAGCAAGATGGCGACCCTGTTCGAGCGCGGCGAGACCATCGAAGCCATTG
GCGTGGGTGTCCCCGGCCCCGTGAATGGCGACGTCGTGGAAGTCGCCGTCAACCTTGGTTGGAATAACATGCCCCTGGGCCACCTGCTGCAGGAGGGTAC
GGGCCTGCCCGTCACCCTGCTCAACGACGCTAACGCAGCGGCTTTGGGCGAAGCCTGGATGGGCGGCGACTCCGCGTTCCGTGCCAGCACGGTTGTCTTT
GTGACGCTGGGCACCGGCGTGGGCGGCGGCGTTATCGTGGACGGCAAGGTTCTCAACGGCCAGCACGGCTGCGCGGGCGAGGTTGGCCACATGCCCGTT
ATGGCGGCGGATACCCGCCAGTGCGGTTGCGGAAAAACCAACTGCCTTGAGTGCTACTGCTCTGCCACCGGTTTTGTGAAGACCGCCAATGATCTCTATGAG
GCGGCTGGCGACACGCGCGCCTACACCGAGGGCAAGGACGTTTTTGACCAGGTGGCTGCGGGCGATGAGATCGCCATCCAAGCGCGCGATATCACGGTCG
ATTACCTCGCGCGCGGCTTGGCCGCGATCATGAGCACCGTCGACCCGCAGGAGGTCATTGTGGGCGGCGGCCTGTCGCATGCGGGCGATTTGCTTATGGG
GCCCTTGGCAGAGAAGTTGGACGAGTACGTGTTCCCCGGCATGCGCGGCAAGTACGTGCTGCGTCGCGCCACGCTCGGCAACGATGCCGGCGCTCTGGGC
GCTGCCTACCAAGCGTTCCAGATGCTCGCCGAAGGCGCGGGAGAGCAGATGATCTAAGGCGTCTGGCTCCGCGGTGAAGTTCTGCGGCGCCTCCGTAAAAC
GCTGAGGCTGGCATTTCGTTCGCAGTCATTATTGTTGCTGCCATCTACGTGAAAAAGCCCGGGTGCTCATTGCTTGTATAAGTGAGCGCCCGGGCTTTGTTGT
GTAACGTAGCCGCATGCTGCGGCTTGCTGTCATATGGGATGGCGCGGTATTTGCCTCTTTAAGGGCGGGAGGGTCGTTGTAGTCCTTATGCGCCACGCTGAT
TAAAATGATAAAAATCTAAGAGACTATGCATATCATGAGAACGGTCTCCGATGGCCTTATGATTGTCTACTGGTATGTAACCAGTAATACCAAGTTGTTATTTAT
GCAGGTACAAGGTTGTAAACATATTATCTGGTATGTACCAGTGAGCAACCGTTTGTCGCTAATTTGGAACCGTTTGCCTATCAGCCTAATTGTATTTTCTTTTGC
TTCCGTAGGATGTACACGTTCTATATGGGTGGATTTTATCTTACGGATGTCAGTGACTTCTTTCTCCAAAGTTTAGCTTTGAGCGCTAGGGGGACATCCAAAAA
TCCCAAACATGGAAAGGGAGGTATGGATGAAGAAGTCTAGGTTGGCTACAGCTGCCGTTGCAGCGGTTGTGGTTGGAACCATCGGTTTCTCTGTCCCCGCGT
TCGGCGCGTCTGGATATGGCGCCGTTAACGTGGAAAAGACCGACAACGTCGTAACTATCGGTAACGACGCTCTGTCCCGTACGTTTAACATCAAAGACAACA
AGCTGAAGCCTGGCAAGATCGACAACAAGCTTGCGGGTAAGGGCACCGCTTTTGATCCTAAAAGCGGTTCCGAAGAGTTTGTTATTCAGCAGCTCACAAAGG
CAACCCGCGTAGAGCCCGATGAGGGCGTGCTGAGCAGCGTCATGCCTGGCGCCATGGGTGCTCGCGTTGTTGTTAGCTCGACGATGGTCGAGGATGGCGT
TTCTGCTGAAGCGGCGATCGATGGAAAAGCCGAAACCTACTGGTGTTCAAACCTGAATACTTCTGGCGTCTCTAACGAGTCGTTCGTCTTGGACTTGGGCTCT
GCTAAATCGGTCAAGAAGGTTAAATACACTCCGCGTTACGACCAGAATGCAAAATACAACTGCACGGGTCGTATCTTTGATTATGCGCTCGATTACTGGGACG
GCTCGAAATGGGTCGAGGTAAAGACCGGCGAGTTTGGAAAAGATGCTGACAAGGGTGAGCAGGAGATTGTCCTCGACTCTGAAATCAGCACGCGTAAGCTTC
GCCTGCGCGCAAAGACGAGCTATTACCCAGGCGATAGCCAAGCTAACCAGGCAATGAACGTTGCGGAGTTCAACGCTTACGACTCTAAGGACGAGTCGGTC
ATTTCCCGTGATGAGACTGAAGCCTGGACGGTAACCTCGACGTCGAATCAGACGAATGACGGCAATGGTCCTCAGTCCCTTATCGACGGCAATGTGAATACG
TATTTCCATTCGGCTTATAAAGACGATGGCAAAGAGGGCACTGTCAGCCAGATGCCCGTGGACGTGACTTTCGATCGCGGCGAGTCGGGCAAGGATGTGGCT
TTCCAGACCATCGGCTATGCCGGTCGTAAAGCTGCGAACGGCGCGAACACCAACGGCAATGCGAAGAAGTTTGAGGTCTATGCTTCGAACGACAAGGATGG
CCTTTTTAACAAGGACAACAAGAAGGGTTCCTTCAAGGTCAATTATGCCGATGCGTATGAAGGCGAAAACCCTAAGATGATCTACTTCGGCCTCAGCGAAAAG
GTCGAAGACGCCCGCTATATCGGCGTCCGCTTCACCGAGGGCACCGGCGGGTCTTTTGCCGCTGGCTCCGAAATTGCATTCTACGATGGCAAGTTCACTTCG
GTTCCTGAGACCGATGAGGGAGCTATCAAGGCAAGCTCTCTTGAGATTGAAAACGTTGAGATCGGAGAAGCCGAGGGCGAATCCGGCAAAATGGTAACCTTT
GATTTCAAGCCTTTCCAGTTCGGATCCGATGCCGACGCTGAGCTTGAGATGAAGGTTGTCATGAATGACAACGAGCATTACATGCGTAAGTGGCTCGAGCTTA
AAACTGATGATAAGGAAACCCGCATCAGCTATATCGATGGCGAGCATCTCGACGTGTCCGACGCCGATGAGGAGTGGACTGTTCCCAACACCAAGGGCGTTG
TTCAGATGGACGCGGCGAAGGCCATTCTTGGTCAGCCTATCTACGTCAACGGTATGTTCTTTGGTTCCGAGTTCCCCGAGTCTGATACCCAGATCGTTGACGA
TATGGGTCGCTCCCGCTATTGGACCGGAAAGAATTTCGCCGACTTTGAGCGCGATGGCCAGTTGACCACGGATGGTGAGTACGTCTCTTGGCAGACCGTTTG
CGGTGCAACACGCTCCGATGGCTCGGATATGAATGTTGTGCAGGCTGACTTCTTCTCTTATATCGATGAGATTGCCAAGCCCAGTGACTTCCGCATTCAGTAC
AACTCCTGGTTCGACAACATGATGTTCATCGATGACGAGAACATCATCGAGTCCTTTAAGGCTGTCGATAAGCAGCTTTCTGAGACTGGCGTTCGTCCGCTCG
AGTCCTATGTTGTGGACGACGGCTGGAACAACTATCGTCCTGCAGCTGATCAGCTGATGGGTAAAGACGATATACGCAGAAACGGTGATGGCGTTAACGAAG
ACGGTTTCTGGACGTTCAACACCAAGTTCCCCGATGGCCTCACTCCTTCTAGCAGCCTGGTGCAGAAGCTAGGCTCTGATTTTGGCGTGTGGATTGGTCCGC
GCGGAGGCTATAACTACTATGGCCAGCTCGCGGGCATCATGGCTAAGGCTGGAACCGGCTCTGCTGCGGGCGGCTCGATCGACGTCGCCGACGCTAGGTA
CGTTAAGAACTTCCAGGACATGGCTGTTAAATGGATGAAAGACTACGGCGTTAACTATTGGAAGTGGGATGGCTTTGCGGACAGGGCCCAGTACAACCACTT
CTCGTCCGGTGACGGTGTGGTTGGCTATAGCGATTCCAATAAGCACATGTATGGTGGCCCCAATGGGTATTACCACTCGACCGATCTTTGGGAGAAGTGGAT
TGTTCTCTTTGAGAAGGTCTGGCAGACCGCTGATGCGGAGGAAATTAACGATCTCTGGATCTCTCTGACCTGCTACACCAATCCCAGCCCGTGGTTCCTGCAG
TGGTCAAACTCTGTCTGGATGCAGTGCACGGGTGACCGCGGCGAGCGCCACAACGGCACCTTGAACAACAAGATGGATAACATGCTCACGTATCGTGATGC
GTGCTACTACGATTTCATCAAGAAGCATGAGTTCCAGTTCCCGCTGGCAAACGTGTACAACCATGACCCCATCTACGGCAAAGAGGGCACGGGCATCAACGC
CGACTCCATGAACGGCGAAGAGTTCCGTAACTATCTGTATATGATGGGCACTCGCGGCACTGCCTTCTGGGAGCTCTACTATTCTGACTCCCTGCTCGATGAT
GAGAAGTACCTGGTCAATGCAGACTTCCTCAAGTGGGAGGAAGAGAACTTTGACATGCTTCGCAACGCCAAGTGGATTGGTGGCAACCCGGATACCAGCGTG

ACTTTGACCGCGGGTCCCGGGGACCAGGCGTCCGGGGAGCAGGACGCATATGGATTTGCTGGCTTTAACGCGGCGGGCAACGAGGGCATCATTTCCATGC
GCAATCCTGCTGCAACTGCTAAAACCATCGAGTTCACGCTGGATGAGGGTATTGGCGTTCGTGCGGACGGCACCTTCAAGCGCGTTATCGACCATTCTTATG
CACGTAAGGGCGCTGAGCTTGGAGCGTCGAACGAGACTTATGCGAAGGGCGATACCGTTGAGGTGACCCTGCAGCCTGGCGAGGTCCAGATTTGGCACCTC
TCCAAGAAGGGCGATACGGTTGCGCCTAAGCTTGAAAAGCTGTATGCCGAGGACAACTCAACCCTGCGCGTTCAGGCCTCCGAGCACGTGTGCAACACTACT
CTCGAGGTGACCGTTAACGGCAAGAAGGTCGATGCGACCGTTAAGGCCTACGCCGACCTCAAGACCTTCGACATCACCCTGCCCAAGGCCTACAAAGACGG
CGACAAAGTTGAGGTCAAGGCACTGACCGGCGCTGATTCTGCCGGCAACAAGCTCGAGGGTTCCATCACTCGCGTTGCCTACACCGATGGCGTGGTCGTGT
CTTCTGATTCCATCACTGGCACCACCGTTTCTCCCTCCAATCGCTCTGTTGAGGGCAAGAACGGCTTCTCCGTTGCTGCCACTGCTGAGACCGTGACCCCTG
GTGCCACCGTCGCGCAGGGCAATGCGTGGAGCCTTTCTATCAACAACGAAAGCAAGGCAGTCTTCACGGTTAACGGTGTCTCCGCCGTCTCTGATACCGAG
GTTAAGGAGCTTTCCACCATCACCGGCGTTCGCGAGAACAACGGCATGCTGAAGGTCTATGTCAACGGCGAGATCTCCGGCAACGGCTACGACGCAAAGAA
GACGCTCGGTTACAGCGTGAAGGCAGCTGCCATCACCGTTTCCGATGCTTCCGCGTTCAGCAGCGTTAAGGTCTACGACAAGTCCCTGGGCTACGACGAGG
TTCCGGCTTCTCCGCTGGCTGACCTTATCAAGACCGTCGAGGGCGTTAAGGACCGCGTTACCGGTGACTCCTGGACCAAGGCAGGCATGGACAGCCTGATC
ACTGCTGCAAAGGAAGCTCTCAAGAGCGATGCCGCTGCACAGAAGGTCGCTTACGAGGCTCTGCTTAAGGGTTACAAGTCCCTTGTTCCCGGCCAGGGCGC
TGTTGAGGTCACCAACGTTGCTAAGGGCATCAACCCCACCGCTTCTTGGCTGCCCGGCAGCAACAAGAATCCTAGCGATGTCATCAACACCGAGGGAGCATC
TCTGTCTAAGGCAACCGATGGTGCTCACGACAACGTGAACAACTACGGCATCTTTGGTAAAGATGCCCACCTGGAGCCTGCTTACATGCAGATCGACCTGGG
TTCTGAGCATGTCATCGATAGTGTGAATCTGTGGCGTTACTGGACCGCTTCTCGTGAGTACGCGGATACCGCGCTGGTTGTGTCTAACGATGCCGATTTTGAG
AACAAGCAGGTTCTGTACTACTCCTCTCTCGATGAGGATATGAACGATACGTTCAAGCTGGGCGAGAATCCCACTGAGCGTACGTATGCCGAATCTTCCGAGG
GAAAGACCCTCTTCGCTGCGAAGTCCGGCAAGACCGTAACCGCACGCTATGTCCGTCTCTATGCTAATGGCTATAAGGGCGGCGAGGGCAAGGAGAACCAC
GTCATCGAGCTGCAGGTCAACGGCTACAAGAAAGCCGACCTCAAGGATCCGTACGGCATGGAGAAGCTCGAGGCCCTCATGGCTGATGCCAAGAAGGCTGT
CGACAGCGGCAAGTACACTGATGCCTCCATCGCCAAGGTCAAGACTGCGCTCGACGTCCTCGAGCCCATGGTCGAGAAGATCAAGGCTGAGCAGAAGACTG
GCAACTACACCCAGACCCTGGGTGAGGTGAGCGATGCCCGCGCCGCGCTTGAGCTGGCCATCAAGCAGCTTGAGACTCGTCCCGTCGCTCCCGCACCGGA
TCAGGGCGAGGACAAGGTGACGGTCGAGAAGCACCCCGACGGATCCCAGACCGTGACCACCACCACTAAGGACGGCACCAGCACCAAGGTGGACGTGGAC
GCCAAGGGCAACGTGACCGCTGTTGACGCGACCGTTTCCAAGGACGCCGCCAAGGACGGCTCCGTTACCCTGCCCATGGATGCTCTGGAGCAGACCTCGTC
CAAAAAGGCGCCGGTCATCTCTGTTAAGGTTCCCGCTTCCGTGAACAAGGATTCCCAGCTTGTCGTGACCATTCCCGTTGCCAAGGACAAGGGTGACAAGAC
CGTCGATCCCGGCCTGGTTGTGATCATGGTGGACAAGGACGGCAAGGAGACCGTGCTGCCCAAGACCGCGGTGGGTGCAGACGGCGTGACCGTTGCCATT
GACGGCAACTGCGACCTTAAGGTTGTGGACAACGCCAAGAAGATGTCCGACGTGTCGGCTTCCGACTGGTTCGCCGAGGGCGTTGTTCCGTTTGCGACCGC
CCGTGGCATTGTGAACGGCGTGGAGCTTGCGGACGGCACCCGCGTGTTCGACGGCTACGGCAAGACGTCGCGCGGCATGTTTGTCGCCATGCTCCACAAC
CTGGAGCTCAATCCTAAGGCTGCCACCGAGGATAGCCTGGCCGATGTTCCGGCCGACGCGTTCTATGCCGGCGCTGCCGCATGGGCCGTTGAGAACGGGC
TGCTTTCCGGCGTGGATATGCCCGACGGCTCCAAGCAGTTCCAGGGCAATGCCGACGTTACCCGCGAGCAGGTGGCCGTGTTCCTGATGCGTTACGCGCAG
AGCCTGGGCATGGACGTTTCCGCGCGTGCCAAGGTGGACTTCCCGGATGGTGGGATCCCCGGGTACCGAGCTCGAATTCGCCCTATAGTGAGTCGTATTAC
AATTCACTGGCCGTCGTTTTACAACGTCGTGACTGGGAAAACCCTGGCGTTACCCAACTTAATCGCCTTGCAGCACATCCCCCTTTCGCCAGCTGGCGTAATA
GCGAAGAGGCCCGCACCGATCGCCCTTCCCAACAGTTGCGCAGCTGAATGGCGAATGGCGCCTGATGCGGTATTTTCTCCTTACGCATCTGTGCGGTATTTC
ACACCGCATATGGTGCACTCTCAGTACAATCTGCTCTGATGCCGCATAGTTAAGCCAGCCCCGACACCCGCCAACACCCGCTGACGCGAACCCCTTGCGGCC
GCATCGAATATAACTTCGTATAATGTATGCTATACGAAGTTATTAGCGATGAGCTCGGACTTCCATTGTTCATTCCACGGACAAAAACAGAGAAAGGAAACGAC
AGAGGCCAAAAAGCTCGCTTTCAGCACCTGTCGTTTCCTTTCTTTTCAGAGGGTATTTTAAATAAAAACATTAAGTTATGACGAAGAAGAACGGAAACGCCTTA
AACCGGAAAATTTTCATAAATAGCGAAAACCCGCGAGGTCGCCGCCCCGTAACCTGTCGGATCACCGGAAAGGACCCGTAAAGTGATAATGATTATCATCTAC
ATATCACAACGTGCGTGGAGGCCATCAAACCACGTCAAATAATCAATTATGACGCAGGTATCGTATTAATTGATCTGCATCAACTTAACGTAAAAGCAACTTCA
GACAATACAAATCAGCGACACTGAATACGGGGCAACCTCATGTCCGAGCTCGCGAGCTCGTCGACAGCGACACACTTGCATCGGATGCAGCCCGGTTAACGT
GCCGGCACGGCCTGGGTAACCAGGTATTTTGTCCACATAACCGTGCGCAAAATGTTGTGGATAAGCAGGACACAGCAGCAATCCACAGCAGGCATACAACCG
CACACCGAGGTTACTCCGTTCTACAGGTTACGACGACATGTCAATACTTGCCCTTGACAGGCATTGATGGAATCGTAGTCTCACGCTGATAGTCTGATCGACA
ATACAAGTGGGACCGTGGTCCCAGACCGATAATCAGACCGACAACACGAGTGGGATCGTGGTCCCAGACTAATAATCAGACCGACGATACGAGTGGGACCG
TGGTCCCAGACTAATAATCAGACCGACGATACGAGTGGGACCGTGGTTCCAGACTAATAATCAGACCGACGATACGAGTGGGACCGTGGTCCCAGACTAATA
ATCAGACCGACGATACGAGTGGGACCATGGTCCCAGACTAATAATCAGACCGACGATACGAGTGGGACCGTGGTCCCAGTCTGATTATCAGACCGACGATAC
GAGTGGGACCGTGGTCCCAGACTAATAATCAGACCGACGATACGAGTGGGACCGTGGTCCCAGACTAATAATCAGACCGACGATACGAGTGGGACCGTGGT
CCCAGTCTGATTATCAGACCGACGATACAAGTGGAACAGTGGGCCCAGAGAGAATATTCAGGCCAGTTATGCTTTCTGGCCTGTAACAAAGGACATTAAGTAA
AGACAGATAAACGTAGACTAAAACGTGGTCGCATCAGGGTGCTGGCTTTTCAAGTTCCTTAAGAATGGCCTCAATTTTCTCTATACACTCAGTTGGAACACGAG
ACCTGTCCAGGTTAAGCACCATTTTATCGCCCTTATACAATACTGTCGCTCCAGGAGCAAACTGATGTCGTGAGCTTAAACTAGTTCTTGATGCAGATGACGTT
TTAAGCACAGAAGTTAAAAGAGTGATAACTTCTTCAGCTTCAAATATCACCCCAGCTTTTTTCTGCTCATGAAGGTTAGATGCCTGCTGCTTAAGTAATTCCTCT
TTATCTGTAAAGGCTTTTTGAAGTGCATCACCTGACCGGGCAGATAGTTCACCGGGGTGAGAAAAAAGAGCAACAACTGATTTAGGCAATTTGGCGGTGTTGA
TACAGCGGGTAATAATCTTACGTGAAATATTTTCCGCATCAGCCAGCGCAGAAATATTTCCAGCAAATTCATTCTGCAATCGGCTTGCATAACGCTGACCACGT
TCATAAGCACTTGTTGGGCGATAATCGTTACCCAATCTGGATAATGCAGCCATCTGCTCATCATCCAGCTCGCCAACCAGAACACGATAATCACTTTCGGTAAG
TGCAGCAGCTTTACGACGGCGACTCCCATCGGCAATTTCTATGACACCAGATACTCTTCGACCGAACGCCGGTGTCTGTTGACCAGTCAGTAGAAAAGAAGG
GATGAGATCATCCAGTGCGTCCTCAGTAAGCAGCTCCTGGTCACGTTCATTACCTGACCATACCCGAGAGGTCTTCTCAACACTATCACCCCGGAGCACTTCA
AGAGTAAACTTCACATCCCGACCACATACAGGCAAAGTAATGGCATTACCGCGAGCCATTACTCCTACGCGCGCAATTAACGAATCCACCATCGGGGCAGCT
GGTGTCGATAACGAAGTATCTTCAACCGGTTGAGTATTGAGCGTATGTTTTGGAATAACAGGCGCACGCTTCATTATCTAATCTCCCAGCGTGGTTTAATCAGA
CGATCGAAAATTTCATTGCAGACAGGTTCCCAAATAGAAAGAGCATTTCTCCAGGCACCAGTTGAAGAGCGTTGATCAATGGCCTGTTCAAAAACAGTTCTCAT
CCGGATCTGACCTTTACCAACTTCATCCGTTTCACGTACAACATTTTTTAGAACCATGCTTCCCCAGGCATCCCGAATTTGCTCCTCCATCCACGGGGACTGAG
AGCCATTACTATTGCTGTATTTGGTAAGCAAAATACGTACATCAGGCTCGAACCCTTTAAGATCAACGTTCTTGAGCAGATCACGAAGCATATCGAAAAACTGC
AGTGCGGAGGTGTAGTCAAACAACTCAGCAGGCGTGGGAACAATCAGCACATCAGCAGCACATACGACATTAATCGTGCCGATACCCAGGTTAGGCGCGCT
GTCAATAACTATGACATCATAGTCATGAGCAACAGTTTCAATGGCCAGTCGGAGCATCAGGTGTGGATCGGTGGGCAGTTTACCTTCATCAAATTTGCCCATTA
ACTCAGTTTCAATACGGTGCAGAGCCAGACAGGAAGGAATAATGTCAAGCCCCGGCCAGCAAGTGGGCTTTATTGCATAAGTGACATCGTCCTTTTCCCCAAG
ATAGAAAGGCAGGAGAGTGTCTTCTGCATGAATATGAAGATCTGGTACCCATCCGTGATACATTGAGGCTGTTCCCTGGGGGTCGTTACCTTCCACGAGCAAA
ACACGTAGCCCCTTCAGAGCCAGATCCTGAGCAAGATGAACAGAAACTGAGGTTTTGTAAACGCCACCTTTATGGGCAGCAACCCCGATCACCGGTGGAAAT
ACGTCTTCAGCACGTCGCAATCGCGTACCAAACACATCACGCATATGATTAATTTGTTCAATTGTATAACCAACACGTTGCTCAACCCGTCCTCGAATTTCCATA
TCCGGGTGCGGTAGTCGCCCTGCTTTCTCGGCATCTCTGATAGCCTGAGAAGAAACCCCAACTAAATCCGCTGCTTCACCTATTCTCCAGCGCCGGGTTATTT
TCCTCGCTTCCGGGCTGTCATCATTAAACTGTGCAATGGCGATAGCCTTCGTCATTTCATGACCAGCGTTTATGCACTGGTTAAGTGTTTCCATGAGTTTCATT
CTGAACATCCTTTAATCATTGCTTTGCGTTTTTTTATTAAATCTTGCAATTTACTGCAAAGCAACAACAAAATCGCAAAGTCATCAAAAAACCGCAAAGTTGTTTA
AAATAAGAGCAACACTACAAAAGGAGATAAGAAGAGCACATACCTCAGTCACTTATTATCACTAGCGCTCGCCGCAGCCGTGTAACCGAGCATAGCGAG

**Nucleotide sequence of the homologous genes of *Bacteroides thetaiotaomicron***
Highlighted yellow: BT2156, Green:BT2157, Turquoise: BT2158, Pink: BT2159, Teal: BT2160

TAACTTTGCAGCGTTTTAATAAAGCAATTGATTGTATTACGTTTTTAAAAAGCAAAGAAAATGAAAAAATTAGTTTTGATGGCCGTAGCTATCGTAGCAGTATCAT
TCGCATCTTGTGGTAACAAAGCAGCTGACGCAGCAAAAGCTCAACAAGATTCTATCCGCATCGCTGACTCAATCGCAGCTGTAGAAGCAGCAGCAGCTGAAG
CAGAACAAGCAGCAGCTCAGGCAGCTGACTCTTTGAATGCAGATAGCACTGCAACTGAAGCTGTAGCTGAATAATTCAGAACAGCCTAGAGAGAATTGAAAGT
CTGACATGCTAAAAGCATGCACAGACTTTTTTTATGCCTTATTTAGCGTATGAGCTTTTAACAAAAGGAGCATCCAGCAAATAATCCAGACTTACCTCGACACTT
TTCTCTACGGGCATGCTGTAGCTTTCTATCTCTGCAACAAGATGCTTCACACCAGCCGTCTTTGCATTCTTGAAAATAGCGTCGAAACCAACCATTCCGCTCTG
CCCGATTTCTCTGTGATCCTTGATATGGAACATTTTAAAACGACCGGGGTATTTATTAAAATAATCTACCGGACTGTTTTGCCCGCGAACTACCCAATAAACATC
CATCTGGAAGAAAACATATTCAGGATTGGTATGCTCAATCATGTAGTCATACATCACTTTATCTTCTACCTTCTGAAATTCATGAGCATGATTATGGTATCCGAA
CGACATCCCCTGCTGTTTACAACGTTTGCCTATTTCATTGTAATAAGCACAATATGTATCCAACTCCTTAAGTGTTTTAGGCACATCCATCCAAGGAGCTACAAT
GTAACTCATACCGGCAGCTTTATGATCAGCAATACACTGGTCCCACCATTGAAGTGAACTTGAAAAATCACCGGAAGCTAATTCTTCTTTCGACAATCCTCTTG
TGCAATGTGAGGATAAAACCTTCAGCCCGGCAGATTCAACATCTTTTTTAAATTGATCGGGAGTTCTGTCGTAAAACTTTCCATTATTATAATTGGCGGCTTCCA
CACTGGTATATCCCATTTTCGCCAAGTTCTTCAGAAGAGTAATATAAGCGGCGTCACATTTGCCGTCCTTATTGTCCACCTTATTCAATATATCCCTGACAGAAT
AGAGTTGAATCGCTACTTCTTTTTTAGCTTTCGCTTTTGTTTTAGACTGTGCCATGGCCGTAAACGGAATCATTAAAAGAAGAGCCAGCGCACATAGATTTATTT
GGAGTTTCATGTTTGTAGATAAAGGGGTTCATGTATTAAAATAAAAAAGCTGTGTCAGAAGTTCGTCGGGCATATCATTCTGTCATTCAGAACTAAATGCCATGT
TGATGTTACTTTTGACACAGCCTATCTTTGAATTATTTATGTGATCAGTCCAGAACTTTCACGCGGATATTGCGGAACCAAACGTCATCACCGTGATCCTGCATT
CCGATAAATCCTTCGTGGTTTTCGCCACCGCAATTATTCAGCAGTTCGAAAGCCAACGGCCATTTGTCCTGGCTGAATTTGCTGGCTTGCAAAAGGTCAGTCC
ATTGTTTGGTCCACAAGTGGTATTCGAGTACATTCTCGTCATTCTGTCCGTGAACAACAGTACCTTTATAAACCATGATCTTAGCCTTGTTCCATTCGCCGAAAG
GTTTCGCATTCTGAGGAACTGCCGGAATCATGTCATATAAAGAAGCCGACTGCCGGTTATTGTCTTTCCCCAATTTAGCATCCGGATGATTGTCGTTGTCCAAT
ACCTGATATTCAGGTGCGGAAATATAAATAGGTTCGAGCACGTCGTTGCCGTCCTTATCTTTAGAAGTCACTTCCTGAGCTAAATAGAAGATACCGGAGTTACC
GCCTTTGGAAACTTTCCATTCCATTTCCAGTTCGAAGTTTTTGAATTTGTGAGCAAAGATCAGGTCTCCGCCATCACCATCCTGTGCTTCACCGCCACCGGAAC
CGTTGAACTTGATGCAACCGTCCTCGATAGTCCACTTGGAAGGAACTCTGTCCTTGCCATAACCACGCCATCCATTAAATGTTTTGCCGTCAAAGATAGTGATG
TAACCATCCGCATCTACCGGTAATTGCAAGCTATCCATCTCCGCTGCTTTTAATGATTTTGAATATGATACTGCTACTTCGTTTGCCGATTGCTCTTCCTGAGCA
CTCCCGGCTTTCTTCTGACCGCTGCAGGAAACTAAAACTCCCGCTGCAAGACAACAAGCTAATGGATAAAATACTTTCTTCATAATAATAACTATTTAATTTTTA
ATCTTTAATTCTTAACTTACTTTTATCGTGTTATCGTGGCATATCAGGCAACTTCCATCCTTCACGATAATTATGTTTCACCAGTTCGGCTGCAAATTGTTTCGCG
TTGATCGGATCTGTCCAGGTCTTATTGAATGACGGGTGACCATCATGAATCGTAAATCCGTCCTTAATGCAGGTTCTGAGTGTTTCGTTATCACCAATATTAGT
GAAGCACATATTAGCGCCGTCCCATTCCAGTGTCTTGTTCAATCCTTGCAGACGGATAGCCAATACACCCATTACCACCATTTCATTCATCGGTCCCGATTCGG
AGAAATCCGCTTTCGGCATCACGCGGTTGGATTTGTCTTCCTTACAAGCGCGTACCCAGTCCATTTCATGAGAACAAGTAACACGGCGGCATACTTTCGGTGC
ATTGGGCACACGGCCTGACAACAAGAAAGGTTGTTCTCCGTAGCAGCCGCAAATCAATGTATCTTTTGTTCCGTGGAAGATAGTCAGACCACCACCGGGGCC
CATCAGTTGTTTTCCTTCCGGGAAACCTTTCGGACGTTCCGGCATCATACCGCCGTCATACCAGTGTACCTCAACTTCCGGCAAGGCAACTTTCGGCATATTG
TCGCGGGCGGGGAAAATCATCTTCACGTGTTGGGCCTGAGGAGCGCAGGCACTCAGTAATAAAGTAGAAGAACCTTCCACCTTGGTAGGATAACCCAGCTTC
AGAGCTCTGAACGGCTGGTGCAGAATATGACAGGCCATATCACCCAGTGCACCTGTACCGTAATCCCACCATCCACGCCAGTTCCAGGGATGATAAACGTTG
TTGTACGGGTTCAGTTTGGCAGGTCCGGTGAAAAGATCCCAGTTCAATGTGTTGGGGATGCGGTCCGCCTTTTCGGGTGCATTCAGTCCTTGCGGCCAGATA
GGACGGTCGGTGGCGCATTCCACTTTGGTTACTTCGCCGATTTCTCCATTCCATATCCATTCGCAAACCAGGTCGGTACCTTCGCCCGATGCTCCCTGATTAC
CCATTTGAGTAACTACTCCGGTAGAGGCGGCCAGCTTGGTCAGCAAACGGGATTCGTAAACGGAGTGAGTCAACGGTTTCTGGCAGTATACGTGCTTGCCCA
TTGTCATGGCGTCTGCCGTGATGATGGCATGAGTATGGTCGGCGGTAGCAATGATTACGCCGTCAATGGATTTGCCCATTTCATCGTACATCTTGCGATAATC
CCAGTATTTCTTCGCATTCGGGAATTCATCGAATACGCCTTTGGCATATTTCCAGTCTACGTCACAAAGGGCTACGATGTTTTCGGTGCCCTTCACGTTGTTGA
TGTTGGCATGACCCATACCGCCGATACCTACGGCTGCAAGGTTCAGTTTGTCGGTTGGGGAAACGTGCCCGTGGCTCATACCTAAAATAGAGCTGGGAGCAA
TGGTGATTCCTGCTAAAGCAGCGGCTCCCGTTTTAAGAAATTTTCTTCTTGAAAAGTCTGACATGGTTTTATTATTTTAAGGAGTTAGTTATTTGTGGTGGATTA
CTATGCAGCAACCCATATCCATTTTTCATGTTTTTTCTGCGTTCGCGTATTTGTTCAAGAGTCTGCAAATTACCGATAGCCGGCAAGTCATTCATACGAGCATCC
TGCAGGAATTTCCACGCATACTCGGATGCAATAGTCGAGTCATACATTAAAGGCAAACGAGGGTTTCGCTTACCGGTCAATATGGAATCGGCGAACAAATCGC
ACAGCACGTCAATATTCTTACCACCAAAAGGTTTTTCCACACGAAGTGTCTGATGTACTCCGTGTAAATCGACAACAGCTGTTTTGAAATCATGCGTCATACGC
ACGATACCTTCCGTACCTATAATATCTACATAAGAATTGTGCGTCTGATCTTTCGATAATTGTCCATAGACAAAACCCTGGGTTATATCAAAGACTACGCCATTT
TGAAAAGTTCCGTGACATTGTACCCACCAGGGATCTTTATAATTCCACATGTTCACGCCTTGTGCATTCCATGTTCTATATTCACTTTCGGCATACCAGCGAAC
GATGTCAACATAATGCATTCCACAATCATGGAAAGCGGGGCCTTCATATTCATGTCCTTCACCCGGTGCCAGTCCCGGAGTCATGTGGCAAATACGGATAATG
GCTAACTCACCTATTTCTCCCTTCTGGATATATTCCTTCATCAGATTATGATACCAGGAATTTCGCAGATATAAGTTTACCGTAGAAAGTACATTGGAGTTTTCTA
TCATATCCACCACCTTCCACTCCTTTTCCATCGTATCTGCAACAGGTTTTTCCGCGATGATATGTTTTCCGTACCGAATAGCCTTTTCTATTTGTTCGAGCCTTG
AGTCTGCCAATGTGAAGAGCCCGACAACCTGTACACTCTCGTCTTCAAAGATTTTTTGATCATTCTCTACTACAAGAGATTCAGGAGATATTTTTTTAGCAAGTT
CGCGCGTGGTAGGATCTGTATCGCAAATATAGGCAATCTCCCATCGCCCGCTTTTCCGCATCGCCTCCCAATAGAATCTTCCCATTCTACCGAAACCGATTATT
CCTACTTTTATTTTTGTGTCAAGTGAAAATTGTGCATCCATAAAAAATTGCGAATTTAAGTACAAACAACATAAAAGGTATCAATTTTCATCTGTAGGCAGATTGT
TCGAAAACTGATATCGCAAAACTAGCCAAATAAATAACGCTTTATTTAGTCTACACAACTTTATGTAGATTATAAAATCATATTACATATTATAAATCAACACTTTA
GATATACACAAGGATATAAAAAATGTAGATTGTTTGGAAAGTATTTTATTTATCGAATGGCACAAATTCGTTTATATTGGTCAGATTTTCATAATTTCATTTTCAAA
TTCATCCCGATCTACTGCAGCCTTGTTACGCATTTTAGTACGATAATTATAAACAGTACTTGTAGAGCAACGAAGGAAATTGGCAATCTTTCCGCTATCCGTAAT
GCCCAAACGTAACAGCGCATAGATACGAAGTTCCCTATTCAACAAAGCGTCCTGTTTAAGAATGATCTTCTCGTCATCTTTCAGTAAAGCATTGAAATCTGAAA
CAAAAGTCGGATATAAATTCAGAAAGACTCTATCGAAGCGAGTATACAGAGCATCAAGTTCATCATCAATCAATGCGGAAGATTTCAGCTTTTTAATCAGTTCCT
CATAGCACTTATTAATGGCTATTTTATACAGCATATTCTGATACTTCTCCATTTTATGAATATAGCTGAAGCACACGTCGAAAAACTGAGCGATATAATGCTCCTT
TATATTATTAATCTCGCACAGTTCATCATTCTTATCATTCAACTCACTATTCATGCTGTTCAGTTTGTCGTTCAGTCTCAAAAGTTCCTCGTTACTTTGTGCCAGC
GCCCGTTTCATCCGCAACGTTTTTTTCATCTGAATATAAATAAATACGACCAACACGATTAAAAGAAAAAGAGAAACGCTGGTTGAGATAAGAAAAGTAATCAGA
TTCGATTTAGACCTAGCTTCTTCCGTCTGATAAGCCGTATTAATAATAGAATAGAATTTATAGATTTCCATTGCCCGGAAATGAATGCCGGAGGAAACGACATCA
TCAATAGCCGACTGAGTAAATTTGAAAGCATCCGCTAAATTATTTTCCTCATATTGTATCAGTGCAAGTGCCTGCAGGGAGGCTGTTTCGCGTGTGGCATTCTG
AATATCAGCAATGGCGGACATCATAAGATATTTCTTCGCATCATCCTCTCTTTTCAAATAGCGGCTCAGCATGGCATACGAATGAGTAATCATCGCATAGTTAG
GAGTGCCTACCTCCTCAGCATTCAGCAACTCATCCAAAATCTTGATCGCCTTTGTTGAATCGTGTCCTGCATAAGAATAAGCCCGTGACAATTTATAGTCAAAG
GAAGTATGATCCATAAGAGCATACAGTGAATCCTGATAGGCTTCCCGTTTTTTTCCGTACTGATTATTGGTAGCAGAAATCGAATAATACTCCCAAAAACGGGC
ATATGTATCATAATAAGTAGCTAATAATGAACGAGGAAATTCTGATGGCTTCATTTTTTCGAGAAGTAATTCTGCATCTCTATATCTGCCGCACATGGAATATACA
AGGCTGAGTTGCAAAGAAGACTGGTATTTCAAATAGTTTCTATTTAACTGACGAGCGATTTCAAGGTTCCGGTCGACATAATGGATGGCAGAGTCTATATTAAA
TTTCTTATACTCATTGTATAATTGGAGATTCACCTTATATTCCCGATCGGGAGTCAACCCAGAGTCCTTCAGTAAACATTTAATCCGAGCGATTCGCTGCTCTTT
CTTCTGGACGAAGAGCGGCTTCGCATCTATCATTTTATGCAGAGAGTCAGCCAGTATTTTCAATTCGTTATTTGCATAAAGAGCATTGCCAATTCCCAGAGCAA
CCAGGAACAGCAAAACGATTCTCTTCAGATAACAACGGTTGAATTTCATATTATTAGCAAAGATGATAAAAAAAAGCGATTTATCCATCAGAACAAATCGCTTTT
TTAATATATGAAGTAAATTTGCTCTTATTCAGCAACTACTTCGAAAGGAATTTCTACAGAAACTTCCTTGTGAAGTTTAACGATAGCTTTGTAGTTACCAACTTCT
TTCACAGCGTCTTTTACAACGATGATCTTTCTGTCAATTTCGTGACCCAGTTTTGCCAATGCTTCAGCAATCTGGATATTGCTAACAGAACCGAAGATAGTACCT
GTTGAACTAACTTTTGTTGCAATAGTCAAAGAAACGCCTTCCAACTTAGCAGCCAATGCTTCAGCATCCTTCTTGATTTTCTCAAGTTTGTGA

#### Protein sequences

**AL1- Alistipes- Isomerase partial, Original, present on P1C11:**
MKRKLFTAVCILLASAIAPCGAFAKAKKDAKKDIGIQLYSVRDLIGSFGRNQHDYKPVLKALADMGYTSIEAASYNDGKFYGNTPEEFRRDVEAVGMKVLSSHVGSPG
TELEFAL*
**Alistipes- Isomerase, Original, NCBI Reference Sequence: WP_055211454.1**:
MKRKLFTAVCILLASAIAPYGAFAKAKKDAKKDIGIQLYSVRDLIGSFGRNQHDYKPVLKALADMGYTSIEAASYNDGKFYGNTPEEFRRDVEAVGMKVLSSHCGKGL
SDEELASGDFSESMKWWDQCIAAHKAAGMEYIVTPYLPVPKTLKEMQTYCDYLNAIGKKCREAGIKYGYHNHAHEFQKIEDKAVMLDYMIENTDPENLFIELDVYWA
VMGKASPVDYFHKYPGRFKMLHIKDRREIGQSGMVGFDAIFENAKTAGVENIIVEVEQYSYDVEKSVKLSLDYLLEAPFVKASYSK*
**AL1, Alistipes- Isomerase, Cloned construct**:
MKAKKDAKKDIGIQLYSVRDLIGSFGRNQHDYKPVLKALADMGYTSIEAASYNDGKFYGNTPEEFRRDVEAVGMKVLSSHCGKGLSDEELASGDFSESMKWWDQCI
AAHKAAGMEYIVTPYLPVPKTLKEMQTYCDYLNAIGKKCREAGIKYGYHNHAHEFQKIEDKAVMLDYMIENTDPENLFIELDVYWAVMGKASPVDYFHKYPGRFKML
HIKDRREIGQSGMVGFDAIFENAKTAGVENIIVEVEQYSYDVEKSVKLSLDYLLEAPFVKASYSKLEHHHHHH*

**AL2 – Alistipes DUF1080, Original:**
MKKVILFSTCIAAAATMAACGGPGAKGKAAADTAAAETKAVPEYTVINNATVDLAGFPQDKDGYYVIFDGKTFNGWRGYGKDHVPSKWTIDNGAIKFNGSGGGEAQ
DGDGGDLIFTHKFKNFELEVEWKVAKGSNSGIFYLGQEVATTKDGKQRMEPIYISCPEYQVLDNENHPDAKLGVDGNRKSASLYDMIPAVPQNAKPYGEWNKAKIM
VYKGTVVHGQNDQNVLEYHLWTPQWTEMLEKSKFSPEKWPLAFELLNNCGGAQREGYIGFQDHGDDVWFRNIRIKVLD*
**AL2 – Cloned construct:**
MGGPGAKGKAAADTAAAETKAVPEYTVINNATVDLAGFPQDKDGYYVIFDGKTFNGWRGYGKDHVPSKWTIDNGAIKFNGSGGGEAQDGDGGDLIFTHKFKNFELE
VEWKVAKGSNSGIFYLGQEVATTKDGKQRMEPIYISCPEYQVLDNENHPDAKLGVDGNRKSASLYDMIPAVPQNAKPYGEWNKAKIMVYKGTVVHGQNDQNVLEY
HLWTPQWTEMLEKSKFSPEKWPLAFELLNNCGGAQREGYIGFQDHGDDVWFRNIRIKVLDHHHHHH*

**AL3 – Alistipes Oxido1, Original:**
MSSEMSRRKFLKKGAAAAIGLAVVPGTVLGKKFGHTAPSDKLNILGVGIGGRGSSVLRGLESQNIIGLCDVDWKYADHVFKRYPAAKKYNDYRKMFDEMLKSADAV
MVATADHTHAIIAADAMTAGKHVYVEKPLTHTVYESRLLTKLADKYKVATQMGNQGASDEGVRKVCEWIWNGEIGEVRKVETFTDRPIWPQGLSRPEDDQRIPKTLN
WDAFIGPAPYRPYNAIYTPWNFRGWWDFGTGALGDMACHILHPVFKGLKLGYPTKVQGSSTLLLNESAPMAQTVKFVFPARDNMPKVAMPEVEVYWYDGGLKPAR
PEGLPAGKDLNMAGGGVIFYGTKDTLICGCYGVNPYLVSGRVPNAPKVLREIKESHQMDWVRACKEDADDRVPSASDFSEAGPFNEMVVMGVLAVRLQNLNRELL
WDGPNMRFTNIPDDATISAVIKDGFHIKDGHPTFDKTWTDPVNAQQFAQELIKHTYRDGWKLPDMPR*
**AL3 – Cloned construct:**
MKKFGHTAPSDKLNILGVGIGGRGSSVLRGLESQNIIGLCDVDWKYADHVFKRYPAAKKYNDYRKMFDEMLKSADAVMVATADHTHAIIAADAMTAGKHVYVEKPLT
HTVYESRLLTKLADKYKVATQMGNQGASDEGVRKVCEWIWNGEIGEVRKVETFTDRPIWPQGLSRPEDDQRIPKTLNWDAFIGPAPYRPYNAIYTPWNFRGWWDF
GTGALGDMACHILHPVFKGLKLGYPTKVQGSSTLLLNESAPMAQTVKFVFPARDNMPKVAMPEVEVYWYDGGLKPARPEGLPAGKDLNMAGGGVIFYGTKDTLICG
CYGVNPYLVSGRVPNAPKVLREIKESHQMDWVRACKEDADDRVPSASDFSEAGPFNEMVVMGVLAVRLQNLNRELLWDGPNMRFTNIPDDATISAVIKDGFHIKDG
HPTFDKTWTDPVNAQQFAQELIKHTYRDGWKLPDMPRHHHHHH*

**AL4 – Alistipes Oxido2, Original:**
MNPIKVGLIGFGRMGGFYLEEMQKSGNWDIVYICDVSPASRELAQRLSPRSKVISDEQIIFNDPEIQAVGLFALADSRPEQIQKAIASGKHVIAEKPIADTVEKEWQTVA
LAEKAKIFSTVNLYLRNSWYHNTIKDFIAQGEIGELAIIRVCHMTPGLAPGEGHEFEGPSFHDCGMHYVDIARWYAESEFKTWNAQAIRMWDYKDPWWLQCHGTFEN
GIVFDITQGHVYGQLAEKQTHNSYIDIIGTKGIARMTHDFKTAIVELHGVTQTHRLIQPYGGKNIDVLCKLFAESIESGKRNPSLPSFRDAATASEYAWRFLQDSRQHDL
PAIGNLETLQEIRERRRTMSNGYGLLRKRA*
**AL4 – Cloned construct:**
MNPIKVGLIGFGRMGGFYLEEMQKSGNWDIVYICDVSPASRELAQRLSPRSKVISDEQIIFNDPEIQAVGLFALADSRPEQIQKAIASGKHVIAEKPIADTVEKEWQTVA
LAEKAKIFSTVNLYLRNSWYHNTIKDFIAQGEIGELAIIRVCHMTPGLAPGEGHEFEGPSFHDCGMHYVDIARWYAESEFKTWNAQAIRMWDYKDPWWLQCHGTFEN
GIVFDITQGHVYGQLAEKQTHNSYIDIIGTKGIARMTHDFKTAIVELHGVTQTHRLIQPYGGKNIDVLCKLFAESIESGKRNPSLPSFRDAATASEYAWRFLQDSRQHDL
PAIGNLETLQEIRERRRTMSNGYGLLRKRAHHHHHH*

**BT1 – BT2156, Original:**
MKLQINLCALALLLMIPFTAMAQSKTKAKAKKEVAIQLYSVRDILNKVDNKDGKCDAAYITLLKNLAKMGYTSVEAANYNNGKFYDRTPDQFKKDVESAGLKVLSSHCT
RGLSKEELASGDFSSSLQWWDQCIADHKAAGMSYIVAPWMDVPKTLKELDTYCAYYNEIGKRCKQQGMSFGYHNHAHEFQKVEDKVMYDYMIEHTNPEYVFFQM
DVYWVVRGQNSPVDYFNKYPGRFKMFHIKDHREIGQSGMVGFDAIFKNAKTAGVKHLVAEIESYSMPVEKSVEVSLDYLLDAPFVKSSYAK*
**BT2 – BT2156, Cloned construct:**
MQSKTKAKAKKEVAIQLYSVRDILNKVDNKDGKCDAAYITLLKNLAKMGYTSVEAANYNNGKFYDRTPDQFKKDVESAGLKVLSSHCTRGLSKEELASGDFSSSLQW
WDQCIADHKAAGMSYIVAPWMDVPKTLKELDTYCAYYNEIGKRCKQQGMSFGYHNHAHEFQKVEDKVMYDYMIEHTNPEYVFFQMDVYWVVRGQNSPVDYFNKY
PGRFKMFHIKDHREIGQSGMVGFDAIFKNAKTAGVKHLVAEIESYSMPVEKSVEVSLDYLLDAPFVKSSYAKHHHHHH*

**BT2 – BT2157, Original:**
MKKVFYPLACCLAAGVLVSCSGQKKAGSAQEEQSANEVAVSYSKSLKAAEMDSLQLPVDADGYITIFDGKTFNGWRGYGKDRVPSKWTIEDGCIKFNGSGGGEAQ
DGDGGDLIFAHKFKNFELEMEWKVSKGGNSGIFYLAQEVTSKDKDGNDVLEPIYISAPEYQVLDNDNHPDAKLGKDNNRQSASLYDMIPAVPQNAKPFGEWNKAKI
MVYKGTVVHGQNDENVLEYHLWTKQWTDLLQASKFSQDKWPLAFELLNNCGGENHEGFIGMQDHGDDVWFRNIRVKVLD*
**BT2 – BT2157, Cloned construct:**
MGSSHHHHHHSSGLVPRGSAQEEQSANEVAVSYSKSLKAAEMDSLQLPVDADGYITIFDGKTFNGWRGYGKDRVPSKWTIEDGCIKFNGSGGGEAQDGDGGDLIF
AHKFKNFELEMEWKVSKGGNSGIFYLAQEVTSKDKDGNDVLEPIYISAPEYQVLDNDNHPDAKLGKDNNRQSASLYDMIPAVPQNAKPFGEWNKAKIMVYKGTVVH
GQNDENVLEYHLWTKQWTDLLQASKFSQDKWPLAFELLNNCGGENHEGFIGMQDHGDDVWFRNIRVKVLD*

**BT3 – BT2158, Original:**
MSDFSRRKFLKTGAAALAGITIAPSSILGMSHGHVSPTDKLNLAAVGIGGMGHANINNVKGTENIVALCDVDWKYAKGVFDEFPNAKKYWDYRKMYDEMGKSIDGVII
ATADHTHAIITADAMTMGKHVYCQKPLTHSVYESRLLTKLAASTGVVTQMGNQGASGEGTDLVCEWIWNGEIGEVTKVECATDRPIWPQGLNAPEKADRIPNTLNW
DLFTGPAKLNPYNNVYHPWNWRGWWDYGTGALGDMACHILHQPFRALKLGYPTKVEGSSTLLLSACAPQAQHVKMIFPARDNMPKVALPEVEVHWYDGGMMPE
RPKGFPEGKQLMGPGGGLTIFHGTKDTLICGCYGEQPFLLSGRVPNAPKVCRRVTCSHEMDWVRACKEDKSNRVMPKADFSESGPMNEMVVMGVLAIRLQGLNK
TLEWDGANMCFTNIGDNETLRTCIKDGFTIHDGHPSFNKTWTDPINAKQFAAELVKHNYREGWKLPDMPR*
**BT3 – BT2158, Cloned construct:**
MGSSHHHHHHSSGLVPRGSHVSPTDKLNLAAVGIGGMGHANINNVKGTENIVALCDVDWKYAKGVFDEFPNAKKYWDYRKMYDEMGKSIDGVIIATADHTHAIITAD
AMTMGKHVYCQKPLTHSVYESRLLTKLAASTGVVTQMGNQGASGEGTDLVCEWIWNGEIGEVTKVECATDRPIWPQGLNAPEKADRIPNTLNWDLFTGPAKLNPY
NNVYHPWNWRGWWDYGTGALGDMACHILHQPFRALKLGYPTKVEGSSTLLLSACAPQAQHVKMIFPARDNMPKVALPEVEVHWYDGGMMPERPKGFPEGKQLM
GPGGGLTIFHGTKDTLICGCYGEQPFLLSGRVPNAPKVCRRVTCSHEMDWVRACKEDKSNRVMPKADFSESGPMNEMVVMGVLAIRLQGLNKTLEWDGANMCFT
NIGDNETLRTCIKDGFTIHDGHPSFNKTWTDPINAKQFAAELVKHNYREGWKLPDMPR*

**BT4 – BT2159, Original:**
MDAQFSLDTKIKVGIIGFGRMGRFYWEAMRKSGRWEIAYICDTDPTTRELAKKISPESLVVENDQKIFEDESVQVVGLFTLADSRLEQIEKAIRYGKHIIAEKPVADTME
KEWKVVDMIENSNVLSTVNLYLRNSWYHNLMKEYIQKGEIGELAIIRICHMTPGLAPGEGHEYEGPAFHDCGMHYVDIVRWYAESEYRTWNAQGVNMWNYKDPWW
VQCHGTFQNGVVFDITQGFVYGQLSKDQTHNSYVDIIGTEGIVRMTHDFKTAVVDLHGVHQTLRVEKPFGGKNIDVLCDLFADSILTGKRNPRLPLMYDSTIASEYAW
KFLQDARMNDLPAIGNLQTLEQIRERRKNMKNGYGLLHSNPPQITNSLK*
**BT4 – BT2159, Cloned construct:**
MDAQFSLDTKIKVGIIGFGRMGRFYWEAMRKSGRWEIAYICDTDPTTRELAKKISPESLVVENDQKIFEDESVQVVGLFTLADSRLEQIEKAIRYGKHIIAEKPVADTME
KEWKVVDMIENSNVLSTVNLYLRNSWYHNLMKEYIQKGEIGELAIIRICHMTPGLAPGEGHEYEGPAFHDCGMHYVDIVRWYAESEYRTWNAQGVNMWNYKDPWW
VQCHGTFQNGVVFDITQGFVYGQLSKDQTHNSYVDIIGTEGIVRMTHDFKTAVVDLHGVHQTLRVEKPFGGKNIDVLCDLFADSILTGKRNPRLPLMYDSTIASEYAW
KFLQDARMNDLPAIGNLQTLEQIRERRKNMKNGYGLLHSNPPQITNSLKHHHHHH*

**P2B11- Iso1, Original**:
MAVISLGGSMGSVRISGFADEISSDFDKQLDVVKKLGMSYICLRSAGTKGVADYSPADFADELWPKMQAAGIGLSSIGSPIGKVGINDEEGFQKQLVSLEGLCQICEM
TGCRYIRVFSFFIPAGEDPDAYYDKVIEKVKRFVEIAERHDVILIHENEKDIFGDIARRCQELFDAIKSDHFKAAFDFANFVQVGQDPVAAWDLLHEHVVYIHIKDAVHGS
NENVVAGTGDGHIEEILKRAIVDEDYEGFLTLEPHLVIFDSLKMLETKDVSDIIRGDKAKDGEEGYTMQYNALVEILGHIGATAS*
**P2B11- Iso1, Cloned construct**:
MAVISLGGSMGSVRISGFADEISSDFDKQLDVVKKLGMSYICLRSAGTKGVADYSPADFADELWPKMQAAGIGLSSIGSPIGKVGINDEEGFQKQLVSLEGLCQICEM
TGCRYIRVFSFFIPAGEDPDAYYDKVIEKVKRFVEIAERHDVILIHENEKDIFGDIARRCQELFDAIKSDHFKAAFDFANFVQVGQDPVAAWDLLHEHVVYIHIKDAVHGS
NENVVAGTGDGHIEEILKRAIVDEDYEGFLTLEPHLVIFDSLKMLETKDVSDIIRGDKAKDGEEGYTMQYNALVEILGHIGATASHHHHHH*

**P2B11- Oxido1, Original**:
MKPGGEKEQEMEKVRYGIIGVGNQGGAYAGFLTGTGNVPGMPAAPCPPHCALGALCDIDPQKEEMCKEKYPDVPFYKDWKDMVASGDVDAVITTVPHYLHTEIAIY
CLEHGMNVLVEKPAGVYAKSVREMNECAAAHPEVTFGIMFNQRTNKLYQKIREIVASGELGEIRRSNWIINNWYRPDSYYRLSDWRATWGGEGGGVLVNQAPHQL
DLWQWICGIPTTVYANCINGSHRDIAVENDVTVLTEYENGATGSFITCTHDLLGTDRFEIDLDGGKIVVEDSKKAYIYRFKETETAVNARDMDWMQIAMLTSSNGNSD
DKMFEVEEFENTDGWGYQHTTVMENFAQHIIDGTPLLAPGSDGINGVRLANAIQLSGWTGEKVANPVDEDKYLAELNKRIEAEGKFPVRE*
**P2B11- Oxido1, Cloned construct**:
MEKVRYGIIGVGNQGGAYAGFLTGTGNVPGMPAAPCPPHCALGALCDIDPQKEEMCKEKYPDVPFYKDWKDMVASGDVDAVITTVPHYLHTEIAIYCLEHGMNVLV
EKPAGVYAKSVREMNECAAAHPEVTFGIMFNQRTNKLYQKIREIVASGELGEIRRSNWIINNWYRPDSYYRLSDWRATWGGEGGGVLVNQAPHQLDLWQWICGIPT
TVYANCINGSHRDIAVENDVTVLTEYENGATGSFITCTHDLLGTDRFEIDLDGGKIVVEDSKKAYIYRFKETETAVNARDMDWMQIAMLTSSNGNSDDKMFEVEEFEN
TDGWGYQHTTVMENFAQHIIDGTPLLAPGSDGINGVRLANAIQLSGWTGEKVANPVDEDKYLAELNKRIEAEGKFPVREHHHHHH*

**P2B11- Oxido2, Original**:
MKRAAIVGMGAIYPIHSAAIQALDGIELVGVCDIDAKKRAAAPEGVPAFENVREMVEQTHPDCVHVCLPHYLHYPISKQVVEMGVNVLCEKPVALNGREALEFRRLEQ
EHPEVKIAVSLQNRLNETTEELVRIIGSGEHGKVMGIRAEVPWYRPLAYYQAGPWRGSWDQAGSGVMMNQAIHTIDLMYLLGGPVQRIKASVEQILDYGIEVEDTVS
ARFQYENGAIGHLYATNANFKNEGVNISVDLECASFRMRDNVLYEVGEGNVETKLVEDQKLPGSKFYYGASHKKLIRGFYDCLEDGSDEYIHVRDAYMSVHLIDVIKD
SGLTGSWANV*
**P2B11- Oxido2, Cloned construct**:
MGSSHHHHHHSSGLVPRGSHMASMTGGQQMGRGSMKRAAIVGMGAIYPIHSAAIQALDGIELVGVCDIDAKKRAAAPEGVPAFENVREMVEQTHPDCVHVCLPHY
LHYPISKQVVEMGVNVLCEKPVALNGREALEFRRLEQEHPEVKIAVSLQNRLNETTEELVRIIGSGEHGKVMGIRAEVPWYRPLAYYQAGPWRGSWDQAGSGVMM
NQAIHTIDLMYLLGGPVQRIKASVEQILDYGIEVEDTVSARFQYENGAIGHLYATNANFKNEGVNISVDLECASFRMRDNVLYEVGEGNVETKLVEDQKLPGSKFYYG
ASHKKLIRGFYDCLEDGSDEYIHVRDAYMSVHLIDVIKDSGLTGSWANV*

**P2B11- Iso2, Original**:
MAEKGLIGVQMSTIAPAKMPKFDAYEAMGKLSDIGYHCVEISQVPMTKENVDGFRRAIDELGLNVSSCTASVGPLMPGVPGETLSDPDDFKKIVEDCHALDCDMLRI
GMLPISCMGSFEKAMDFAAQAEECAAKLKEEGIDLYYHNHHVEFVRYNGEYLLDIIRDHAPHVGFELDTHWIHRGGEDPVSFIKKYAGRIRLLHLKDYRVVEPKFPEG
NFDPAGFMQAFTSNIEFAEVGEGTLDIKGCIEAGLAGGGEYFLVEQDDTYGRDPFESLKISHDNLVKLGYEDWF*
**P2B11- Iso2, Cloned construct**:
MAEKGLIGVQMSTIAPAKMPKFDAYEAMGKLSDIGYHCVEISQVPMTKENVDGFRRAIDELGLNVSSCTASVGPLMPGVPGETLSDPDDFKKIVEDCHALDCDMLRI
GMLPISCMGSFEKAMDFAAQAEECAAKLKEEGIDLYYHNHHVEFVRYNGEYLLDIIRDHAPHVGFELDTHWIHRGGEDPVSFIKKYAGRIRLLHLKDYRVVEPKFPEG
NFDPAGFMQAFTSNIEFAEVGEGTLDIKGCIEAGLAGGGEYFLVEQDDTYGRDPFESLKISHDNLVKLGYEDWFHHHHHH*
